# Morphological Profiling Dataset of EU-OPENSCREEN Bioactive Compounds Over Multiple Imaging Sites and Cell Lines

**DOI:** 10.1101/2024.08.27.609964

**Authors:** Christopher Wolff, Martin Neuenschwander, Carsten Jörn Beese, Divya Sitani, Maria C. Ramos, Alzbeta Srovnalova, María José Varela, Pavel Polishchuk, Katholiki E. Skopelitou, Ctibor Škuta, Bahne Stechmann, José Brea, Mads Hartvig Clausen, Petr Dzubak, Rosario Fernández-Godino, Olga Genilloud, Marian Hajduch, María Isabel Loza, Martin Lehmann, Jens Peter von Kries, Han Sun, Christopher Schmied

## Abstract

Morphological profiling with the Cell Painting assay has emerged as a promising method in drug discovery research. The assay captures morphological changes across various cellular compartments enabling the rapid identification of the effect of compounds. We present a comprehensive morphological profiling dataset using the carefully curated and well-annotated EU-OPENSCREEN Bioactive Compound Set.

Our profiling dataset was generated across multiple imaging sites with high-throughput confocal microscopes using the Hep G2 as well as the U2 OS cell line. We employed an extensive assay optimization process to achieve high data quality across the different imaging sites. An analysis of the four replicates validates the robustness of the generated data. We compare morphological features of the different cell lines and map the profiles to activity, toxicity, and basic compound targets to further describe the dataset as well as to demonstrate the potential of this dataset to be used for mechanism of action exploration.

## Introduction

High-throughput morphological profiling of small molecule libraries, also known as the Cell Painting assay, has received increasing attention in drug discovery research (Bray et al. 2016; Gustafsdottir et al. 2013; Ziegler, Sievers, and Waldmann 2021). Compared to conventional high-throughput screening of a single biological target, morphological profiling with high content imaging offers the advantage of identifying multiple biological activities of small chemical compounds simultaneously, promising to substantially accelerate the early stage of the drug discovery process. Moreover, this method enables to predict toxicity and more specifically the of mechanism of action (MOA) of drug-like compounds at the cellular or subcellular levels in a non-invasive manner (Boutros, Heigwer, and Laufer 2015). A typical Cell Painting assay uses six fluorescent stains imaged over multiple channels, revealing morphological changes upon perturbation of cells in eight major cellular compartments, namely DNA, cytoplasmic RNA, nucleoli, actin, Golgi apparatus, plasma membrane, endoplasmic reticulum, and mitochondria (Bray et al. 2016).

To analyze the morphological changes in cells, induced by small chemical compounds in these different cellular compartments, high dimensional image features need to be extracted from the generated images (Caicedo et al. 2017; Chandrasekaran et al. 2021; Pratapa, Doron, and Caicedo 2021). In classical Cell Painting analysis, hundreds of handcrafted image features are extracted using computational tools such as CellProfiler (Stirling et al. 2021). The extracted features serve as fingerprints or profiles that quantitatively characterize the induced cellular phenotypes. Dimension reduction and clustering of the profiles enable the identification of biological activity of uncharacterized chemical compounds. These methods have been successfully employed in recent years to identify chemical probes and drug-like molecules for various biological targets (Foley et al. 2020; Hughes et al. 2020; Laraia et al. 2020; Schneidewind et al. 2020) such as the Sigma 1 receptor antagonist (Wilke et al. 2021) and mitotic kinesin inhibitors (Liu et al. 2023), as well as for treating SARS-CoV-2 infections (Mirabelli et al. 2021).

For an exhaustive characterization of small molecules and their activity, it is crucial that large high quality data sources exist that systematically assay under as many experimental conditions, e.g. compound concentrations and cell models, as possible. One large data source is the Joint Undertaking for Morphological Profiling (JUMP) Cell Painting Consortium, which has very recently published and released a large collection of the Cell Painting data using U-2 OS cells stemming from a joint effort of various academia and industrial partner sites (Chandrasekaran et al. 2023; Cimini et al. 2022). Here, we present a Cell Painting dataset for one of the compound sets from EU-OPENSCREEN. EU-OPENSCREEN is a European Research Infrastructure Consortium (ERIC) dedicated to accelerating the discovery of small molecule compounds for new biological targets by providing academic research groups with open-access to high-throughput screening technologies (Frank 2014). EU-OPENSCREEN hosts multiple compound collections such as the European Chemical Biology Library (ECBL), which includes more than 100,000 chemically diverse and commercially available compounds (Horvath et al. 2014). Part of the ECBL is the Bioactive Compound set. This smaller collection consists of 2,464 bioactive compounds carefully chosen for their diverse biological activity including 681 approved drugs and 385 highly selective probes. This set is also part of the pilot library of the ECBL, thus serves as an ideal reference for the Cell Painting assay.

Within this study, we generated multiple datasets of the EU-OPENSCREEN Bioactive Compound Set over four different imaging sites, primarily using the Hep G2 cell line and for comparison with other datasets in the Cell Painting community in the U-2 OS cell line. Image acquisition was performed using high throughput confocal microscopes. To validate our assay, we performed multiple replicates, which demonstrated the high reproducibility and robustness of the method. To facilitate further research in this field, we have made the data freely available via the Cell Painting Gallery (Weisbart et al. 2024), which will inspire the development of novel computational approaches for identifying the biological activity of chemical compounds. Moreover, this dataset will serve as an important reference for future high-throughput screening based on the larger EU-OPENSCREEN collection.

## Data and code Availability

Annotations for the EU-OPENSCREEN Bioactive Compound Set are available via the Probes & Drugs portal (Skuta et al. 2017): https://www.probes-drugs.org/compounds/standardized#compoundset=353@AND

Raw images are available via the Cell Painting Gallery (Weisbart et al. 2024): cpg0036-EU-OS-bioactives: https://cellpainting-gallery.s3.amazonaws.com/index.html#cpg0036-EU-OS-bioactives/

Aggregated profiles, processed profiles, code for data analysis as well as plots for figures are available via a zenodo repository: https://doi.org/10.5281/zenodo.13309566

Analysis code for aggregated profiles and results are available via: https://github.com/schmiedc/EU-OS_bioactives

## Results

### EU-OPENSCREEN Bioactive Compound Set

The EU-OPENSCREEN Bioactive Compounds are a densely annotated small compound set that consists of 2464 compounds, with 96% of the compounds having at least a single annotated target (**Figure 1A**). This set is enriched for approved drugs, chemical probes and compounds with known MOA (See Methods for selection strategy). The majority of compounds have multiple targets annotated in the literature, with a median number of six targets per compound and one compound with a maximum of 272 targets annotated, demonstrating the capability of small compounds to interact with multiple targets (i.e. polypharmacology, **Figure 1B**). The set was generated with the intention of a wide proteome coverage and therefore the targets range over many different target classes (**Figure 1C**, **Suppl. Table 1**) and are involved in many different pathways (**Figure 1D**, **Suppl. Table 2**).

**Figure 1:**
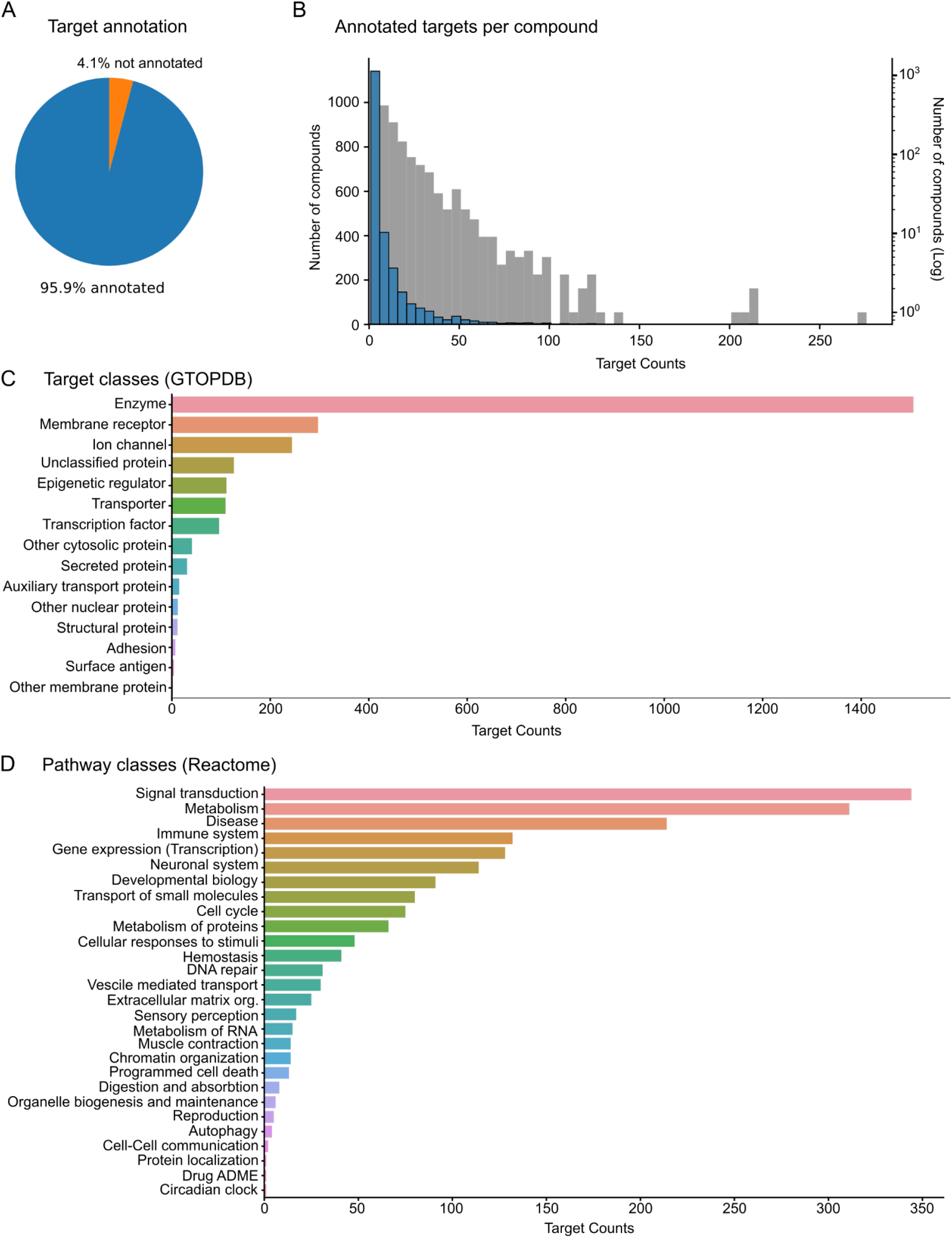
EU-OPENSCREEN Bioactive Compound Set. (A) Almost all of the 2464 compounds of the bioactive set have at least a single annotated target. (B) Histograms of the number of compounds over the number of targets (bin size of 5, log scale in gray). (C) The annotated targets range over a diverse number of target classes based on their GTOPDB annotation. (D) The targets are implicated in a diverse range of pathway classes based on their Reactome annotation.

### Cell Painting assay

To characterize the EU-OPENSCREEN Bioactive Compound Set we employed a Cell Painting assay based on an established protocol (Bray et al. 2016). The assay was carried out at four different imaging sites (FMP - Leibniz-Forschungsinstitut für Molekulare Pharmakologie, Germany; IMTM - Institute of Molecular and Translational Medicine; MEDINA - Fundación MEDINA; USC – Universidad de Santiago de Compostela, Spain) with the 2464 compounds of the set distributed on seven 384 well plates. The assay was performed over four replicates per dataset from the different sites. After cell seeding, the cells were grown for 24 hours before being incubated for 24 hours with the compounds at 10 µM concentration (**Figure 2A**). DMSO was used as a negative control and reference for the plate normalization. We used Tetrandrine and Nocodazole at 5 µM concentration as positive controls since these compounds show a strong phenotypic response in different cell lines.

**Figure 2:**
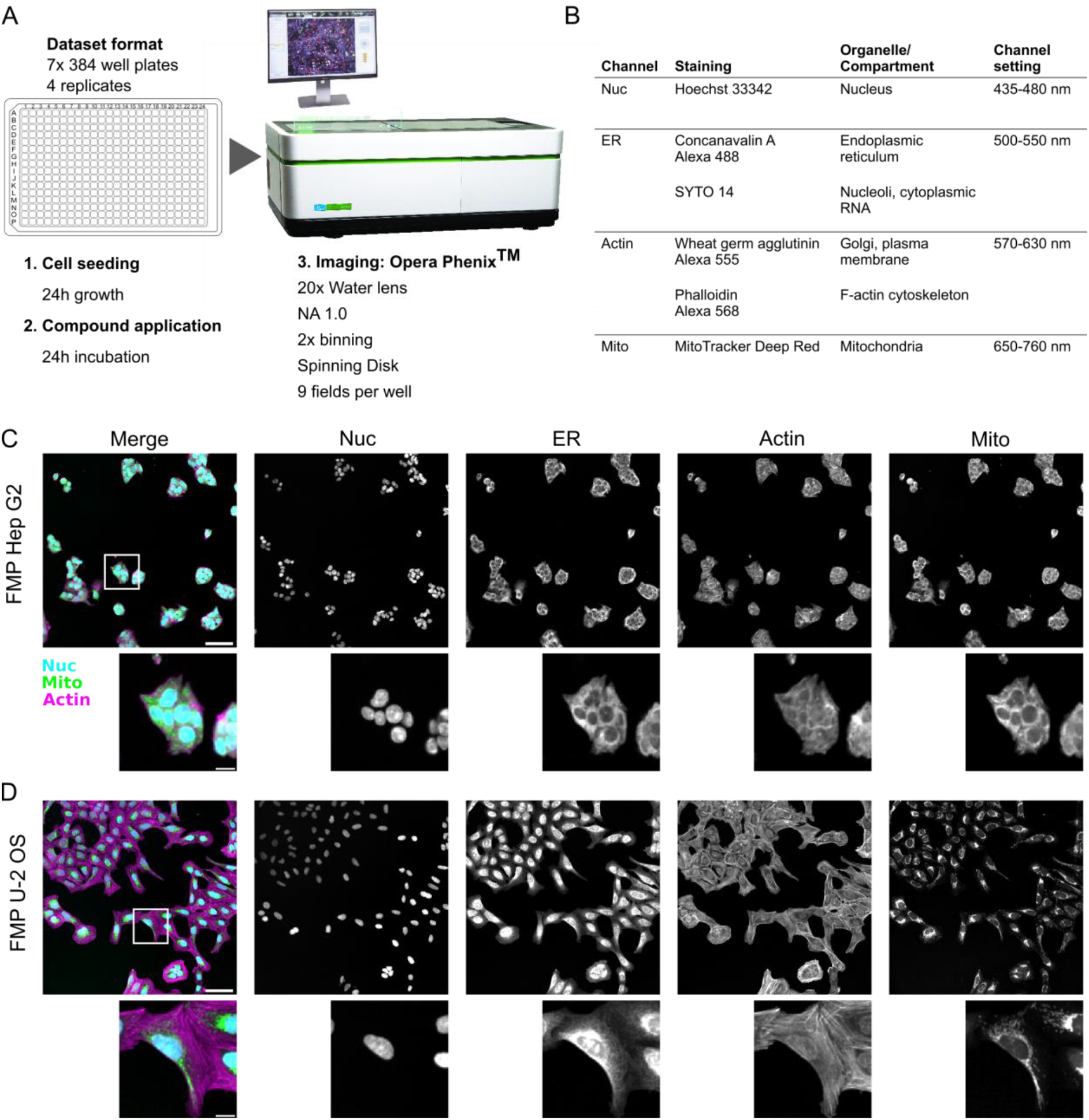
Cell Painting assay performed on EU-OPENSCREEN Bioactive Compound Set. (A) Cell Painting approach using high-throughput confocal imaging. Image of Opera Phenix, credit and ©PerkinElmer. (B) Four imaging channels with the associated staining labeling different cellular compartments. (C) Composite image of the nucleus (Nuc, Cyan), mitochondria (Mito, green) and the actin (Actin, magenta) channels. Individual channels in gray-scale of Hep G2 cells treated with DMSO are shown; a cluster of cells is shown below. (D) Merge of the nucleus (Nuc, Cyan), mitochondria (Mito, green) and the actin (Actin, magenta) channels and the individual channels in gray-scale of U2OS cells treated with DMSO. Enlarged single cell cropped close to the center of the field of view is shown below. Note that the individual channels between the different cell lines were adjusted using different brightness contrast settings. (C, D) Scale bars in merge panels correspond to 100 µm in the full field of view and 20 µm in the crop.

The cells were then fixed and stained with six stains labeling different cellular compartments. Imaging acquisition was performed using spinning disk confocal systems over nine fields per well (**Figure 2A**). The six cellular stains were then acquired in four separate channels (**Figure 2B**). The assay was performed primarily on the Hep G2 cell line generating 387072 images per dataset (**Figure 2C**). One site (FMP) additionally applied the Cell Painting assay with the bioactive set on the U-2 OS cell line (**Figure 2D**), a cell line with many already existing Cell Painting datasets (Akbarzadeh et al. 2022; Bray et al. 2017; Chandrasekaran et al. 2023; Christoforow et al. 2019; Grigalunas et al. 2021). We were able to directly compare our data from the Hep G2 cell line with data based on this very widespread cell line and will enable the Cell Painting community to compare the Cell Painting data from the Bioactive Compound Set with their own datasets.

Overall, we aimed to produce high quality data, increasing comparability across sites keeping the experimental conditions as consistent as technically feasible. To achieve this we performed an extensive evaluation and validation process. First, we selected suitable imaging sites based on submitted proposals and validation data, which were quantitatively evaluated by two external reviewers (**See Methods: Assay optimization and standardization, Supplemental Methods**). Further, we optimized the protocol to reduce variability across sites. In our experience the cell culture as well as staining conditions were responsible for most of the variability and thus we supplied the same cell culture serum, cells and the same lot fluorescent dyes centrally. For practical and technical reasons some parameters remained different across sites. For instance, the four imaging sites employed three different microscopy systems (**See Methods: Image acquisition**). Since the same compound dataset has been acquired with four replicates, the dataset will provide an opportunity to study and develop methods to overcome experimental variability during downstream processing.

### Extraction of morphological profiles

For analyzing the image data we performed an established JUMP-CP CellProfiler based pipeline (Chandrasekaran et al. 2023) extracting 2977 handcrafted image features within single cells based on three cell areas (nucleus, cell, cytoplasm). The features based on single cells were filtered using a Histogram Based outlier selection (HBOS) (Rezvani, Bigverdi, and Rohban 2022), as well as missing and infinite values. Single cell features were aggregated using a median per well. A quality control visualization was based on the intensity in the individual acquired channel as well as the cell count plotted as a heatmap for each plate (**Suppl_QC_FMP_U2OS.pdf, Suppl_QC_FMP_HepG2.pdf, Suppl_QC_IMTM_HepG2.pdf, Suppl_QC_MEDI_HepG2.pdf, Suppl_QC_USC_HepG2.pdf**). The heatmaps indicate minimal plate-based artifacts (e.g. drift, edge effects). Additionally, calculating the median of the four replicates for most channels effectively reduces the majority of these artifacts (**Suppl_QC_FMP_U2OS.pdf, Suppl_QC_FMP_HepG2.pdf, Suppl_QC_IMTM_HepG2.pdf, Suppl_QC_MEDI_HepG2.pdf, Suppl_QC_USC_HepG2.pdf**). The only exception was the image data acquired using MitoTracker Deep, which for the FMP and USC datasets consistently showed noticeable edge effects in all plates.

Analyzing the raw cell numbers per well revealed that most of the wells show an increase of cell number from the initial number of seeded cells. A small number of wells were exhibiting a significant reduction in cell number, indicating severe cytotoxic effects of some compounds in the set at the given concentration (**Figure 3A**). The median cell number, acquired from the 9 fields per well and after cell filtering, was varying between the different imaging sites from 691 to 1659 cells (**Figure 3A**, **Suppl. Table 3**). We also visualized the cell numbers of the control wells, showing the expected reduction of average cell numbers in the positive controls compared to the negative controls, with Nocodazole showing a larger overall reduction in cell number than Tetrandrine in both cell lines (**Figure 3B**). Of note is the more pronounced reduction in the overall cell number at the given concentration in the U-2 OS dataset (**Figure 3B**: **FMP U-2 OS**), which was much more pronounced as compared to the Hep G2 cells from all imaging sites (**Figure 3B**: **Hep G2**). The potent cytotoxic effect of Nocodazole in U-2 OS cells was also evident through the increased number of dead cells, very small and round cells with small and bright nucleus, in the images of the U-2 OS cells (**Suppl. Figure 1B**) as compared to the images of the Hep G2 cells (**Suppl. Figure 2B**).

**Figure 3:**
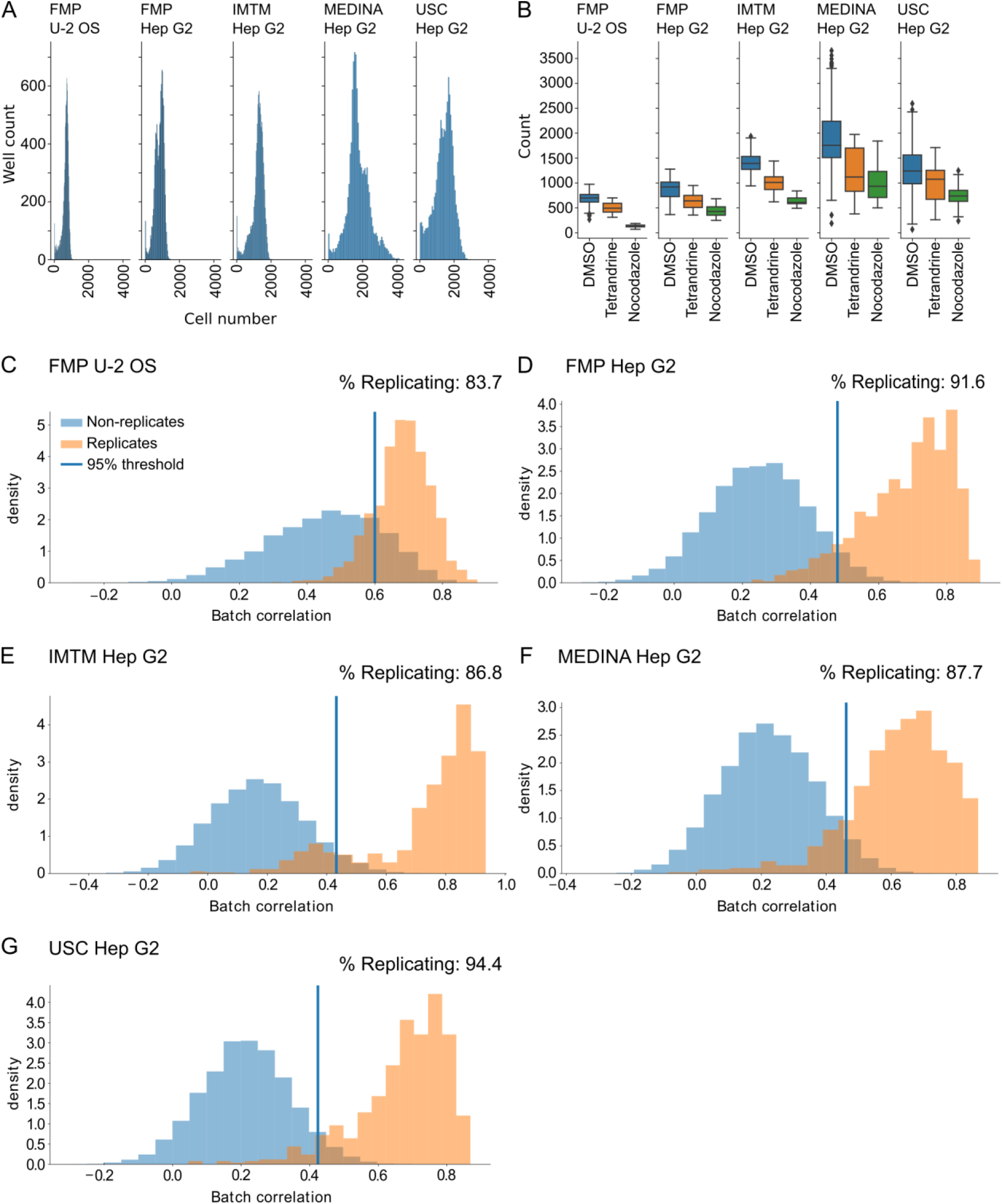
Cell numbers and profile reproducibility in each dataset. (A) Raw cell count over all plates and wells in U-2 OS and Hep G2 from four different sites. (B) Cell count in negative and positive controls in U-2 OS and Hep G2 cells from four imaging sites. (C) Percent replication based on 1600 highly active and non-toxic compounds in FMP U-2 OS cells. (D) Percent replication on 861 highly active and non-toxic compounds in FMP Hep G2 cells. (E) Percent replication of 676 highly active and non-toxic compounds in IMTM Hep G2 cells. (F) Percent replication of 845 highly active and non-toxic compounds in MEDI Hep G2 cells. (G) Percent replication of 607 highly active and non-toxic compounds in USC Hep G2 cells.

For further analysis we reduced the total 2977 features to around 700 features per dataset (**Suppl. Table 4**) based on established feature reduction approaches (Bray et al. 2016; Caicedo et al. 2017; Chandrasekaran et al. 2023; Serrano et al. 2023). Feature reduction was performed by removing features with missing values, low variance, outlier features and most importantly reducing highly correlated features. We observed that the adopted feature selection, generated a feature set that was balanced across various feature types and cellular areas (**Suppl_FeatureReduction_FMP_U2OS.txt, Suppl_FeatureReduction_FMP_HepG2.txt, Suppl_FeatureReduction_IMTM_HepG2.txt, Suppl_FeatureReduction_MEDI_HepG2.txt, Suppl_FeatureReduction_USC_HepG2.txt**). For a qualitative assessment, we visualized the median consensus morphological profiles over the replicates per plate for the positive controls. As the positive controls are present across the seven plates of the Cell Painting assay, the visualization showed that the selected features were highly consistent across the different plates and produced distinct patterns for the two different control compounds (**Supplemental Figure 3A & B**).

### Toxicity, activity and reproducibility of imaged compounds

We then proceeded with a general characterization of the datasets and the compound set by assessing the overall activity, cytotoxicity, as well as the reproducibility of the compound profiles. Highly toxic compounds, albeit bioactive, usually show nonspecific mechanisms of action (Dahlin et al. 2023). Conversely, compounds with very low activity, at the given concentration in the specific cell line, may lead to unspecific morphological profiles. Toxicity was assessed based on cell number, where compounds were defined as acting toxic if they reduced the cell number to below 2.5 standard deviations of the median cell number of the entire dataset. Here, we show that only 3-7% of compounds in the bioactive set had a toxic effect on the cells, with Hep G2 showing a slightly higher resistance against the toxic effects of compounds at the used concentration (**Table 2**).

**Table 2:** Number of compounds per processing step in each dataset.

| Site | Cell line | All | Toxic | % toxic | low active | % low active | Non replicating | % non-replicating | After filtering |
| --- | --- | --- | --- | --- | --- | --- | --- | --- | --- |
| FMP | U-2 OS | 2467 | 177 | 7% | 690 | 28% | 261 | 11% | 1339 |
| FMP | Hep G2 | 2467 | 103 | 4% | 1503 | 61% | 72 | 3% | 789 |
| IMTM | Hep G2 | 2467 | 118 | 5% | 1673 | 68% | 89 | 4% | 587 |
| MEDINA | Hep G2 | 2467 | 80 | 3% | 1542 | 63% | 104 | 4% | 741 |
| USC | Hep G2 | 2467 | 92 | 4% | 1768 | 72% | 34 | 1% | 573 |

**Table 2:** Devices for cell staining.

|  | FMP | IMTM | Medina | USC |
| --- | --- | --- | --- | --- |
| cell seeding | BioTek EL406 | Multidrop | Multidrop | Multidrop |
| compound transfer | Echo 650<br>Biomek i7 | Echo 550 | Echo 550 | Echo 550 |
| aspiration | BioTek EL406 | BioTek EL406<br>Blue Washer | BioTek EL406 | BioTek EL406 |
| dispensing | BioTek EL406<br>Integra VIAFLO 384 | CERTUS FLEX | Multidrop | Multidrop<br>Integra VIAFLO 384 |

For assessing the activity of compounds, we used the induction score (Christoforow et al. 2019), defining compounds exhibiting lower activity in the given cell line at the used concentration when less than 5% of their features were positively or negatively deviating compared to the DMSO negative control. Using this induction score 28% of the total number of compounds in the case of the U-2 OS dataset show lower activity. In the case of the Hep G2 datasets the z-scores were in general much lower compared to the U-2 OS cells. This difference is also apparent when applying the same induction threshold, which describes 61-72% of compounds in the Hep G2 cell lines as having lower activity (**Table 2**). The fact that the Hep G2 cell line shows a much lower response than other cell lines has been recently described (Heinrich et al. 2023). However, one must note that the induction threshold applied here to assess activity was optimized for U-2 OS cells using a handpicked feature set (Christoforow et al. 2019). In this work, we aimed to specifically compare the Hep G2 datasets to the U-2 OS dataset and thus did not change this threshold, other analysis approaches might require adjusting such an analysis to the specific cell lines and feature set and used concentration.

We further computed the percentage of replicating compounds within the datasets after applying toxicity and induction filters. This analysis revealed that the morphological profiles are highly replicating in both cell lines over the different imaging sites with datasets showing a percent replication from 84-94% (**Figure 3C-G**). Thus, only 1-11% of compounds exhibit non-replicating profiles from the original 2467 compounds before toxicity and induction filter (**Table 2**). The percent replication scores of these datasets are in line with the published scores of comparable datasets (Cimini et al. 2022). For a final quality control of the dataset, we also filtered the non-replicating compounds and performed dimensionality reduction using Uniform Manifold Approximation and Projection (UMAP) (McInnes et al. 2018). This allows the visual detection of any batch effects in the data (Arevalo et al. 2024). Indeed, the U-2 OS data produced at the FMP imaging site exhibits some batch effects (**Suppl. Figure 4 A & B**). The Hep G2 datasets from all imaging sites show no apparent batch effects using this visualization (**Suppl. Figure 4 C & D, Suppl. Figure 5 A-F).**

### Comparison of Hep G2 and U-2 OS

We further analyzed the available datasets by comparing the U-2 OS and the Hep G2 cell lines. For a direct comparison, we focus on biologically relevant differences between these cell lines. We thus compare the data from one imaging site to exclude confounding factors that arise from any technical differences (e.g. microscopy, precise staining protocols and devices). Directly comparing the features after feature reduction revealed that most features are consistent across the different cell lines (**Figure 4A**). The morphological profiles of the 428 overlapping features in the positive controls revealed that some features exhibit consistent responses across cell types, while many others displayed varying responses (**Suppl. Figure 3 C & D**), in line with previous findings in the literature (Willis, Nyffeler, and Harrill 2020).

**Figure 4:**
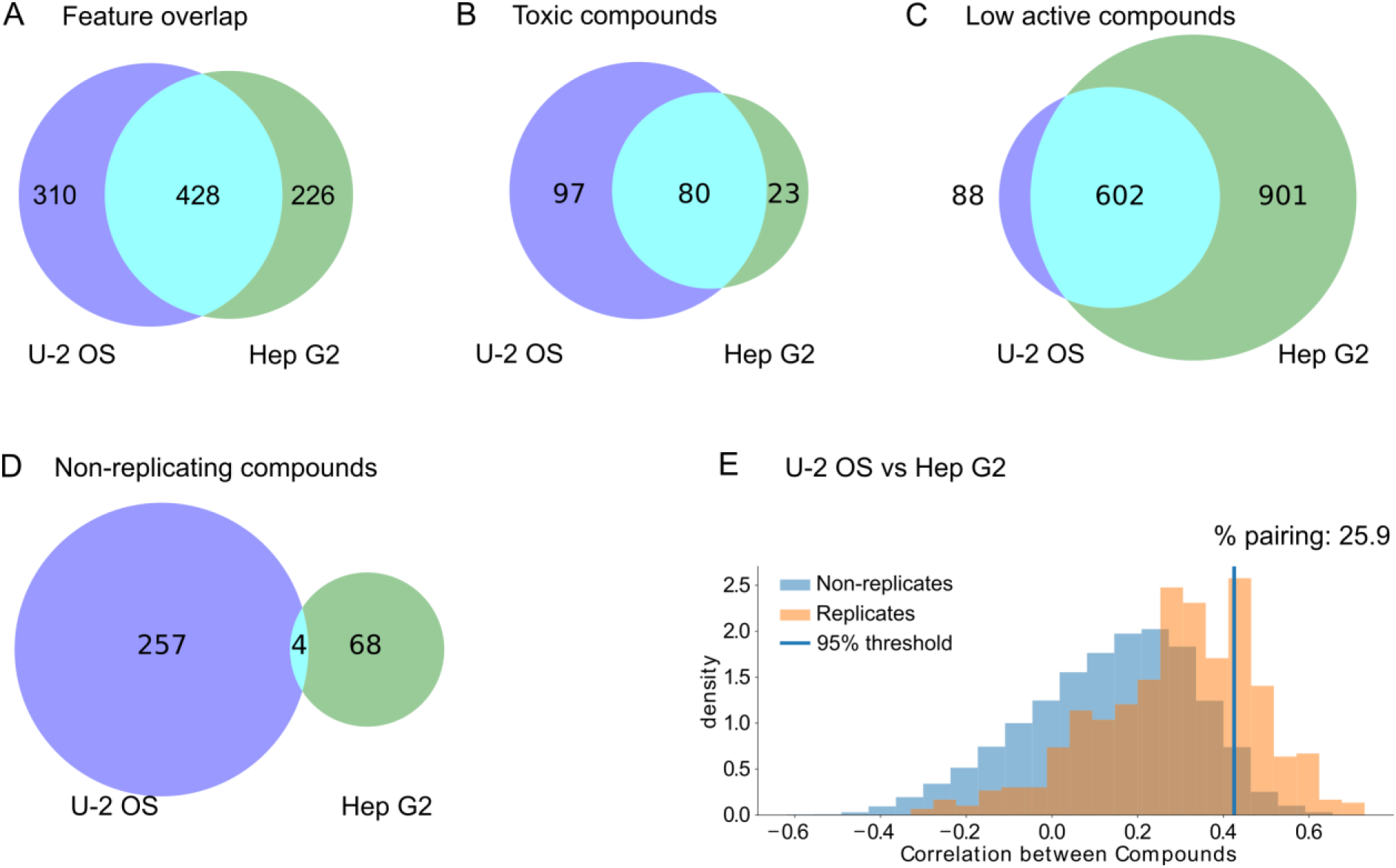
Comparison across cell lines based on FMP datasets. (A) Number of image features after feature reduction and feature overlap between U-2 OS and Hep G2 cells. (B) The number of toxic compounds and overlap between different cell types. (C) Number of compounds that did not pass the induction filter (lower active compounds) and the overlap between U-2 OS and Hep G2 cells. (D) Number of non-replicating compounds and overlap between different cell lines. (E) Percent pairing of 564 non-toxic and highly active compounds across U-2 OS and Hep G2 datasets from the FMP imaging site compared over 427 overlapping features.

We have already noted that Hep G2 cells exhibited greater resistance to toxic compounds at the given concentration when compared to U-2 OS cells (**Figure 3A & B**, **Table 2**). We subsequently investigated the extent of overlap between toxic compounds in the different cell lines and found that 78% of toxic compounds in Hep G2 cells overlapped with compounds defined as toxic in U-2 OS cells (**Figure 4B**). Conversely, a higher response to the compounds was observed in the U-2 OS cell lines. According to the induction filter, more than 60% of the compounds displayed lower activity in the Hep G2 cell line (**Table 2**). Additionally, we observed that 87% of compounds with low activity in U-2 OS also had low activity in the Hep G2 cell line (**Figure 4C**). In summary, it appears that the Hep G2 cell line exhibits a smaller morphological response to the same compounds at the given concentration of 10μM, encompassing both their overall activity and toxicity.

Overall, the datasets show a high technical replication after filtering highly toxic and lower active compounds (**Table 2**). When comparing the non-replicating compounds in both datasets we find as expected that they do not share a large overlap, as this will be determined by random technical variability (**Figure 4D**). We further made a direct quantitative comparison of the profiles across 564 non-toxic and highly active compounds in the different cell lines. To this end we used a variation of the percent replicating compounds, which we term percent pairing to differentiate it from technical replication and other metrics such as percent matching (Way et al. 2021). We found that about 26% of the compared compounds have a correlating profile significantly elevated from random samples in the dataset (**Figure 4E**).

### Analysis of morphological profiles

To visualize the wealth of data in the extracted profiles, we performed dimensionality reduction using UMAP. For a description of the entire datasets, we only filtered the datasets for the non-replicating compounds, not filtering the toxic and low active compounds. We then proceeded to map the control compounds onto these visualizations (**Suppl. Figure 7A-F**). We further projected the toxic and low active compounds as well one basic MOA based on compounds having Tubulin as an annotated target onto these feature maps (**Figure 5A-F**).

**Figure 5:**
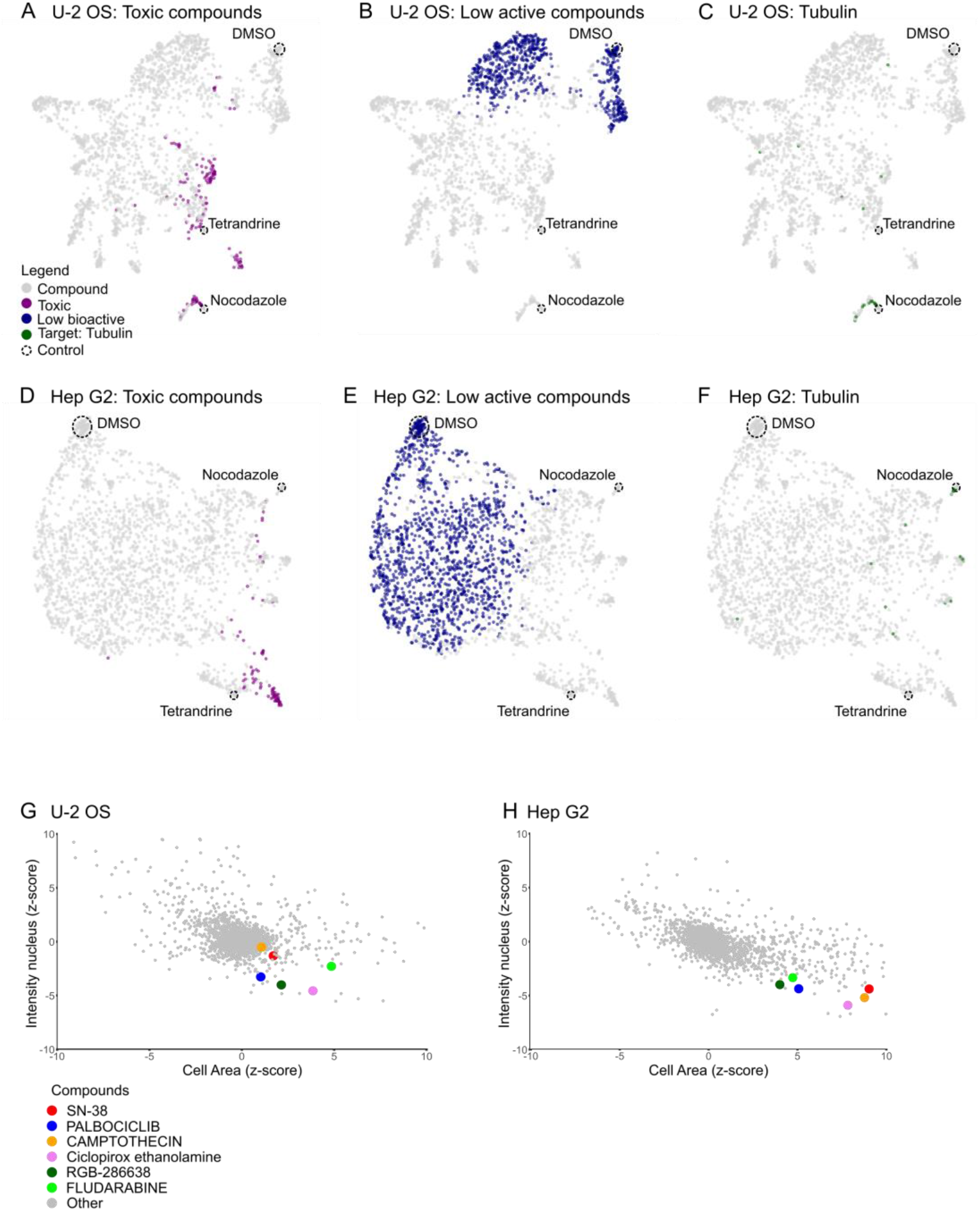
Visualization of morphological feature space based on FMP datasets. (A-C) Visualizations of morphological feature space using UMAP based on U-2 OS cells after feature reduction and filtering of non-reproducible compounds. The location of the controls (DMSO, Tetrandrine and Nocodazole) are labelled using a dashed circle. Color labeling of lower active (A), toxic (B), and compounds annotated for Tubulin as target (C). (D-F) UMAP of morphological profiles based on Hep G2 cells after feature reduction and filtering of non-reproducible compounds, with labeling of low activity (D), toxic (E), and compounds with an annotation for Tubulin as target (F). (G-H) Assessment of specific parameters indicative of cellular senescence based on U2 OS (G) and Hep G2 cells (H).

The visualization based on the U-2 OS cells reveal that toxic compounds are spread over a large part of the feature space. Toxic compounds are closely associated with both positive controls in particular with Nocodazole (**Figure 5A**). This observation confirms the toxic effect of both Nocodazole as well as Tetrandrine treatments previously observed based on the cell numbers (**Figure 3B**). The negative control DMSO is associated with lower active compounds (**Figure 5B**). From the visualization it appears that there could be two sets of lower active compounds. A potential cluster associated with the DMSO control and other lower active compounds not associated with the negative control. However, this split could be also artificial as dimensionality reduction as UMAPs are known for not necessarily preserving global structure perfectly (Huang et al. 2022). It is however conceivable that some compounds could indeed be without any measureable activity in the tested cell line. While compounds with activity likely will exhibit a gradient of activity ranging from a very small to a very large impact on the given cell line. Finally, compounds with at least one target annotated against Tubulin are in the same cluster to both the positive control Nocodazole (**Figure 5C**) and an accumulation of compounds labeled as toxic (**Figure 5A**). In the case of U-2 OS cells it is therefore very likely that the morphological profiles pick up on the very distinct morphology of dead or dying cells (**Suppl. Figure 1B**). This means that many compounds in this cluster might not be necessarily specifically acting against tubulin, as severe cytotoxicity produces similar strongly correlating morphologies via diverse mechanisms that are not associated with specific MOAs (Dahlin et al. 2023).

For the morphological profiles in the Hep G2 we can see that the overall smaller number of toxic compounds are located over a smaller area (**Figure 5D**). In contrast to the result in the U-2 OS cells, the positive control Nocodazole is not associated with any toxic compounds and forms a separate cluster with many compounds labeled with Tubulin as target (**Figure 5F**). Thus, in the Hep G2 cells the compounds that cluster actually might show a phenotype related to a MOA against Tubulin, rather than being a general cytotoxic phenotype. In the Hep G2 cells, many compounds are labeled low active (**Table 2**). However, similar to U-2 OS some low active compounds fall tightly around the negative control DMSO with many extending further away (**Figure 5E**). Again this is indicating that a smaller number of compounds might have no activity, while the majority of compounds in this category has some measureable activity in Hep G2.

Alternatively, to the analysis of the entire feature space using unsupervised machine learning we can also use specific features for measuring distinct cellular phenomena, such as cellular senescence. Cellular senescence is an arrest in the cell cycle with cells entering a stage without cell division (Hayflick 1965; Kuilman et al. 2010). The morphological hallmarks are increased cell and nuclear size (Cristofalo and Pignolo 1993) with abnormal nuclear morphology, in particular a decrease in density of the DAPI signal (Zhao et al. 2010). We thus can use the intensity nucleus as well as the cell area measurements as a robust read out of cellular senescence. With this we can show that compounds known to induce cellular senescence (Petrova et al. 2016; Herrmann et al. 2021; Lukasova et al. 2019; Wang et al. 2023) indeed have increased cell size combined with a decrease in nuclear intensity in Hep G2 cells and to a lower extent in U-2 cells (**Figure 5G & H**).

## Discussion

We presented here a comprehensive Cell Painting dataset based on the EU-OPENSCREEN Bioactive Compound Set. The data were acquired from four different imaging sites and using two different cell lines. The advantage of this well characterized compound set lies in the diversity of selected compounds in terms of their chemical space, as well as well-annotated biological effects, including MOAs and targets. Such comprehensive annotation should not only facilitate downstream analysis and applications, such as MOA identification through unsupervised techniques (e.g. clustering) and other machine learning approaches (Caicedo et al. 2018; Moshkov et al. 2022; Perakis et al. 2021), but more importantly, it also serves as a crucial reference for future morphological characterization of unknown compounds.

Our datasets have been generated with a full set of replicates across multiple imaging sites in three European countries. The data generation was preceded by an extensive selection, validation and protocol optimization process towards establishing common processes across these different laboratories. Care was taken to standardize particularly on cell culture material. A systematic comparison revealed that these datasets exhibit high reproducibility and quality within the data produced at each imaging site, comparable to other published datasets (Chandrasekaran et al. 2022; Cimini et al. 2022). We envision that this dataset will not only be useful for assessing the influence of different technical confounding factors on the morphological profiles but can also be effectively used to develop methods for normalizing datasets acquired from different imaging sites. The knowledge gained will be important for developing strategies to better merge and analyze data from multiple sources, which is currently a major challenge in the field (Arevalo et al. 2024).

The dataset is focusing primarily on the Hep G2 cell line, which is an established cellular model in high throughput screening (O’Brien et al. 2006). Hep G2 is a liver cancer cell line and thus aside from its relevance for liver diseases it is highly relevant as a cellular model for studying the toxic effect of small compounds (Schoonen, de Roos, et al. 2005; Schoonen, Westerink, et al. 2005). To allow a comparison of our Cell Painting approach with other existing datasets, we further generated a dataset based on the U-2 OS cell line, as many datasets in the community have been generated using this cell line. We provide a brief qualitative and quantitative comparison between the morphological effects of the compounds on the different cell lines (**Figure 4 & 5**).

We found that the Hep G2 cell line is overall less sensitive to toxic effects of compounds compared to U-2 OS at the relatively high used concentration of 10 µM (**Table 2**, **Figure 4B**). Important to note is that toxicity has been assessed based on the overall cell number per well and does not reflect a comprehensive assessment of toxicity. In addition we have observed that the tested compounds at the given concentration show overall a smaller morphological response in Hep G2 (**Figure 4C**). Furthermore, the cell line due to its cellular morphology, particularly its compact clustered growth, can pose a challenge for the image analysis and particularly feature extraction. This in part could also explain the observed lower overall activity of the tested bioactive compounds at the used concentration (Heinrich et al. 2023).

When visualizing the feature space using UMAPs we were able to delineated compounds annotated as Tubulin modulators in the Hep G2 cell line closely associated the positive control Nocodazole, a known disruptor of microtubule assembly/disassembly (De Brabander et al. 1976) (**Figure 5F**). Interestingly, the same analysis U-2 OS Tubulin modulators were closely associated with many compounds lacking this specific MOA and rather showing a general cell toxic effect (**Figure 5C**). This highlights that small chemical compounds can produce varying effects across different cell lines and emphasizes the need to perform Cell Painting in multiple cell lines for a more comprehensive and robust characterization.

The EU-OPENSCREEN Bioactive Compound Set is part of much larger compound library. The ECBL is an open source compound library based on more than 100,000 commercially available compounds designed to cover a larger and diverse chemical space. Most of the compounds within the ECBL are poorly characterized. Furthermore, EU-OPENSCREEN is collecting novel compounds synthesized from academic researchers from around the world in the European Academic Compound Library (EACL). Currently, this Cell Painting consortium is applying the presented Cell Painting approach to both the ECBL as well as the EACL. On the one hand, the presented dataset based on the Bioactives compounds served as an important milestone to the consortium to validate the approach and the feasibility to apply it on a much larger compound set. On the other hand, the very well characterized Bioactive dataset will serve as an important foundation and reference map for discovery of novel compound properties and MOAs within the ECBL and EACL.

Finally, the Cell Painting project is part of a much larger effort for a comprehensive characterization of the provided compounds based on the EU-OPENSCREEN Bioprofiling project. Here general physico-chemical properties such as solubility, light absorbance, and fluorescence as well as biological properties such as cell viability, anti-bacterial, and anti-fungal are tested using common assay panels. The data produced by Cell Painting and the wider Bioprofiling project will be provided open source to the scientific community in public data repositories such as the Cell Painting Gallery (Weisbart et al. 2024) and Bioimage Archive (Hartley et al. 2022) with the morphological profiles and other numeric data fully integrated in dedicated databases (ECBD). This data will provided a rich source for powerful computational approaches (Sanchez-Fernandez et al. 2023) that promise to unlock the hidden potential of many small chemical compounds and thereby will accelerate early drug discovery.

## Methods

### Bioactive Set

The EU-OPENSCREEN Bioactive Compound Set comprises 2,464 compounds selected utilizing data from the Probes & Drugs Portal (P&D) (Skuta, Southan, and Bartunek 2021) ver. 07.2018. (Skuta et al. 2017). P&D is a hub for the integration of high-quality bioactive compound sets with a focus on chemical probes and drugs. The set was created by the combination of manually selected high-quality chemical tools, such as chemical probes from the Structural Genomics Consortium [Structural Genomics Consortium, https://www.thesgc.org/] or Chemical probes portal (Antolin et al. 2023), and an automatically generated set of, predominantly, target-selective compounds (prioritizing such with known mechanism of action), in order to achieve wide proteome coverage. The EU-OPENSCREEN Bioactive Set contains 385 compounds labeled as chemical probes and 681 as approved drugs. It is also a part of the P&D compound set list (probes-drugs.org/compoundsets) and therefore, can be accessed/worked with at P&D (https://www.probes-drugs.org/compounds/standardized#compoundset=353@AND). The full list of compounds with basic annotations is a part of the supplementary material (**Suppl. Table 5**).

### Cell culture

Hep G2 cells (ATCC, HB-8065) were cultured in Roswell Park Memorial Institute Medium (RPMI 1640, Gibco, 61870044) supplemented with 10% (v/v) fetal bovine serum (Sigma Aldrich, S0615). U-2 OS cells (DSMZ, ACC785) were cultured in Dulbecco’s Minimum Essential Medium (DMEM, Gibco, 61965026) also supplemented with 10% FBS. The cell lines were tested for Mycoplasma using a luminescence-based MycoAlert kit (Lonza, LT07-418) and maintained at 37 °C under 5% CO2. When cells reached a confluence of 70 to 80%, cells were washed with DPBS (Gibco, 14190250), dissociated with Trypsin-EDTA, 0.05% (Gibco; 25300025) and reseeded into a new cell culture flask with fresh complete medium or seeded for experiments.

### Cell seeding and compound transfer

Hep G2 and U-2 OS cells are counted and seeded into 384 well plates (PhenoPlate 384-well, Perkin Elmer, 6057328) using a Biotek or Multidrop microplate dispenser. Cells are seeded in a volume of 40 µl per well with 2000 cells/well and 700 cells/well, respectively and kept at room temperature for at least 30 min to aid homogeneous spreading. Plates were incubated for 24 h at 37 °C at 5% CO2 atmosphere to allow for cell attachment and propagation. Compounds were transferred from library plates to cell plates using Echo 650 acoustic dispenser or Biomek I7 liquid handler. Plates were incubated for another 24 h at 37 °C at 5% CO2 atmosphere.

The EU-OPENSCREEN Bioactives Compound Set comprises seven 384 well plates with randomly distributed compounds located in column 1-22. Controls are located in column 23 and 24, which include DMSO (0.1%) as vehicle control as well as Nocodazole (5 µM) and Tetrandrine (5 µM) as reproducibility control as they showed strong consistent profiles even over different cell lines. Compound screen was performed at 10 µM in four replicates. For the replicates we used a new independent cell seeding event (biological replicate).

### Cell Painting

The Cell Painting staining protocol is based on (Bray et al. 2016) with minor modifications. The protocol was adjusted by each site according to the available equipment. A list of all devices used by each site can be found in **Table 2**: **Devices for cell staining**. In general, the medium was first aspirated from the plates to a residual volume of 10 µl. Subsequently 30 µl of MitoTracker (Invitrogen, M22426) solution in pre-warmed medium were added to the cells with a final concentration of 500 nM and incubated for 30 min at 37°C. For fixation, the MitoTracker solution was removed and replaced with 30 µl paraformaldehyde (4%, Roth, 0335) and incubated in the dark at room temperature (RT) for 20 min. After fixation, cells were washed with 70µl PBS and permeabilized by adding 30 μL of 0.1% (v/v) Triton X-100/PBS (Sigma Aldrich, T8787) solution to each well for 20 min at RT. Triton X-100 was removed followed by two washes with 70 µl PBS. Cells were then stained for 30 min at RT in the dark with of the staining solution containing HOECHST 33342 (Invitrogen, H3570), SYTO14 green (Invitrogen, S7576), Concanavalin A/Alexa Fluor 488 (Invitrogen, C11252), Wheat Germ Agglutinin/Alexa Fluor 555 (Invitrogen, W32464) and Phalloidin/Alexa Fluor 568 (Invitrogen, A12380) in PBS with 1% (m/v) BSA (Sigma Aldrich, A7030). Adding 30 µl of the staining solution to each well resulted in the final well concentration of 4 μM HOECHST, 25 μg/ml Concanavalin A, 3 μM SYTO14, 1U/ml Phalloidin and 1.5μg/ml Wheat germ agglutinin. Finally, cells were washed three times with 70 µl PBS and sealed with adhesive foil. Plates were stored in the dark at 4°C until image acquisition.

### Assay optimization and standardization

We employed an extensive evaluation and validation process to select suitable imaging sites for performing the cell painting assay and achieve standardization and comparability.

#### First validation phase: Selection of imaging sites

An initial protocol based on an established Cell Painting Protocol was first optimized at the FMP site (**Suppl. Methods: Cell Painting assay**). We then invited imaging sites in an open call to submit a proposal and validation data. This initial proposal required to include an estimation of costs and duration as well as information about the available instrumentation and a track record of performing automated screening assays.

To generate the validation data the developed protocol with the standardized staining and cell culture protocol was shared with imaging sites. To achieve further standardization and optimal comparability across the sites key reagents were shared. Reference compounds were prepared and aliquoted from the same lot and distributed to all sites. The same cell culture serum lot was acquired and provided to all sites. Finally the cells were prepared and distributed by the FMP site.

The evaluation of the proposals and the validation data was performed in two stages based on pre-defined criteria by two external reviewers (**Suppl. Methods: Evaluation Criteria, Evaluation scoring**). The reviewers were selected based on their expertise in the field of image-based screening and early drug discovery. In the first stage the provided data was evaluated based on how well the candidate sites implemented the protocol, the overall quality of the generated data, as well as for intra-as well as inter-plate consistency (**Suppl. Methods: Evaluation stage 1**). To pass the first evaluation more or equal to 65% of the first selection criteria needed to be reached. In the second stage, the candidate sites that passed the first criterion were evaluated for assay cost, the duration for screening 100,000 compounds, clarity of the proposal as well as a track record (**Suppl. Methods: Evaluation stage 2**). The score of the assay validation was considered with a factor of 10% in the second evaluation. Based on the participant’s final score four imaging sites were selected for performing the assay on the EU-OPENSCREEN Bioactive Compound Set.

#### Second evaluation phase: Protocol optimization

To reduce variability across sites we implemented additional measures and experiments. We reasoned that the main sources of variability observed could originate from differences in compound preparation, the cell culture, staining methods, or microscopy (Arevalo et al. 2024).

To assess the effect of the microscopy on the result of our Cell Painting assays we performed an additional validation experiment. The FMP site shared a fully stained plate with the other three partner sites. We further developed a microscopy guide for the imaging sites so that the final images were in the same intensity range in each channel. In this experiment, we found no improvement in comparability across sites. To address any variability stemming from differences in compound preparation we distributed pre-prepared assay-ready plates (compound plates). This also did not improve the comparability. Thus differences in cell culture as well as staining drive this variability.

To exclude variability due to cell culture and staining material, we acquired, prepared and distributed the same serum lot, cells, and lots of fluorescent dyes centrally.

### Image acquisition

Cell images were acquired using automated confocal microscopes equipped with water-immersion 20x objectives (1.0 NA). Offsets were determined for each cell line and kept constant throughout the experiment. For each well of the 384 well plate, nine fields in a 3 × 3 array, located in the center of the well, were imaged using 2x binning and four fluorescence channels to capture HOECHST 33342, Concanavalin A and SYTO14, Wheat Germ Agglutinin and Phalloidin, as well as MitoTracker. Excitation and emission wavelengths of these four channels vary based on the imaging system used by the different partner sites (**Table 3**).

**Table 3:** Image acquisition settings.

|  | FMP<br>(Opera Phenix) |  | IMTM<br>(Yokogawa CV8000) |  | Medina<br>(Operetta CLS) |  | USC<br>(Operetta CLS) |  |
| --- | --- | --- | --- | --- | --- | --- | --- | --- |
|  | Excitation | Emission | Excitation | Emission | Excitation | Emission | Excitation | Emission |
| HOECHST 33342 | 375 nm | 435-480 nm | 405 nm | 400-490 nm | 405 nm | 430-500 nm | 370 nm | 430-500 nm |
| Concanavalin A + SYTO14 | 488 nm | 500-550 nm | 488 nm | 500-550 nm | 475 nm | 500-550 nm | 475 nm | 500-550 nm |
| Phalloidin + Wheat germ Agglutinin | 561 nm | 570-630 nm | 561 nm | 563-637 nm | 550 nm | 570-650 nm | 550 nm | 570-650 nm |
| MitoTracker Deep red | 640 nm | 650-760 nm | 640 nm | 647-705 nm | 630 nm | 655-760 nm | 630 nm | 655-760 nm |

### Image feature extraction

We used CellProfiler version 4.1.3 (Stirling et al. 2021) with the JUMP analysis pipeline version 3 (Chandrasekaran et al. 2022), original available here: https://github.com/broadinstitute/imaging-platform-pipelines/tree/master/JUMP_production) for feature extraction. A total of 2977 features were extracted from each segmented cell in all fields per well. The pipeline was executed in parallel on 5 nodes of our high performance computing cluster (HPC) (**Table 4**).

**Table 4:**
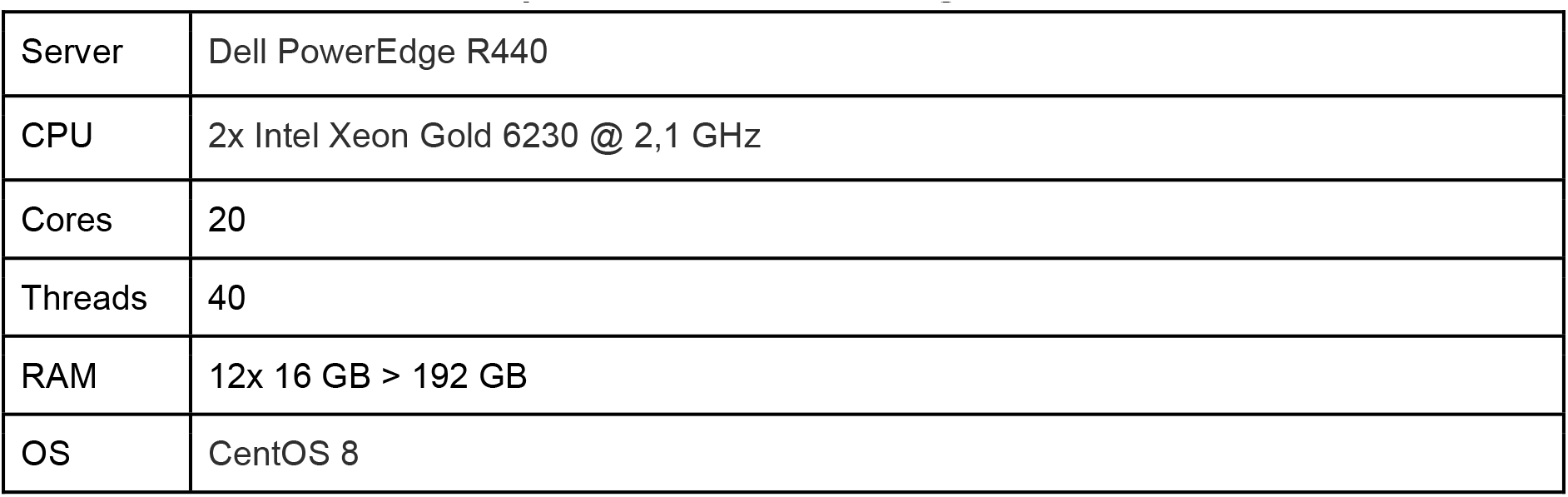
Single node of high performance computing cluster.

|  |  |
| --- | --- |
| Server | Dell PowerEdge R440 |
| CPU | 2x Intel Xeon Gold 6230 @ 2,1 GHz |
| Cores | 20 |
| Threads | 40 |
| RAM | 12x 16 GB > 192 GB |
| OS | CentOS 8 |

We adjusted the feature extraction pipeline for our four channel dataset (DNA, ER, AGP and Mito). The illumination function was calculated on the individual channels with a median filter set to a kernel size of 20 px. The function was computed on all images across cycles with rescaling. The illumination function was applied using a division. No images were removed based on the image quality control of the CellProfiler workflow.

Nuclei in the DNA channel were segmented with a global minimum cross-entropy threshold using a threshold smoothing scale of 1 and a threshold correction factor of 1. The lower and upper bounds of the threshold were set to 0.005 and 1.0 respectively. No log transform of the image intensity values was performed before thresholding. The shape of the objects was used to separate objects with a smoothing filter using a radius of 10 px applied before separation. Local maxima with a distance smaller than 8 px were suppressed. Holes were filled after de-clumping. Nuclei with a diameter of 15 px - 90 px were kept in U-2 OS and Hep G2 cells.

Cells were segmented based on the ER channel using a marker-controlled watershed using the segmented nuclei as input. A global intensity based threshold using the Otsu thresholding method was used to compute a three class threshold, assigning the pixels of the middle class to foreground. No smoothing was applied to the images. The threshold correction factor was set to 0.7. The lower and upper bounds of the threshold were fixed to 0.005 and 0.6 respectively. A log transformation was applied before thresholding. Cell and nuclei objects that were touching the image border were filtered for some of the feature computation. To create a mask of only the cytoplasm the nuclei were subtracted from the cell mask.

The following image features were computed based on the AGP, DNA, ER, and Mito channel within the nuclei, cell, and cytoplasm mask. All correlation metrics with a threshold of 15% of the maximum intensity. The Faster method was used to compute a Costes thresholding. Granularity was measured with a subsampling factor of 0.5 and a subsampling factor of 0.5 for background reduction. The radius of the structuring element was set to 10 and the range of the granular spectrum set to 16. The intensity in the illumination corrected images was measured. The intensity distribution within the objects was computed by scaling the bins and using 4 bins.

Textures were measured in nuclei, cells and cytoplasm in the DNA, ER, Mito channel using 256 gray level bins with a scale of 3, 5, and 10. Size and shape of nuclei, cells, and cytoplasm including Zernike features were extracted. The number of cell neighbors was measured within a 5 px distance including cells that were touching the image border. Further the number of all adjacent cells were measured including cells that were touching the image border. The number of nuclei neighbors were measured within a 1 px distance including nuclei that were touching the image border. For the Mito channel the Neurites feature type score was computed enhancing the tubeness enhanced Mito channel with a smoothing scale of 1. Based on the tubeness enhanced Mito image, the intensity distribution of cells with nuclei as center, the cytoplasm with nuclei as center was computed by scaling the bins and using 16 and 20 bins with a maximum radius of 200 px.

On the tubeness enhanced Mito channel a global minimum cross-entropy threshold was applied using a smoothing scale of 1.3488 and a correction factor of 1.0. The lower and upper bounds for the threshold were set to 0.0 and 1.0 respectively. No log transform was performed before the thresholding. On the mask of the Mito channel a skeletonization algorithm was applied with filling in small holes of maximum 10 px. Finally, skeleton features were extracted.

Overall image intensities were extracted from the illumination corrected images. Background in each channel was computed in the image content outside of the segmented objects.

### Single cell filter

Before aggregation, the measurements for individual cells were filtered to remove cells with any missing or infinite values. Furthermore, an HBOS filter was performed to remove objects that had a feature vector that was different from the distribution of features over the entire plate (Rezvani, Bigverdi, and Rohban 2022). In our tests, this was well suited to particularly remove artifacts in segmentation as well as dead cells. The outlier selection was computed with the HBOS function of pyod version 1.0.9 (Zhao, Nasrullah, and Li 2019) with python version 3.9.16. The histogram was computed using a static number of bins of 10, an alpha of 0.1, the flexibility parameter (tol) set to 0.5, and the proportion of outliers (contamination) set to 0.1. The HBOS model was then applied to classify each individual feature vector into outlier and non-outlier.

After the outlier, missing and infinite values filters, median values were computed for each well based on the up to nine fields per well. Note that for some fields no segmentation was achieved or they were removed in the filter step before data aggregation. Some particular toxic compounds had no extracted features over all replicates; these were counted towards the toxic compounds number.

### Data processing and analysis

#### Normalization, feature reduction and profile aggregation

We used pycytominer version 0.2.0 (Serrano et al. 2023) in python 3.9.16 on Ubuntu 22.04.3 LTS for further processing of the profiles. The image features per well were normalized per plate to the DMSO controls using the robust median absolute deviation function (mad_robustize). The epsilon value was set to the default value of 1e-06. Feature reduction was performed by first removing columns with NaN values. Then features with a low variance were removed via a variance frequency cut-off of 0.1 and a variance unique cut-off of 0.1. Feature outliers were removed with the outlier cut-off set to 100. Finally, features with high correlation were reduced using a correlation threshold of 0.9. For aggregating the profiles over the four replicates the median function was used.

#### Toxicity filter

We determined toxic compounds as these have been shown to produce highly similar features with unspecific MOAs (Dahlin et al. 2023). We first compute the consensus median cell number for each well per plate over the four replicates. We then defined compounds as toxic with per well consensus cell count smaller than the median of the population of the entire dataset subtracted by 2.5 standard deviations of the population.

#### Percent replication

The percent of replicating compounds was computed based on the precent replication score developed in the JUMP-CP consortium (Way et al. 2021). For each compound the pairwise Pearson correlation over all available replicates was determined and a median replicate correlation computed. To compute a null distribution, 10000 random samples of four randomly chosen compound feature vectors (non-replicates) were drawn from the dataset. The pairwise Pearson correlation of these feature vectors were computed and used to calculate a median non-replicate correlation. Compounds were defined as replicating if their median replicate correlation was more than 95% of the null distribution computed from the samples of the median non-replicate correlation.

#### Activity filter

The percent of replicating compounds over the compound set after the toxicity filter was initially ranging from 60.1 to 77.5% (**Suppl. Figure 6 A-D**). We also observed that many of the compounds in both cell lines gave very small responses in their morphological profiles compared to the DMSO negative control. We suspected that many of the compounds with small phenotypic response in the specific cell line also exhibit low reproducibility. To determine compounds with lower activity we applied an induction filter (Christoforow et al. 2019). To determined induction first median features per compound over the four replicates were computed. Features were defined responding when deviating three times from the median absolute deviation from the median of the DMSO controls. For each compound the fraction of active features was then computed and a threshold of 5% applied to define a compound with an active response.

#### Plate quality controls

The aggregated data files containing the median values for each well per plate were further aggregated into a single dataset using a KNIME workflow (Berthold et al. 2009). For visualization of plate artifacts five CellProfiler features were selected: Metadata_Object_Count, reflecting the number of detected cell objects. Further the mean fluorescence intensity values of following compartments and stains, reflecting the dispense quality of the dyes: Nuc_Intensity_MeanIntensity_DNA (HOECHST 33342), Nuc_Intensity_MeanIntensity_ER and Cyto_Intensity_MeanIntensity_ER (Concanavalin A & SYTO14), Cyto_Intensity_MeanIntensity_AGP (Phalloidin & Wheat germ agglutinin), Cyto_Intensity_MeanIntensity_Mito (Mitotracker Deep red).

The data for each plate and feature was visualized (McInnes et al. 2018) using heat maps that were generated within a KNIME workflow using R snippets and the R library “ggplot2”. KNIME version 5.1.0, R version 4.3.1, R ggplot2 package version 3.4.3 was used. The generated plots were then transferred into the integrated reporting tool of KNIME, and assembled into a single printable report file for each dataset. Heatmaps of Metadata_Object_Count were scaled as follows: the global maximum and median were determined, and rounded up to two significant digits. The color yellow was assigned to the median object count, blue to the maximum, red to zero objects. This way it is possible to spot differences in absolute cell number that was dispensed across the different plates. Heatmaps of the mean fluorescence features were scaled as follows: for each plate, the values were divided by their median for data normalization, since we do not have plate controls at hand that are specific for those features. The color yellow was assigned to the median value, blue to 1.5 fold the median value, and red to 0.5 fold the median value.

The plots of the four technical replicates were combined on a single report page for each screened library plate. The median Z-score values of these four technical replicates was added as a fifth heatmap. This way, it is possible to spot whether taking the median of the replicates would reduce plate artifacts that are visible on individual technical replicates.

#### Percent paring

For comparing the profiles between the two cell lines, the percent replication metric was computed over the corresponding compounds after toxicity, activity and reproducibility filtering. We used the 427 overlapping features of the cell lines after feature reduction (**Figure 4A**). To differentiate this metric from the percent replication metric we changed the name of this analysis to percent pairing. The null distribution was computed over randomly selected compound pairs. The same compound pairs over the two different cell lines were defined as pairing if above the 95% correlation threshold defined on the null distribution.

#### Dimensionality reduction

We used Uniform Manifold Approximation and Projections (UMAP)(McInnes et al. 2018) to create visualizations of the morphological feature space in 2D. For the batch control quality figures, we used the data filtered for toxic, lower activity and non-replicating compounds. To perform the UMAPs for the batch quality control we used the default settings using the Euclidean metric, with the number of neighbors set to 15 and the minimum distance set to 0.1. The data points in the visualizations were then labeled by their replicate or plate number.

For the analysis of the morphological features space, we used the data only filtered for non-replicating compounds leaving in the toxic as well as the lower active compound. The number of neighbors was set to 30 with a minimum distance of 0.1. We then labeled on the UMAP visualizations the control compounds (DMSO, Nocodazole and Tetrandrine). The data points were then labeled for toxic as well as lower active compounds. For the basic MOA analysis for compounds, acting against Tubulin we did a basic search and asked if a single target in the annotated targets of any compound contained the string ‘Tubulin’.

#### Senescence analysis

For the analysis of cellular senescence, we extracted the cell area as well as the nuclear intensities for all compounds of each FMP dataset. For this analysis we computed Z-Scores based on the median and median absolute deviation of the entire plate. The normalized parameters were then plotted against each other in a 2D dot plot. Finally we labeled compounds that are known to induce cellular senescence (Petrova et al. 2016) (e.g. PALBOCICLIB, CAMPTOTHECIN and SN-38 an active metabolite of a CAMPTOTHECIN analog) on these plots.

#### Image figures

Image figures were prepared with established processing and image visualization standards (Schmied and Jambor 2020; Schmied et al. 2023) using Fiji (Schindelin et al. 2012). In detail for the image figures, randomly selected wells from the negative and positive controls were used. For Figure 2 C & D the fifth (middle) field of the selected well was used. For Supplemental Figure 1 & 2 the first field of the selected well was used. The individual images for each channel were corrected for illumination using the same method as the HPC CellProfiler cluster workflow. The lower bound of the brightness contrast function was then set in all treatments to an empirically determined camera background in the DMSO treated images within each individual channel. Three large rectangular ROIs were drawn in an area of the image without cells. The mean gray value was measured in the ROIs and an average was computed. The upper bound was adjusted based on the brightest treatment in each individual channel and applied over all treatments to ensure intensity values can be compared over the treatments within a single cell line (**Table 5 & 6**).

**Table 5:** Brightness contrast settings Figure 2.

| Site | Cell Type | Figure | Channel | Lower | Upper |
| --- | --- | --- | --- | --- | --- |
| FMP | U-2 OS | 2C | DNA | 65 | 1400 |
| FMP | U-2 OS | 2C | ER | 109 | 2000 |
| FMP | U-2 OS | 2C | AGP | 135 | 3000 |
| FMP | U-2 OS | 2C | Mito | 536 | 3000 |
| FMP | Hep G2 | 2D | DNA | 82 | 2500 |
| FMP | Hep G2 | 2D | ER | 141 | 8000 |
| FMP | Hep G2 | 2D | AGP | 144 | 4000 |
| FMP | Hep G2 | 2D | Mito | 552 | 4000 |

**Table 6:**
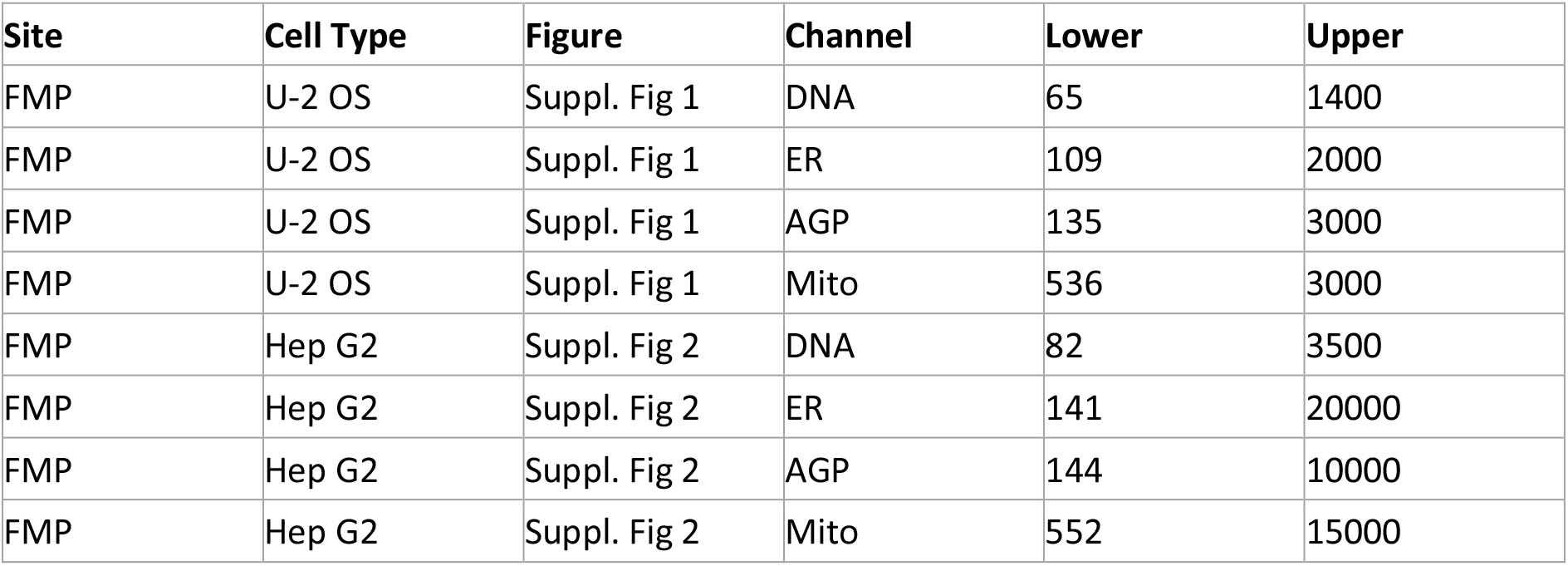
Brightness contrast settings Suppl. Figure 1 & 2.

After brightness and contrast adjustments, the images were converted to 8-bit and saved as PNG. For an overview the DNA, AGP and Mito channels were merged with Cyan, Magenta and Green LUTs respectively. A ROI for the inset was chosen in the middle of the field of view. ROIs as well as scale bars are shown on the overview images.

## Acknowledgements

We want to thank Sonja Sievers and Arnaud Ogier for their reviews in the first project evaluation phase. Beth Cimini for critical feedback for the project. Utesch Tillmann for HPC support and maintenance. Ankur Kumar and Erin Weisbart for support in uploading the data to the Cell Painting Gallery. The FMP IT Department, particularly Ingo Breng for support. Michael Ebner for critical feedback to the image figures. Edgar Specker for input towards the compound collection. Nathaniel Smith for lively discussion concerning the data analysis.

This project was supported by the Leibniz-Forschungsinstitut für Molekulare Pharmakologie via the Integrated Project titled: “Machine Learning Enhanced Cell Morphology Profiling in Molecular Pharmacology” awarded to Jens Peter von Kries, Han Sun and Christopher Schmied.

This project was funded by the German Federal Ministry for Education and Research under grant number AZA 16KX1816.

The selection of Bioactive compounds was supported by the Ministry of Education, Youth and Sports of the Czech Republic (LM2023052).

This work was supported by the EGI DataHub and EGI Check-in services from CYFRONET and GRNET, provided from the EGI-ACE project (Horizon 2020) under Grant number 101017567.

## Notes

https://doi.org/10.5281/zenodo.13309566

https://github.com/schmiedc/EU-OS_bioactives

https://cellpainting-gallery.s3.amazonaws.com/index.html#cpg0036-EU-OS-bioactives/

https://www.probes-drugs.org/compounds/standardized#compoundset=353@AND

### Competing Interest Statement

The authors have declared no competing interest.

### Summary of Updates

Fixed the broken link the Zenodo Repository in the article.

