## Supplementary material for "Morphological Profiling Dataset of EU-OPENSCREEN Bioactive Compounds Over Multiple Imaging Sites and Cell Lines": Suppl_Figures.docx

### Supplementary figures

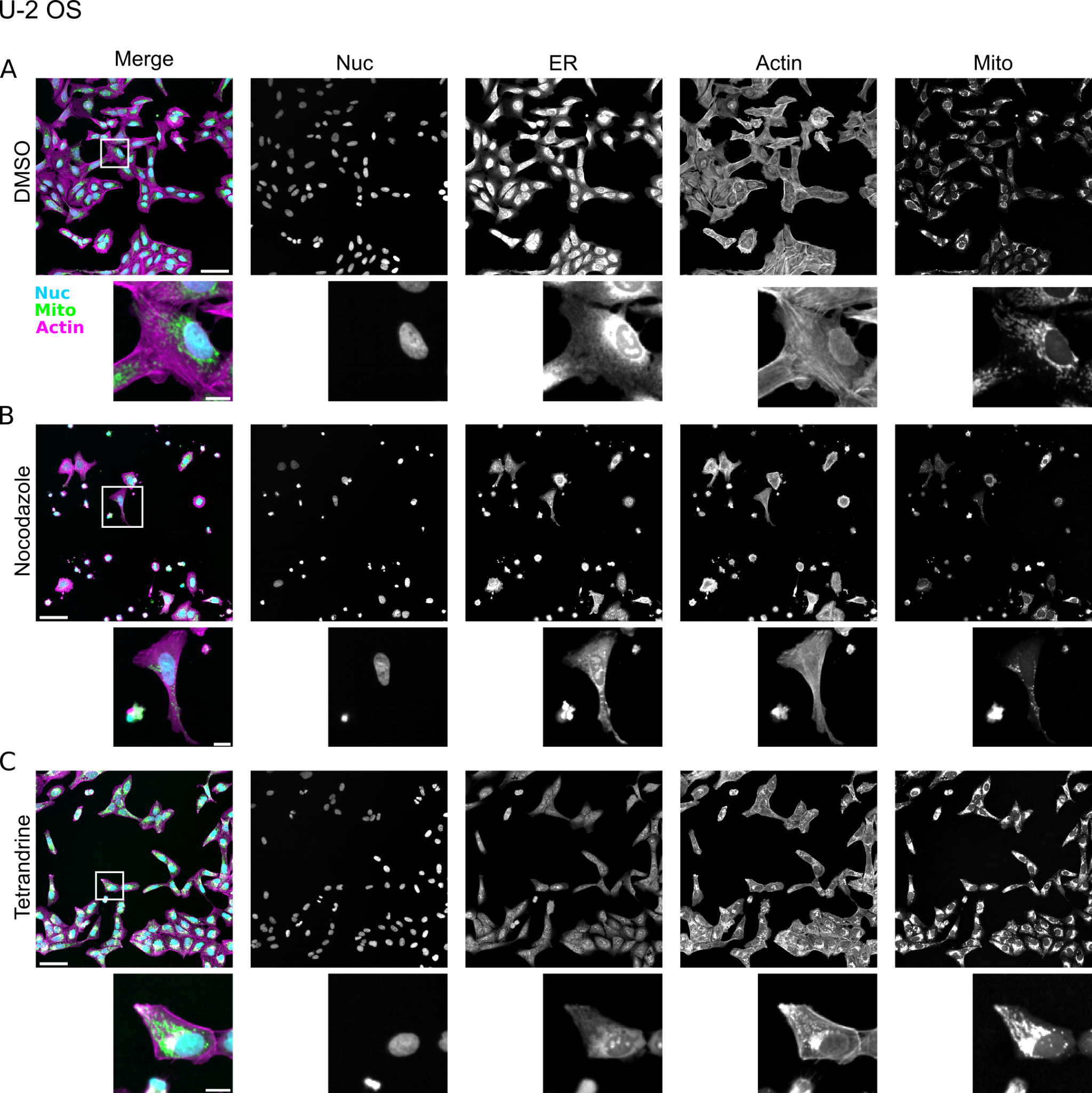

**Suppl. Figure 1: Examples of images of the positive controls of the FMP U-2 OS cell dataset.** (A) Example images of U-2 OS cells treated with DMSO. The merge is a composite of the nucleus (Nuc, cyan), mitochondria (Mito, green) and actin (Actin, magenta) channels. Gray scale images of the individual channels with crops of single cells from the middle of the field of view below. (B) Images of Nocodazole treated U-2 OS cells. Note many dead or dying cells in the overview images (small round cells with small bright nuclei). (C) Tetrandrine treated U-2 OS cells. (A-C) Scale bars in the merged panels correspond to 100 µm in the overview and 20 µm in the crop.

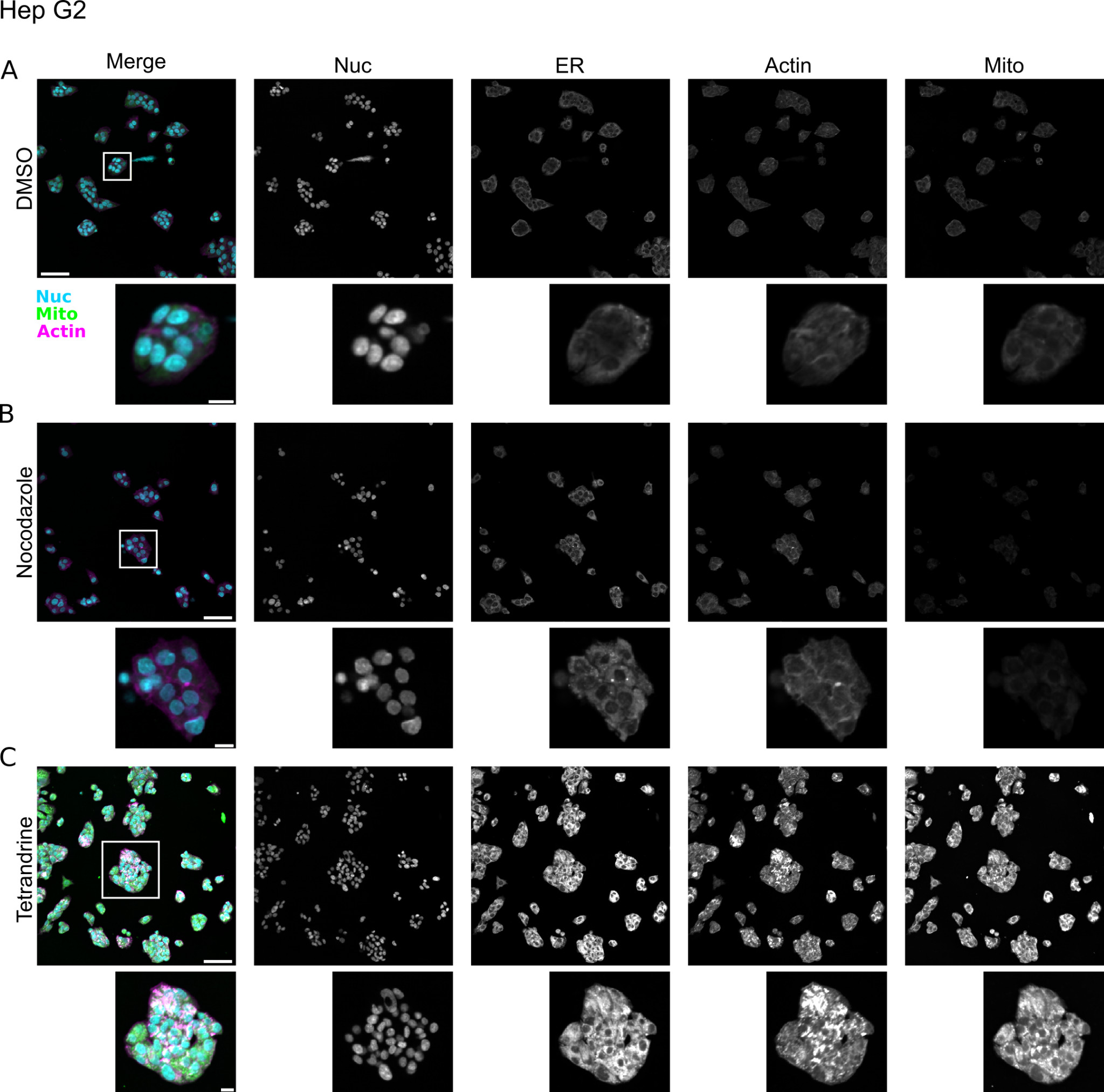

**Suppl. Figure 2: Examples of images of the positive controls of the FMP Hep G2 cell dataset.** (A) Example images of Hep G2 cells treated with DMSO. The merge is a composite of the nucleus (Nuc, cyan), mitochondria (Mito, green) and actin (Actin, magenta) channels. Gray scale images of the individual channels with crops of single cells from the middle of the field of view below. Note the dense clusters of relatively small cells that are typical for this cell line. (B) Images of Nocodazole treated Hep G2 cells. (C) Tetrandrine treated Hep G2 cells. (A-C) Scale bars in the merged panels correspond to 100 µm in the overview and 20 µm in the crop.

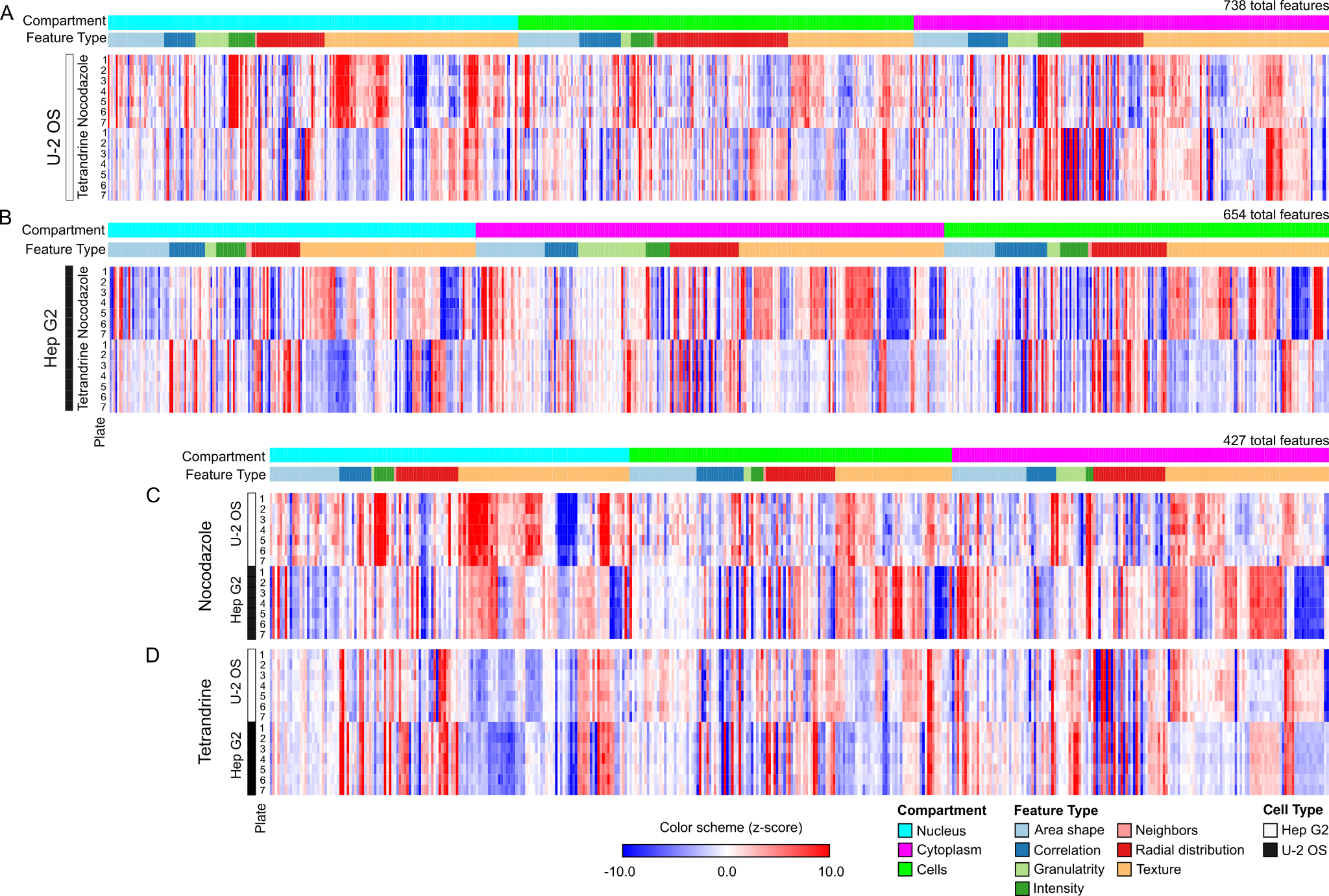

**Suppl. Figure 3: Cellpainting profiles of positive controls for the FMP dataset.** (A) Morphological profiles of positive controls Nocodazole and Tetrandrine over different plates after feature selection sorted by compartment and feature type in U-2 OS. (B) Morphological profiles over the seven plates of Nocodazole and Tetrandrine after features selection in Hep G2 cells. (C) Overlapping features between U-2 OS and Hep G2 cells treated with Nocodazole. (D) Overlapping features between U-2 OS and Hep G2 cells treated with Tetrandrine. Profiles visualized with Morpheus, <https://software.broadinstitute.org/morpheus>. The color LUT was set with a minimum z-score of -10 and a maximum z-score of +10 (See color scheme in legend).

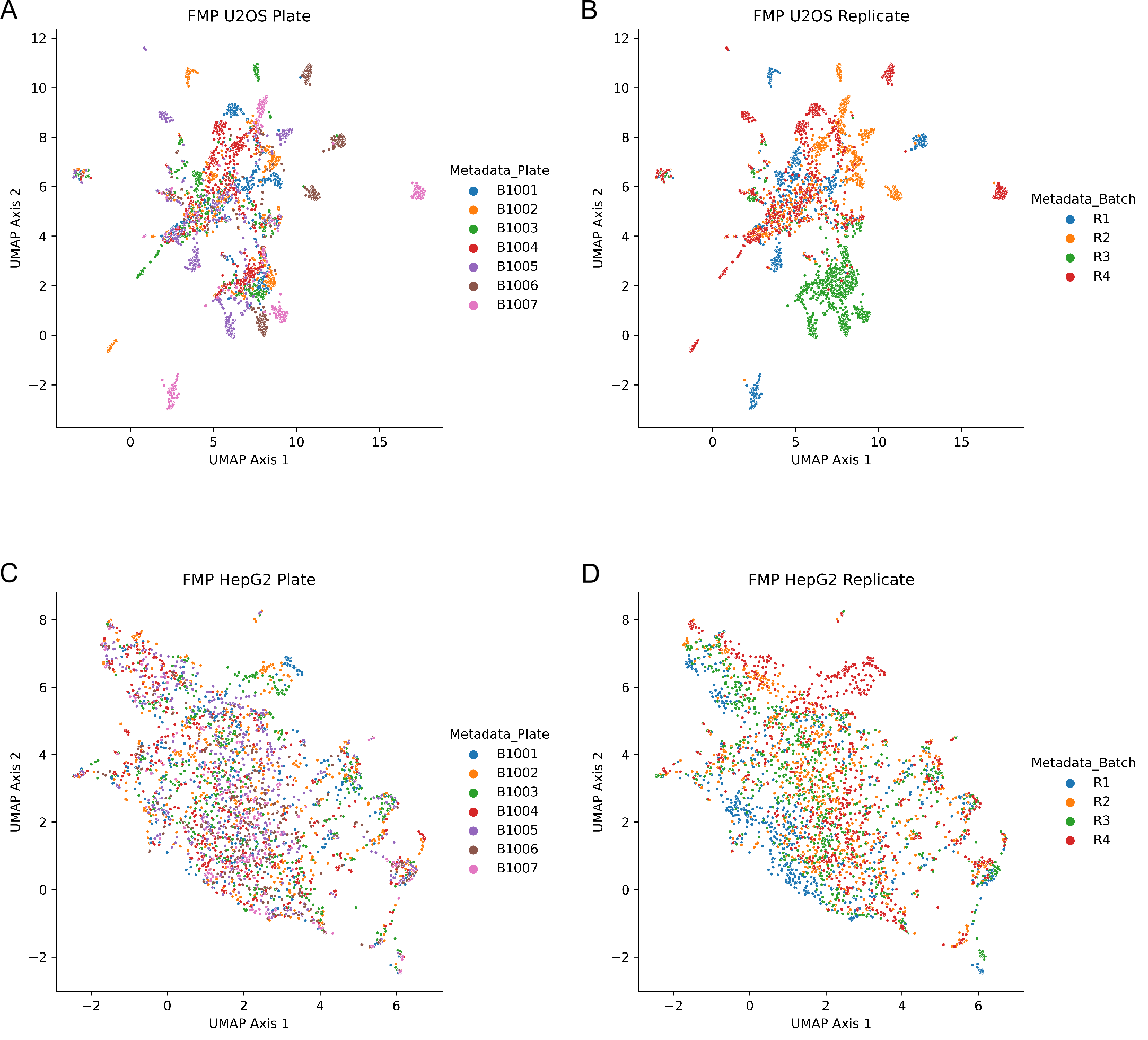
**Suppl. Figure 4: Batch effect quality control visualization based on FMP datasets.** (A-D) 2D UMAP for assessing plate and batch effects of the FMP datasets after normalization, feature reduction and compound filtering. Each point corresponds to a single well from an individual replicate. (A) U-2 OS dataset labeled per plate. (B) U-2 OS dataset labeled per replicate. (C) Hep G2 dataset labeled per plate. (D) Hep G2 dataset labeled per replicate.

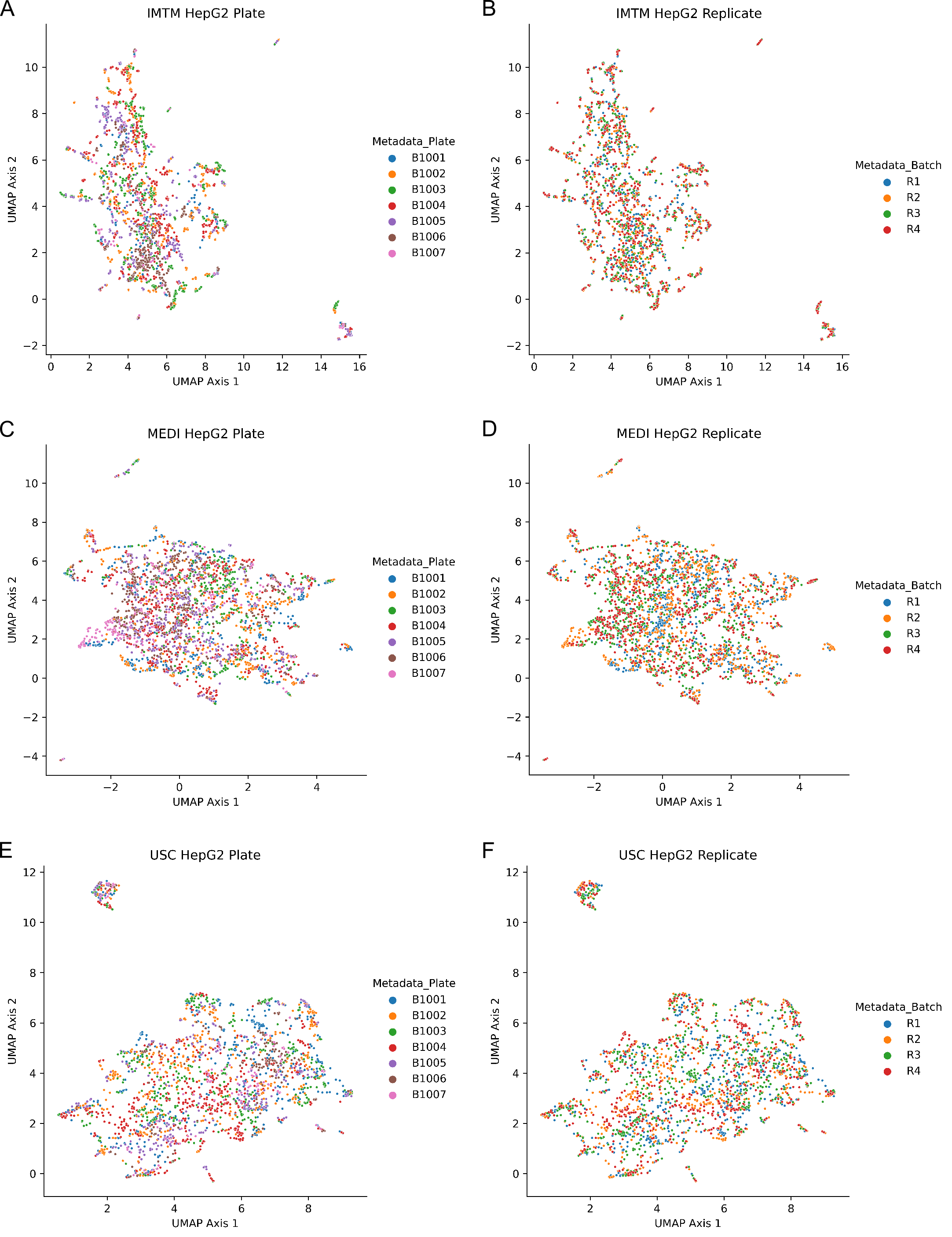

**Suppl. Figure 5: Batch effect quality control visualization based on IMTM, MEDI and USC datasets.** (A-F) 2D UMAP for assessing plate and batch effects of the datasets from IMTM, MEDI and USC after normalization, feature reduction and compound filtering. Each point corresponds to a single well from an individual replicate. (A) IMTM Hep G2 dataset labeled per plate. (B) IMTM Hep G2 dataset labeled per replicate. (C) MEDI Hep G2 dataset labeled per plate. (D) MEDI Hep G2 dataset labeled per replicate. (E) USC Hep G2 dataset labeled per plate. (F) USC Hep G2 dataset labeled per replicate.

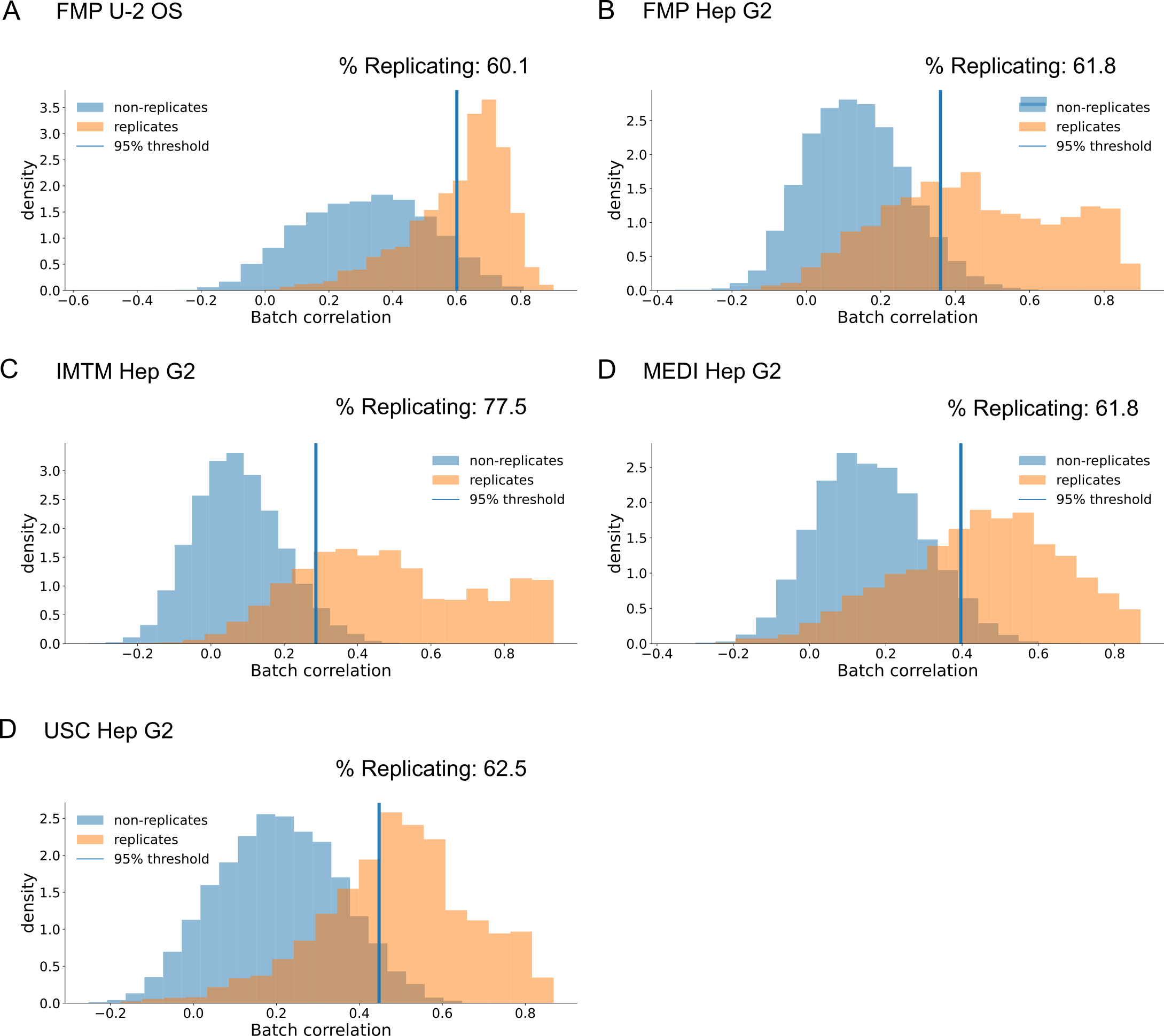

**Suppl. Figure 6: Percent replication before induction filter for the FMP dataset.** (A-D) percent replication of compounds of the U-2 OS and Hep G2 data from all the imaging sites before induction filter.

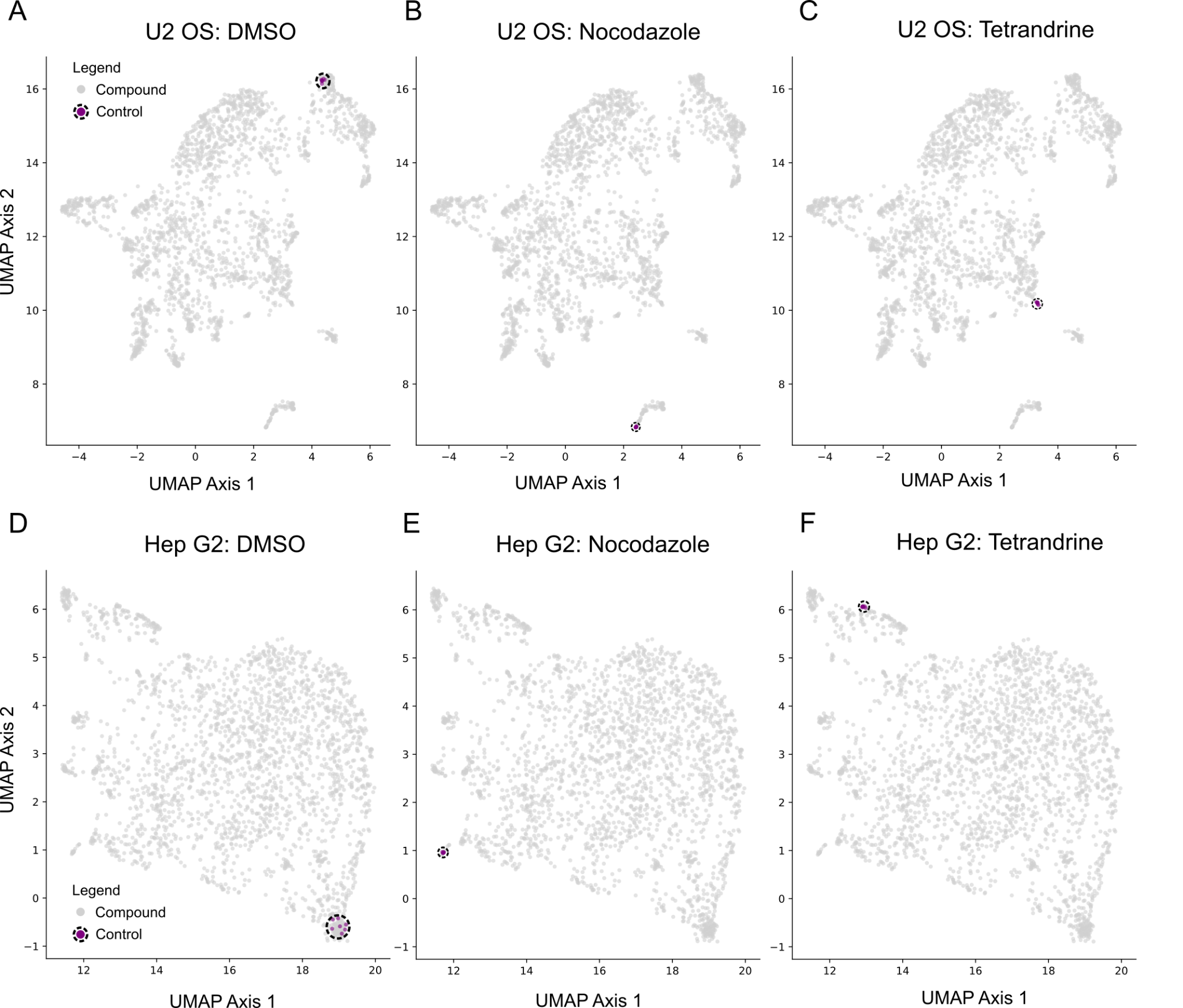

**Suppl. Figure 7: Visualization of morphological feature space in FMP datasets labeling control compounds.** (A-C) 2D UMAPs based on the FMP U-2 OS dataset labeling DMSO (A), Nocodazole (B) and Tetrandrine (C). (D-F) 2D UMAPs based on the FMP U-2 OS dataset labeling DMSO (D), Nocodazole (E) and Tetrandrine (F).
