## Supplementary material for "Morphological Profiling Dataset of EU-OPENSCREEN Bioactive Compounds Over Multiple Imaging Sites and Cell Lines": Suppl_Methods.docx

### Supplemental Methods

### Introduction

The goal of the cell-painting assay is to create a large dataset of morphological profiles in order to assess the biological impact of compounds on cells, and group them based on their mode of action. The complexity of the assay requires a specific and aligned work-flow involving different cell lines, thus the cell-painting assay should be conducted by more than one EU-OS partner site in two separate rounds. The first round involves the screening of the EU-OPENSCREEN bioactives library of about 2.500 compounds with one cell-line (HepG2) which will act as a proof-of-principle study. The aim is to show that this sophisticated assay can be performed at several labs with a consistent quality, delivering comparable data between the different sites. The ability to replicate the scientific findings with such a complex assay is the most important challenge. Additionally, since bioactives have a known mode of action, this specific proof-of-principle study should deliver representative and indicative data on the molecular and cellular effects of these compounds. The final number of the partners that will enter the first round will be defined after the submission and evaluation of all proposals, based on the evaluation criteria.
The successful sites will conduct the first screening round, and the results derived from this proof-of-principle study will be evaluated. The evaluation to qualify for this first screening round will be based on the replicate reproducibility of the reference compounds. For this purpose, the reference compounds will be added as controls to the bioactives library. The concentration with the highest correlation factor in the validation should be used for each reference compound. The comparison between the replicates of each reference compound should yield a correlation factor >0.95 to ensure also a high reproducibility of the morphological profiles when the bioactives library will be screened. The EU-OS partner sites which can then match the reproducibility criterium for the 2.500 compounds, will qualify for the second round of the cell-painting assay which can involve the test of additional cell-lines (number will depend on participating partner sites) and the screen of up to 100.000 compounds from the EU-OPENSCREEN compound collection. Based on the number of EU-OS partner sites which were successful in the first round, the cell-lines that will be tested during the second round can be divided according to the following scheme: two cell lines/partner site will be used while the second cell line tested by one partner site will also be tested as a first cell line by another partner site, in order to ensure reproducibility of screening data for the same number of compounds between at least two sites. For example: if 6 partners sites will qualify for the second round then 6 cell lines could be tested (A, B, C, D, E, and F) and divided between the 6 partners such that partner 1 will test A+B cell lines, partner 2 will test B+C cell lines, partner 3 will test C+D cell lines, partner 4 will test D+E cell lines, partner 5 will test E+F cell lines, and partner 6 will test F+A cell lines.

### Cell Painting assay

The cell painting assay should be based on the method proposed by (Bray et al. 2016) but with the necessary modifications to accommodate the available instrumentation at the different partner sites. A general experimental workflow with six fluorescent stains for imaging in four channels is shown in Figure 1. The protocol aims at analyzing the compound-induced changes in eight cellular compartments. As described above, the assay requires a validation with a small set of reference compounds which are shown in Table 1.

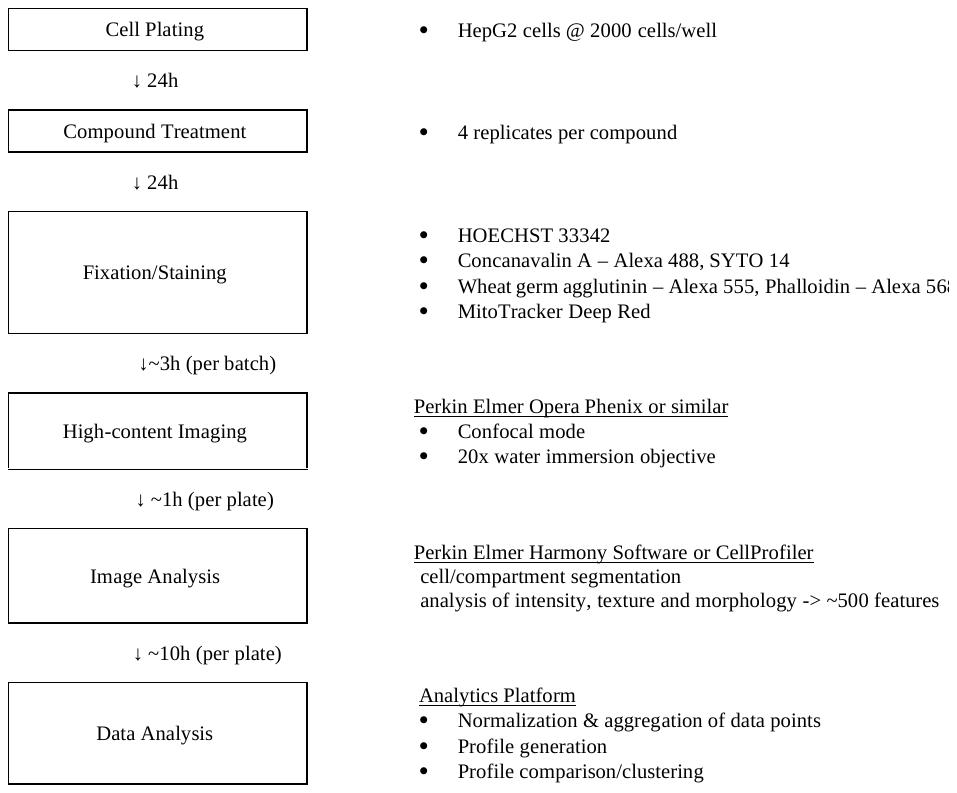

Figure 1: Suggested workflow for the cell-painting assay.
In summary, all compounds will be tested at a final concentration of 10µM with DMSO as vehicle control and reference compounds as evaluation control. Compounds will be transferred to the cells 24h after seeding. Staining procedures will be performed 24h after compound transfer as described in Bray et al. using the following final dye concentrations: 500 nM Mitotracker, 4 µM Hoechst, 25 µg/ml Concanavalin A, 3 µM SYTO14, 1U/ml Phalloidin and 1.5µg/ml Wheat germ agglutinin. For each compound, 4 replicates are required. Replicates have to be spread over different plates. Reference compounds, also to be used to generate the validation data for the assay proposal, induce the toxic effects via the cellular targets shown in Table 1.

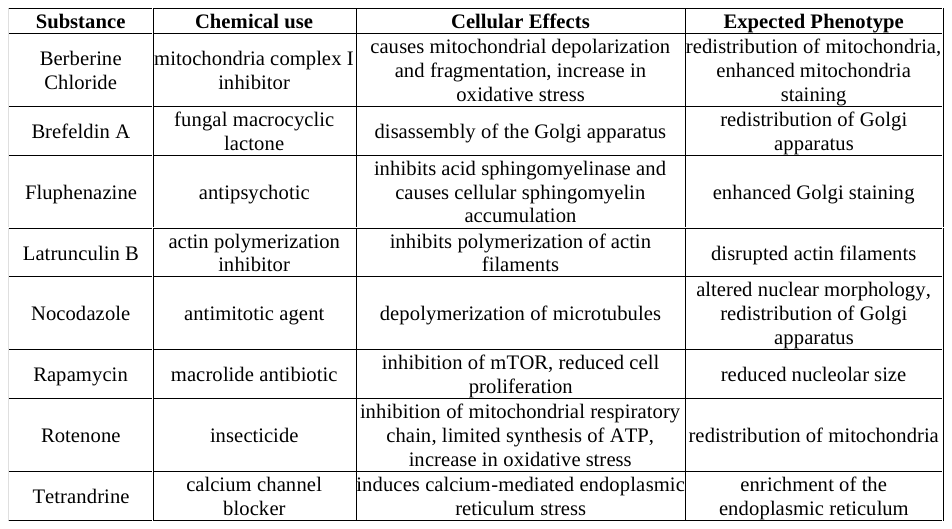

Table 1: Reference compounds with known toxic activities.
The data analysis can be performed either with the Harmony high-content imaging and analysis software from Perkin Elmer, or with the CellProfiler software which can also be used for single-cell feature detection. These single-cell feature values will be aggregated over the whole well, and all replicates with median, and then normalized to the DMSO control using z-scores. Morphological profiles for each compound can then be generated using these z-scores (Figure 2).
Compounds with high toxicity (changes in more than 80% of the morphological features) have to be tested again with concentrations of 3uM, 1uM and 0.3uM. In addition, compounds with a distinctive morphological profile should also be tested with similar concentration response curves to provide additional information on the concentration dependency as well as the possibility to better compare profiles of compounds with differing efficiencies. The number of compounds tested this way should not surpass 5% of the bioactives library. To facilitate the cost comparison of different proposals please assume a ‘hit’ rate of 1% and concentration response curves with 8 concentrations in duplicate, using 50uM as the highest concentration.
Similarity analysis and hierarchical clustering will allow identifying and grouping compounds of similar morphological effect. In addition, the resulting clustering based on the use of Harmony or CellProfiler (Stirling et al. 2021; Chandrasekaran et al. 2022) will be compared to estimate the comparability of cell painting data published from the different labs.

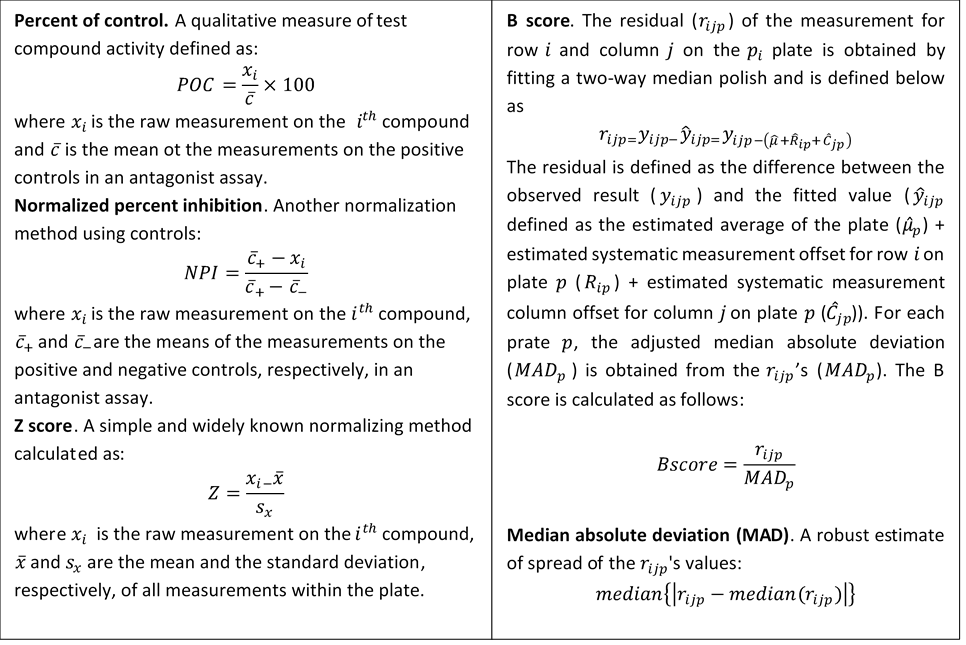
Figure 2: formulae for normalization

### Evaluation Criteria

#### The evaluation of the submitted proposals will be based on the following pre-defined criteria and the scoring system described in detail (Scoring system)

#### Adequate description and clarity of the assay (Weight: 20%)

The description of the assay should include the experimental protocol that will be followed including definition of compound concentration, type and number of cell lines where applicable, replicates, reagents, instruments etc.

#### Assay validation experiments/Assay performance and sensitivity (Weight: 30%)

The validation experiments will serve as small-scale demonstration of the suitability of the candidates to perform the large-scale assay(s) (testing the full library of compounds) of their interest if their proposal(s) is successful. Thus, the results of the validation experiments should be included in their submitted proposals. For the assay validation experiments the candidates will have to order all the necessary reagents in advance in order to perform the experiments before the submission of their proposal(s). Due to the complexity of the cell-painting assay, and in order to ensure compatibility of the candidates with the requirements of the assay, the reagents (HepG2 cells, reference compounds, and serum) for the cell-painting assay validation experiments will be provided by EU-OPENSCREEN ERIC and sent to the partner sites which indicated an interest in participation. The fluorescent stains need to be ordered by the participants under specific guidelines that will be provided by EU-OPENSCREEN regarding LOT numbers, vendor, etc.

#### Availability of assay validation data for Cell-painting assay

**Cell preparation and measurement.** Due to the complexity of the cell-painting assay, and in order to ensure compatibility of the candidates with the requirements of the assay, the reagents (HepG2 cells, reference compounds, serum) for the cell-painting assay validation experiments will be provided by EU-OPENSCREEN ERIC. Cell-painting assay validation experiments will be conducted with HepG2 cells and should be used within 10 passages. The cells should be cultured under the conditions described in the provided cell culture protocol to ensure growth without substantial clumping. Cells are grown until a confluency of 80% and either passaged or seeded for the experiment at a seeding density of 2.000 cells per 384-well. The experimental design of the cell painting assay follows the protocol (Bray et al. 2016) using the six fluorescent stains - MitoTracker Deep Red, Wheat germ agglutinin Alexa 555 conjugate, Phalloidin Alexa 568 conjugate, Concavalin A Alexa 488 conjugate, Hoechst 33342, SYTO 14 – which will be imaged in four channels to analyze the eight different cellular components or compartments. A small set of eight reference compounds (Berberine chloride, Brefeldin A, Fluphenazine, Latrunculin B, Nocodazole, Rapamycin, Rotenone, Tetrandrine) needs to be tested for obtaining the validation data. A 1:2 dilution series has to be prepared for testing of these compounds in cell assays at final concentrations starting from 50 µM using dilutions in 4 replicates (blue) plus DMSO controls (yellow) on one 384-well and identical replicate on a second plate. For this purpose, stock solutions are added to a pre-dilution plate and step wise diluted using DMSO as depicted in Figure 3. DMSO will also serve as vehicle control.

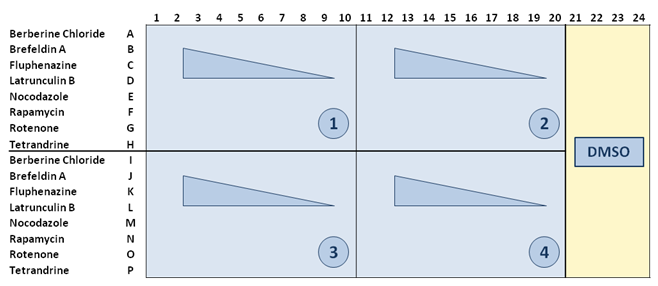

Figure 3: Schematic representation of the replicates and DMSO controls in 384-well plate.
24h after seeding the cells, compounds will be transferred to the cells resulting in a final concentration of 50 µM for the highest dilution. Use a pre-dilution of compounds in RPMI medium with 10% FBS if needed. Staining procedures will be performed 24h after compound transfer as described in Bray et al. using the following final dye concentrations: 500 nM Mitotracker, 4 µM Hoechst, 25 µg/ml Concanavalin A, 3 µM SYTO14, 1U/ml Phalloidin and 1.5µg/ml Wheat germ agglutinin. At least 2 plates with 4 replicates each should be performed to ensure data quality. All plates can be prepared with the same batch of cells.
**Image analysis**. To extract cell morphology feature data, one of two analysis softwares could be used: either CellProfiler (Bray et al. 2016; Stirling et al. 2021), or Harmony (PerkinElmer) following a pipeline provided by EU-OPENSCREEN. For the CellProfiler pipeline, output of the measured features should be adjusted to a comma-delimited file (csv) using the *ExportToSpreadsheet* module. Extracted features in Harmony should be saved as txt file. The result files (txt or csv) will have to be included in your proposal and should be provided in electronic format.
The results will be evaluated by an external reviewers’ committee using similarity checks between the resulting morphological profiles and similar profiles generated by EU-OPENSCREEN using the appropriate software (KNIME, R or others) to check how similar the profiles are to each other. Each extracted feature value will first be normalized by Z’-score normalization (using the feature values of the DMSO-wells as background population) (Figure 2).
The correlation of those z-scores between the two technical replicates (averages of each plate) will be calculated using the Pearson correlation equation (Figure. 4). For each compound, at least one concentration of each reference compound should yield a correlation factor > 0.95.

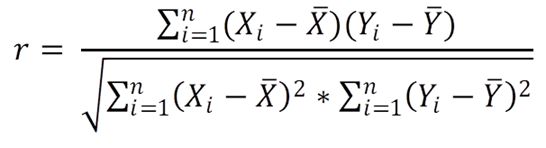

Figure 4: The Pearson correlation equation.
Where:
**X**: vector of Z’-scores of the features, replicate 1
**Y**: vector of Z’-scores of the features, replicate 2
**r**: Pearson correlation coefficient

#### Assay cost and timelines (Weight: 40%)

A justified and reasonable total cost breakdown analysis for a full assay, including reagents, replicates, personnel and time for testing 10.000 and 100.000 compounds should be provided. This should be accompanied with a detailed plan with timelines for screening 100.000 compounds. If indicated in the assay descriptions (e.g., for assays 3 and 4), estimations for lower number of compounds can be given.

#### Proven track record of the services provider, excellence of the provider (Weight: 5%)

Please include a statement on experience with similar assays in the past, reference to publications, established protocols etc. Please also state the required scientific and technical expertise for performing and analyzing the data related to the assay(s) for which you have applied.

#### Consumption of tested compounds (Weight: 5%)

Please indicate the highest compound concentration in the assay and the approximate need of stock solution (x µl of 10mM in DMSO) for each compound to perform the assay, as well as data about reagent stability and compatibility with DMSO.

### Evaluation scoring

#### Evaluation stage 1

As a first step, the participant should reach a score ≥65% for Criterion Nr.1 and validation data for the Cell Painting assay for each of the submitted proposals. The calculation of the score for this step is described in Table 2. In that case, the evaluation of the proposal continues to the second step. If the score for Criterion Nr.1: validation data is ˂65% the submitted proposal is rejected and excluded from the bioprofiling tender process. The scoring system for the cell-painting assay validation data is described in Table 2 & 3.

Table 2: Scoring system for Criterion 1.

| **Validation data for assays** | | | |  |  |
| --- | --- | --- | --- | --- | --- |
| **Criterion Nr.** | **Description** | **Answer** | **Weight factor** |  |  |
| **1** | Validation data for assay. |  | **100%** | **Points** | **Comments** |
| 1.1 | 8 positive and minimum of 64 DMSO background controls. | **☐Yes ☐No** | 10**%** |  |  |
| 1.2 | At least two independent experiments on different plates. | One experiment: 0 points  Two experiments or more: 100 points | 5% |  |  |
| 1.3 | Controls should reflect the absence and presence of the anticipated signal caused by interfering or modulating compounds | **☐Yes ☐No** | 5**%** |  |  |
| 1.4 | Plate patterns should give no indication of signal drift or edge effects. | **☐Yes ☐No** | 5**%** |  |  |
| 1.5 | Quantitative assessment of the assay demonstrated by Z’ factor calculation. | Not applicable | **-** |  |  |
| 1.6 | The correlation of the z-scores between the two technical replicates (averages of each plate) will be calculated using the Pearson correlation equation. | For each compound, one or more concentrations of 8 reference compounds with correlation factor > 0.95: 100 points  For 7 out of 8 compounds, one or more concentrations with a correlation factor > 0.95: 75 points  Correlation factors do not reach above numbers, but are not significantly lower as those from other groups: 50 points  Correlation factors do not meet above numbers and are significantly lower as those from other groups: exclusion from tender. | 70% |  |  |
| 1.7 | Please provide documentation on the stability of the critical (crucial to assay performance) reagents of the assay. For assistance you may refer to [The Principles of Good Laboratory Practice](https://www.researchgate.net/profile/Miroslav_Cervinka/publication/281570234_The_Principles_of_Good_Laboratory_Practice_Application_to_In_Vitro_Toxicology_Studies_The_Report_and_Recommendations_of_ECVAM_Workshop_3712/links/59d25f344585150177f63475/The-Principles-of-Good-Laboratory-Practice-Application-to-In-Vitro-Toxicology-Studies-The-Report-and-Recommendations-of-ECVAM-Workshop-371-2.pdf) | **☐Yes ☐No** | 5**%** |  |  |
| 1.8 | Tolerance to DMSO:  typical % final DMSO concentration in biochemical and cell-based assays. | Not applicable | **-** |  |  |
|  |  |  |  | **Total points** | **Overall review** |

#### Evaluation stage 2

After the initial step, the evaluation of the submitted proposals will be based on five additional criteria: assay cost, assay time needed for screening 100.000 compounds, description/clarity of the assay, required compound consumption and proven track record of the services/excellence of the provider. However, the score of the assay validation data from the previous step will also be taken into account (with a weight factor of 10%) in the participant’s final score. The scoring system for the above-mentioned criteria is presented in Table 2.

Table 3: Scoring system for the calculation of the final score of the applicant.

| **Criterion Nr.** | **Description** | **Answer** | **Weight factor** | **Points** | **Comments** |
| --- | --- | --- | --- | --- | --- |
| **1** | **Assay validation data** | **Please refer to Step 1** | **10%** |  |  |
| **2** | **Assay cost**  A justified and reasonable total cost breakdown analysis for a full assay, including reagents, replicates, personnel and time for testing 100.000 compounds, should be provided. | For each of the assays: score = 40% *(LP/OP)  LP: lowest price of all quotes  OP: offered price of this quote | **40%** |  |  |
| **3** | **Time needed for 100.000 compounds**  Justified and reasonable detailed plan with timelines for screening 100.000 compounds | For each of the assays: Maximum data delivery time 12 months (delivery time˃12 months: 0 points)  score = 20% *(LT/OT)  LT: lowest data delivery time of all quotes  OT: offered data delivery time of quote | **20%** |  |  |
| **4** | **Description/clarity of the assay** |  | **20%** |  |  |
| 4.1 | -Description of positive and negative assay controls (compound, dissolver, concentration, etc.)  -Compound and reagents concentrations  -Replicates  -Plating format |  | 5% |  |  |
| 4.2 | **Protocol details:**  -Automation flowchart, which is a graphical layout of the different operational steps, such as dispensing, incubation, compound-transfer and plate reading  -Automation instruments (manufacturer, model, instrument parameters) |  | 5% |  |  |
| 4.3 | -Reagent details such as vendor, catalogue and lot number (if applicable), cost, storage conditions, sensitivity to light/temperature (if known), shelf life, preparation  -Cell line details. Please provide SOP or key points of experimental protocol. (e.g., source, catalogue number, phenotype, passage number, split ratio, media, culture protocol, etc.) |  | 5% |  |  |
| 4.4 | **Analysis details. Please refer to specific values for:**  -Readout (including time of readout)  -Normalisation method (e.g., percent of control, normalized percent inhibition, Z score, B score, median absolute deviation). Please refer to Fig.4.  -Test statistics for hit detection with replicates |  | 5% |  |  |
| **5** | **Consumption of tested compounds.**  Please indicate the highest compound concentration in the assay and the approximate need of stock solution (x ul of 10mM in DMSO) to perform the assay. | For each of the assays:  score = 5% *(LV/OV)  LV: lowest required volume of all quotes  OV: offered required volume of this quote | **5%** |  |  |
| **6** | **Proven track record of the services provider.** Please include a statement on:  -experience with similar assays in the past  -reference to publications  -established protocols (clear SOPs).  -scientific and technical expertise for performing and analyzing the data related to the assay(s) for which you have applied. | For the publications:  0 publications relevant to the assay: 0 points  ≥5 publications: 100 points, linear increase for between 0 and 5. | **5%** |  |  |
|  |  |  |  | **Total points** | **Overall review** |

Chandrasekaran, Srinivas Niranj, Beth A. Cimini, Amy Goodale, Lisa Miller, Maria Kost-Alimova, Nasim Jamali, John G. Doench, Briana Fritchman, Adam Skepner, Michelle Melanson, Alexandr A. Kalinin, John Arevalo, Marzieh Haghighi, Juan Caicedo, Daniel Kuhn, Desiree Hernandez, Jim Berstler, Hamdah Shafqat-Abbasi, David Root, Susanne E. Swalley, Sakshi Garg, Shantanu Singh, and Anne E. Carpenter. 2022. "Three million images and morphological profiles of cells treated with matched chemical and genetic perturbations." In.: Bioinformatics.

Stirling, D. R., M. J. Swain-Bowden, A. M. Lucas, A. E. Carpenter, B. A. Cimini, and A. Goodman. 2021. 'CellProfiler 4: improvements in speed, utility and usability', *BMC Bioinformatics*, 22: 433.
