## Supplementary material for "Morphological Profiling Dataset of EU-OPENSCREEN Bioactive Compounds Over Multiple Imaging Sites and Cell Lines": Suppl_QC_FMP_HepG2.pdf

### HepG2 FMP

1

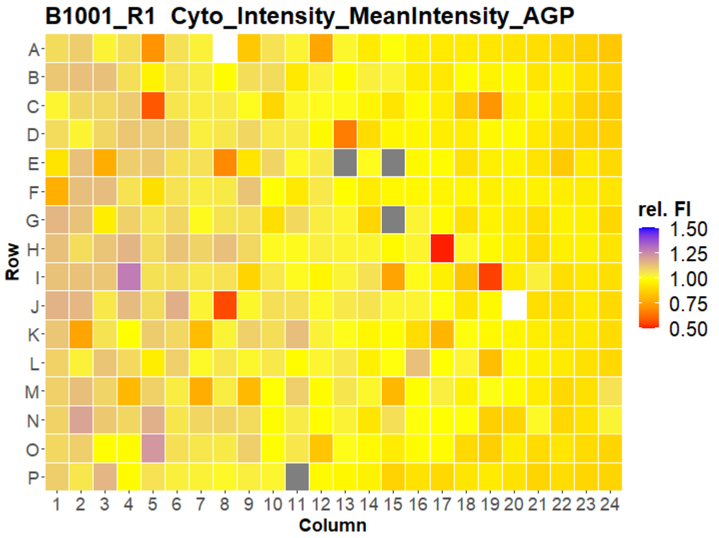


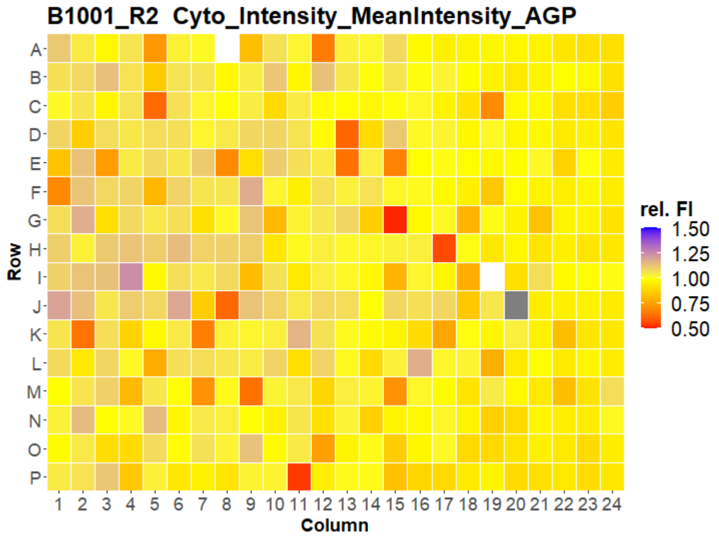


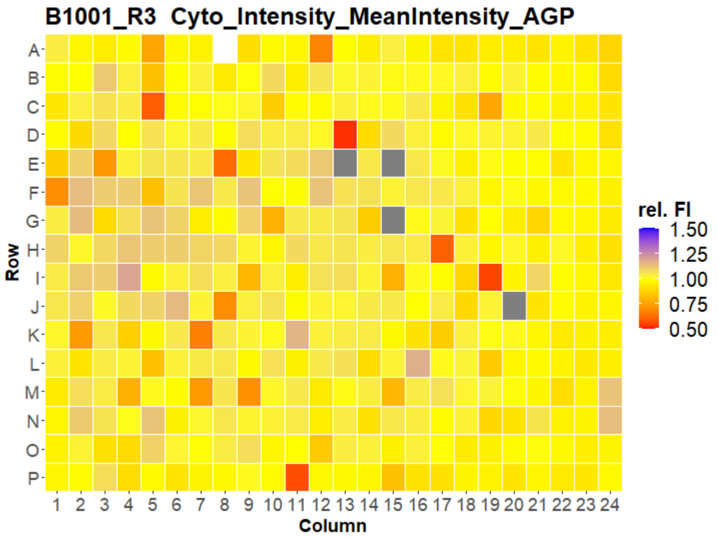


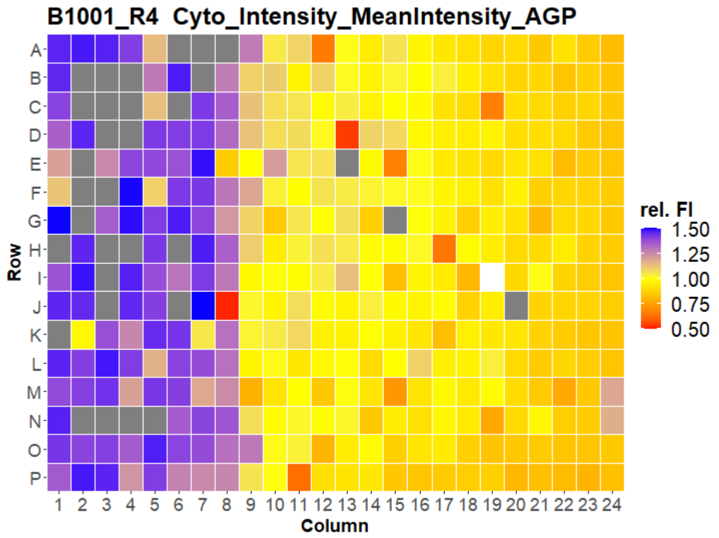


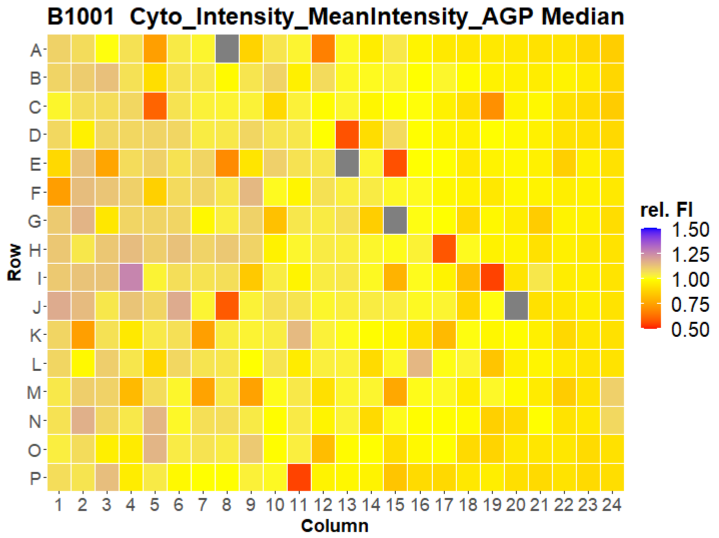


Created with KNIME Analytics Platform www.knime.com

Open for Innovation KNIME

### HepG2 FMP

2

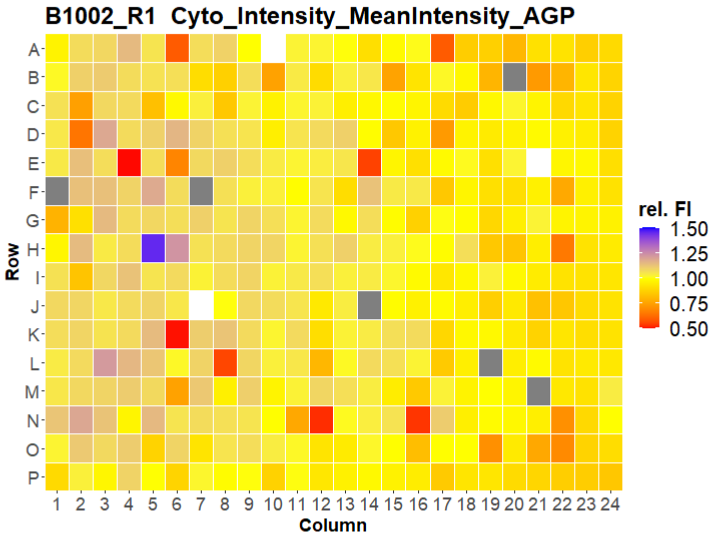
B1002_R1 Cyto_Intensity_MeanIntensity_AGP
A B C D E F G H I J K L M N O P
Row
1 2 3 4 5 6 7 8 9 10 11 12 13 14 15 16 17 18 19 20 21 22 23 24
Column
rel. FI
1.50
1.25
1.00
0.75
0.50

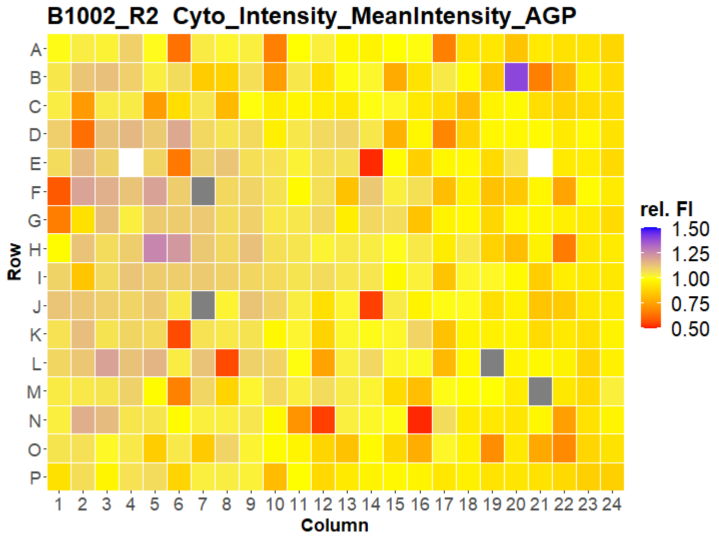
B1002_R2 Cyto_Intensity_MeanIntensity_AGP
A B C D E F G H I J K L M N O P
Row
1 2 3 4 5 6 7 8 9 10 11 12 13 14 15 16 17 18 19 20 21 22 23 24
Column
rel. FI
1.50
1.25
1.00
0.75
0.50

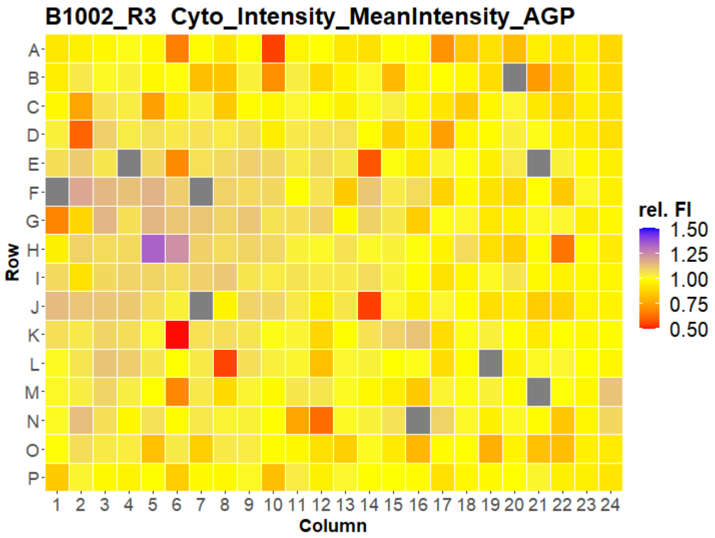
B1002_R3 Cyto_Intensity_MeanIntensity_AGP
A B C D E F G H I J K L M N O P
Row
1 2 3 4 5 6 7 8 9 10 11 12 13 14 15 16 17 18 19 20 21 22 23 24
Column
rel. FI
1.50
1.25
1.00
0.75
0.50

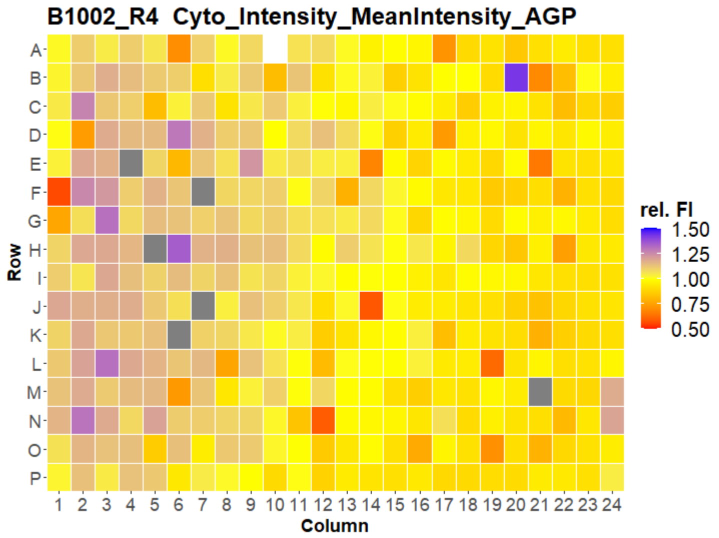
B1002_R4 Cyto_Intensity_MeanIntensity_AGP
A B C D E F G H I J K L M N O P
Row
1 2 3 4 5 6 7 8 9 10 11 12 13 14 15 16 17 18 19 20 21 22 23 24
Column
rel. FI
1.50
1.25
1.00
0.75
0.50

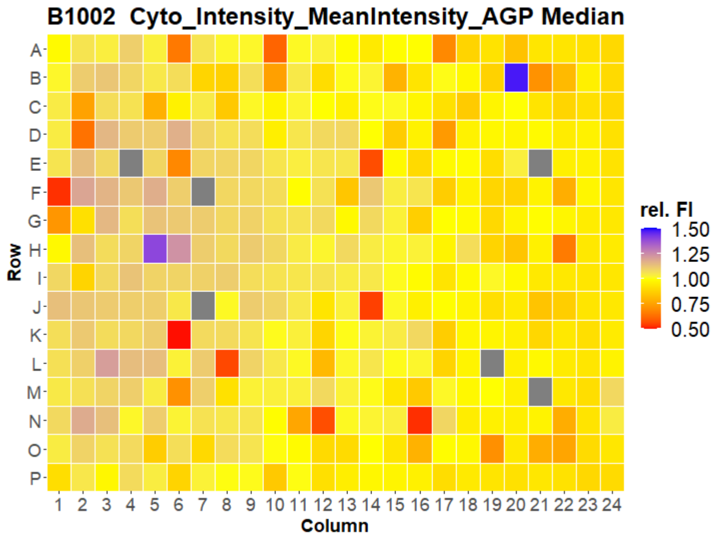
B1002 Cyto_Intensity_MeanIntensity_AGP Median
A B C D E F G H I J K L M N O P
Row
1 2 3 4 5 6 7 8 9 10 11 12 13 14 15 16 17 18 19 20 21 22 23 24
Column
rel. FI
1.50
1.25
1.00
0.75
0.50

### HepG2 FMP

3

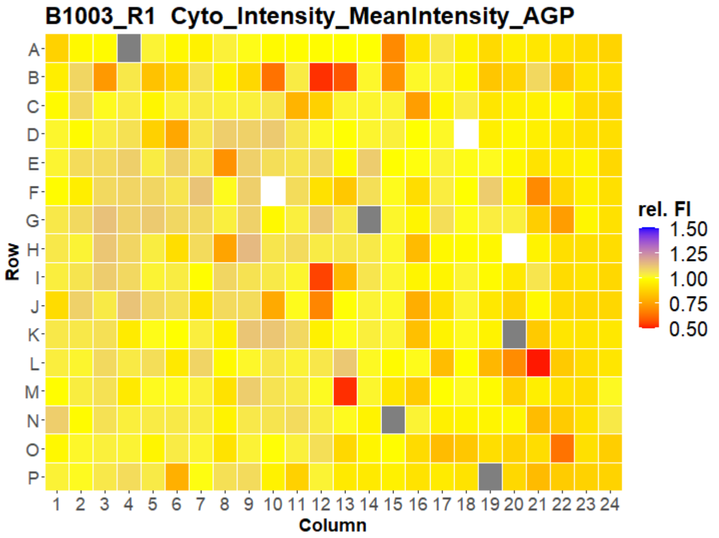
B1003_R1 Cyto_Intensity_MeanIntensity_AGP
A B C D E F G H I J K L M N O P
Row
1 2 3 4 5 6 7 8 9 10 11 12 13 14 15 16 17 18 19 20 21 22 23 24
Column
rel. FI
1.50
1.25
1.00
0.75
0.50

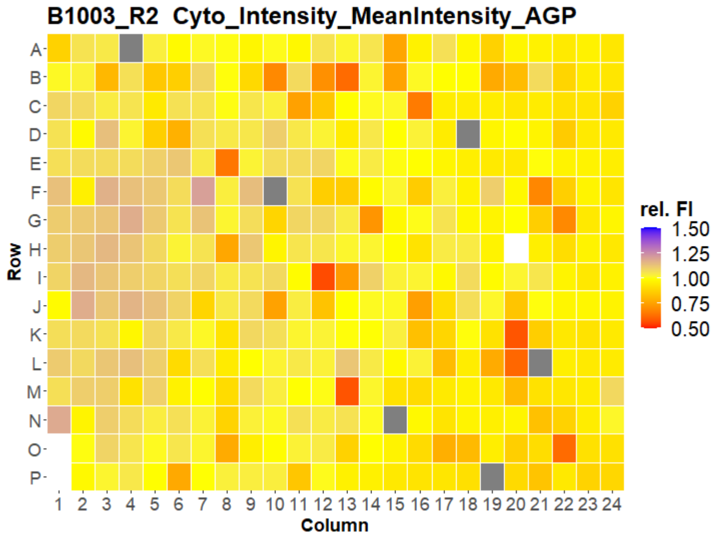
B1003_R2 Cyto_Intensity_MeanIntensity_AGP
A B C D E F G H I J K L M N O P
Row
1 2 3 4 5 6 7 8 9 10 11 12 13 14 15 16 17 18 19 20 21 22 23 24
Column
rel. FI
1.50
1.25
1.00
0.75
0.50

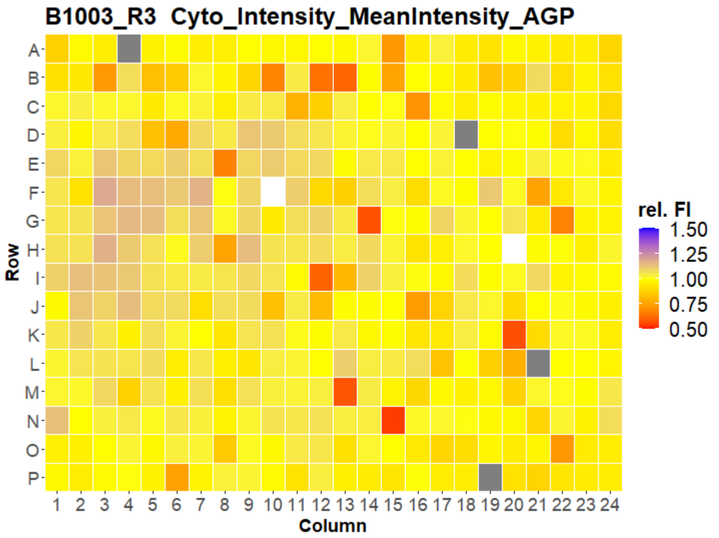
B1003_R3 Cyto_Intensity_MeanIntensity_AGP
A B C D E F G H I J K L M N O P
Row
1 2 3 4 5 6 7 8 9 10 11 12 13 14 15 16 17 18 19 20 21 22 23 24
Column
rel. FI
1.50
1.25
1.00
0.75
0.50

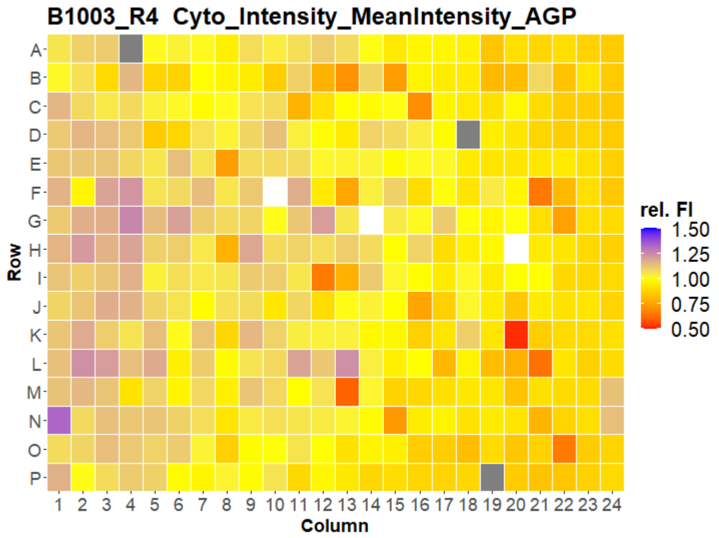
B1003_R4 Cyto_Intensity_MeanIntensity_AGP
A B C D E F G H I J K L M N O P
Row
1 2 3 4 5 6 7 8 9 10 11 12 13 14 15 16 17 18 19 20 21 22 23 24
Column
rel. FI
1.50
1.25
1.00
0.75
0.50

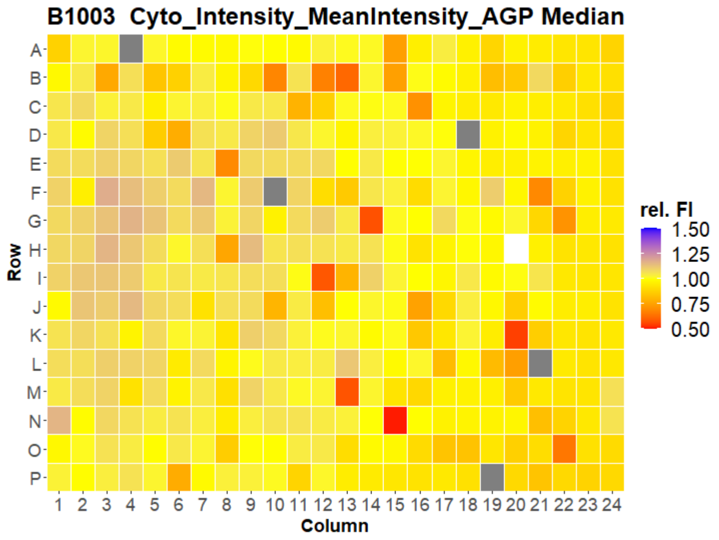
B1003 Cyto_Intensity_MeanIntensity_AGP Median
A B C D E F G H I J K L M N O P
Row
1 2 3 4 5 6 7 8 9 10 11 12 13 14 15 16 17 18 19 20 21 22 23 24
Column
rel. FI
1.50
1.25
1.00
0.75
0.50

### HepG2 FMP



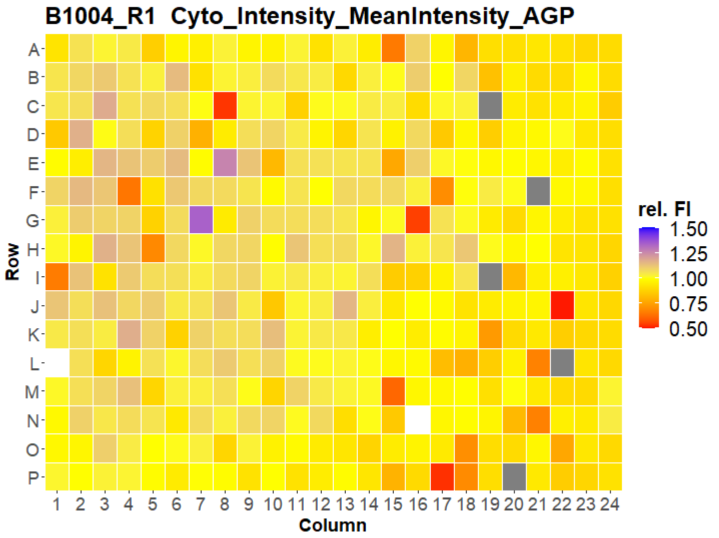
B1004_R1 Cyto_Intensity_MeanIntensity_AGP
Row
Column
rel. FI
1.50
1.25
1.00
0.75
0.50

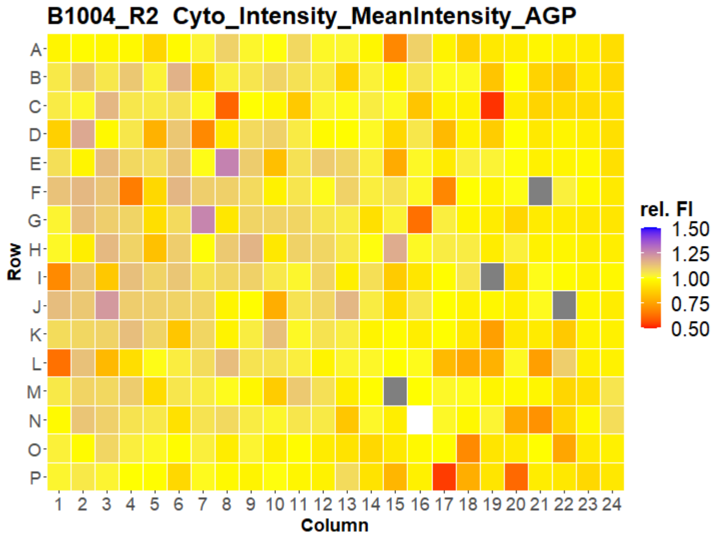
B1004_R2 Cyto_Intensity_MeanIntensity_AGP
Row
Column
rel. FI
1.50
1.25
1.00
0.75
0.50

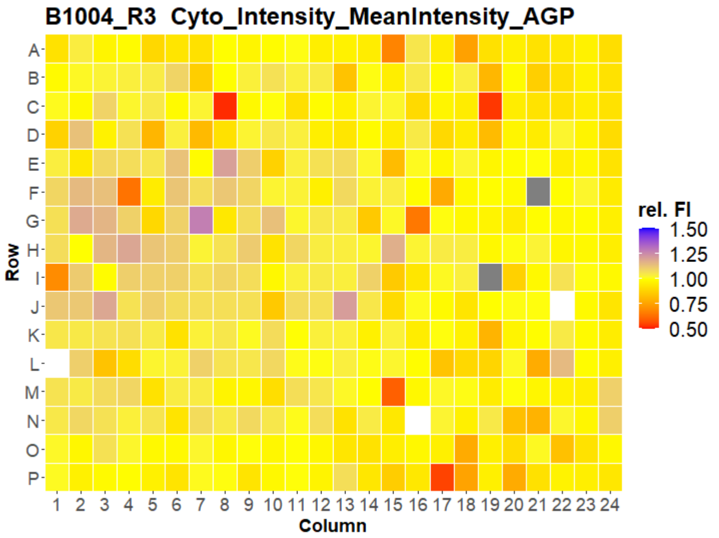
B1004_R3 Cyto_Intensity_MeanIntensity_AGP
Row
Column
rel. FI
1.50
1.25
1.00
0.75
0.50

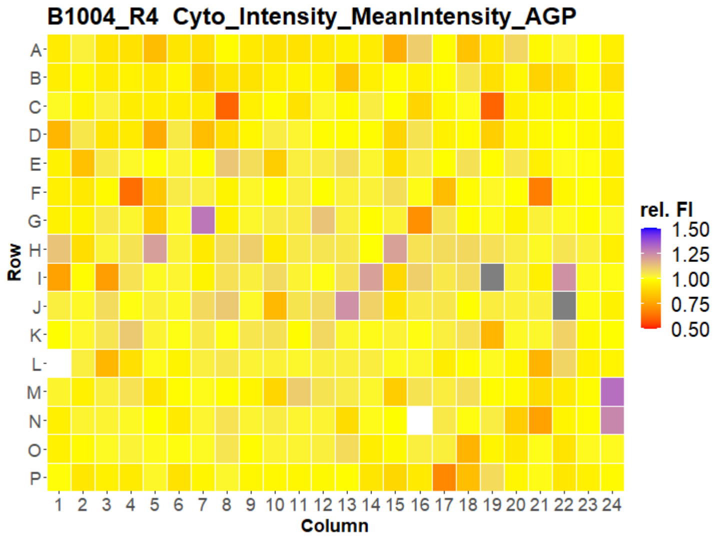
B1004_R4 Cyto_Intensity_MeanIntensity_AGP
Row
Column
rel. FI
1.50
1.25
1.00
0.75
0.50

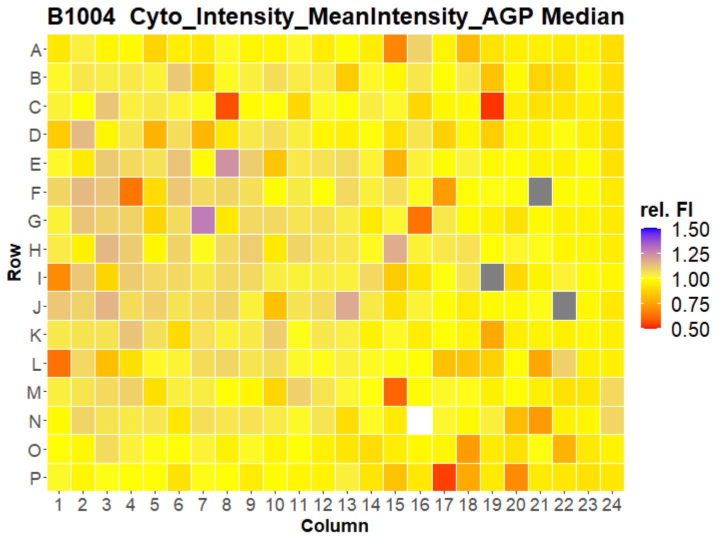
B1004 Cyto_Intensity_MeanIntensity_AGP Median
Row
Column
rel. FI
1.50
1.25
1.00
0.75
0.50

### HepG2 FMP



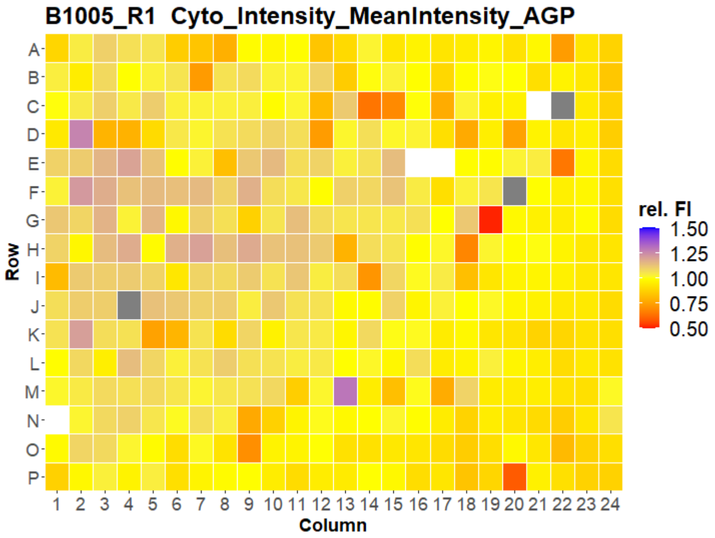
B1005_R1 Cyto_Intensity_MeanIntensity_AGP
Row
A B C D E F G H I J K L M N O P
Column
1 2 3 4 5 6 7 8 9 10 11 12 13 14 15 16 17 18 19 20 21 22 23 24
rel. FI
1.50
1.25
1.00
0.75
0.50

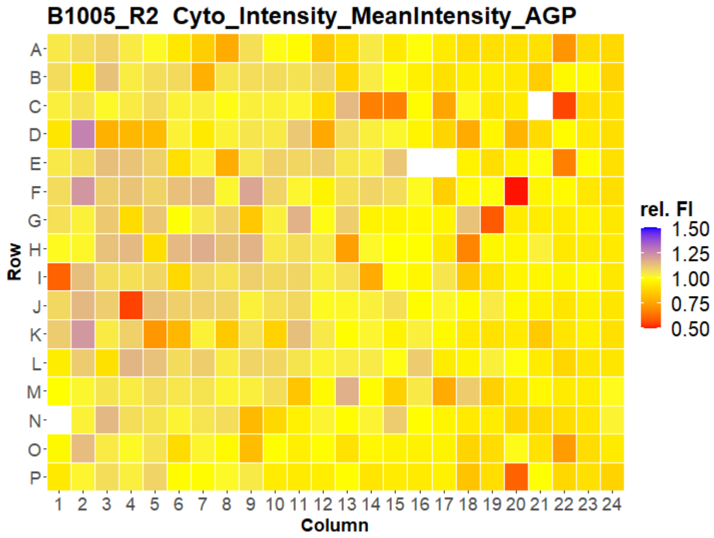
B1005_R2 Cyto_Intensity_MeanIntensity_AGP
Row
A B C D E F G H I J K L M N O P
Column
1 2 3 4 5 6 7 8 9 10 11 12 13 14 15 16 17 18 19 20 21 22 23 24
rel. FI
1.50
1.25
1.00
0.75
0.50

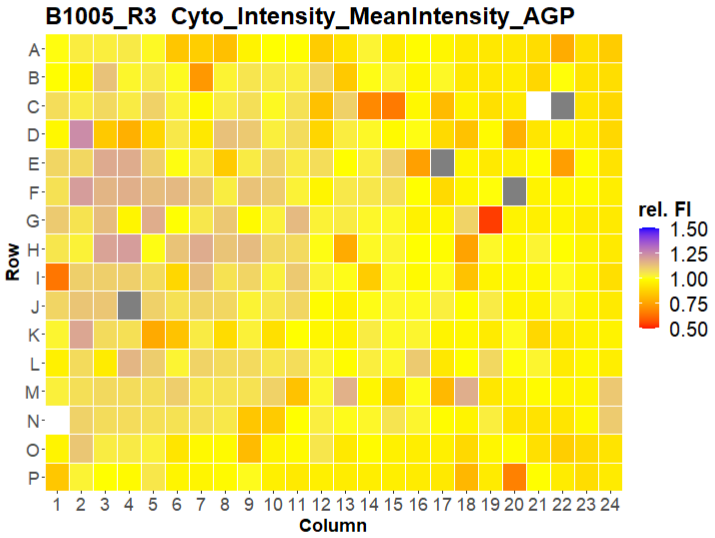
B1005_R3 Cyto_Intensity_MeanIntensity_AGP
Row
A B C D E F G H I J K L M N O P
Column
1 2 3 4 5 6 7 8 9 10 11 12 13 14 15 16 17 18 19 20 21 22 23 24
rel. FI
1.50
1.25
1.00
0.75
0.50

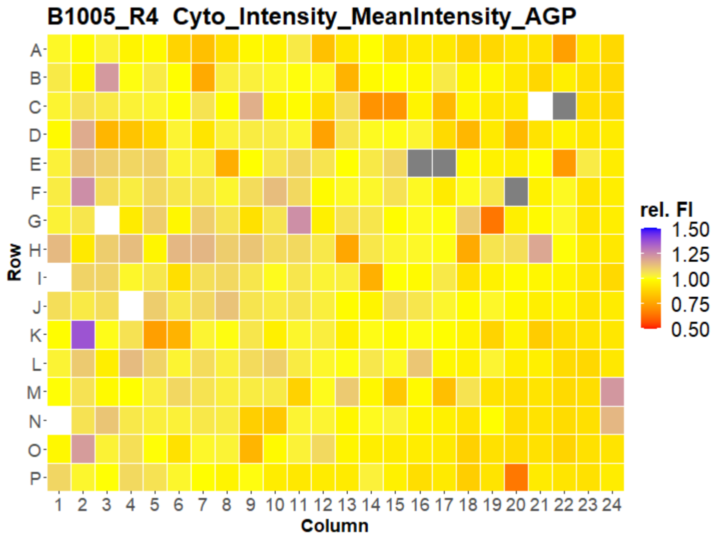
B1005_R4 Cyto_Intensity_MeanIntensity_AGP
Row
A B C D E F G H I J K L M N O P
Column
1 2 3 4 5 6 7 8 9 10 11 12 13 14 15 16 17 18 19 20 21 22 23 24
rel. FI
1.50
1.25
1.00
0.75
0.50

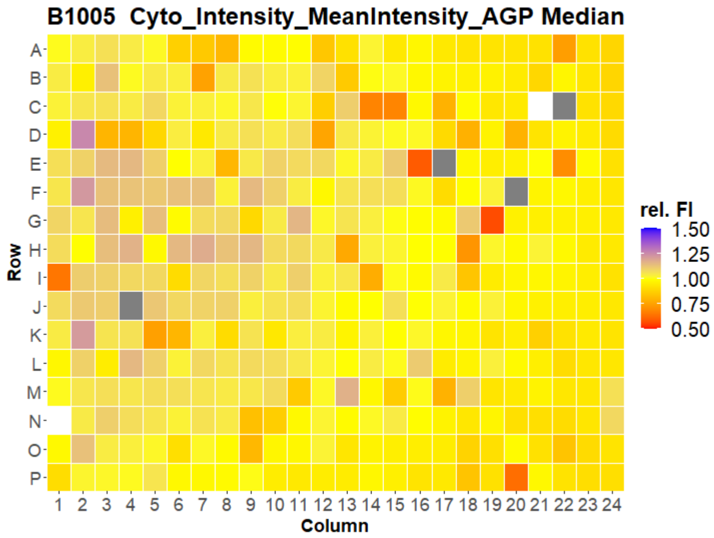
B1005 Cyto_Intensity_MeanIntensity_AGP Median
Row
A B C D E F G H I J K L M N O P
Column
1 2 3 4 5 6 7 8 9 10 11 12 13 14 15 16 17 18 19 20 21 22 23 24
rel. FI
1.50
1.25
1.00
0.75
0.50

### HepG2 FMP

6

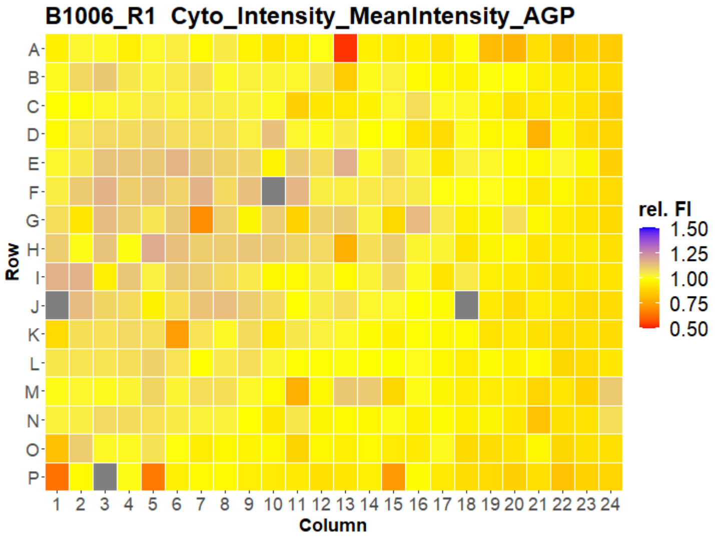
B1006_R1 Cyto_Intensity_MeanIntensity_AGP
Row
Column
rel. FI
1.50
1.25
1.00
0.75
0.50

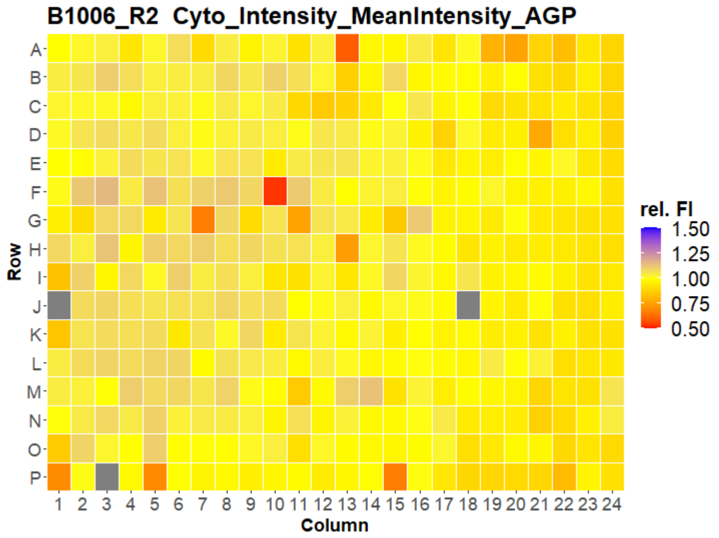
B1006_R2 Cyto_Intensity_MeanIntensity_AGP
Row
Column
rel. FI
1.50
1.25
1.00
0.75
0.50

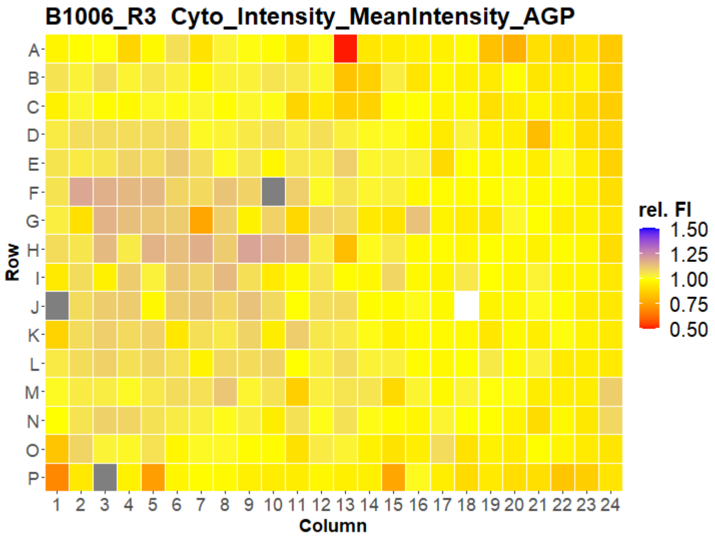
B1006_R3 Cyto_Intensity_MeanIntensity_AGP
Row
Column
rel. FI
1.50
1.25
1.00
0.75
0.50

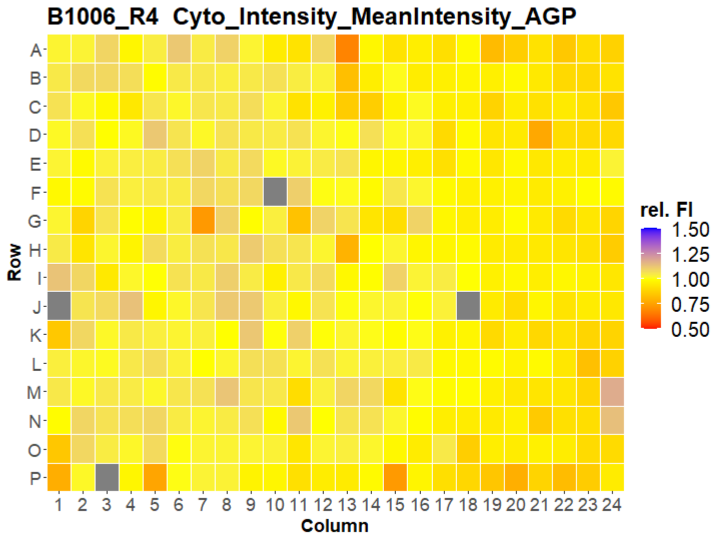
B1006_R4 Cyto_Intensity_MeanIntensity_AGP
Row
Column
rel. FI
1.50
1.25
1.00
0.75
0.50

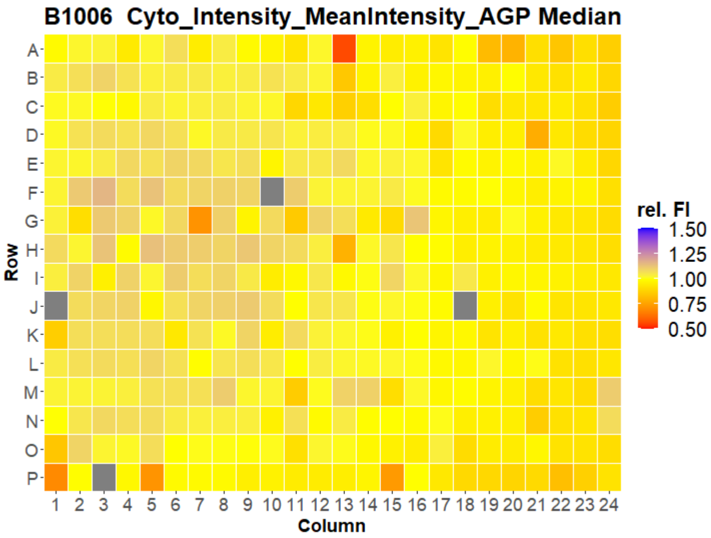
B1006 Cyto_Intensity_MeanIntensity_AGP Median
Row
Column
rel. FI
1.50
1.25
1.00
0.75
0.50

### HepG2 FMP

7

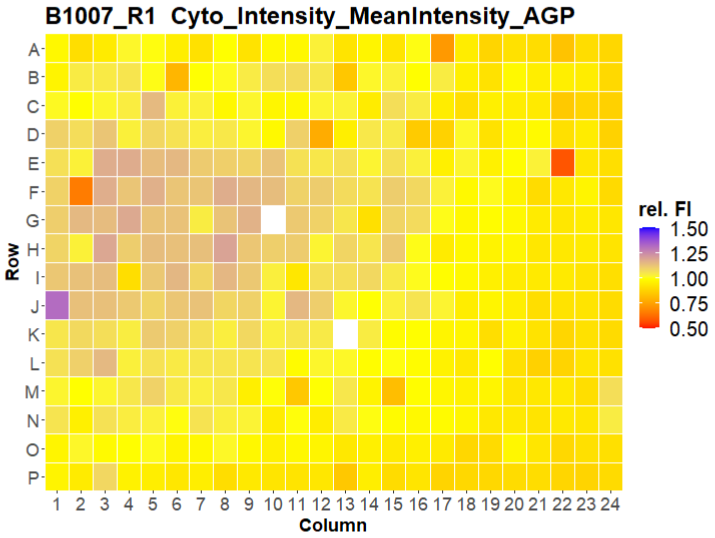
B1007_R1 Cyto_Intensity_MeanIntensity_AGP
Row
Column
rel. FI
1.50
1.25
1.00
0.75
0.50

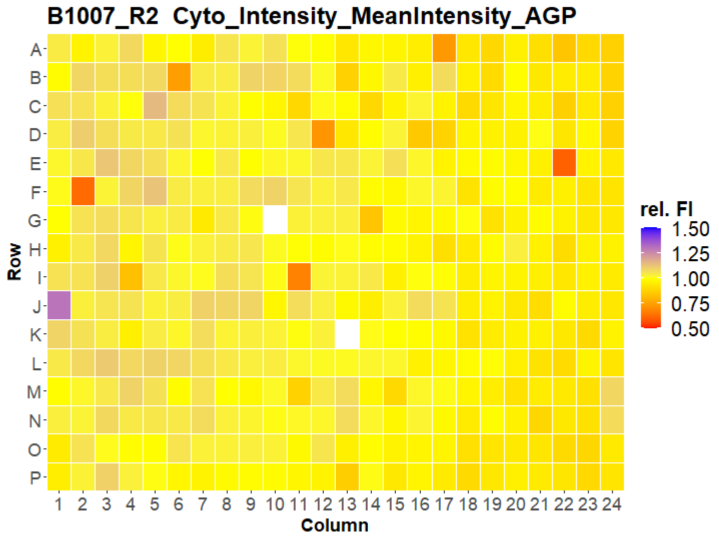
B1007_R2 Cyto_Intensity_MeanIntensity_AGP
Row
Column
rel. FI
1.50
1.25
1.00
0.75
0.50

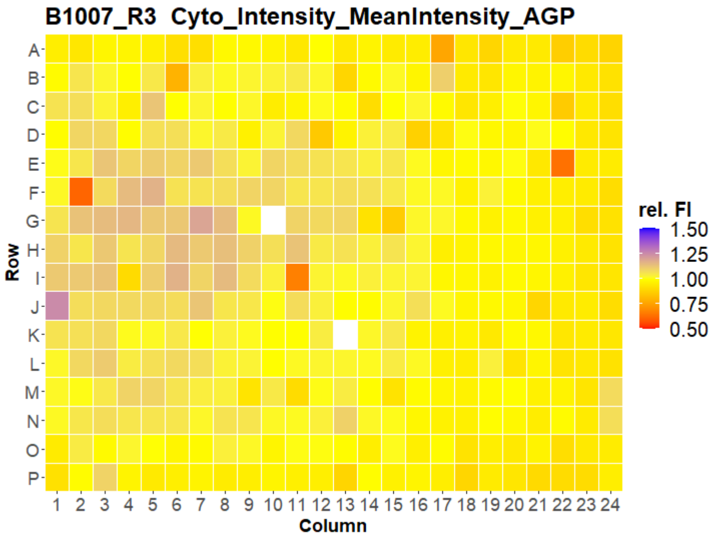
B1007_R3 Cyto_Intensity_MeanIntensity_AGP
Row
Column
rel. FI
1.50
1.25
1.00
0.75
0.50

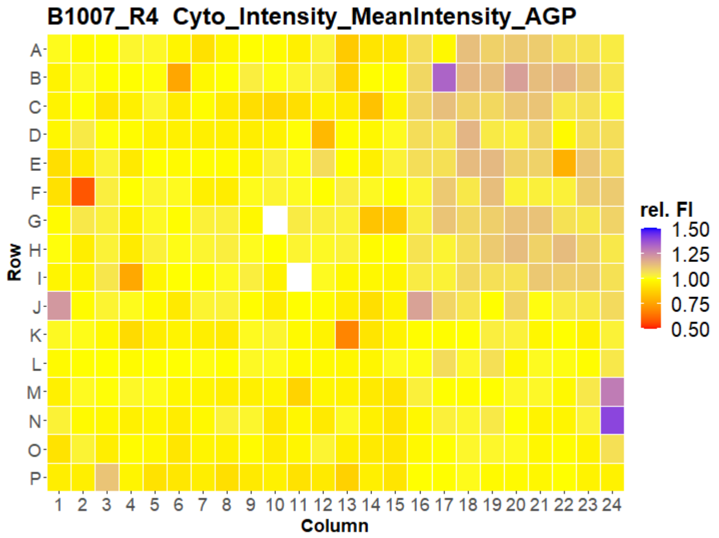
B1007_R4 Cyto_Intensity_MeanIntensity_AGP
Row
Column
rel. FI
1.50
1.25
1.00
0.75
0.50

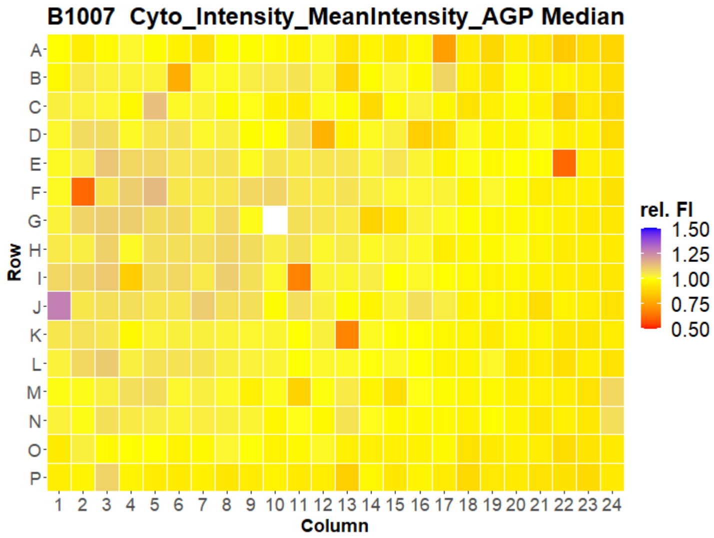
B1007 Cyto_Intensity_MeanIntensity_AGP Median
Row
Column
rel. FI
1.50
1.25
1.00
0.75
0.50

### HepG2 FMP

8

B1001_R1 Cyto_Intensity_MeanIntensity_ER

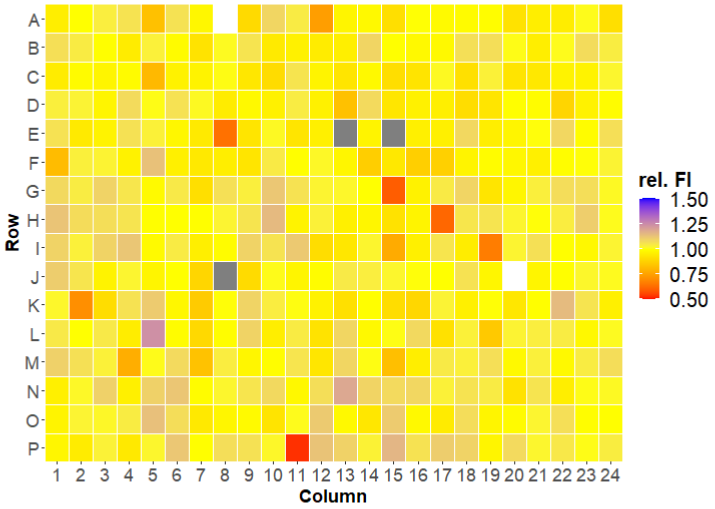


B1001_R2 Cyto_Intensity_MeanIntensity_ER

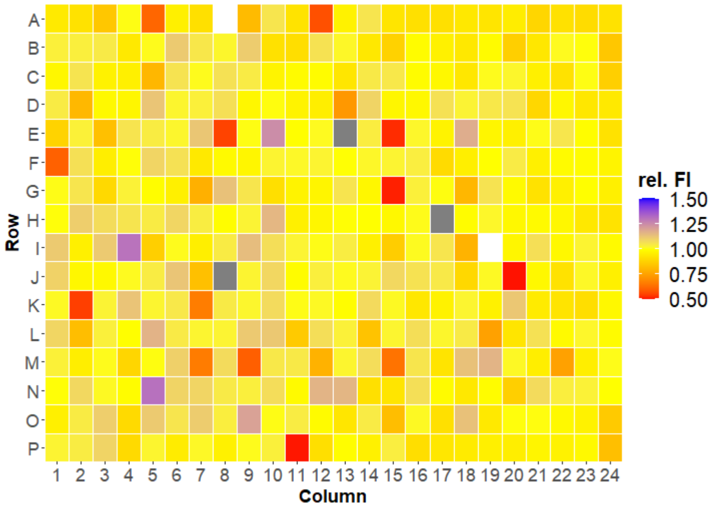


B1001_R3 Cyto_Intensity_MeanIntensity_ER

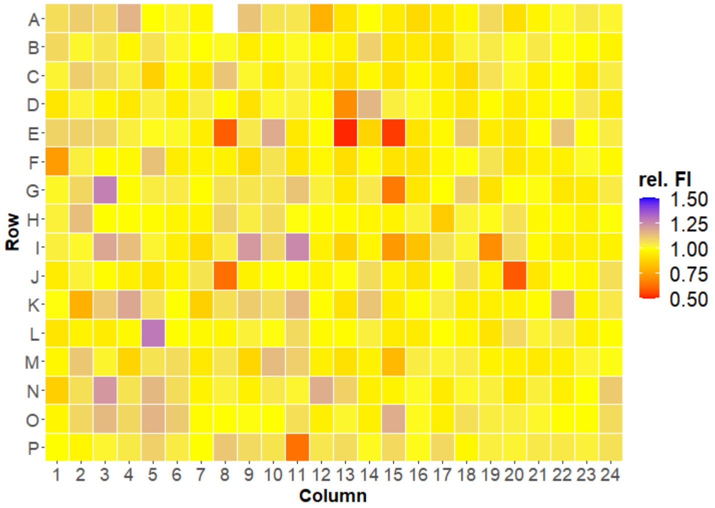


B1001_R4 Cyto_Intensity_MeanIntensity_ER

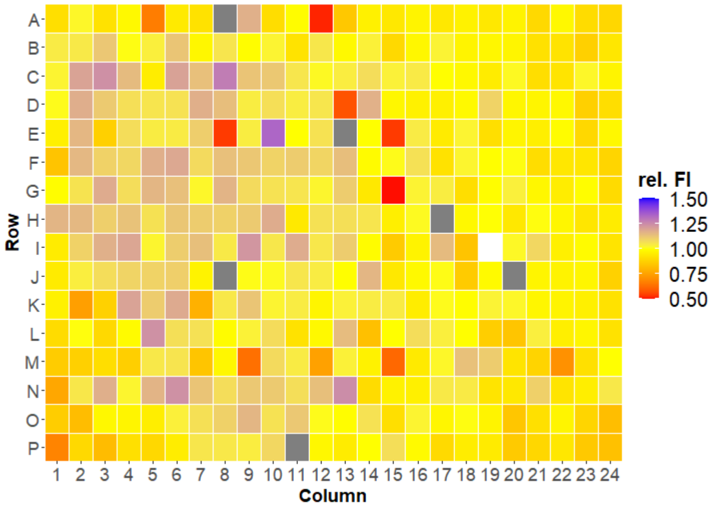


B1001 Cyto_Intensity_MeanIntensity_ER Median

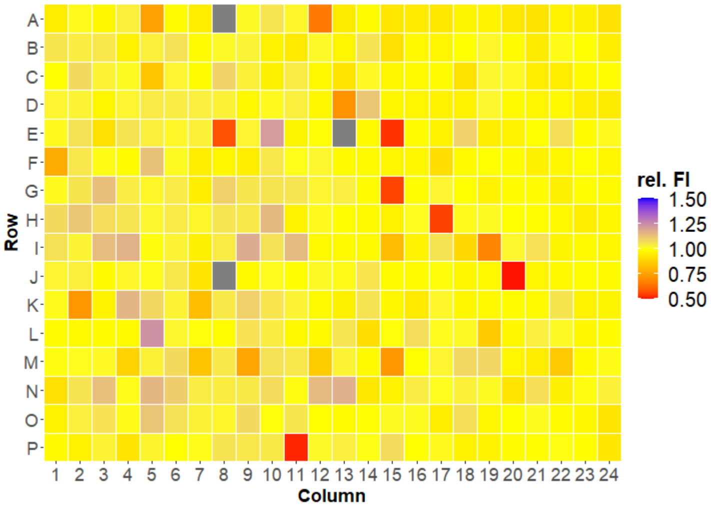



Open for Innovation KNIME

### HepG2 FMP



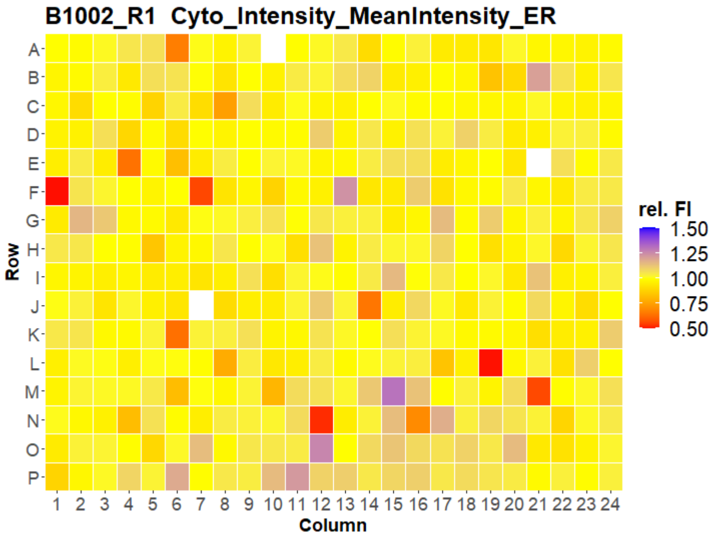
B1002_R1 Cyto_Intensity_MeanIntensity_ER
Row
Column
rel. FI
1.50
1.25
1.00
0.75
0.50

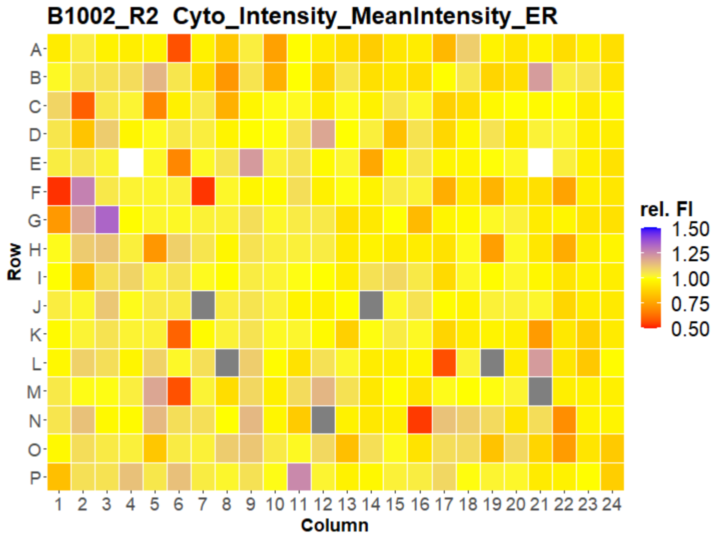
B1002_R2 Cyto_Intensity_MeanIntensity_ER
Row
Column
rel. FI
1.50
1.25
1.00
0.75
0.50

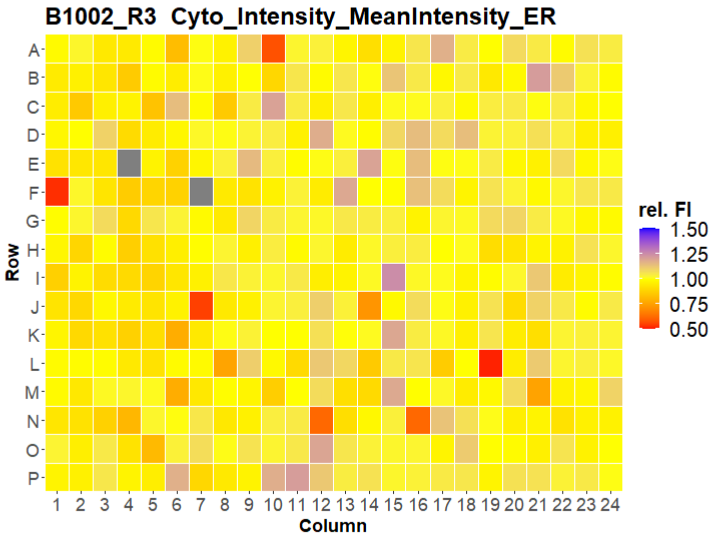
B1002_R3 Cyto_Intensity_MeanIntensity_ER
Row
Column
rel. FI
1.50
1.25
1.00
0.75
0.50

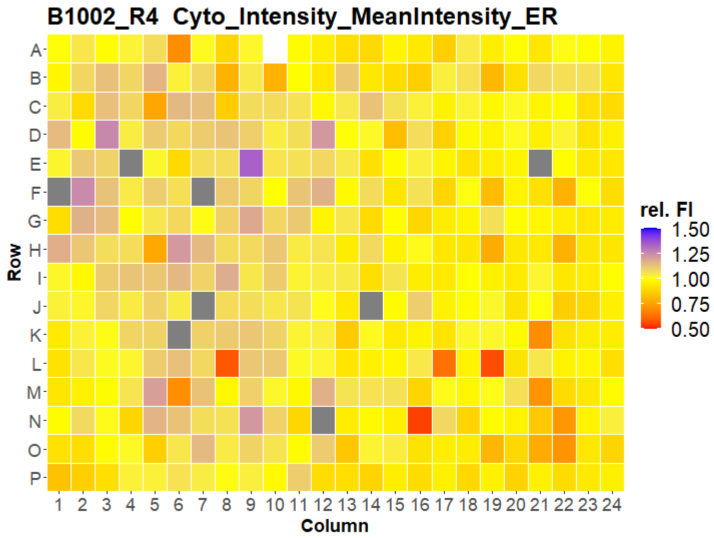
B1002_R4 Cyto_Intensity_MeanIntensity_ER
Row
Column
rel. FI
1.50
1.25
1.00
0.75
0.50

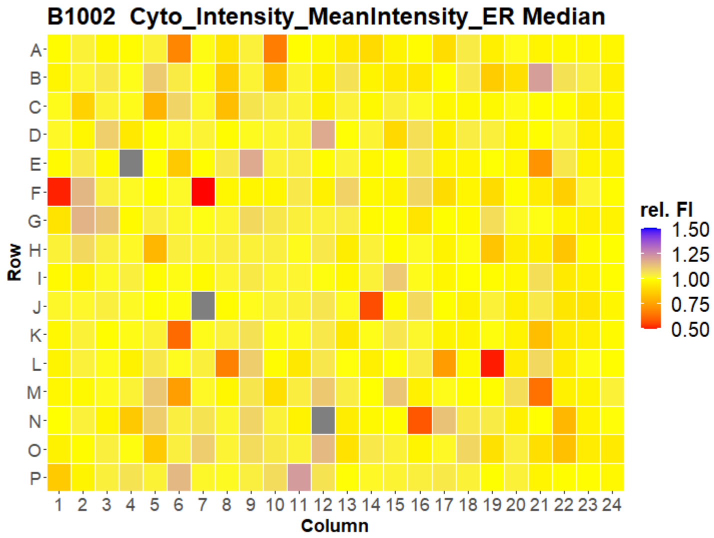
B1002 Cyto_Intensity_MeanIntensity_ER Median
Row
Column
rel. FI
1.50
1.25
1.00
0.75
0.50

### HepG2 FMP



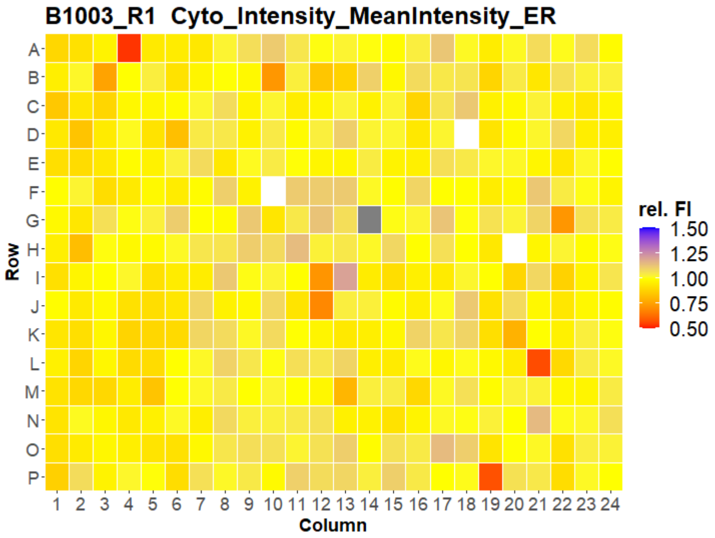


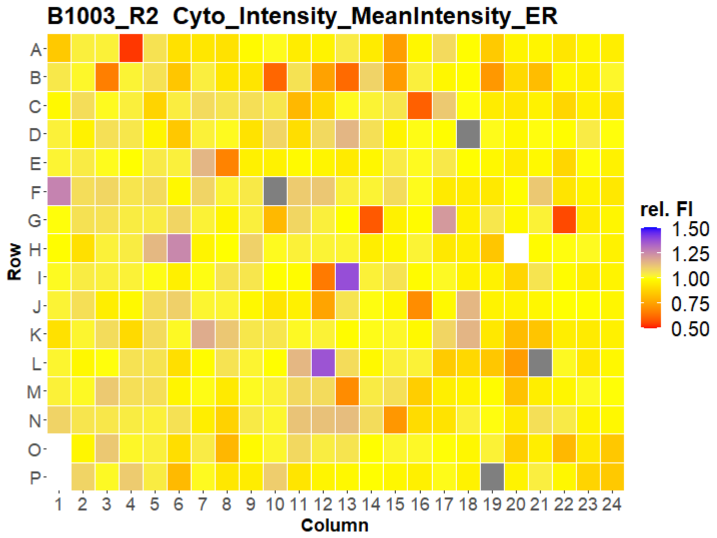


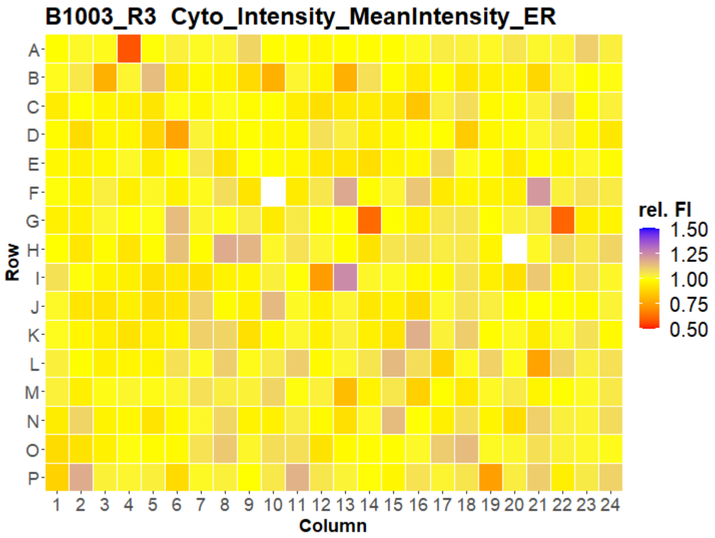


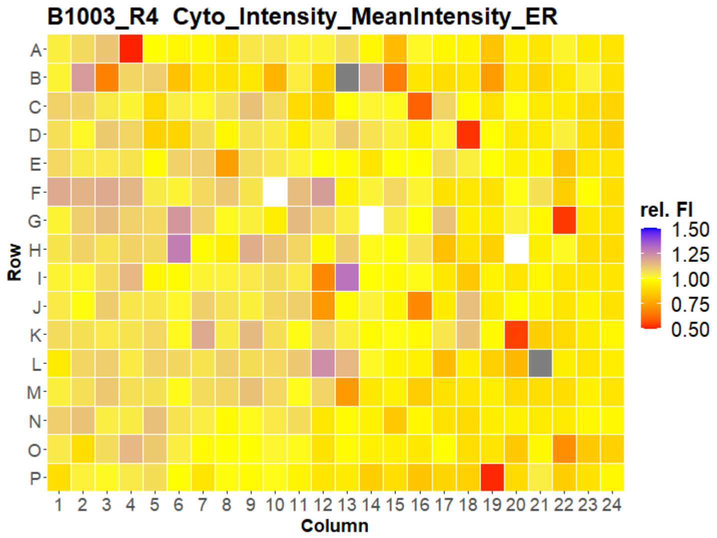


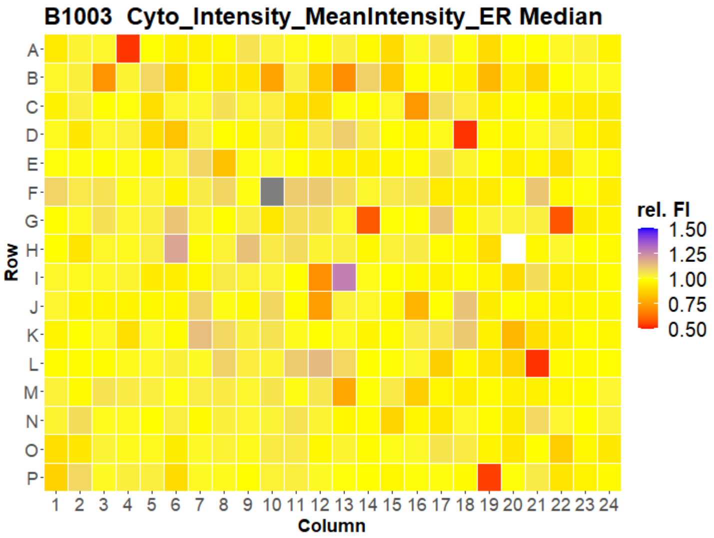



Open for Innovation KNIME

### HepG2 FMP



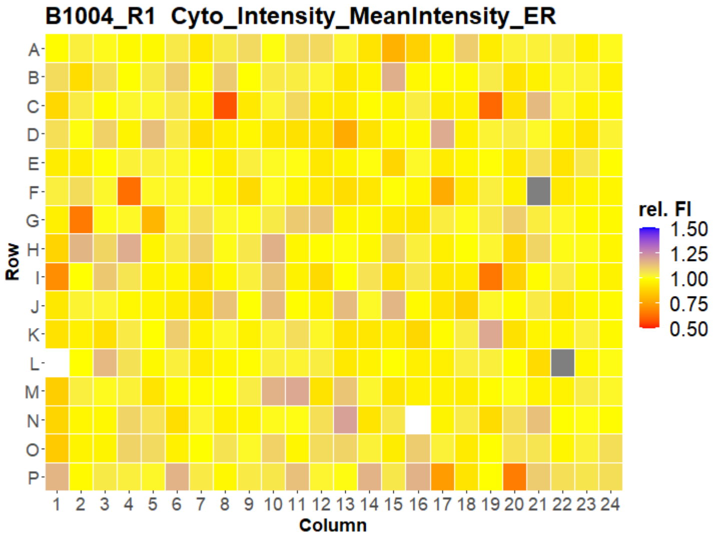
B1004_R1 Cyto_Intensity_MeanIntensity_ER
Row
Column
rel. FI
1.50
1.25
1.00
0.75
0.50

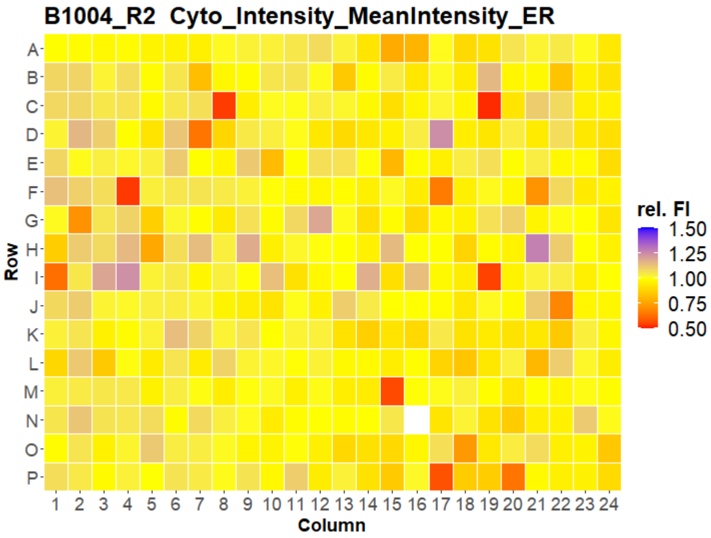
B1004_R2 Cyto_Intensity_MeanIntensity_ER
Row
Column
rel. FI
1.50
1.25
1.00
0.75
0.50

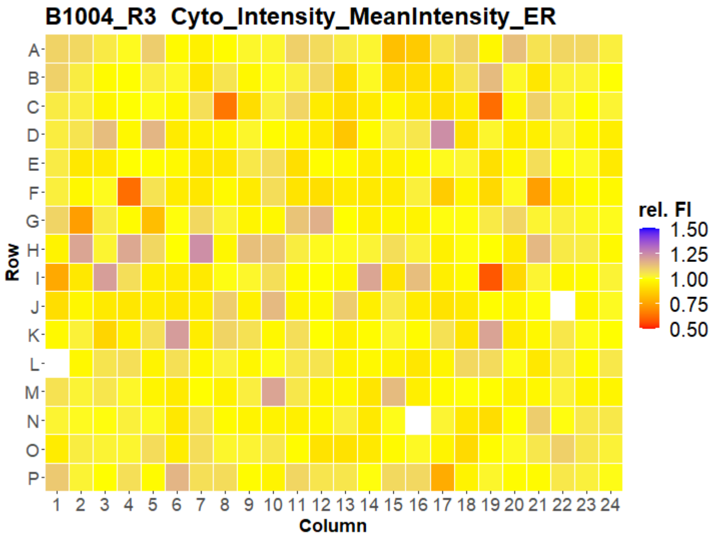
B1004_R3 Cyto_Intensity_MeanIntensity_ER
Row
Column
rel. FI
1.50
1.25
1.00
0.75
0.50

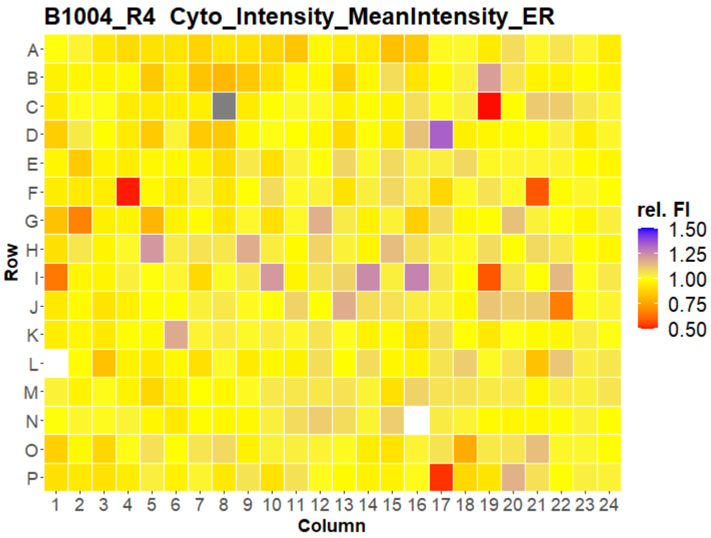
B1004_R4 Cyto_Intensity_MeanIntensity_ER
Row
Column
rel. FI
1.50
1.25
1.00
0.75
0.50

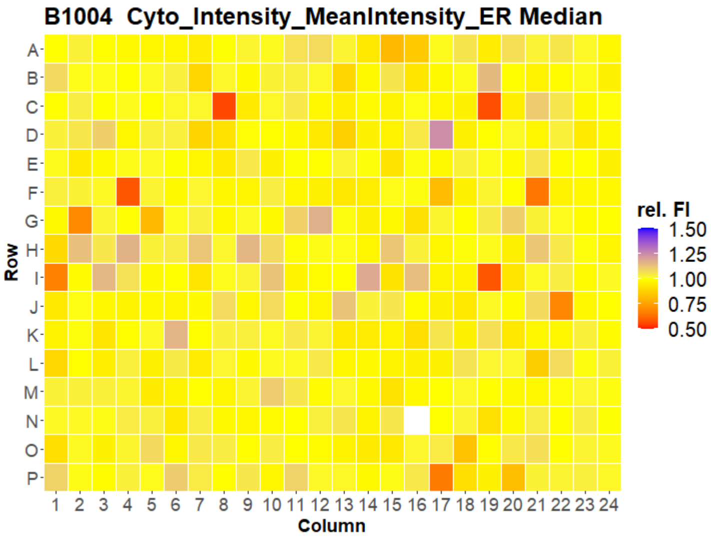
B1004 Cyto_Intensity_MeanIntensity_ER Median
Row
Column
rel. FI
1.50
1.25
1.00
0.75
0.50

### HepG2 FMP

12

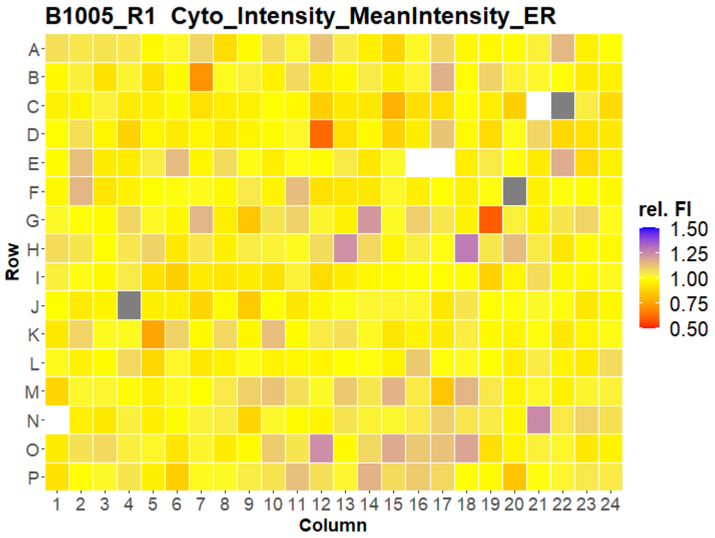
B1005_R1 Cyto_Intensity_MeanIntensity_ER
Row
Column
rel. FI
1.50
1.25
1.00
0.75
0.50

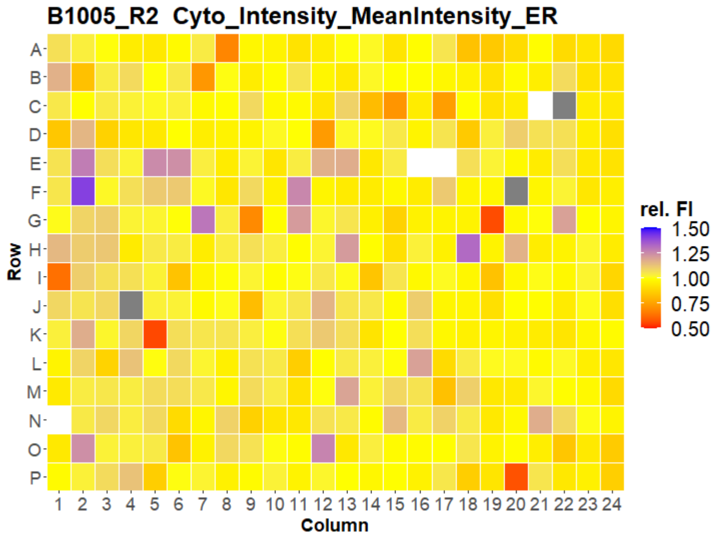
B1005_R2 Cyto_Intensity_MeanIntensity_ER
Row
Column
rel. FI
1.50
1.25
1.00
0.75
0.50

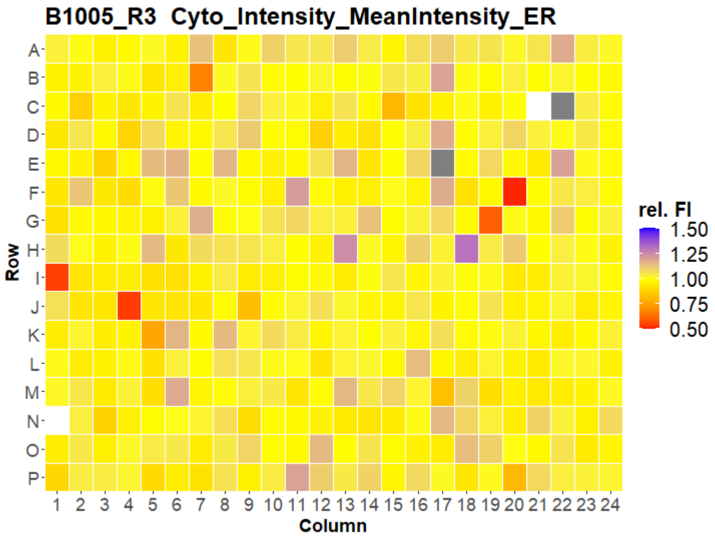
B1005_R3 Cyto_Intensity_MeanIntensity_ER
Row
Column
rel. FI
1.50
1.25
1.00
0.75
0.50

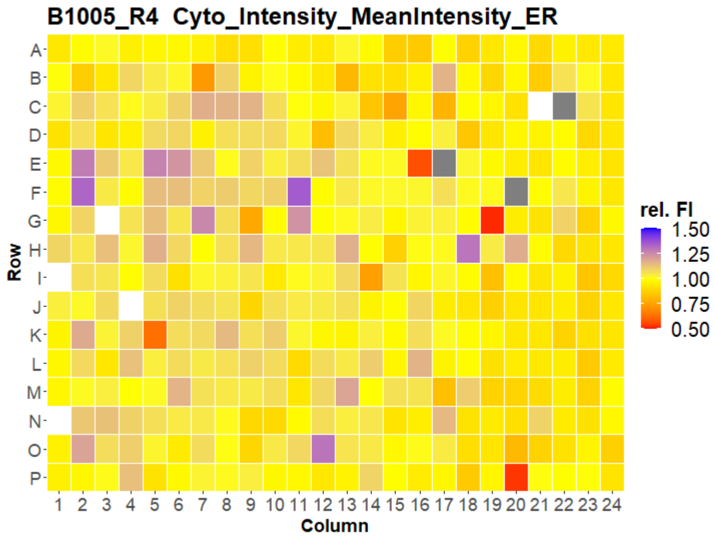
B1005_R4 Cyto_Intensity_MeanIntensity_ER
Row
Column
rel. FI
1.50
1.25
1.00
0.75
0.50

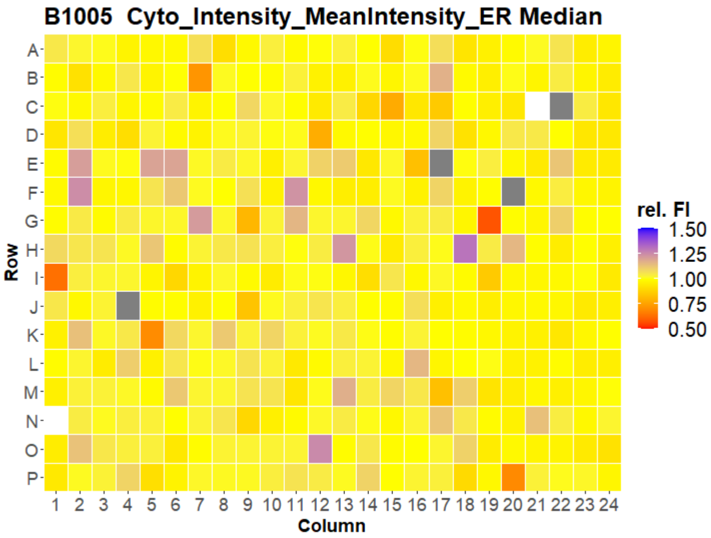
B1005 Cyto_Intensity_MeanIntensity_ER Median
Row
Column
rel. FI
1.50
1.25
1.00
0.75
0.50

### HepG2 FMP

13

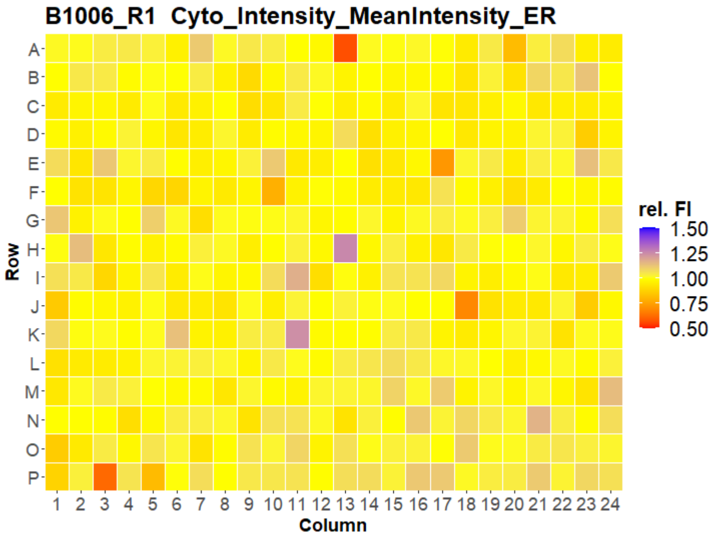


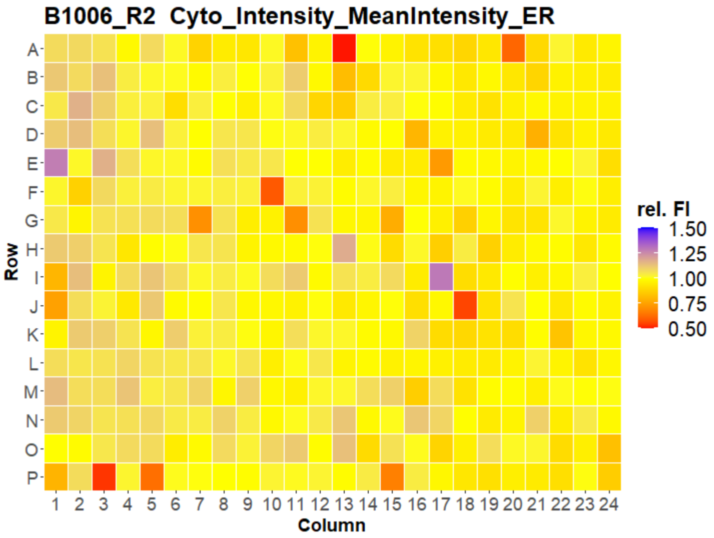


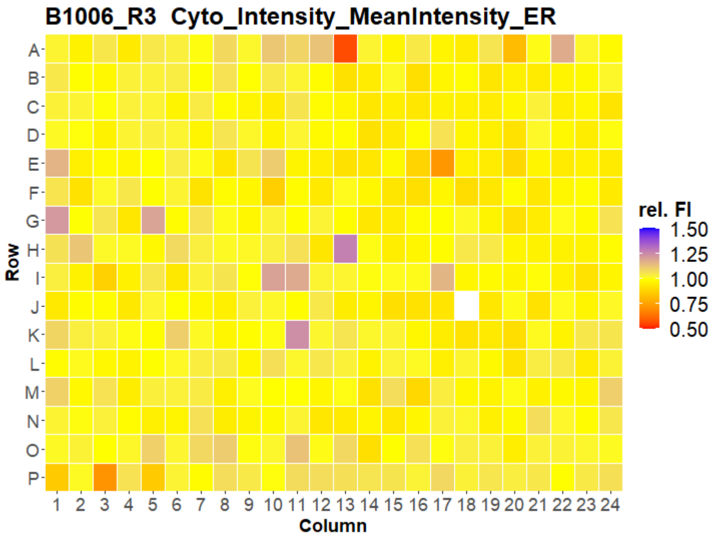


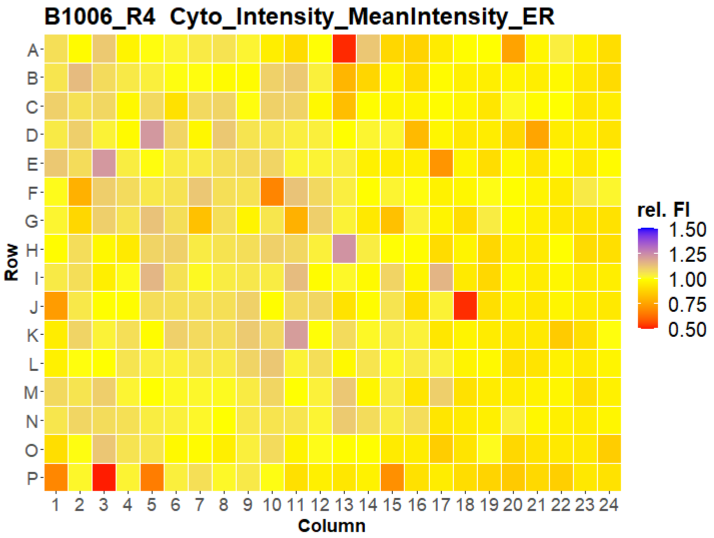


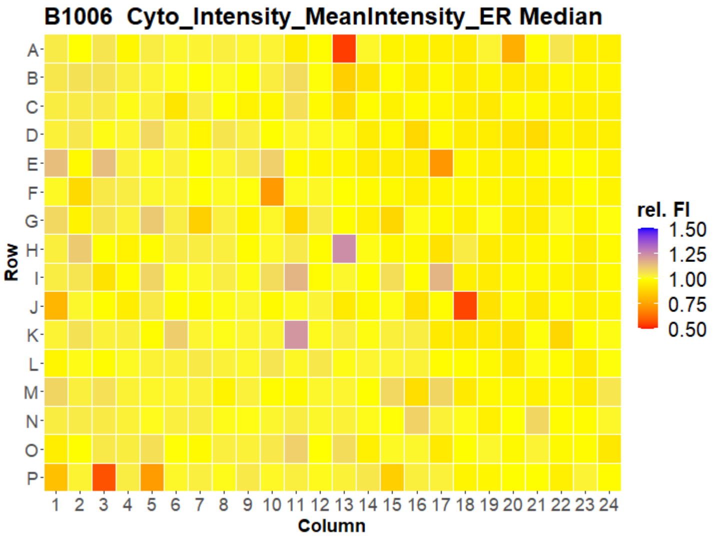




Open for Innovation KNIME

### HepG2 FMP

14

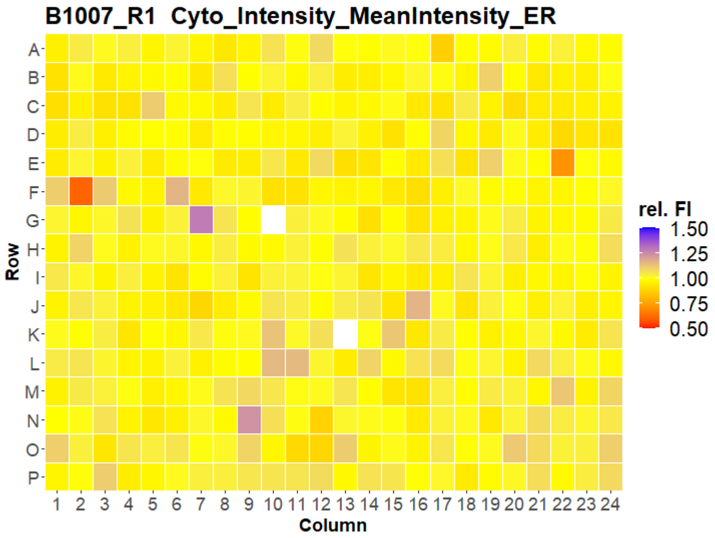
B1007_R1 Cyto_Intensity_MeanIntensity_ER
Row
A B C D E F G H I J K L M N O P
1 2 3 4 5 6 7 8 9 10 11 12 13 14 15 16 17 18 19 20 21 22 23 24
Column
rel. FI
1.50
1.25
1.00
0.75
0.50

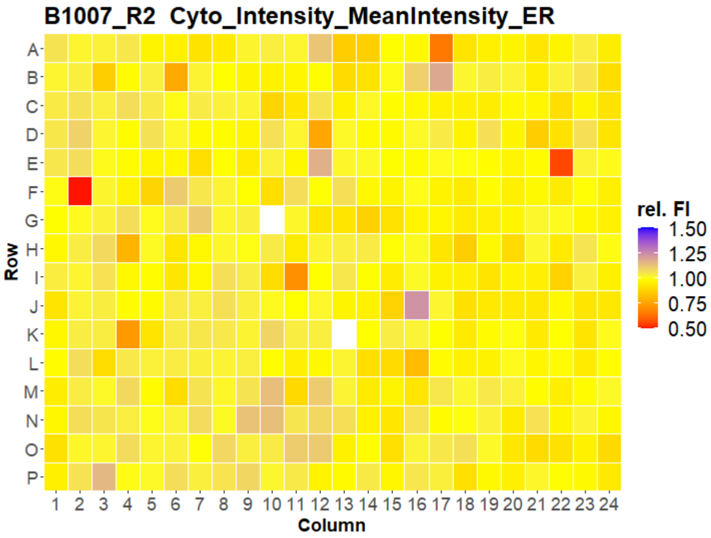
B1007_R2 Cyto_Intensity_MeanIntensity_ER
Row
A B C D E F G H I J K L M N O P
1 2 3 4 5 6 7 8 9 10 11 12 13 14 15 16 17 18 19 20 21 22 23 24
Column
rel. FI
1.50
1.25
1.00
0.75
0.50

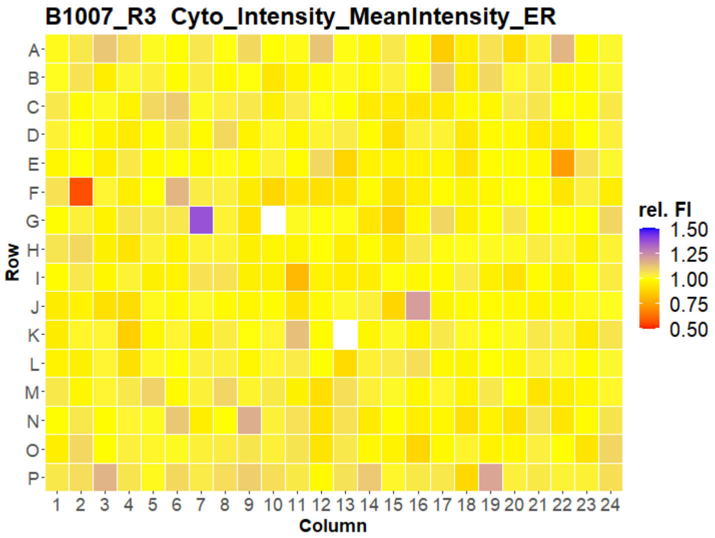
B1007_R3 Cyto_Intensity_MeanIntensity_ER
Row
A B C D E F G H I J K L M N O P
1 2 3 4 5 6 7 8 9 10 11 12 13 14 15 16 17 18 19 20 21 22 23 24
Column
rel. FI
1.50
1.25
1.00
0.75
0.50

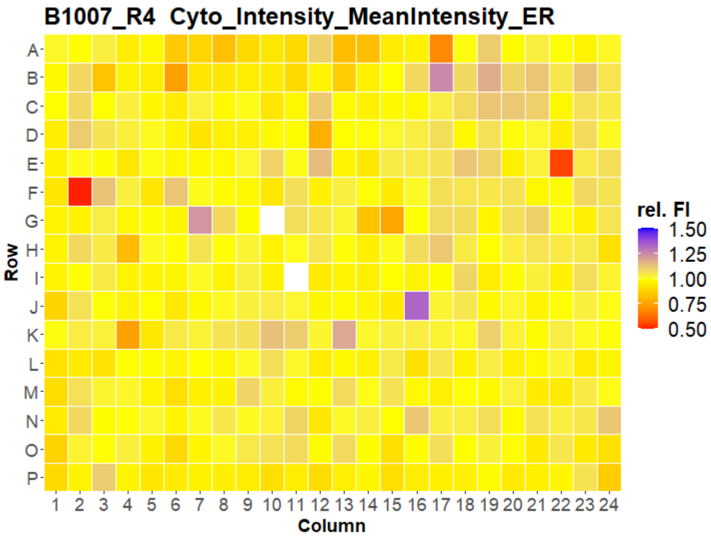
B1007_R4 Cyto_Intensity_MeanIntensity_ER
Row
A B C D E F G H I J K L M N O P
1 2 3 4 5 6 7 8 9 10 11 12 13 14 15 16 17 18 19 20 21 22 23 24
Column
rel. FI
1.50
1.25
1.00
0.75
0.50

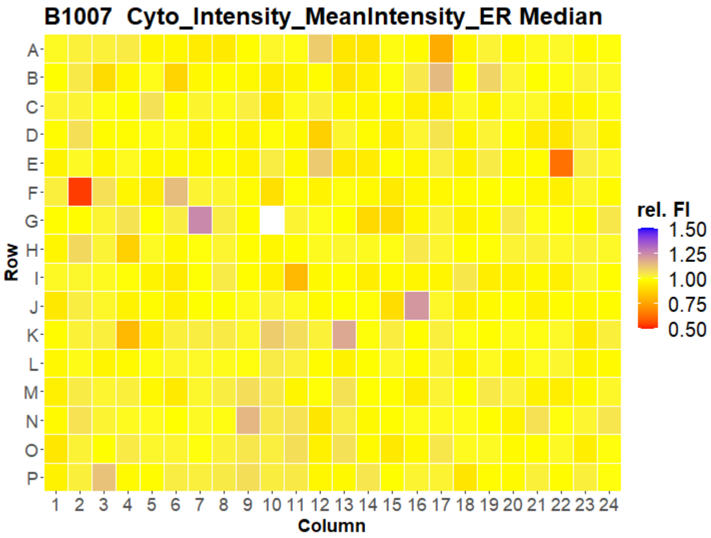
B1007 Cyto_Intensity_MeanIntensity_ER Median
Row
A B C D E F G H I J K L M N O P
1 2 3 4 5 6 7 8 9 10 11 12 13 14 15 16 17 18 19 20 21 22 23 24
Column
rel. FI
1.50
1.25
1.00
0.75
0.50

### HepG2 FMP

15

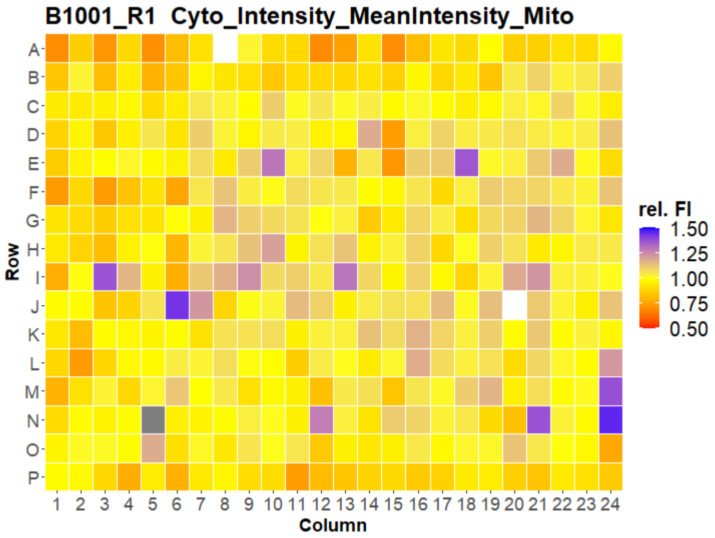
B1001_R1 Cyto_Intensity_MeanIntensity_Mito
Row
Column
rel. FI
1.50
1.25
1.00
0.75
0.50

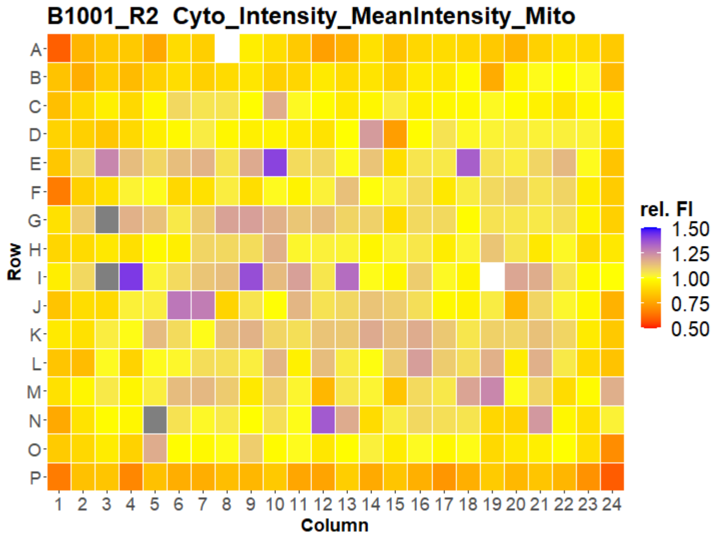
B1001_R2 Cyto_Intensity_MeanIntensity_Mito
Row
Column
rel. FI
1.50
1.25
1.00
0.75
0.50

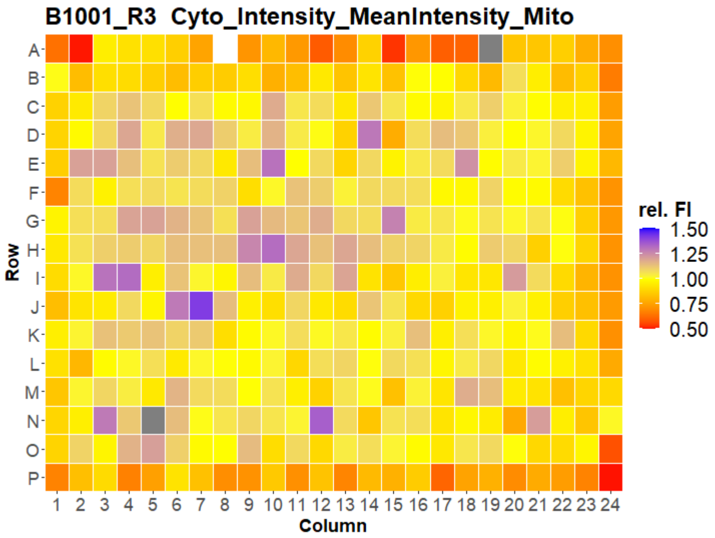
B1001_R3 Cyto_Intensity_MeanIntensity_Mito
Row
Column
rel. FI
1.50
1.25
1.00
0.75
0.50

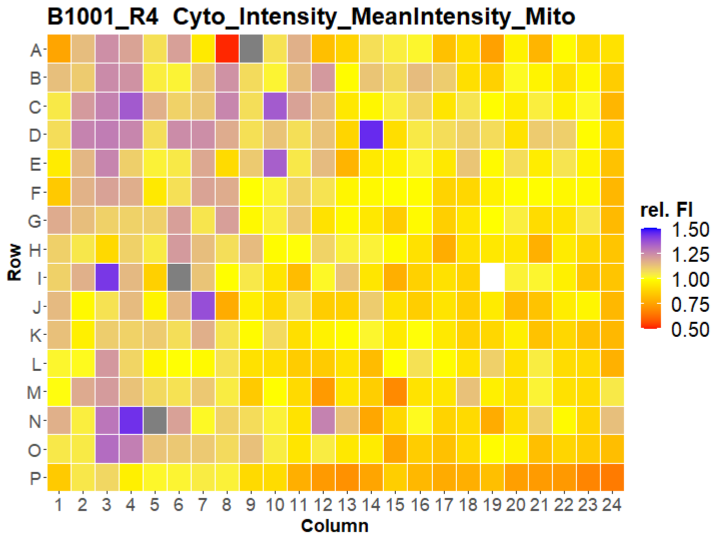
B1001_R4 Cyto_Intensity_MeanIntensity_Mito
Row
Column
rel. FI
1.50
1.25
1.00
0.75
0.50

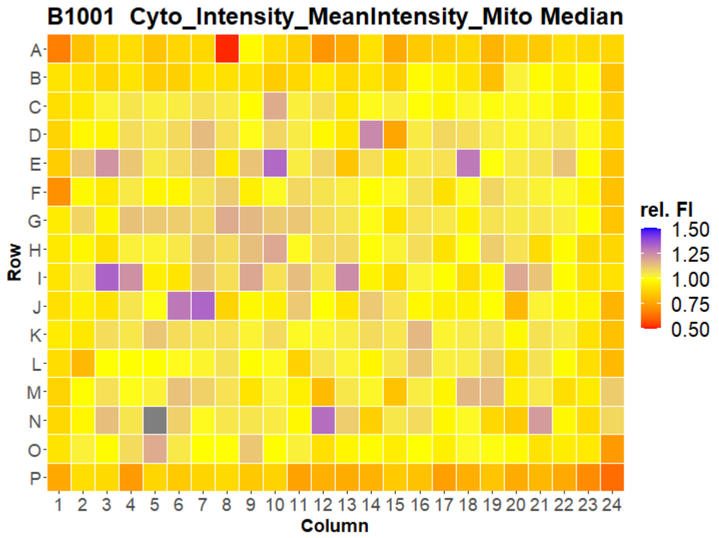
B1001 Cyto_Intensity_MeanIntensity_Mito Median
Row
Column
rel. FI
1.50
1.25
1.00
0.75
0.50

### HepG2 FMP

16

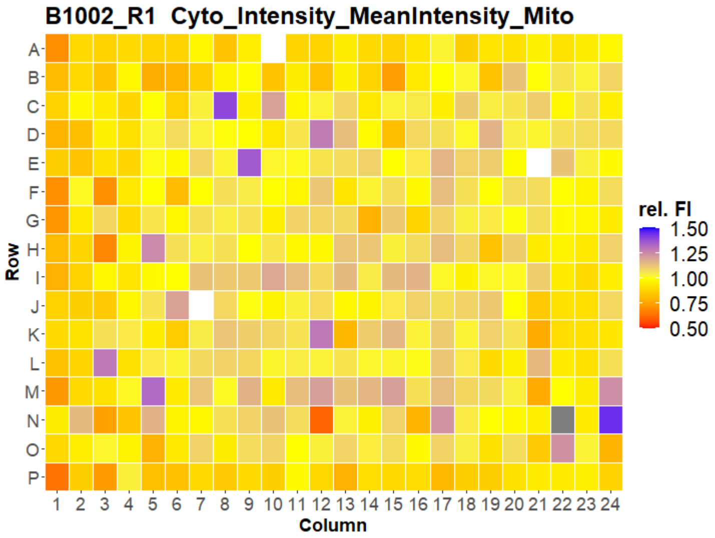
B1002_R1 Cyto_Intensity_MeanIntensity_Mito
Row
Column
rel. FI
1.50
1.25
1.00
0.75
0.50

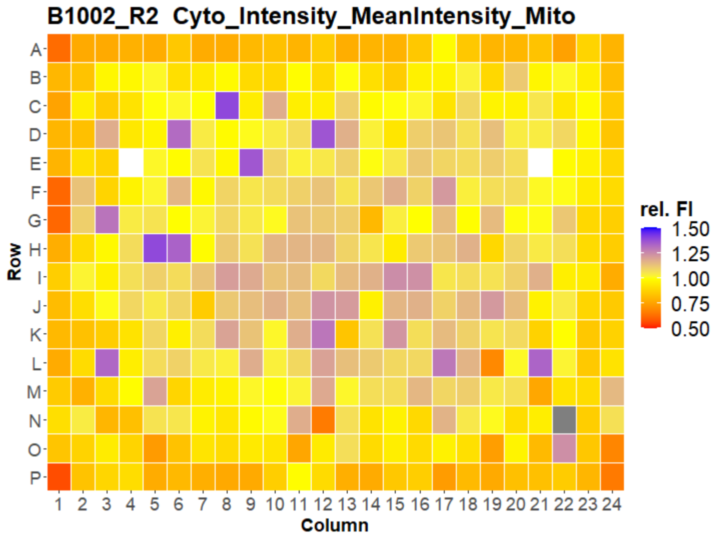
B1002_R2 Cyto_Intensity_MeanIntensity_Mito
Row
Column
rel. FI
1.50
1.25
1.00
0.75
0.50

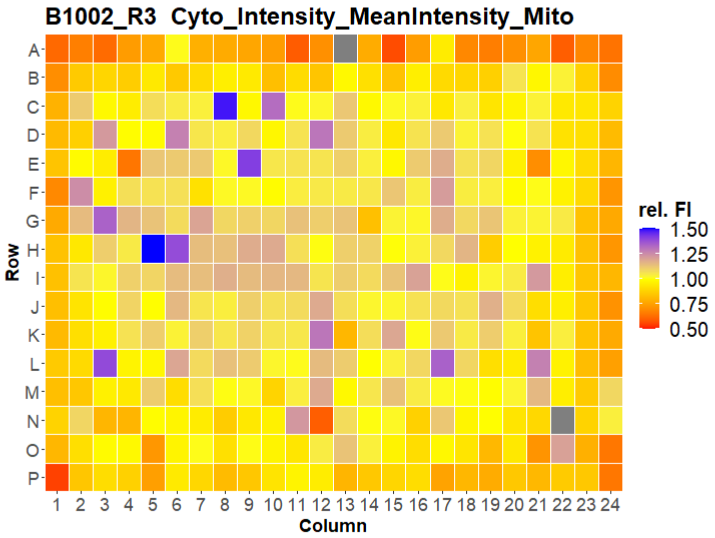
B1002_R3 Cyto_Intensity_MeanIntensity_Mito
Row
Column
rel. FI
1.50
1.25
1.00
0.75
0.50

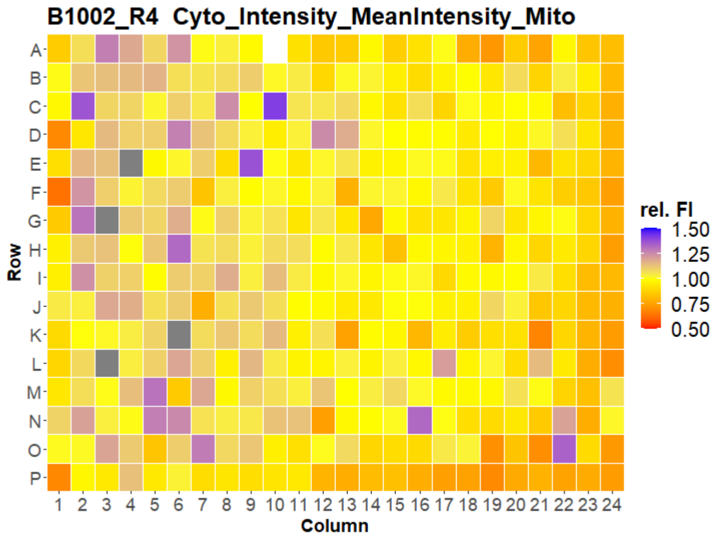
B1002_R4 Cyto_Intensity_MeanIntensity_Mito
Row
Column
rel. FI
1.50
1.25
1.00
0.75
0.50

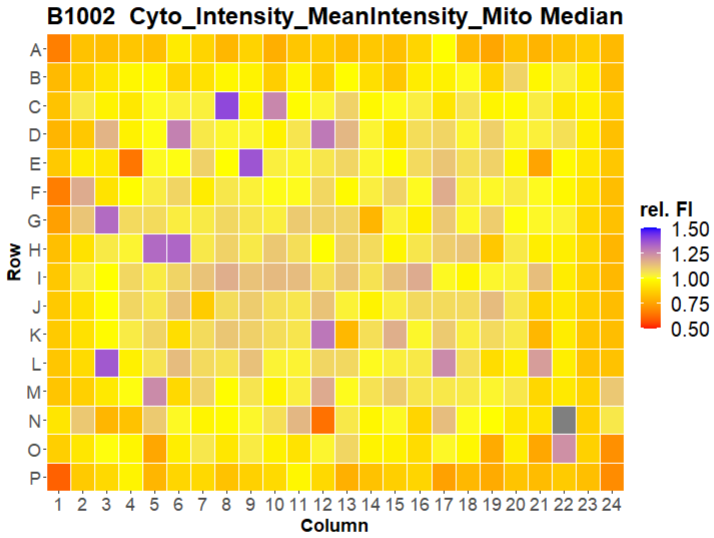
B1002 Cyto_Intensity_MeanIntensity_Mito Median
Row
Column
rel. FI
1.50
1.25
1.00
0.75
0.50

### HepG2 FMP

17

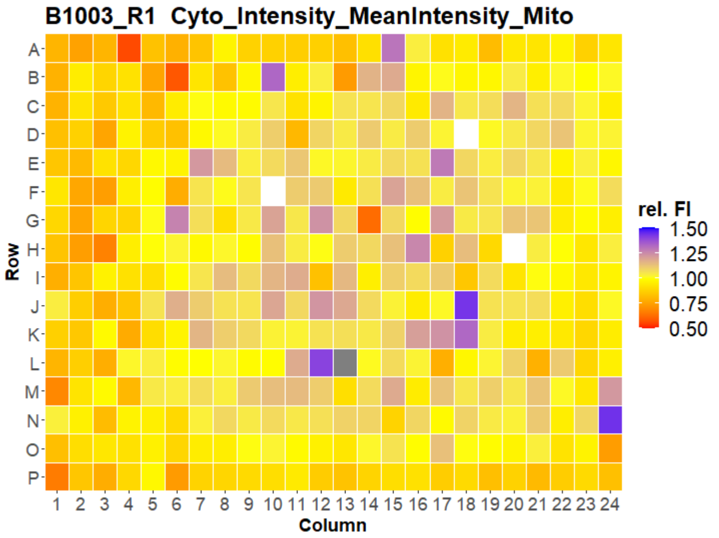
B1003_R1 Cyto_Intensity_MeanIntensity_Mito
Row
A B C D E F G H I J K L M N O P
Column
1 2 3 4 5 6 7 8 9 10 11 12 13 14 15 16 17 18 19 20 21 22 23 24
rel. FI
1.50
1.25
1.00
0.75
0.50

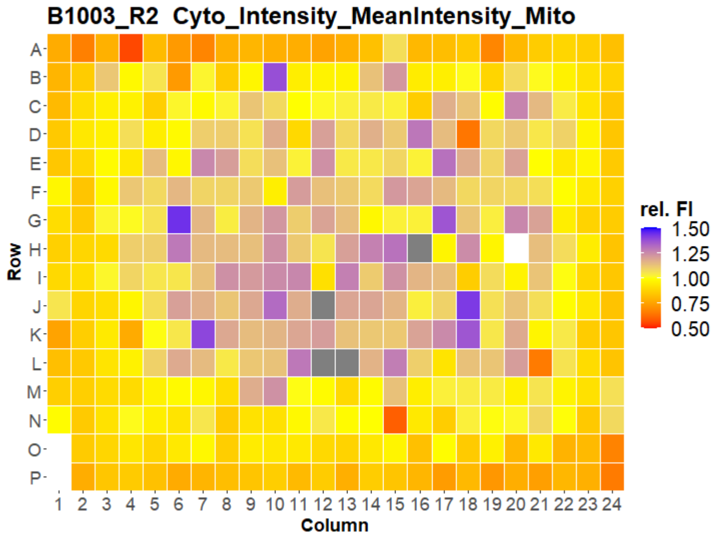
B1003_R2 Cyto_Intensity_MeanIntensity_Mito
Row
A B C D E F G H I J K L M N O P
Column
1 2 3 4 5 6 7 8 9 10 11 12 13 14 15 16 17 18 19 20 21 22 23 24
rel. FI
1.50
1.25
1.00
0.75
0.50

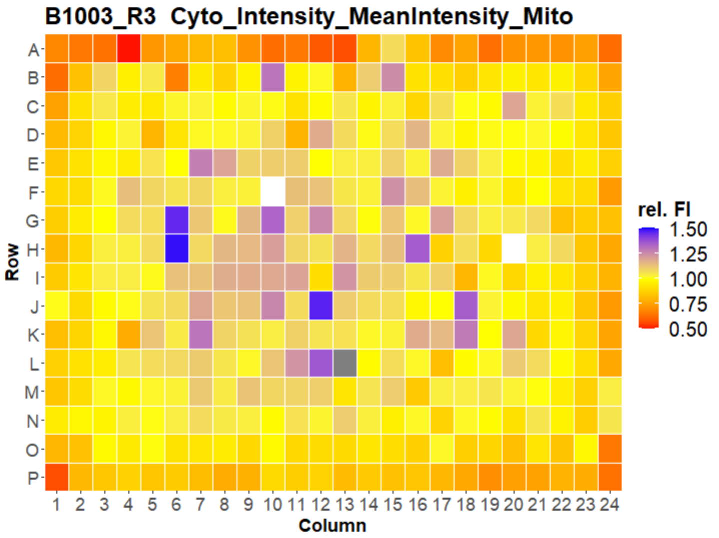
B1003_R3 Cyto_Intensity_MeanIntensity_Mito
Row
A B C D E F G H I J K L M N O P
Column
1 2 3 4 5 6 7 8 9 10 11 12 13 14 15 16 17 18 19 20 21 22 23 24
rel. FI
1.50
1.25
1.00
0.75
0.50

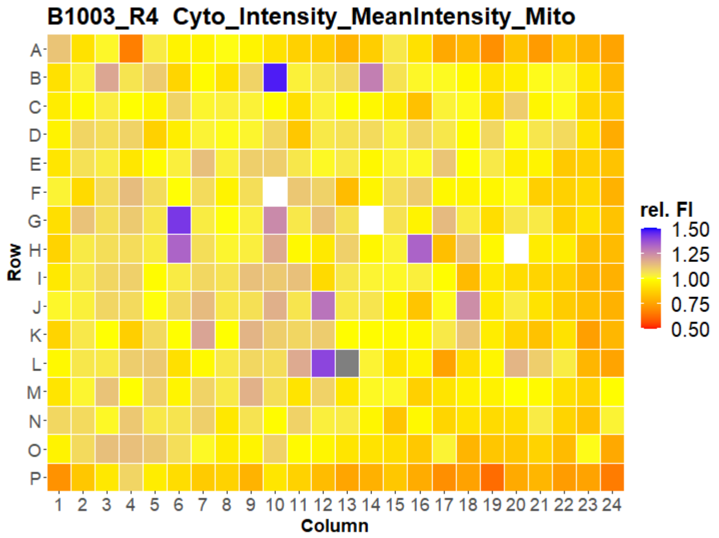
B1003_R4 Cyto_Intensity_MeanIntensity_Mito
Row
A B C D E F G H I J K L M N O P
Column
1 2 3 4 5 6 7 8 9 10 11 12 13 14 15 16 17 18 19 20 21 22 23 24
rel. FI
1.50
1.25
1.00
0.75
0.50

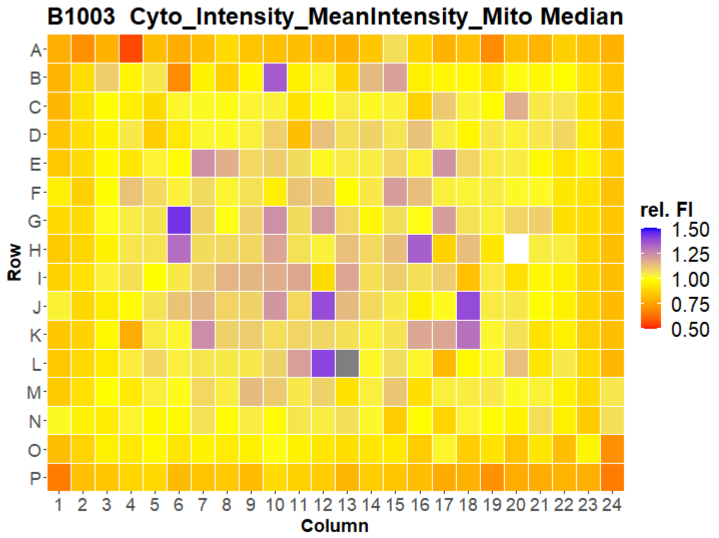
B1003 Cyto_Intensity_MeanIntensity_Mito Median
Row
A B C D E F G H I J K L M N O P
Column
1 2 3 4 5 6 7 8 9 10 11 12 13 14 15 16 17 18 19 20 21 22 23 24
rel. FI
1.50
1.25
1.00
0.75
0.50

### HepG2 FMP



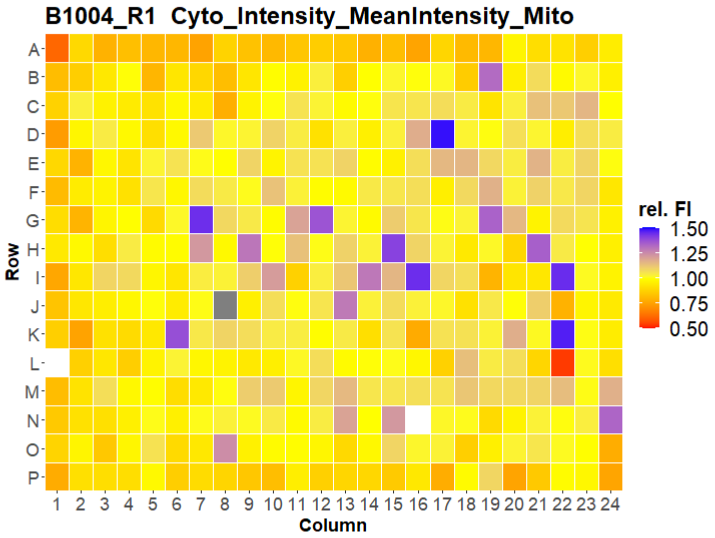
B1004_R1 Cyto_Intensity_MeanIntensity_Mito
Row
Column
rel. FI
1.50
1.25
1.00
0.75
0.50

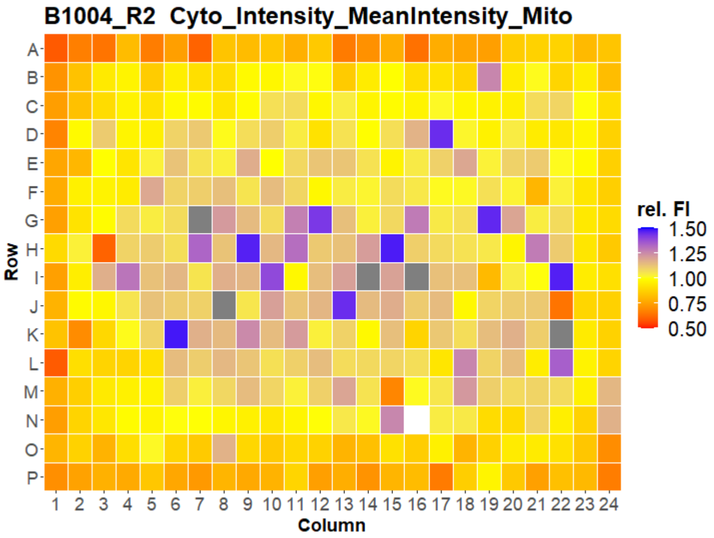
B1004_R2 Cyto_Intensity_MeanIntensity_Mito
Row
Column
rel. FI
1.50
1.25
1.00
0.75
0.50

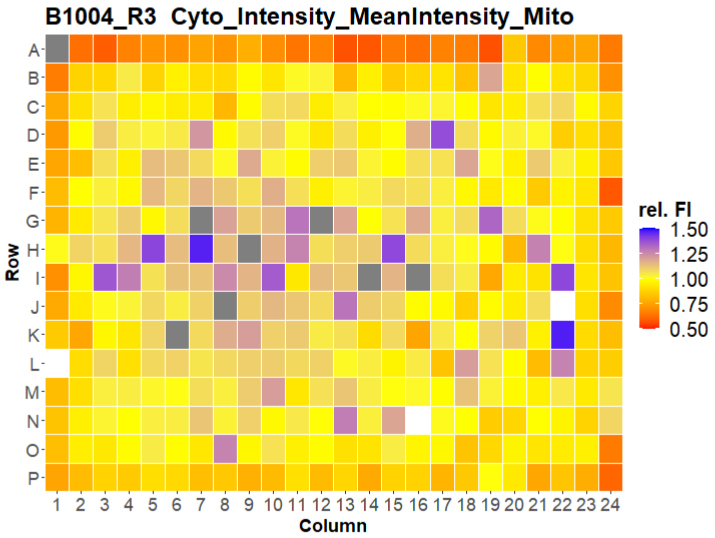
B1004_R3 Cyto_Intensity_MeanIntensity_Mito
Row
Column
rel. FI
1.50
1.25
1.00
0.75
0.50

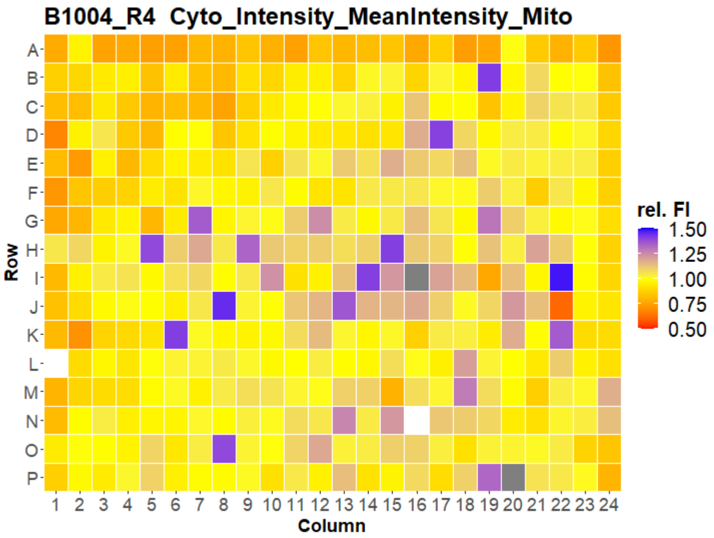
B1004_R4 Cyto_Intensity_MeanIntensity_Mito
Row
Column
rel. FI
1.50
1.25
1.00
0.75
0.50

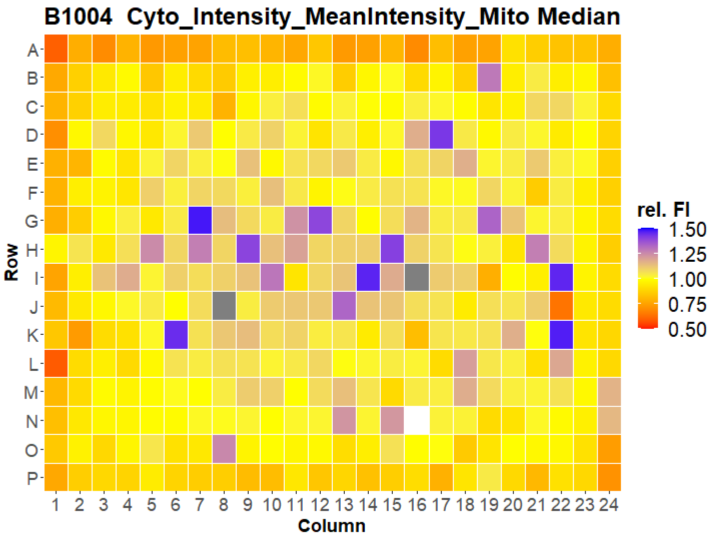
B1004 Cyto_Intensity_MeanIntensity_Mito Median
Row
Column
rel. FI
1.50
1.25
1.00
0.75
0.50

### HepG2 FMP

19

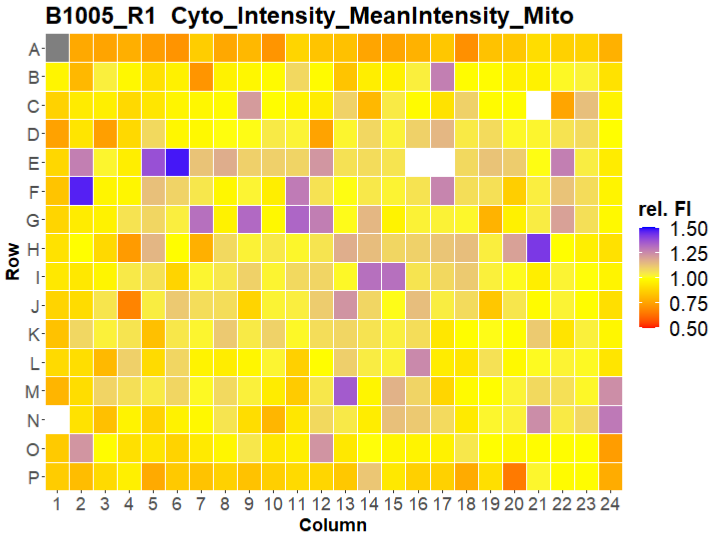

B1005_R1 Cyto_Intensity_MeanIntensity_Mito
Row
Column
rel. FI
1.50
1.25
1.00
0.75
0.50


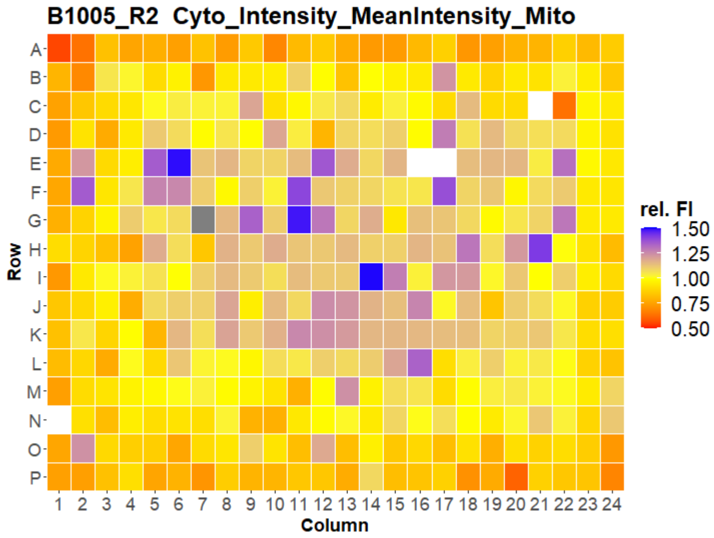

B1005_R2 Cyto_Intensity_MeanIntensity_Mito
Row
Column
rel. FI
1.50
1.25
1.00
0.75
0.50


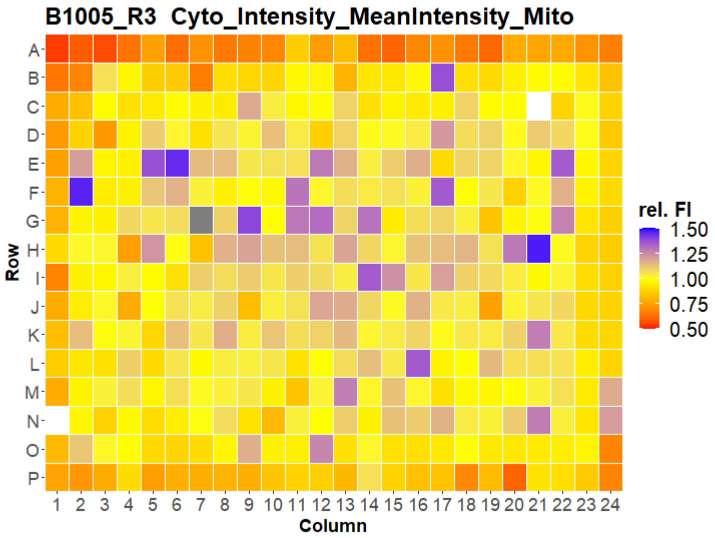

B1005_R3 Cyto_Intensity_MeanIntensity_Mito
Row
Column
rel. FI
1.50
1.25
1.00
0.75
0.50


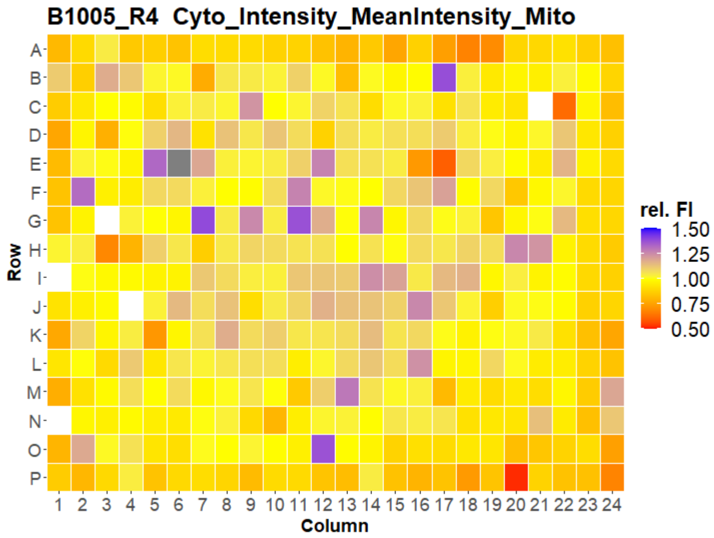

B1005_R4 Cyto_Intensity_MeanIntensity_Mito
Row
Column
rel. FI
1.50
1.25
1.00
0.75
0.50


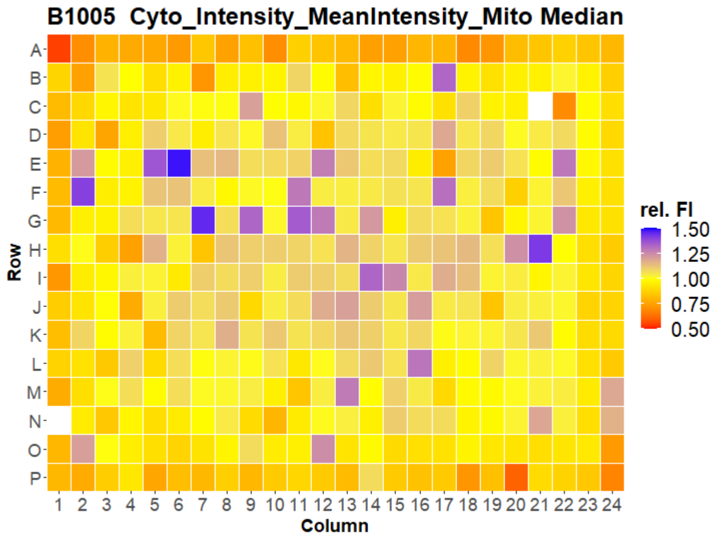

B1005 Cyto_Intensity_MeanIntensity_Mito Median
Row
Column
rel. FI
1.50
1.25
1.00
0.75
0.50

### HepG2 FMP



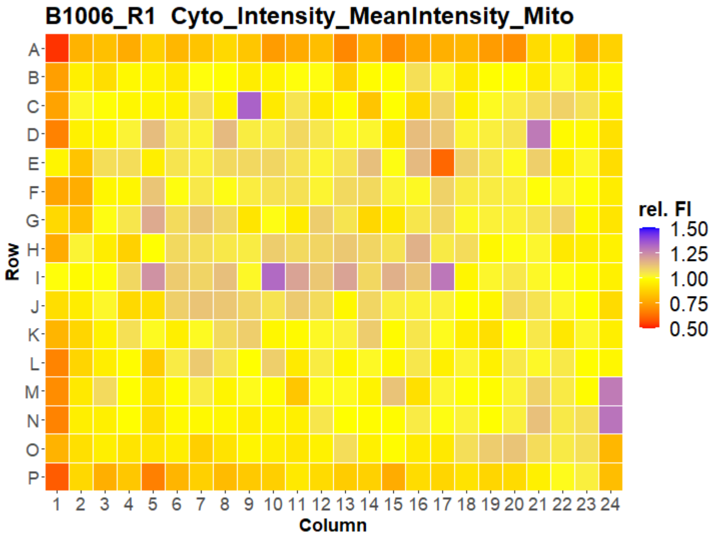
B1006_R1 Cyto_Intensity_MeanIntensity_Mito
Row
Column
rel. FI
1.50
1.25
1.00
0.75
0.50

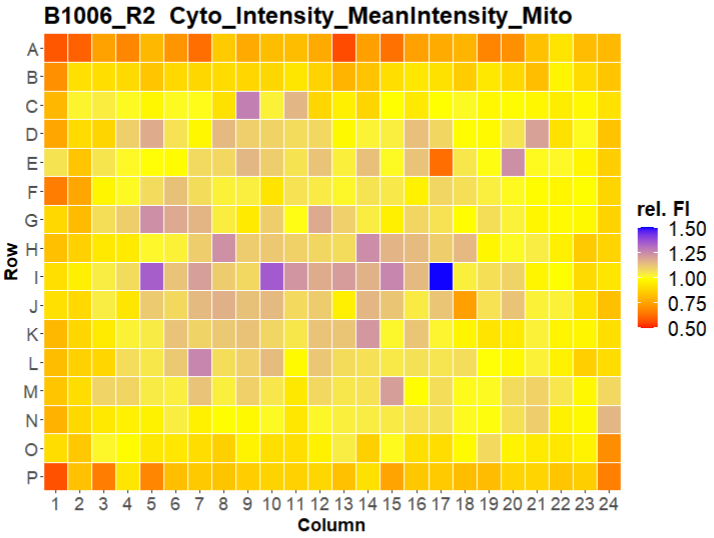
B1006_R2 Cyto_Intensity_MeanIntensity_Mito
Row
Column
rel. FI
1.50
1.25
1.00
0.75
0.50

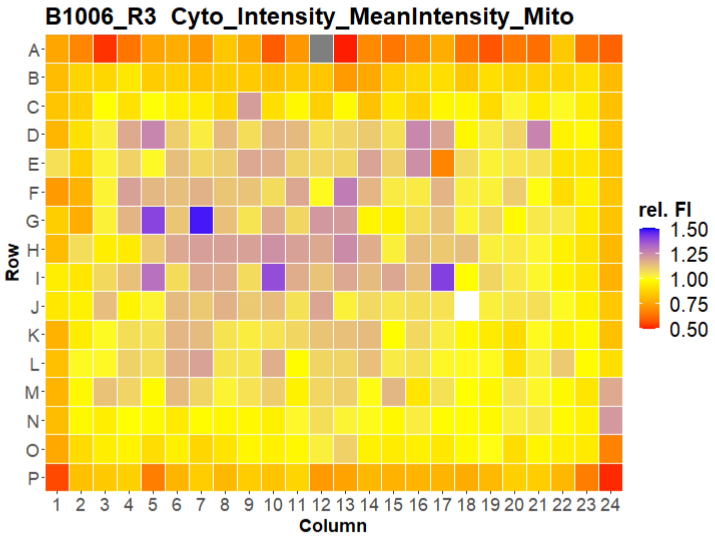
B1006_R3 Cyto_Intensity_MeanIntensity_Mito
Row
Column
rel. FI
1.50
1.25
1.00
0.75
0.50

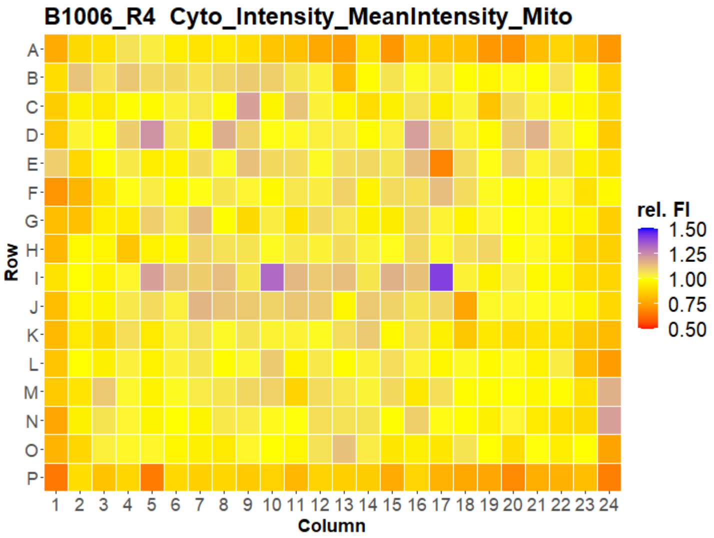
B1006_R4 Cyto_Intensity_MeanIntensity_Mito
Row
Column
rel. FI
1.50
1.25
1.00
0.75
0.50

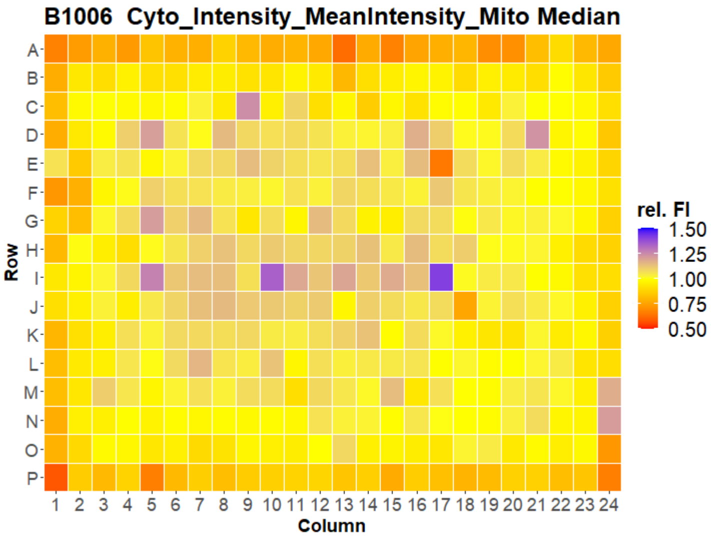
B1006 Cyto_Intensity_MeanIntensity_Mito Median
Row
Column
rel. FI
1.50
1.25
1.00
0.75
0.50

### HepG2 FMP



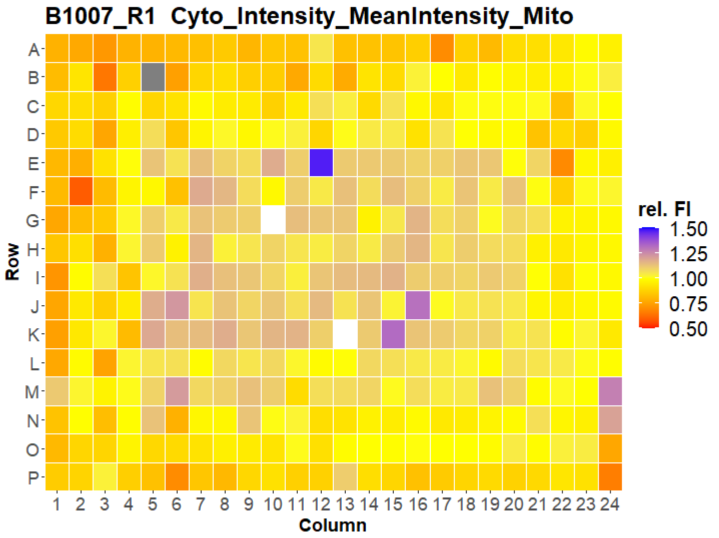
B1007_R1 Cyto_Intensity_MeanIntensity_Mito
A B C D E F G H I J K L M N O P
Row
1 2 3 4 5 6 7 8 9 10 11 12 13 14 15 16 17 18 19 20 21 22 23 24
Column
rel. FI
1.50
1.25
1.00
0.75
0.50

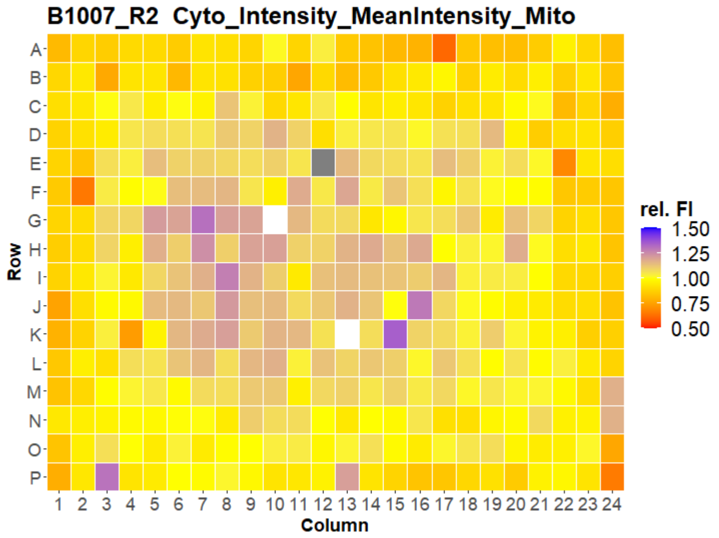
B1007_R2 Cyto_Intensity_MeanIntensity_Mito
A B C D E F G H I J K L M N O P
Row
1 2 3 4 5 6 7 8 9 10 11 12 13 14 15 16 17 18 19 20 21 22 23 24
Column
rel. FI
1.50
1.25
1.00
0.75
0.50

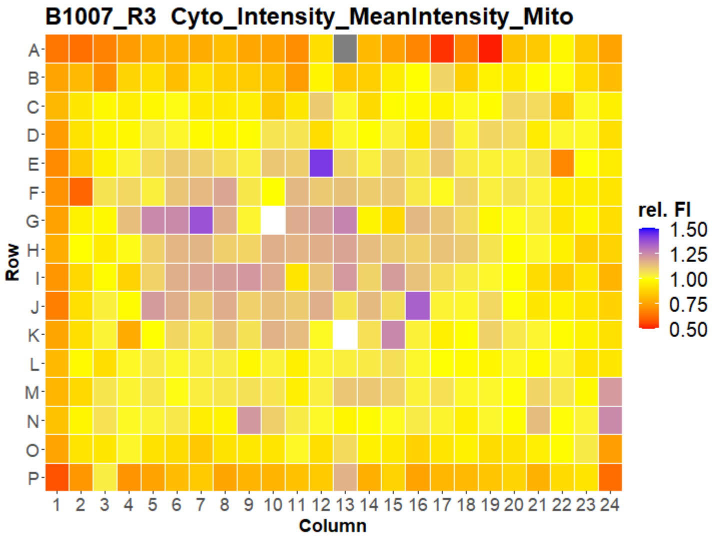
B1007_R3 Cyto_Intensity_MeanIntensity_Mito
A B C D E F G H I J K L M N O P
Row
1 2 3 4 5 6 7 8 9 10 11 12 13 14 15 16 17 18 19 20 21 22 23 24
Column
rel. FI
1.50
1.25
1.00
0.75
0.50

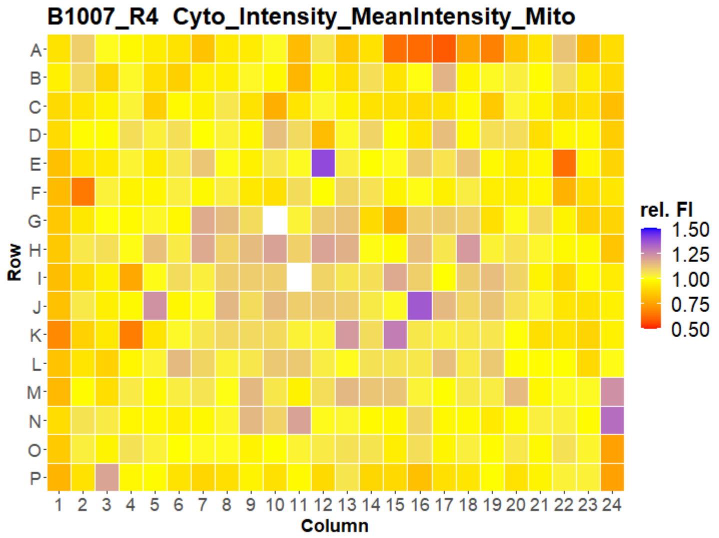
B1007_R4 Cyto_Intensity_MeanIntensity_Mito
A B C D E F G H I J K L M N O P
Row
1 2 3 4 5 6 7 8 9 10 11 12 13 14 15 16 17 18 19 20 21 22 23 24
Column
rel. FI
1.50
1.25
1.00
0.75
0.50

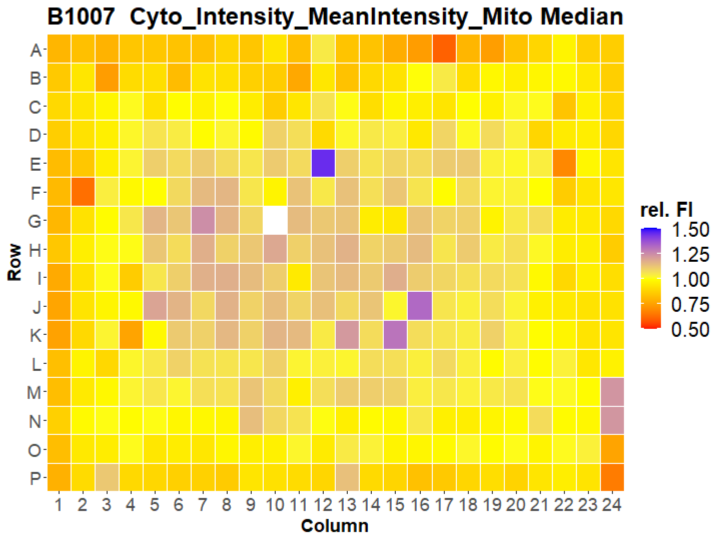
B1007 Cyto_Intensity_MeanIntensity_Mito Median
A B C D E F G H I J K L M N O P
Row
1 2 3 4 5 6 7 8 9 10 11 12 13 14 15 16 17 18 19 20 21 22 23 24
Column
rel. FI
1.50
1.25
1.00
0.75
0.50

### HepG2 FMP

22

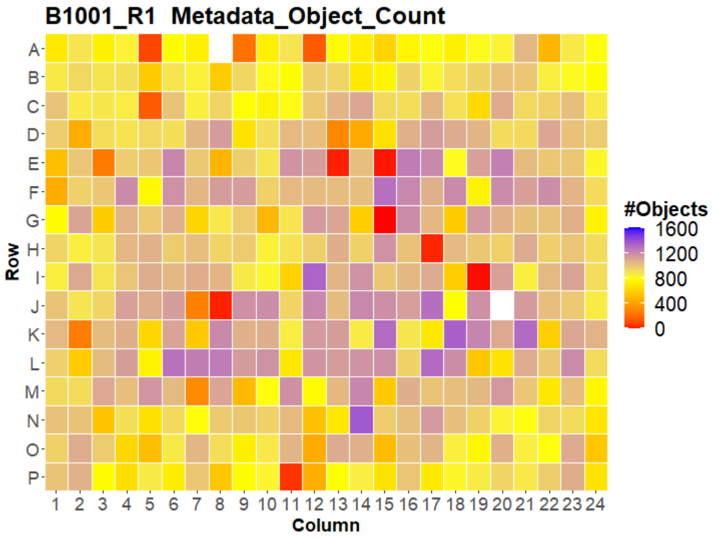
B1001_R1 Metadata_Object_Count
Row
Column
#Objects
1600
1200
800
400
0

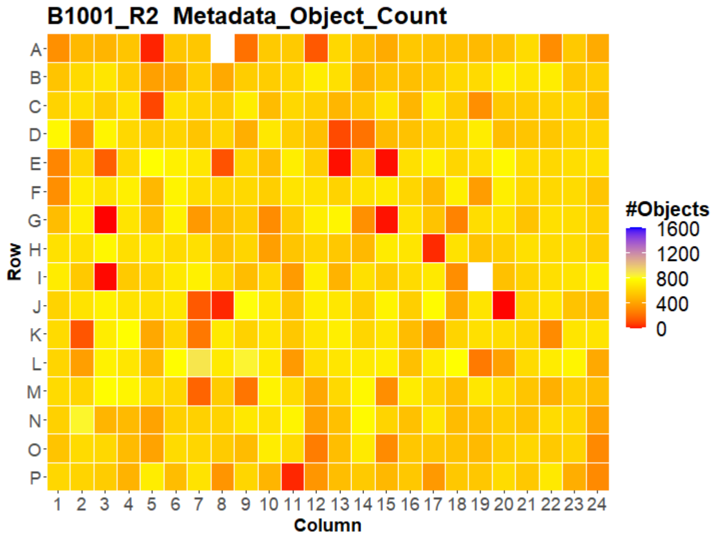
B1001_R2 Metadata_Object_Count
Row
Column
#Objects
1600
1200
800
400
0

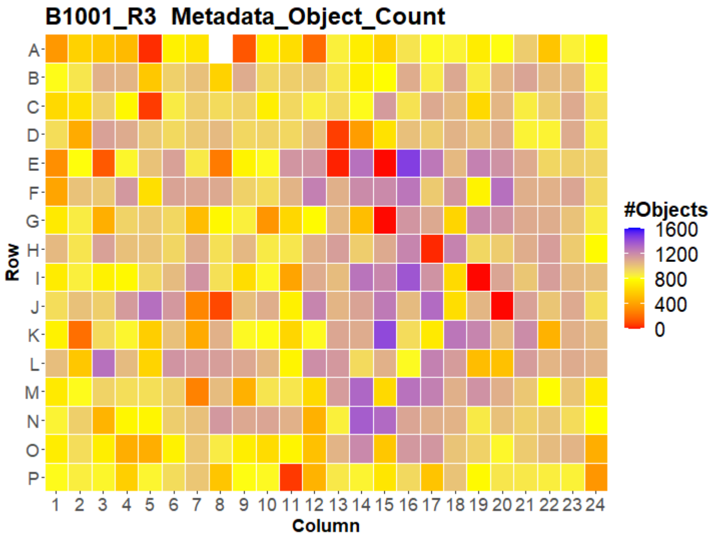
B1001_R3 Metadata_Object_Count
Row
Column
#Objects
1600
1200
800
400
0

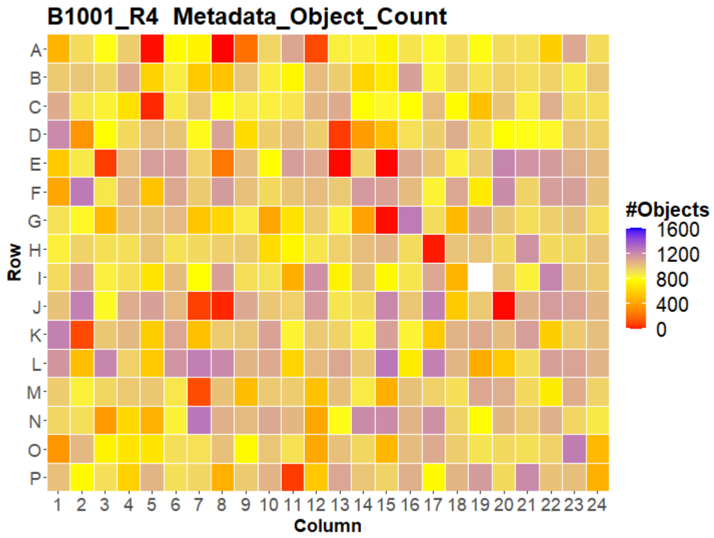
B1001_R4 Metadata_Object_Count
Row
Column
#Objects
1600
1200
800
400
0

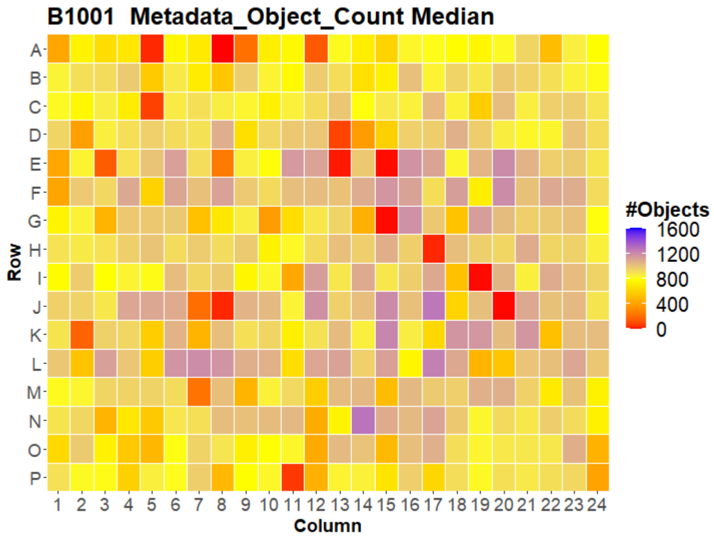
B1001 Metadata_Object_Count Median
Row
Column
#Objects
1600
1200
800
400
0

### HepG2 FMP



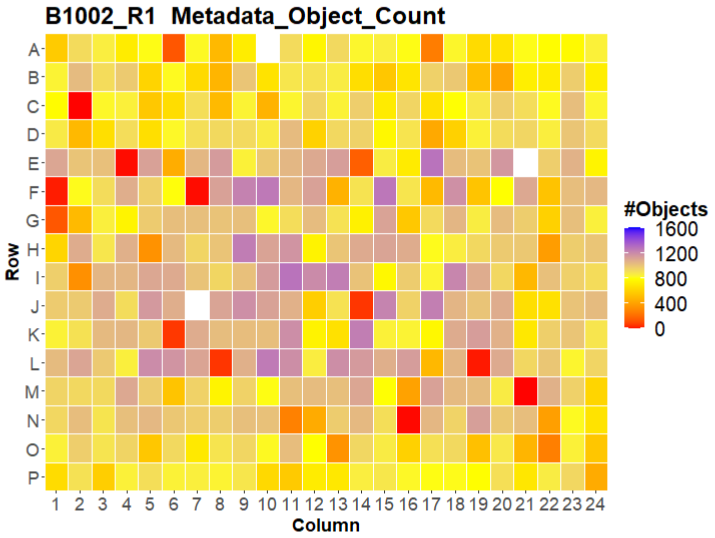
B1002_R1 Metadata_Object_Count
Row
A B C D E F G H I J K L M N O P
1 2 3 4 5 6 7 8 9 10 11 12 13 14 15 16 17 18 19 20 21 22 23 24
Column
#Objects
1600
1200
800
400
0

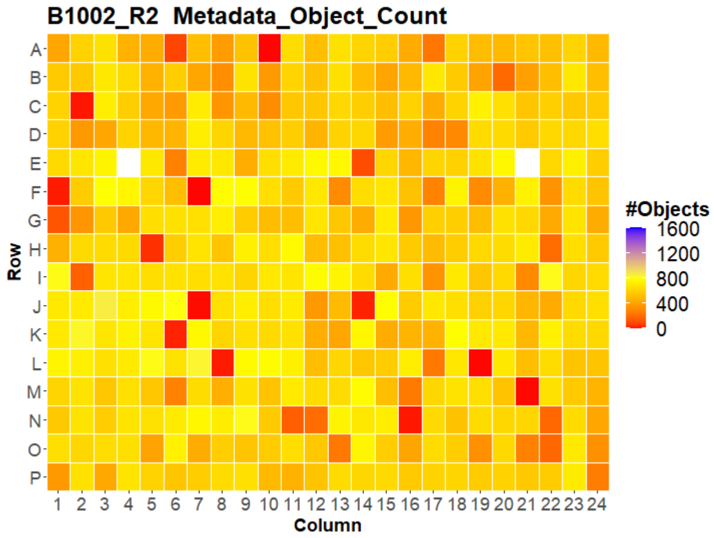
B1002_R2 Metadata_Object_Count
Row
A B C D E F G H I J K L M N O P
1 2 3 4 5 6 7 8 9 10 11 12 13 14 15 16 17 18 19 20 21 22 23 24
Column
#Objects
1600
1200
800
400
0

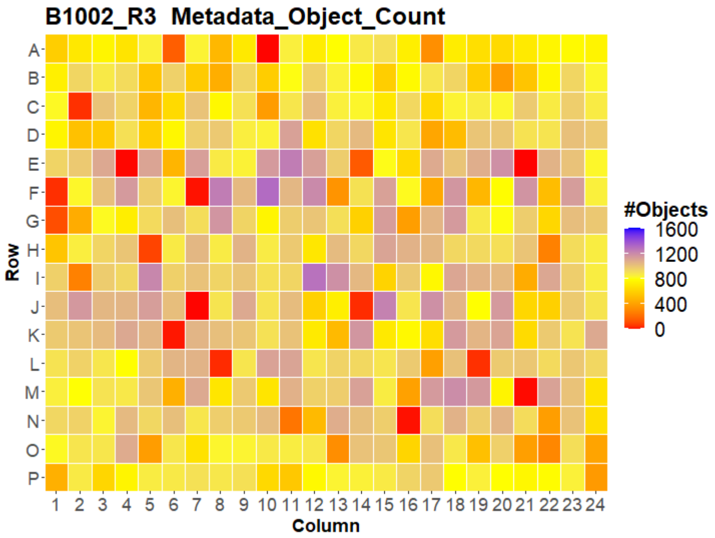
B1002_R3 Metadata_Object_Count
Row
A B C D E F G H I J K L M N O P
1 2 3 4 5 6 7 8 9 10 11 12 13 14 15 16 17 18 19 20 21 22 23 24
Column
#Objects
1600
1200
800
400
0

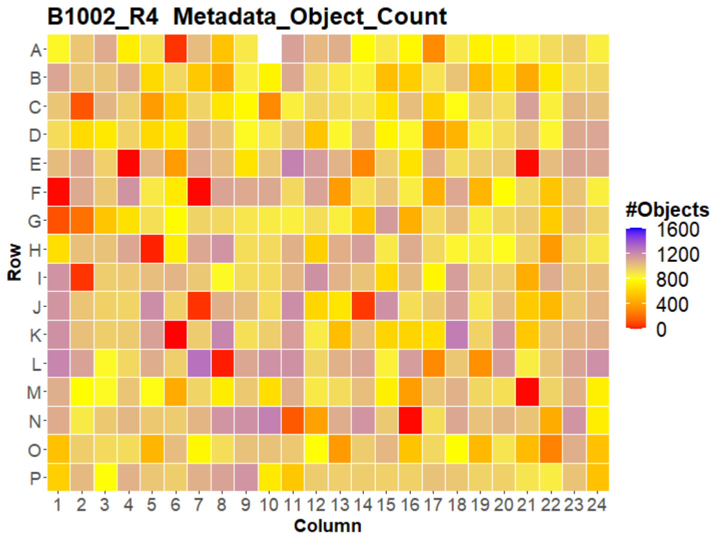
B1002_R4 Metadata_Object_Count
Row
A B C D E F G H I J K L M N O P
1 2 3 4 5 6 7 8 9 10 11 12 13 14 15 16 17 18 19 20 21 22 23 24
Column
#Objects
1600
1200
800
400
0

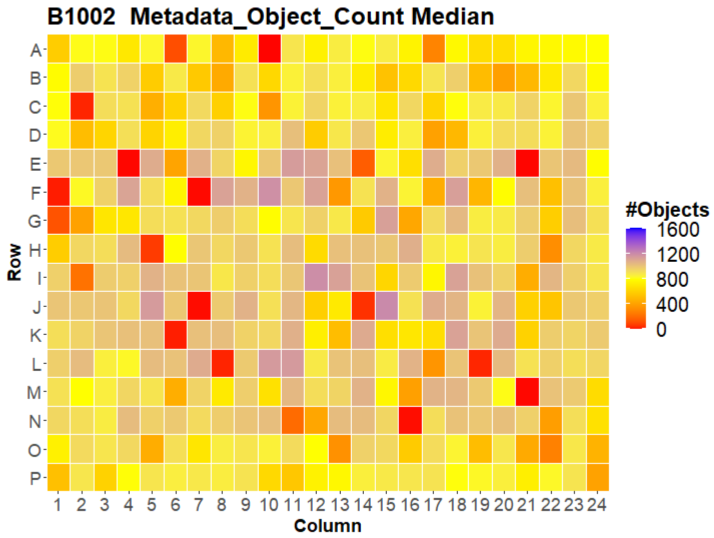
B1002 Metadata_Object_Count Median
Row
A B C D E F G H I J K L M N O P
1 2 3 4 5 6 7 8 9 10 11 12 13 14 15 16 17 18 19 20 21 22 23 24
Column
#Objects
1600
1200
800
400
0

### HepG2 FMP

24

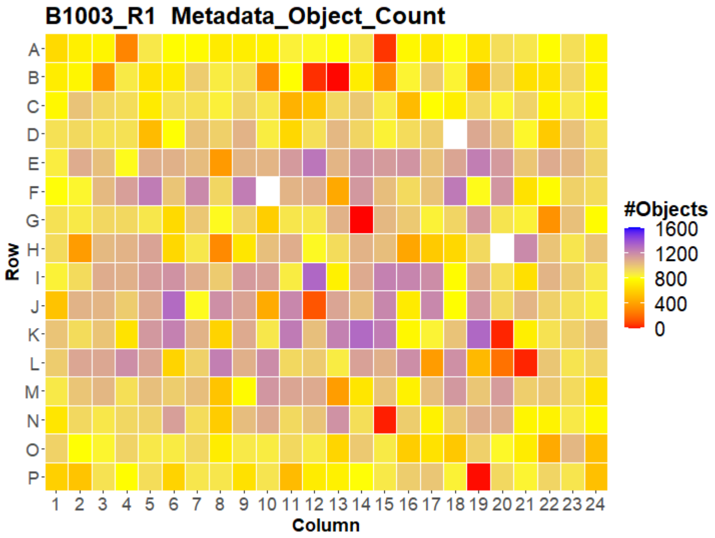
B1003_R1 Metadata_Object_Count
Row
A B C D E F G H I J K L M N O P
Column
1 2 3 4 5 6 7 8 9 10 11 12 13 14 15 16 17 18 19 20 21 22 23 24
#Objects
1600
1200
800
400
0

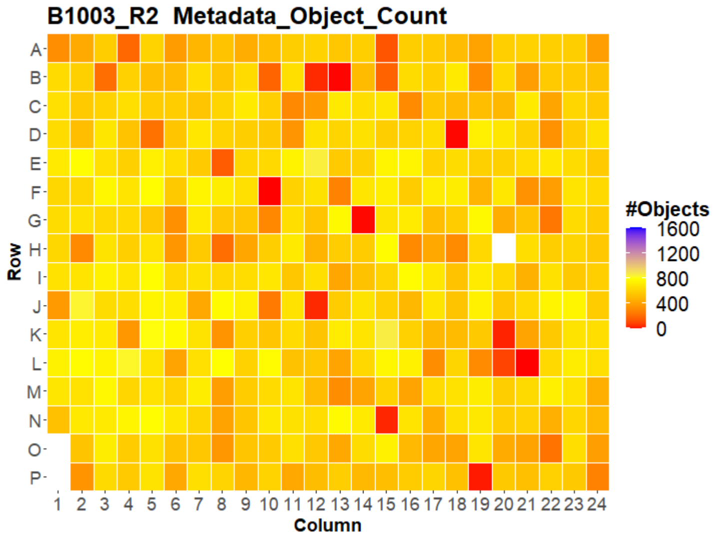
B1003_R2 Metadata_Object_Count
Row
A B C D E F G H I J K L M N O P
Column
1 2 3 4 5 6 7 8 9 10 11 12 13 14 15 16 17 18 19 20 21 22 23 24
#Objects
1600
1200
800
400
0

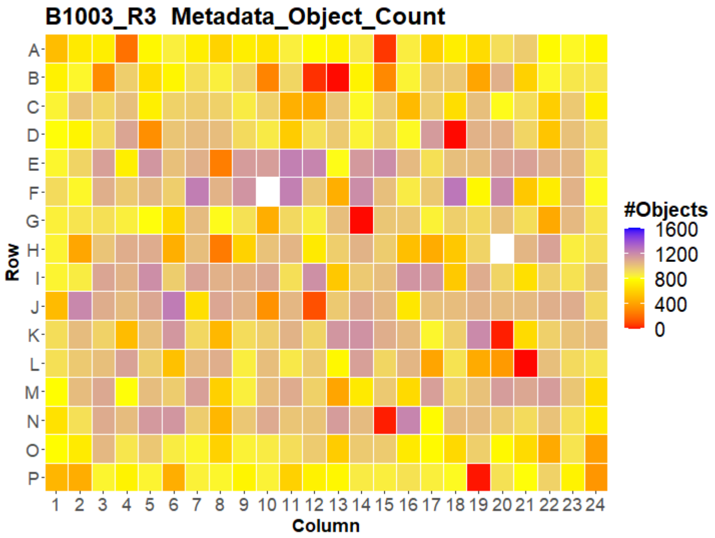
B1003_R3 Metadata_Object_Count
Row
A B C D E F G H I J K L M N O P
Column
1 2 3 4 5 6 7 8 9 10 11 12 13 14 15 16 17 18 19 20 21 22 23 24
#Objects
1600
1200
800
400
0

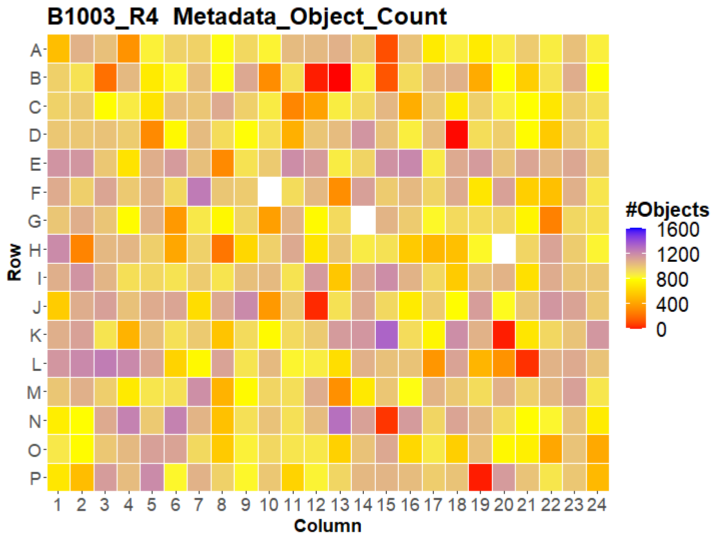
B1003_R4 Metadata_Object_Count
Row
A B C D E F G H I J K L M N O P
Column
1 2 3 4 5 6 7 8 9 10 11 12 13 14 15 16 17 18 19 20 21 22 23 24
#Objects
1600
1200
800
400
0

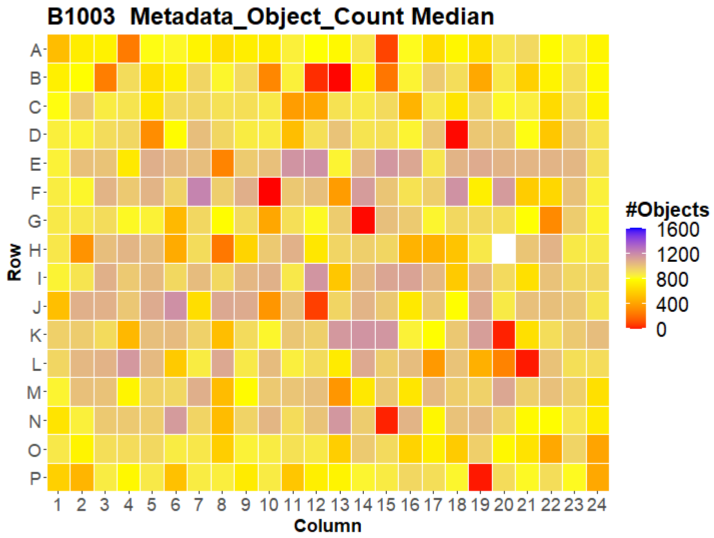
B1003 Metadata_Object_Count Median
Row
A B C D E F G H I J K L M N O P
Column
1 2 3 4 5 6 7 8 9 10 11 12 13 14 15 16 17 18 19 20 21 22 23 24
#Objects
1600
1200
800
400
0

### HepG2 FMP



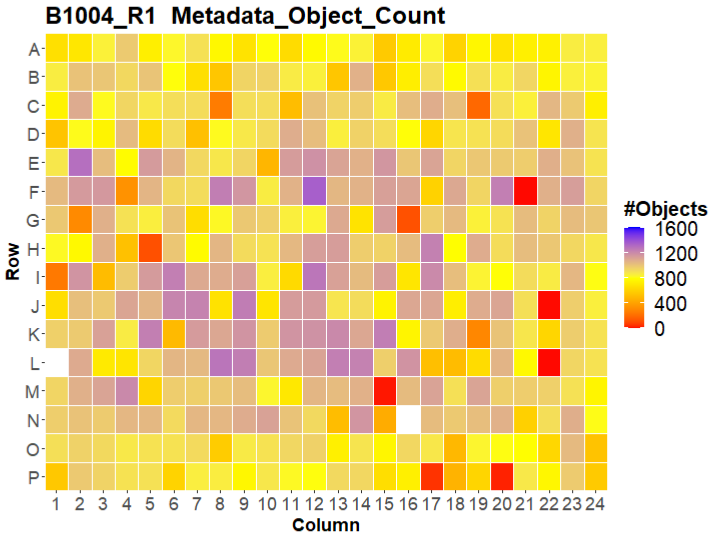
B1004_R1 Metadata_Object_Count
A B C D E F G H I J K L M N O P
Row
1 2 3 4 5 6 7 8 9 10 11 12 13 14 15 16 17 18 19 20 21 22 23 24
Column
#Objects
1600
1200
800
400
0

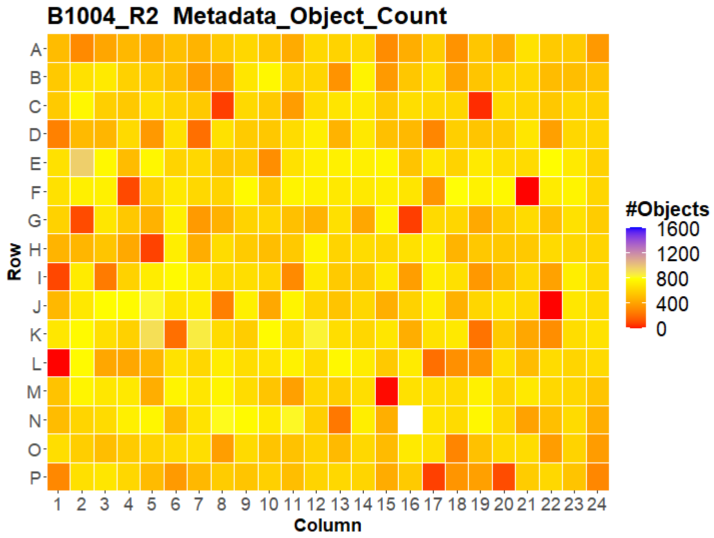
B1004_R2 Metadata_Object_Count
A B C D E F G H I J K L M N O P
Row
1 2 3 4 5 6 7 8 9 10 11 12 13 14 15 16 17 18 19 20 21 22 23 24
Column
#Objects
1600
1200
800
400
0

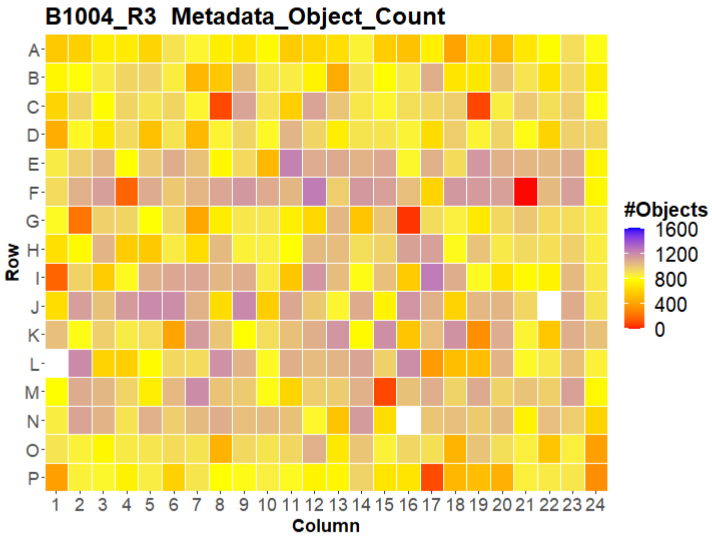
B1004_R3 Metadata_Object_Count
A B C D E F G H I J K L M N O P
Row
1 2 3 4 5 6 7 8 9 10 11 12 13 14 15 16 17 18 19 20 21 22 23 24
Column
#Objects
1600
1200
800
400
0

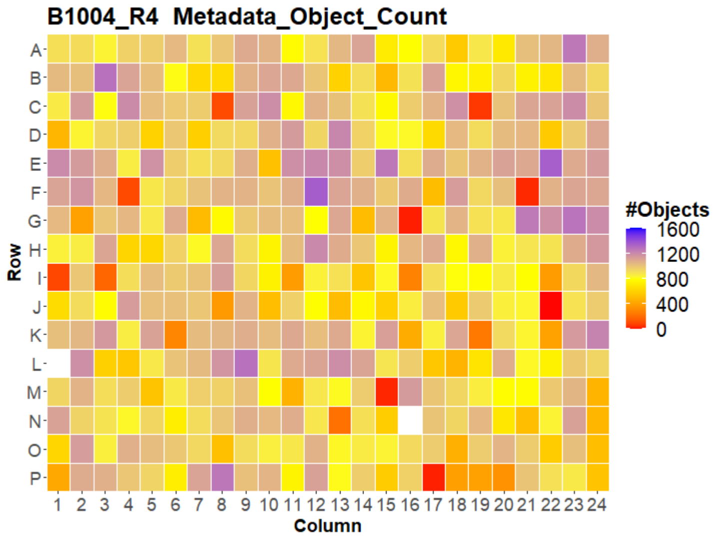
B1004_R4 Metadata_Object_Count
A B C D E F G H I J K L M N O P
Row
1 2 3 4 5 6 7 8 9 10 11 12 13 14 15 16 17 18 19 20 21 22 23 24
Column
#Objects
1600
1200
800
400
0

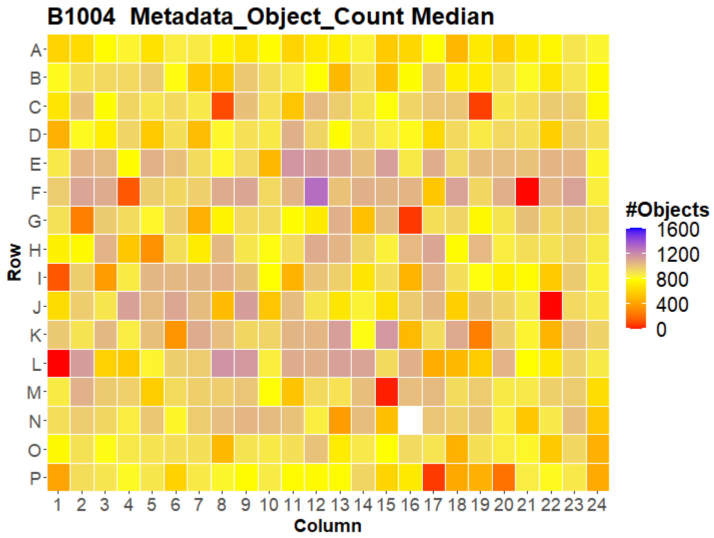
B1004 Metadata_Object_Count Median
A B C D E F G H I J K L M N O P
Row
1 2 3 4 5 6 7 8 9 10 11 12 13 14 15 16 17 18 19 20 21 22 23 24
Column
#Objects
1600
1200
800
400
0

### HepG2 FMP



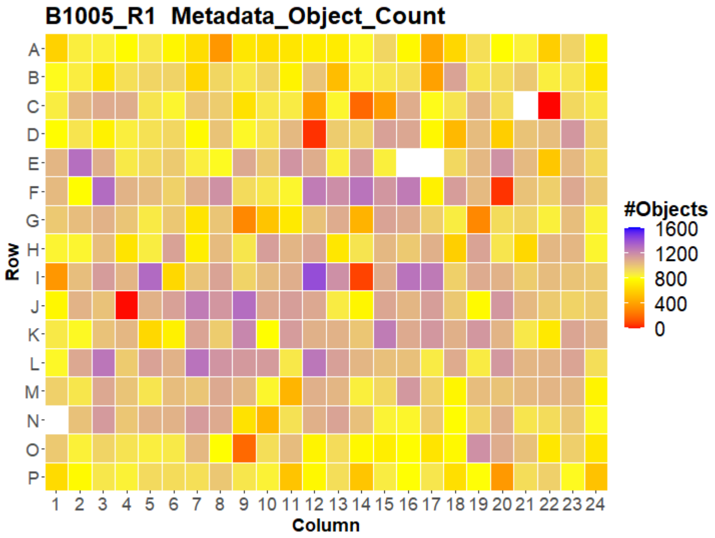
B1005_R1 Metadata_Object_Count
Row
Column
#Objects
1600
1200
800
400
0

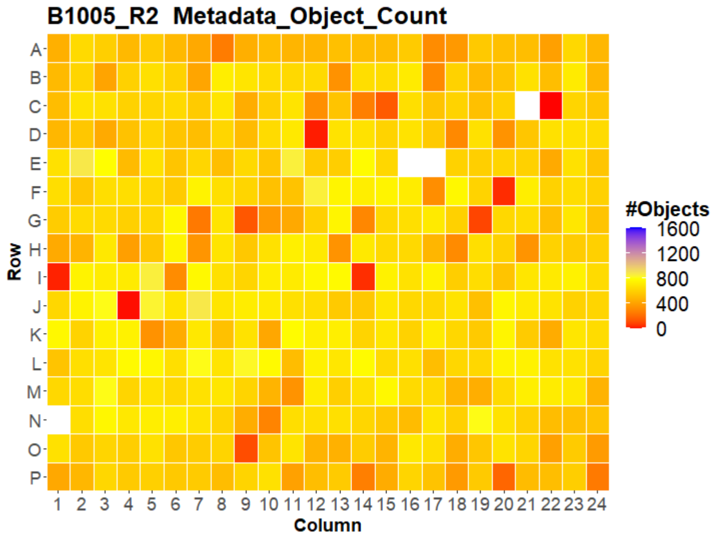
B1005_R2 Metadata_Object_Count
Row
Column
#Objects
1600
1200
800
400
0

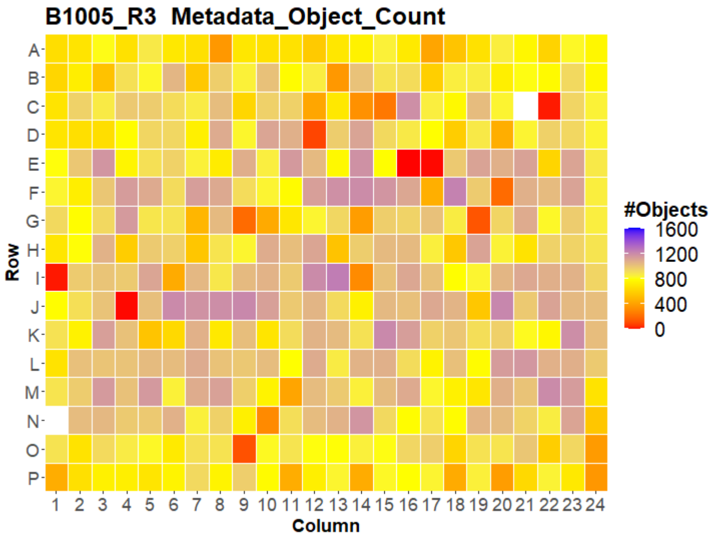
B1005_R3 Metadata_Object_Count
Row
Column
#Objects
1600
1200
800
400
0

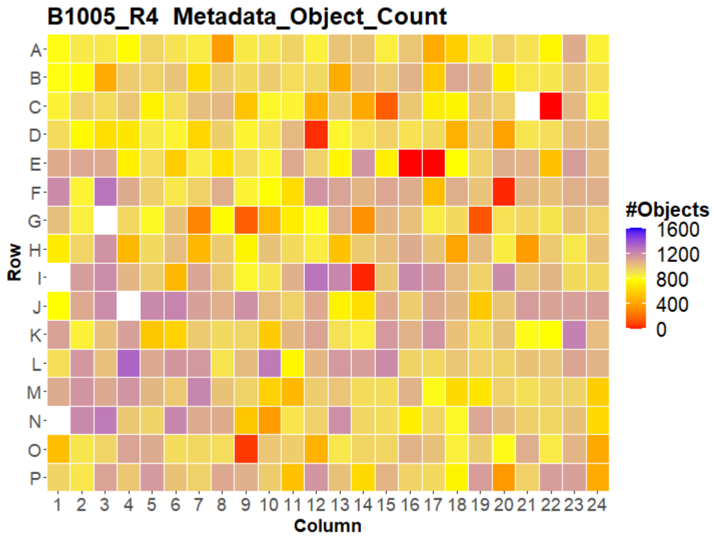
B1005_R4 Metadata_Object_Count
Row
Column
#Objects
1600
1200
800
400
0

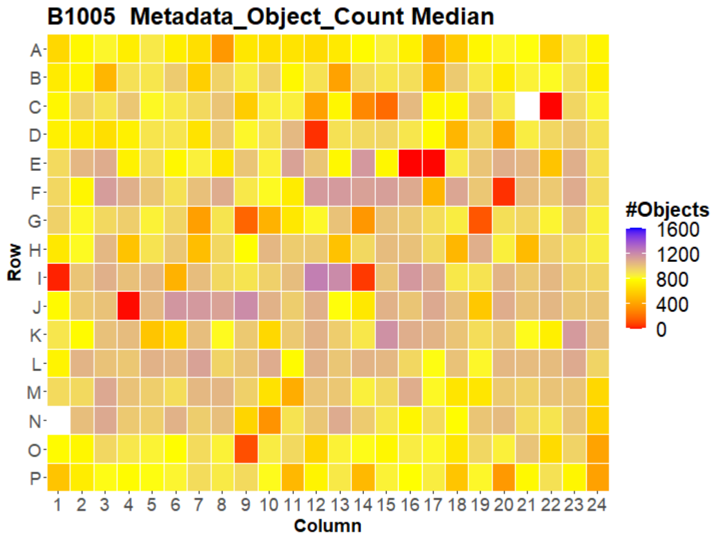
B1005 Metadata_Object_Count Median
Row
Column
#Objects
1600
1200
800
400
0

### HepG2 FMP

27

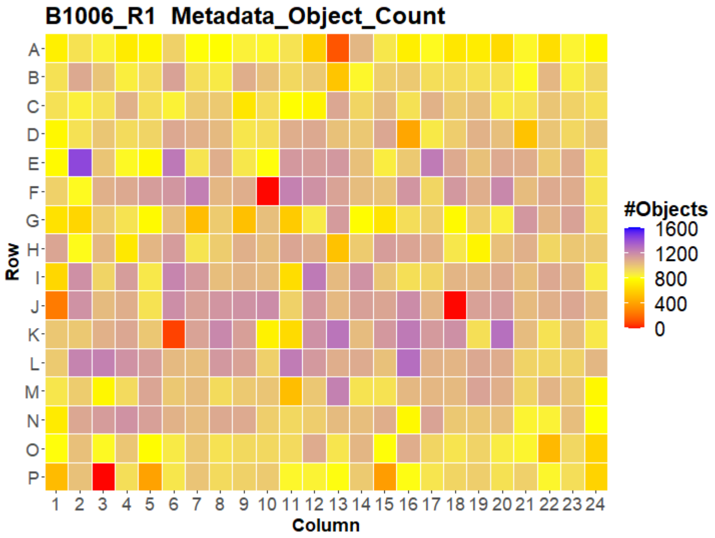


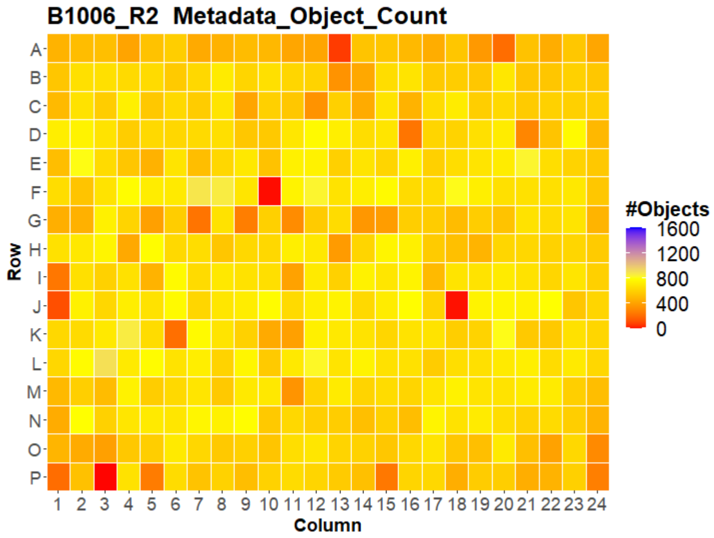


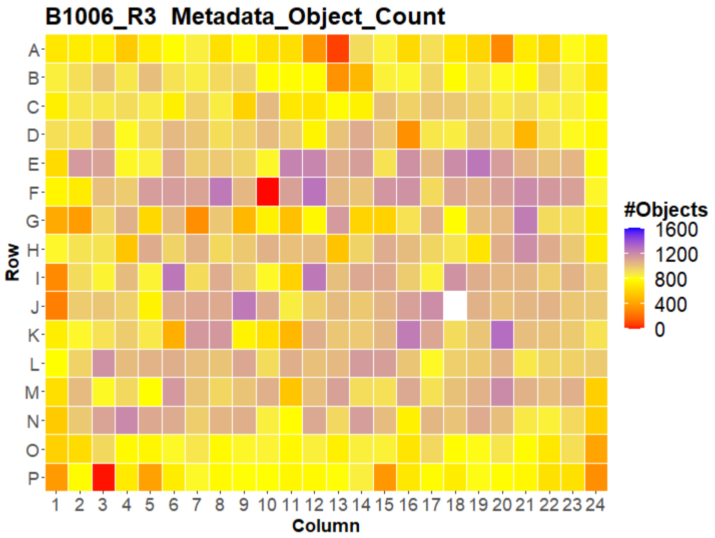


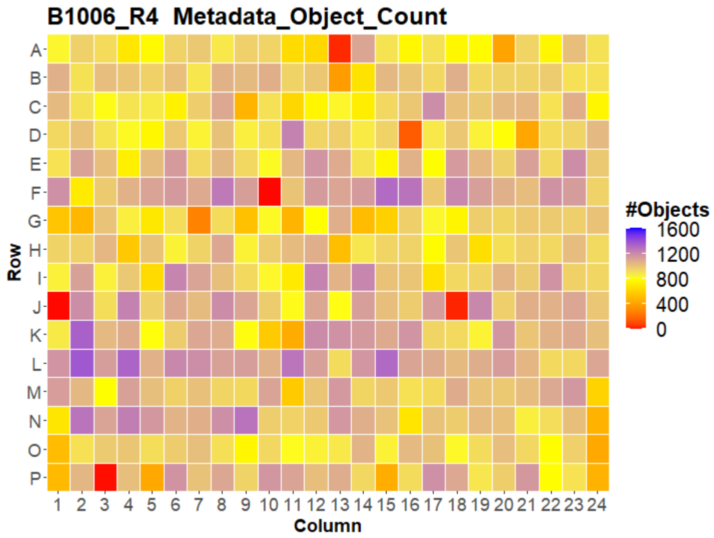


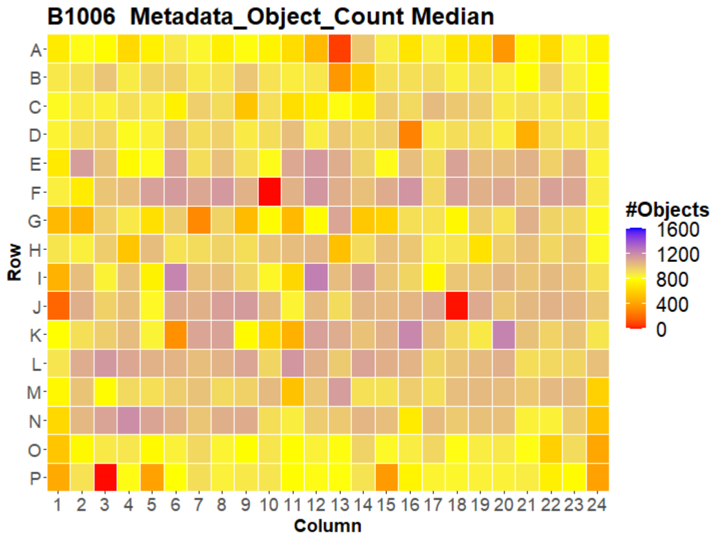



Open for Innovation KNIME

### HepG2 FMP



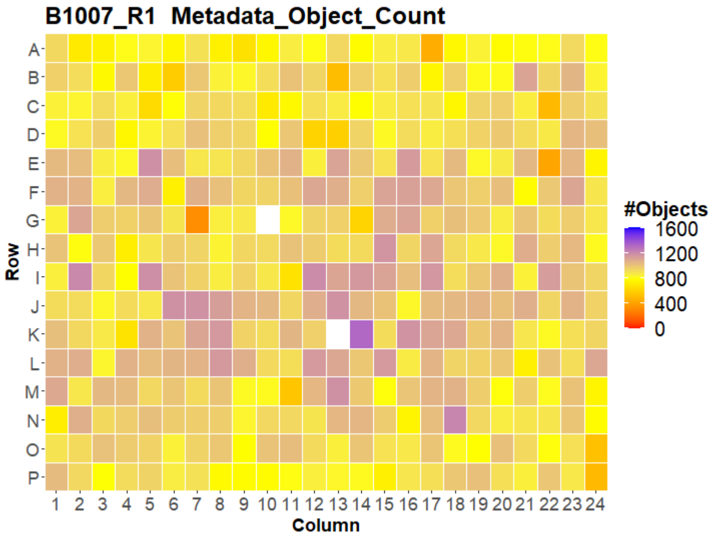
B1007_R1 Metadata_Object_Count
Row
A B C D E F G H I J K L M N O P
1 2 3 4 5 6 7 8 9 10 11 12 13 14 15 16 17 18 19 20 21 22 23 24
Column
#Objects
1600
1200
800
400
0

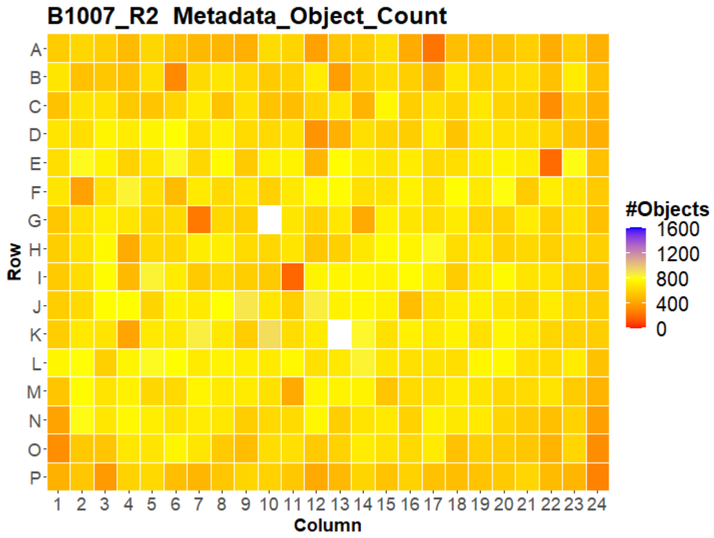
B1007_R2 Metadata_Object_Count
Row
A B C D E F G H I J K L M N O P
1 2 3 4 5 6 7 8 9 10 11 12 13 14 15 16 17 18 19 20 21 22 23 24
Column
#Objects
1600
1200
800
400
0

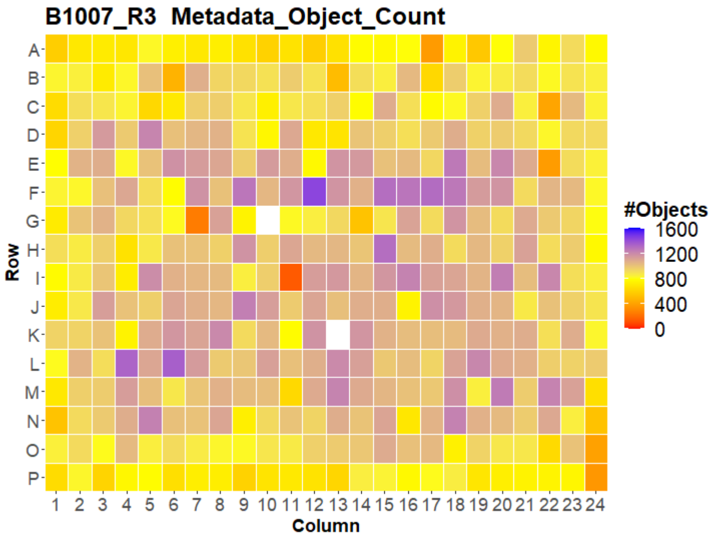
B1007_R3 Metadata_Object_Count
Row
A B C D E F G H I J K L M N O P
1 2 3 4 5 6 7 8 9 10 11 12 13 14 15 16 17 18 19 20 21 22 23 24
Column
#Objects
1600
1200
800
400
0

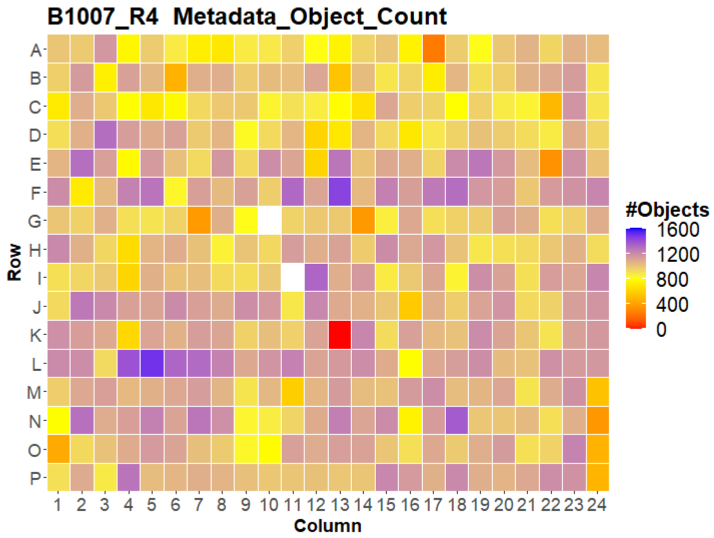
B1007_R4 Metadata_Object_Count
Row
A B C D E F G H I J K L M N O P
1 2 3 4 5 6 7 8 9 10 11 12 13 14 15 16 17 18 19 20 21 22 23 24
Column
#Objects
1600
1200
800
400
0

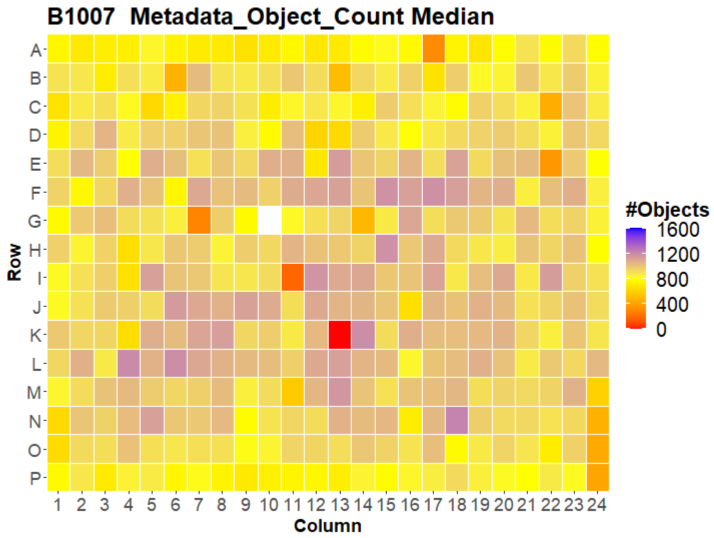
B1007 Metadata_Object_Count Median
Row
A B C D E F G H I J K L M N O P
1 2 3 4 5 6 7 8 9 10 11 12 13 14 15 16 17 18 19 20 21 22 23 24
Column
#Objects
1600
1200
800
400
0

### HepG2 FMP

29

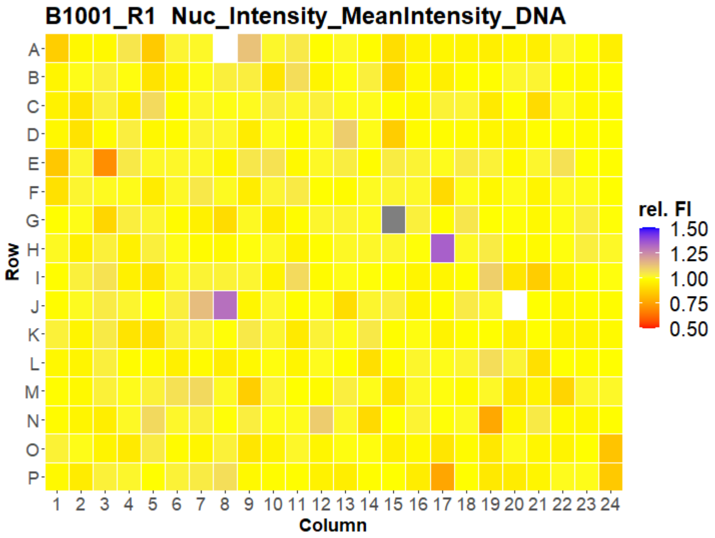
B1001_R1 Nuc_Intensity_MeanIntensity_DNA
Row
Column
rel. FI
1.50
1.25
1.00
0.75
0.50

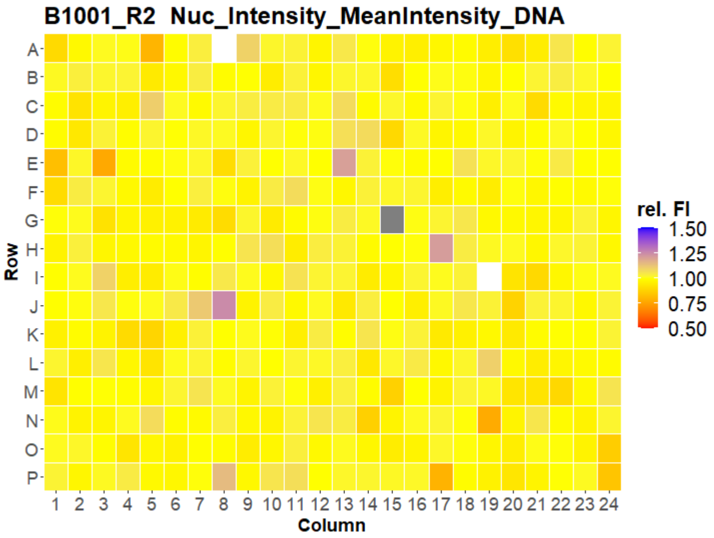
B1001_R2 Nuc_Intensity_MeanIntensity_DNA
Row
Column
rel. FI
1.50
1.25
1.00
0.75
0.50

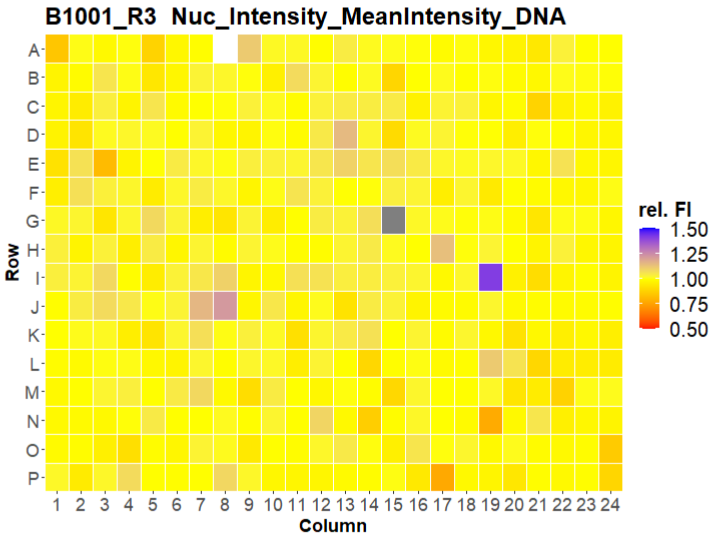
B1001_R3 Nuc_Intensity_MeanIntensity_DNA
Row
Column
rel. FI
1.50
1.25
1.00
0.75
0.50

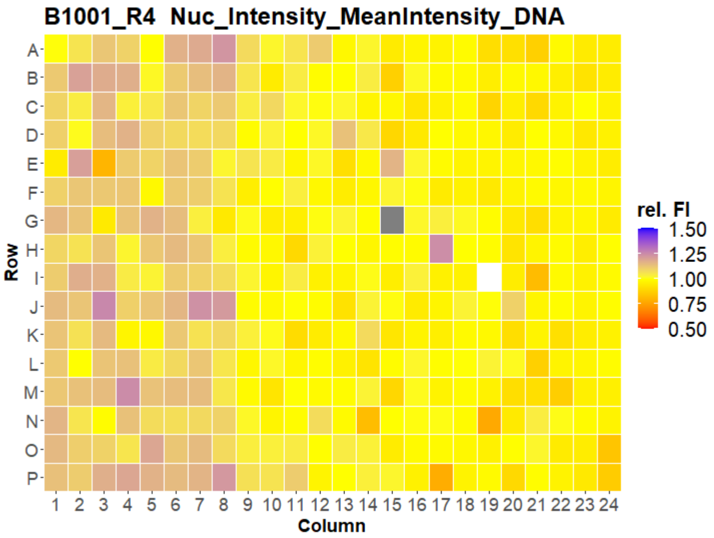
B1001_R4 Nuc_Intensity_MeanIntensity_DNA
Row
Column
rel. FI
1.50
1.25
1.00
0.75
0.50

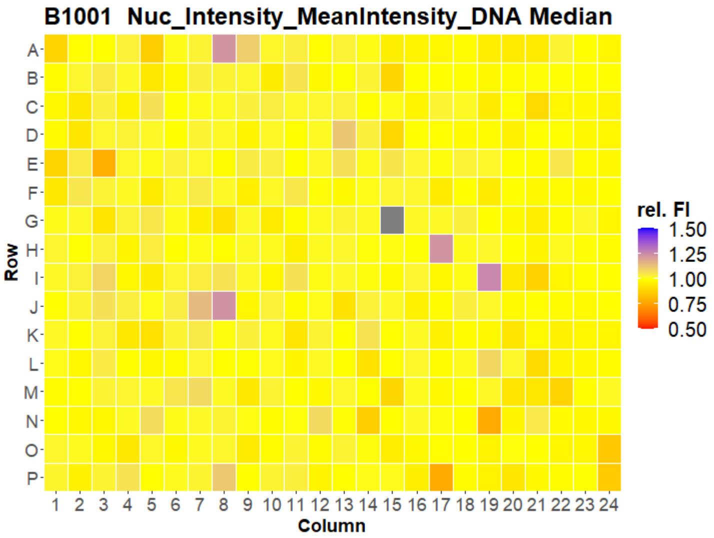
B1001 Nuc_Intensity_MeanIntensity_DNA Median
Row
Column
rel. FI
1.50
1.25
1.00
0.75
0.50

### HepG2 FMP



B1002_R1 Nuc_Intensity_MeanIntensity_DNA

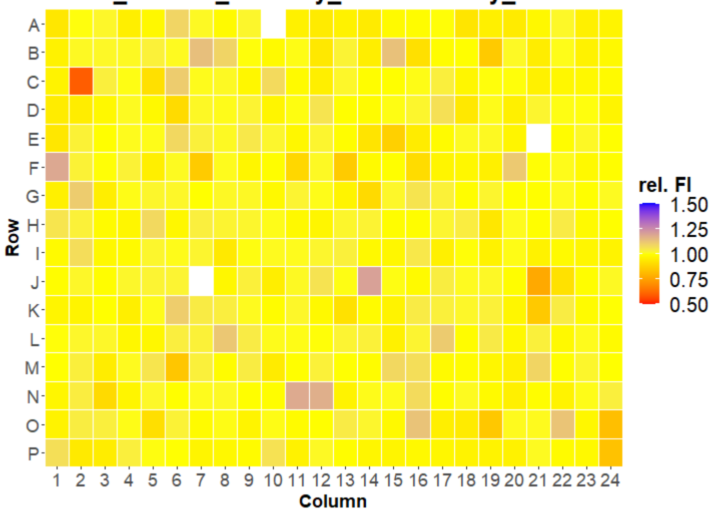


B1002_R2 Nuc_Intensity_MeanIntensity_DNA

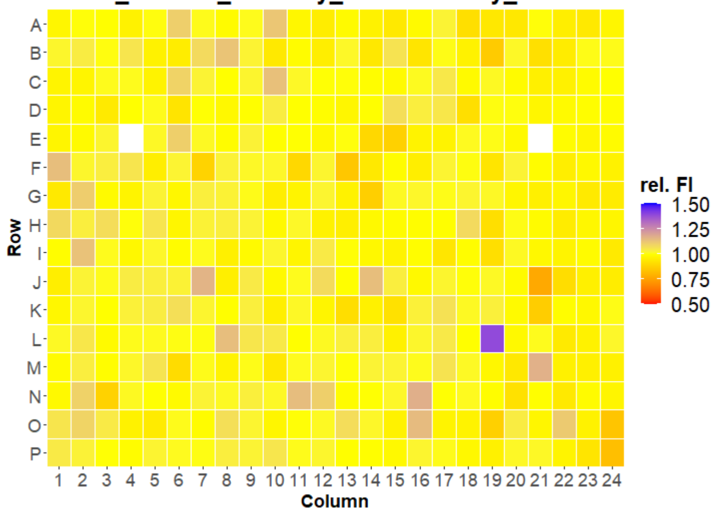


B1002_R3 Nuc_Intensity_MeanIntensity_DNA

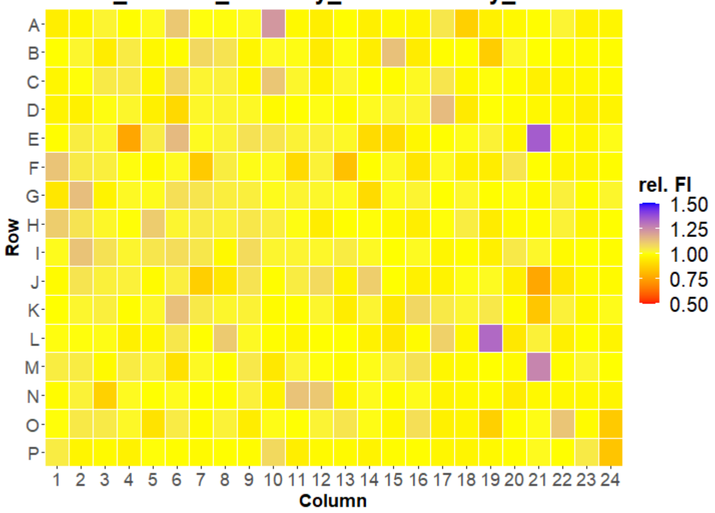


B1002_R4 Nuc_Intensity_MeanIntensity_DNA

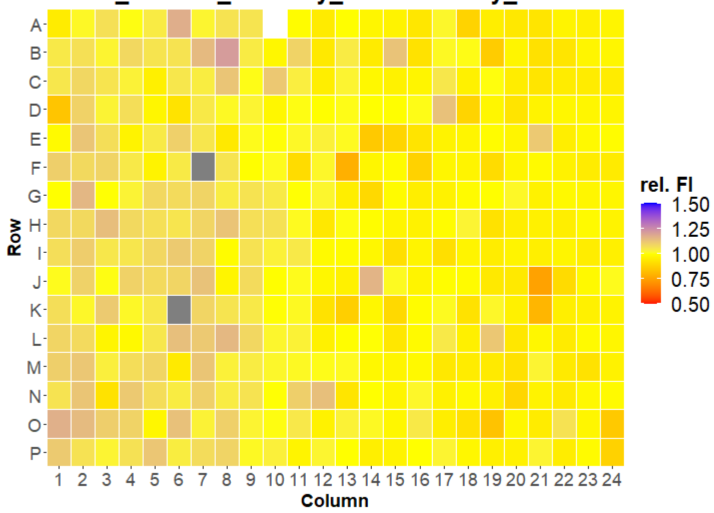


B1002 Nuc_Intensity_MeanIntensity_DNA Median

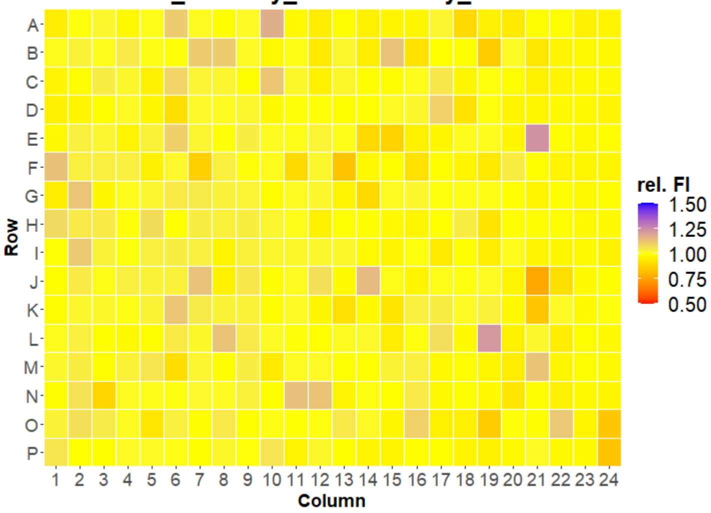

### HepG2 FMP

31

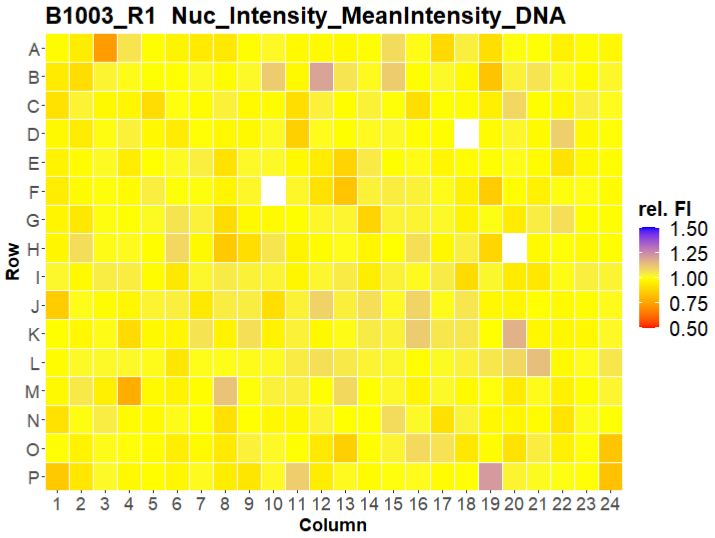

B1003_R1 Nuc_Intensity_MeanIntensity_DNA
Row
Column
rel. FI
1.50
1.25
1.00
0.75
0.50


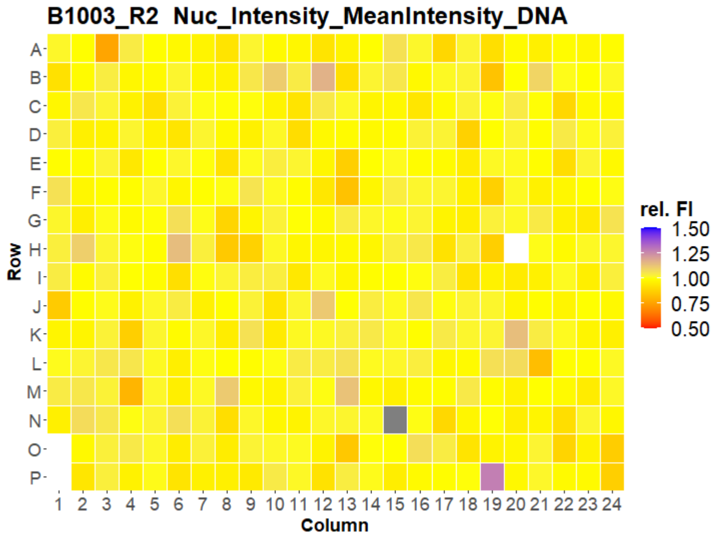

B1003_R2 Nuc_Intensity_MeanIntensity_DNA
Row
Column
rel. FI
1.50
1.25
1.00
0.75
0.50


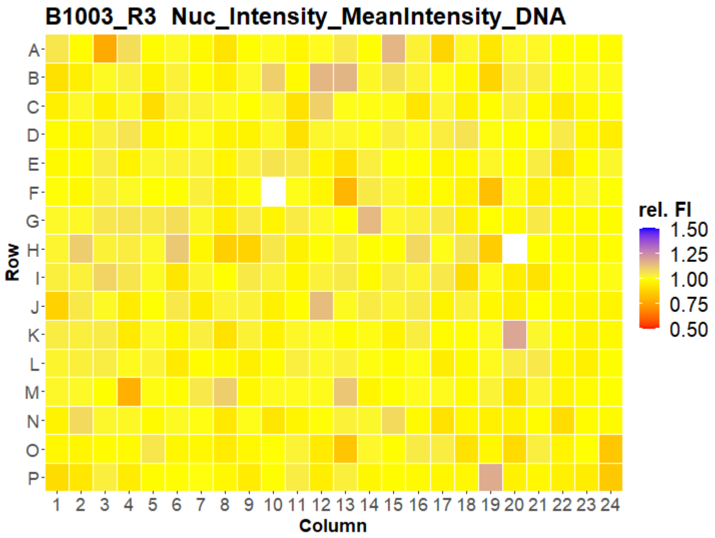

B1003_R3 Nuc_Intensity_MeanIntensity_DNA
Row
Column
rel. FI
1.50
1.25
1.00
0.75
0.50


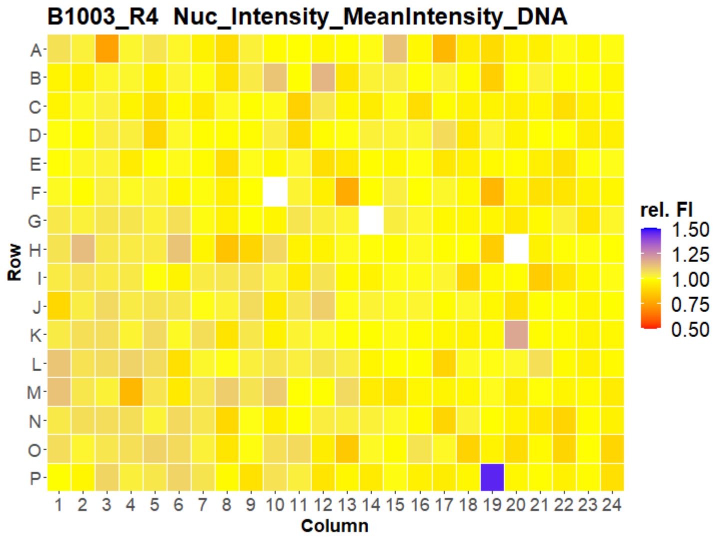

B1003_R4 Nuc_Intensity_MeanIntensity_DNA
Row
Column
rel. FI
1.50
1.25
1.00
0.75
0.50


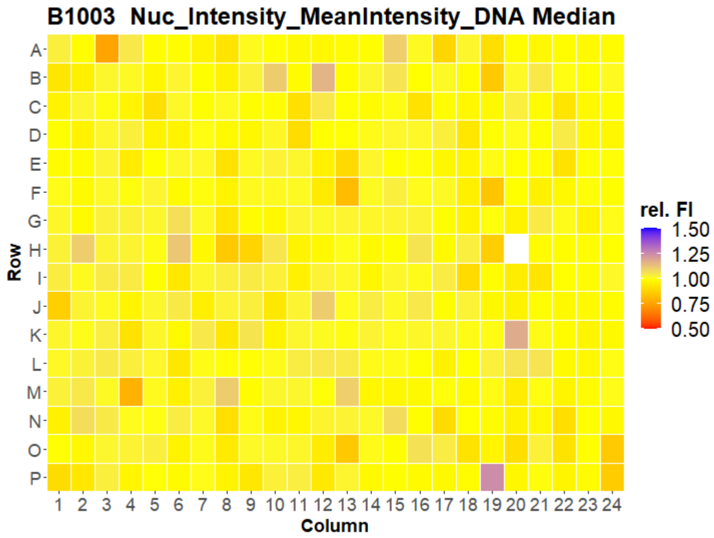

B1003 Nuc_Intensity_MeanIntensity_DNA Median
Row
Column
rel. FI
1.50
1.25
1.00
0.75
0.50

### HepG2 FMP



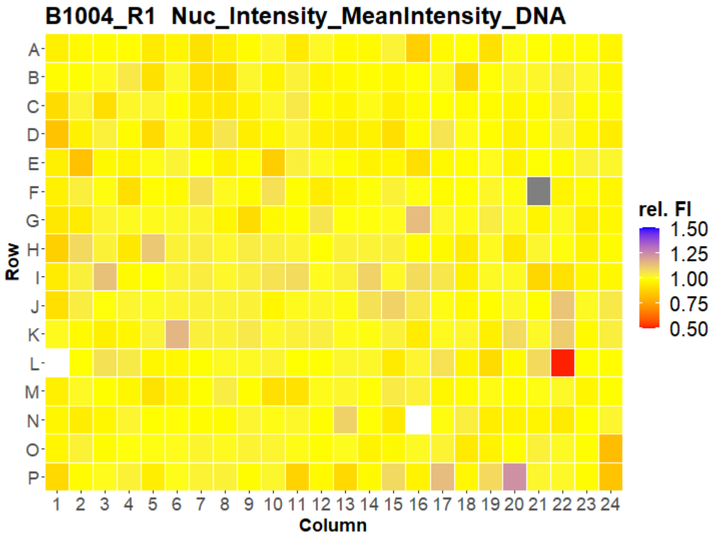
B1004_R1 Nuc_Intensity_MeanIntensity_DNA
Row
Column
rel. FI
1.50
1.25
1.00
0.75
0.50

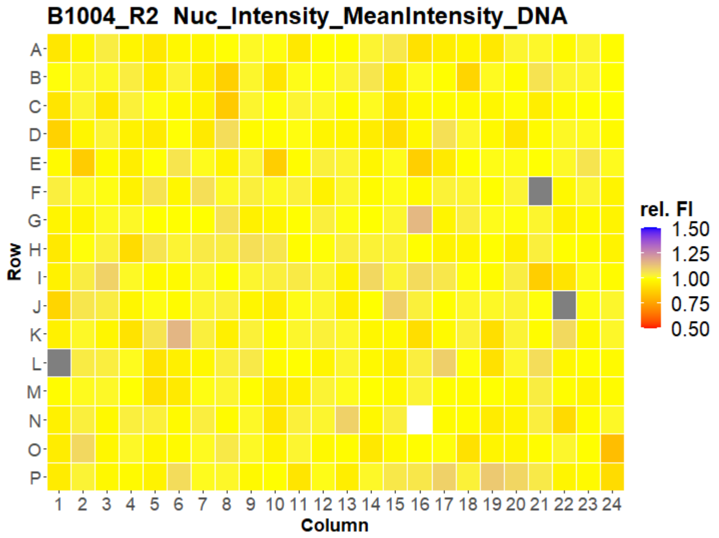
B1004_R2 Nuc_Intensity_MeanIntensity_DNA
Row
Column
rel. FI
1.50
1.25
1.00
0.75
0.50

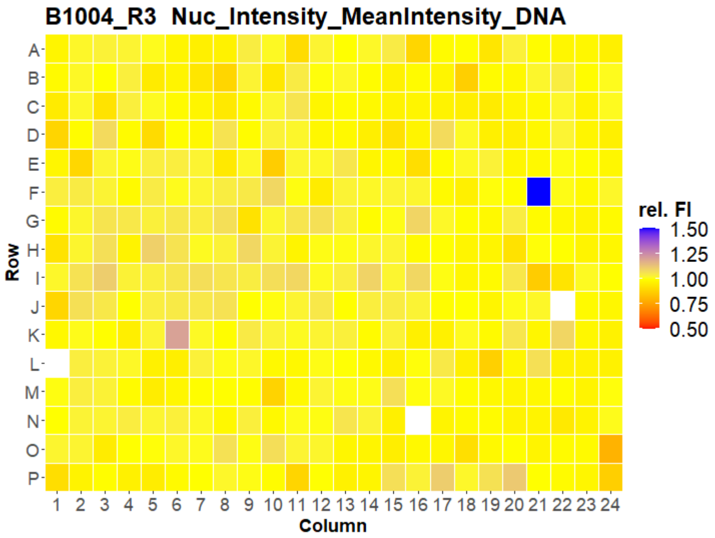
B1004_R3 Nuc_Intensity_MeanIntensity_DNA
Row
Column
rel. FI
1.50
1.25
1.00
0.75
0.50

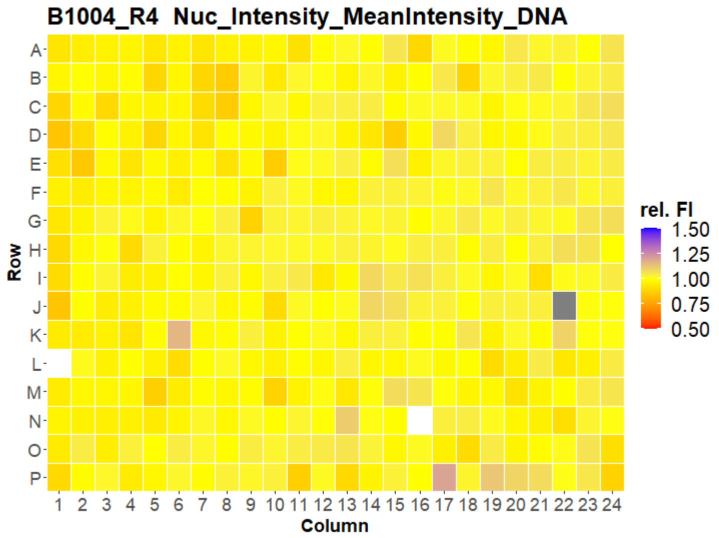
B1004_R4 Nuc_Intensity_MeanIntensity_DNA
Row
Column
rel. FI
1.50
1.25
1.00
0.75
0.50

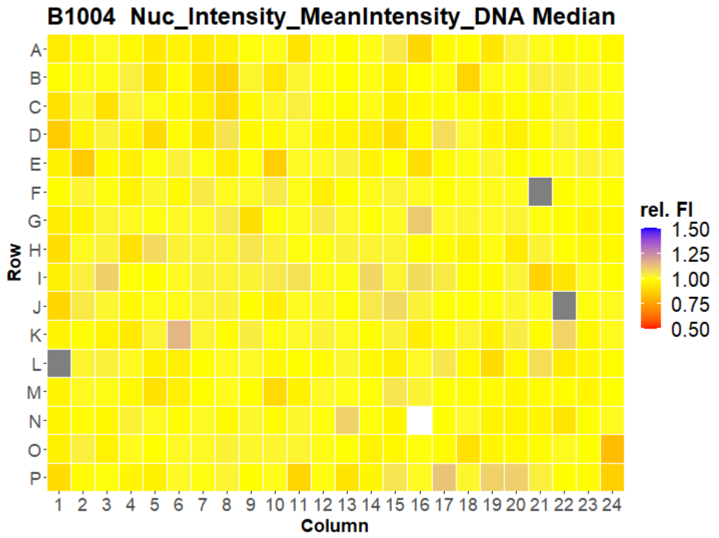
B1004 Nuc_Intensity_MeanIntensity_DNA Median
Row
Column
rel. FI
1.50
1.25
1.00
0.75
0.50

### HepG2 FMP

33

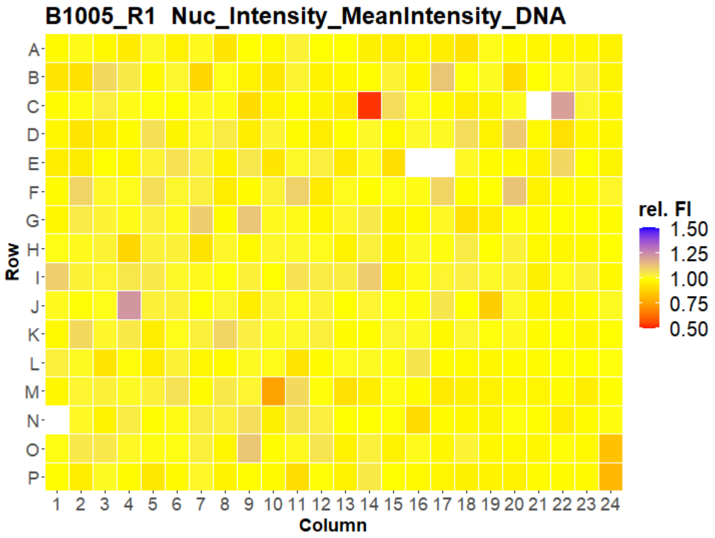


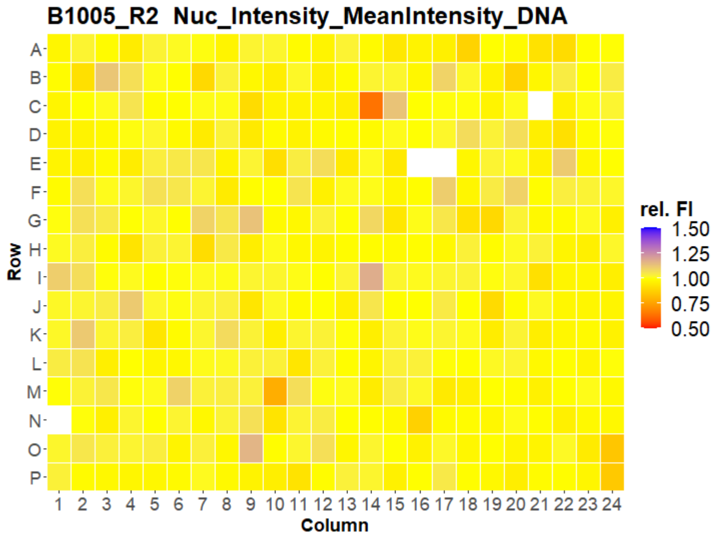


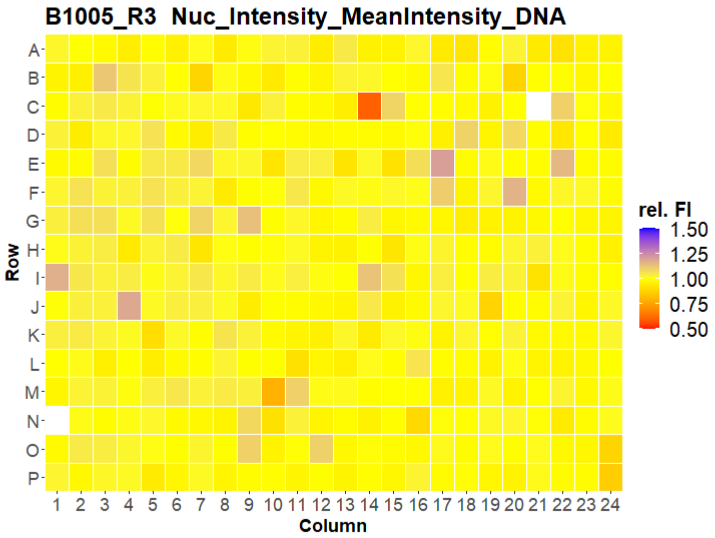


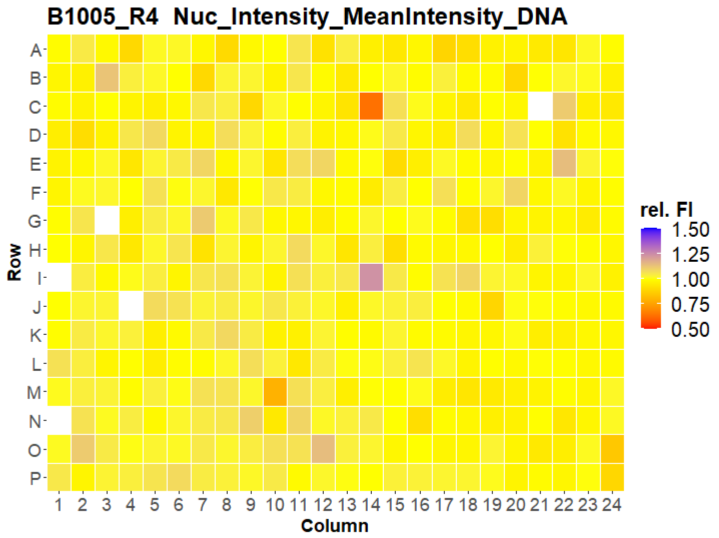


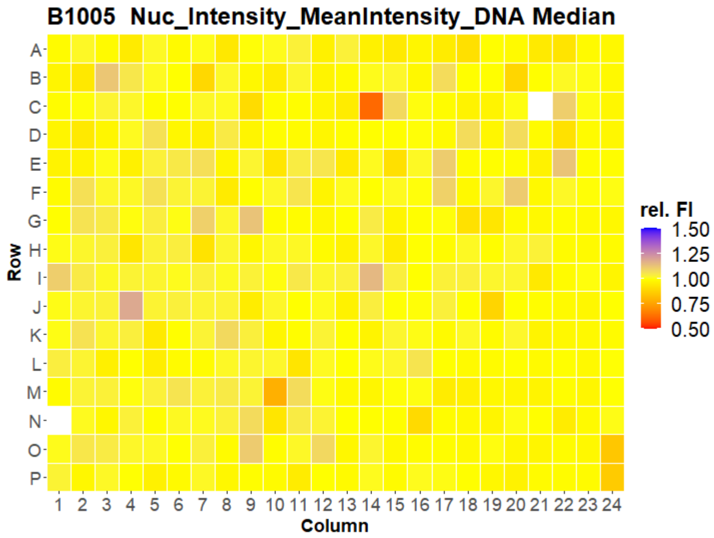



Open for Innovation KNIME

### HepG2 FMP

34

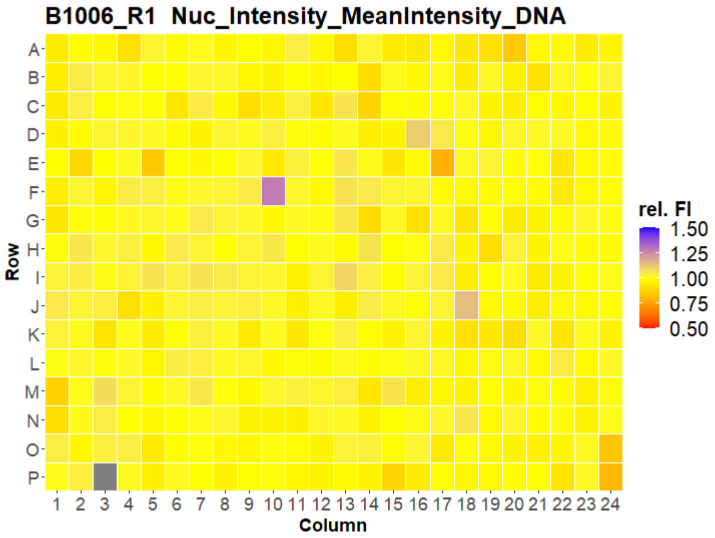


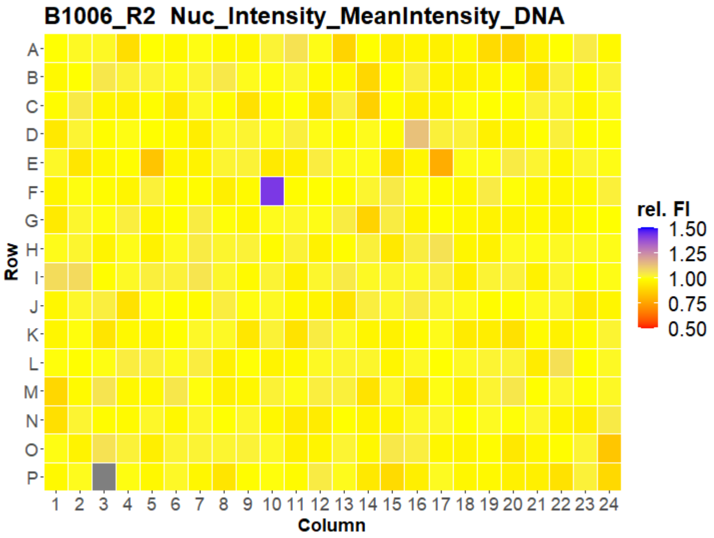


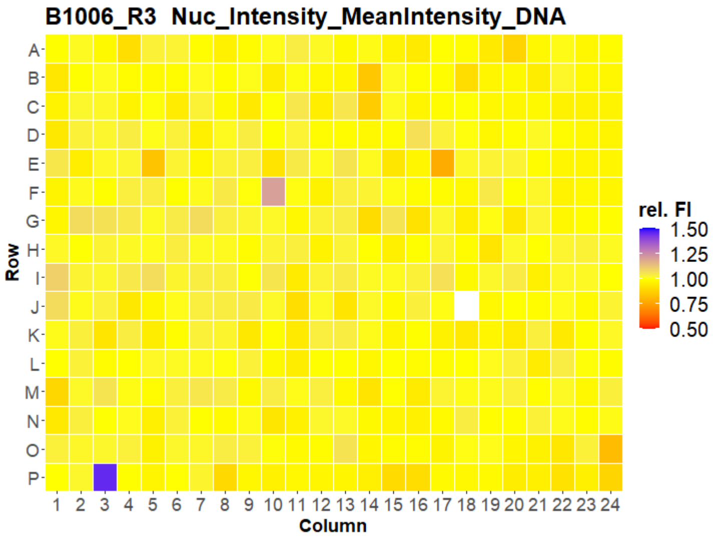


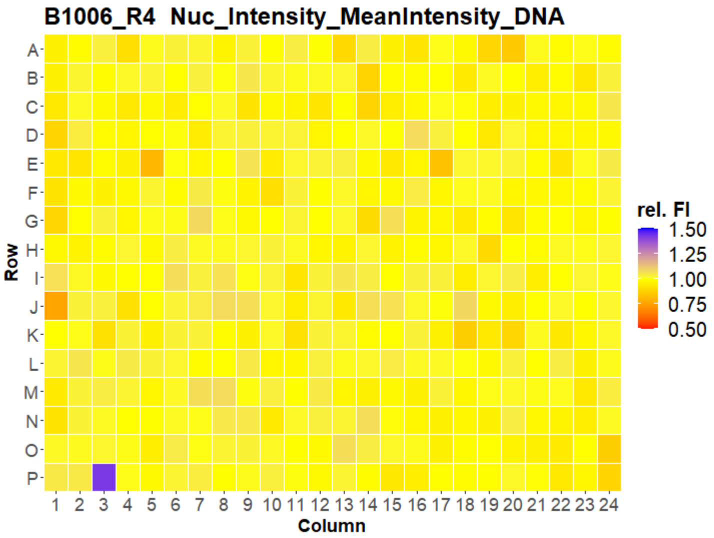


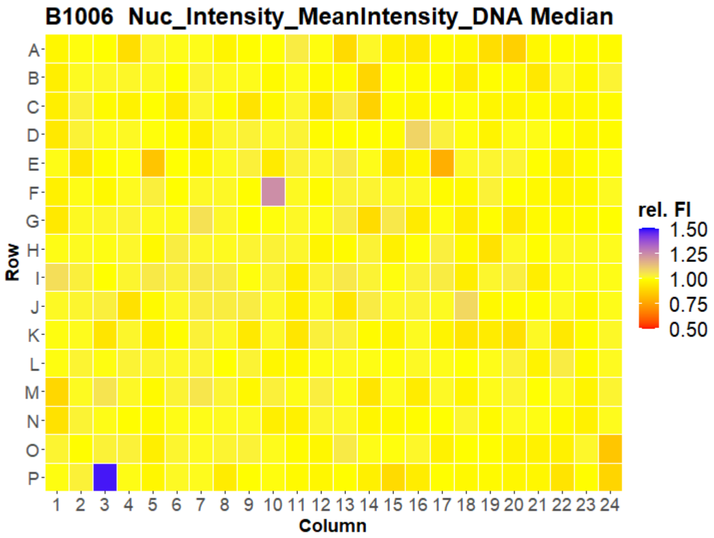



Open for Innovation KNIME

### HepG2 FMP

35

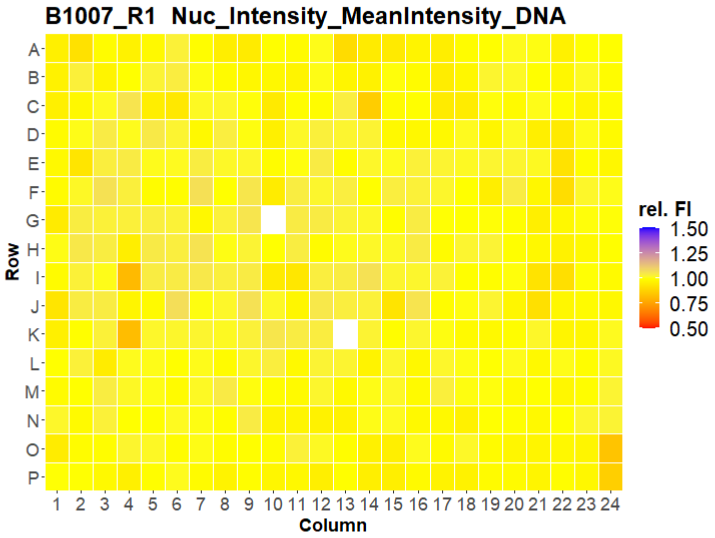
B1007_R1 Nuc_Intensity_MeanIntensity_DNA
Row
Column
rel. FI
1.50
1.25
1.00
0.75
0.50

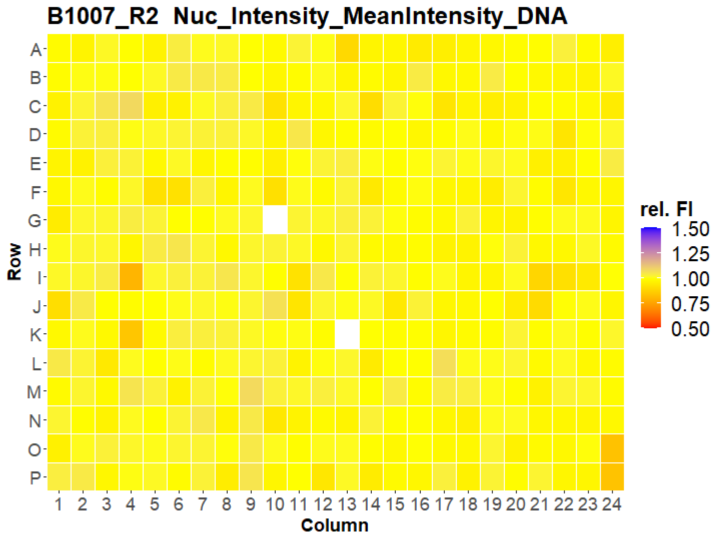
B1007_R2 Nuc_Intensity_MeanIntensity_DNA
Row
Column
rel. FI
1.50
1.25
1.00
0.75
0.50

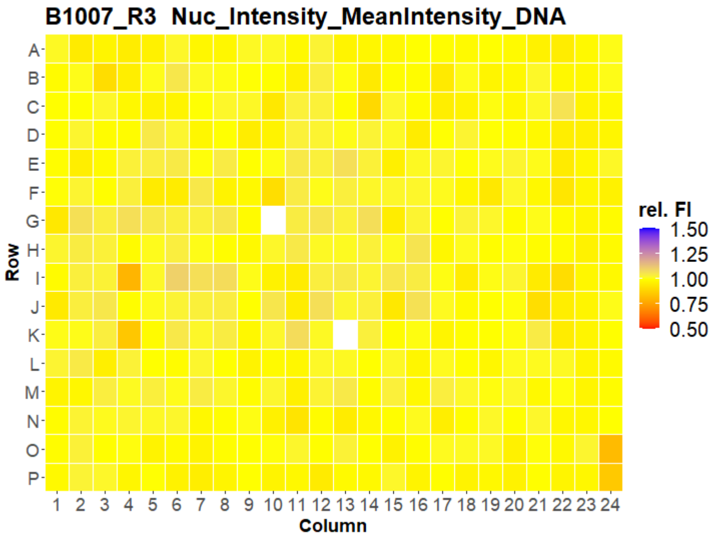
B1007_R3 Nuc_Intensity_MeanIntensity_DNA
Row
Column
rel. FI
1.50
1.25
1.00
0.75
0.50

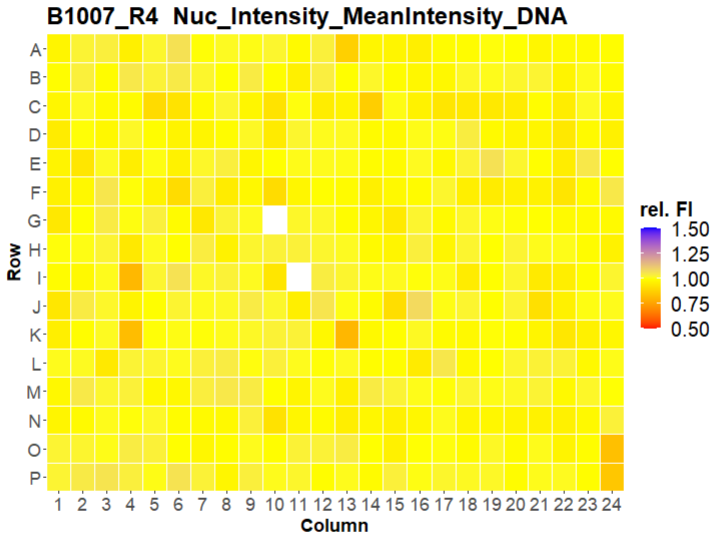
B1007_R4 Nuc_Intensity_MeanIntensity_DNA
Row
Column
rel. FI
1.50
1.25
1.00
0.75
0.50

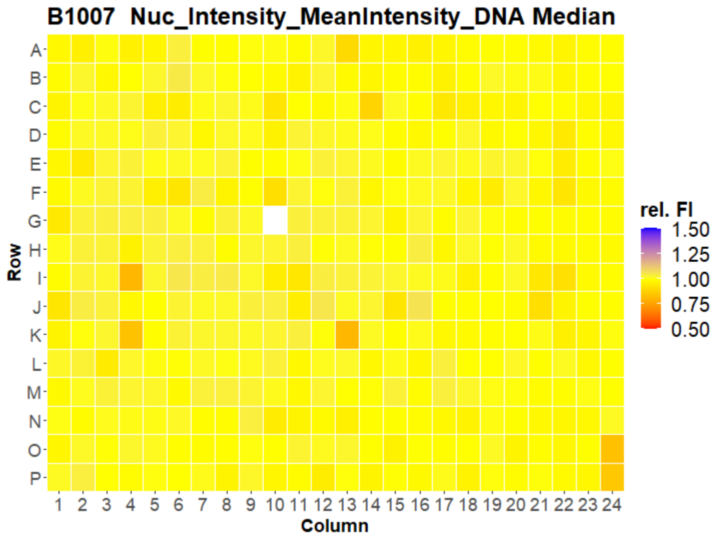
B1007 Nuc_Intensity_MeanIntensity_DNA Median
Row
Column
rel. FI
1.50
1.25
1.00
0.75
0.50

### HepG2 FMP

36

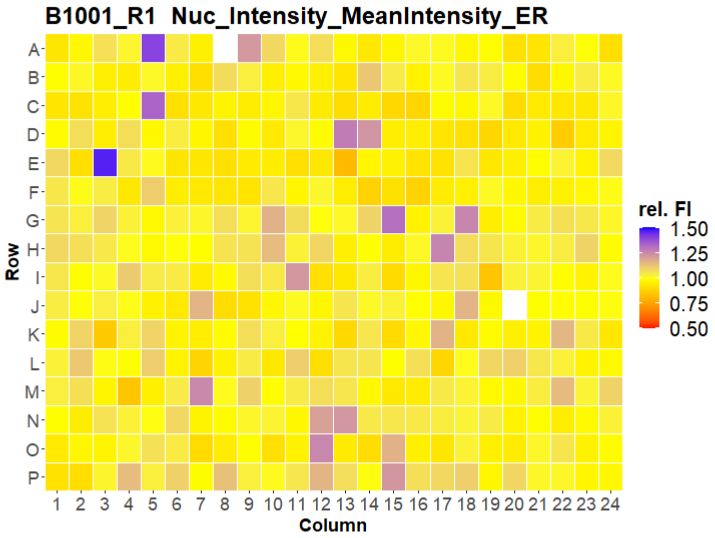
B1001_R1 Nuc_Intensity_MeanIntensity_ER
Row
A B C D E F G H I J K L M N O P
Column
1 2 3 4 5 6 7 8 9 10 11 12 13 14 15 16 17 18 19 20 21 22 23 24
rel. FI
1.50
1.25
1.00
0.75
0.50

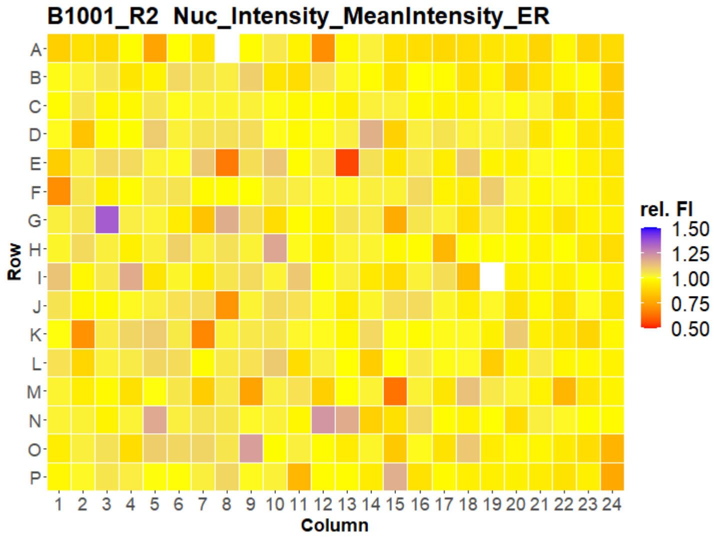
B1001_R2 Nuc_Intensity_MeanIntensity_ER
Row
A B C D E F G H I J K L M N O P
Column
1 2 3 4 5 6 7 8 9 10 11 12 13 14 15 16 17 18 19 20 21 22 23 24
rel. FI
1.50
1.25
1.00
0.75
0.50

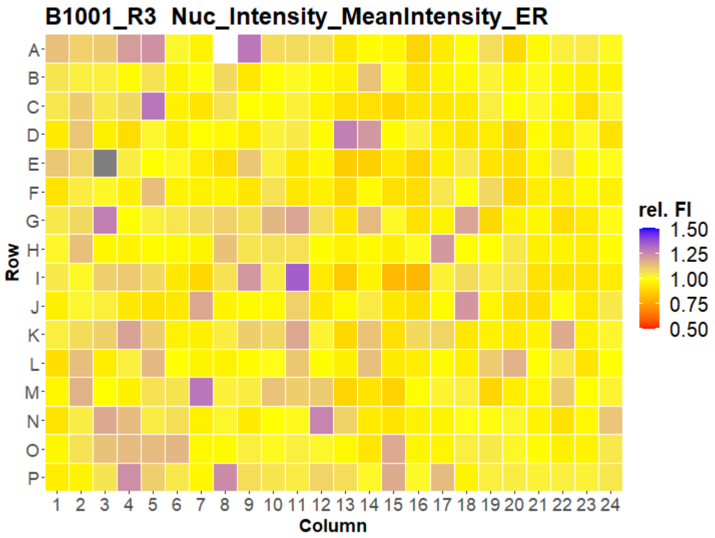
B1001_R3 Nuc_Intensity_MeanIntensity_ER
Row
A B C D E F G H I J K L M N O P
Column
1 2 3 4 5 6 7 8 9 10 11 12 13 14 15 16 17 18 19 20 21 22 23 24
rel. FI
1.50
1.25
1.00
0.75
0.50

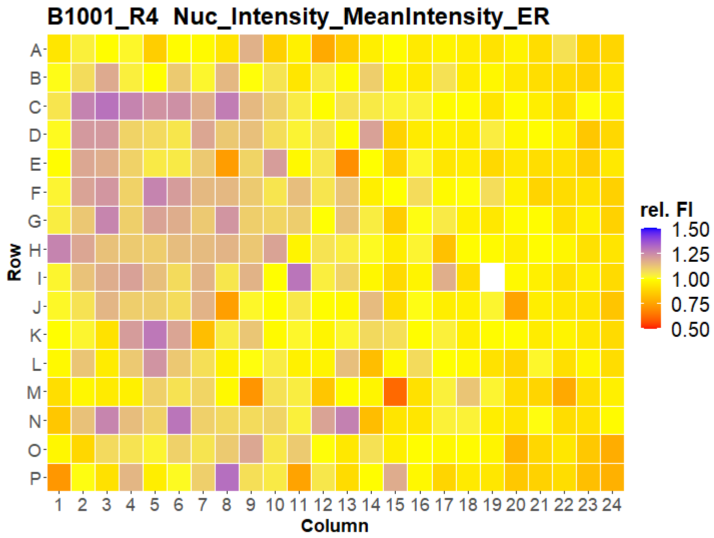
B1001_R4 Nuc_Intensity_MeanIntensity_ER
Row
A B C D E F G H I J K L M N O P
Column
1 2 3 4 5 6 7 8 9 10 11 12 13 14 15 16 17 18 19 20 21 22 23 24
rel. FI
1.50
1.25
1.00
0.75
0.50

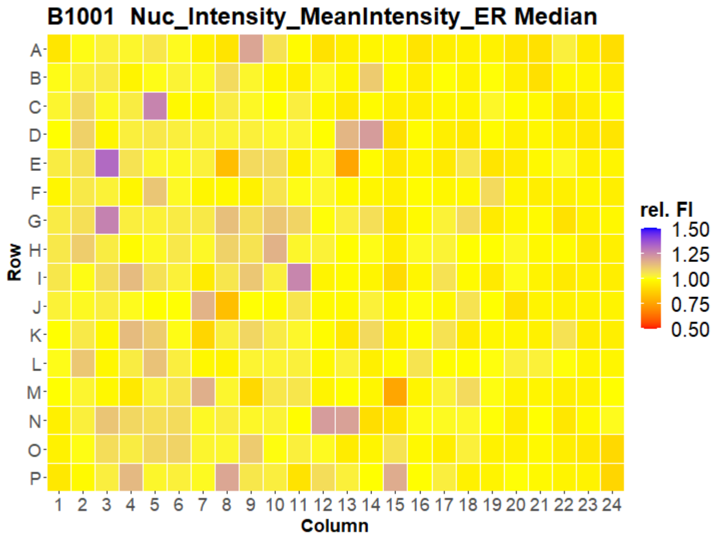
B1001 Nuc_Intensity_MeanIntensity_ER Median
Row
A B C D E F G H I J K L M N O P
Column
1 2 3 4 5 6 7 8 9 10 11 12 13 14 15 16 17 18 19 20 21 22 23 24
rel. FI
1.50
1.25
1.00
0.75
0.50

### HepG2 FMP



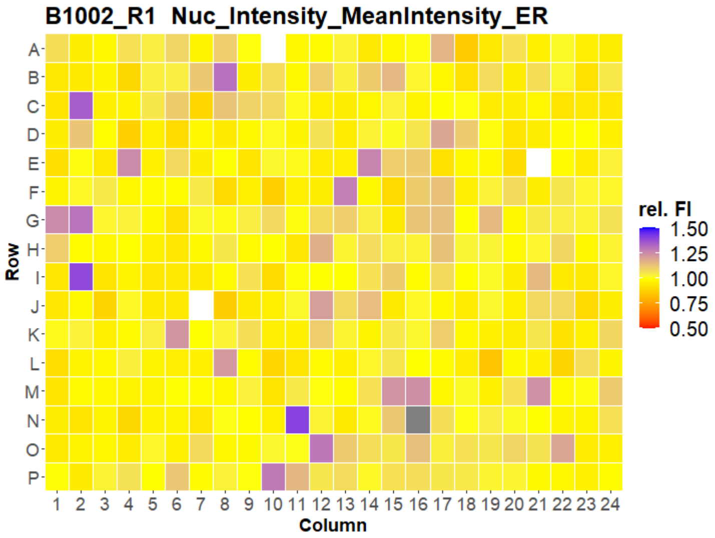
B1002_R1 Nuc_Intensity_MeanIntensity_ER
Row
A B C D E F G H I J K L M N O P
Column
1 2 3 4 5 6 7 8 9 10 11 12 13 14 15 16 17 18 19 20 21 22 23 24
rel. FI
1.50
1.25
1.00
0.75
0.50

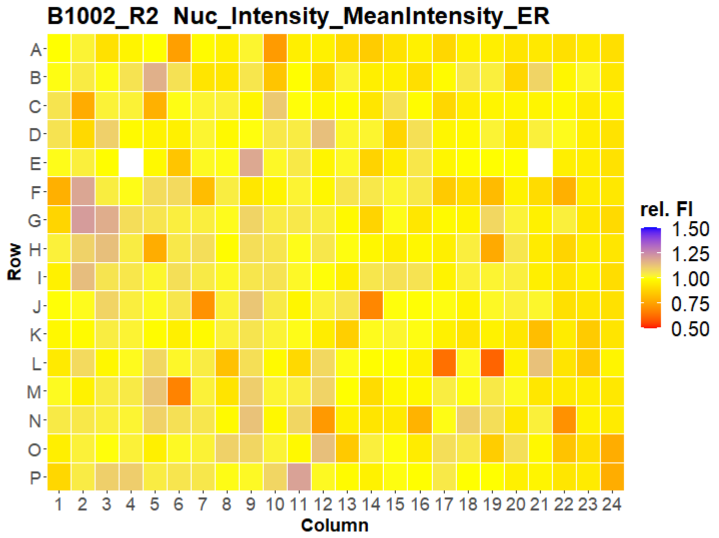
B1002_R2 Nuc_Intensity_MeanIntensity_ER
Row
A B C D E F G H I J K L M N O P
Column
1 2 3 4 5 6 7 8 9 10 11 12 13 14 15 16 17 18 19 20 21 22 23 24
rel. FI
1.50
1.25
1.00
0.75
0.50

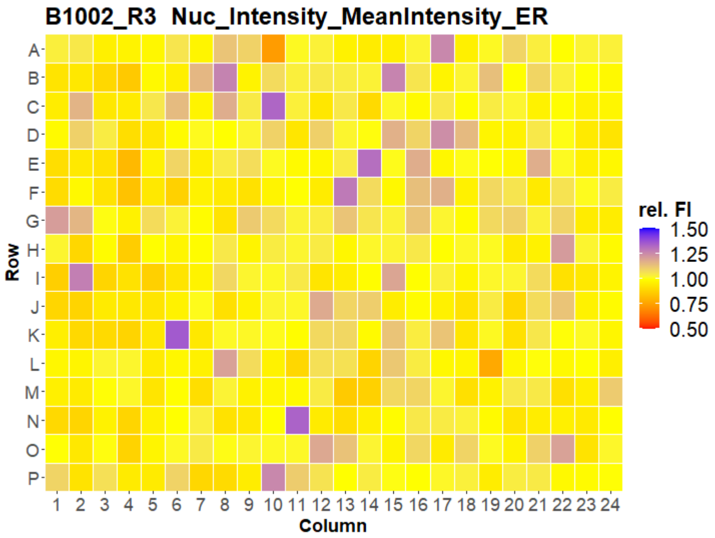
B1002_R3 Nuc_Intensity_MeanIntensity_ER
Row
A B C D E F G H I J K L M N O P
Column
1 2 3 4 5 6 7 8 9 10 11 12 13 14 15 16 17 18 19 20 21 22 23 24
rel. FI
1.50
1.25
1.00
0.75
0.50

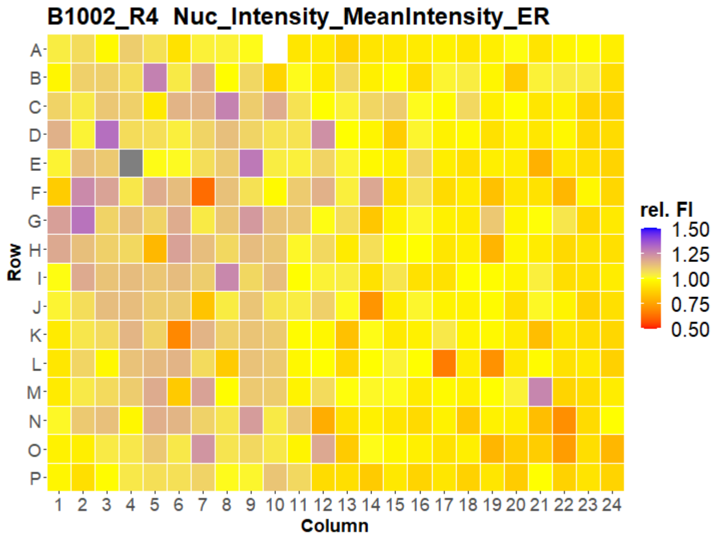
B1002_R4 Nuc_Intensity_MeanIntensity_ER
Row
A B C D E F G H I J K L M N O P
Column
1 2 3 4 5 6 7 8 9 10 11 12 13 14 15 16 17 18 19 20 21 22 23 24
rel. FI
1.50
1.25
1.00
0.75
0.50

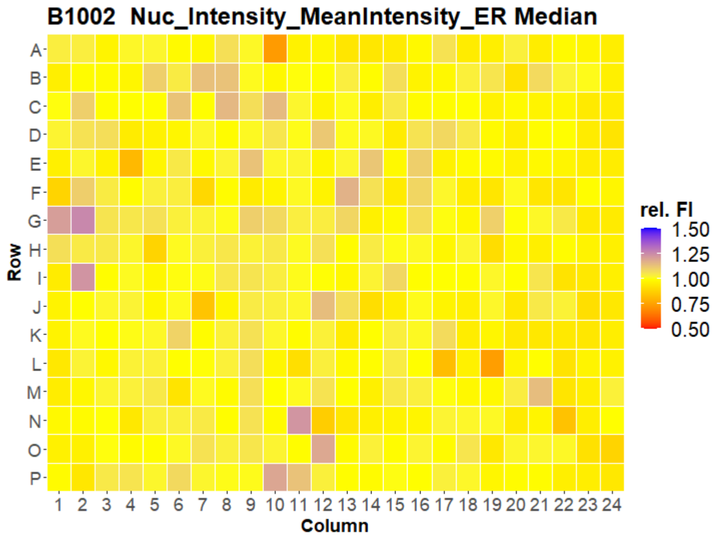
B1002 Nuc_Intensity_MeanIntensity_ER Median
Row
A B C D E F G H I J K L M N O P
Column
1 2 3 4 5 6 7 8 9 10 11 12 13 14 15 16 17 18 19 20 21 22 23 24
rel. FI
1.50
1.25
1.00
0.75
0.50

### HepG2 FMP



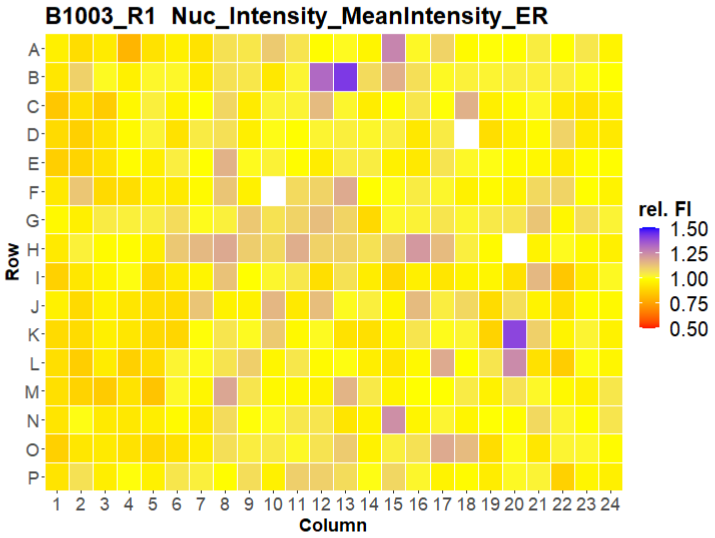
B1003_R1 Nuc_Intensity_MeanIntensity_ER
Row
Column
rel. FI
1.50
1.25
1.00
0.75
0.50

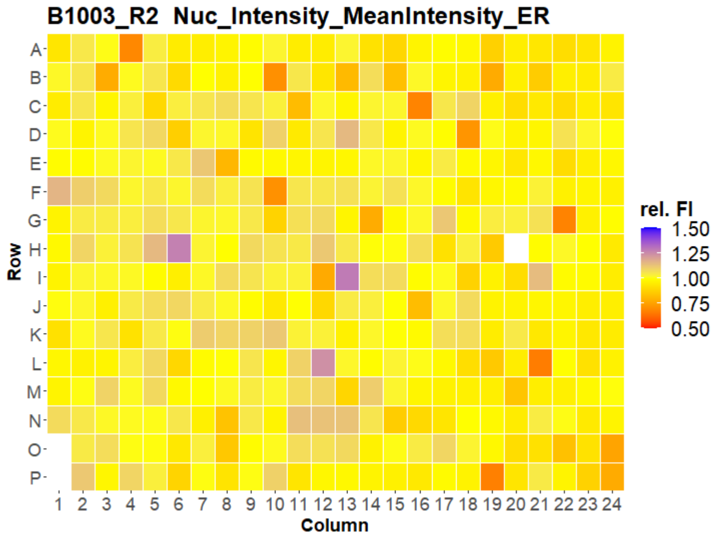
B1003_R2 Nuc_Intensity_MeanIntensity_ER
Row
Column
rel. FI
1.50
1.25
1.00
0.75
0.50

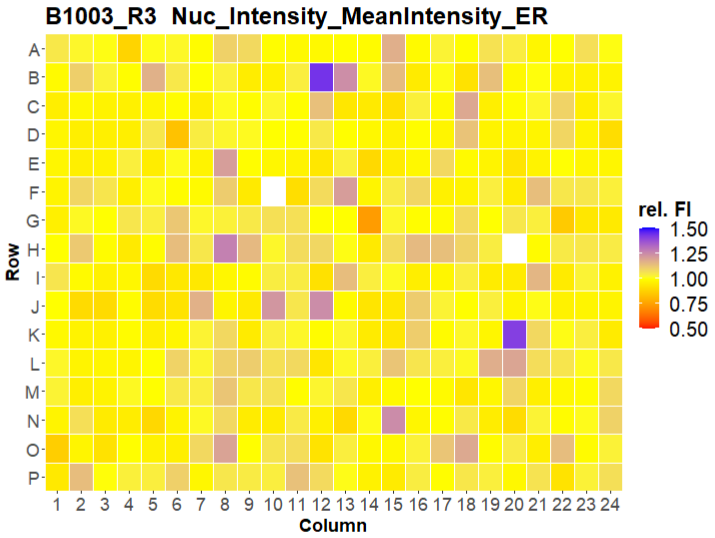
B1003_R3 Nuc_Intensity_MeanIntensity_ER
Row
Column
rel. FI
1.50
1.25
1.00
0.75
0.50

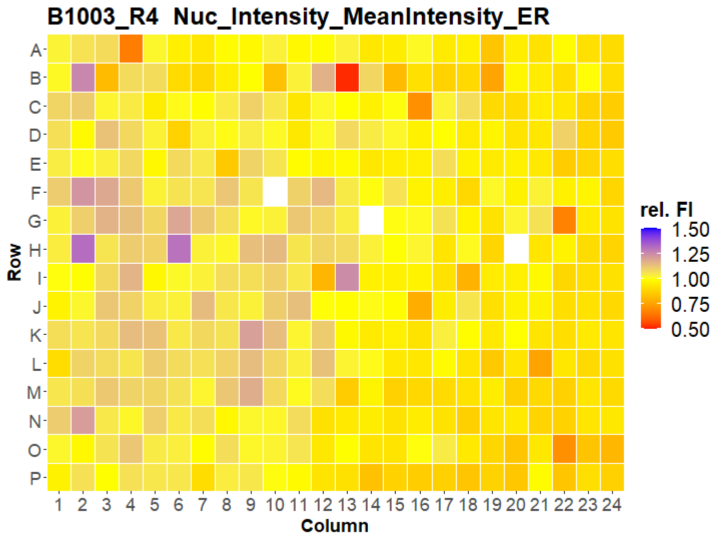
B1003_R4 Nuc_Intensity_MeanIntensity_ER
Row
Column
rel. FI
1.50
1.25
1.00
0.75
0.50

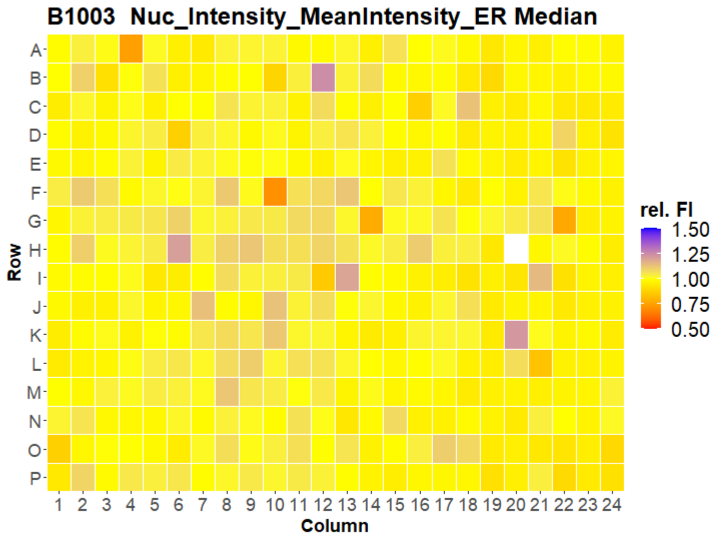
B1003 Nuc_Intensity_MeanIntensity_ER Median
Row
Column
rel. FI
1.50
1.25
1.00
0.75
0.50

### HepG2 FMP

39

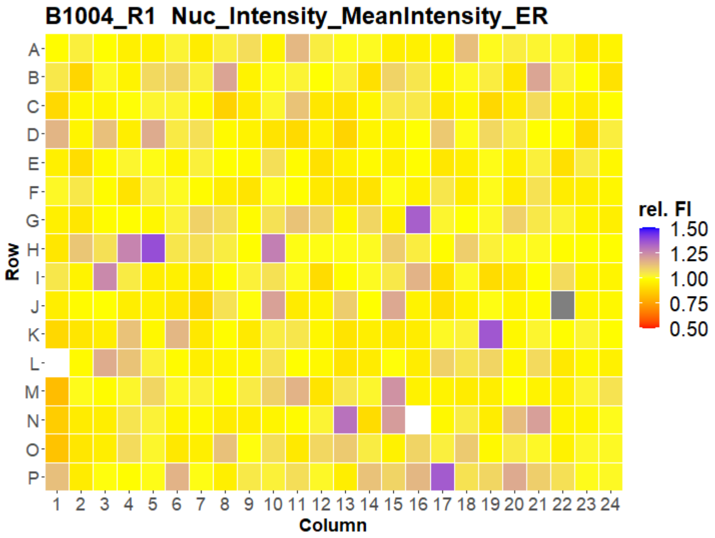

B1004_R1 Nuc_Intensity_MeanIntensity_ER
Row
Column
rel. FI
1.50
1.25
1.00
0.75
0.50


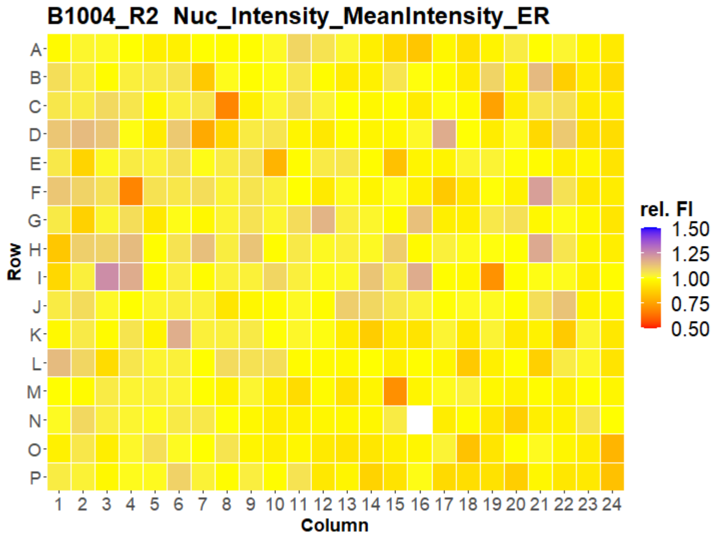

B1004_R2 Nuc_Intensity_MeanIntensity_ER
Row
Column
rel. FI
1.50
1.25
1.00
0.75
0.50


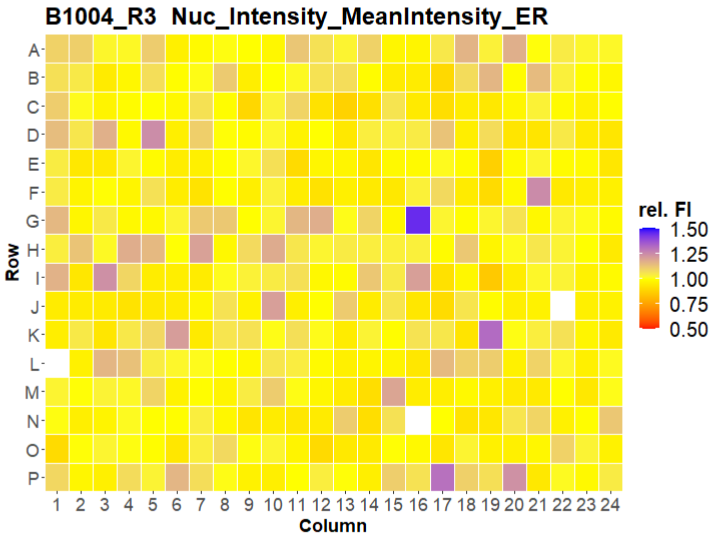

B1004_R3 Nuc_Intensity_MeanIntensity_ER
Row
Column
rel. FI
1.50
1.25
1.00
0.75
0.50


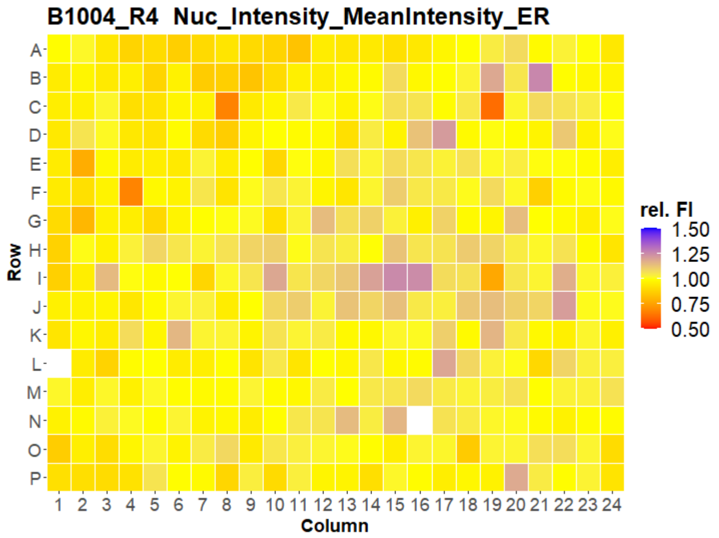

B1004_R4 Nuc_Intensity_MeanIntensity_ER
Row
Column
rel. FI
1.50
1.25
1.00
0.75
0.50


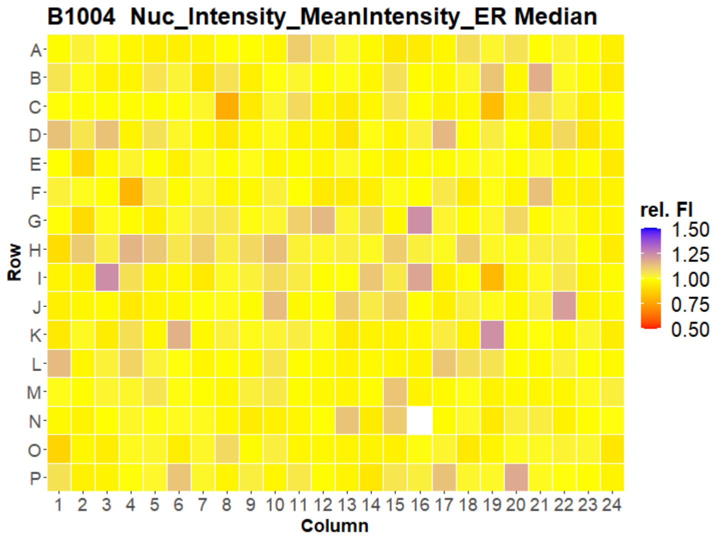

B1004 Nuc_Intensity_MeanIntensity_ER Median
Row
Column
rel. FI
1.50
1.25
1.00
0.75
0.50

### HepG2 FMP

40

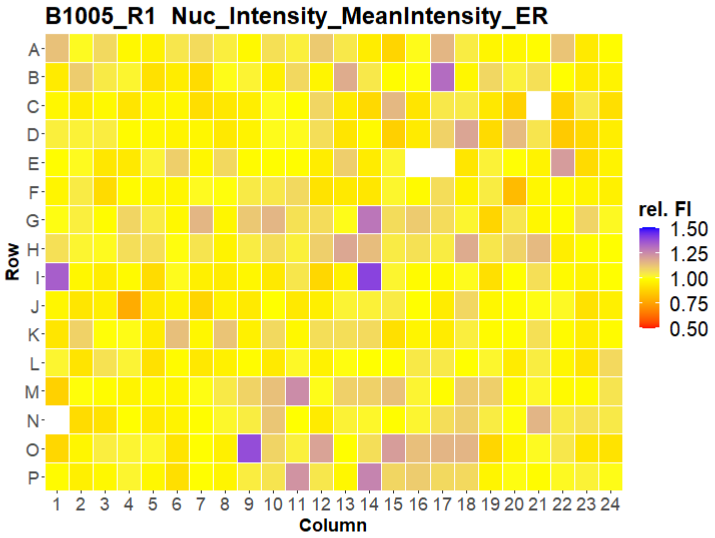
B1005_R1 Nuc_Intensity_MeanIntensity_ER
Row
A B C D E F G H I J K L M N O P
Column
1 2 3 4 5 6 7 8 9 10 11 12 13 14 15 16 17 18 19 20 21 22 23 24
rel. FI
1.50
1.25
1.00
0.75
0.50

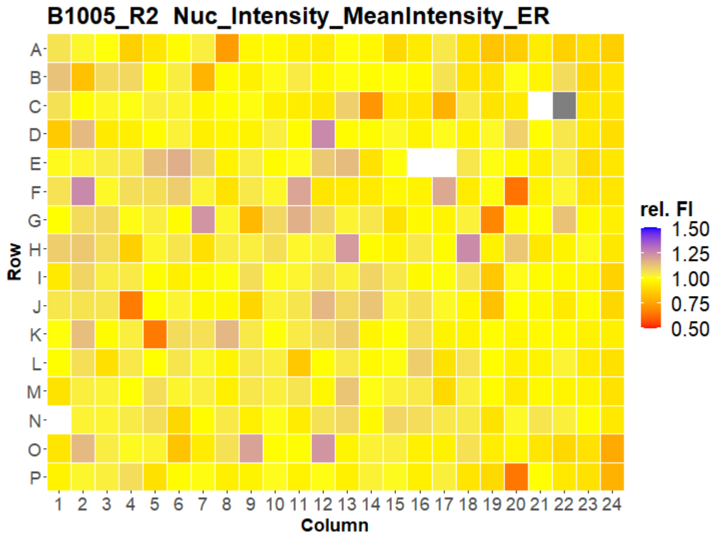
B1005_R2 Nuc_Intensity_MeanIntensity_ER
Row
A B C D E F G H I J K L M N O P
Column
1 2 3 4 5 6 7 8 9 10 11 12 13 14 15 16 17 18 19 20 21 22 23 24
rel. FI
1.50
1.25
1.00
0.75
0.50

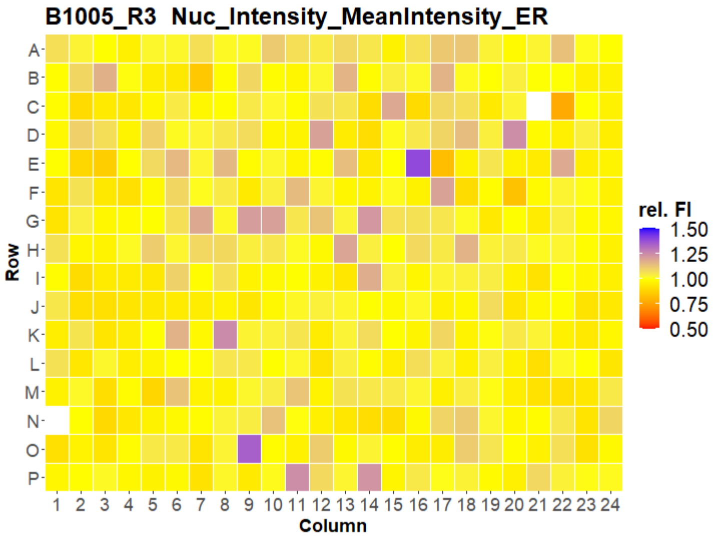
B1005_R3 Nuc_Intensity_MeanIntensity_ER
Row
A B C D E F G H I J K L M N O P
Column
1 2 3 4 5 6 7 8 9 10 11 12 13 14 15 16 17 18 19 20 21 22 23 24
rel. FI
1.50
1.25
1.00
0.75
0.50

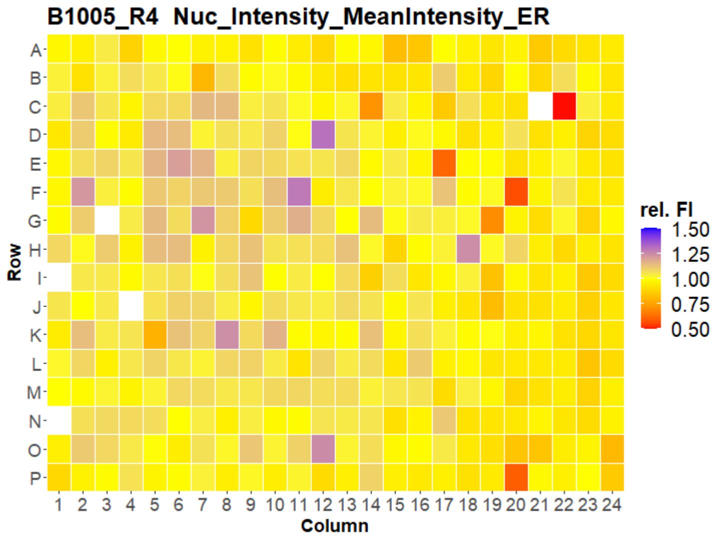
B1005_R4 Nuc_Intensity_MeanIntensity_ER
Row
A B C D E F G H I J K L M N O P
Column
1 2 3 4 5 6 7 8 9 10 11 12 13 14 15 16 17 18 19 20 21 22 23 24
rel. FI
1.50
1.25
1.00
0.75
0.50

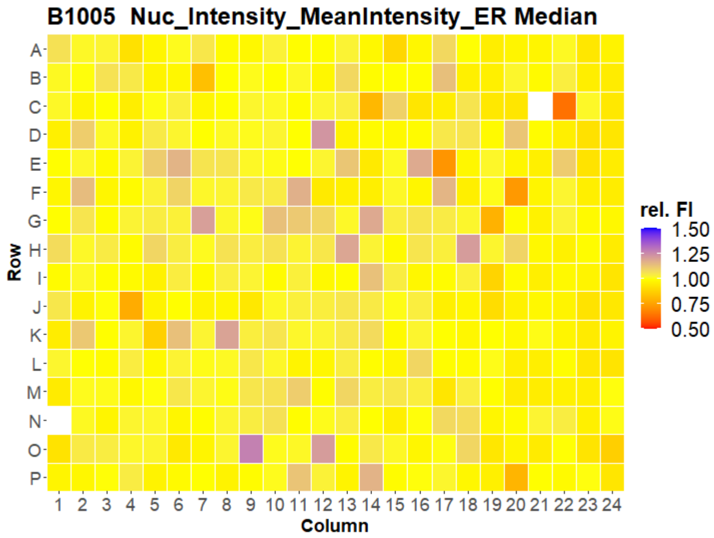
B1005 Nuc_Intensity_MeanIntensity_ER Median
Row
A B C D E F G H I J K L M N O P
Column
1 2 3 4 5 6 7 8 9 10 11 12 13 14 15 16 17 18 19 20 21 22 23 24
rel. FI
1.50
1.25
1.00
0.75
0.50

### HepG2 FMP

41

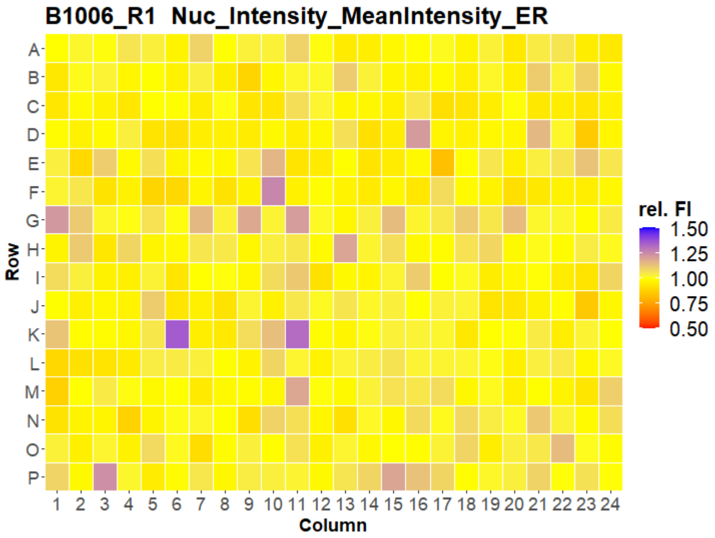
B1006_R1 Nuc_Intensity_MeanIntensity_ER
A B C D E F G H I J K L M N O P
Row
1 2 3 4 5 6 7 8 9 10 11 12 13 14 15 16 17 18 19 20 21 22 23 24
Column
rel. FI
1.50
1.25
1.00
0.75
0.50

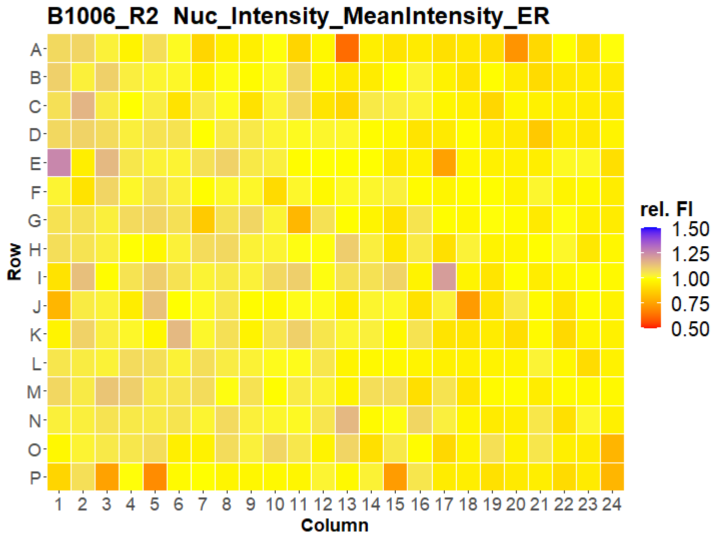
B1006_R2 Nuc_Intensity_MeanIntensity_ER
A B C D E F G H I J K L M N O P
Row
1 2 3 4 5 6 7 8 9 10 11 12 13 14 15 16 17 18 19 20 21 22 23 24
Column
rel. FI
1.50
1.25
1.00
0.75
0.50

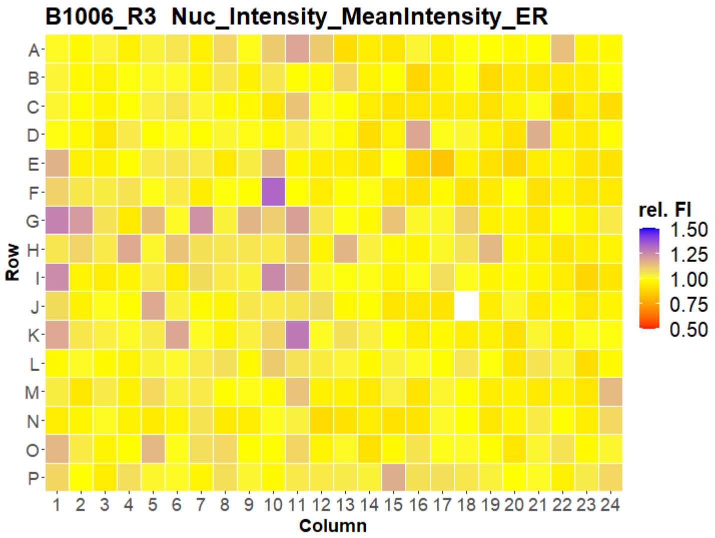
B1006_R3 Nuc_Intensity_MeanIntensity_ER
A B C D E F G H I J K L M N O P
Row
1 2 3 4 5 6 7 8 9 10 11 12 13 14 15 16 17 18 19 20 21 22 23 24
Column
rel. FI
1.50
1.25
1.00
0.75
0.50

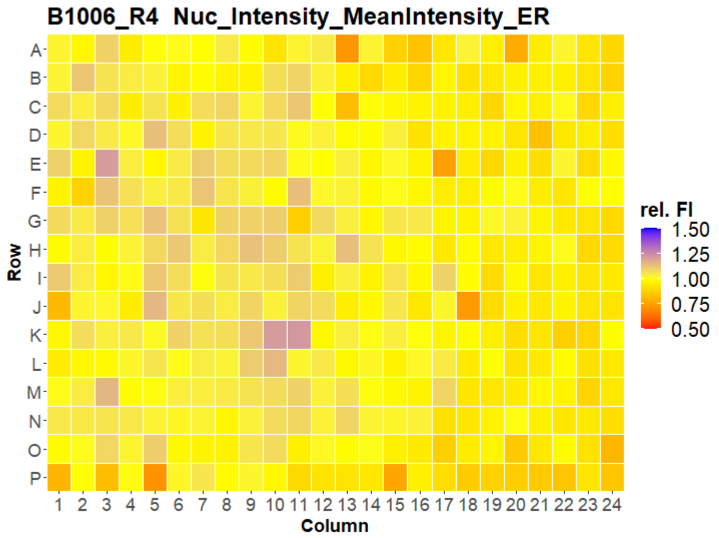
B1006_R4 Nuc_Intensity_MeanIntensity_ER
A B C D E F G H I J K L M N O P
Row
1 2 3 4 5 6 7 8 9 10 11 12 13 14 15 16 17 18 19 20 21 22 23 24
Column
rel. FI
1.50
1.25
1.00
0.75
0.50

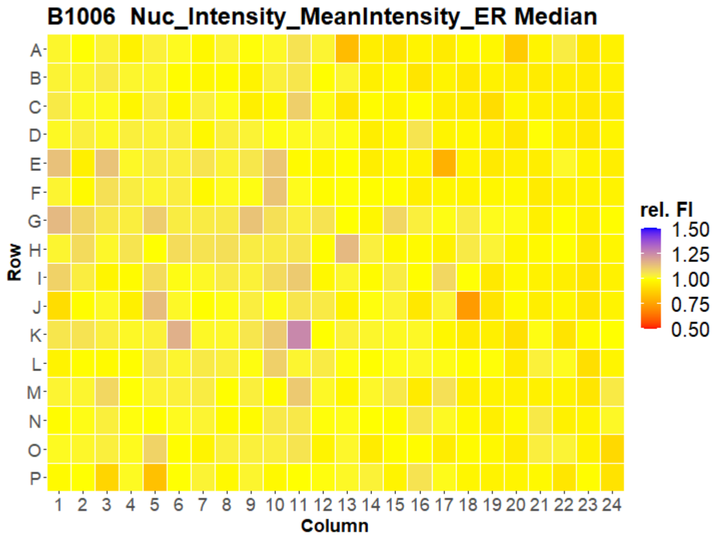
B1006 Nuc_Intensity_MeanIntensity_ER Median
A B C D E F G H I J K L M N O P
Row
1 2 3 4 5 6 7 8 9 10 11 12 13 14 15 16 17 18 19 20 21 22 23 24
Column
rel. FI
1.50
1.25
1.00
0.75
0.50

### HepG2 FMP



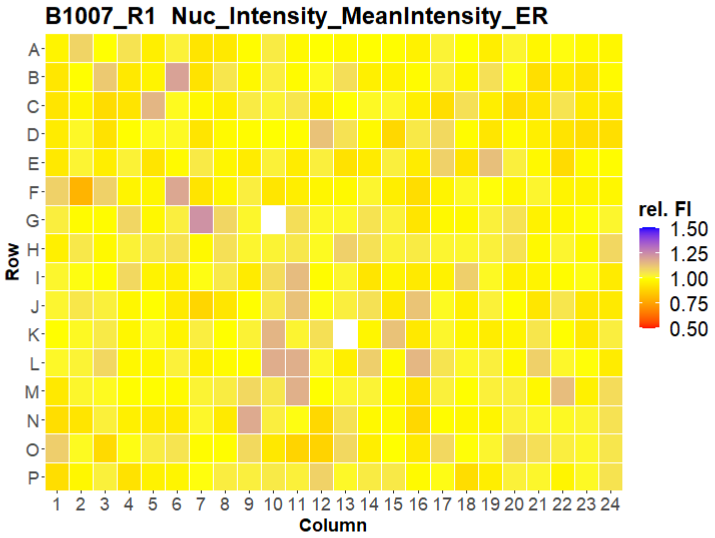
B1007_R1 Nuc_Intensity_MeanIntensity_ER
Row
Column
rel. FI
1.50
1.25
1.00
0.75
0.50

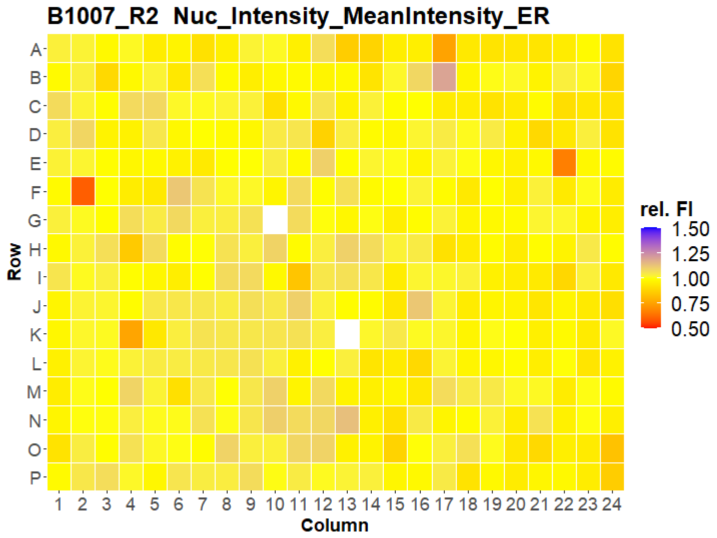
B1007_R2 Nuc_Intensity_MeanIntensity_ER
Row
Column
rel. FI
1.50
1.25
1.00
0.75
0.50

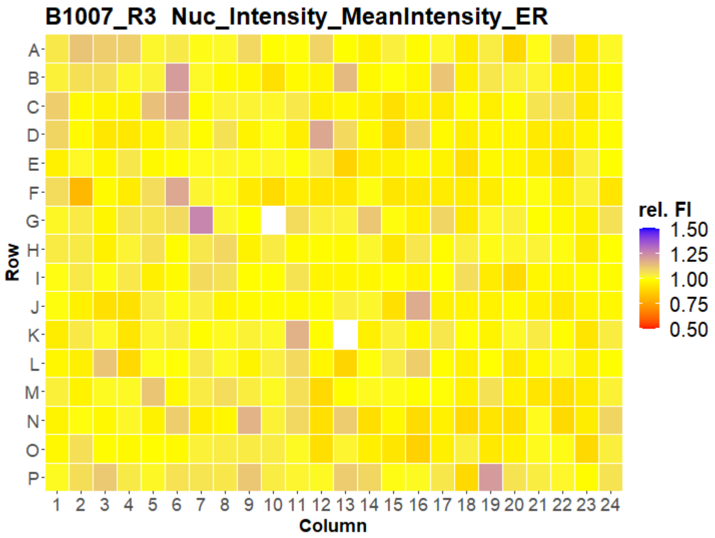
B1007_R3 Nuc_Intensity_MeanIntensity_ER
Row
Column
rel. FI
1.50
1.25
1.00
0.75
0.50

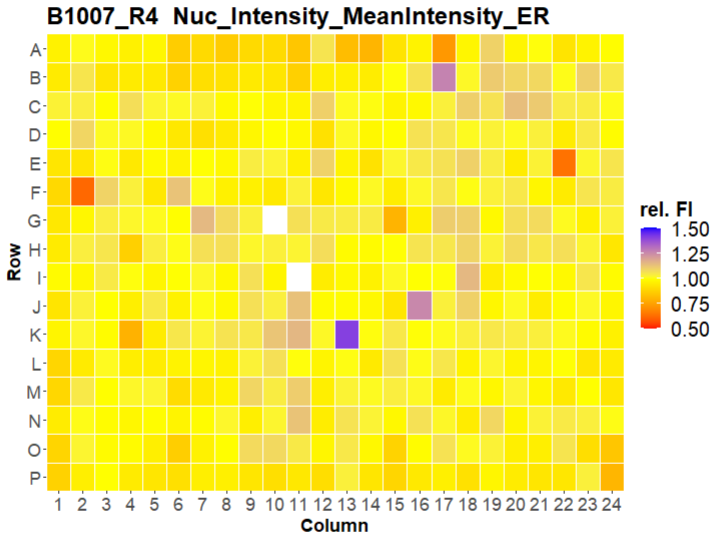
B1007_R4 Nuc_Intensity_MeanIntensity_ER
Row
Column
rel. FI
1.50
1.25
1.00
0.75
0.50

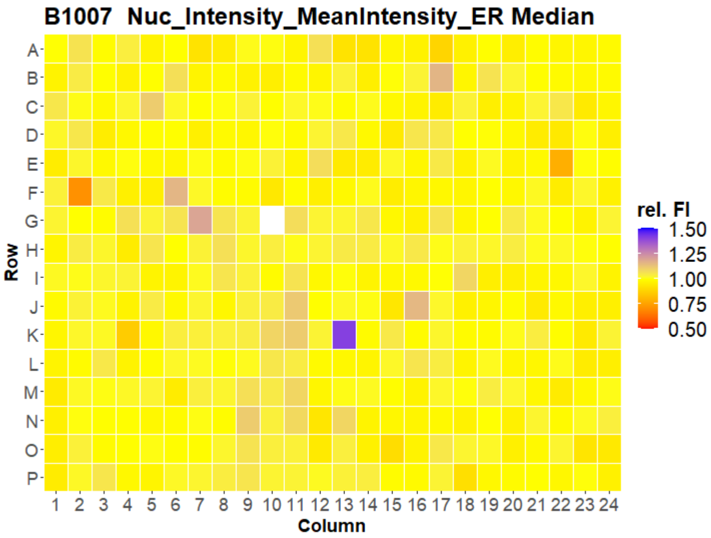
B1007 Nuc_Intensity_MeanIntensity_ER Median
Row
Column
rel. FI
1.50
1.25
1.00
0.75
0.50

### HepG2 FMP



Open for Innovation KNIME
