## Supplementary material for "Morphological Profiling Dataset of EU-OPENSCREEN Bioactive Compounds Over Multiple Imaging Sites and Cell Lines": Suppl_QC_FMP_U2OS.pdf

### U2OS FMP

1

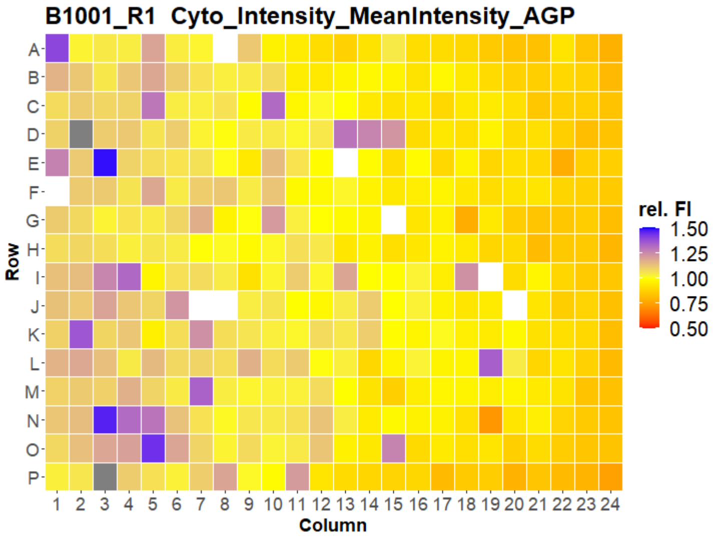
B1001_R1 Cyto_Intensity_MeanIntensity_AGP
rel. FI
1.50
1.25
1.00
0.75
0.50
Row
Column

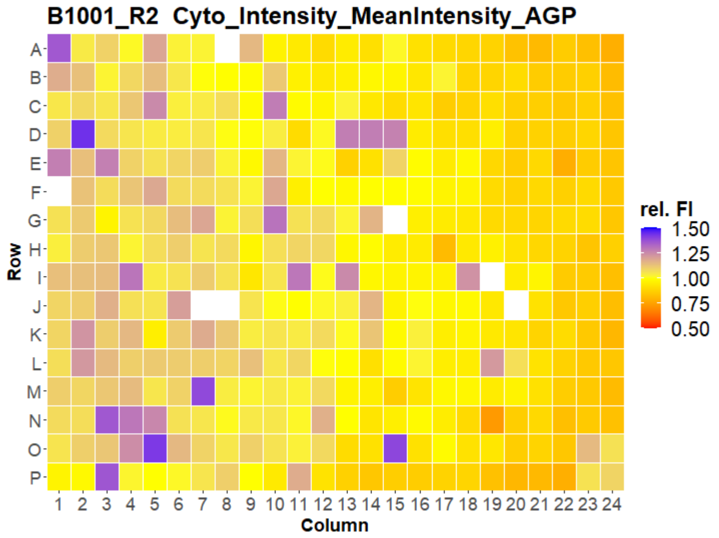
B1001_R2 Cyto_Intensity_MeanIntensity_AGP
rel. FI
1.50
1.25
1.00
0.75
0.50
Row
Column

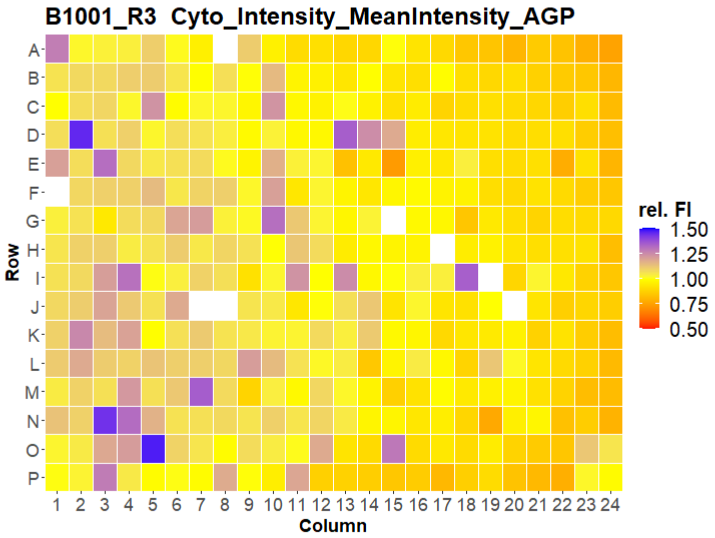
B1001_R3 Cyto_Intensity_MeanIntensity_AGP
rel. FI
1.50
1.25
1.00
0.75
0.50
Row
Column

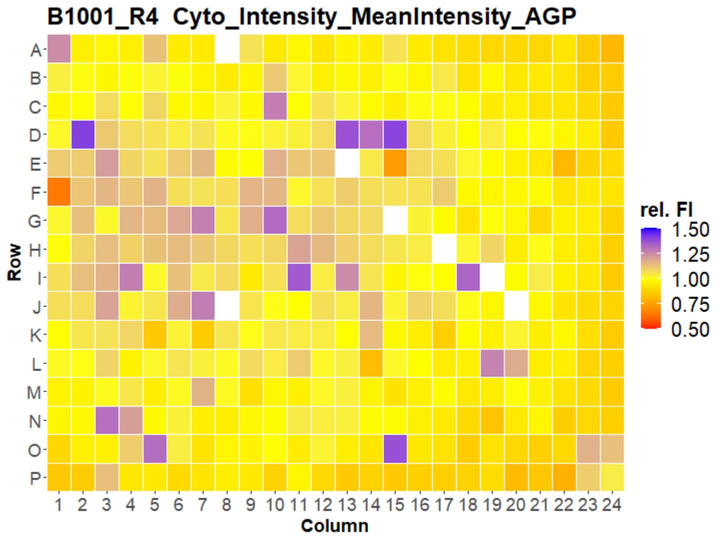
B1001_R4 Cyto_Intensity_MeanIntensity_AGP
rel. FI
1.50
1.25
1.00
0.75
0.50
Row
Column

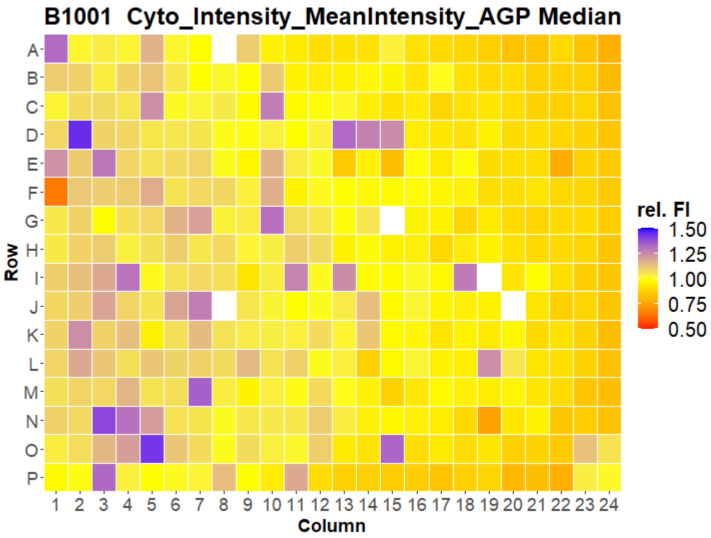
B1001 Cyto_Intensity_MeanIntensity_AGP Median
rel. FI
1.50
1.25
1.00
0.75
0.50
Row
Column

### U2OS FMP

2

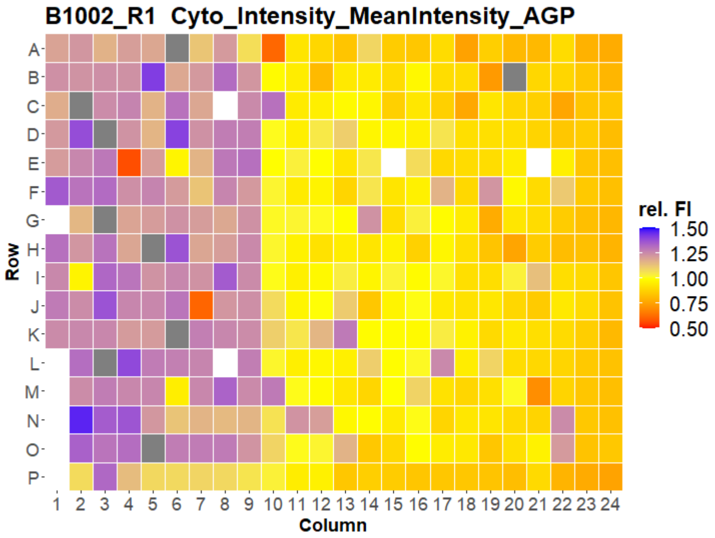

B1002_R1 Cyto_Intensity_MeanIntensity_AGP
Row
Column
rel. FI
1.50
1.25
1.00
0.75
0.50


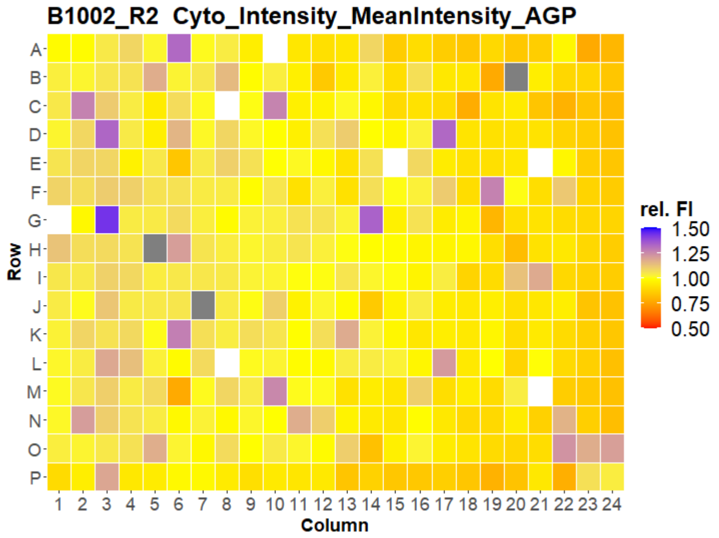

B1002_R2 Cyto_Intensity_MeanIntensity_AGP
Row
Column
rel. FI
1.50
1.25
1.00
0.75
0.50


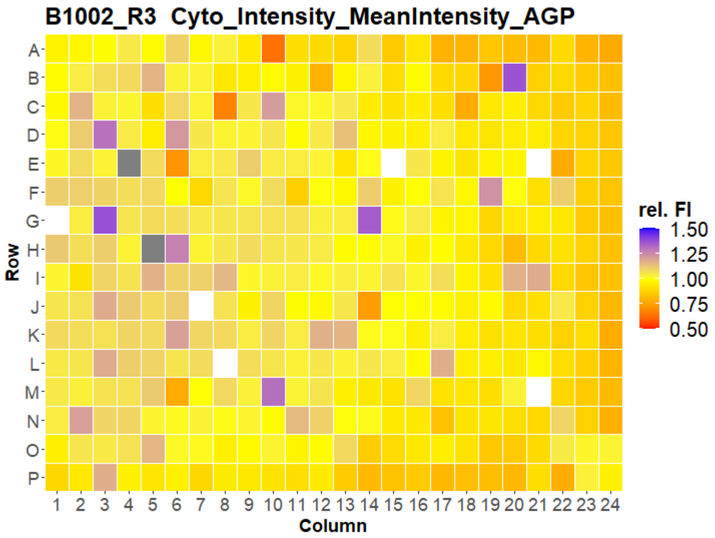

B1002_R3 Cyto_Intensity_MeanIntensity_AGP
Row
Column
rel. FI
1.50
1.25
1.00
0.75
0.50


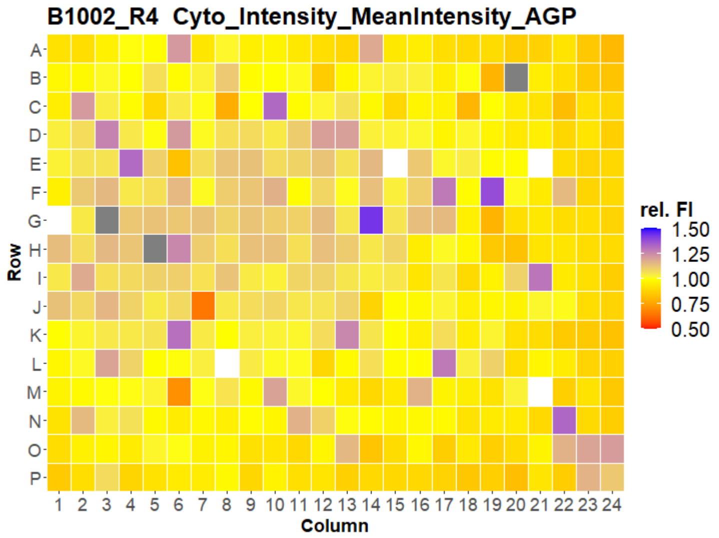

B1002_R4 Cyto_Intensity_MeanIntensity_AGP
Row
Column
rel. FI
1.50
1.25
1.00
0.75
0.50


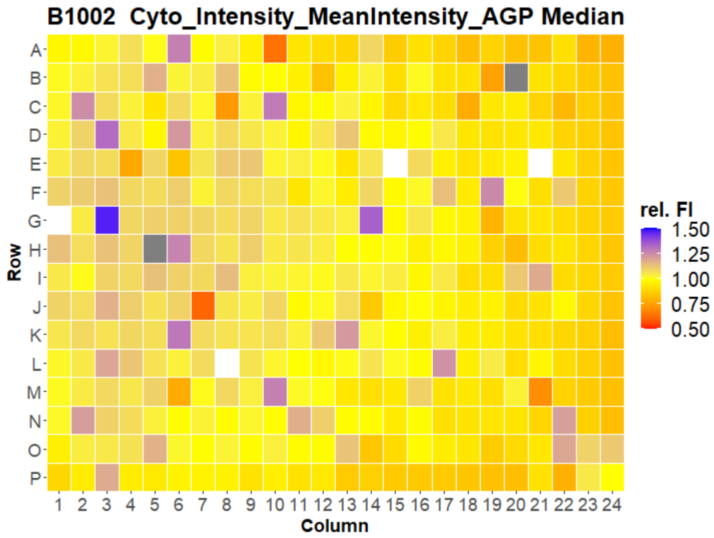

B1002 Cyto_Intensity_MeanIntensity_AGP Median
Row
Column
rel. FI
1.50
1.25
1.00
0.75
0.50

### U2OS FMP

3

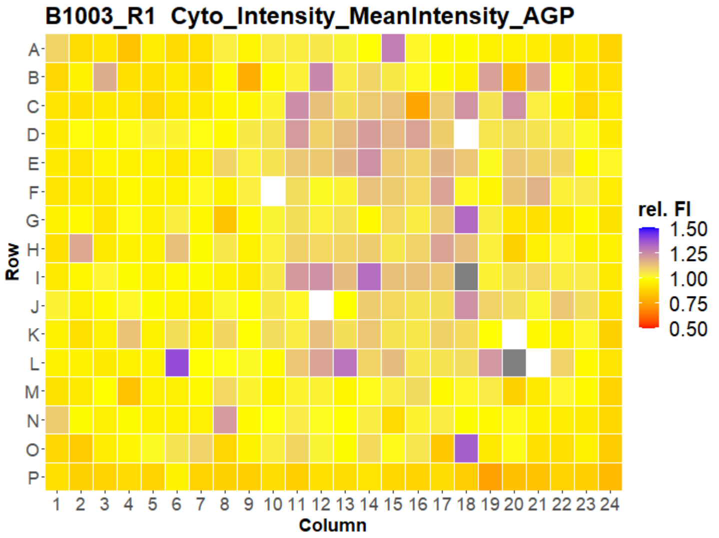


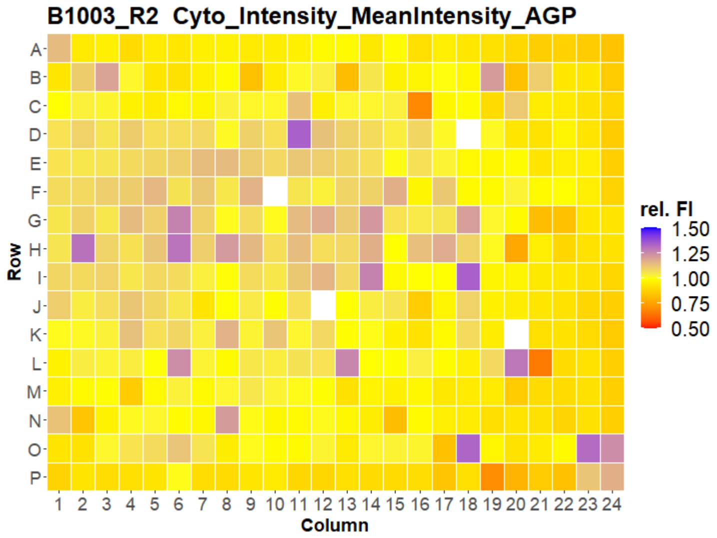


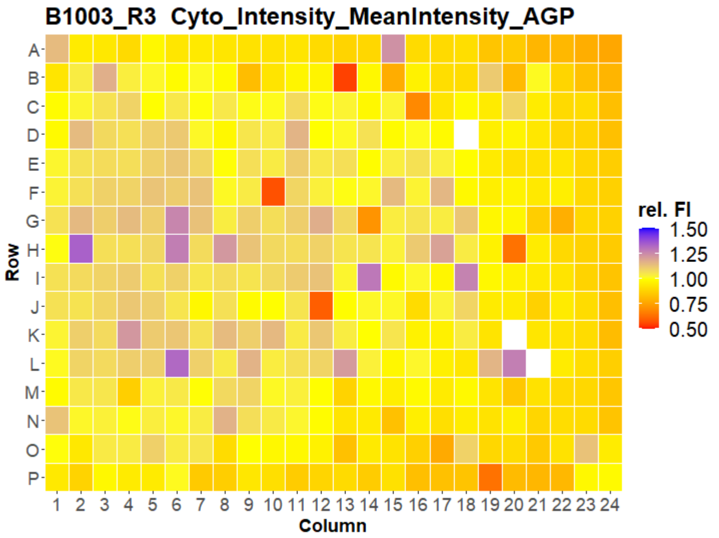


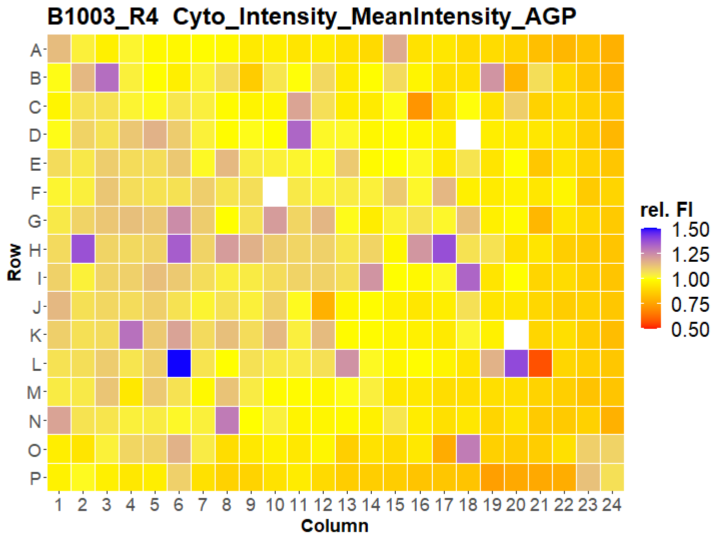


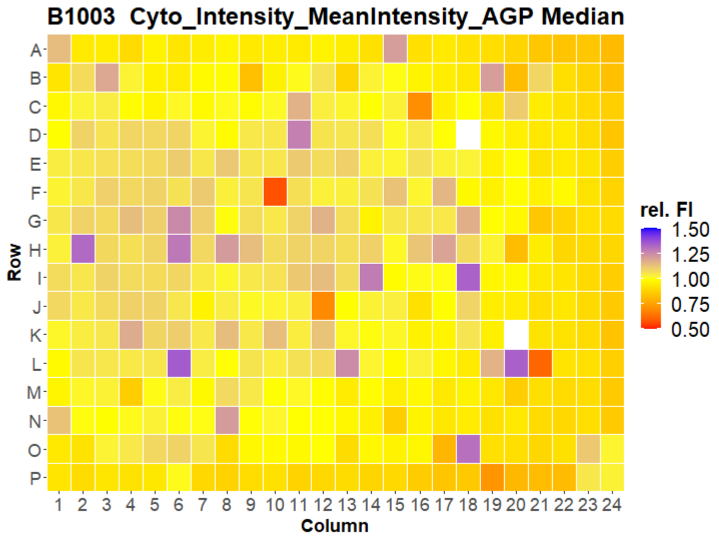



Open for Innovation KNIME

### U2OS FMP



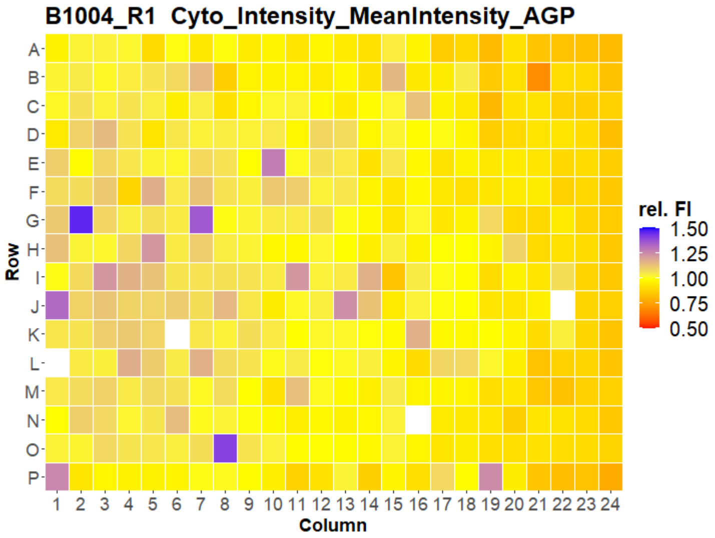
B1004_R1 Cyto_Intensity_MeanIntensity_AGP
Row
Column
rel. FI
1.50
1.25
1.00
0.75
0.50

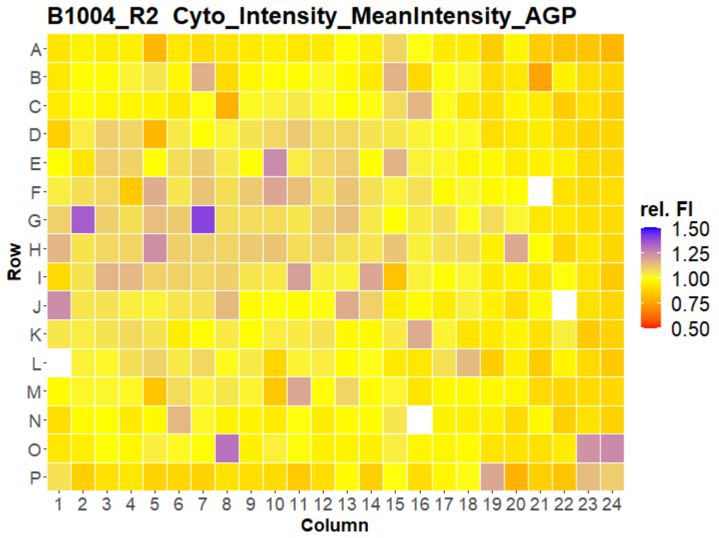
B1004_R2 Cyto_Intensity_MeanIntensity_AGP
Row
Column
rel. FI
1.50
1.25
1.00
0.75
0.50

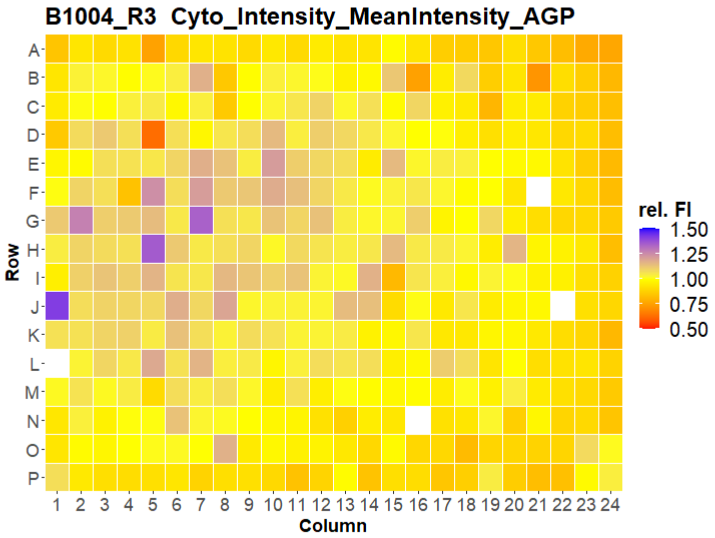
B1004_R3 Cyto_Intensity_MeanIntensity_AGP
Row
Column
rel. FI
1.50
1.25
1.00
0.75
0.50

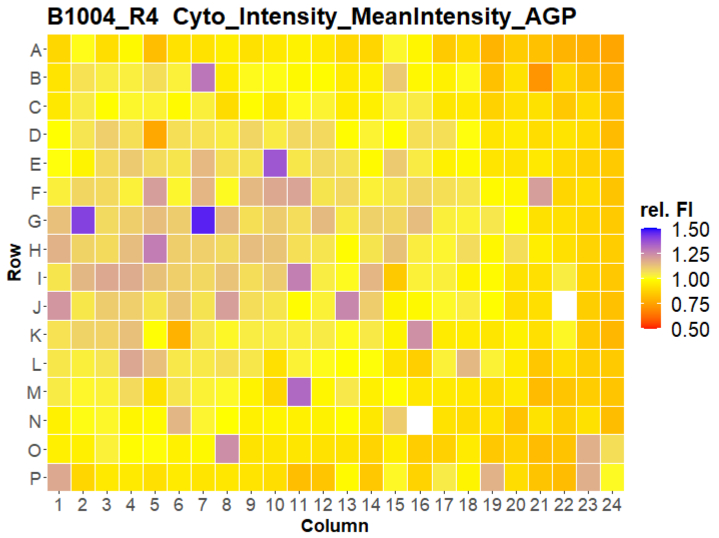
B1004_R4 Cyto_Intensity_MeanIntensity_AGP
Row
Column
rel. FI
1.50
1.25
1.00
0.75
0.50

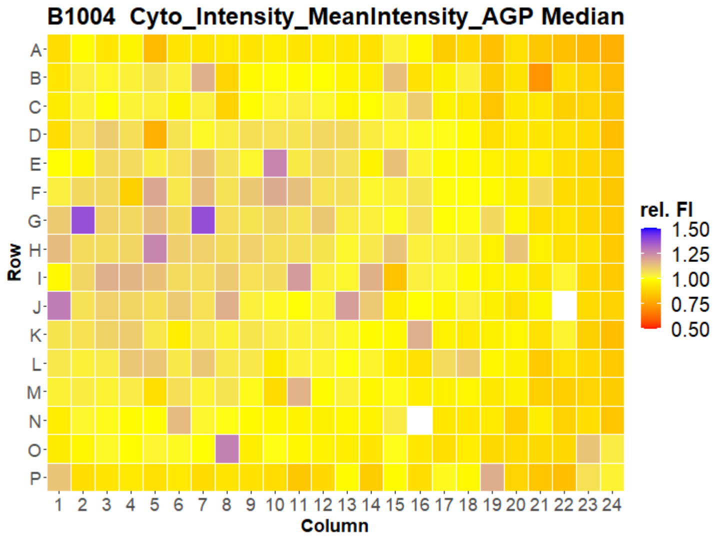
B1004 Cyto_Intensity_MeanIntensity_AGP Median
Row
Column
rel. FI
1.50
1.25
1.00
0.75
0.50

### U2OS FMP

5

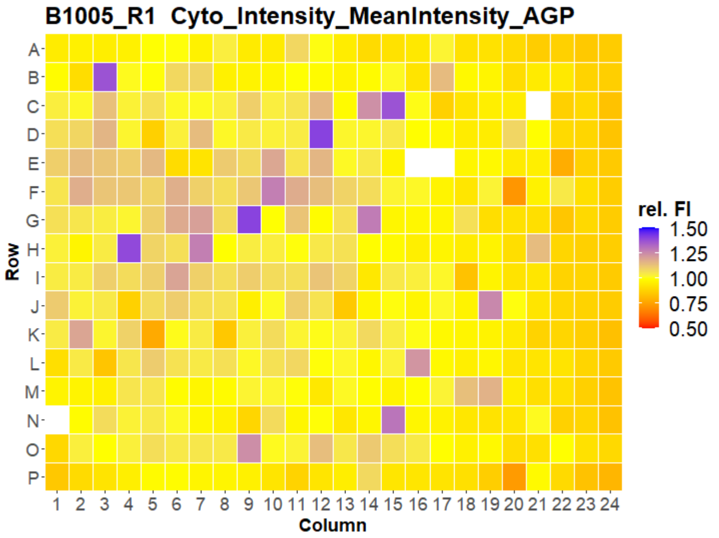


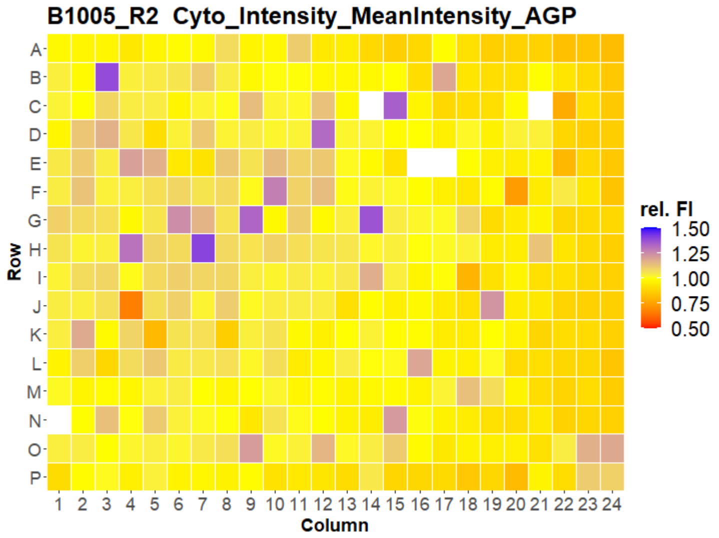


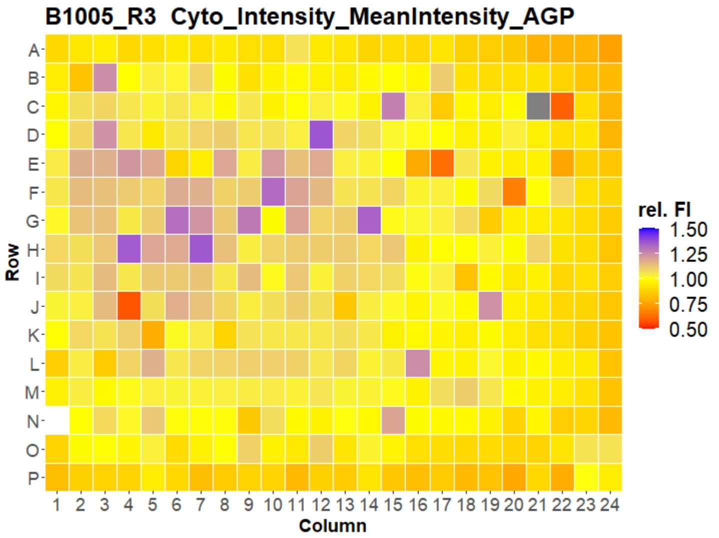


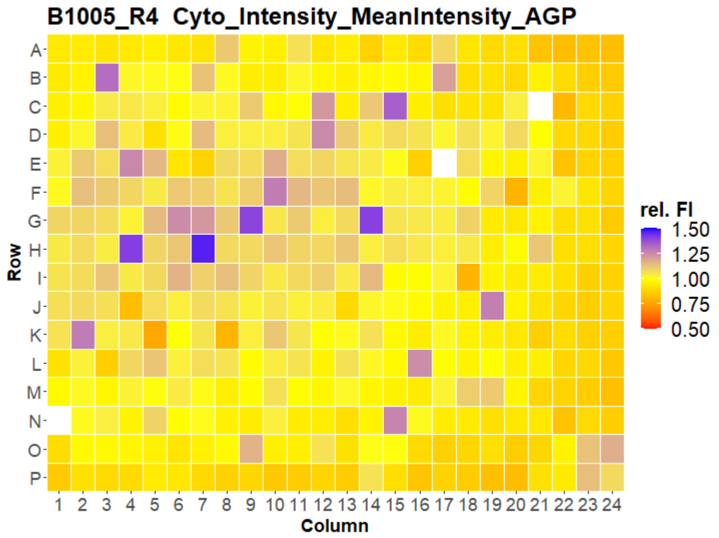


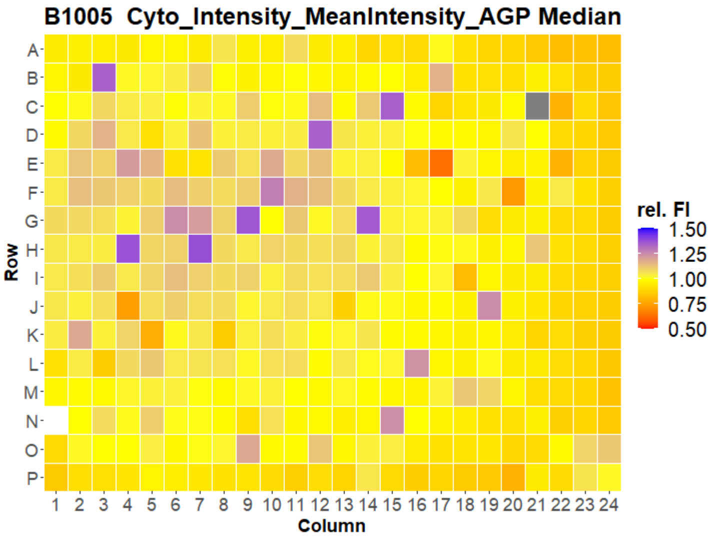



Open for Innovation KNIME

### U2OS FMP

6

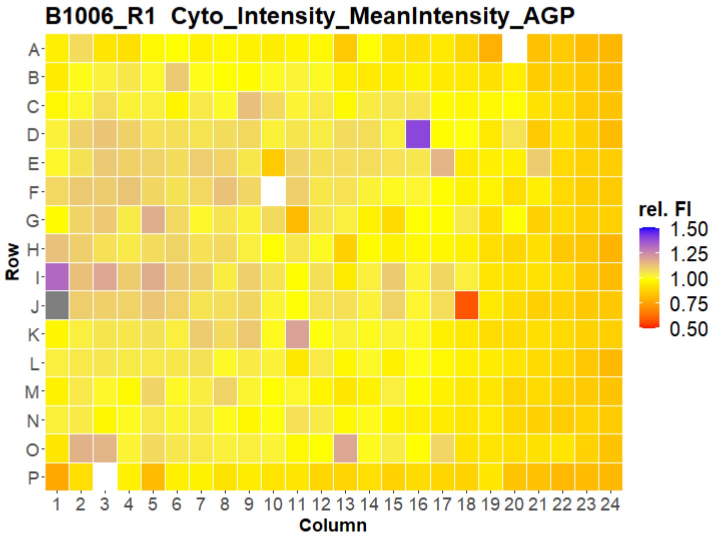
B1006_R1 Cyto_Intensity_MeanIntensity_AGP
Row
Column
rel. FI
1.50
1.25
1.00
0.75
0.50

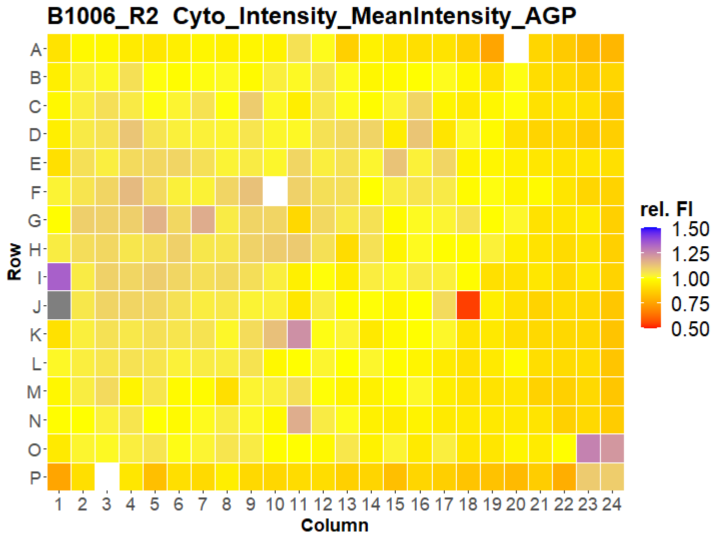
B1006_R2 Cyto_Intensity_MeanIntensity_AGP
Row
Column
rel. FI
1.50
1.25
1.00
0.75
0.50

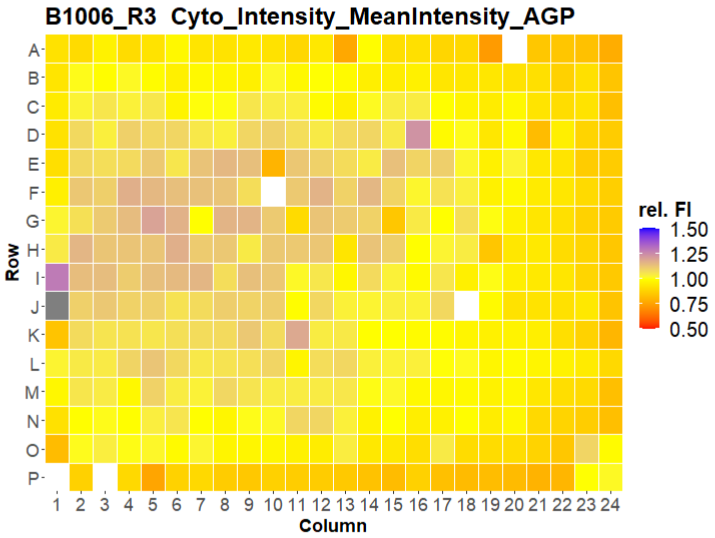
B1006_R3 Cyto_Intensity_MeanIntensity_AGP
Row
Column
rel. FI
1.50
1.25
1.00
0.75
0.50

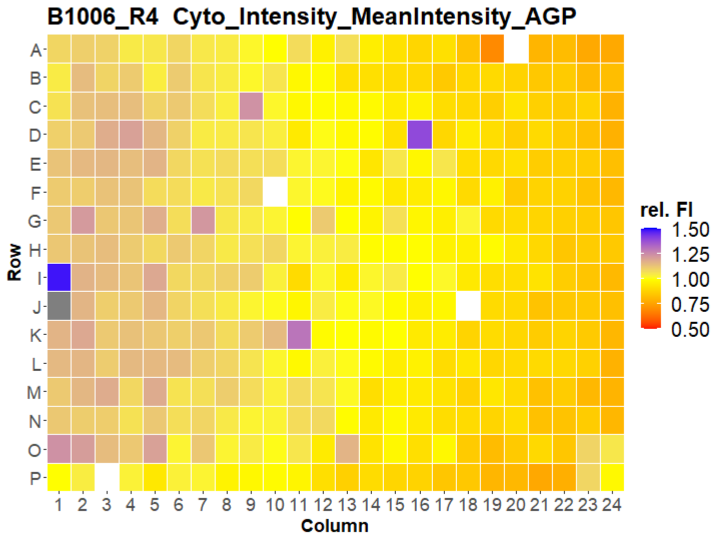
B1006_R4 Cyto_Intensity_MeanIntensity_AGP
Row
Column
rel. FI
1.50
1.25
1.00
0.75
0.50

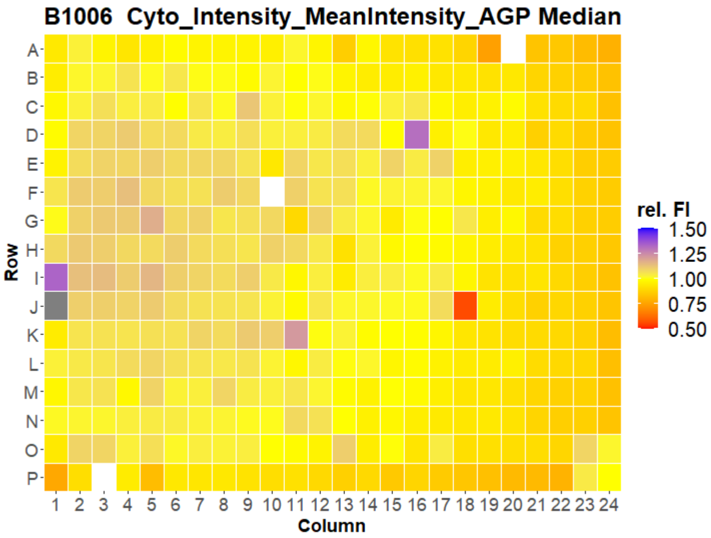
B1006 Cyto_Intensity_MeanIntensity_AGP Median
Row
Column
rel. FI
1.50
1.25
1.00
0.75
0.50

### U2OS FMP

7

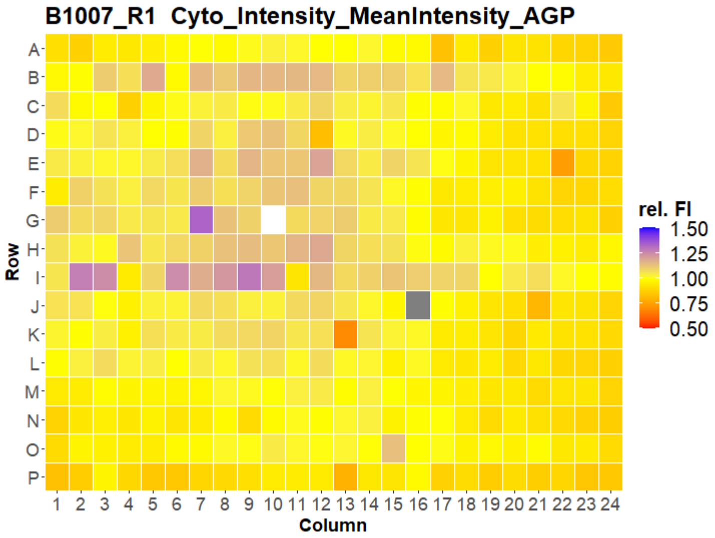
B1007_R1 Cyto_Intensity_MeanIntensity_AGP
Row
Column
rel. FI
1.50
1.25
1.00
0.75
0.50

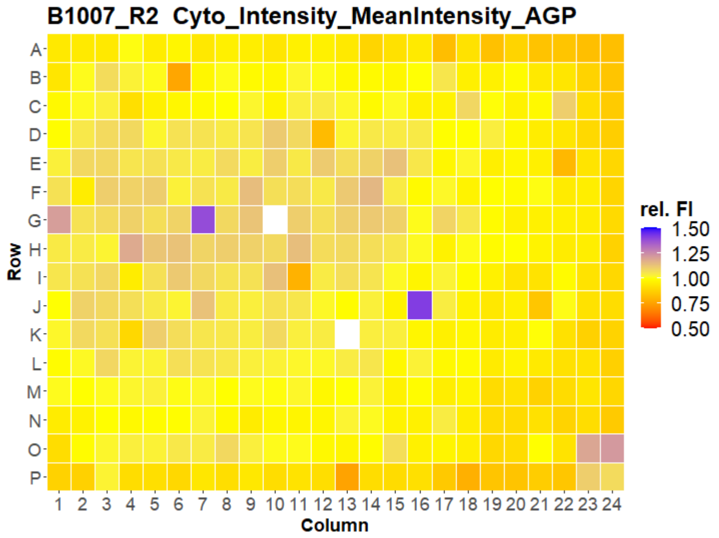
B1007_R2 Cyto_Intensity_MeanIntensity_AGP
Row
Column
rel. FI
1.50
1.25
1.00
0.75
0.50

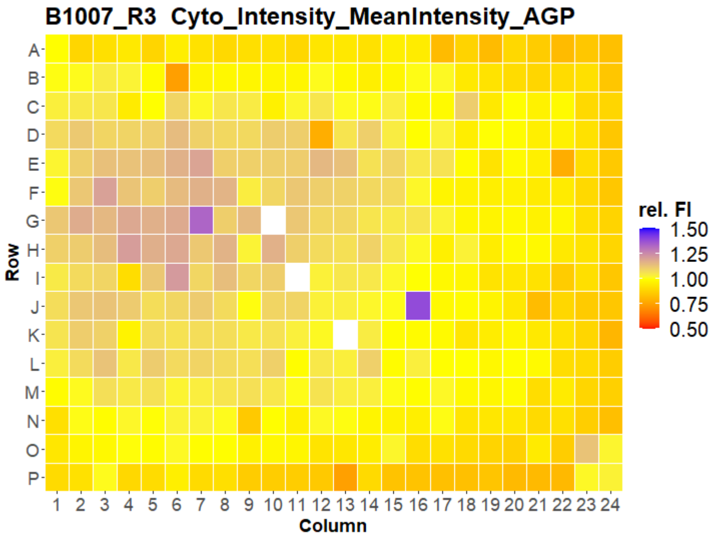
B1007_R3 Cyto_Intensity_MeanIntensity_AGP
Row
Column
rel. FI
1.50
1.25
1.00
0.75
0.50

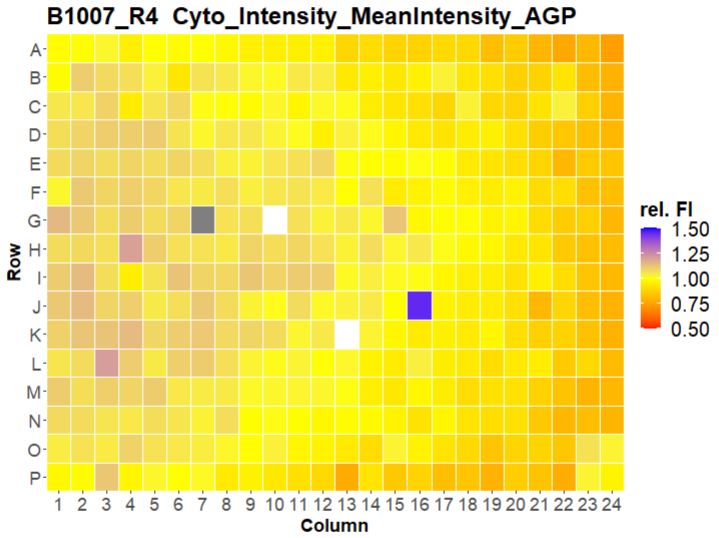
B1007_R4 Cyto_Intensity_MeanIntensity_AGP
Row
Column
rel. FI
1.50
1.25
1.00
0.75
0.50

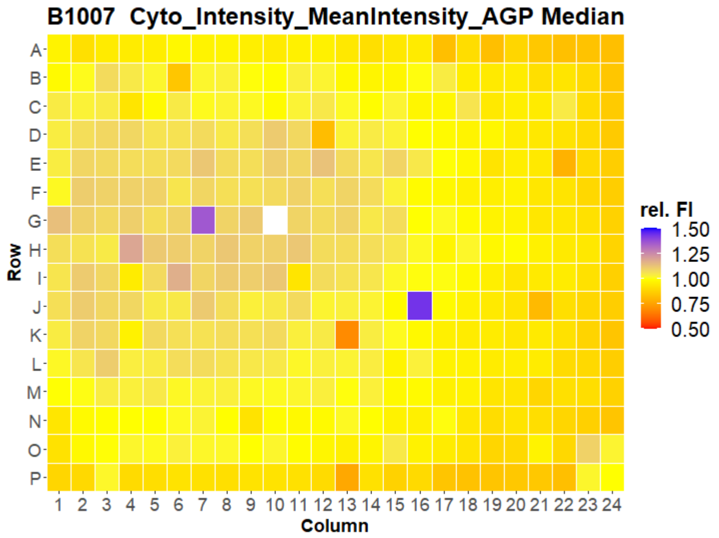
B1007 Cyto_Intensity_MeanIntensity_AGP Median
Row
Column
rel. FI
1.50
1.25
1.00
0.75
0.50

### U2OS FMP

8

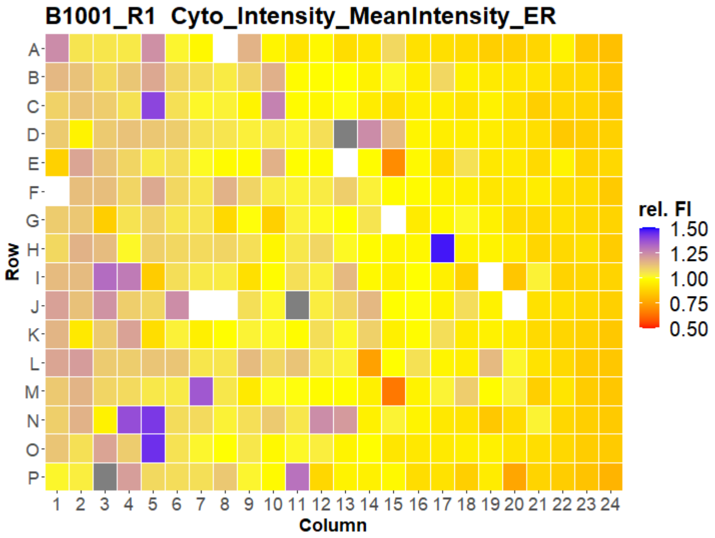
B1001_R1 Cyto_Intensity_MeanIntensity_ER
A B C D E F G H I J K L M N O P
Row
1 2 3 4 5 6 7 8 9 10 11 12 13 14 15 16 17 18 19 20 21 22 23 24
Column
rel. FI
1.50
1.25
1.00
0.75
0.50

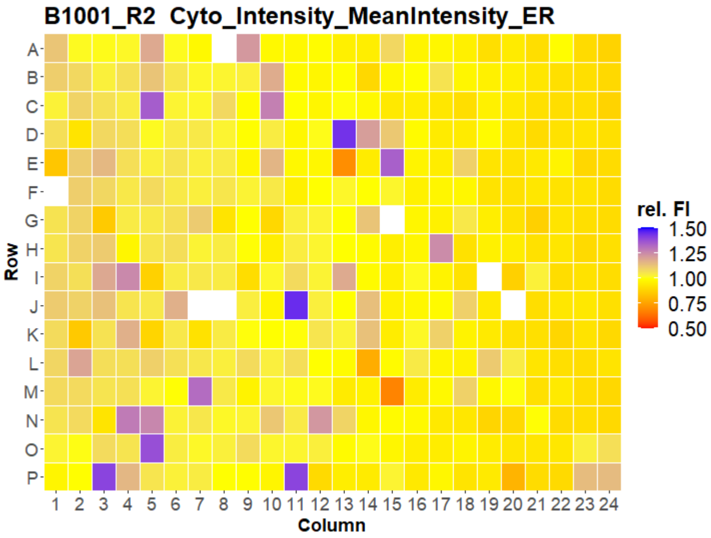
B1001_R2 Cyto_Intensity_MeanIntensity_ER
A B C D E F G H I J K L M N O P
Row
1 2 3 4 5 6 7 8 9 10 11 12 13 14 15 16 17 18 19 20 21 22 23 24
Column
rel. FI
1.50
1.25
1.00
0.75
0.50

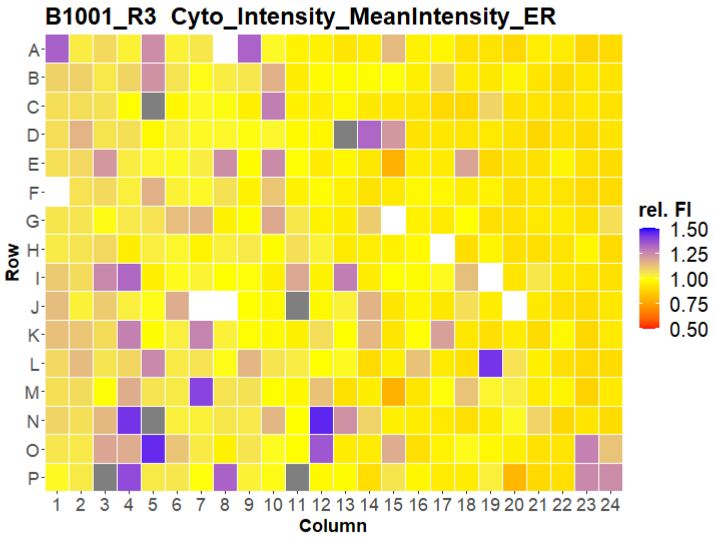
B1001_R3 Cyto_Intensity_MeanIntensity_ER
A B C D E F G H I J K L M N O P
Row
1 2 3 4 5 6 7 8 9 10 11 12 13 14 15 16 17 18 19 20 21 22 23 24
Column
rel. FI
1.50
1.25
1.00
0.75
0.50

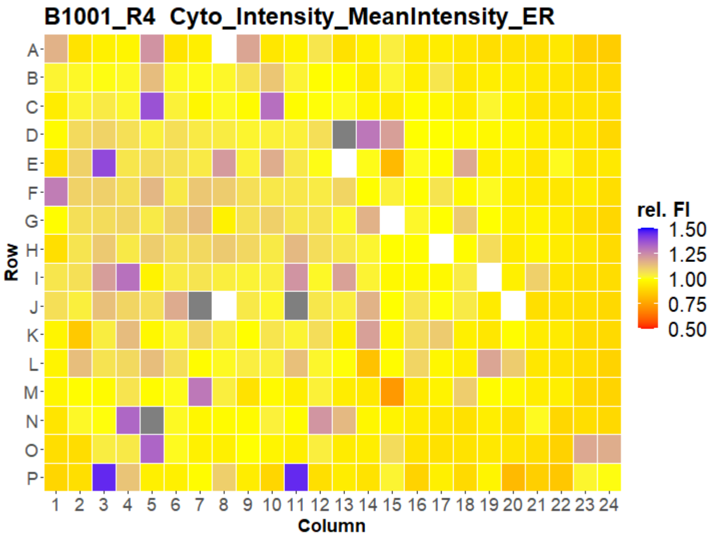
B1001_R4 Cyto_Intensity_MeanIntensity_ER
A B C D E F G H I J K L M N O P
Row
1 2 3 4 5 6 7 8 9 10 11 12 13 14 15 16 17 18 19 20 21 22 23 24
Column
rel. FI
1.50
1.25
1.00
0.75
0.50

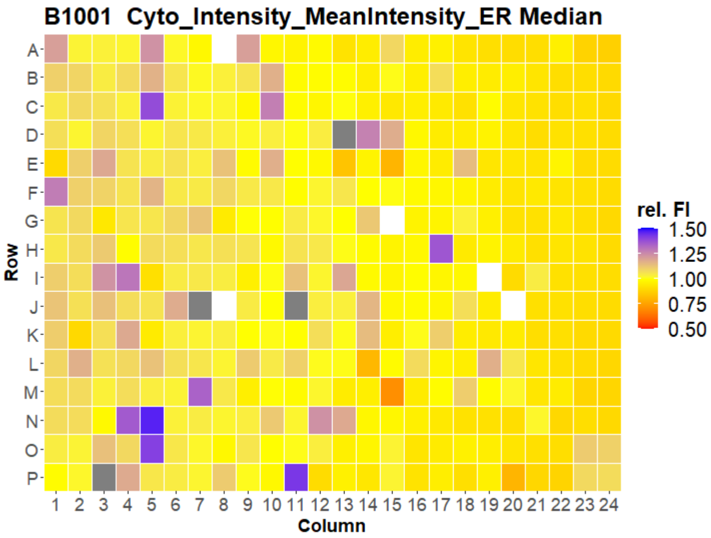
B1001 Cyto_Intensity_MeanIntensity_ER Median
A B C D E F G H I J K L M N O P
Row
1 2 3 4 5 6 7 8 9 10 11 12 13 14 15 16 17 18 19 20 21 22 23 24
Column
rel. FI
1.50
1.25
1.00
0.75
0.50

### U2OS FMP

9

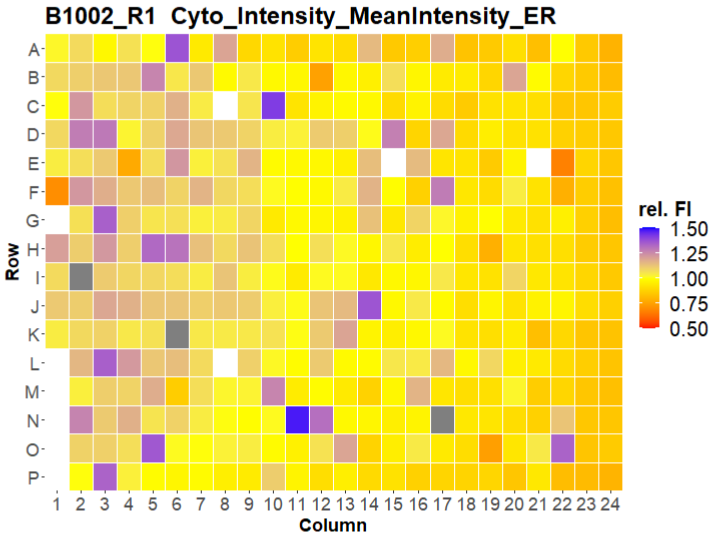


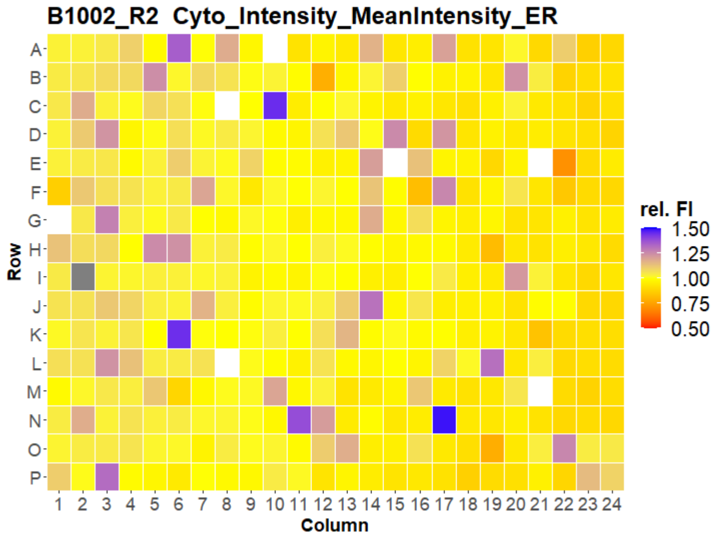


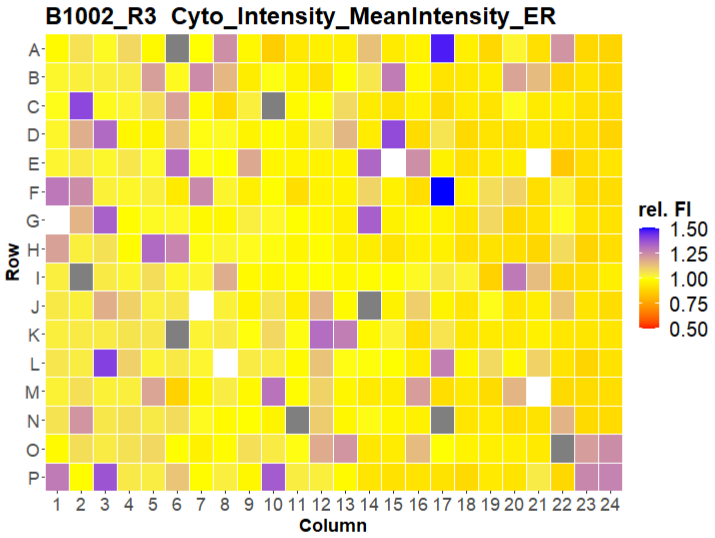


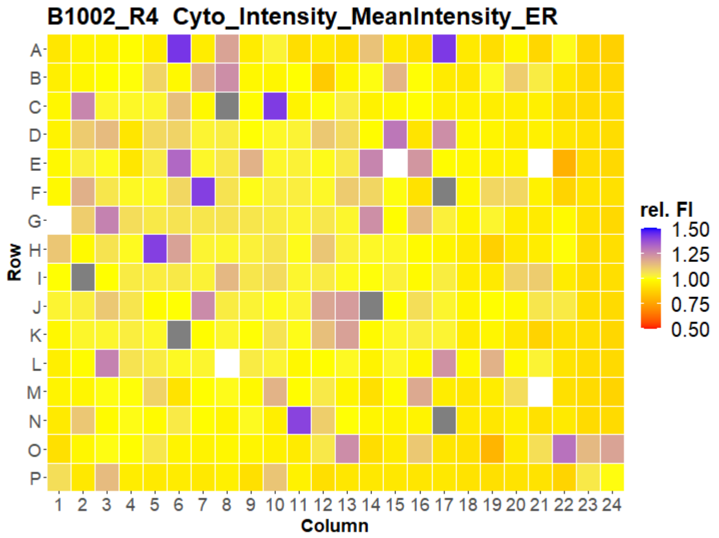


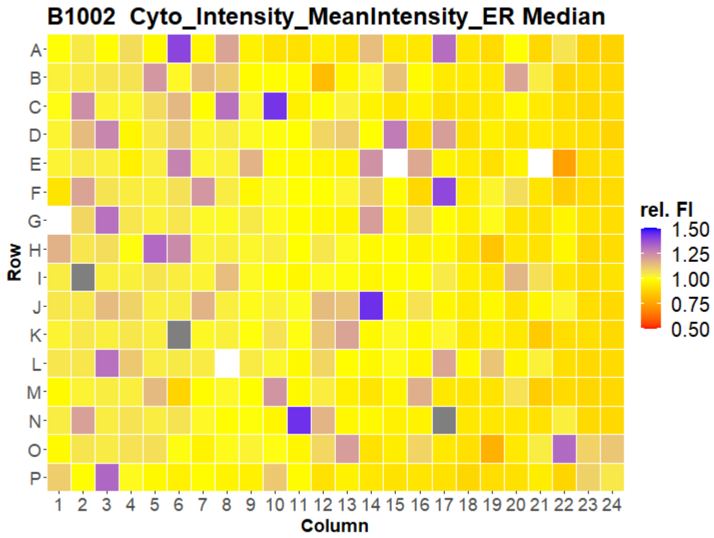



Open for Innovation KNIME

### U2OS FMP



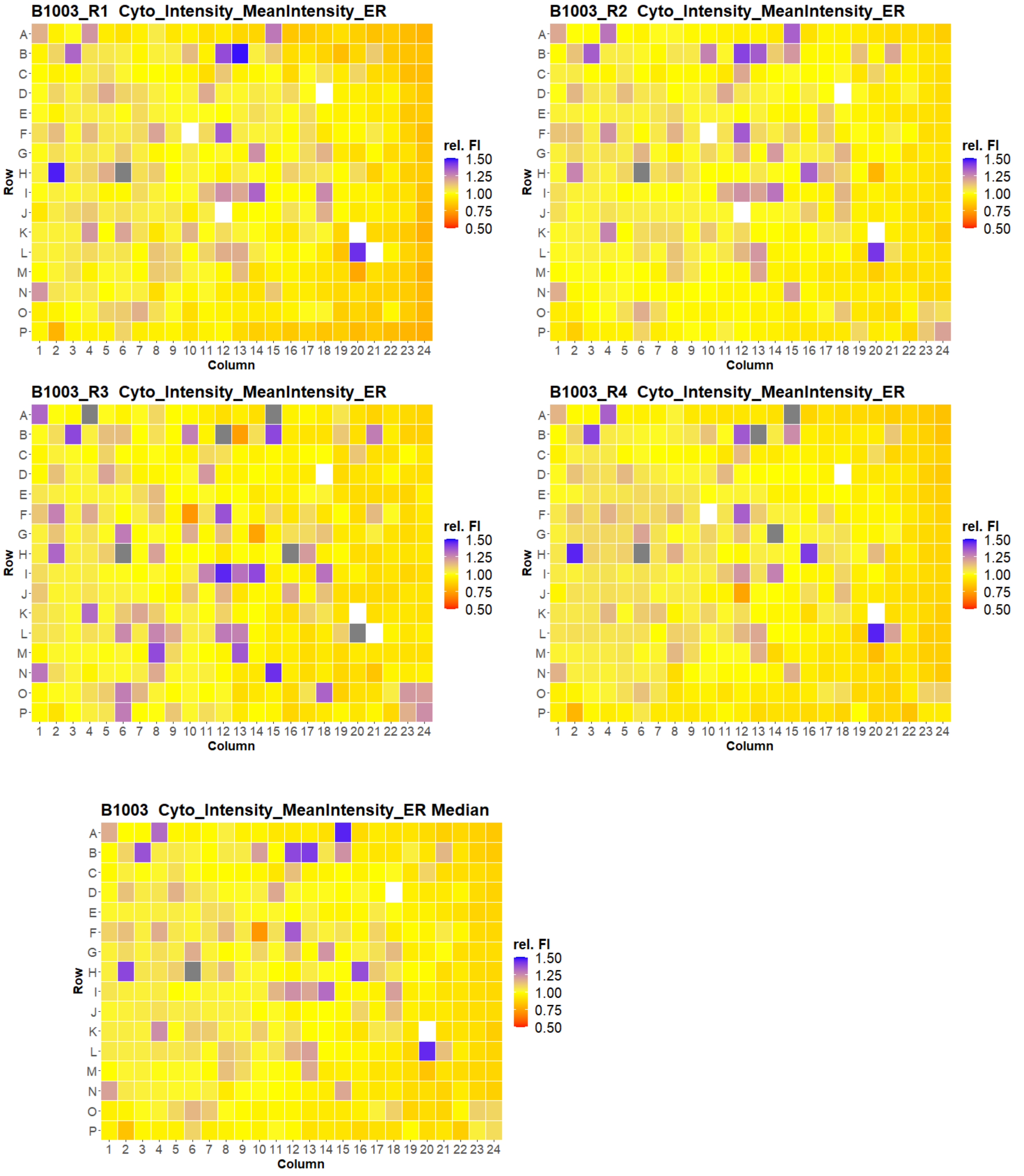
B1003_R1 Cyto_Intensity_MeanIntensity_ER
B1003_R2 Cyto_Intensity_MeanIntensity_ER
B1003_R3 Cyto_Intensity_MeanIntensity_ER
B1003_R4 Cyto_Intensity_MeanIntensity_ER
B1003 Cyto_Intensity_MeanIntensity_ER Median
Row
Column
rel. FI
1.50
1.25
1.00
0.75
0.50

### U2OS FMP

11

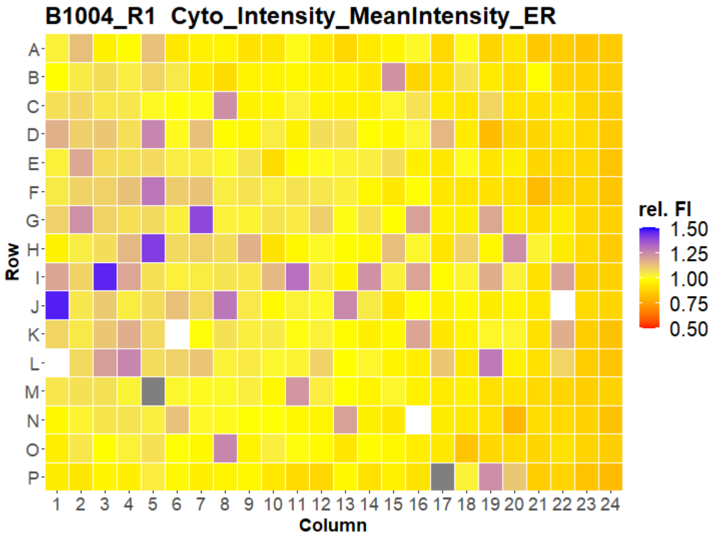
B1004_R1 Cyto_Intensity_MeanIntensity_ER
Row
Column
rel. FI
1.50
1.25
1.00
0.75
0.50

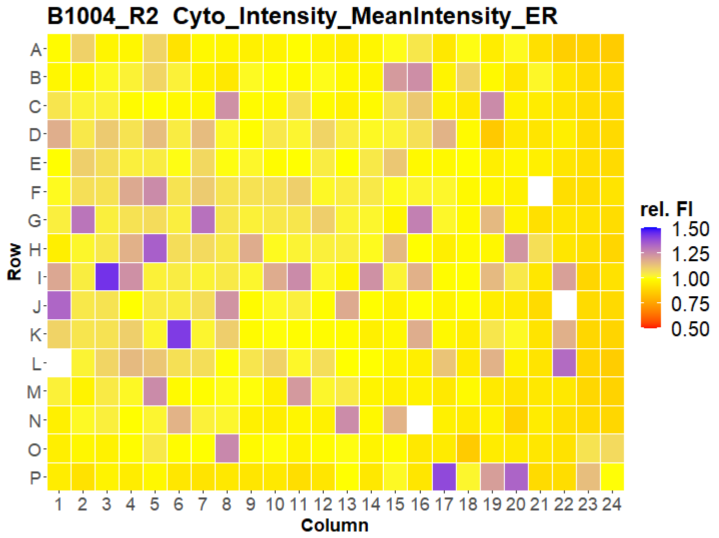
B1004_R2 Cyto_Intensity_MeanIntensity_ER
Row
Column
rel. FI
1.50
1.25
1.00
0.75
0.50

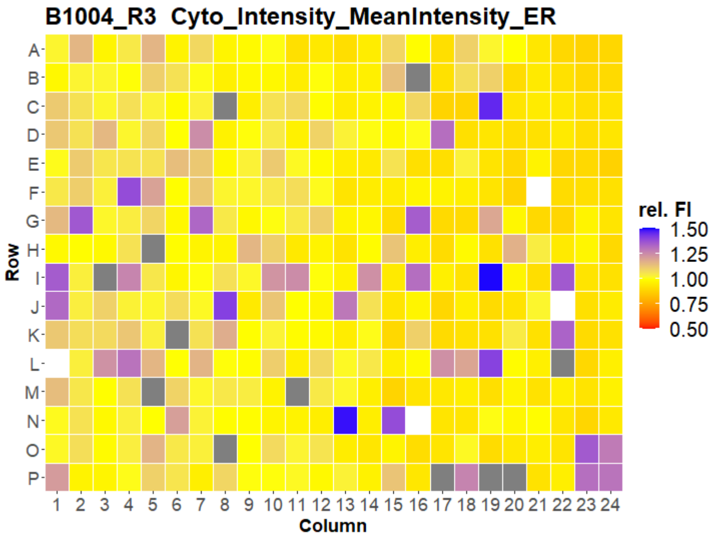
B1004_R3 Cyto_Intensity_MeanIntensity_ER
Row
Column
rel. FI
1.50
1.25
1.00
0.75
0.50

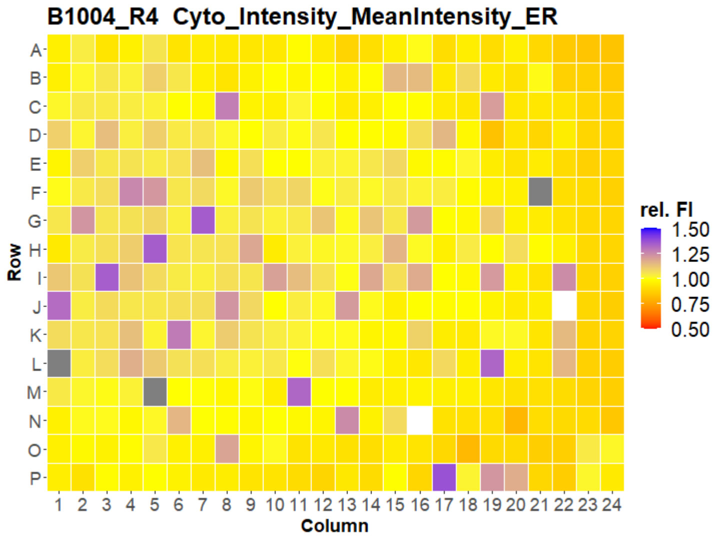
B1004_R4 Cyto_Intensity_MeanIntensity_ER
Row
Column
rel. FI
1.50
1.25
1.00
0.75
0.50

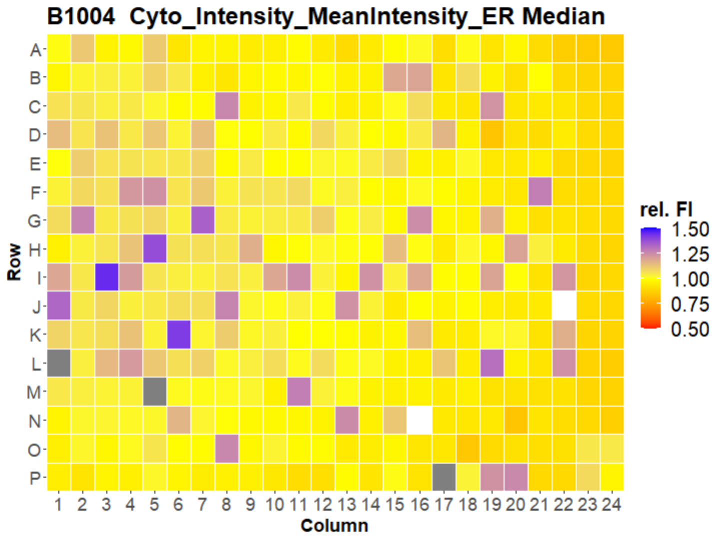
B1004 Cyto_Intensity_MeanIntensity_ER Median
Row
Column
rel. FI
1.50
1.25
1.00
0.75
0.50

### U2OS FMP



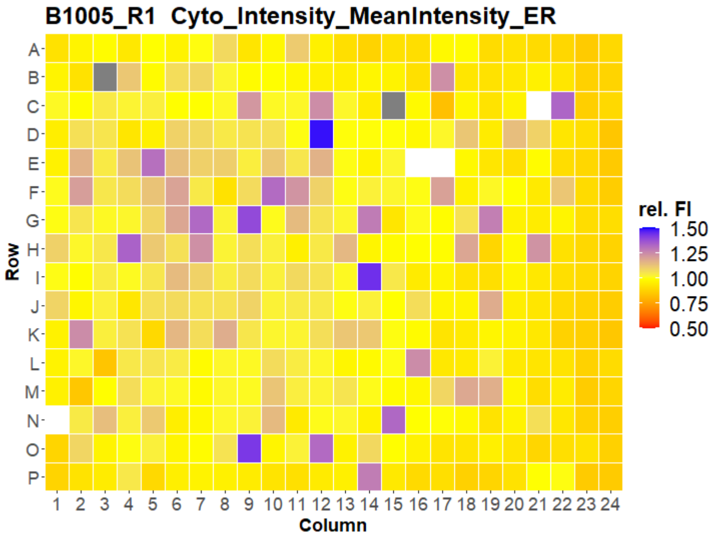
B1005_R1 Cyto_Intensity_MeanIntensity_ER
Row
A B C D E F G H I J K L M N O P
1 2 3 4 5 6 7 8 9 10 11 12 13 14 15 16 17 18 19 20 21 22 23 24
Column
rel. FI
1.50
1.25
1.00
0.75
0.50

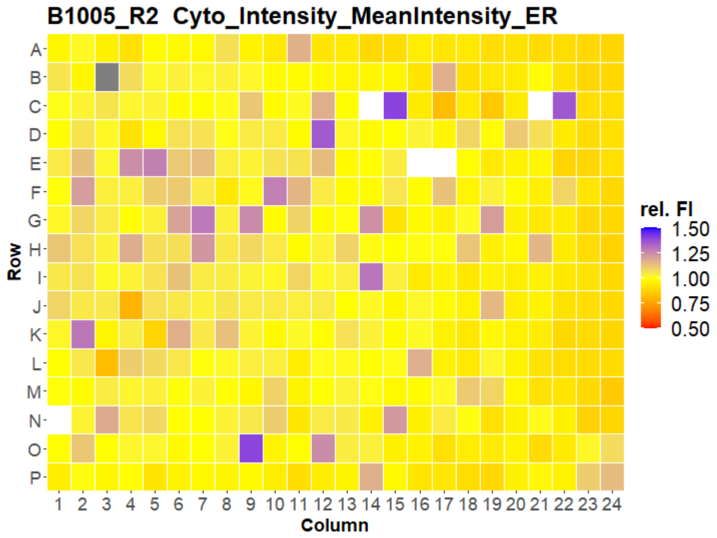
B1005_R2 Cyto_Intensity_MeanIntensity_ER
Row
A B C D E F G H I J K L M N O P
1 2 3 4 5 6 7 8 9 10 11 12 13 14 15 16 17 18 19 20 21 22 23 24
Column
rel. FI
1.50
1.25
1.00
0.75
0.50

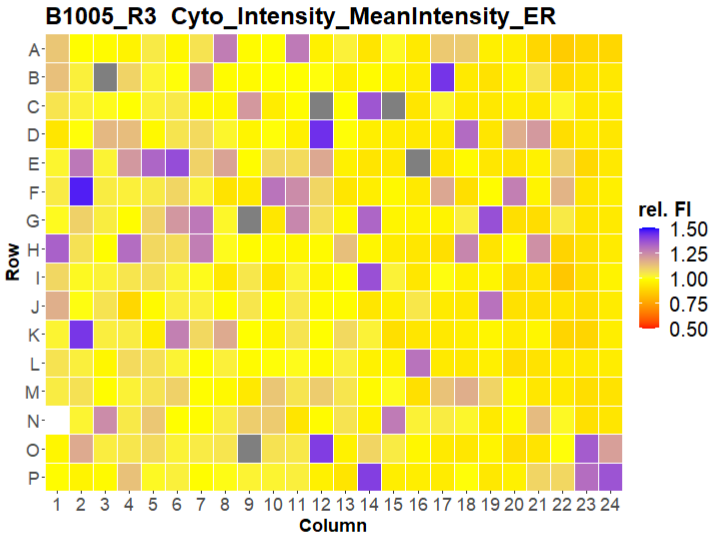
B1005_R3 Cyto_Intensity_MeanIntensity_ER
Row
A B C D E F G H I J K L M N O P
1 2 3 4 5 6 7 8 9 10 11 12 13 14 15 16 17 18 19 20 21 22 23 24
Column
rel. FI
1.50
1.25
1.00
0.75
0.50

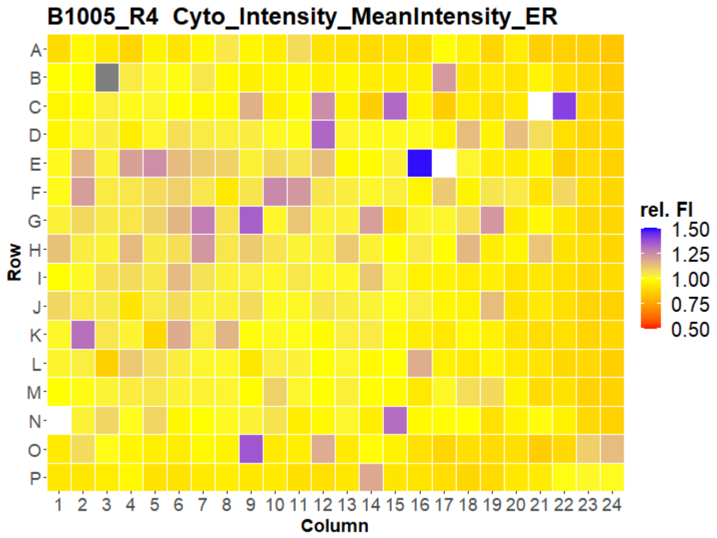
B1005_R4 Cyto_Intensity_MeanIntensity_ER
Row
A B C D E F G H I J K L M N O P
1 2 3 4 5 6 7 8 9 10 11 12 13 14 15 16 17 18 19 20 21 22 23 24
Column
rel. FI
1.50
1.25
1.00
0.75
0.50

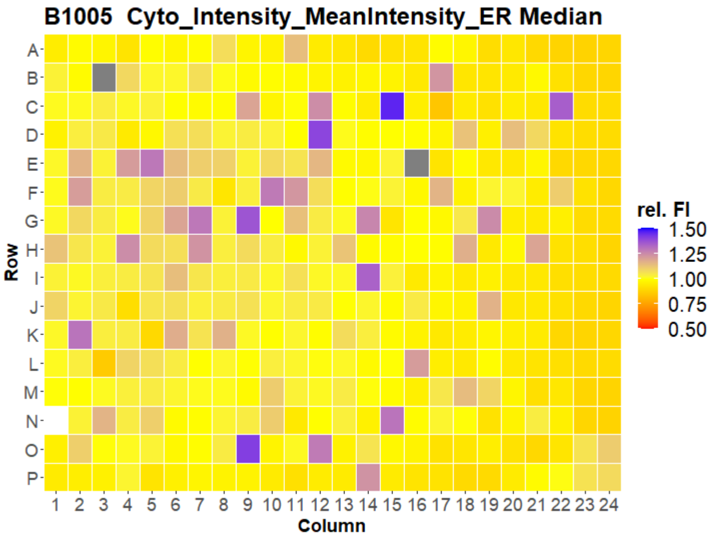
B1005 Cyto_Intensity_MeanIntensity_ER Median
Row
A B C D E F G H I J K L M N O P
1 2 3 4 5 6 7 8 9 10 11 12 13 14 15 16 17 18 19 20 21 22 23 24
Column
rel. FI
1.50
1.25
1.00
0.75
0.50

### U2OS FMP

13

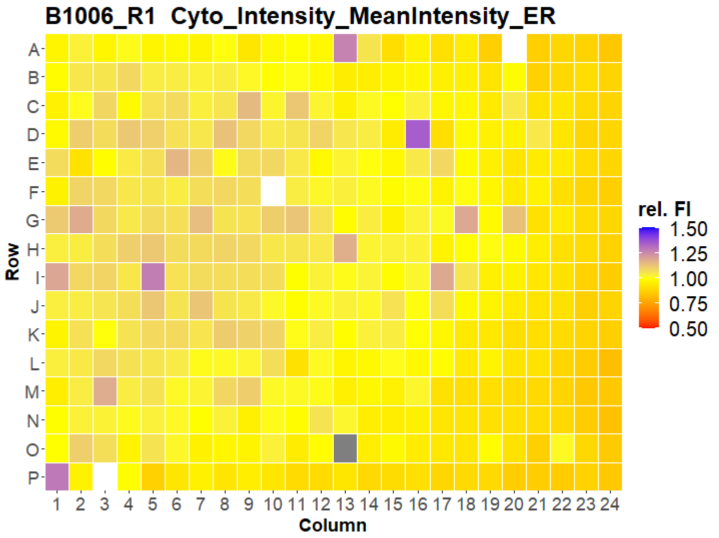
B1006_R1 Cyto_Intensity_MeanIntensity_ER
Row
Column
rel. FI
1.50
1.25
1.00
0.75
0.50

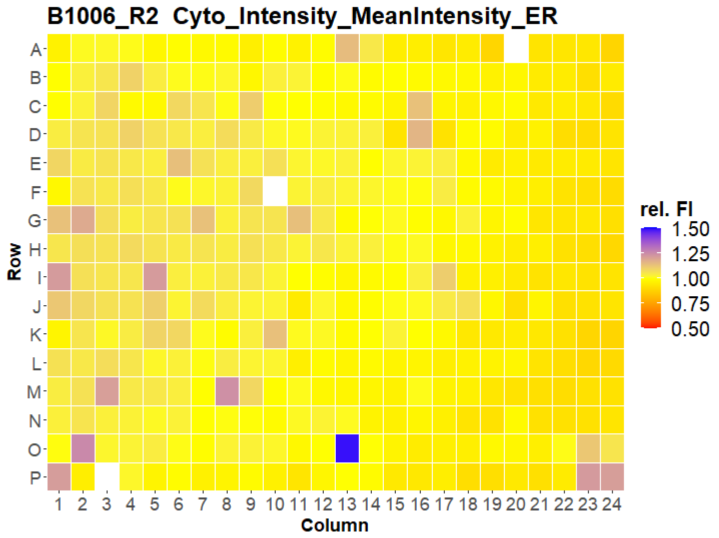
B1006_R2 Cyto_Intensity_MeanIntensity_ER
Row
Column
rel. FI
1.50
1.25
1.00
0.75
0.50

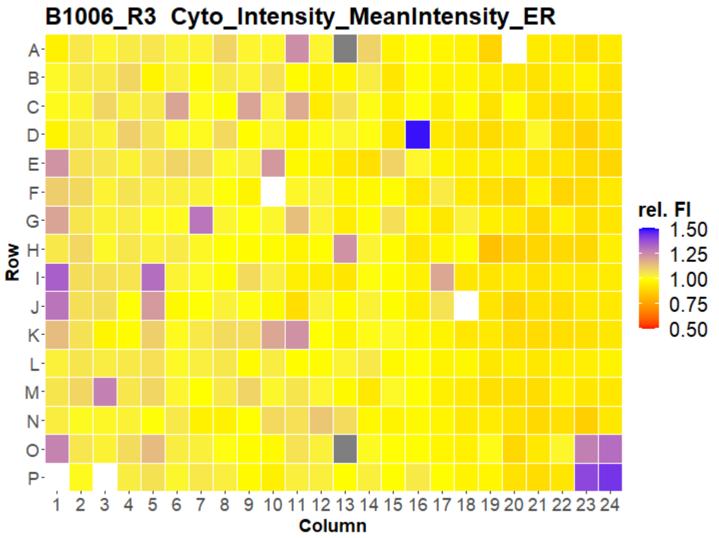
B1006_R3 Cyto_Intensity_MeanIntensity_ER
Row
Column
rel. FI
1.50
1.25
1.00
0.75
0.50

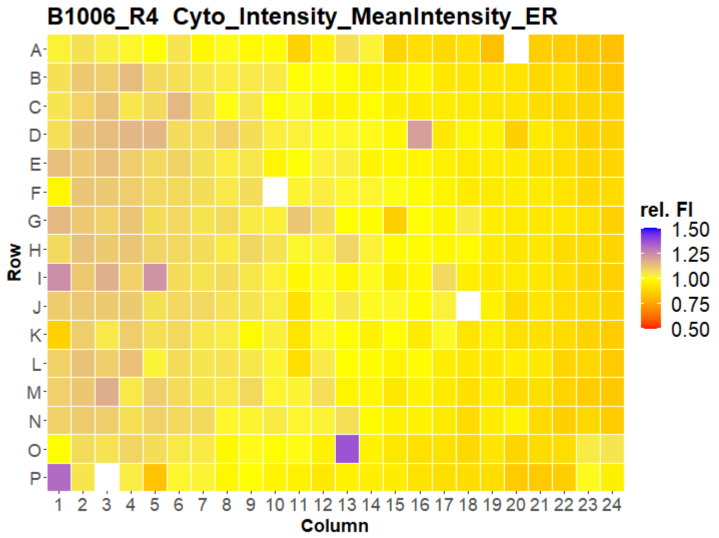
B1006_R4 Cyto_Intensity_MeanIntensity_ER
Row
Column
rel. FI
1.50
1.25
1.00
0.75
0.50

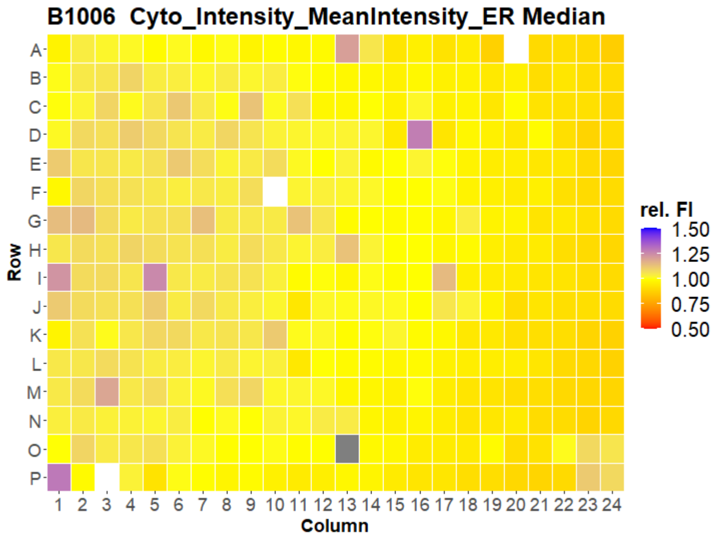
B1006 Cyto_Intensity_MeanIntensity_ER Median
Row
Column
rel. FI
1.50
1.25
1.00
0.75
0.50

### U2OS FMP

14

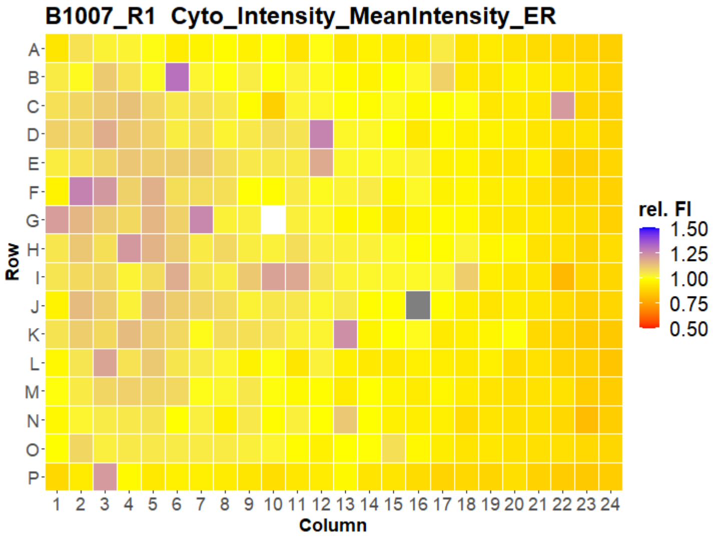
B1007_R1 Cyto_Intensity_MeanIntensity_ER
Row
Column
rel. FI
1.50
1.25
1.00
0.75
0.50

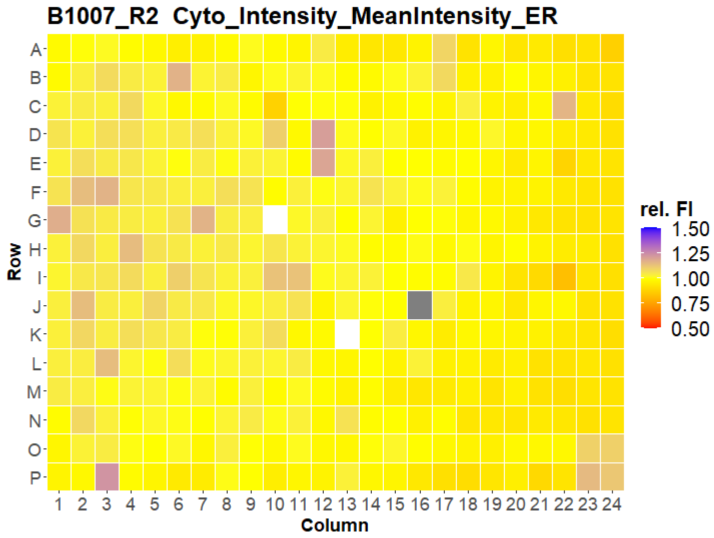
B1007_R2 Cyto_Intensity_MeanIntensity_ER
Row
Column
rel. FI
1.50
1.25
1.00
0.75
0.50

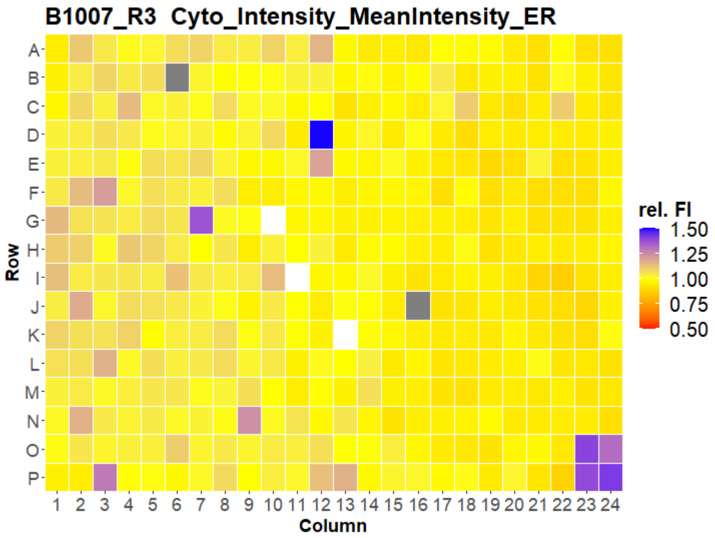
B1007_R3 Cyto_Intensity_MeanIntensity_ER
Row
Column
rel. FI
1.50
1.25
1.00
0.75
0.50

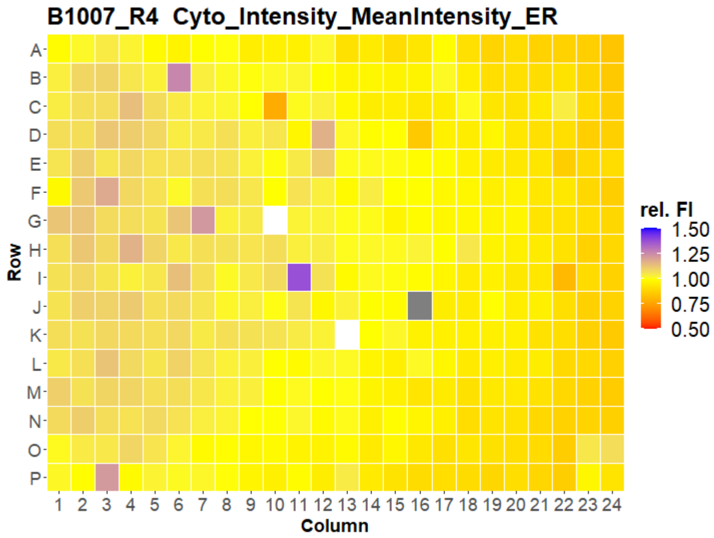
B1007_R4 Cyto_Intensity_MeanIntensity_ER
Row
Column
rel. FI
1.50
1.25
1.00
0.75
0.50

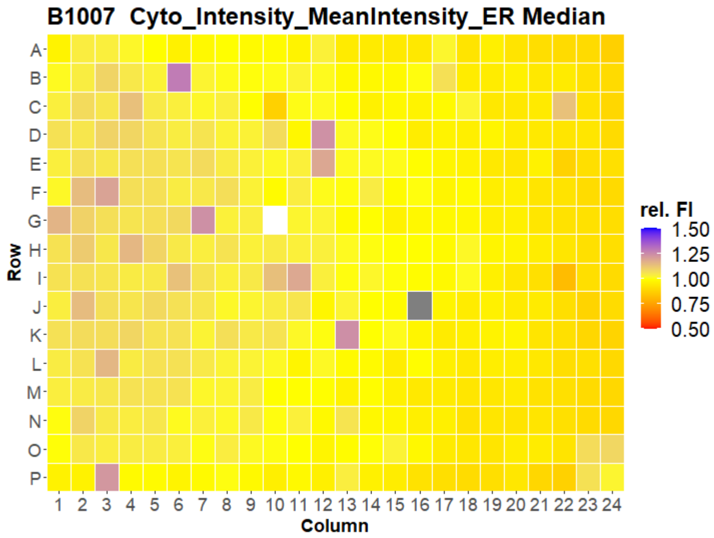
B1007 Cyto_Intensity_MeanIntensity_ER Median
Row
Column
rel. FI
1.50
1.25
1.00
0.75
0.50

### U2OS FMP



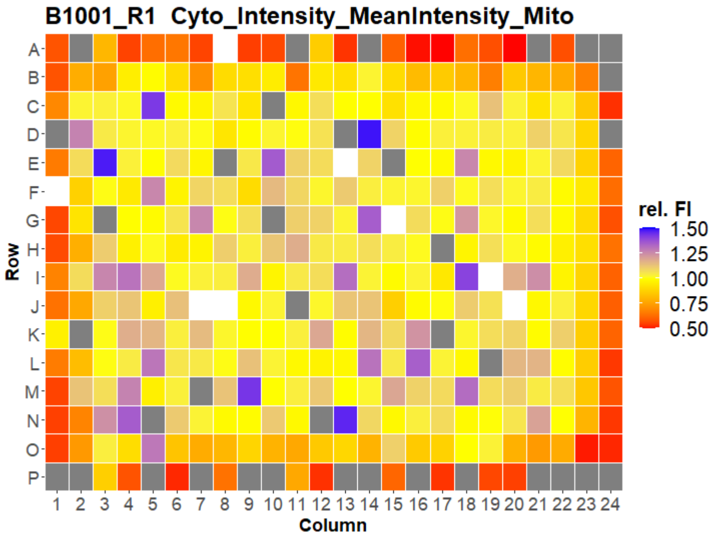
B1001_R1 Cyto_Intensity_MeanIntensity_Mito
Row
Column
rel. FI
1.50
1.25
1.00
0.75
0.50

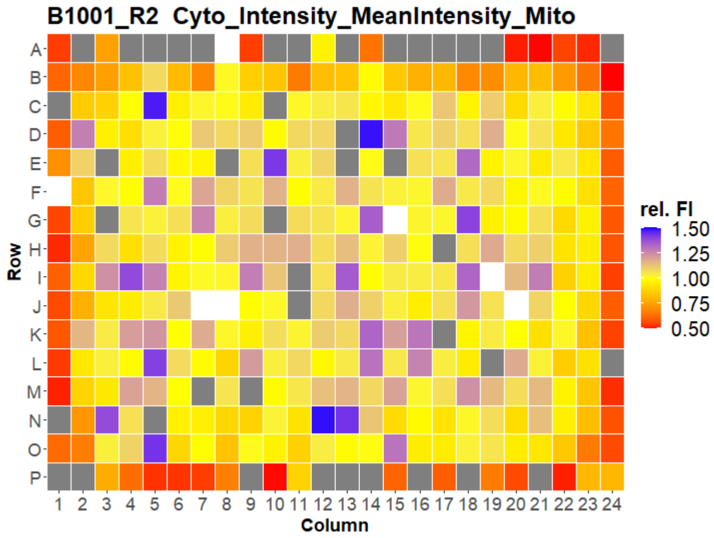
B1001_R2 Cyto_Intensity_MeanIntensity_Mito
Row
Column
rel. FI
1.50
1.25
1.00
0.75
0.50

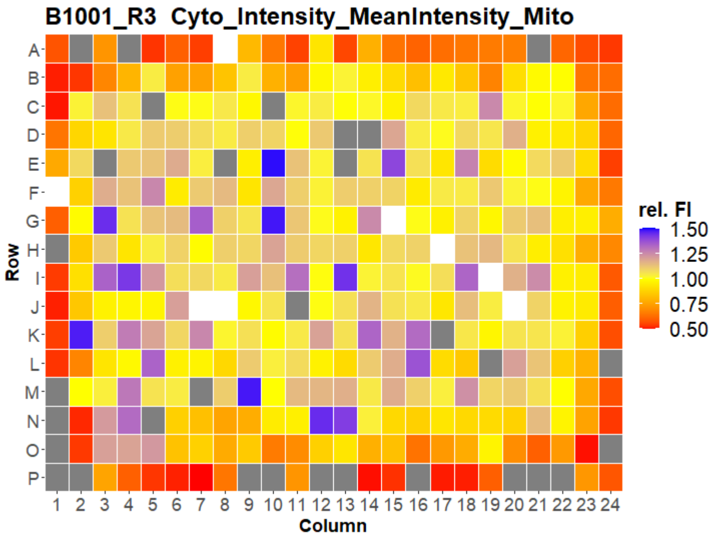
B1001_R3 Cyto_Intensity_MeanIntensity_Mito
Row
Column
rel. FI
1.50
1.25
1.00
0.75
0.50

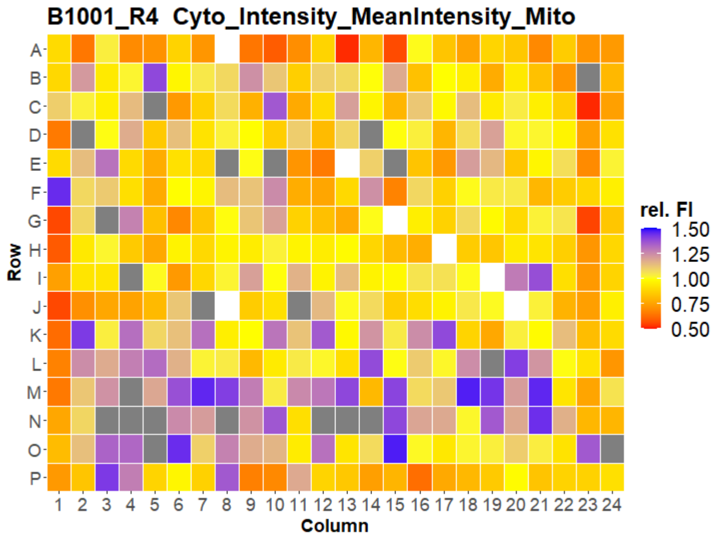
B1001_R4 Cyto_Intensity_MeanIntensity_Mito
Row
Column
rel. FI
1.50
1.25
1.00
0.75
0.50

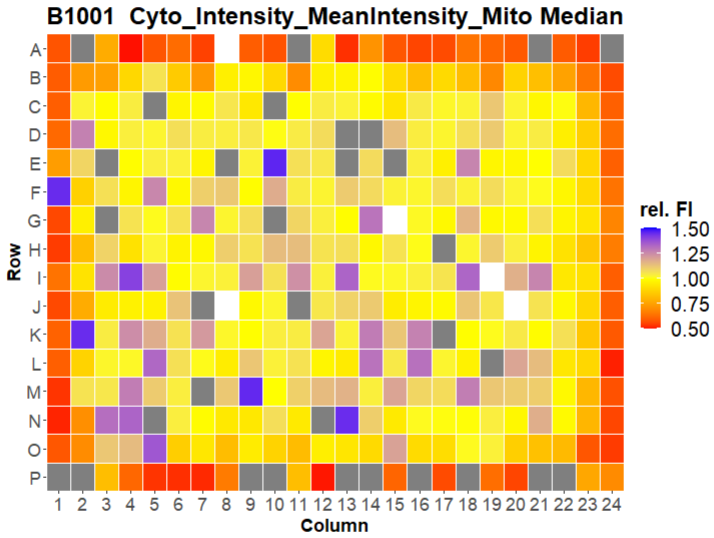
B1001 Cyto_Intensity_MeanIntensity_Mito Median
Row
Column
rel. FI
1.50
1.25
1.00
0.75
0.50

### U2OS FMP

16

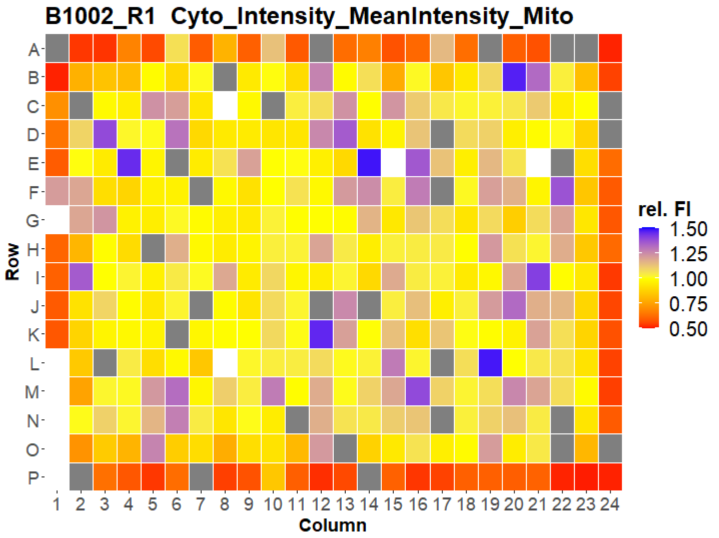
B1002_R1 Cyto_Intensity_MeanIntensity_Mito
Row
Column
rel. FI
1.50
1.25
1.00
0.75
0.50

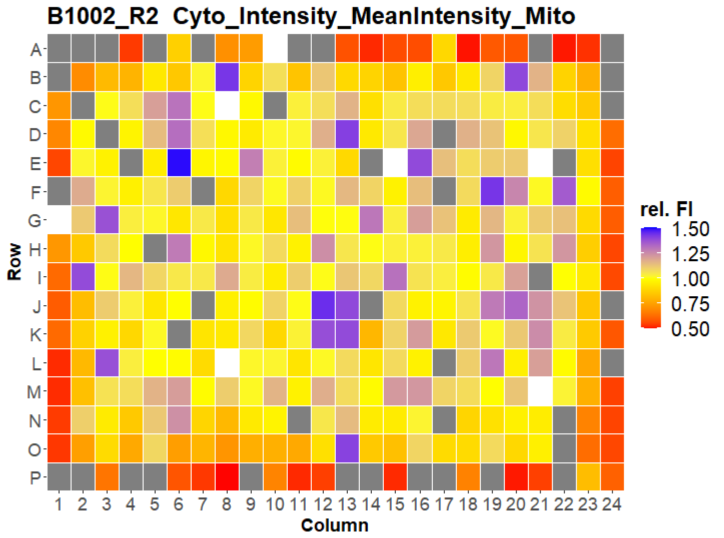
B1002_R2 Cyto_Intensity_MeanIntensity_Mito
Row
Column
rel. FI
1.50
1.25
1.00
0.75
0.50

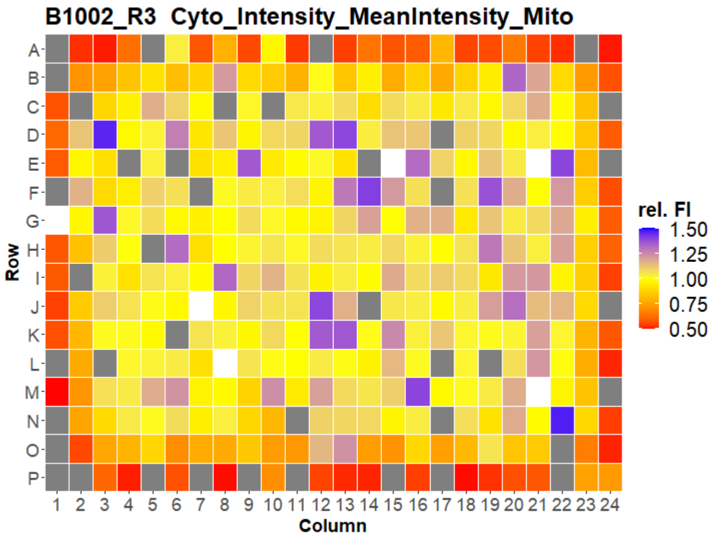
B1002_R3 Cyto_Intensity_MeanIntensity_Mito
Row
Column
rel. FI
1.50
1.25
1.00
0.75
0.50

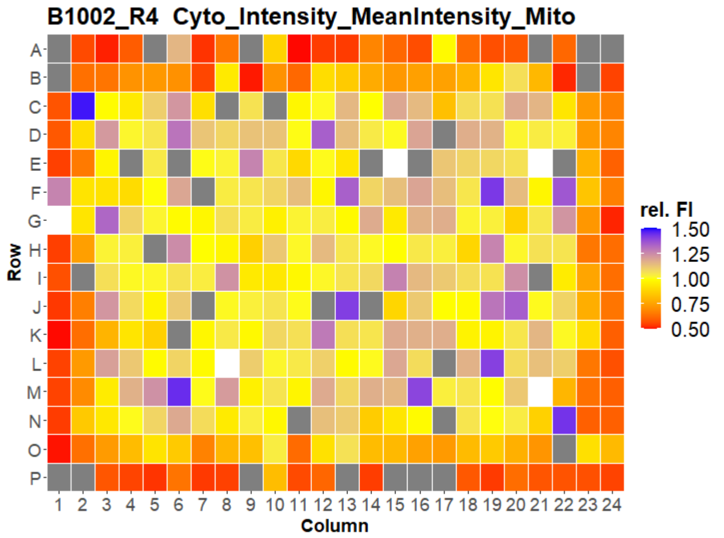
B1002_R4 Cyto_Intensity_MeanIntensity_Mito
Row
Column
rel. FI
1.50
1.25
1.00
0.75
0.50

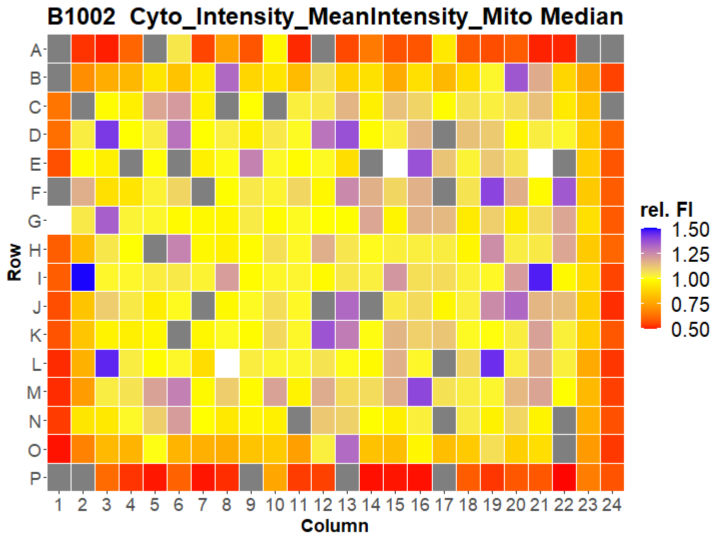
B1002 Cyto_Intensity_MeanIntensity_Mito Median
Row
Column
rel. FI
1.50
1.25
1.00
0.75
0.50

### U2OS FMP

17

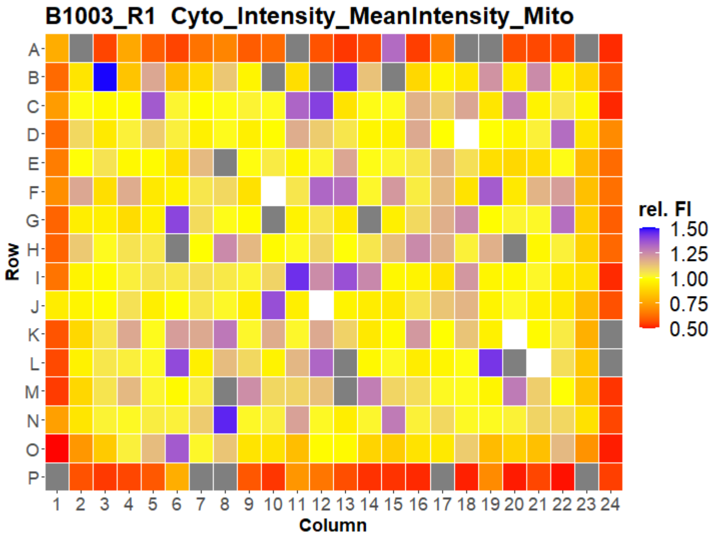


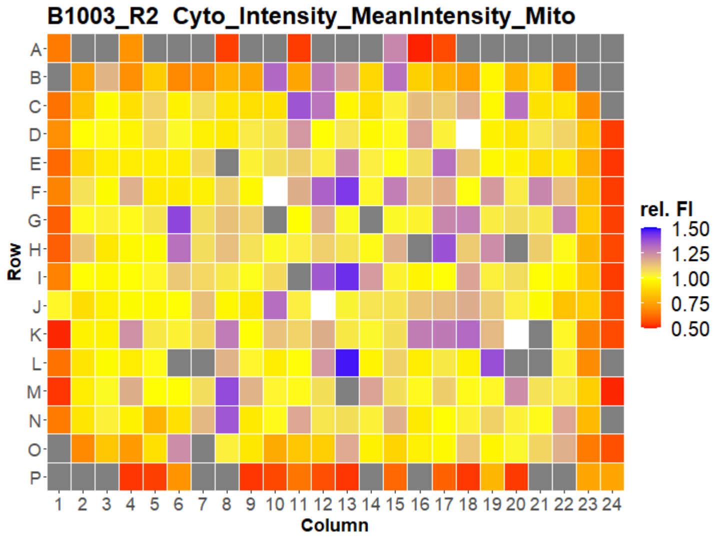


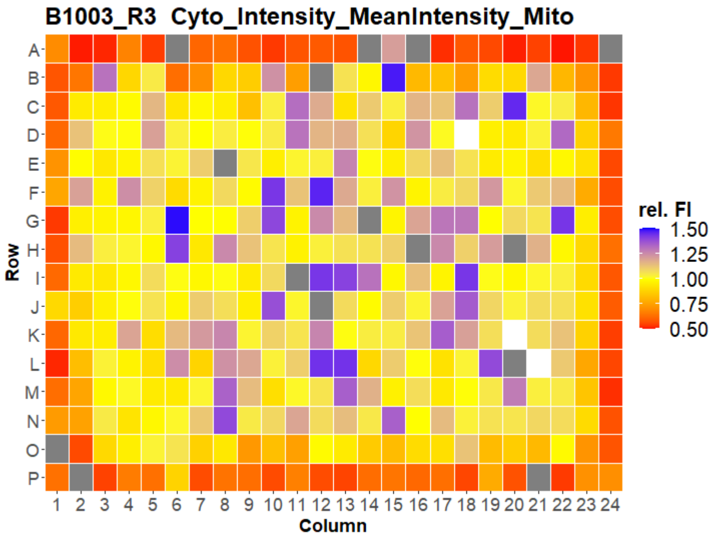


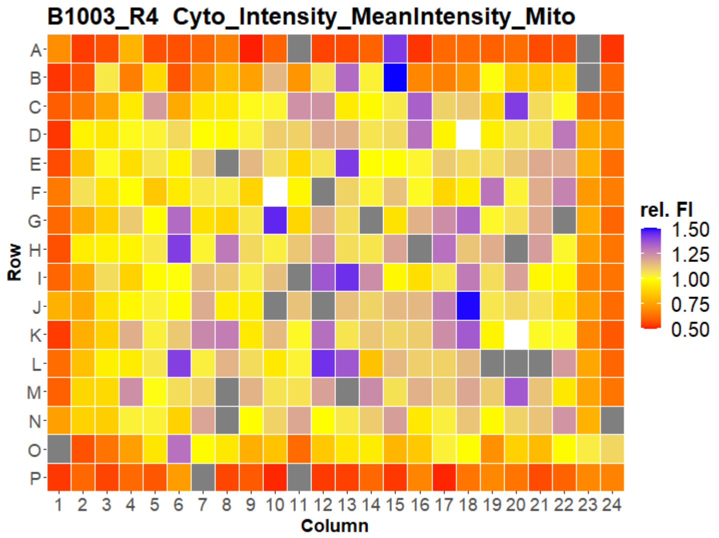


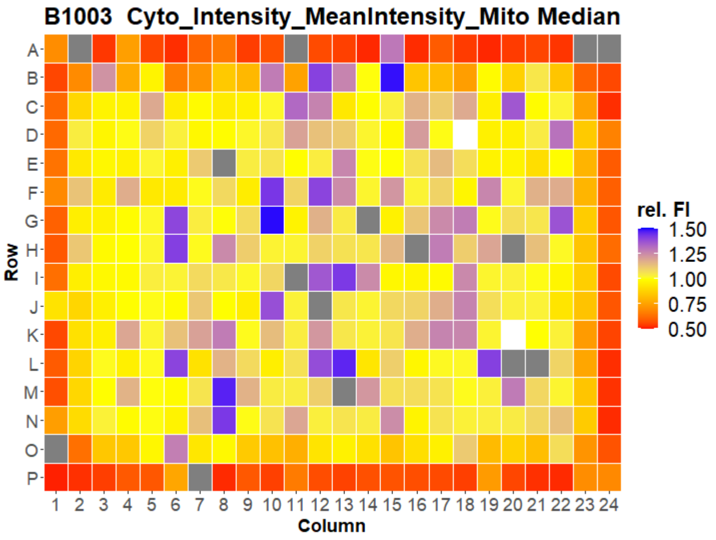



Open for Innovation KNIME

### U2OS FMP

18

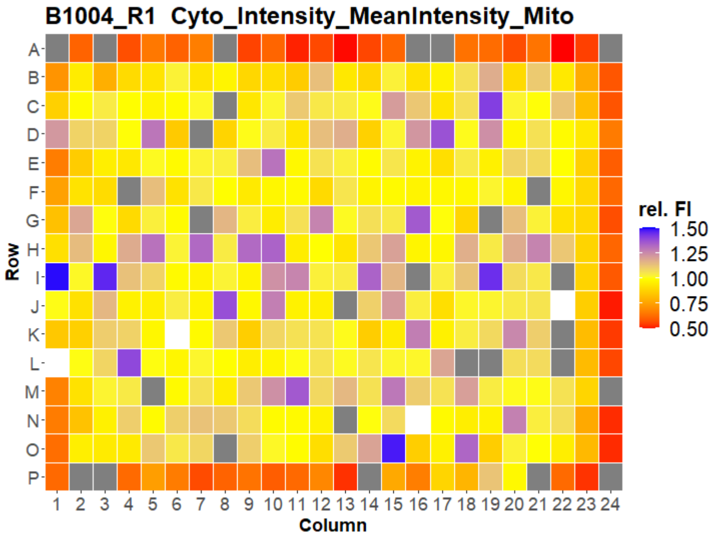
B1004_R1 Cyto_Intensity_MeanIntensity_Mito
Row
Column
rel. FI
1.50
1.25
1.00
0.75
0.50

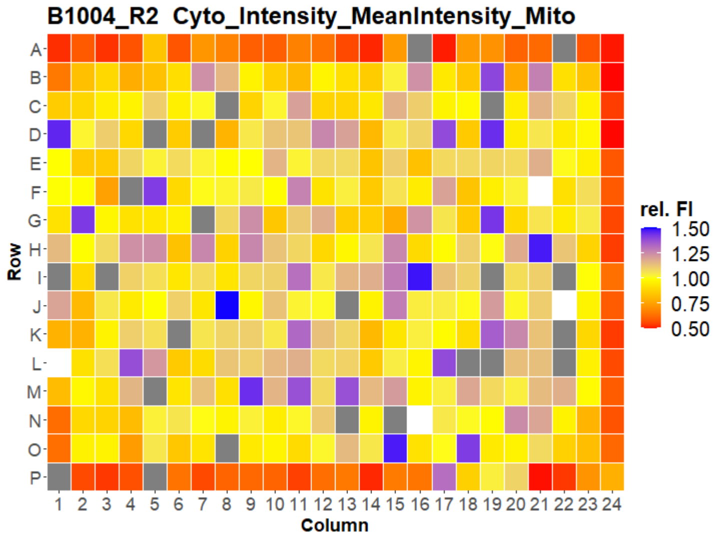
B1004_R2 Cyto_Intensity_MeanIntensity_Mito
Row
Column
rel. FI
1.50
1.25
1.00
0.75
0.50

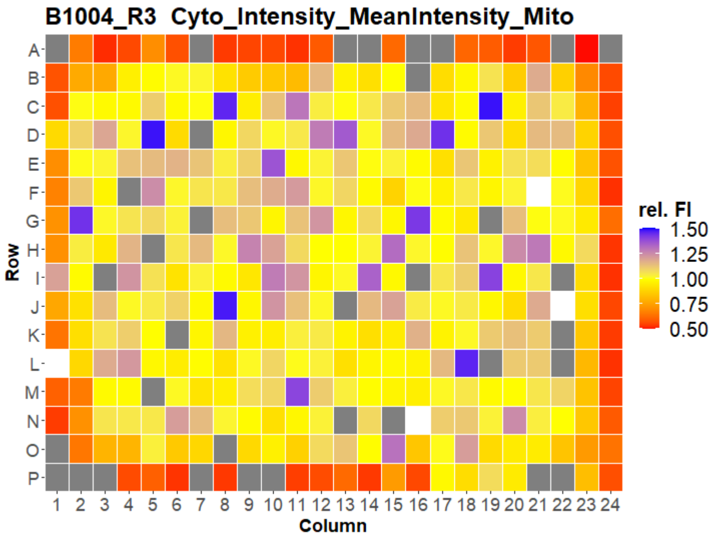
B1004_R3 Cyto_Intensity_MeanIntensity_Mito
Row
Column
rel. FI
1.50
1.25
1.00
0.75
0.50

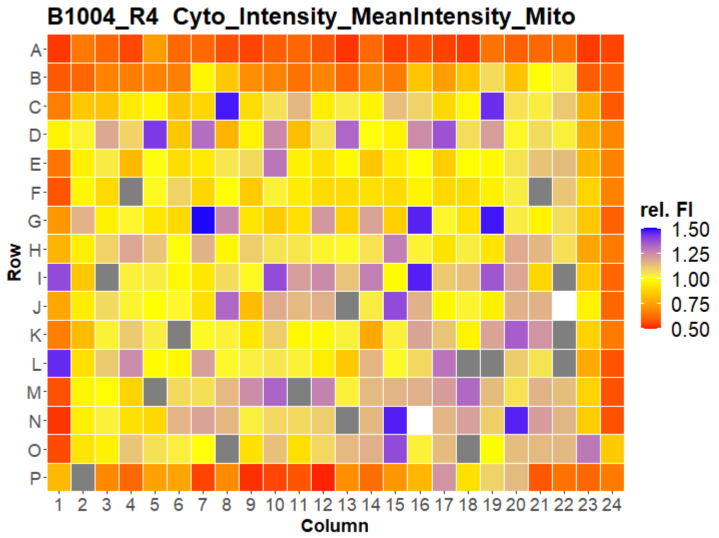
B1004_R4 Cyto_Intensity_MeanIntensity_Mito
Row
Column
rel. FI
1.50
1.25
1.00
0.75
0.50

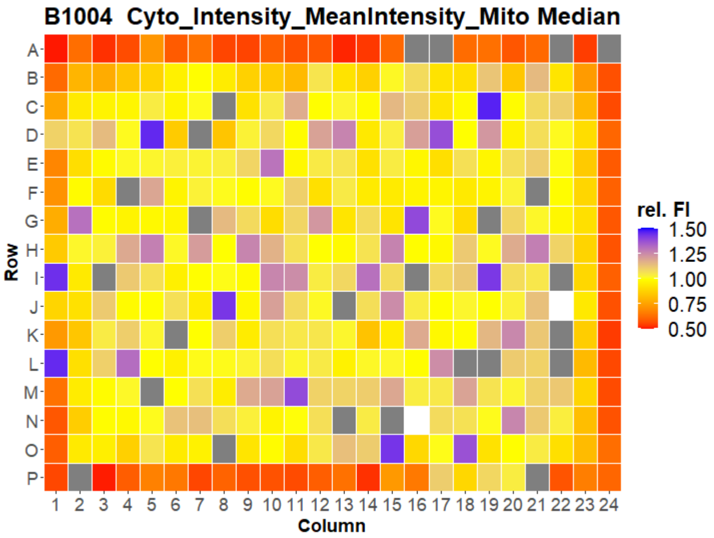
B1004 Cyto_Intensity_MeanIntensity_Mito Median
Row
Column
rel. FI
1.50
1.25
1.00
0.75
0.50

### U2OS FMP

19

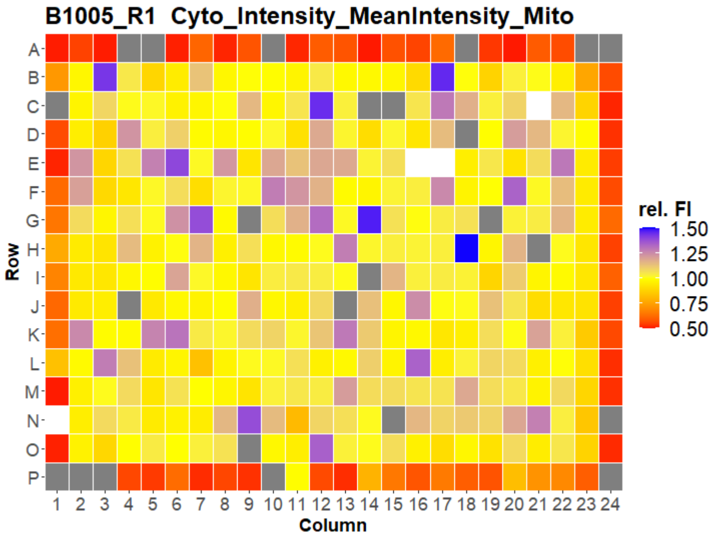
B1005_R1 Cyto_Intensity_MeanIntensity_Mito
Row
Column
rel. FI
1.50
1.25
1.00
0.75
0.50

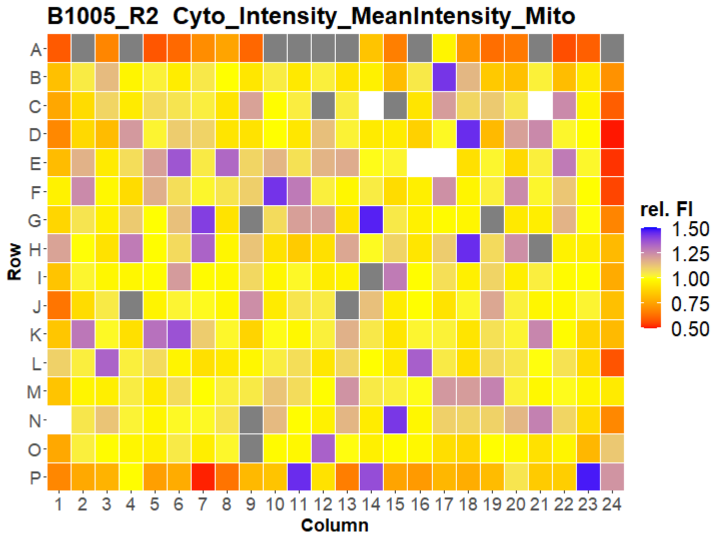
B1005_R2 Cyto_Intensity_MeanIntensity_Mito
Row
Column
rel. FI
1.50
1.25
1.00
0.75
0.50

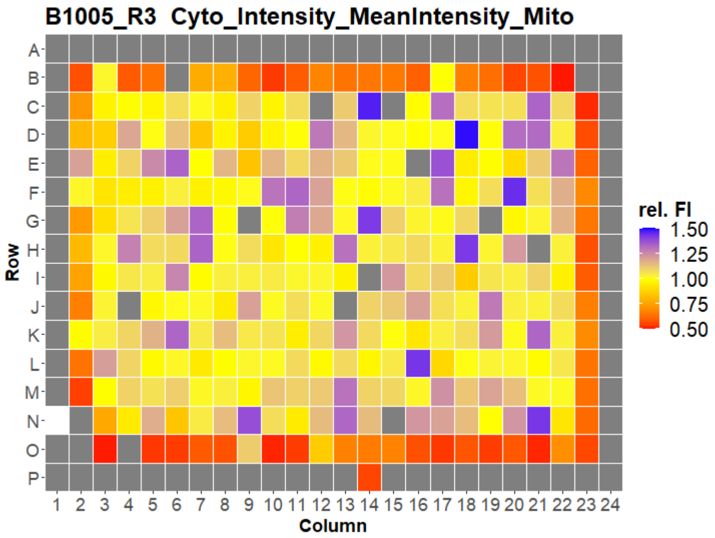
B1005_R3 Cyto_Intensity_MeanIntensity_Mito
Row
Column
rel. FI
1.50
1.25
1.00
0.75
0.50

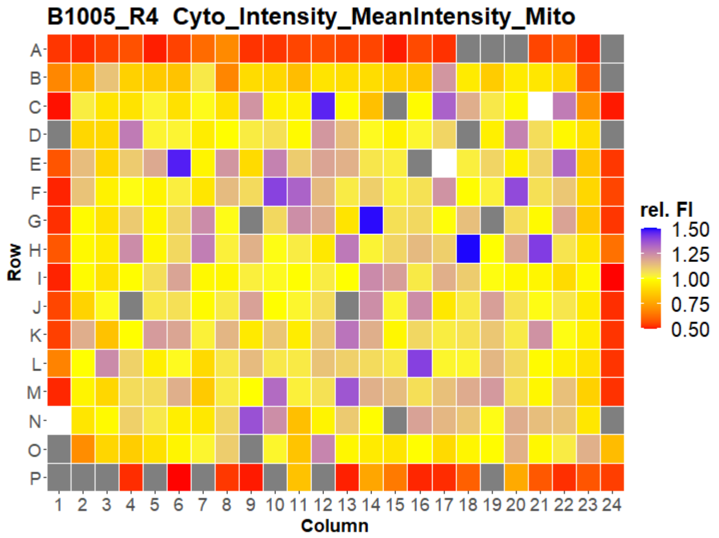
B1005_R4 Cyto_Intensity_MeanIntensity_Mito
Row
Column
rel. FI
1.50
1.25
1.00
0.75
0.50

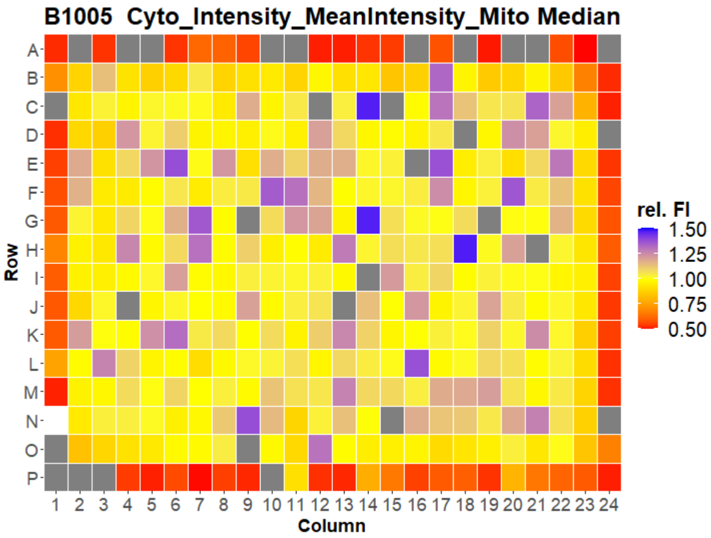
B1005 Cyto_Intensity_MeanIntensity_Mito Median
Row
Column
rel. FI
1.50
1.25
1.00
0.75
0.50

### U2OS FMP

20

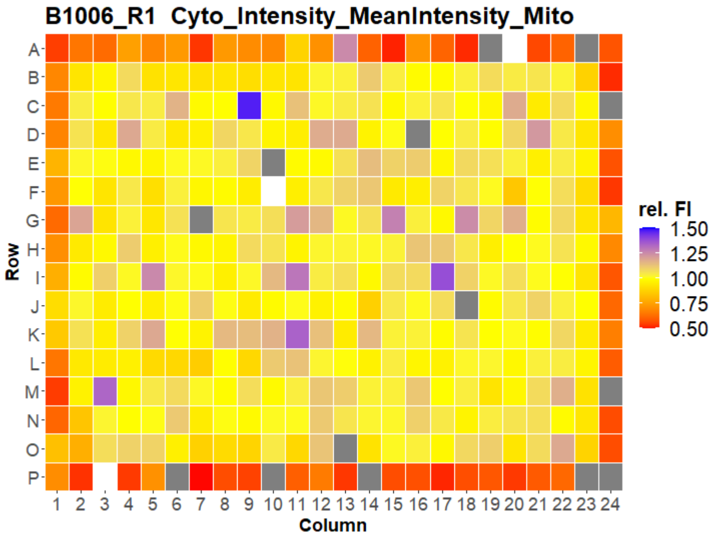


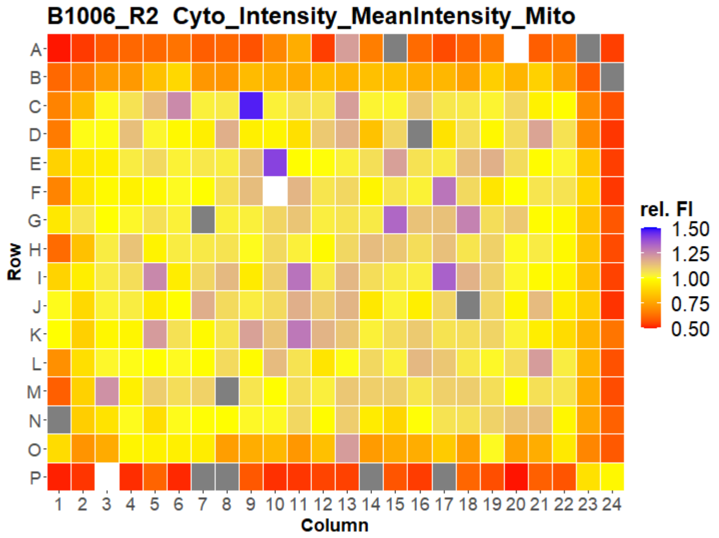


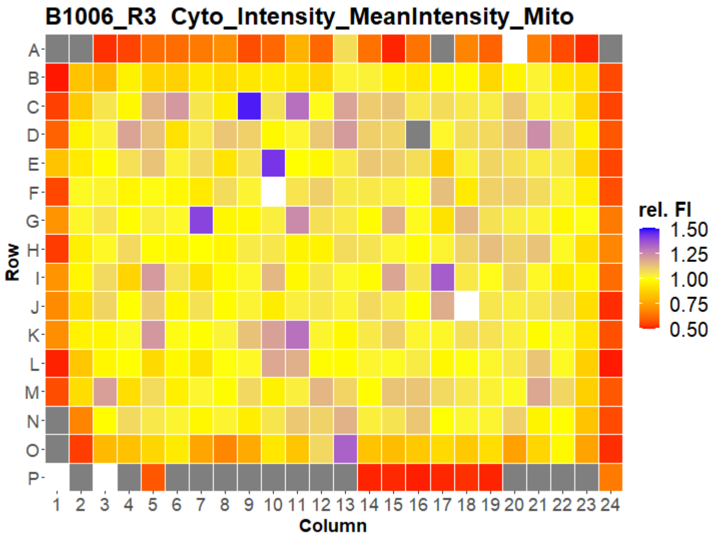


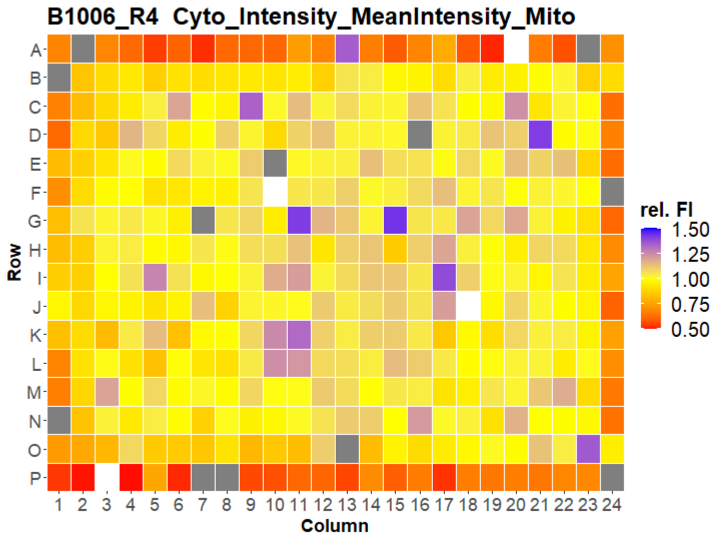


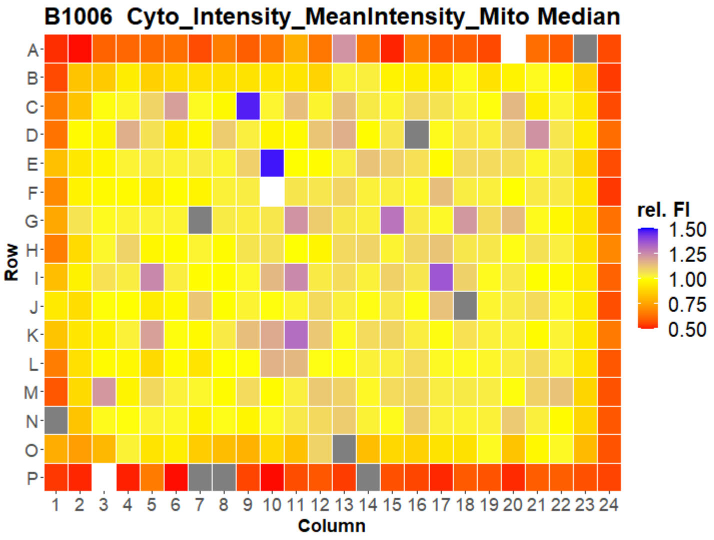



Open for Innovation KNIME

### U2OS FMP

21

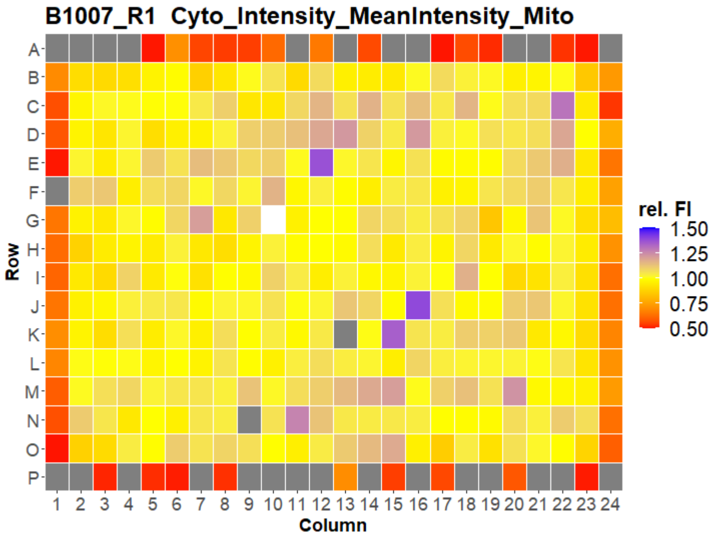
B1007_R1 Cyto_Intensity_MeanIntensity_Mito
Row
A B C D E F G H I J K L M N O P
Column
1 2 3 4 5 6 7 8 9 10 11 12 13 14 15 16 17 18 19 20 21 22 23 24
rel. FI
1.50
1.25
1.00
0.75
0.50

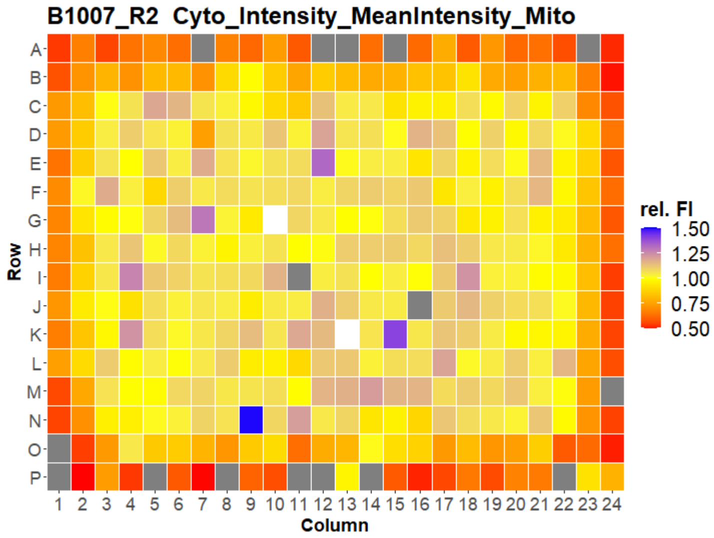
B1007_R2 Cyto_Intensity_MeanIntensity_Mito
Row
A B C D E F G H I J K L M N O P
Column
1 2 3 4 5 6 7 8 9 10 11 12 13 14 15 16 17 18 19 20 21 22 23 24
rel. FI
1.50
1.25
1.00
0.75
0.50

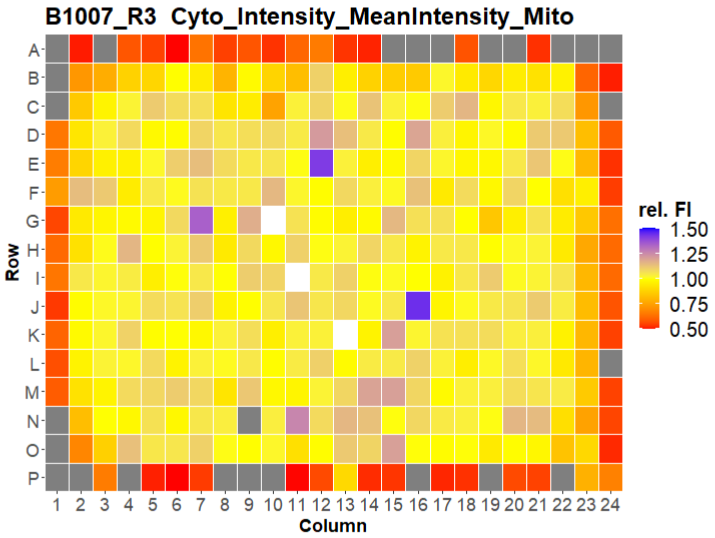
B1007_R3 Cyto_Intensity_MeanIntensity_Mito
Row
A B C D E F G H I J K L M N O P
Column
1 2 3 4 5 6 7 8 9 10 11 12 13 14 15 16 17 18 19 20 21 22 23 24
rel. FI
1.50
1.25
1.00
0.75
0.50

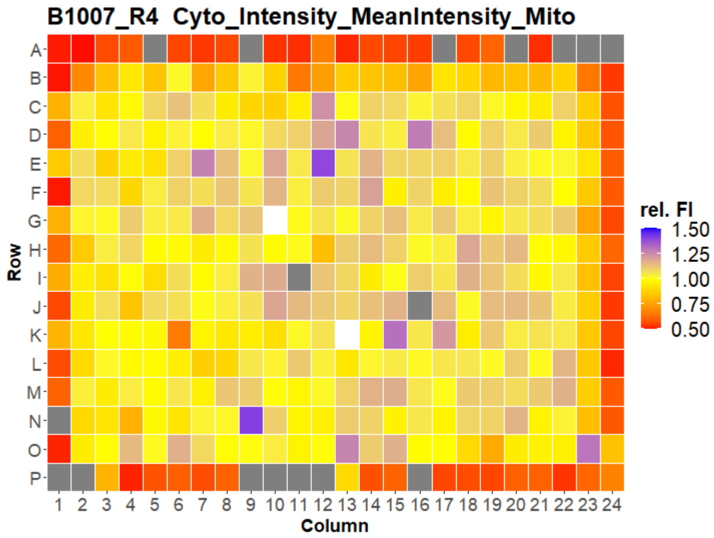
B1007_R4 Cyto_Intensity_MeanIntensity_Mito
Row
A B C D E F G H I J K L M N O P
Column
1 2 3 4 5 6 7 8 9 10 11 12 13 14 15 16 17 18 19 20 21 22 23 24
rel. FI
1.50
1.25
1.00
0.75
0.50

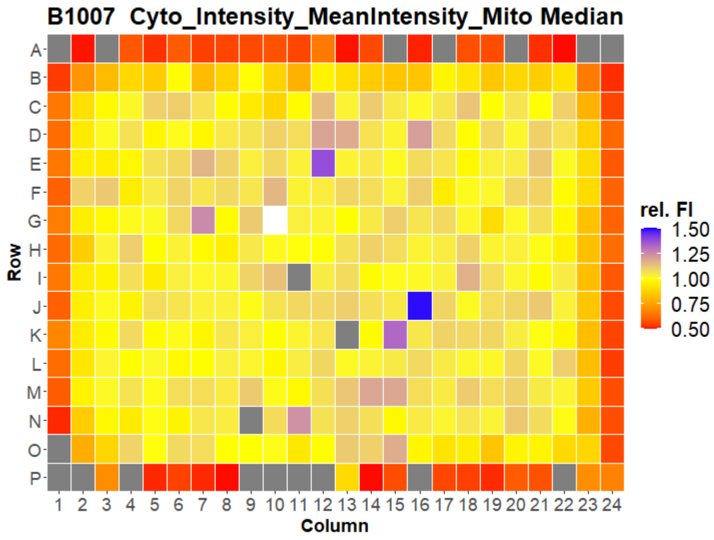
B1007 Cyto_Intensity_MeanIntensity_Mito Median
Row
A B C D E F G H I J K L M N O P
Column
1 2 3 4 5 6 7 8 9 10 11 12 13 14 15 16 17 18 19 20 21 22 23 24
rel. FI
1.50
1.25
1.00
0.75
0.50

### U2OS FMP

22

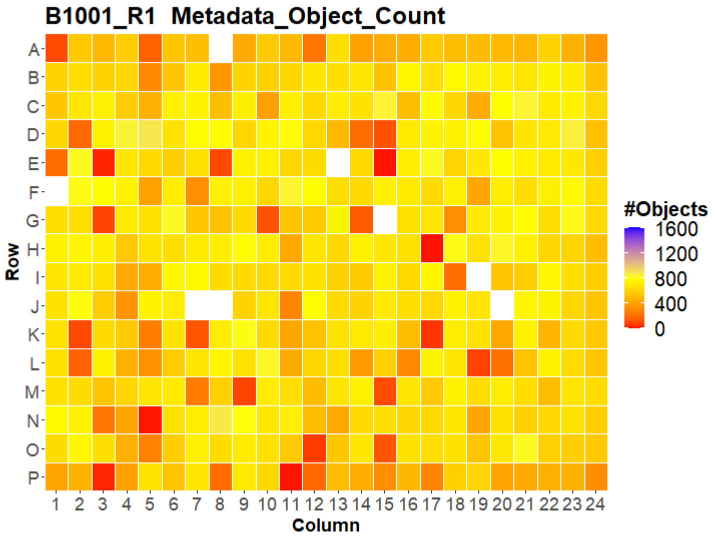


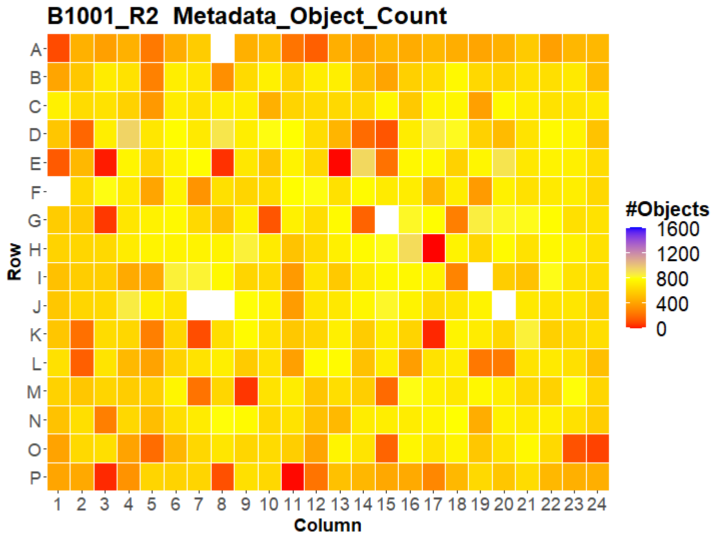


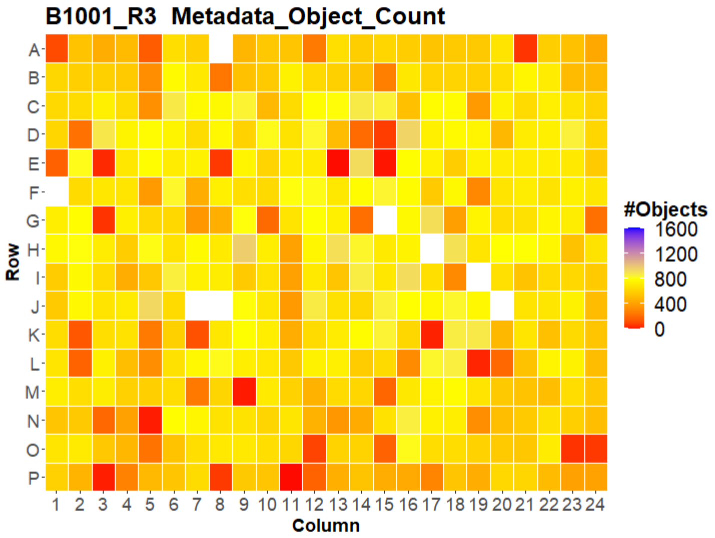


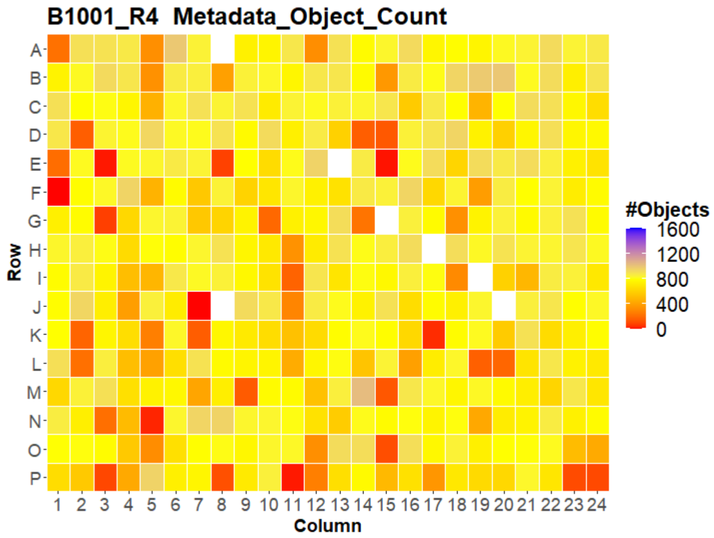


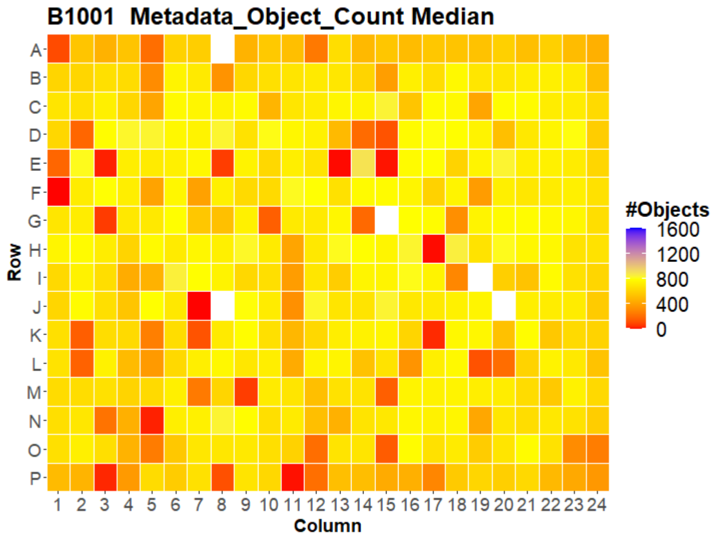




Open for Innovation KNIME

### U2OS FMP

23

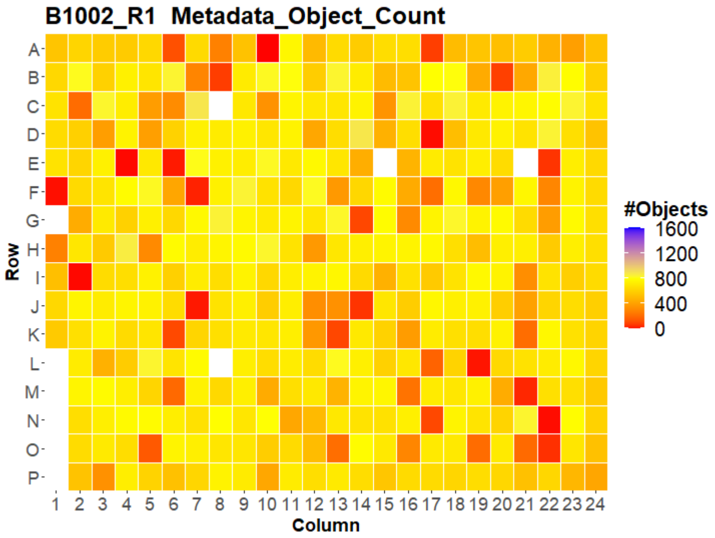
B1002_R1 Metadata_Object_Count
A B C D E F G H I J K L M N O P
Row
1 2 3 4 5 6 7 8 9 10 11 12 13 14 15 16 17 18 19 20 21 22 23 24
Column
#Objects
1600
1200
800
400
0

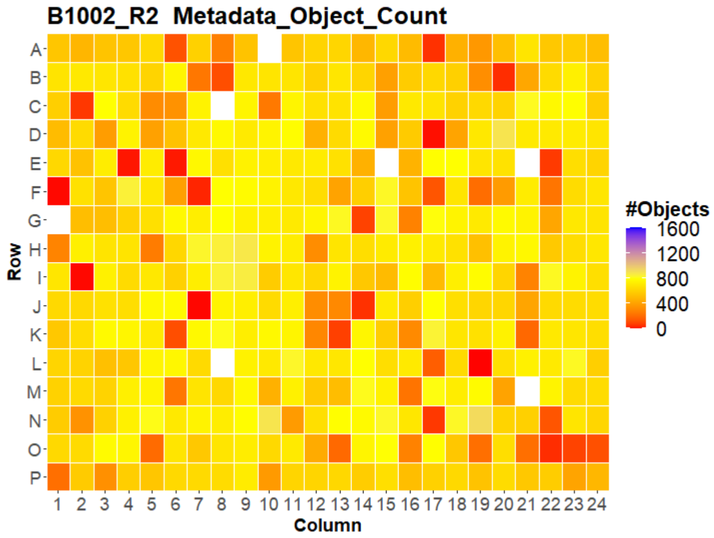
B1002_R2 Metadata_Object_Count
A B C D E F G H I J K L M N O P
Row
1 2 3 4 5 6 7 8 9 10 11 12 13 14 15 16 17 18 19 20 21 22 23 24
Column
#Objects
1600
1200
800
400
0

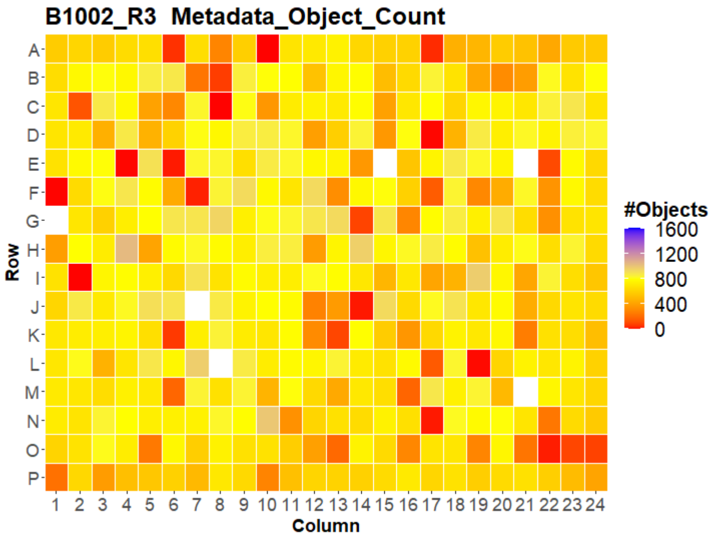
B1002_R3 Metadata_Object_Count
A B C D E F G H I J K L M N O P
Row
1 2 3 4 5 6 7 8 9 10 11 12 13 14 15 16 17 18 19 20 21 22 23 24
Column
#Objects
1600
1200
800
400
0

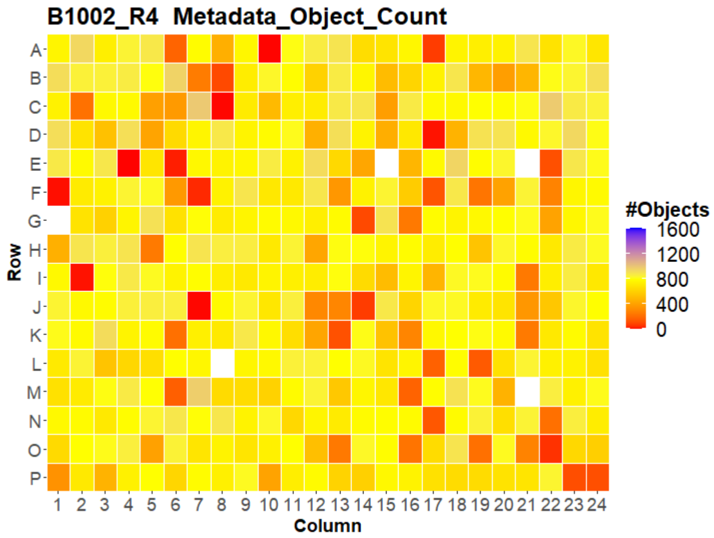
B1002_R4 Metadata_Object_Count
A B C D E F G H I J K L M N O P
Row
1 2 3 4 5 6 7 8 9 10 11 12 13 14 15 16 17 18 19 20 21 22 23 24
Column
#Objects
1600
1200
800
400
0

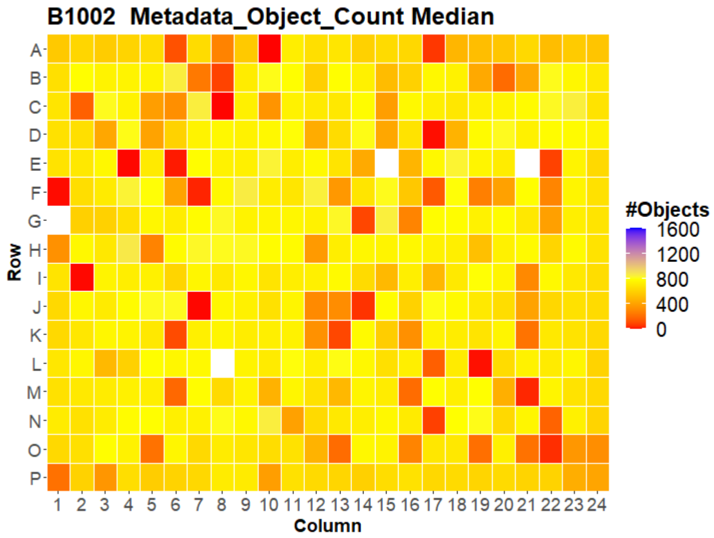
B1002 Metadata_Object_Count Median
A B C D E F G H I J K L M N O P
Row
1 2 3 4 5 6 7 8 9 10 11 12 13 14 15 16 17 18 19 20 21 22 23 24
Column
#Objects
1600
1200
800
400
0

### U2OS FMP



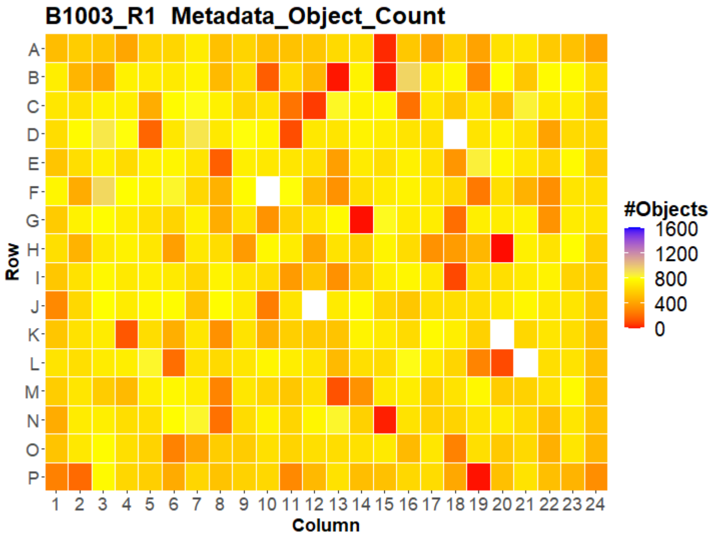
B1003_R1 Metadata_Object_Count
#Objects
1600
1200
800
400
0
Row
Column

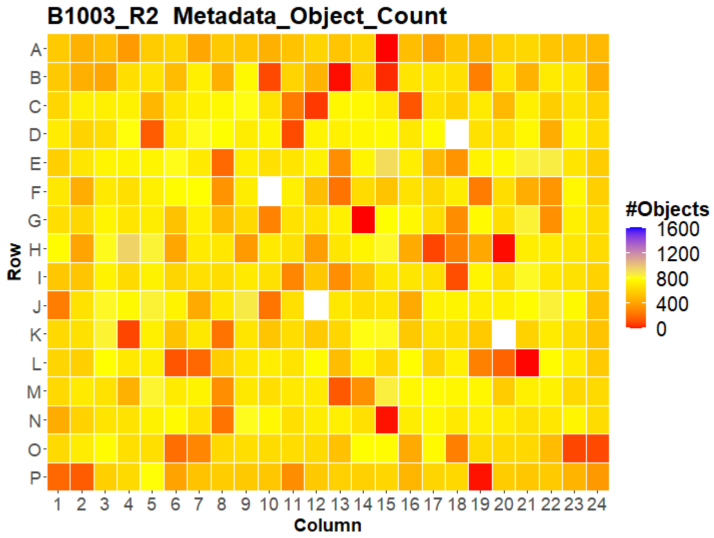
B1003_R2 Metadata_Object_Count
#Objects
1600
1200
800
400
0
Row
Column

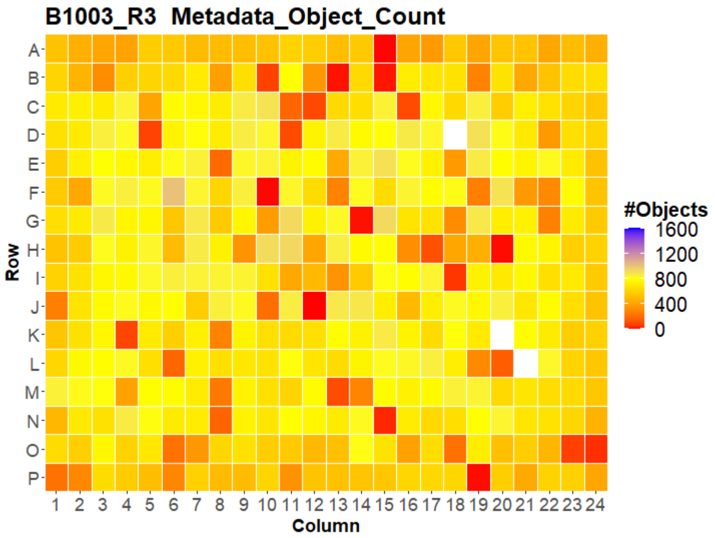
B1003_R3 Metadata_Object_Count
#Objects
1600
1200
800
400
0
Row
Column

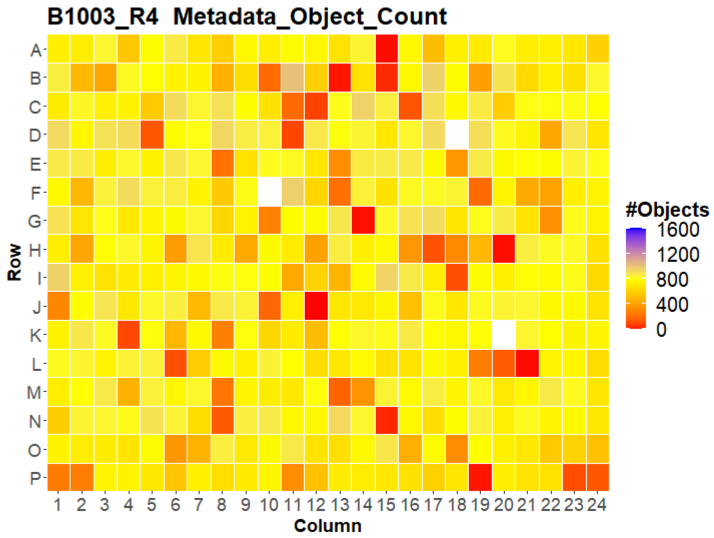
B1003_R4 Metadata_Object_Count
#Objects
1600
1200
800
400
0
Row
Column

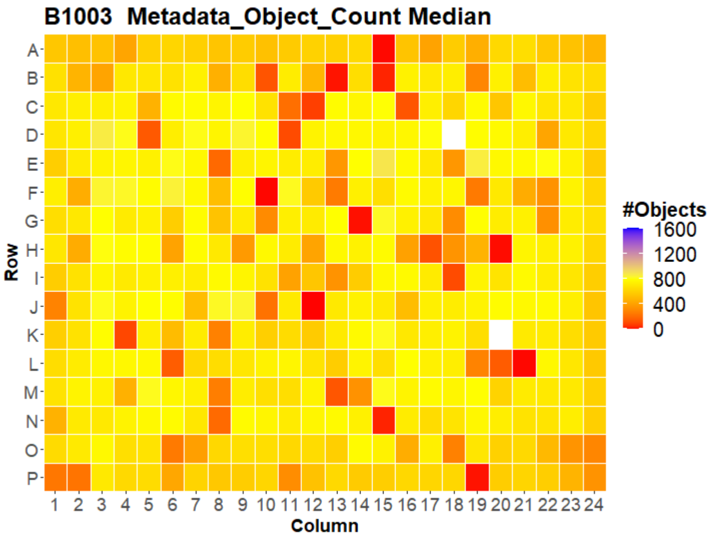
B1003 Metadata_Object_Count Median
#Objects
1600
1200
800
400
0
Row
Column

### U2OS FMP



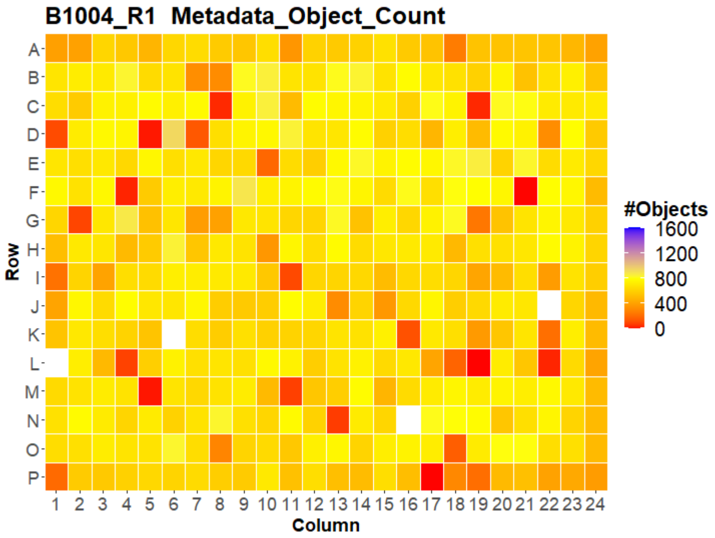
B1004_R1 Metadata_Object_Count
Row
Column
#Objects
1600
1200
800
400
0

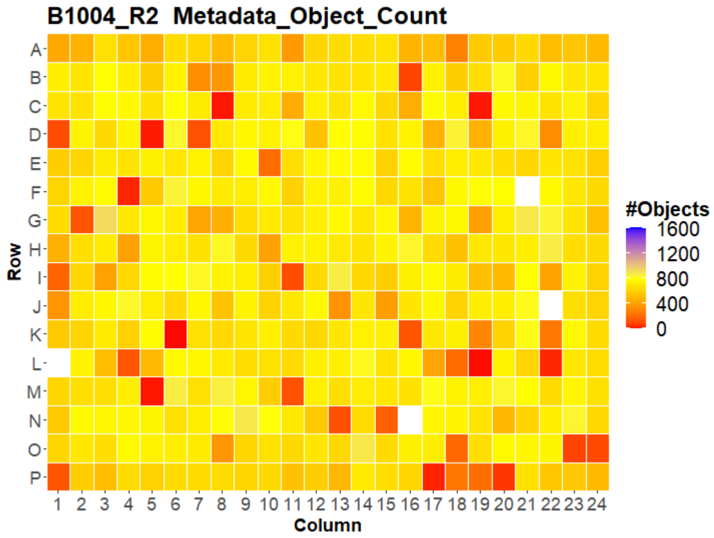
B1004_R2 Metadata_Object_Count
Row
Column
#Objects
1600
1200
800
400
0

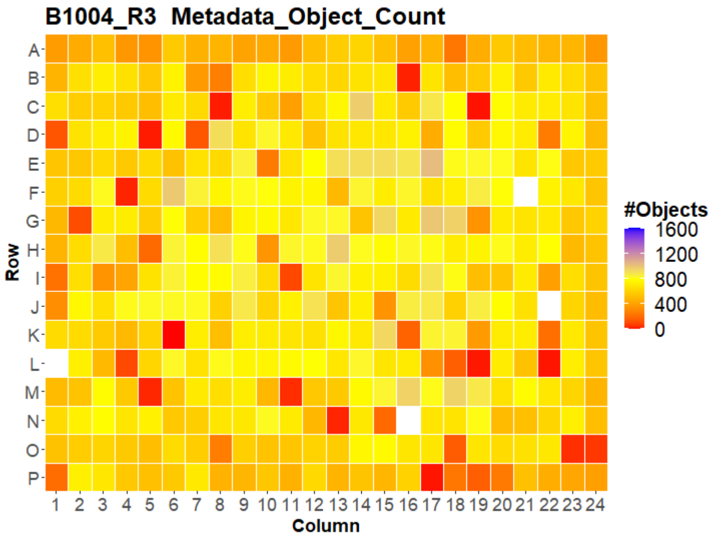
B1004_R3 Metadata_Object_Count
Row
Column
#Objects
1600
1200
800
400
0

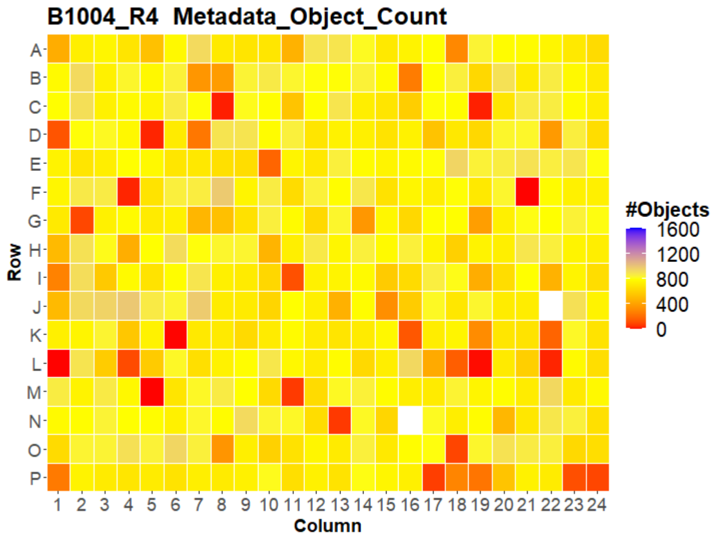
B1004_R4 Metadata_Object_Count
Row
Column
#Objects
1600
1200
800
400
0

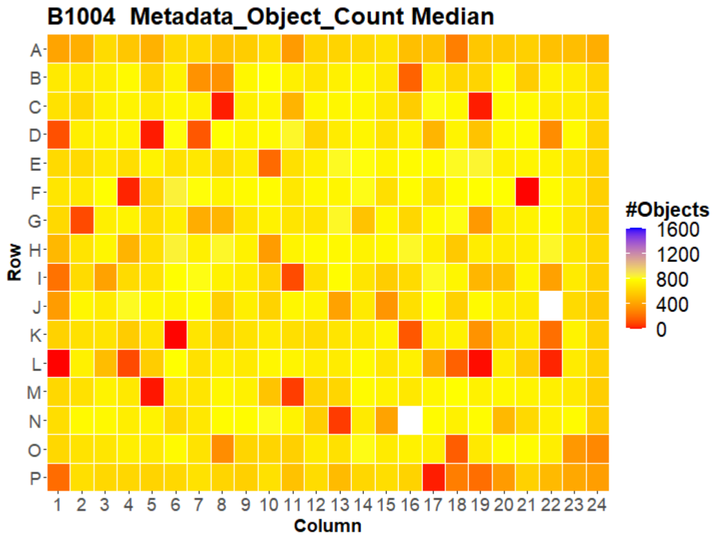
B1004 Metadata_Object_Count Median
Row
Column
#Objects
1600
1200
800
400
0

### U2OS FMP

26

B1005_R1 Metadata_Object_Count

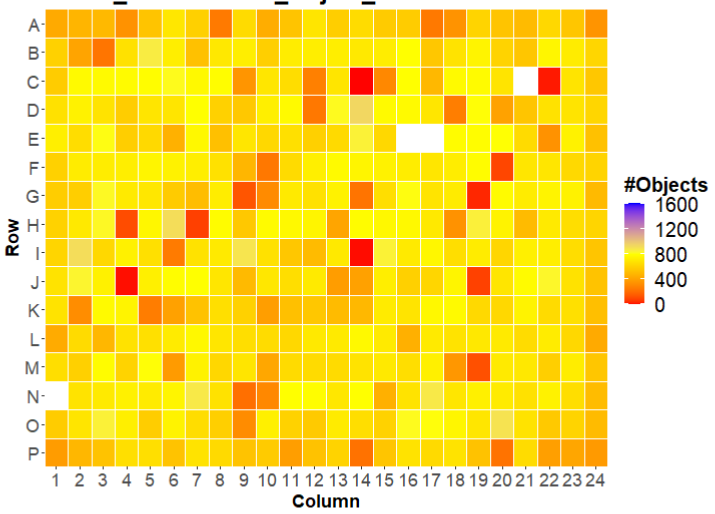


B1005_R2 Metadata_Object_Count

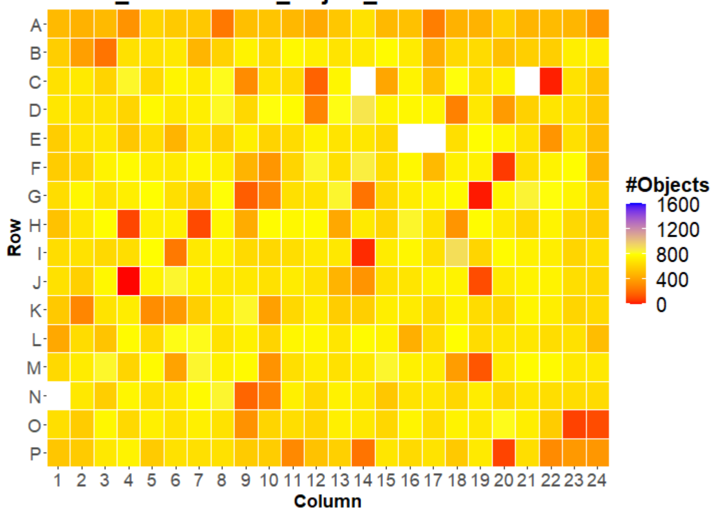


B1005_R3 Metadata_Object_Count

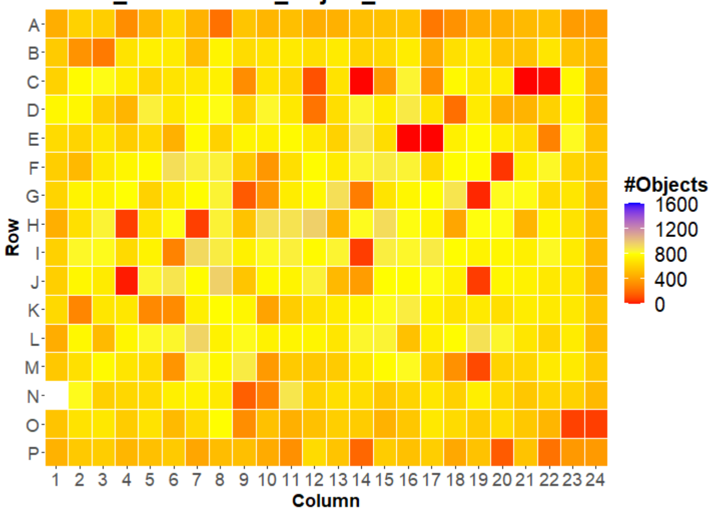


B1005_R4 Metadata_Object_Count

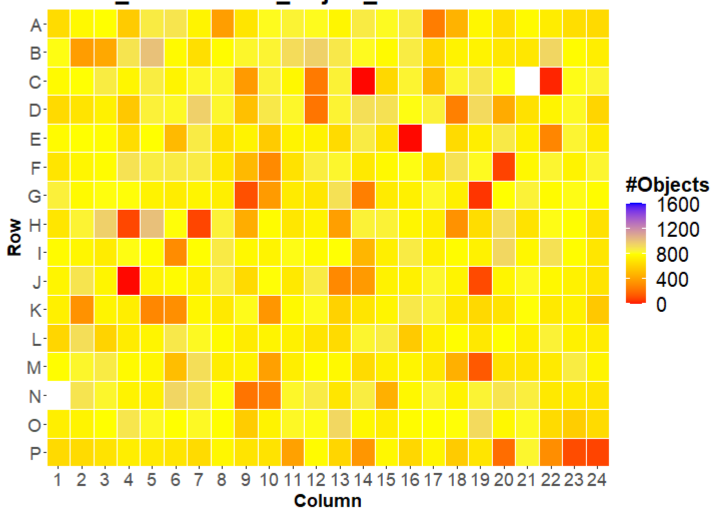


B1005 Metadata_Object_Count Median

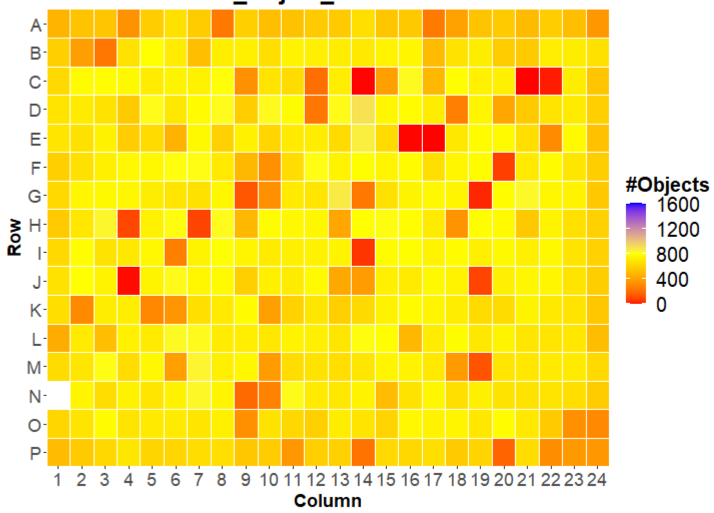

### U2OS FMP

27

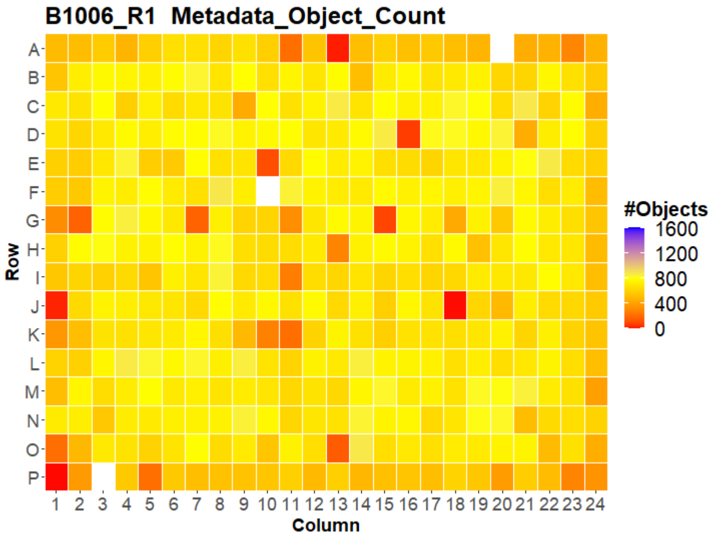
B1006_R1 Metadata_Object_Count
Row
Column
#Objects
1600
1200
800
400
0

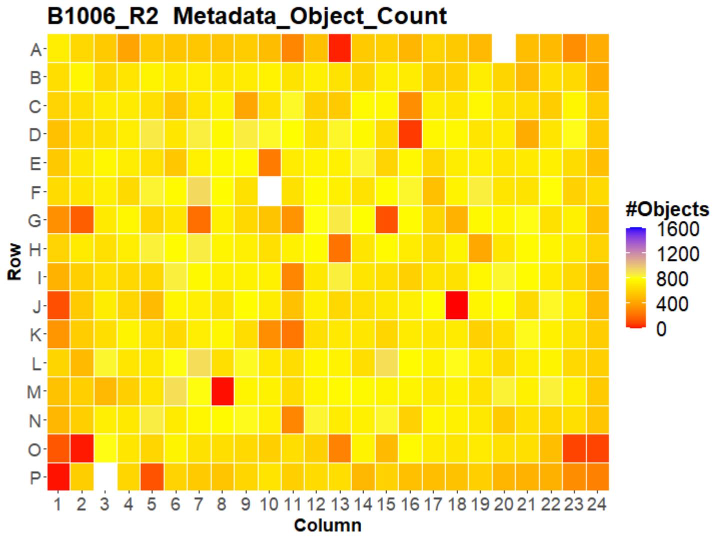
B1006_R2 Metadata_Object_Count
Row
Column
#Objects
1600
1200
800
400
0

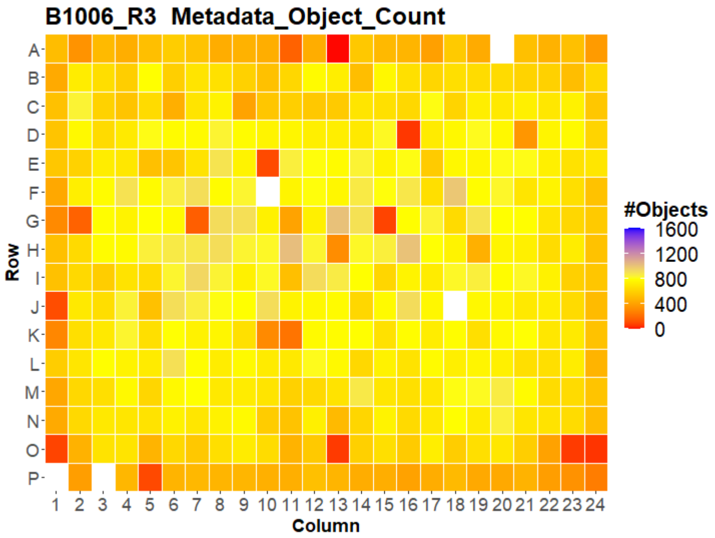
B1006_R3 Metadata_Object_Count
Row
Column
#Objects
1600
1200
800
400
0

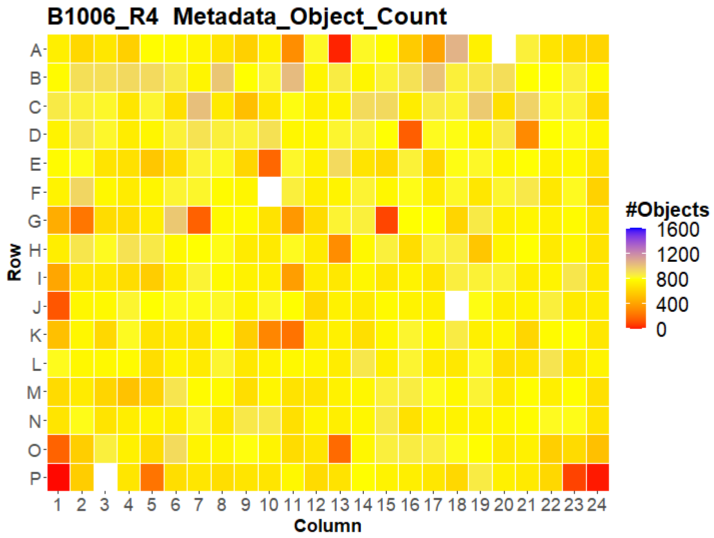
B1006_R4 Metadata_Object_Count
Row
Column
#Objects
1600
1200
800
400
0

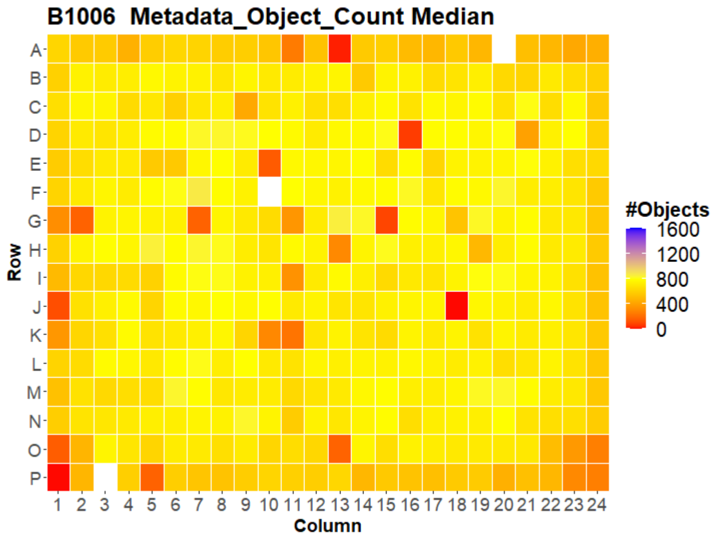
B1006 Metadata_Object_Count Median
Row
Column
#Objects
1600
1200
800
400
0

### U2OS FMP

28

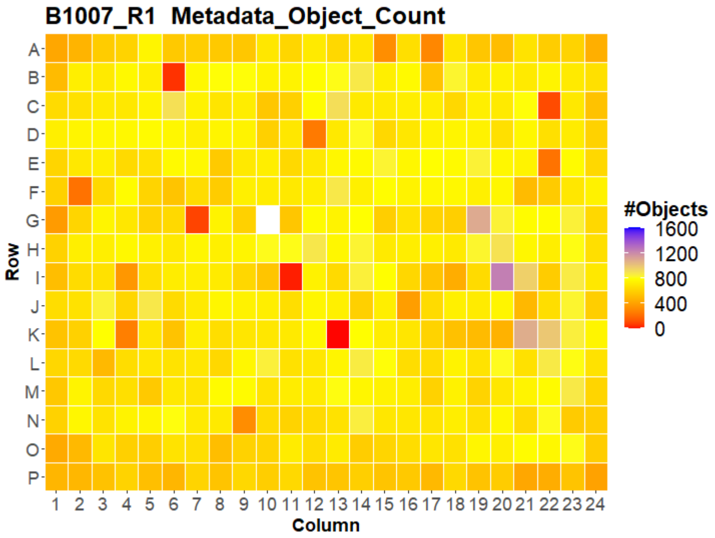
B1007_R1 Metadata_Object_Count
A
B
C
D
E
F
G
H
I
J
K
L
M
N
O
P
Row
1 2 3 4 5 6 7 8 9 10 11 12 13 14 15 16 17 18 19 20 21 22 23 24
Column
#Objects
1600
1200
800
400
0

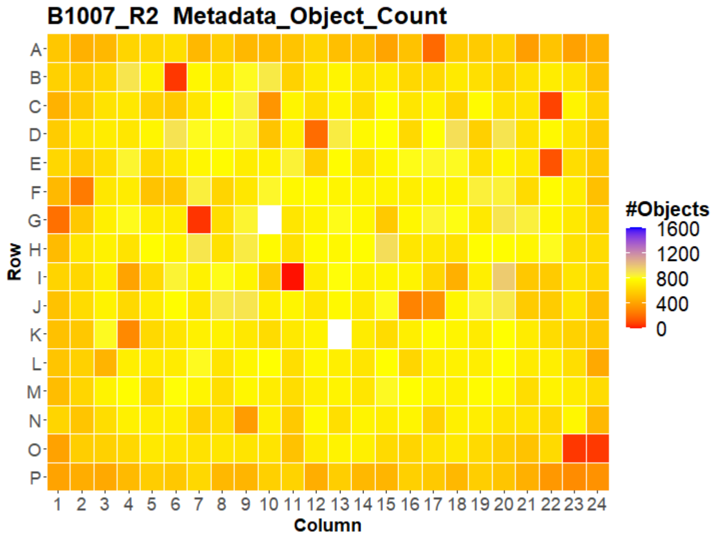
B1007_R2 Metadata_Object_Count
A
B
C
D
E
F
G
H
I
J
K
L
M
N
O
P
Row
1 2 3 4 5 6 7 8 9 10 11 12 13 14 15 16 17 18 19 20 21 22 23 24
Column
#Objects
1600
1200
800
400
0

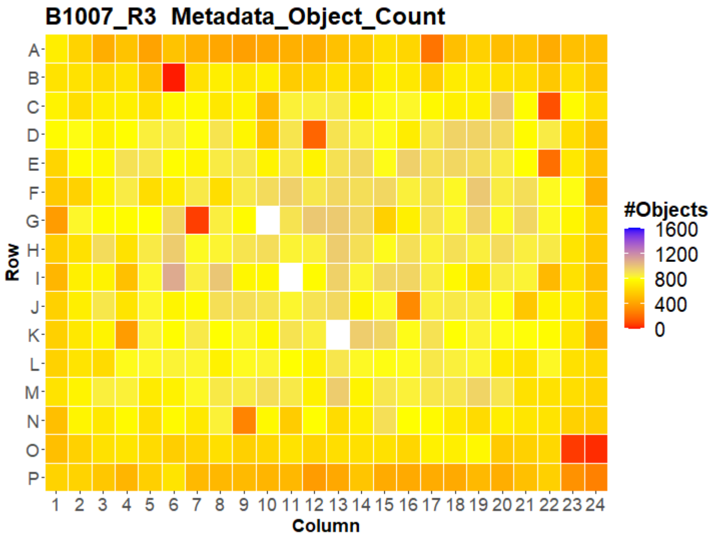
B1007_R3 Metadata_Object_Count
A
B
C
D
E
F
G
H
I
J
K
L
M
N
O
P
Row
1 2 3 4 5 6 7 8 9 10 11 12 13 14 15 16 17 18 19 20 21 22 23 24
Column
#Objects
1600
1200
800
400
0

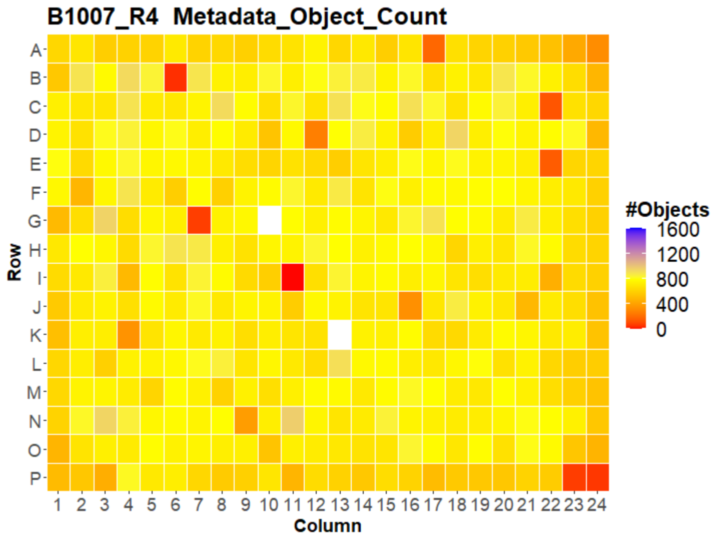
B1007_R4 Metadata_Object_Count
A
B
C
D
E
F
G
H
I
J
K
L
M
N
O
P
Row
1 2 3 4 5 6 7 8 9 10 11 12 13 14 15 16 17 18 19 20 21 22 23 24
Column
#Objects
1600
1200
800
400
0

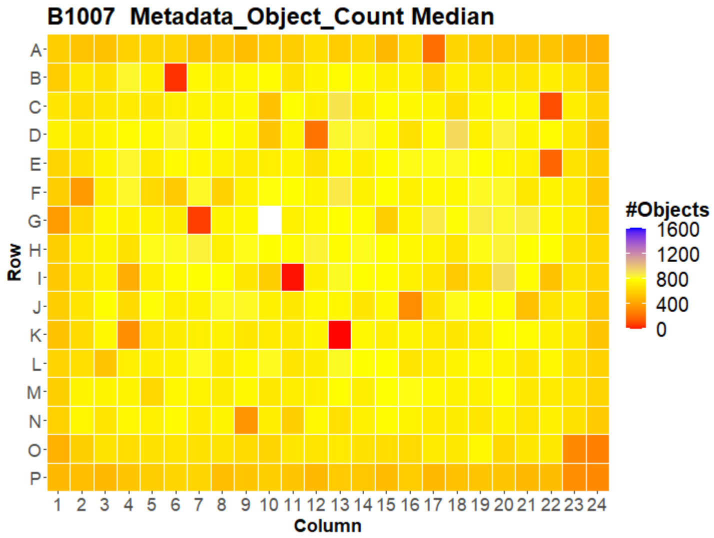
B1007 Metadata_Object_Count Median
A
B
C
D
E
F
G
H
I
J
K
L
M
N
O
P
Row
1 2 3 4 5 6 7 8 9 10 11 12 13 14 15 16 17 18 19 20 21 22 23 24
Column
#Objects
1600
1200
800
400
0

### U2OS FMP

29

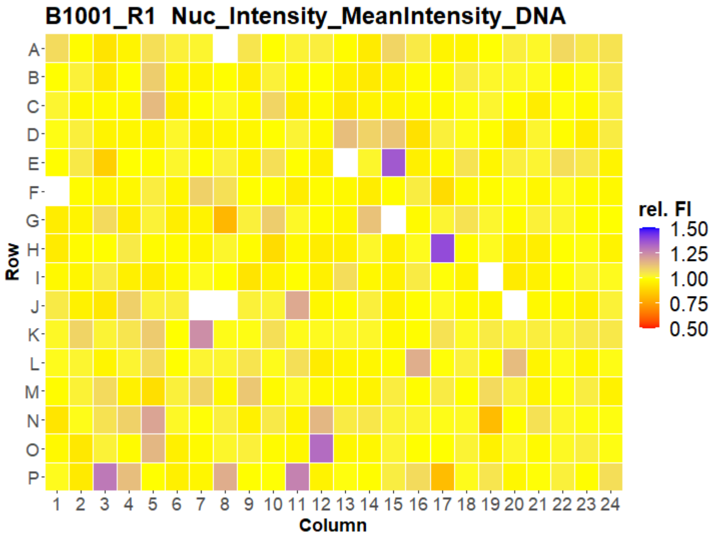
B1001_R1 Nuc_Intensity_MeanIntensity_DNA
Row
Column
rel. FI
1.50
1.25
1.00
0.75
0.50

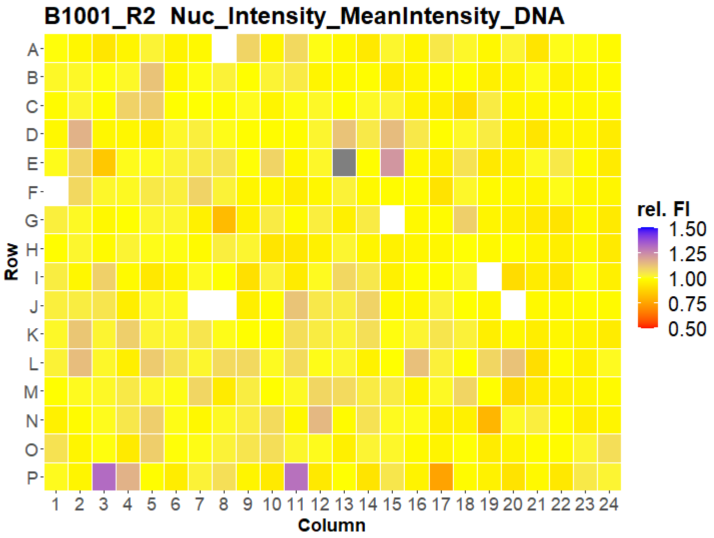
B1001_R2 Nuc_Intensity_MeanIntensity_DNA
Row
Column
rel. FI
1.50
1.25
1.00
0.75
0.50

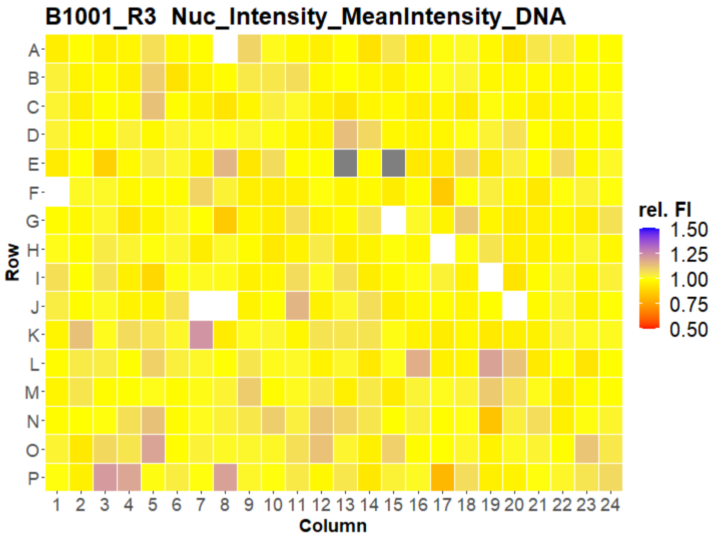
B1001_R3 Nuc_Intensity_MeanIntensity_DNA
Row
Column
rel. FI
1.50
1.25
1.00
0.75
0.50

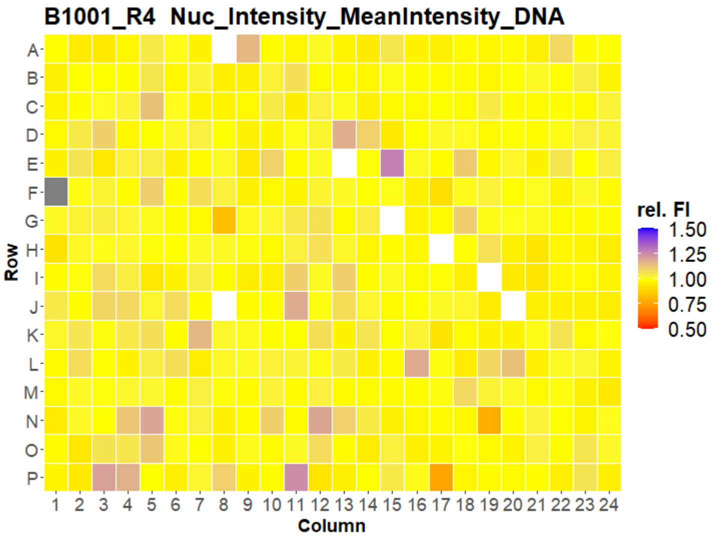
B1001_R4 Nuc_Intensity_MeanIntensity_DNA
Row
Column
rel. FI
1.50
1.25
1.00
0.75
0.50

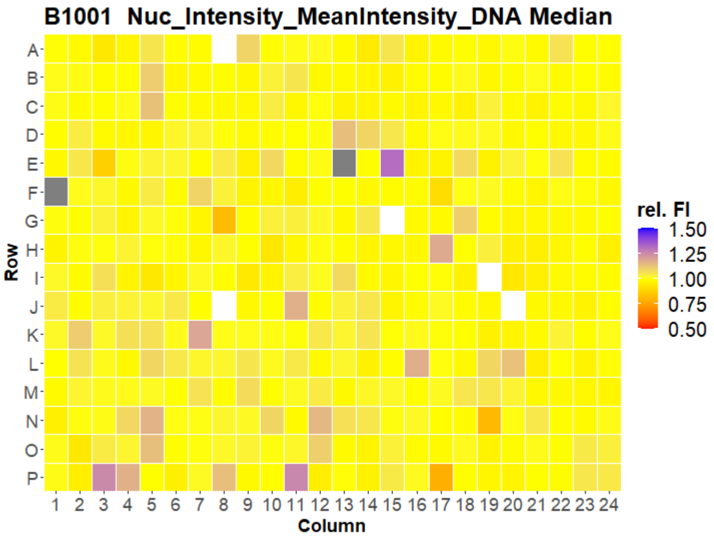
B1001 Nuc_Intensity_MeanIntensity_DNA Median
Row
Column
rel. FI
1.50
1.25
1.00
0.75
0.50

### U2OS FMP

30

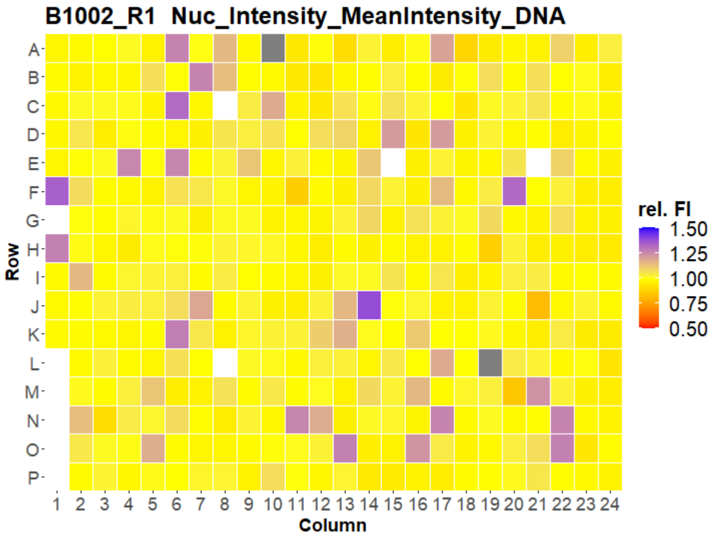
B1002_R1 Nuc_Intensity_MeanIntensity_DNA
Row
A B C D E F G H I J K L M N O P
1 2 3 4 5 6 7 8 9 10 11 12 13 14 15 16 17 18 19 20 21 22 23 24
Column
rel. FI
1.50
1.25
1.00
0.75
0.50

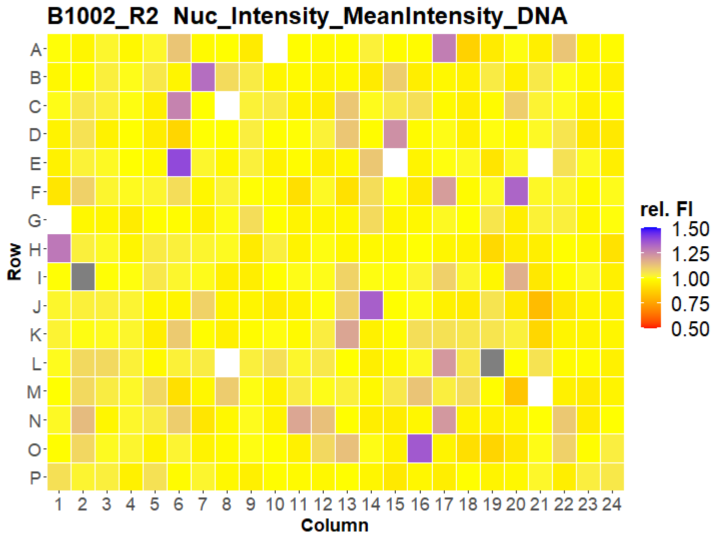
B1002_R2 Nuc_Intensity_MeanIntensity_DNA
Row
A B C D E F G H I J K L M N O P
1 2 3 4 5 6 7 8 9 10 11 12 13 14 15 16 17 18 19 20 21 22 23 24
Column
rel. FI
1.50
1.25
1.00
0.75
0.50

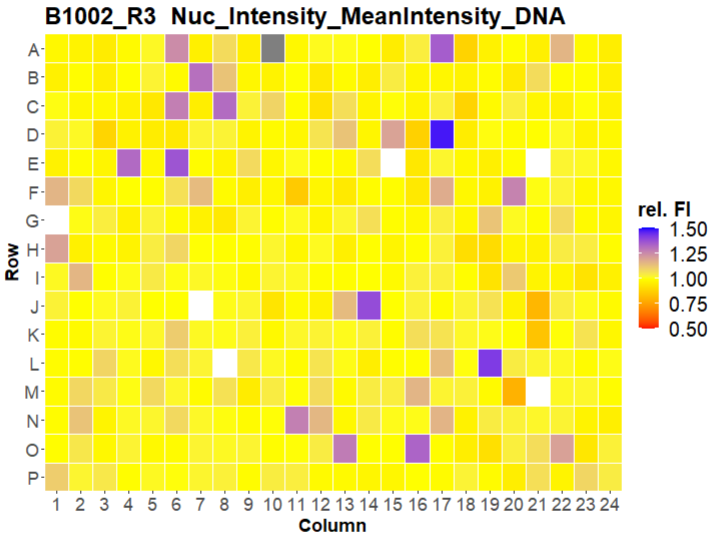
B1002_R3 Nuc_Intensity_MeanIntensity_DNA
Row
A B C D E F G H I J K L M N O P
1 2 3 4 5 6 7 8 9 10 11 12 13 14 15 16 17 18 19 20 21 22 23 24
Column
rel. FI
1.50
1.25
1.00
0.75
0.50

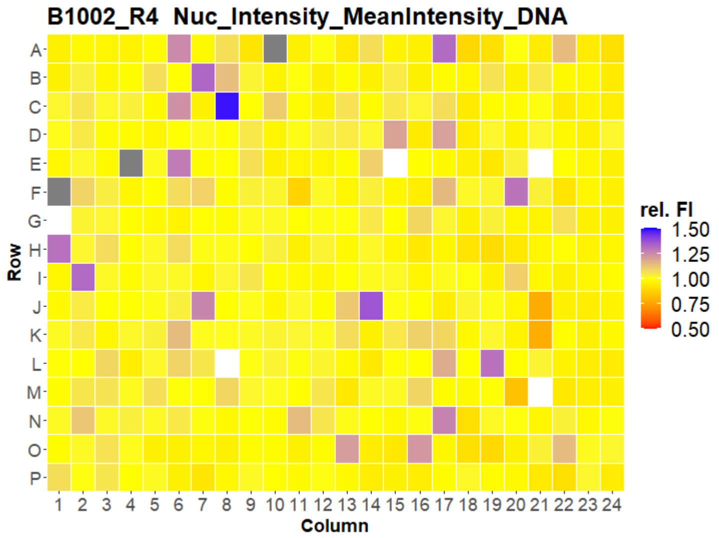
B1002_R4 Nuc_Intensity_MeanIntensity_DNA
Row
A B C D E F G H I J K L M N O P
1 2 3 4 5 6 7 8 9 10 11 12 13 14 15 16 17 18 19 20 21 22 23 24
Column
rel. FI
1.50
1.25
1.00
0.75
0.50

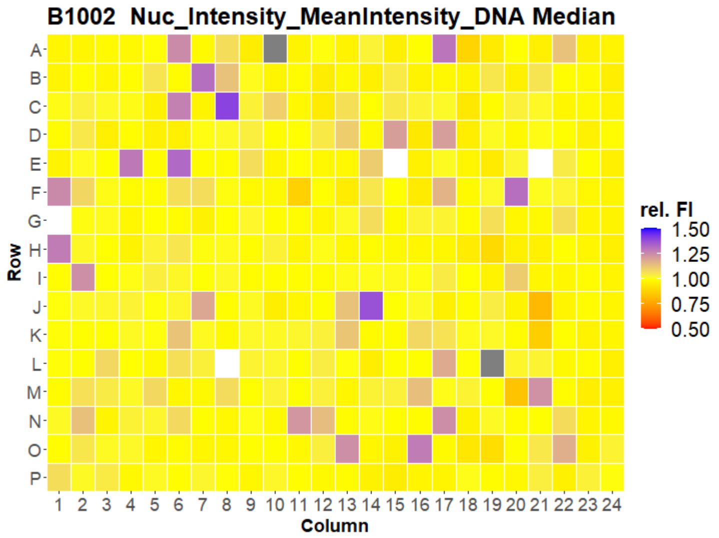
B1002 Nuc_Intensity_MeanIntensity_DNA Median
Row
A B C D E F G H I J K L M N O P
1 2 3 4 5 6 7 8 9 10 11 12 13 14 15 16 17 18 19 20 21 22 23 24
Column
rel. FI
1.50
1.25
1.00
0.75
0.50

### U2OS FMP

31

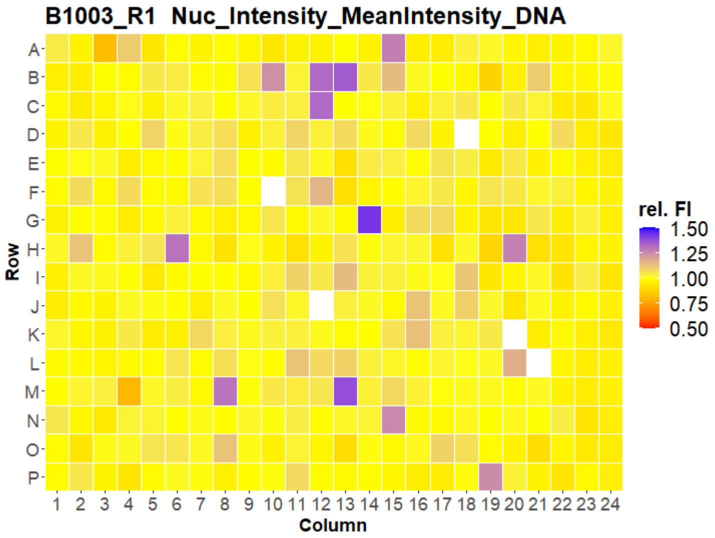
B1003_R1 Nuc_Intensity_MeanIntensity_DNA
Row
A B C D E F G H I J K L M N O P
Column
1 2 3 4 5 6 7 8 9 10 11 12 13 14 15 16 17 18 19 20 21 22 23 24
rel. FI
1.50
1.25
1.00
0.75
0.50

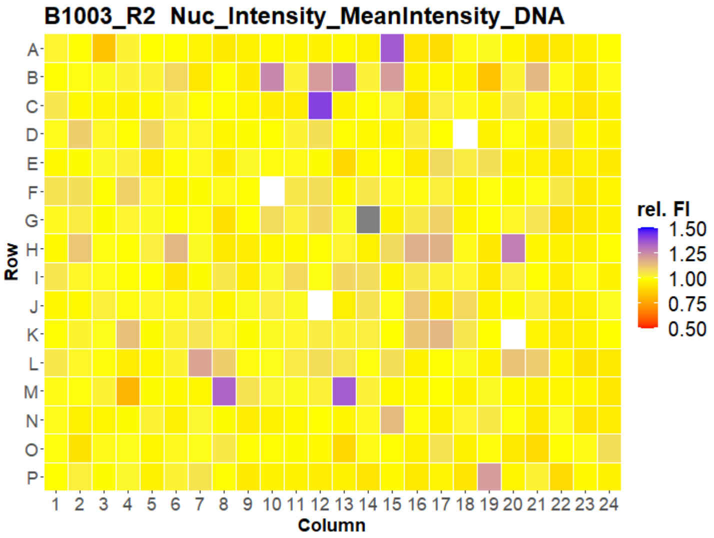
B1003_R2 Nuc_Intensity_MeanIntensity_DNA
Row
A B C D E F G H I J K L M N O P
Column
1 2 3 4 5 6 7 8 9 10 11 12 13 14 15 16 17 18 19 20 21 22 23 24
rel. FI
1.50
1.25
1.00
0.75
0.50

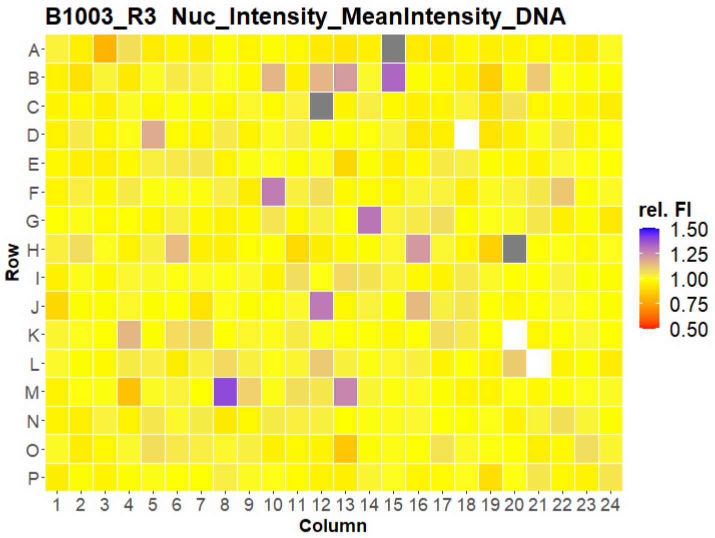
B1003_R3 Nuc_Intensity_MeanIntensity_DNA
Row
A B C D E F G H I J K L M N O P
Column
1 2 3 4 5 6 7 8 9 10 11 12 13 14 15 16 17 18 19 20 21 22 23 24
rel. FI
1.50
1.25
1.00
0.75
0.50

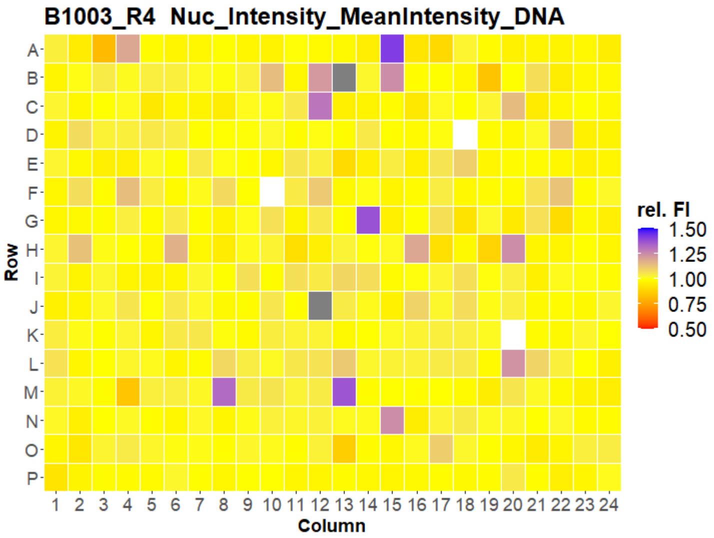
B1003_R4 Nuc_Intensity_MeanIntensity_DNA
Row
A B C D E F G H I J K L M N O P
Column
1 2 3 4 5 6 7 8 9 10 11 12 13 14 15 16 17 18 19 20 21 22 23 24
rel. FI
1.50
1.25
1.00
0.75
0.50

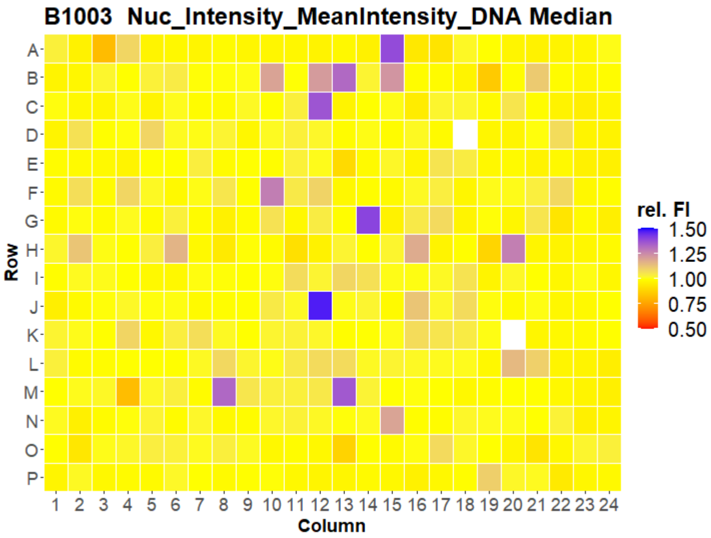
B1003 Nuc_Intensity_MeanIntensity_DNA Median
Row
A B C D E F G H I J K L M N O P
Column
1 2 3 4 5 6 7 8 9 10 11 12 13 14 15 16 17 18 19 20 21 22 23 24
rel. FI
1.50
1.25
1.00
0.75
0.50

### U2OS FMP



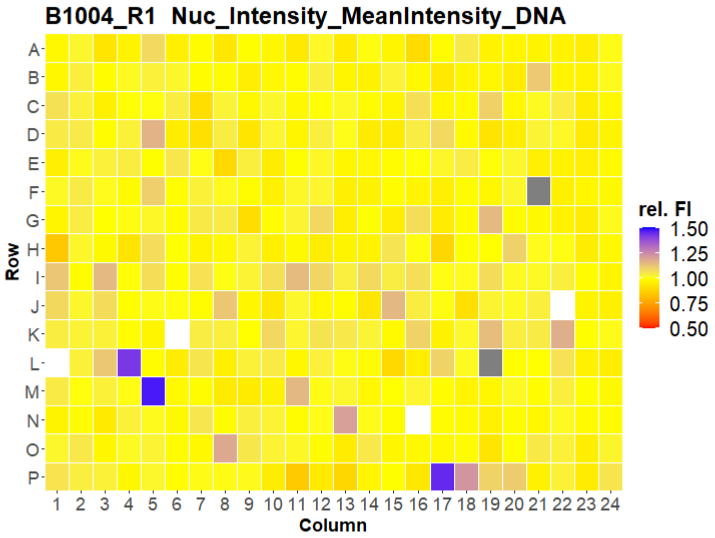
B1004_R1 Nuc_Intensity_MeanIntensity_DNA
Row
Column
rel. FI
1.50
1.25
1.00
0.75
0.50

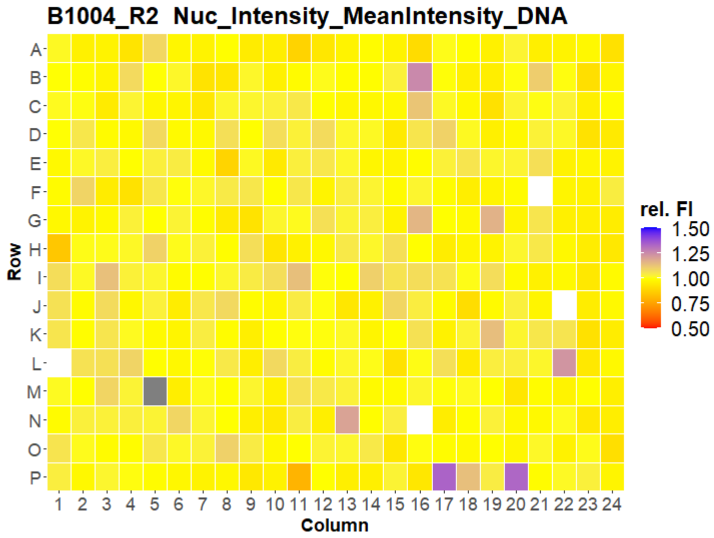
B1004_R2 Nuc_Intensity_MeanIntensity_DNA
Row
Column
rel. FI
1.50
1.25
1.00
0.75
0.50

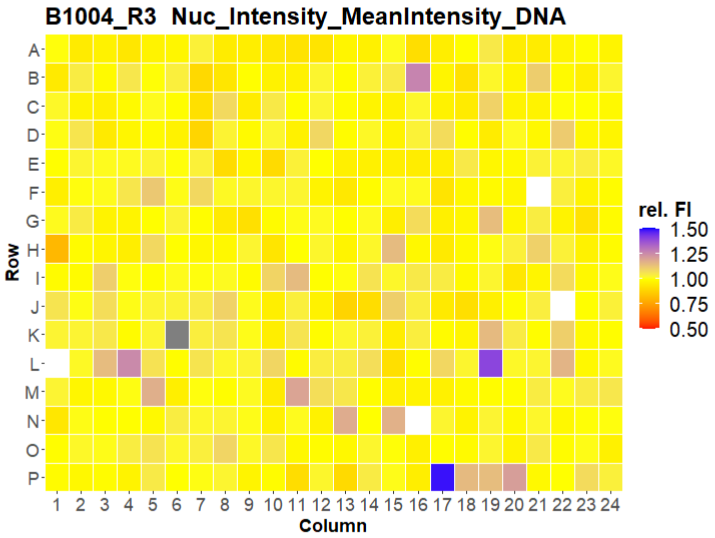
B1004_R3 Nuc_Intensity_MeanIntensity_DNA
Row
Column
rel. FI
1.50
1.25
1.00
0.75
0.50

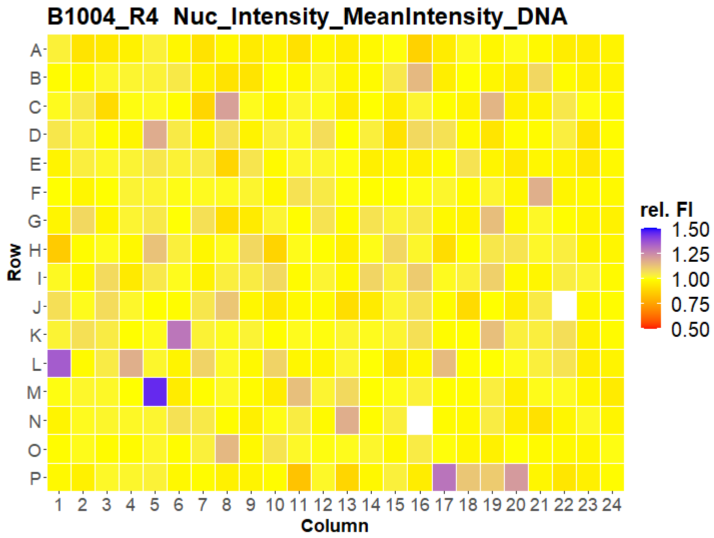
B1004_R4 Nuc_Intensity_MeanIntensity_DNA
Row
Column
rel. FI
1.50
1.25
1.00
0.75
0.50

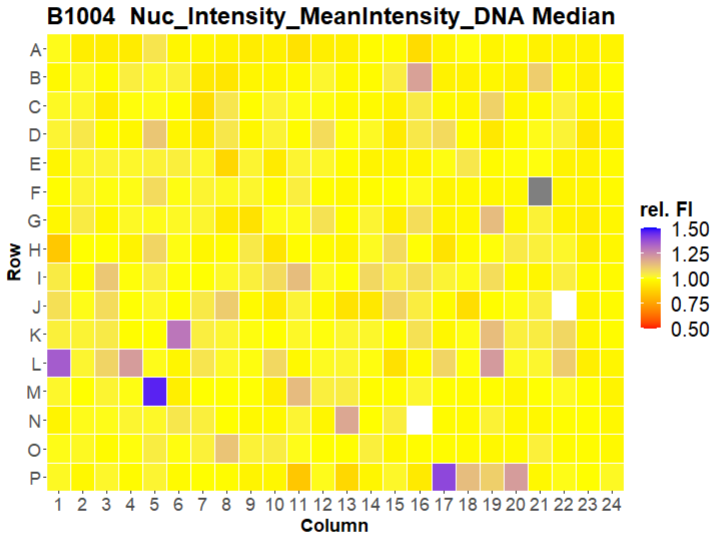
B1004 Nuc_Intensity_MeanIntensity_DNA Median
Row
Column
rel. FI
1.50
1.25
1.00
0.75
0.50

### U2OS FMP



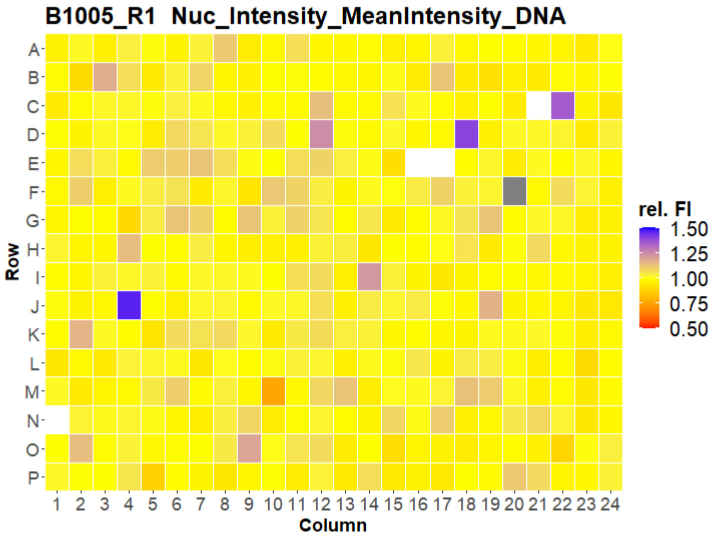
B1005_R1 Nuc_Intensity_MeanIntensity_DNA
Row
A B C D E F G H I J K L M N O P
1 2 3 4 5 6 7 8 9 10 11 12 13 14 15 16 17 18 19 20 21 22 23 24
Column
rel. FI
1.50
1.25
1.00
0.75
0.50

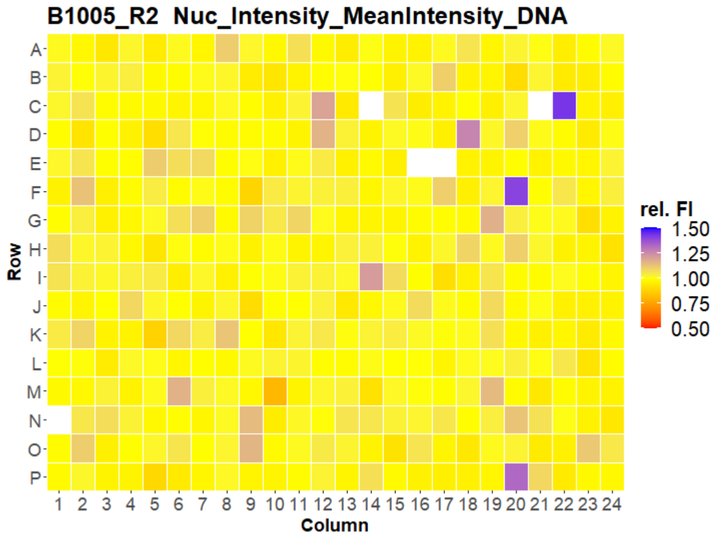
B1005_R2 Nuc_Intensity_MeanIntensity_DNA
Row
A B C D E F G H I J K L M N O P
1 2 3 4 5 6 7 8 9 10 11 12 13 14 15 16 17 18 19 20 21 22 23 24
Column
rel. FI
1.50
1.25
1.00
0.75
0.50

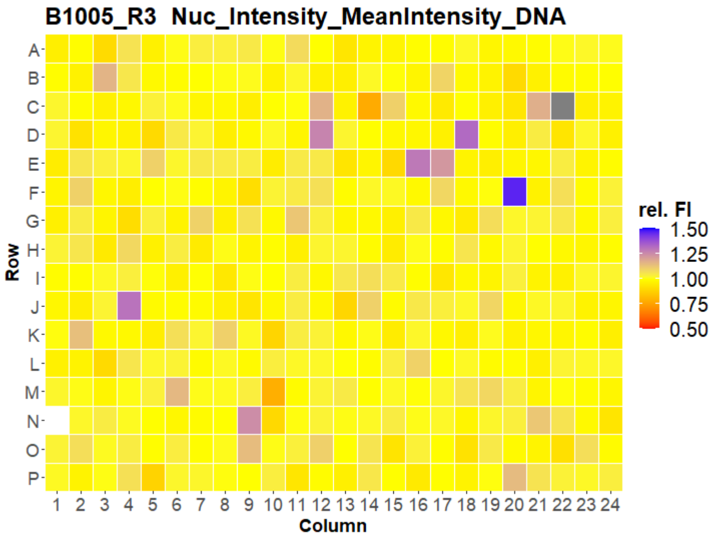
B1005_R3 Nuc_Intensity_MeanIntensity_DNA
Row
A B C D E F G H I J K L M N O P
1 2 3 4 5 6 7 8 9 10 11 12 13 14 15 16 17 18 19 20 21 22 23 24
Column
rel. FI
1.50
1.25
1.00
0.75
0.50

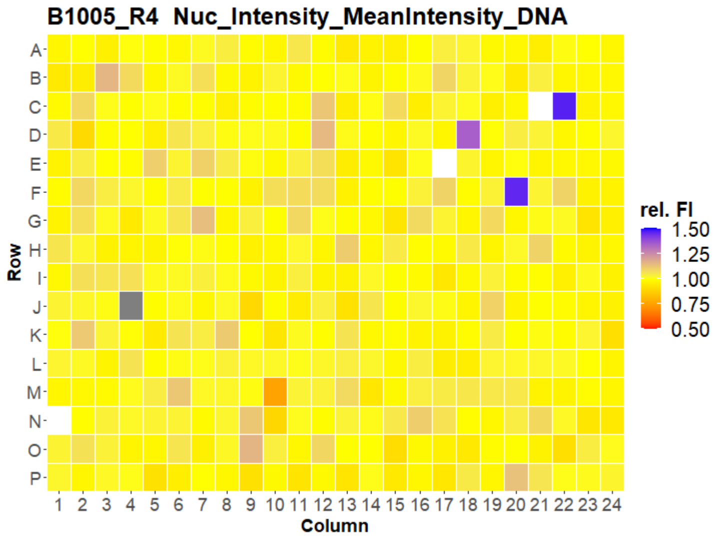
B1005_R4 Nuc_Intensity_MeanIntensity_DNA
Row
A B C D E F G H I J K L M N O P
1 2 3 4 5 6 7 8 9 10 11 12 13 14 15 16 17 18 19 20 21 22 23 24
Column
rel. FI
1.50
1.25
1.00
0.75
0.50

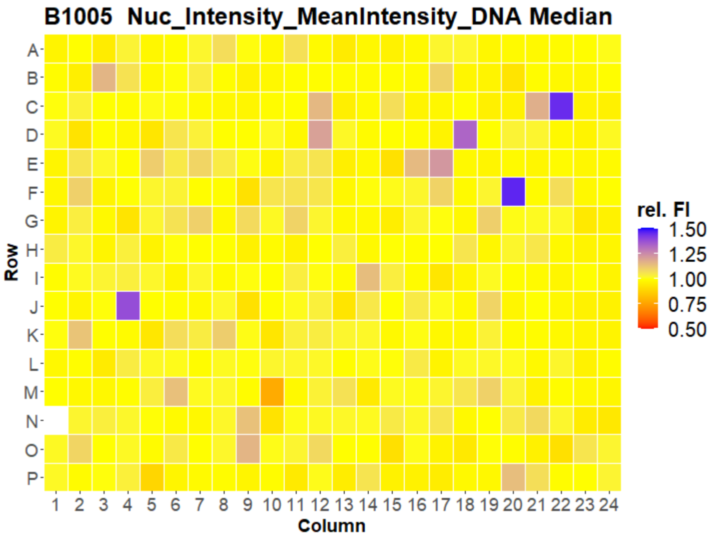
B1005 Nuc_Intensity_MeanIntensity_DNA Median
Row
A B C D E F G H I J K L M N O P
1 2 3 4 5 6 7 8 9 10 11 12 13 14 15 16 17 18 19 20 21 22 23 24
Column
rel. FI
1.50
1.25
1.00
0.75
0.50

### U2OS FMP



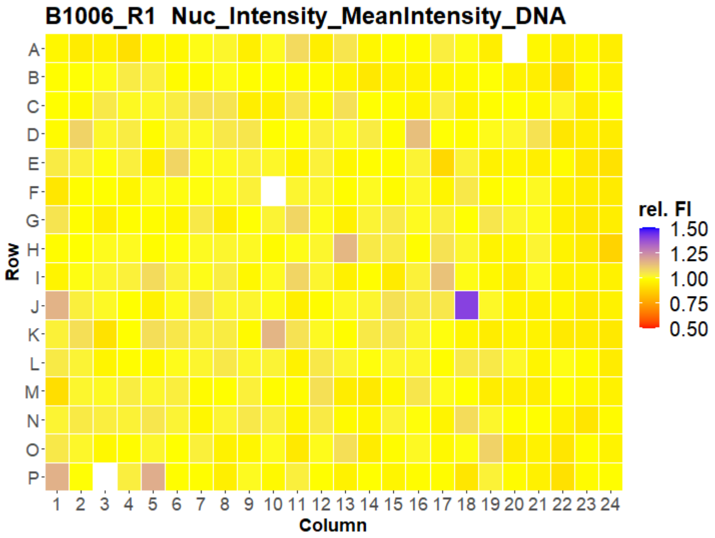
B1006_R1 Nuc_Intensity_MeanIntensity_DNA
Row
Column
rel. FI
1.50
1.25
1.00
0.75
0.50

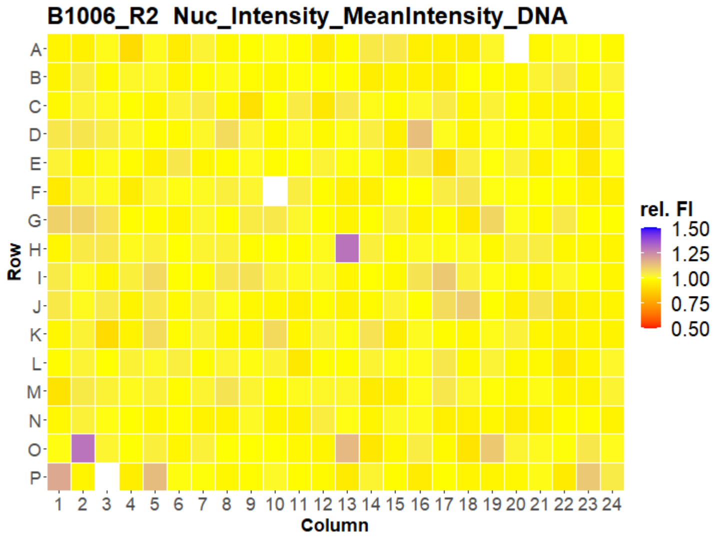
B1006_R2 Nuc_Intensity_MeanIntensity_DNA
Row
Column
rel. FI
1.50
1.25
1.00
0.75
0.50

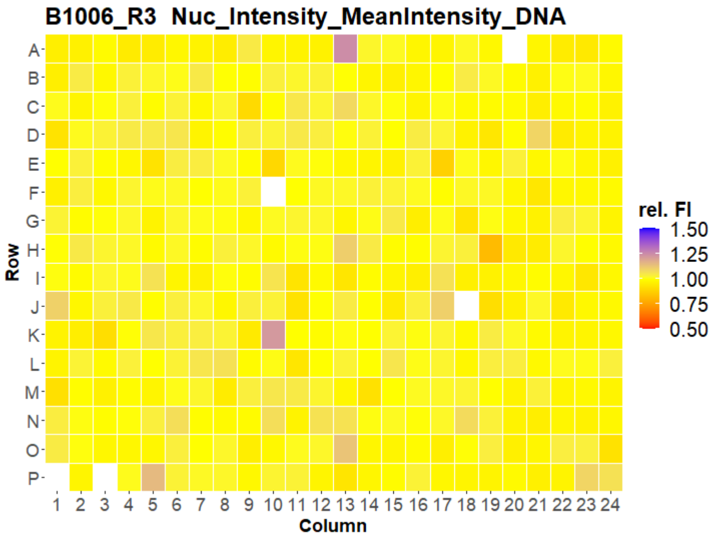
B1006_R3 Nuc_Intensity_MeanIntensity_DNA
Row
Column
rel. FI
1.50
1.25
1.00
0.75
0.50

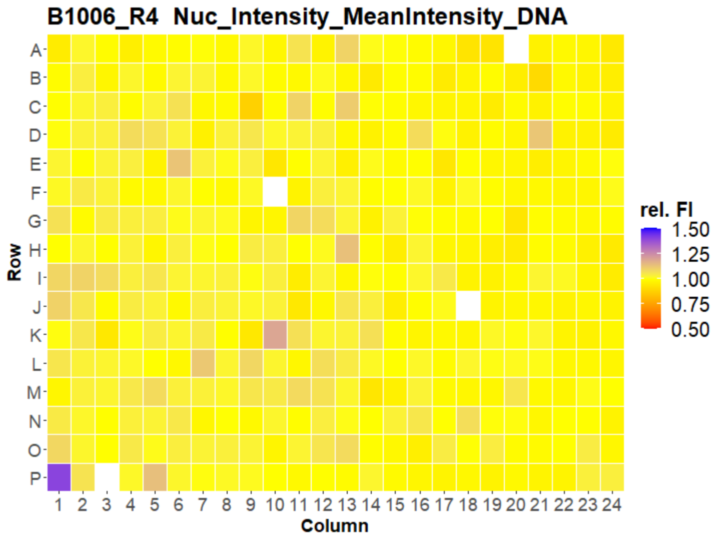
B1006_R4 Nuc_Intensity_MeanIntensity_DNA
Row
Column
rel. FI
1.50
1.25
1.00
0.75
0.50

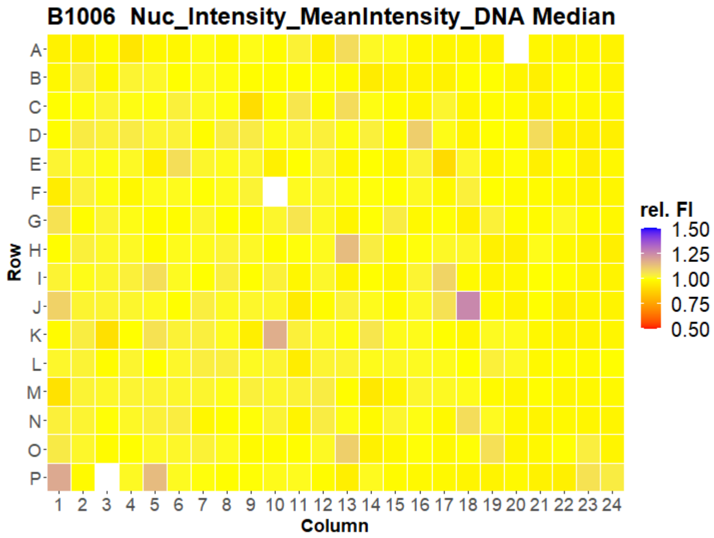
B1006 Nuc_Intensity_MeanIntensity_DNA Median
Row
Column
rel. FI
1.50
1.25
1.00
0.75
0.50

### U2OS FMP

35

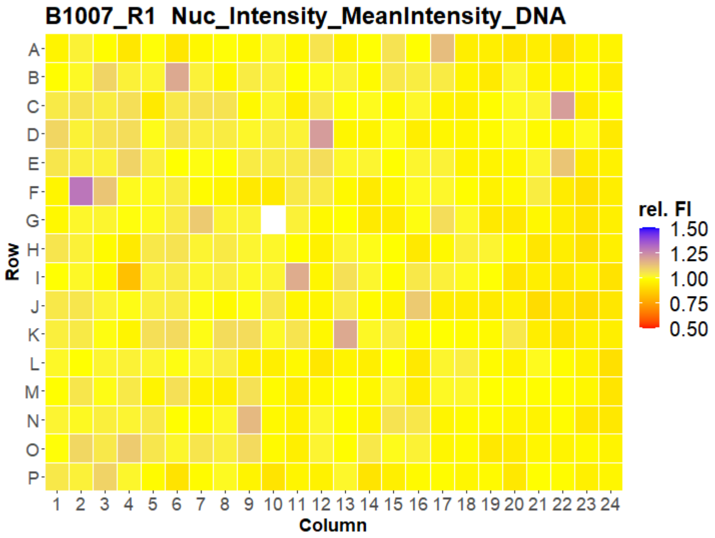
B1007_R1 Nuc_Intensity_MeanIntensity_DNA
Row
Column
rel. FI
1.50
1.25
1.00
0.75
0.50

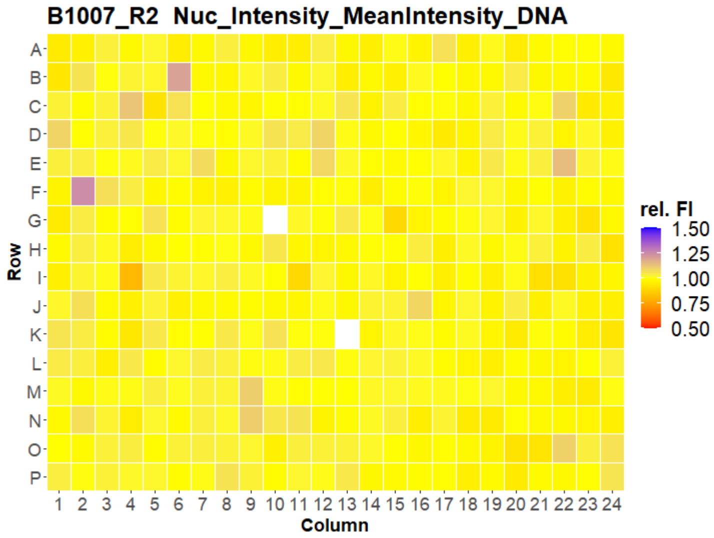
B1007_R2 Nuc_Intensity_MeanIntensity_DNA
Row
Column
rel. FI
1.50
1.25
1.00
0.75
0.50

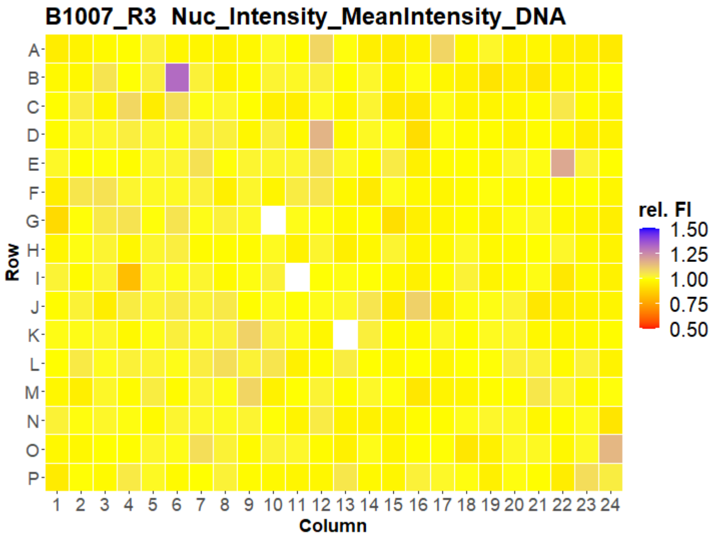
B1007_R3 Nuc_Intensity_MeanIntensity_DNA
Row
Column
rel. FI
1.50
1.25
1.00
0.75
0.50

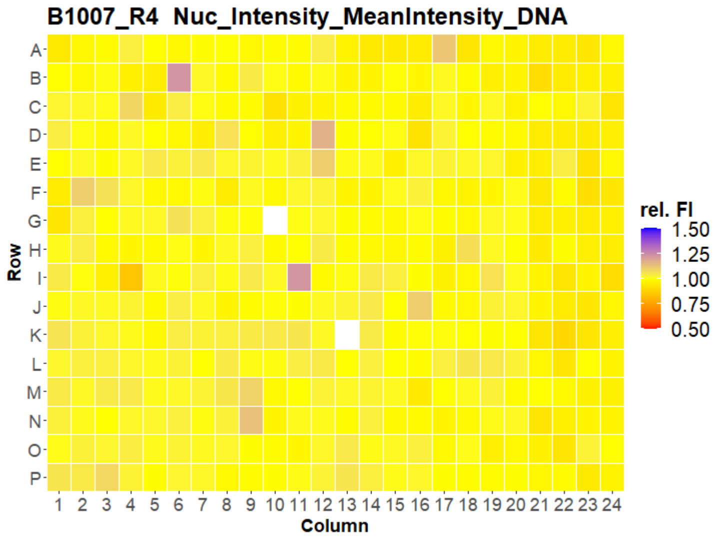
B1007_R4 Nuc_Intensity_MeanIntensity_DNA
Row
Column
rel. FI
1.50
1.25
1.00
0.75
0.50

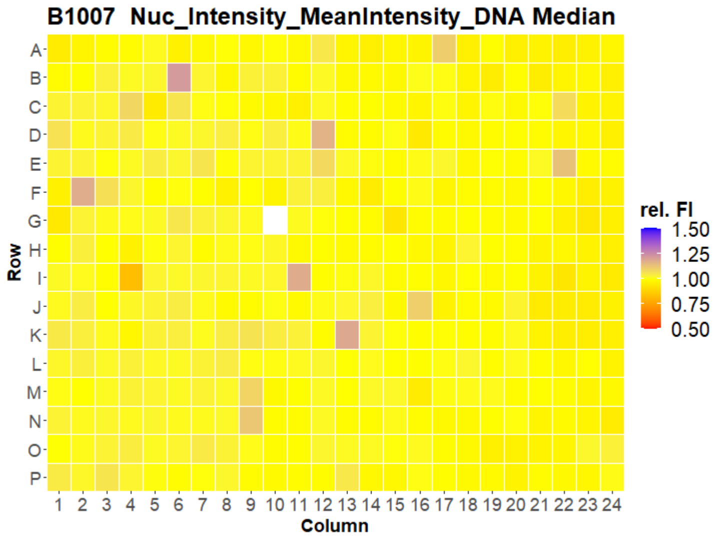
B1007 Nuc_Intensity_MeanIntensity_DNA Median
Row
Column
rel. FI
1.50
1.25
1.00
0.75
0.50

### U2OS FMP



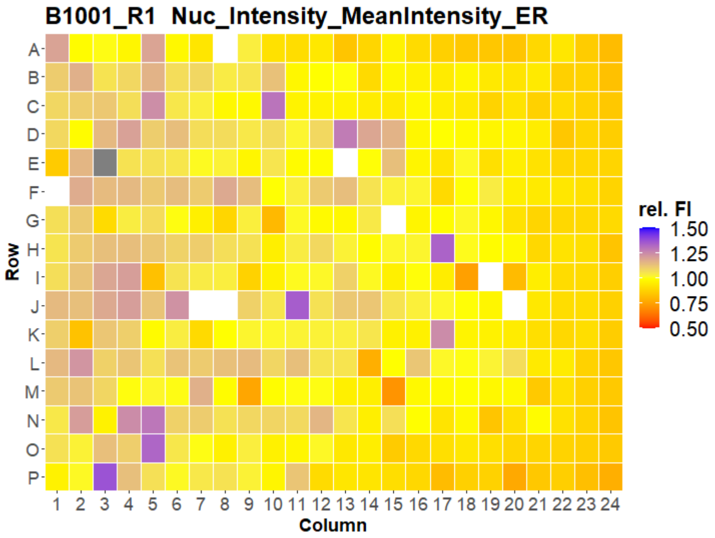
B1001_R1 Nuc_Intensity_MeanIntensity_ER
Row
Column
rel. FI
1.50
1.25
1.00
0.75
0.50

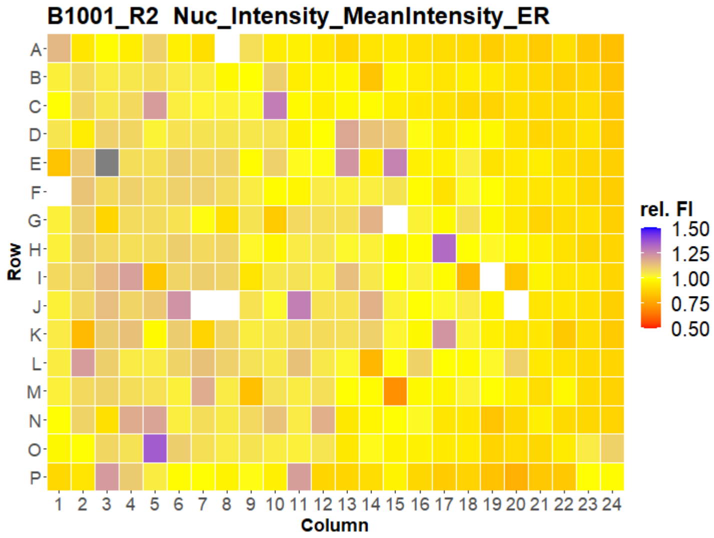
B1001_R2 Nuc_Intensity_MeanIntensity_ER
Row
Column
rel. FI
1.50
1.25
1.00
0.75
0.50

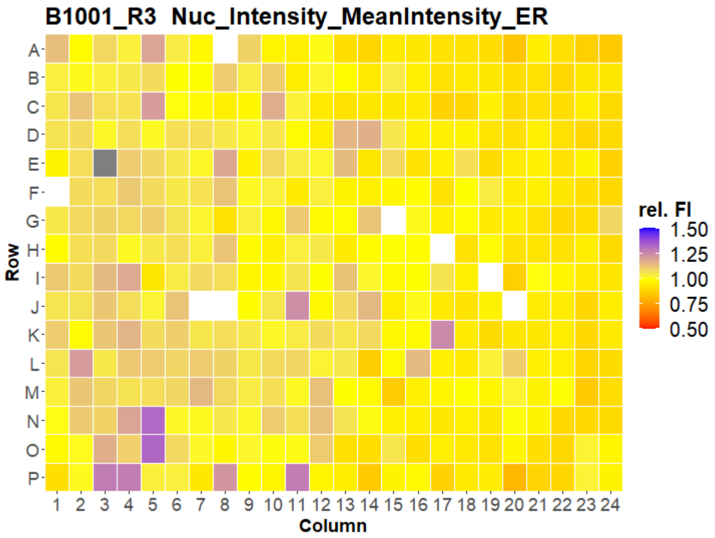
B1001_R3 Nuc_Intensity_MeanIntensity_ER
Row
Column
rel. FI
1.50
1.25
1.00
0.75
0.50

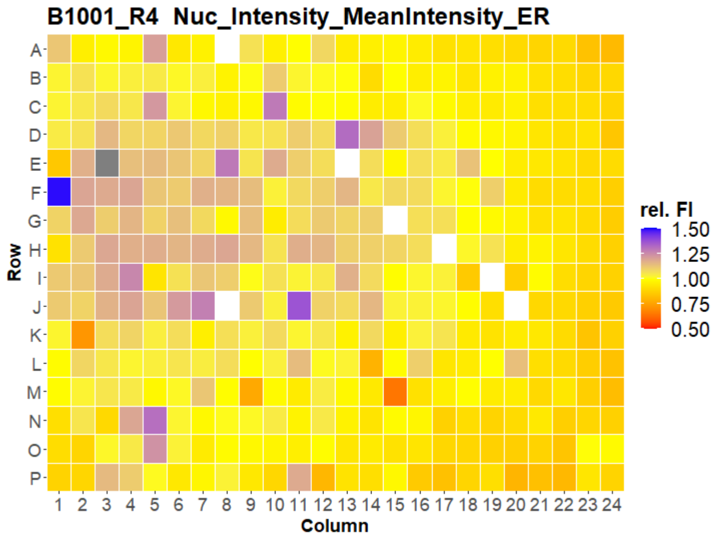
B1001_R4 Nuc_Intensity_MeanIntensity_ER
Row
Column
rel. FI
1.50
1.25
1.00
0.75
0.50

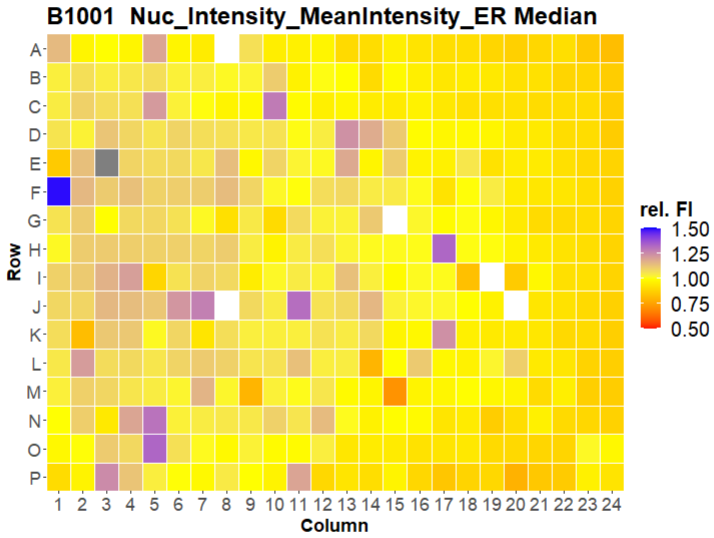
B1001 Nuc_Intensity_MeanIntensity_ER Median
Row
Column
rel. FI
1.50
1.25
1.00
0.75
0.50

### U2OS FMP

37

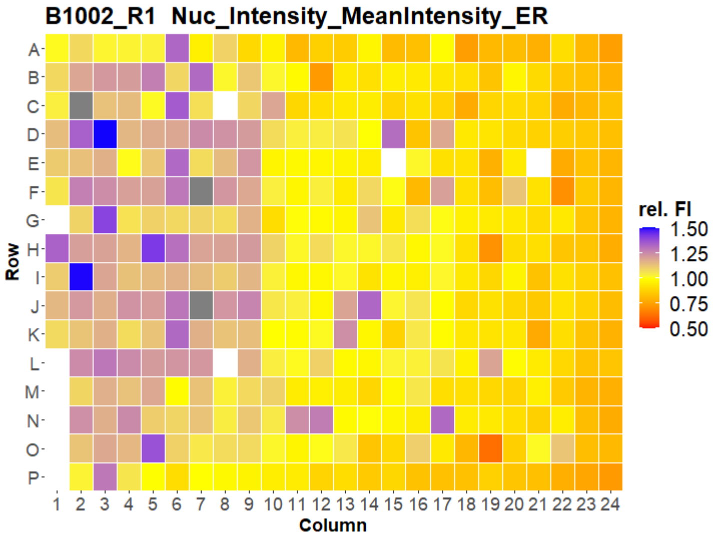
B1002_R1 Nuc_Intensity_MeanIntensity_ER
rel. FI
1.50
1.25
1.00
0.75
0.50
Row
Column

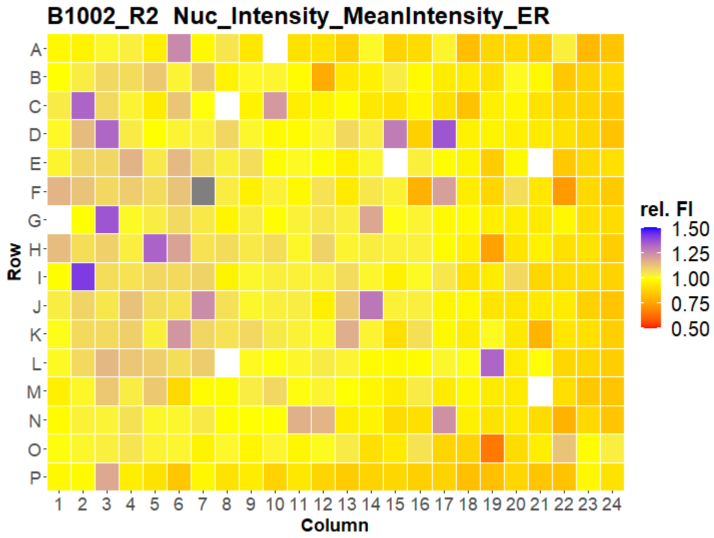
B1002_R2 Nuc_Intensity_MeanIntensity_ER
rel. FI
1.50
1.25
1.00
0.75
0.50
Row
Column

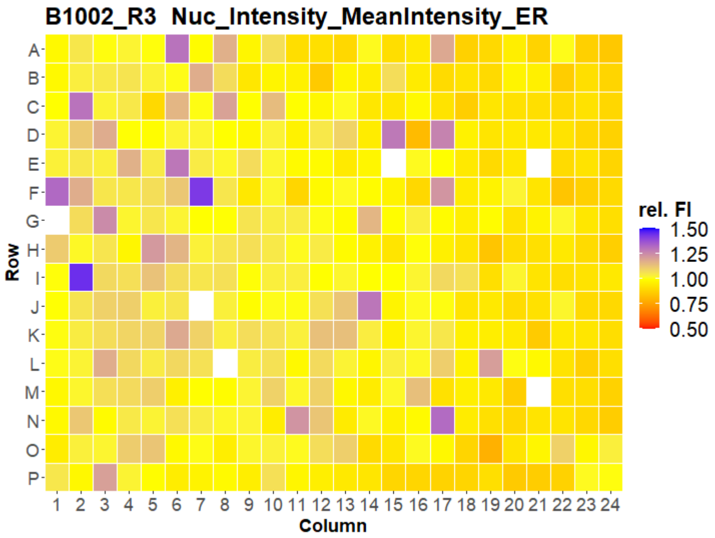
B1002_R3 Nuc_Intensity_MeanIntensity_ER
rel. FI
1.50
1.25
1.00
0.75
0.50
Row
Column

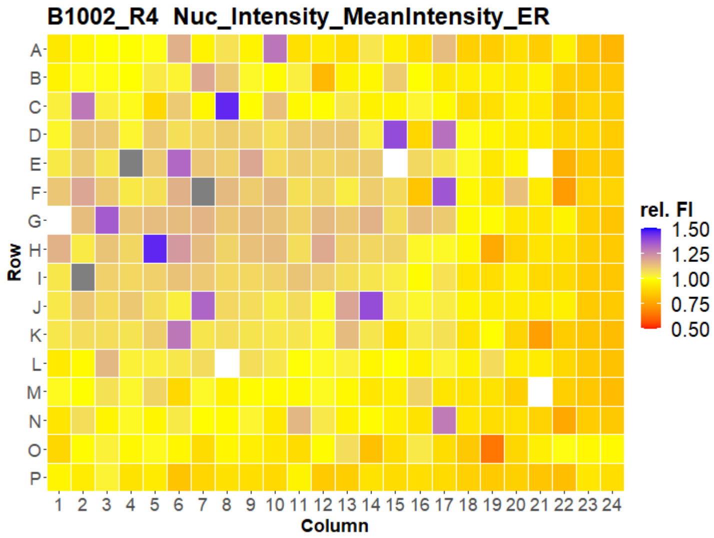
B1002_R4 Nuc_Intensity_MeanIntensity_ER
rel. FI
1.50
1.25
1.00
0.75
0.50
Row
Column

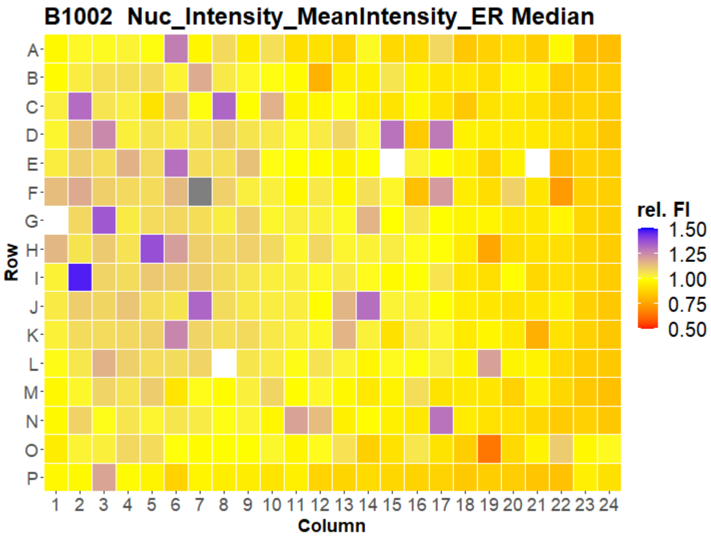
B1002 Nuc_Intensity_MeanIntensity_ER Median
rel. FI
1.50
1.25
1.00
0.75
0.50
Row
Column

### U2OS FMP



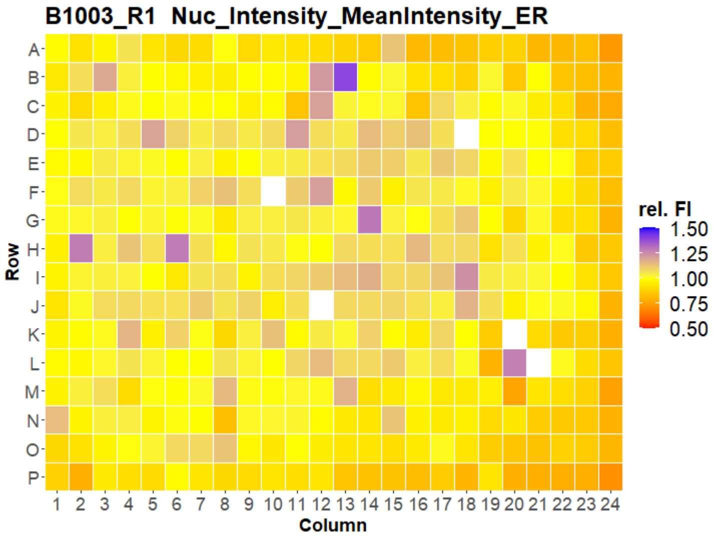
B1003_R1 Nuc_Intensity_MeanIntensity_ER
Row
Column
rel. FI
1.50
1.25
1.00
0.75
0.50

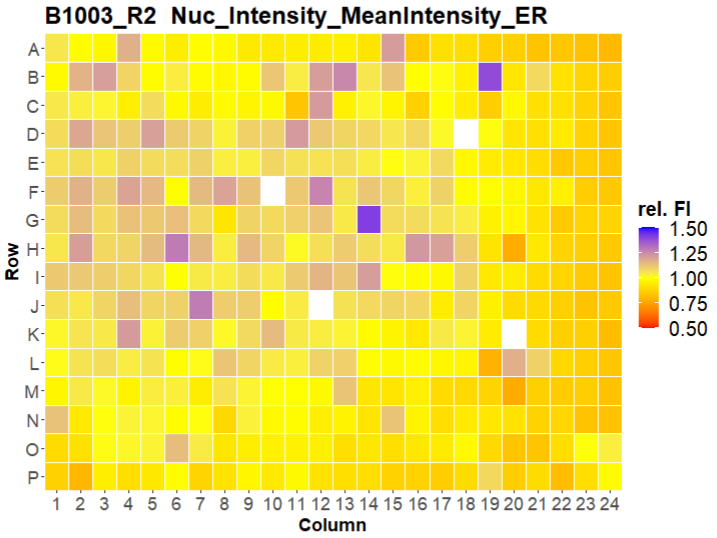
B1003_R2 Nuc_Intensity_MeanIntensity_ER
Row
Column
rel. FI
1.50
1.25
1.00
0.75
0.50

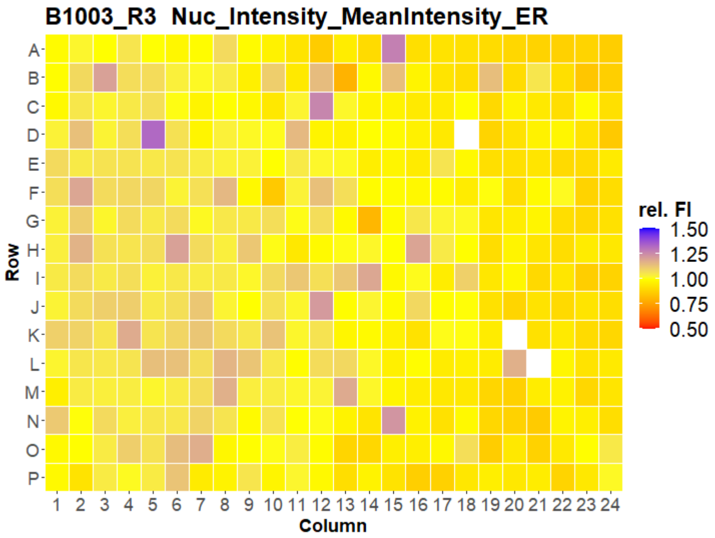
B1003_R3 Nuc_Intensity_MeanIntensity_ER
Row
Column
rel. FI
1.50
1.25
1.00
0.75
0.50

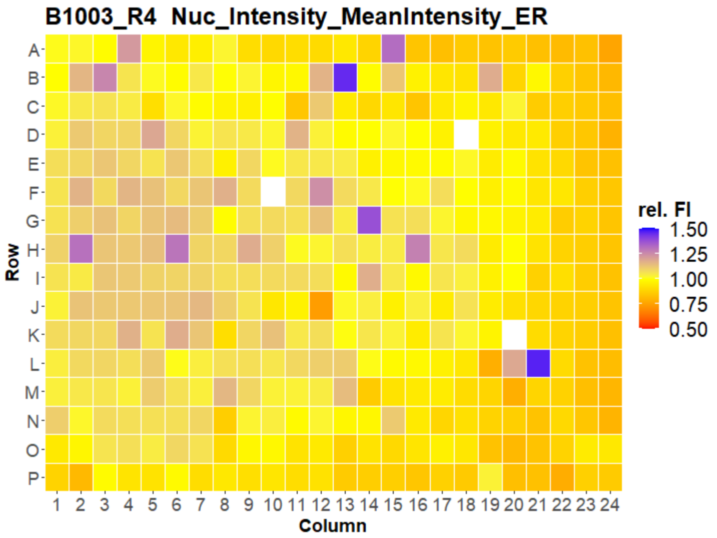
B1003_R4 Nuc_Intensity_MeanIntensity_ER
Row
Column
rel. FI
1.50
1.25
1.00
0.75
0.50

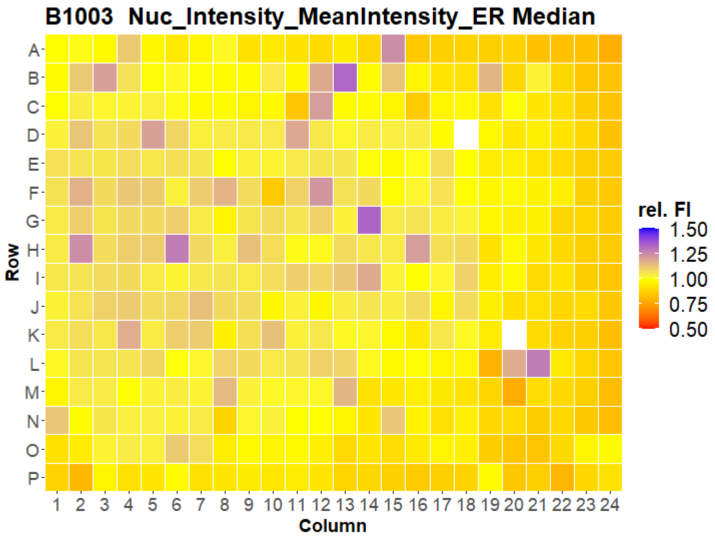
B1003 Nuc_Intensity_MeanIntensity_ER Median
Row
Column
rel. FI
1.50
1.25
1.00
0.75
0.50

### U2OS FMP

39

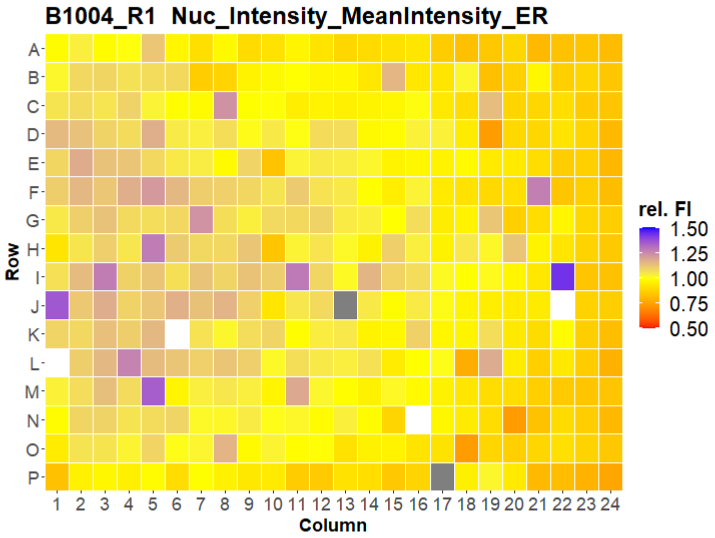
B1004_R1 Nuc_Intensity_MeanIntensity_ER
Row
Column
rel. FI
1.50
1.25
1.00
0.75
0.50

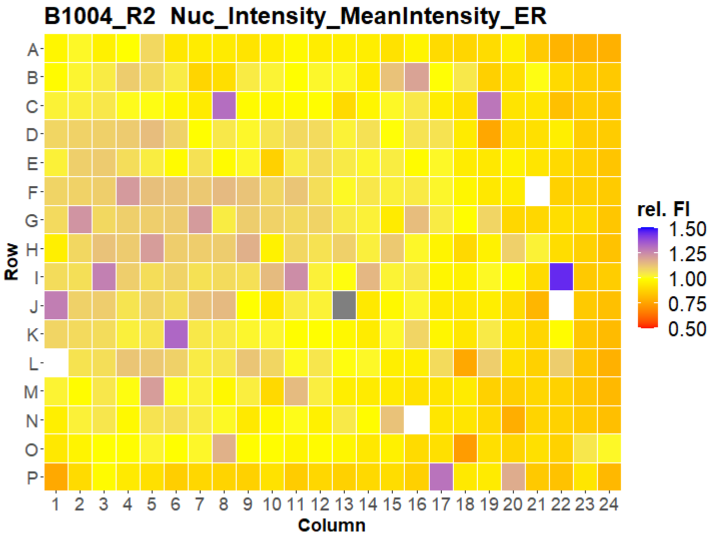
B1004_R2 Nuc_Intensity_MeanIntensity_ER
Row
Column
rel. FI
1.50
1.25
1.00
0.75
0.50

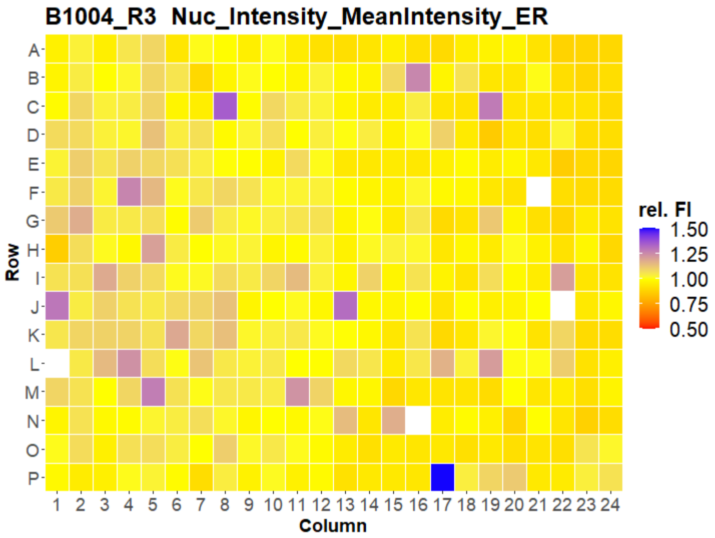
B1004_R3 Nuc_Intensity_MeanIntensity_ER
Row
Column
rel. FI
1.50
1.25
1.00
0.75
0.50

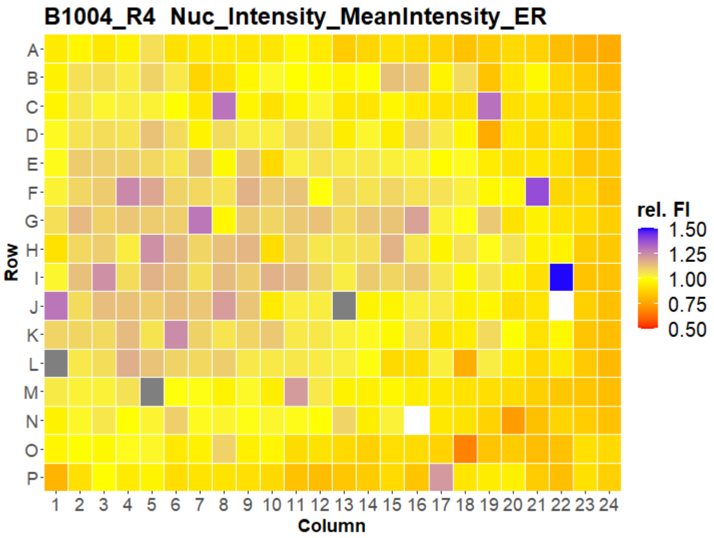
B1004_R4 Nuc_Intensity_MeanIntensity_ER
Row
Column
rel. FI
1.50
1.25
1.00
0.75
0.50

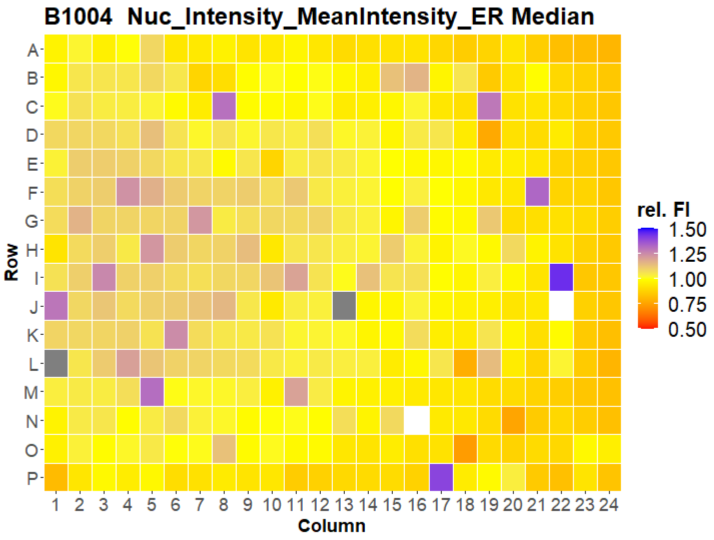
B1004 Nuc_Intensity_MeanIntensity_ER Median
Row
Column
rel. FI
1.50
1.25
1.00
0.75
0.50

### U2OS FMP



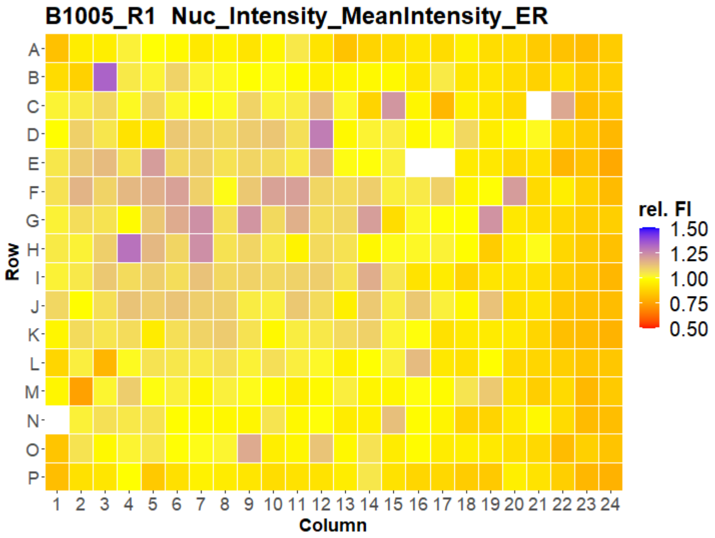


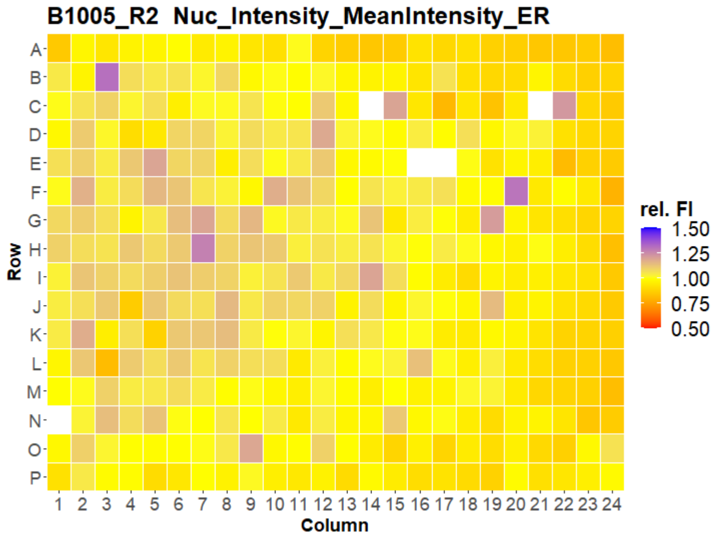


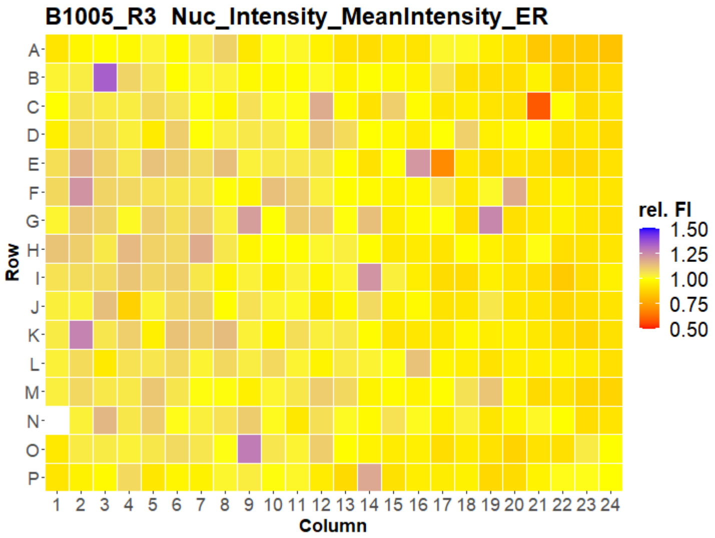


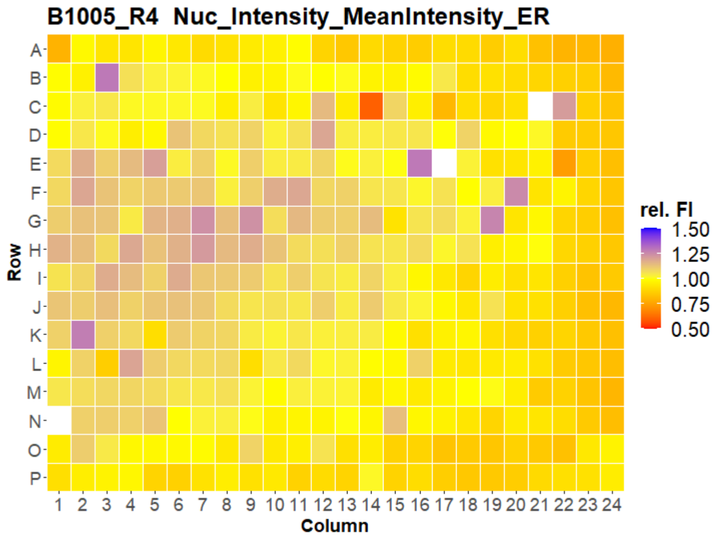


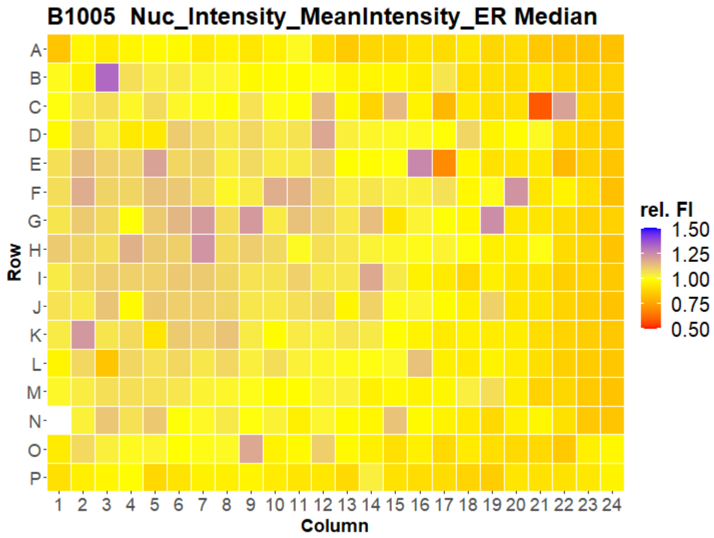




Open for Innovation KNIME

### U2OS FMP



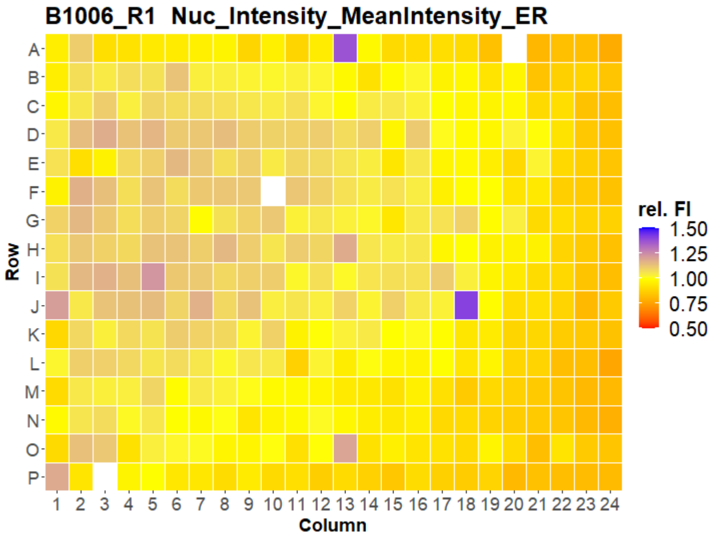
B1006_R1 Nuc_Intensity_MeanIntensity_ER
Row
Column
rel. FI
1.50
1.25
1.00
0.75
0.50

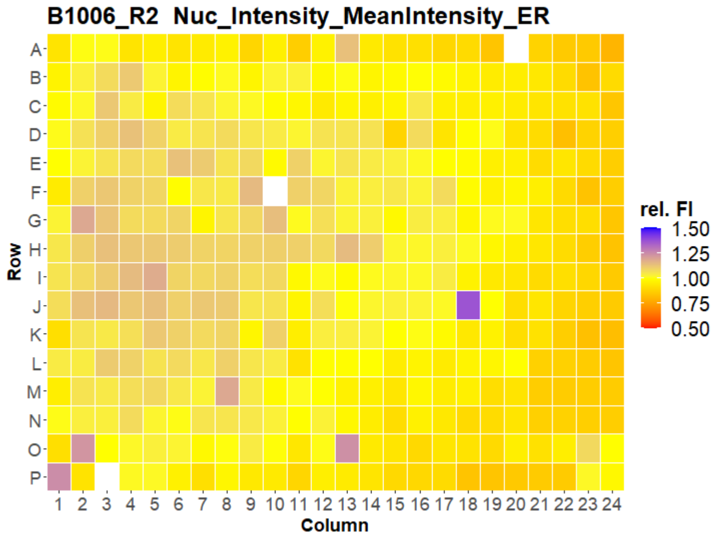
B1006_R2 Nuc_Intensity_MeanIntensity_ER
Row
Column
rel. FI
1.50
1.25
1.00
0.75
0.50

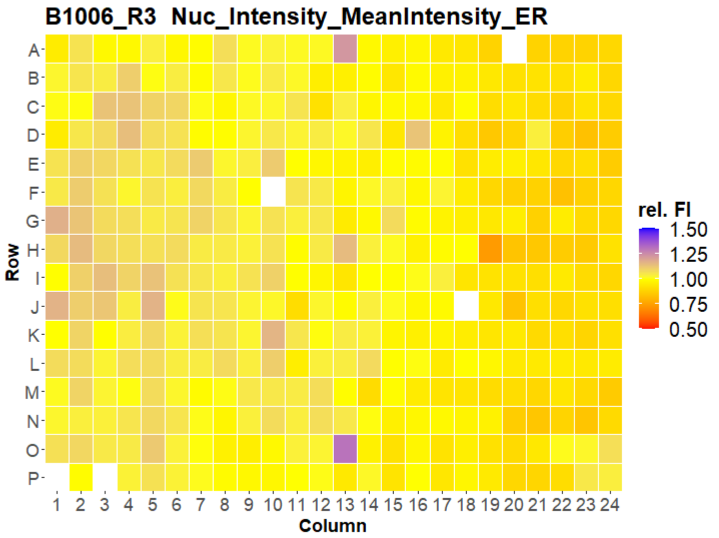
B1006_R3 Nuc_Intensity_MeanIntensity_ER
Row
Column
rel. FI
1.50
1.25
1.00
0.75
0.50

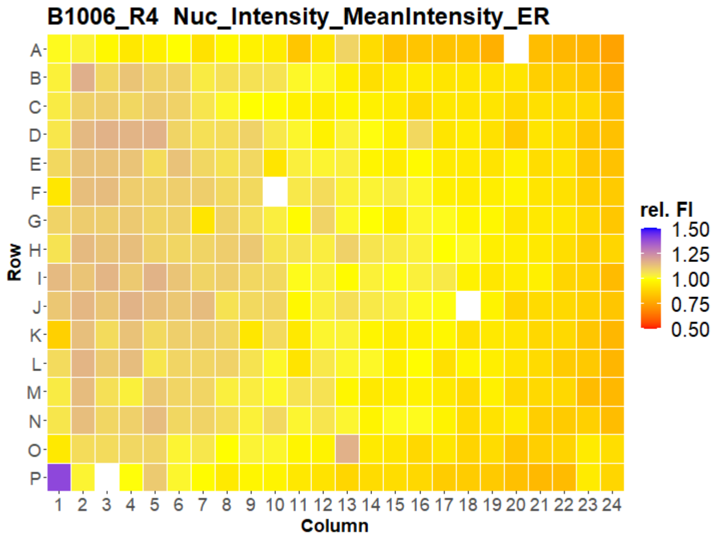
B1006_R4 Nuc_Intensity_MeanIntensity_ER
Row
Column
rel. FI
1.50
1.25
1.00
0.75
0.50

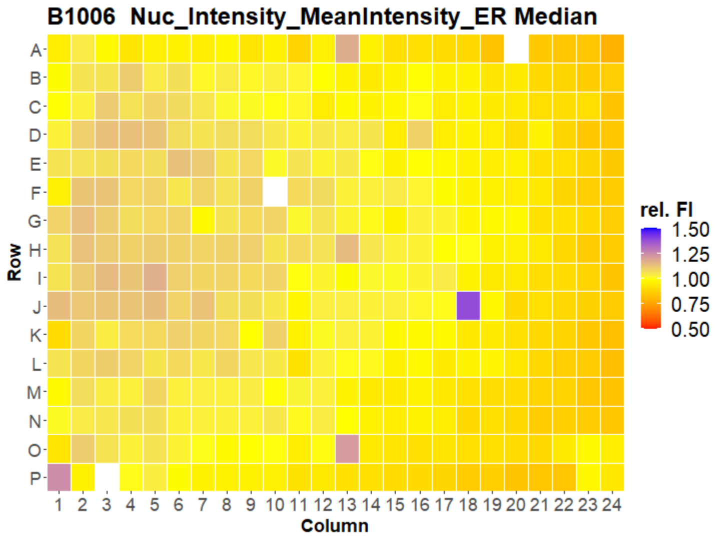
B1006 Nuc_Intensity_MeanIntensity_ER Median
Row
Column
rel. FI
1.50
1.25
1.00
0.75
0.50

### U2OS FMP



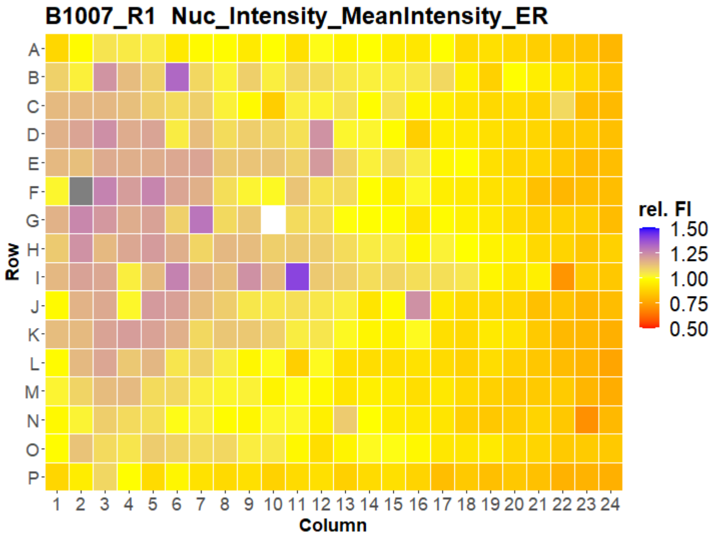
B1007_R1 Nuc_Intensity_MeanIntensity_ER
Row
Column
rel. FI
1.50
1.25
1.00
0.75
0.50

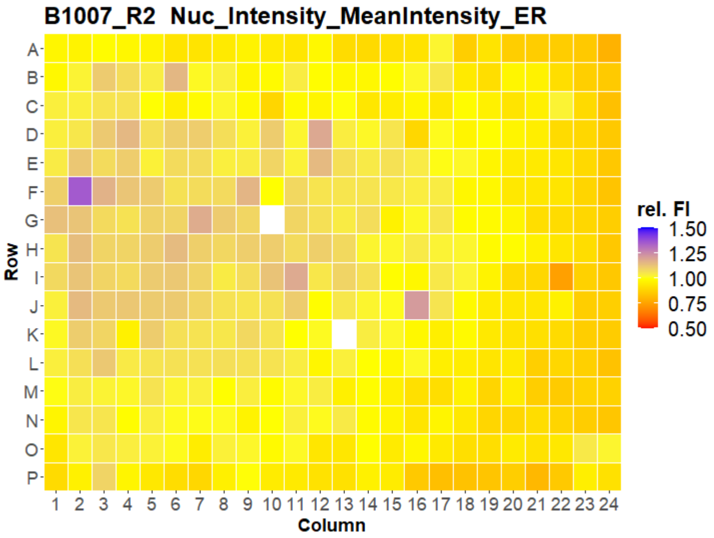
B1007_R2 Nuc_Intensity_MeanIntensity_ER
Row
Column
rel. FI
1.50
1.25
1.00
0.75
0.50

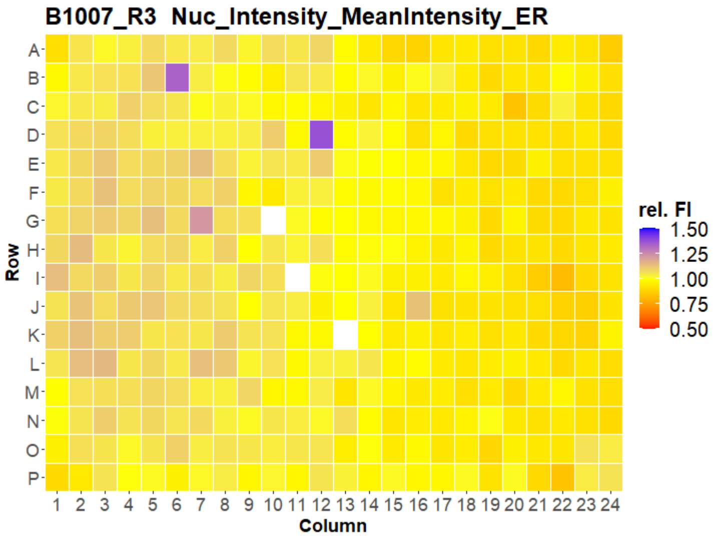
B1007_R3 Nuc_Intensity_MeanIntensity_ER
Row
Column
rel. FI
1.50
1.25
1.00
0.75
0.50

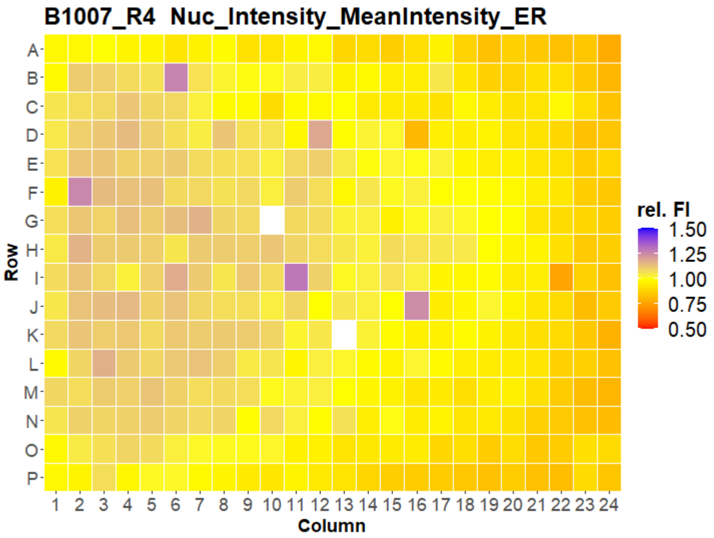
B1007_R4 Nuc_Intensity_MeanIntensity_ER
Row
Column
rel. FI
1.50
1.25
1.00
0.75
0.50

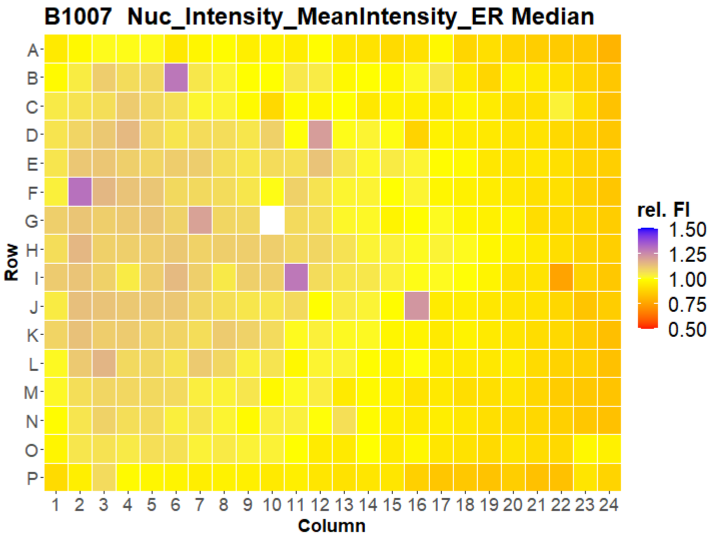
B1007 Nuc_Intensity_MeanIntensity_ER Median
Row
Column
rel. FI
1.50
1.25
1.00
0.75
0.50

### U2OS FMP
