## Supplementary material for "Morphological Profiling Dataset of EU-OPENSCREEN Bioactive Compounds Over Multiple Imaging Sites and Cell Lines": Suppl_QC_IMTM_HepG2.pdf

### HepG2 IMTM

1

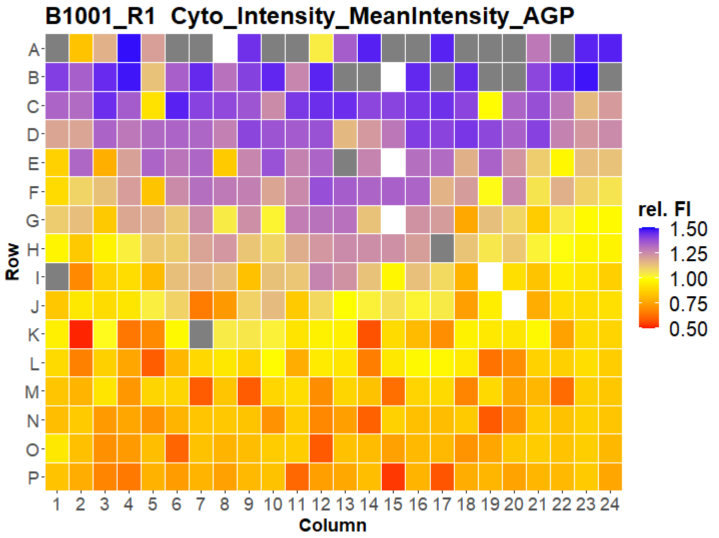


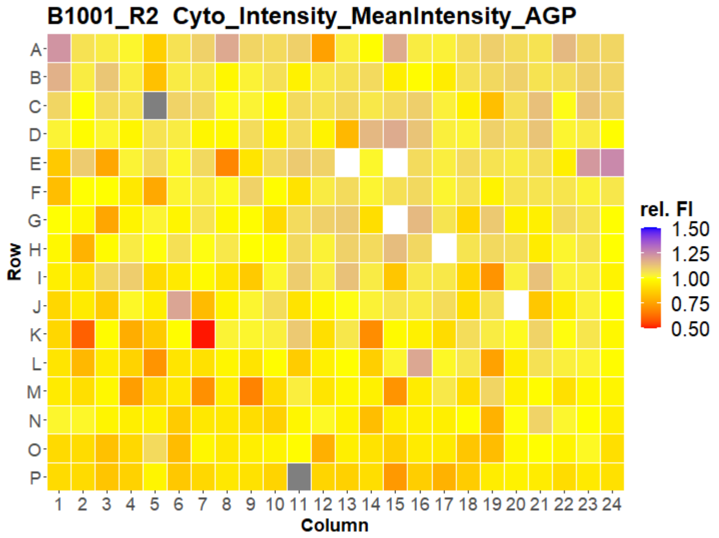


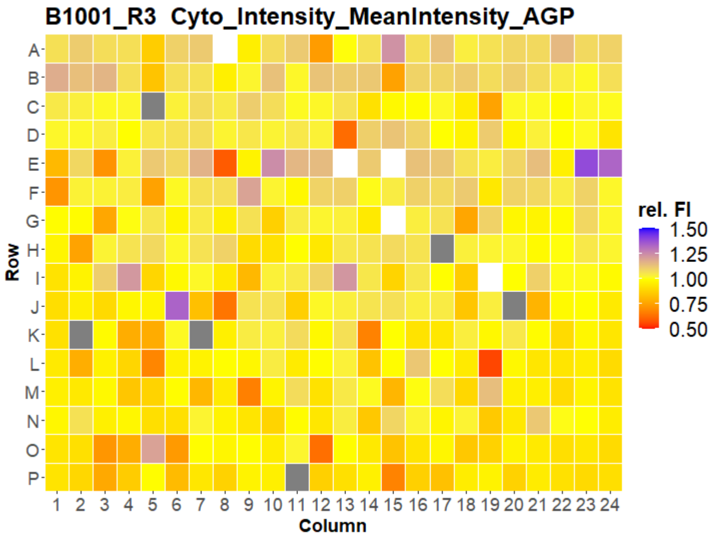


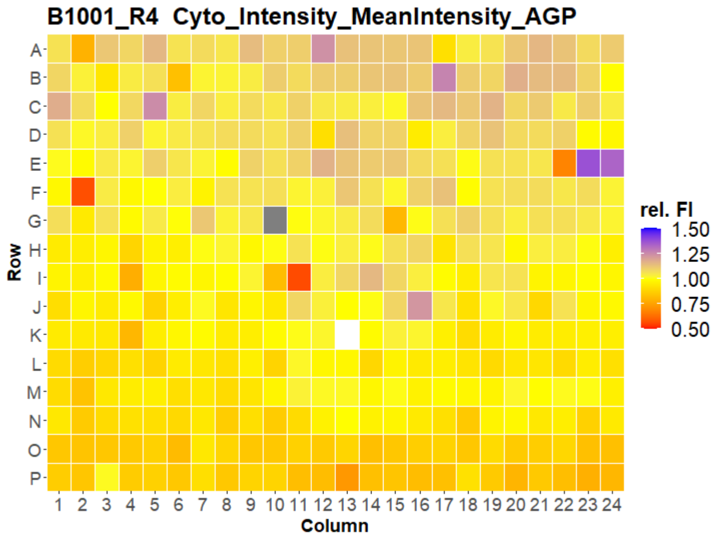


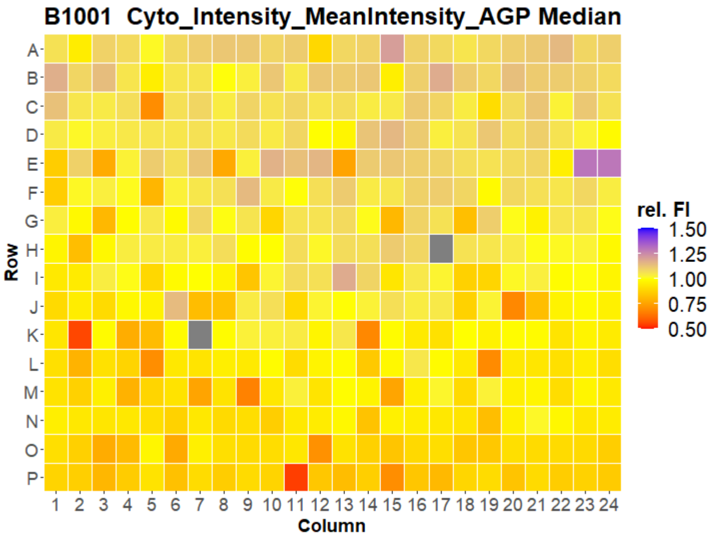



Open for Innovation KNIME

### HepG2 IMTM

2

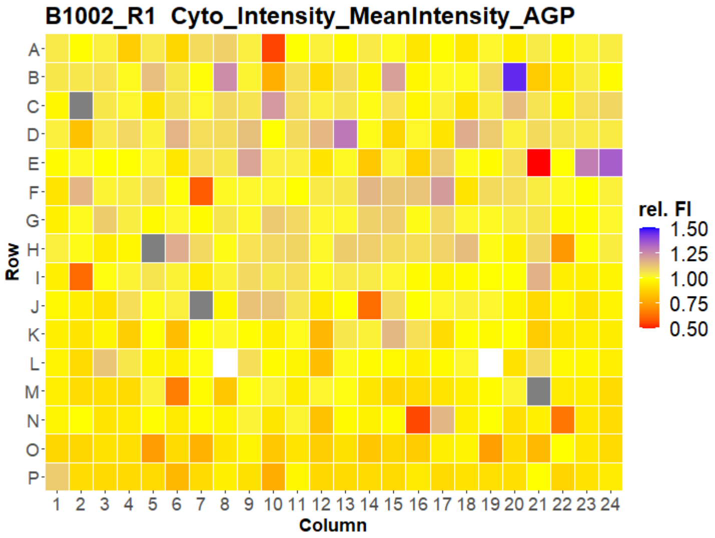
B1002_R1 Cyto_Intensity_MeanIntensity_AGP
Row
Column
rel. FI
1.50
1.25
1.00
0.75
0.50

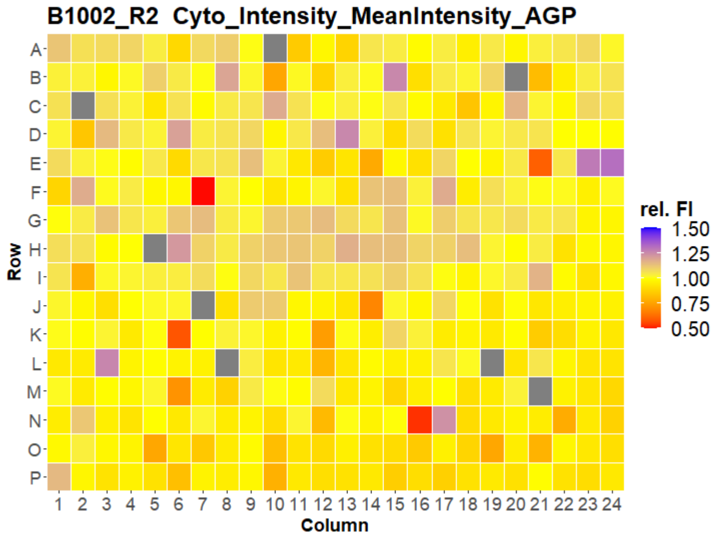
B1002_R2 Cyto_Intensity_MeanIntensity_AGP
Row
Column
rel. FI
1.50
1.25
1.00
0.75
0.50

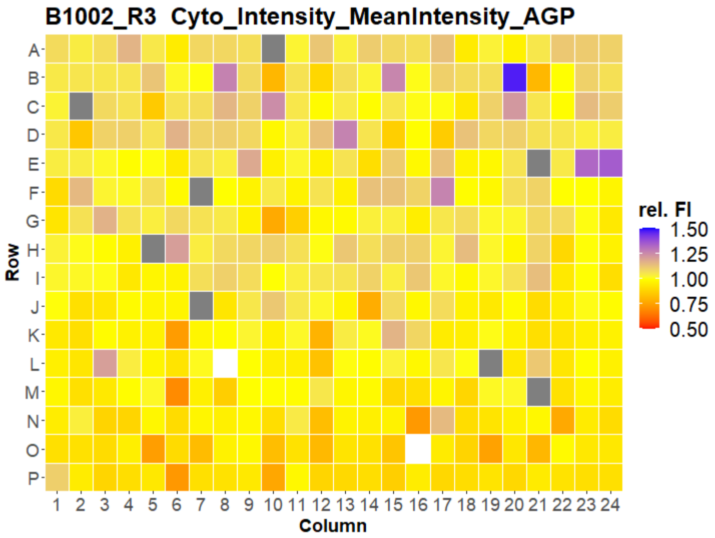
B1002_R3 Cyto_Intensity_MeanIntensity_AGP
Row
Column
rel. FI
1.50
1.25
1.00
0.75
0.50

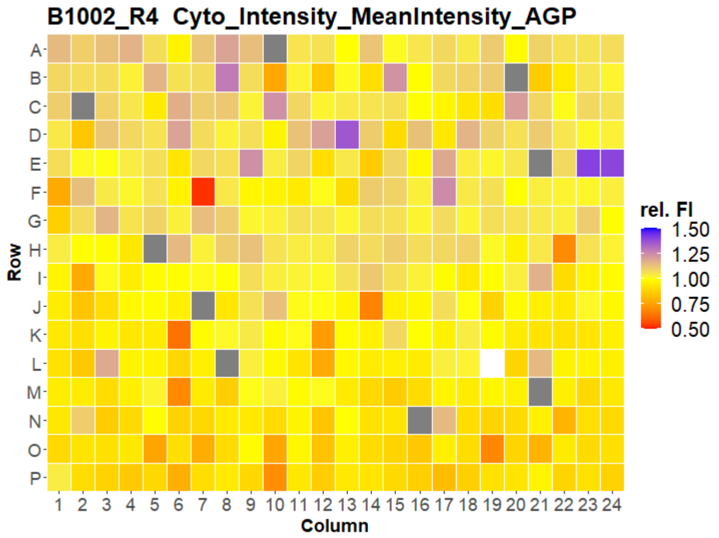
B1002_R4 Cyto_Intensity_MeanIntensity_AGP
Row
Column
rel. FI
1.50
1.25
1.00
0.75
0.50

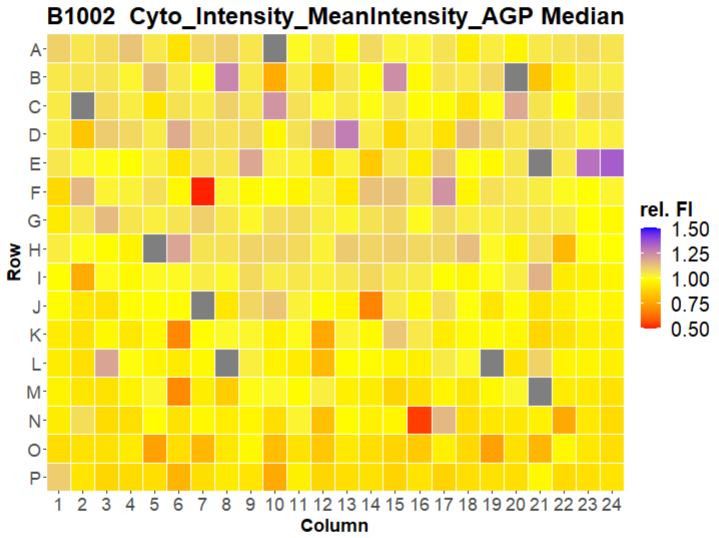
B1002 Cyto_Intensity_MeanIntensity_AGP Median
Row
Column
rel. FI
1.50
1.25
1.00
0.75
0.50

### HepG2 IMTM

3

B1003_R1  Cyto_Intensity_MeanIntensity_AGP

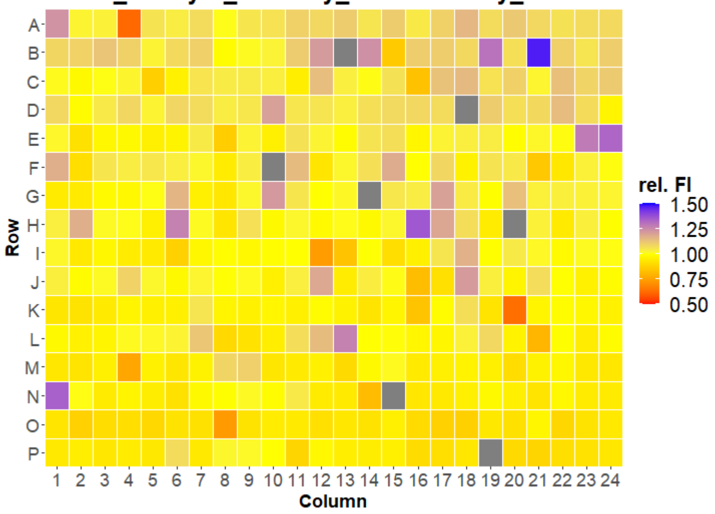


B1003_R2  Cyto_Intensity_MeanIntensity_AGP

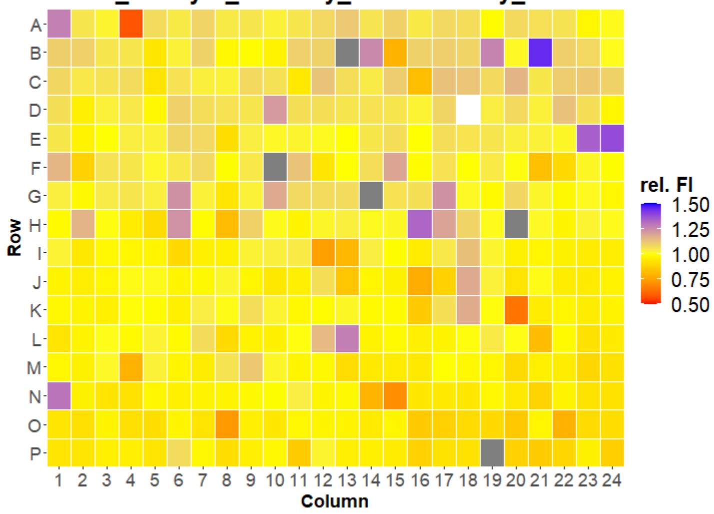


B1003_R3  Cyto_Intensity_MeanIntensity_AGP

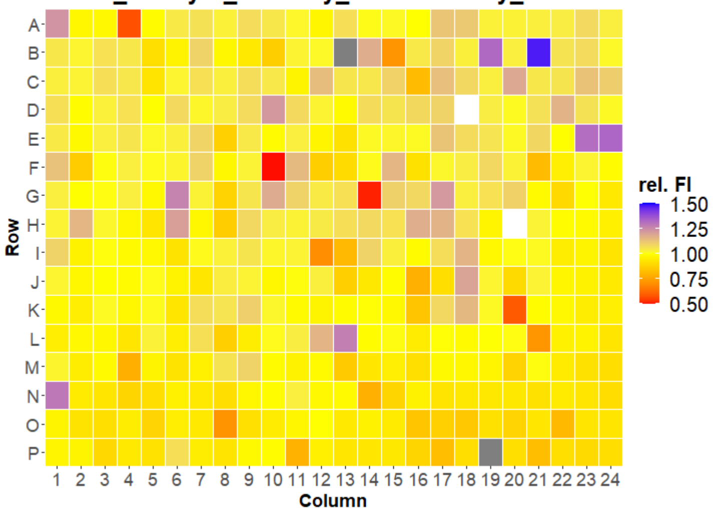


B1003_R4  Cyto_Intensity_MeanIntensity_AGP

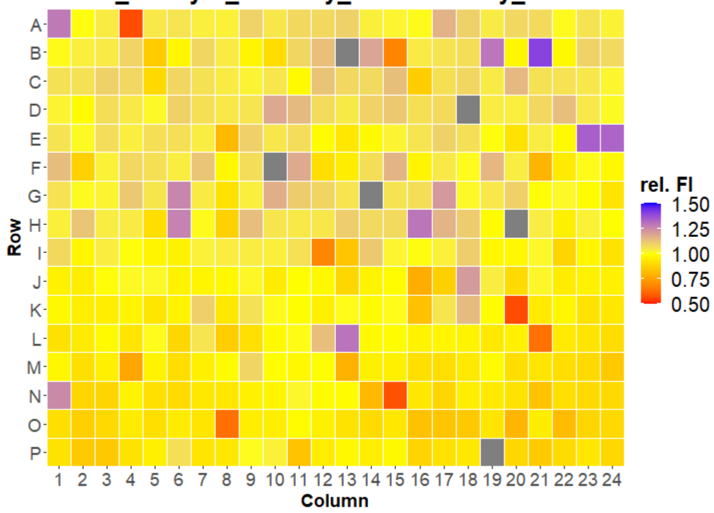


B1003  Cyto_Intensity_MeanIntensity_AGP Median

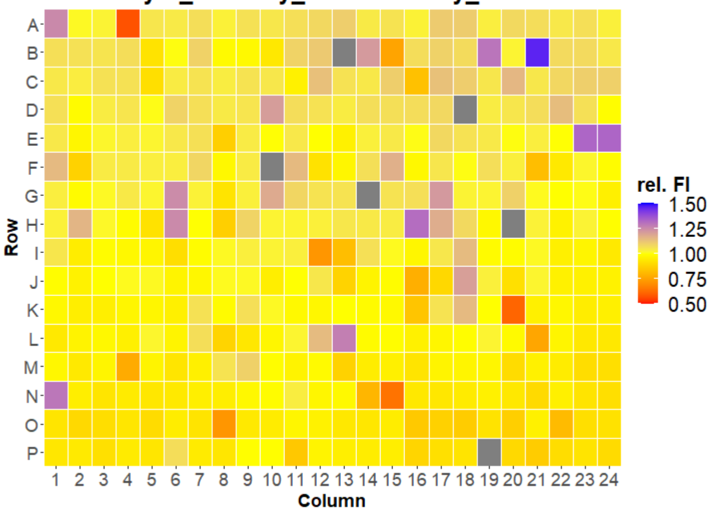

### HepG2 IMTM



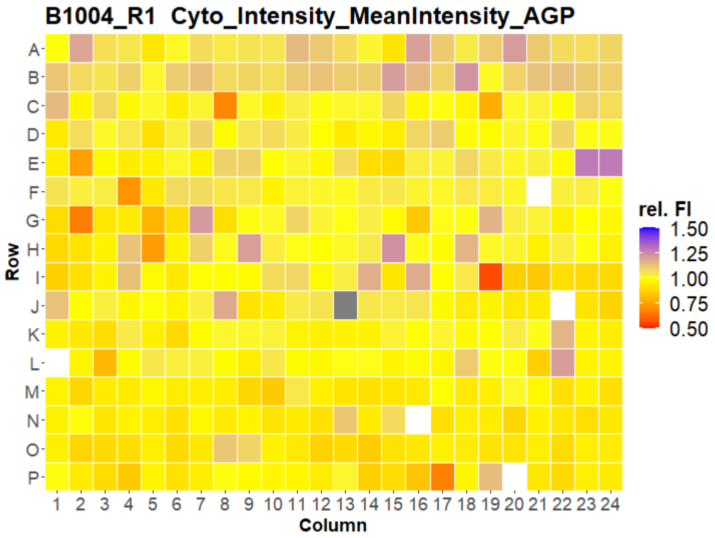
B1004_R1 Cyto_Intensity_MeanIntensity_AGP
Row
Column
rel. FI
1.50
1.25
1.00
0.75
0.50

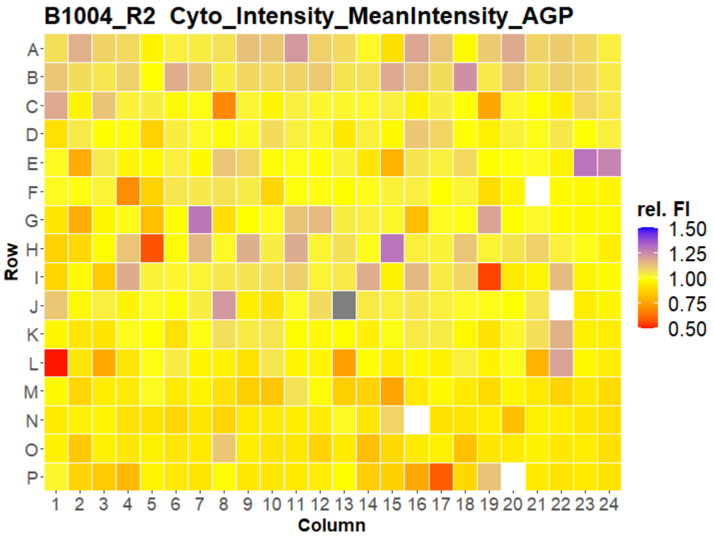
B1004_R2 Cyto_Intensity_MeanIntensity_AGP
Row
Column
rel. FI
1.50
1.25
1.00
0.75
0.50

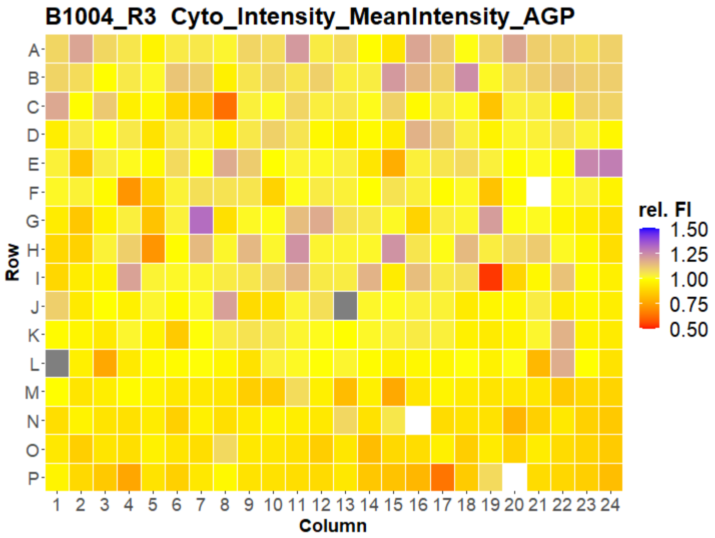
B1004_R3 Cyto_Intensity_MeanIntensity_AGP
Row
Column
rel. FI
1.50
1.25
1.00
0.75
0.50

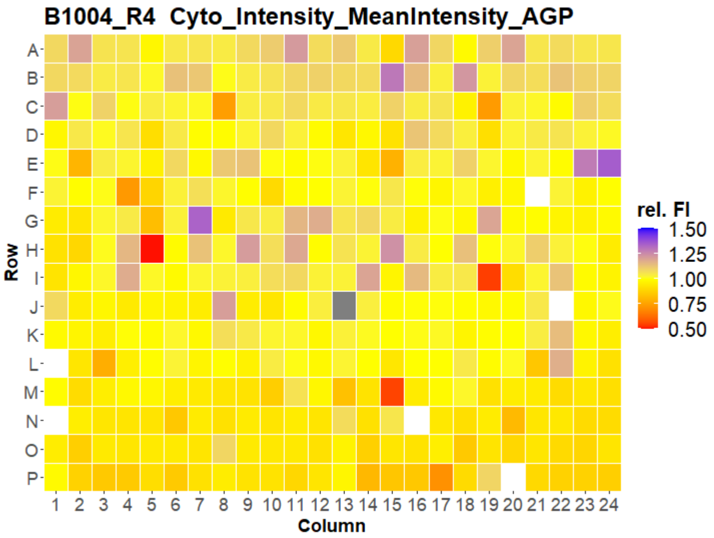
B1004_R4 Cyto_Intensity_MeanIntensity_AGP
Row
Column
rel. FI
1.50
1.25
1.00
0.75
0.50

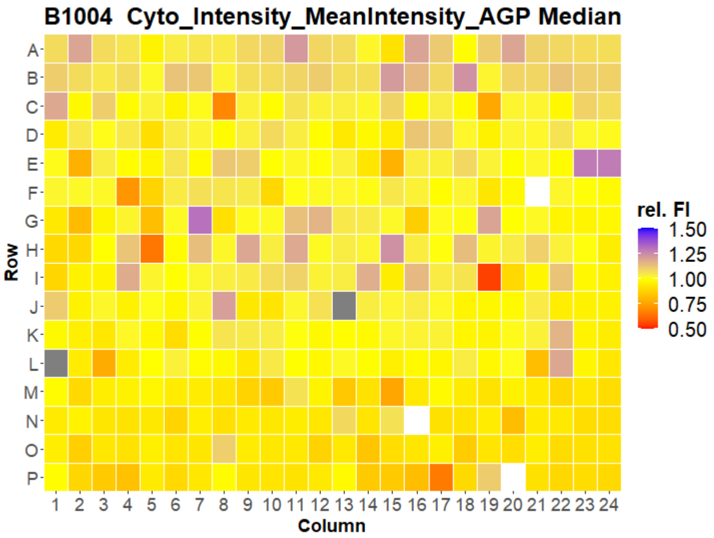
B1004 Cyto_Intensity_MeanIntensity_AGP Median
Row
Column
rel. FI
1.50
1.25
1.00
0.75
0.50

### HepG2 IMTM



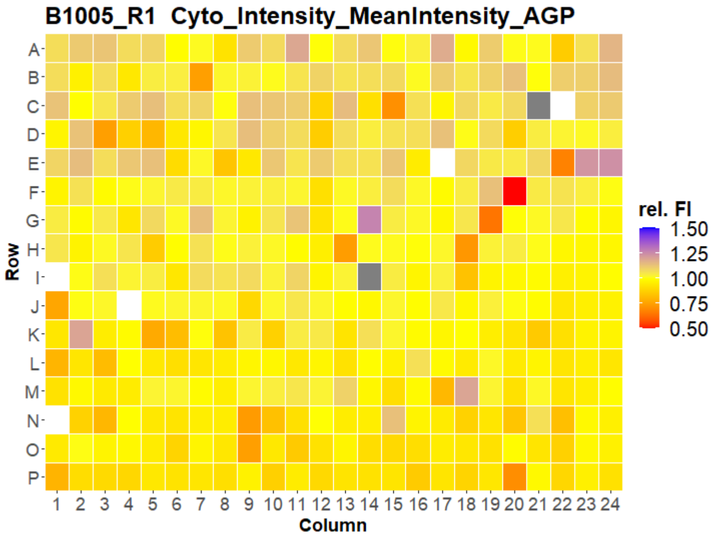
B1005_R1 Cyto_Intensity_MeanIntensity_AGP
Row
Column
rel. FI
1.50
1.25
1.00
0.75
0.50

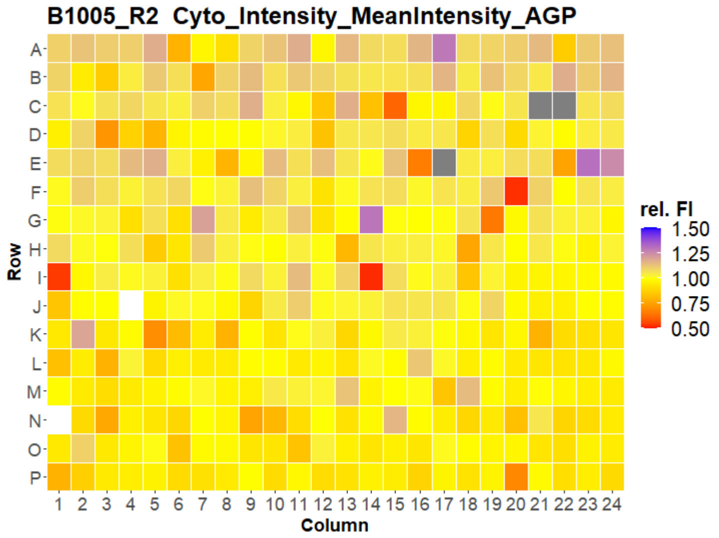
B1005_R2 Cyto_Intensity_MeanIntensity_AGP
Row
Column
rel. FI
1.50
1.25
1.00
0.75
0.50

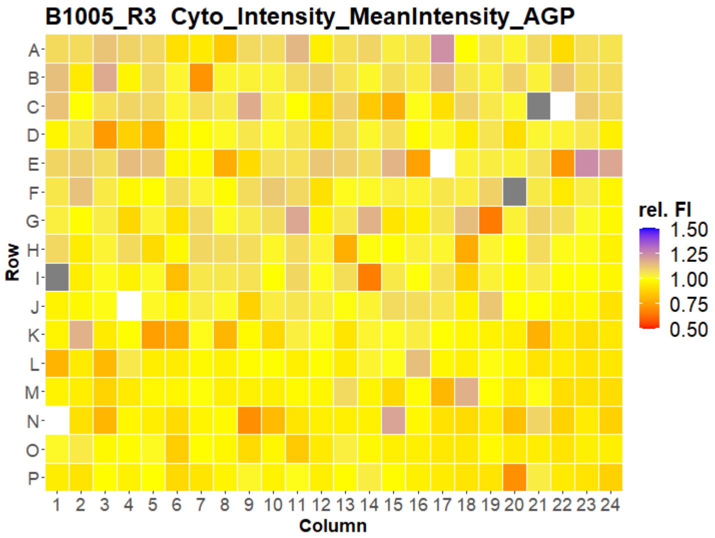
B1005_R3 Cyto_Intensity_MeanIntensity_AGP
Row
Column
rel. FI
1.50
1.25
1.00
0.75
0.50

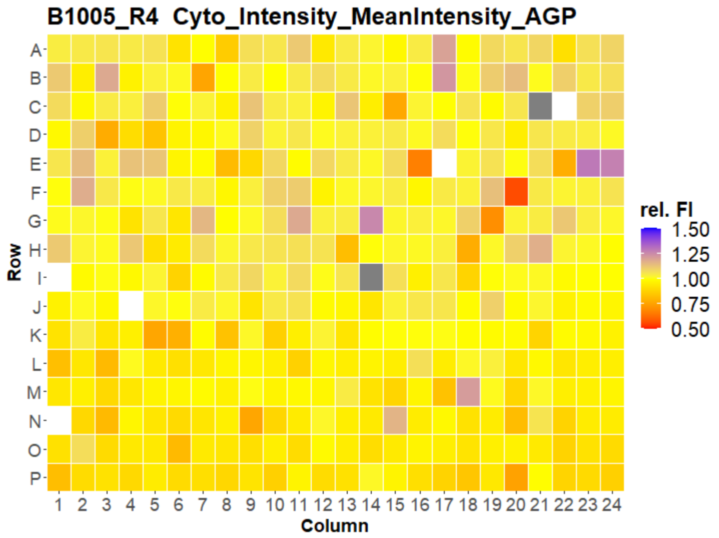
B1005_R4 Cyto_Intensity_MeanIntensity_AGP
Row
Column
rel. FI
1.50
1.25
1.00
0.75
0.50

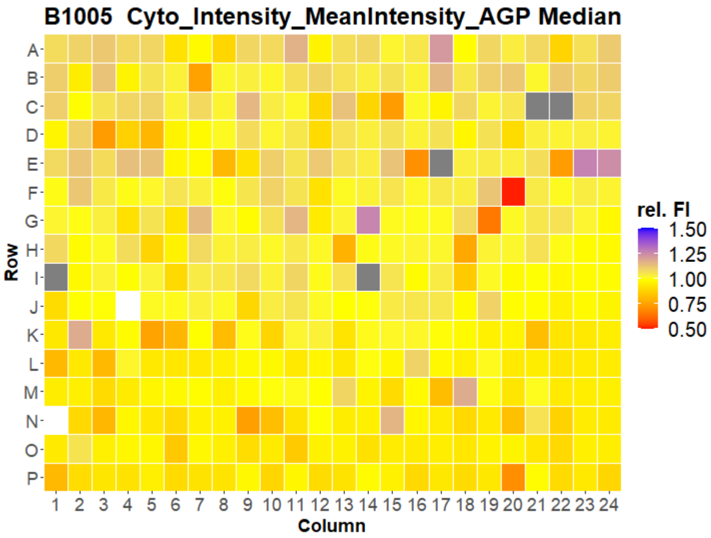
B1005 Cyto_Intensity_MeanIntensity_AGP Median
Row
Column
rel. FI
1.50
1.25
1.00
0.75
0.50

### HepG2 IMTM



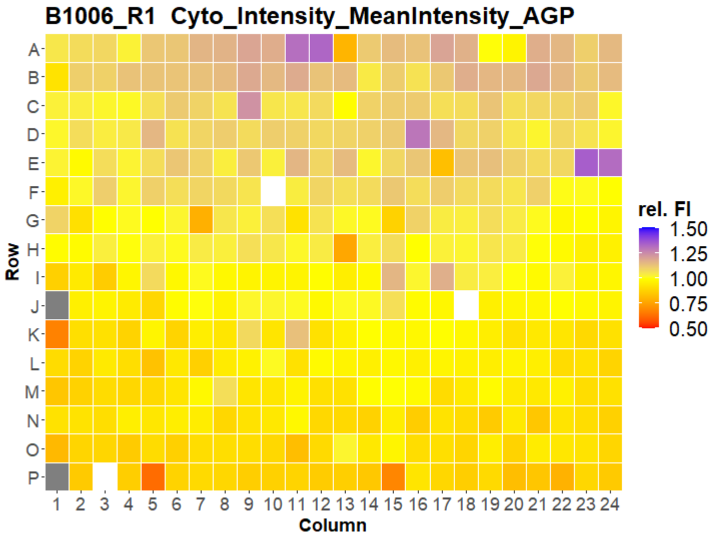
B1006_R1 Cyto_Intensity_MeanIntensity_AGP
Row
Column
rel. FI
1.50
1.25
1.00
0.75
0.50

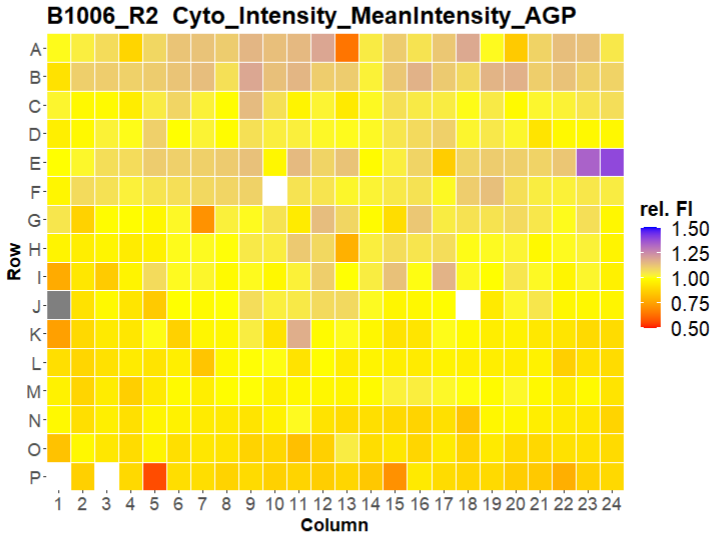
B1006_R2 Cyto_Intensity_MeanIntensity_AGP
Row
Column
rel. FI
1.50
1.25
1.00
0.75
0.50

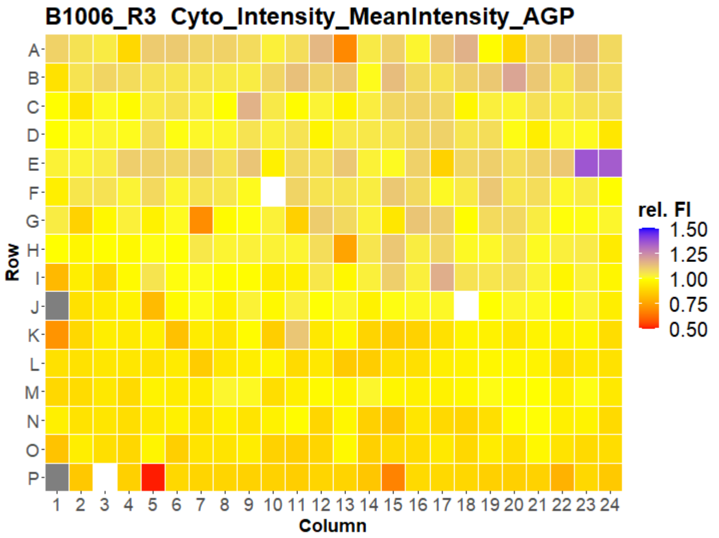
B1006_R3 Cyto_Intensity_MeanIntensity_AGP
Row
Column
rel. FI
1.50
1.25
1.00
0.75
0.50

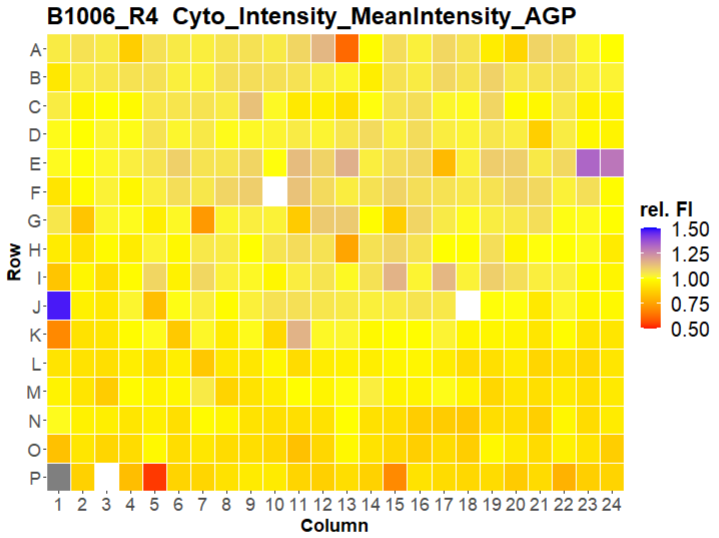
B1006_R4 Cyto_Intensity_MeanIntensity_AGP
Row
Column
rel. FI
1.50
1.25
1.00
0.75
0.50

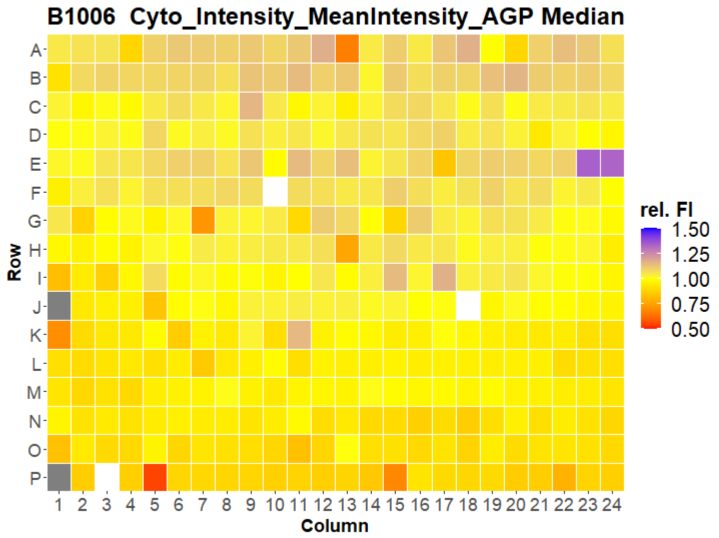
B1006 Cyto_Intensity_MeanIntensity_AGP Median
Row
Column
rel. FI
1.50
1.25
1.00
0.75
0.50

### HepG2 IMTM



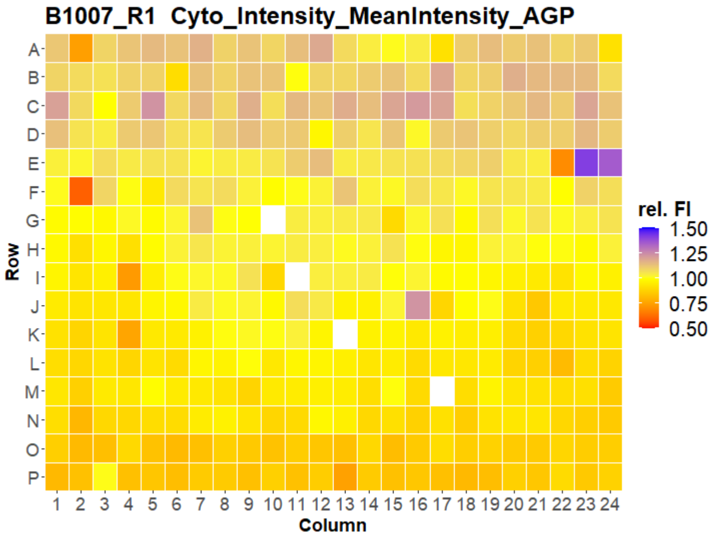


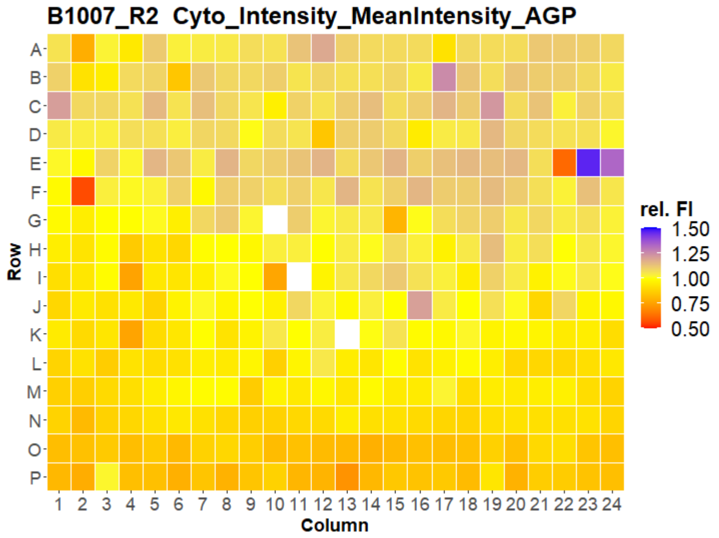


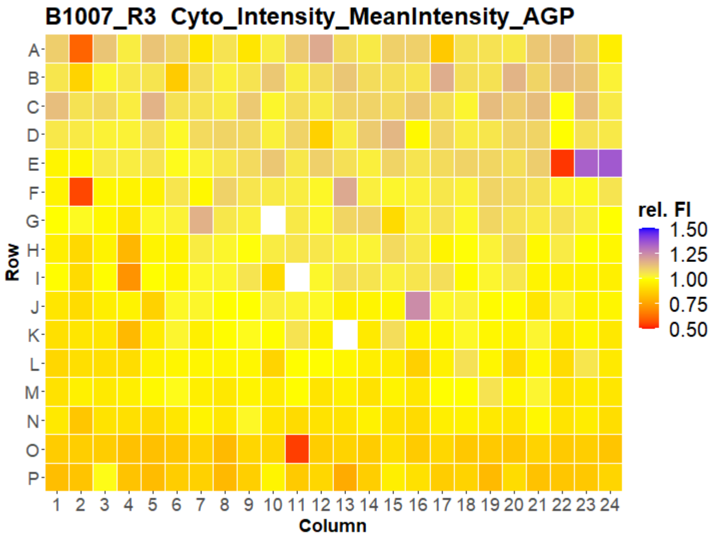


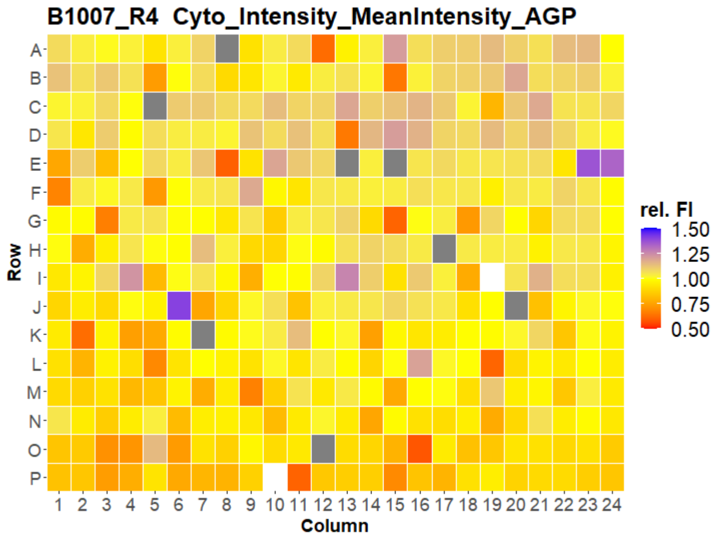


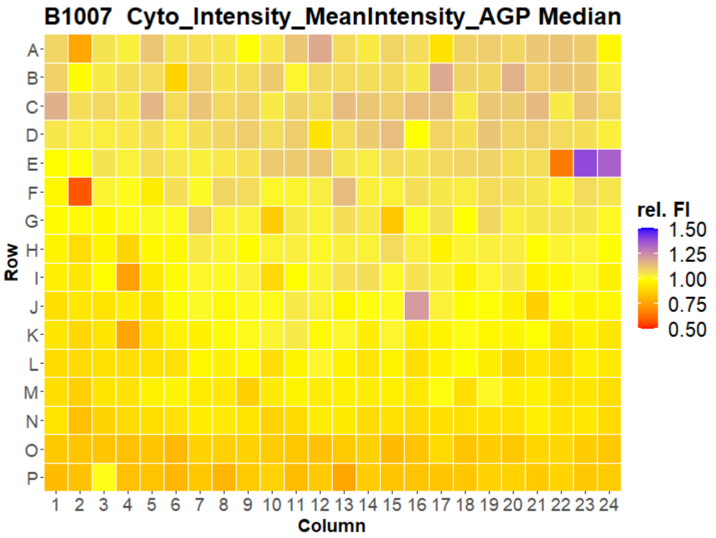



Open for Innovation KNIME

### HepG2 IMTM



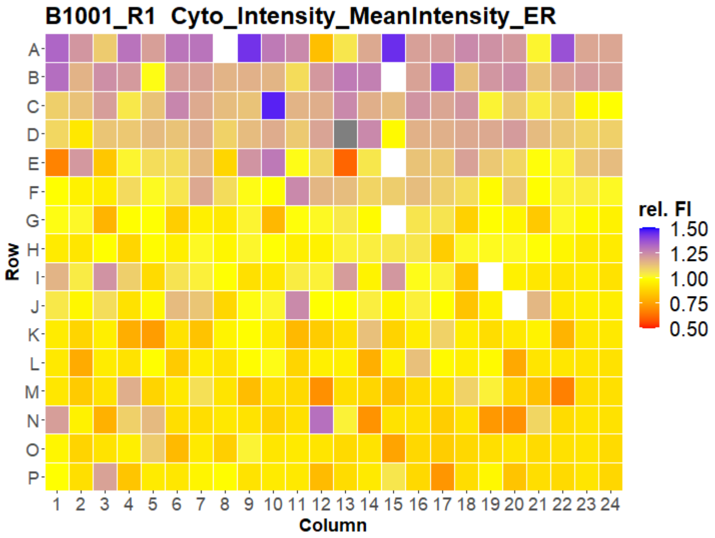
B1001_R1 Cyto_Intensity_MeanIntensity_ER
Row
Column
rel. FI
1.50
1.25
1.00
0.75
0.50

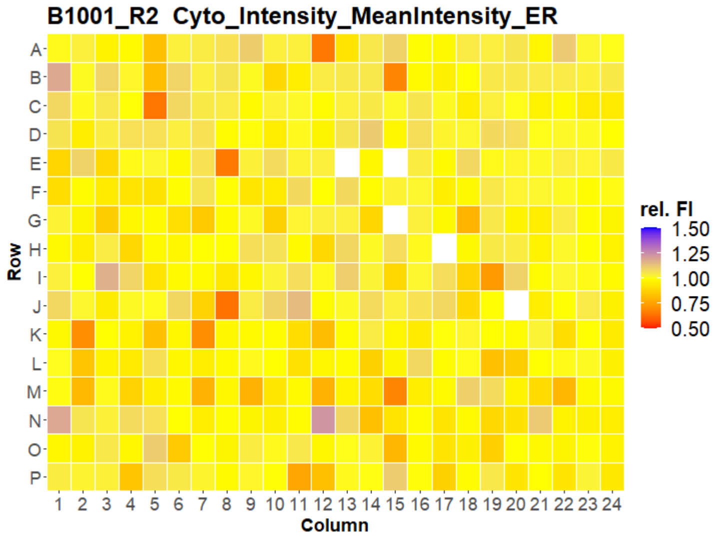
B1001_R2 Cyto_Intensity_MeanIntensity_ER
Row
Column
rel. FI
1.50
1.25
1.00
0.75
0.50

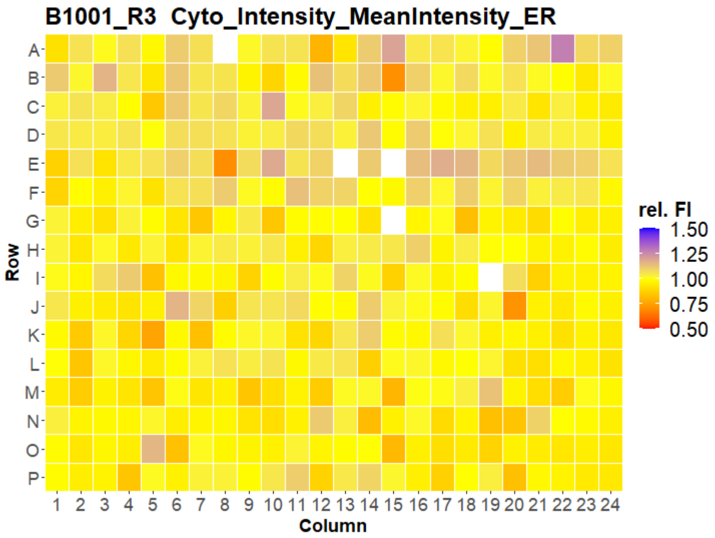
B1001_R3 Cyto_Intensity_MeanIntensity_ER
Row
Column
rel. FI
1.50
1.25
1.00
0.75
0.50

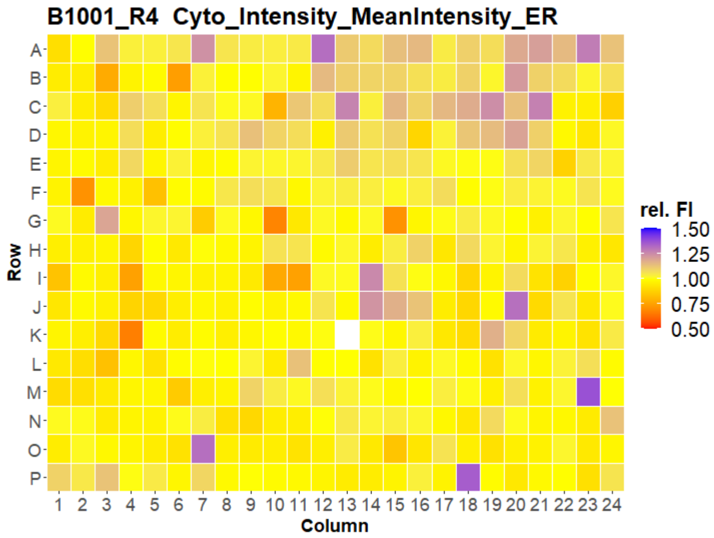
B1001_R4 Cyto_Intensity_MeanIntensity_ER
Row
Column
rel. FI
1.50
1.25
1.00
0.75
0.50

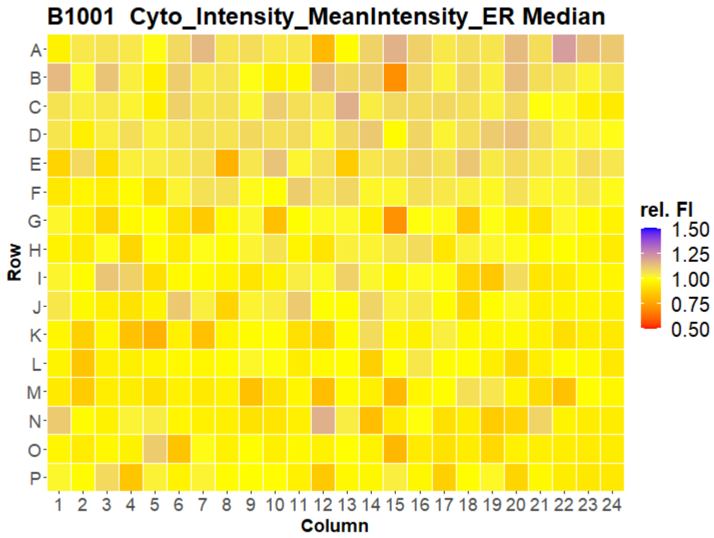
B1001 Cyto_Intensity_MeanIntensity_ER Median
Row
Column
rel. FI
1.50
1.25
1.00
0.75
0.50

### HepG2 IMTM



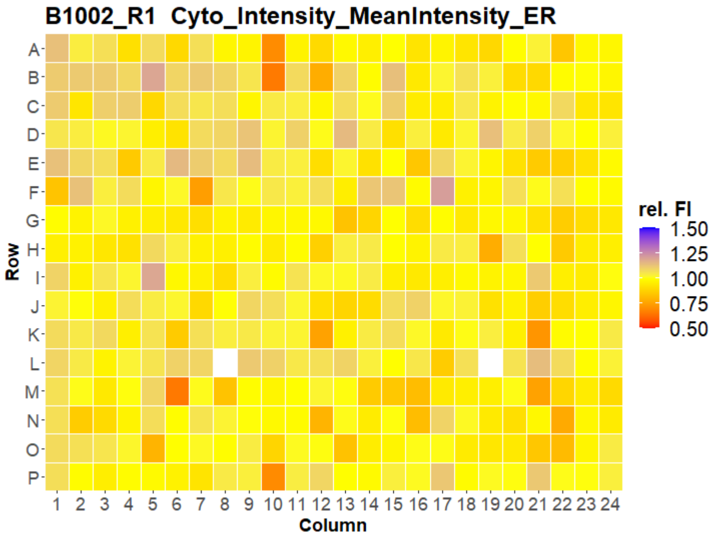
B1002_R1 Cyto_Intensity_MeanIntensity_ER
Row
A B C D E F G H I J K L M N O P
1 2 3 4 5 6 7 8 9 10 11 12 13 14 15 16 17 18 19 20 21 22 23 24
Column
rel. FI
1.50
1.25
1.00
0.75
0.50

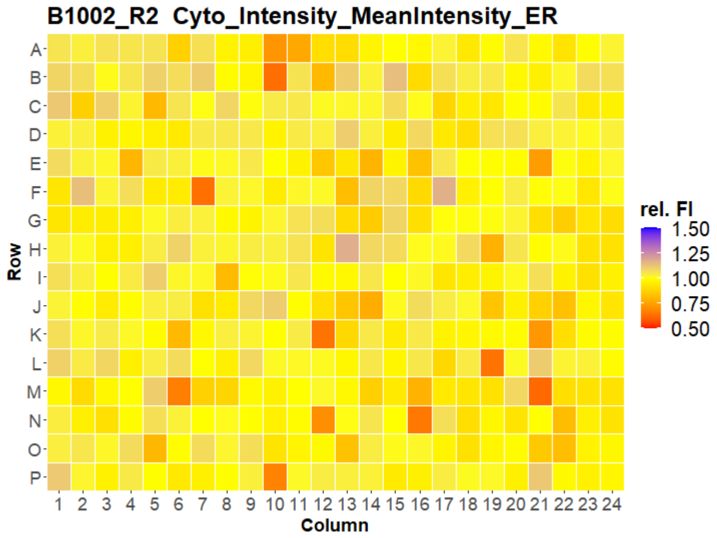
B1002_R2 Cyto_Intensity_MeanIntensity_ER
Row
A B C D E F G H I J K L M N O P
1 2 3 4 5 6 7 8 9 10 11 12 13 14 15 16 17 18 19 20 21 22 23 24
Column
rel. FI
1.50
1.25
1.00
0.75
0.50

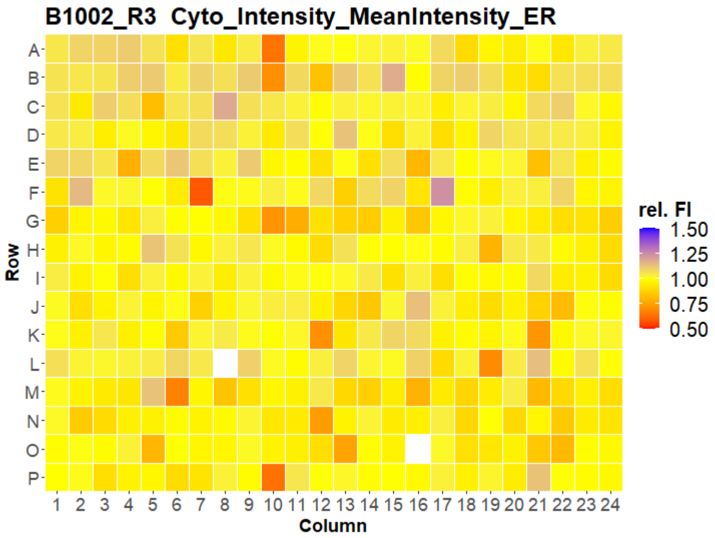
B1002_R3 Cyto_Intensity_MeanIntensity_ER
Row
A B C D E F G H I J K L M N O P
1 2 3 4 5 6 7 8 9 10 11 12 13 14 15 16 17 18 19 20 21 22 23 24
Column
rel. FI
1.50
1.25
1.00
0.75
0.50

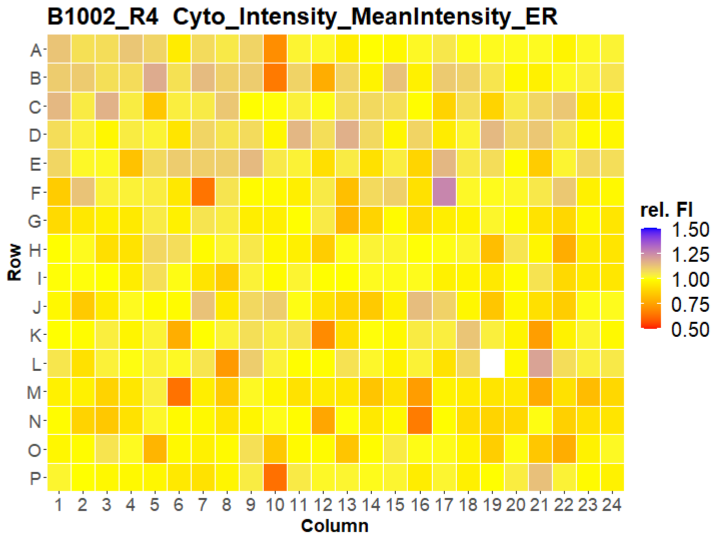
B1002_R4 Cyto_Intensity_MeanIntensity_ER
Row
A B C D E F G H I J K L M N O P
1 2 3 4 5 6 7 8 9 10 11 12 13 14 15 16 17 18 19 20 21 22 23 24
Column
rel. FI
1.50
1.25
1.00
0.75
0.50

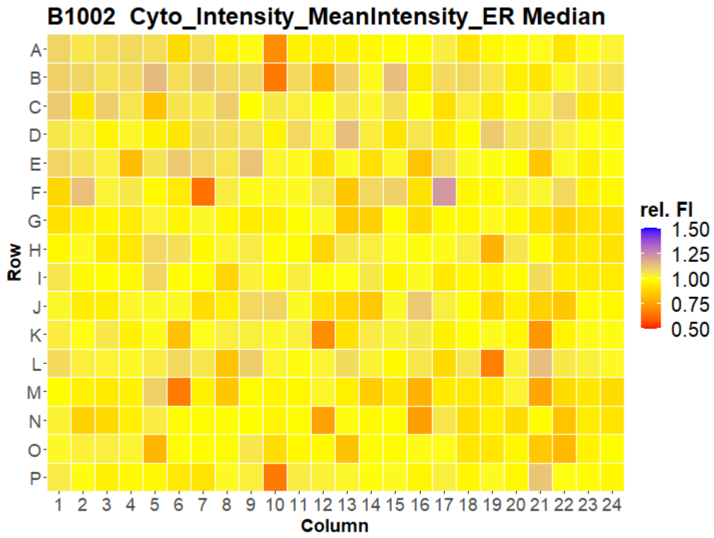
B1002 Cyto_Intensity_MeanIntensity_ER Median
Row
A B C D E F G H I J K L M N O P
1 2 3 4 5 6 7 8 9 10 11 12 13 14 15 16 17 18 19 20 21 22 23 24
Column
rel. FI
1.50
1.25
1.00
0.75
0.50

### HepG2 IMTM

10

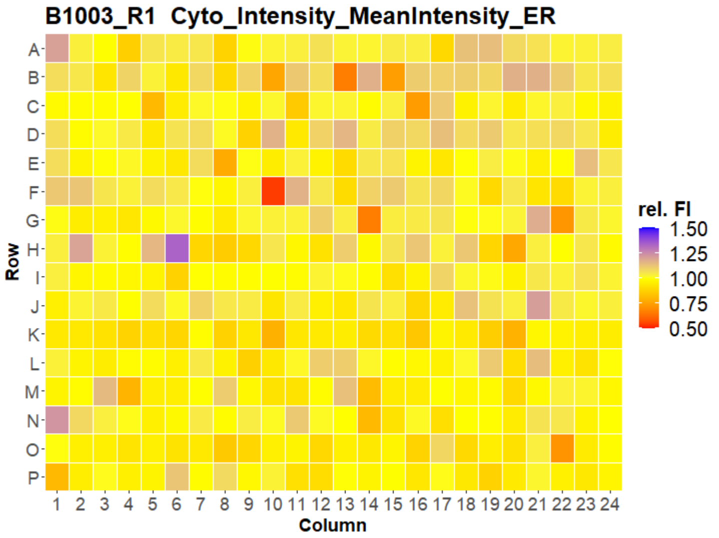


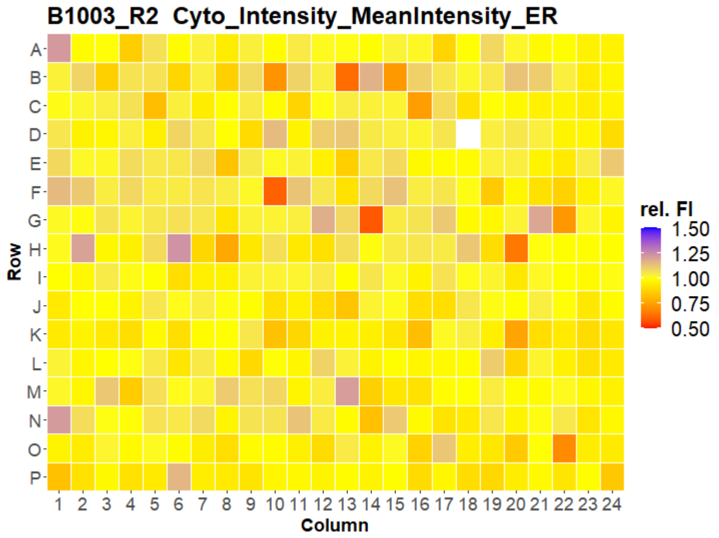


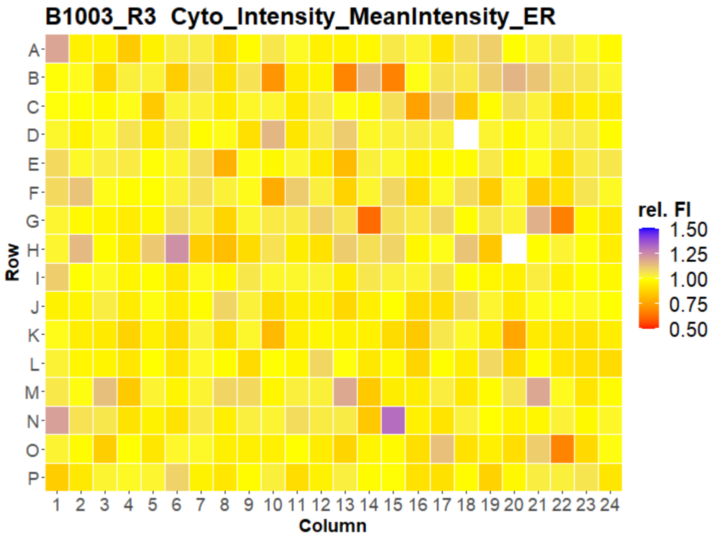


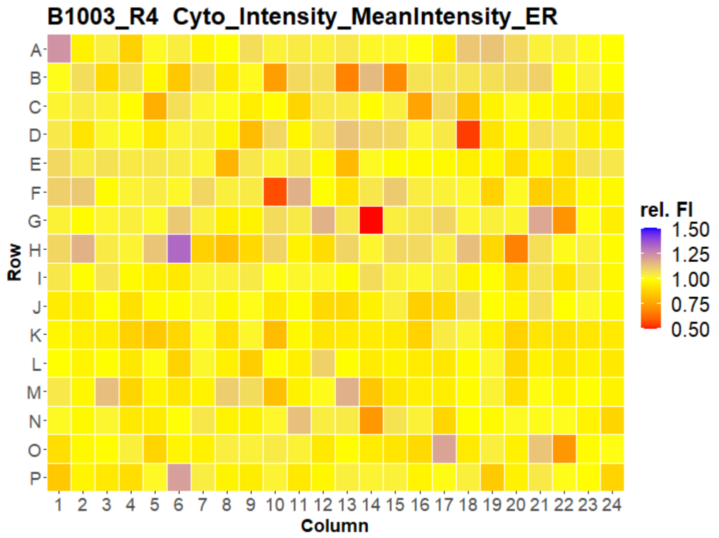


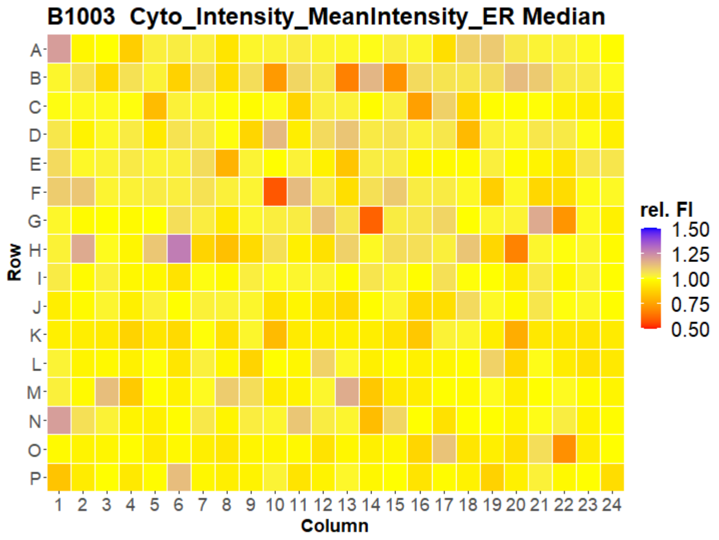

### HepG2 IMTM



**B1004_R1 Cyto_Intensity_MeanIntensity_ER**

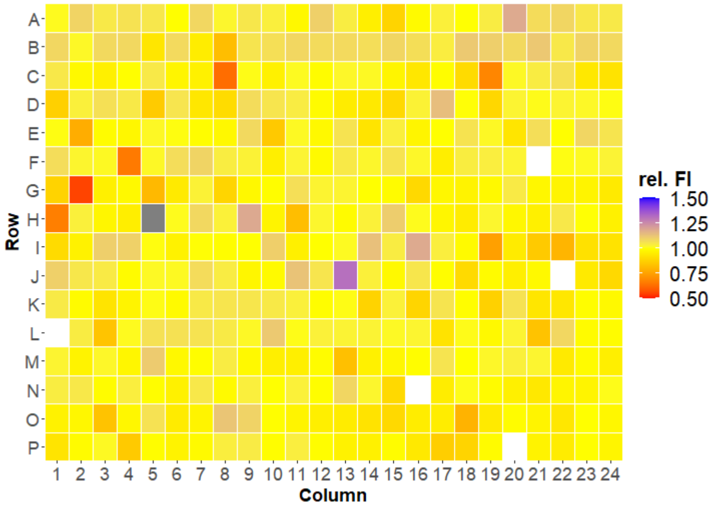


**B1004_R2 Cyto_Intensity_MeanIntensity_ER**

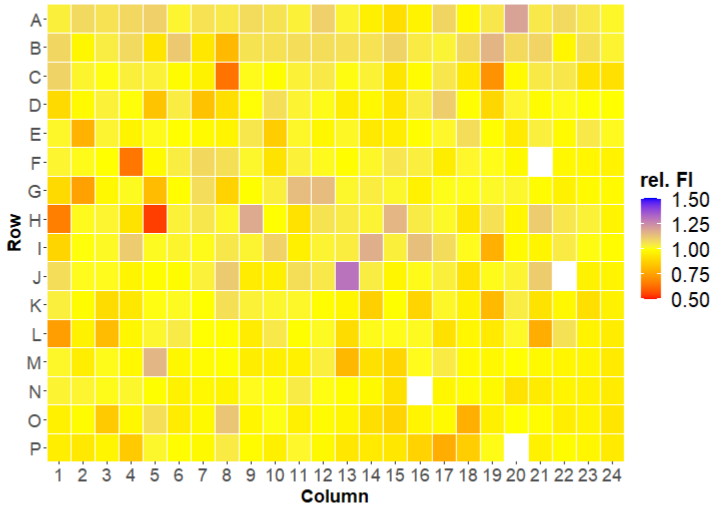


**B1004_R3 Cyto_Intensity_MeanIntensity_ER**

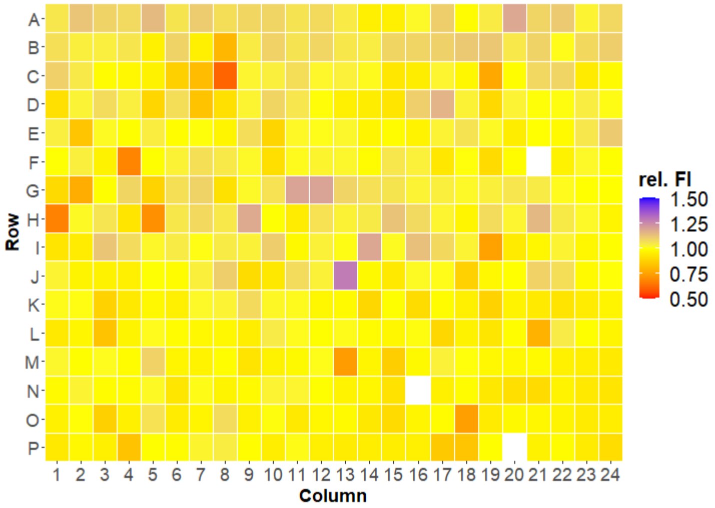


**B1004_R4 Cyto_Intensity_MeanIntensity_ER**

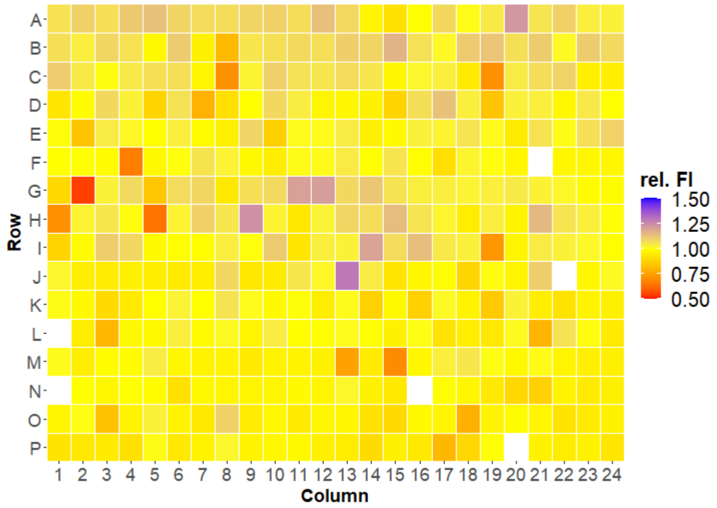


**B1004 Cyto_Intensity_MeanIntensity_ER Median**

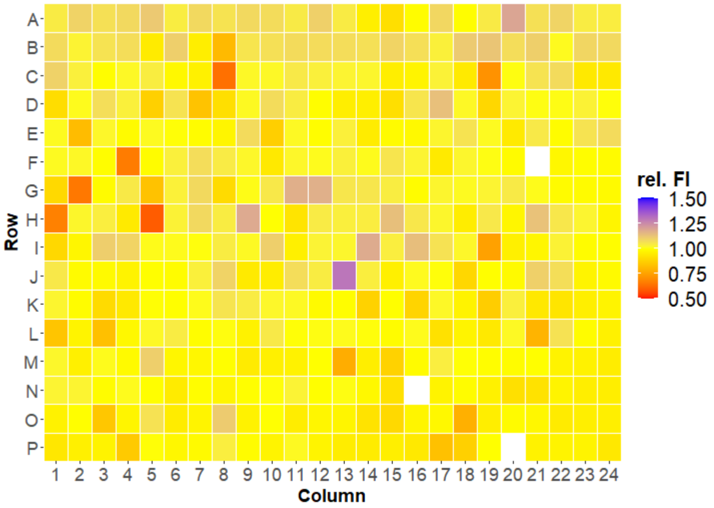

### HepG2 IMTM

12

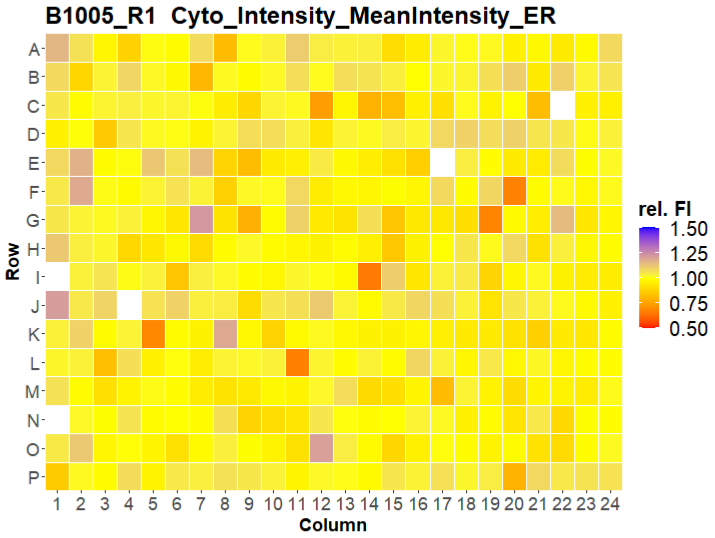


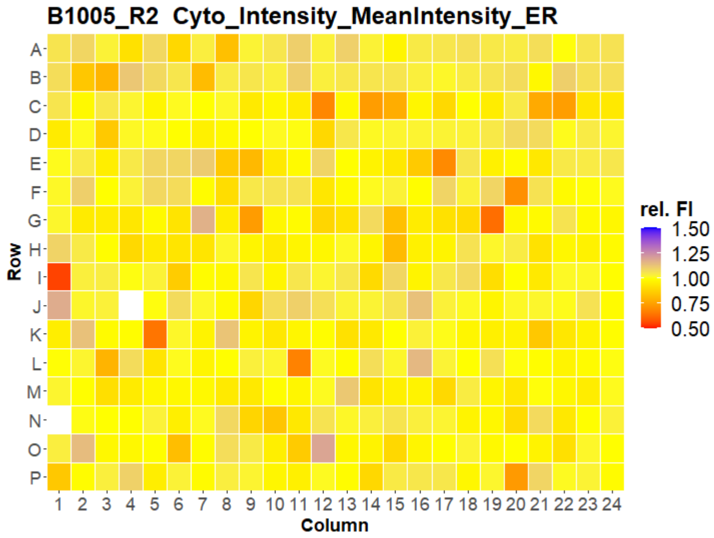


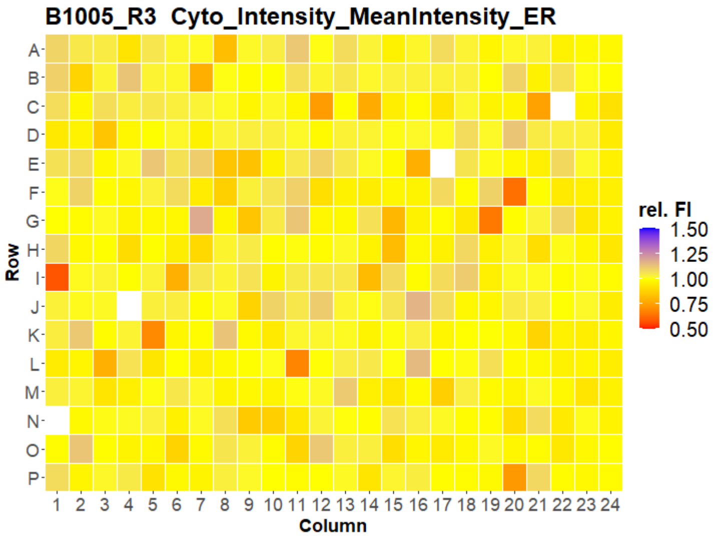


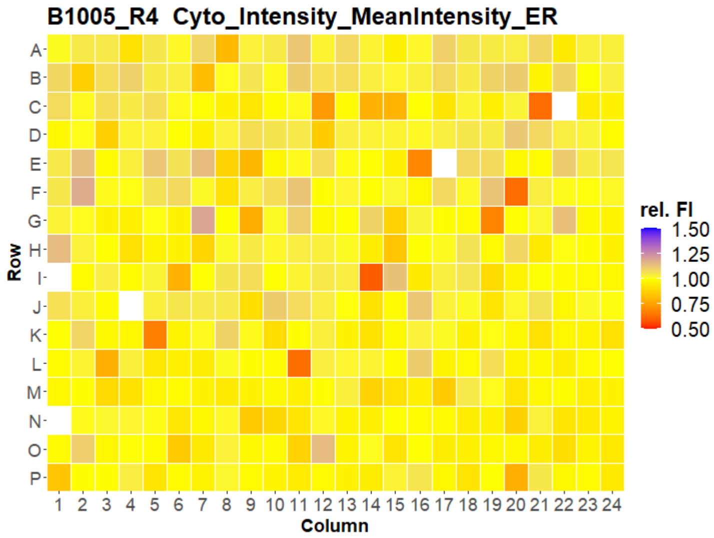


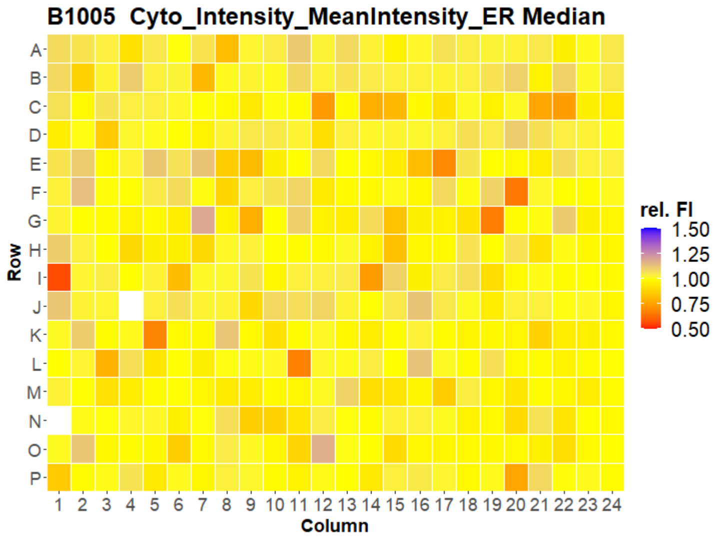



Open for Innovation KNIME

### HepG2 IMTM

13

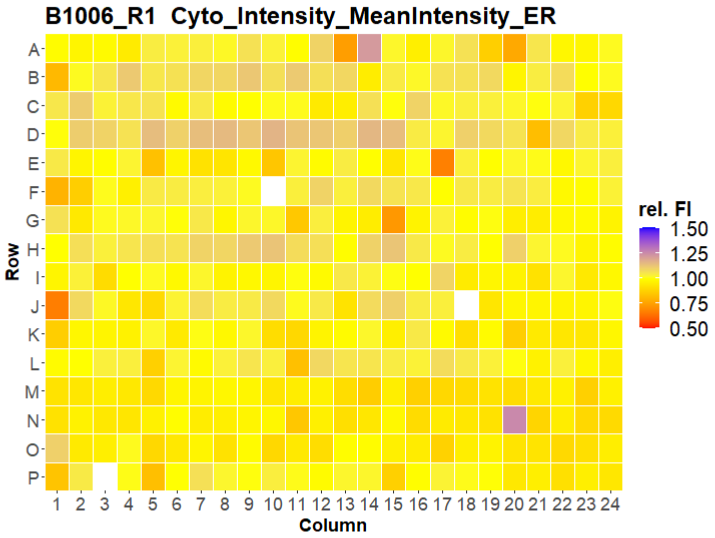


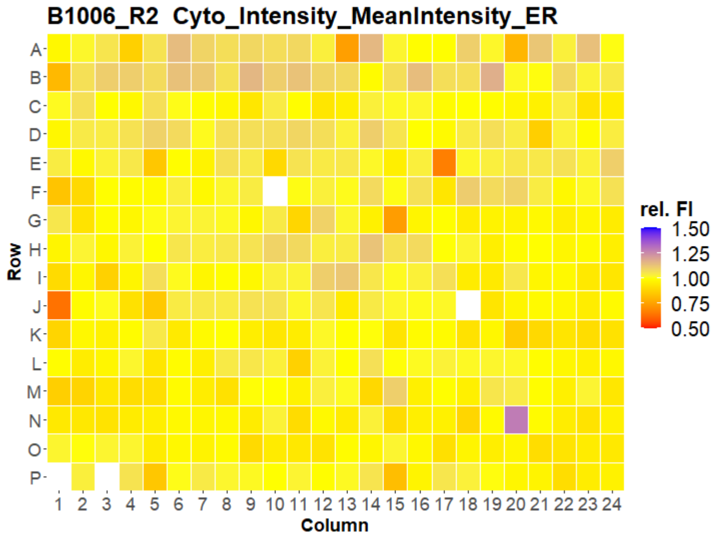


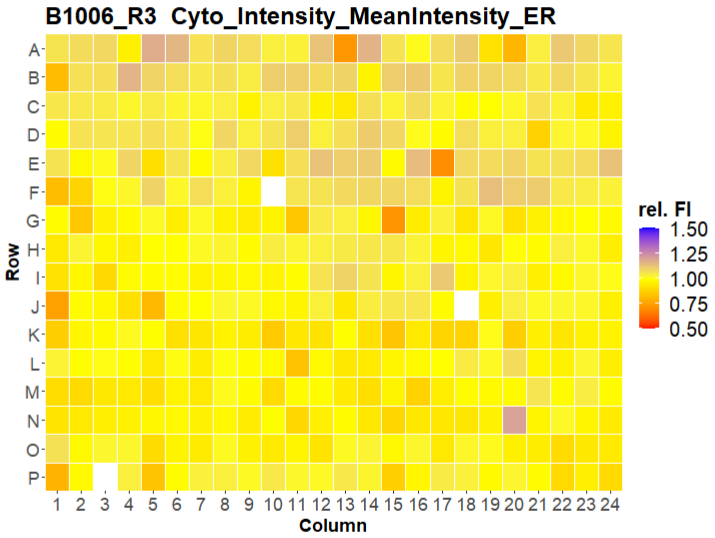


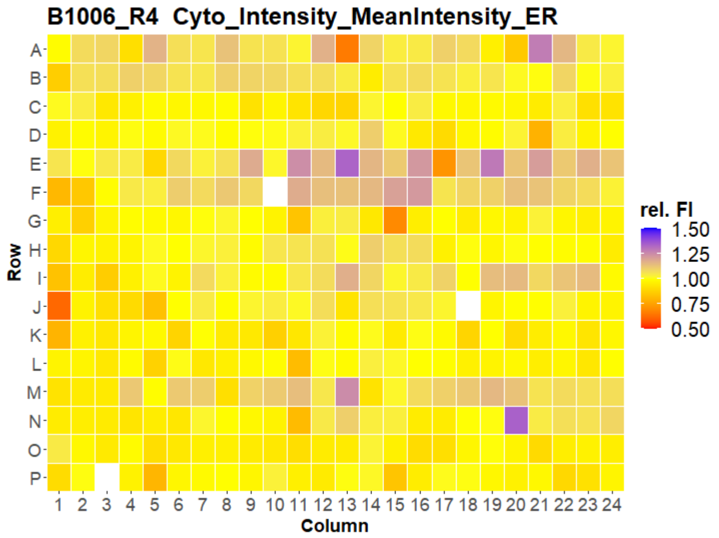


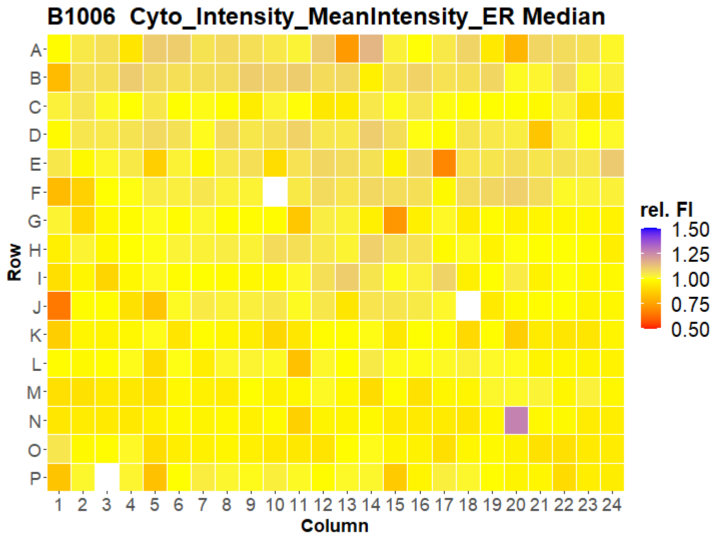

### HepG2 IMTM



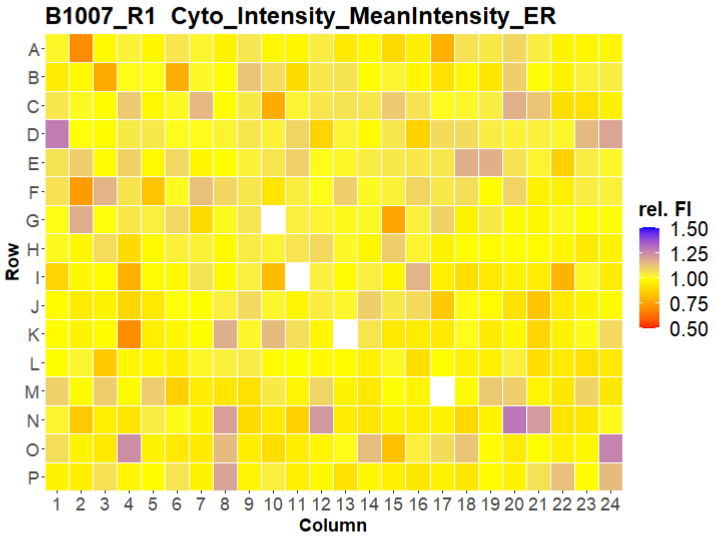
B1007_R1 Cyto_Intensity_MeanIntensity_ER
Row
A B C D E F G H I J K L M N O P
1 2 3 4 5 6 7 8 9 10 11 12 13 14 15 16 17 18 19 20 21 22 23 24
Column
rel. FI
1.50
1.25
1.00
0.75
0.50

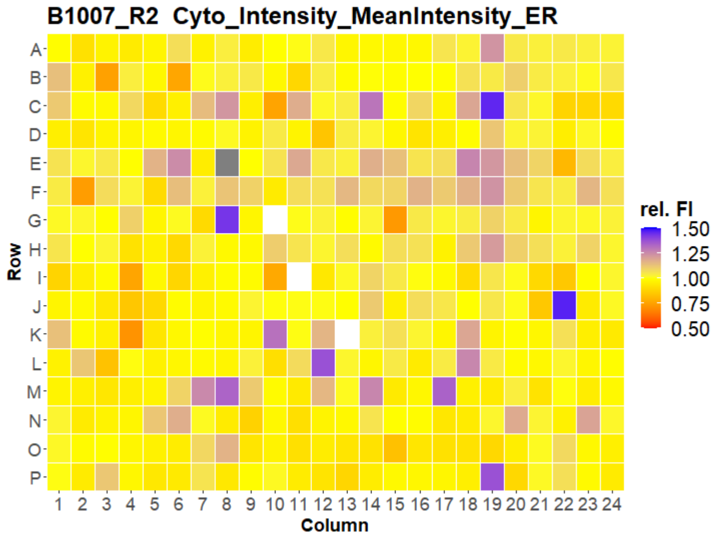
B1007_R2 Cyto_Intensity_MeanIntensity_ER
Row
A B C D E F G H I J K L M N O P
1 2 3 4 5 6 7 8 9 10 11 12 13 14 15 16 17 18 19 20 21 22 23 24
Column
rel. FI
1.50
1.25
1.00
0.75
0.50

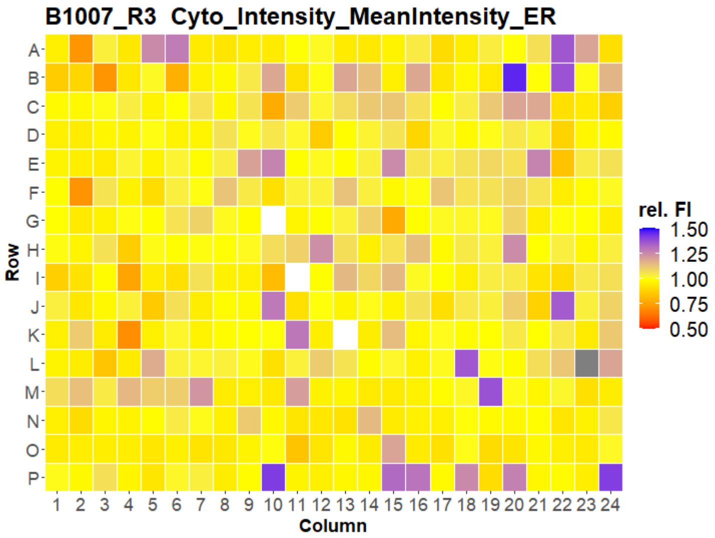
B1007_R3 Cyto_Intensity_MeanIntensity_ER
Row
A B C D E F G H I J K L M N O P
1 2 3 4 5 6 7 8 9 10 11 12 13 14 15 16 17 18 19 20 21 22 23 24
Column
rel. FI
1.50
1.25
1.00
0.75
0.50

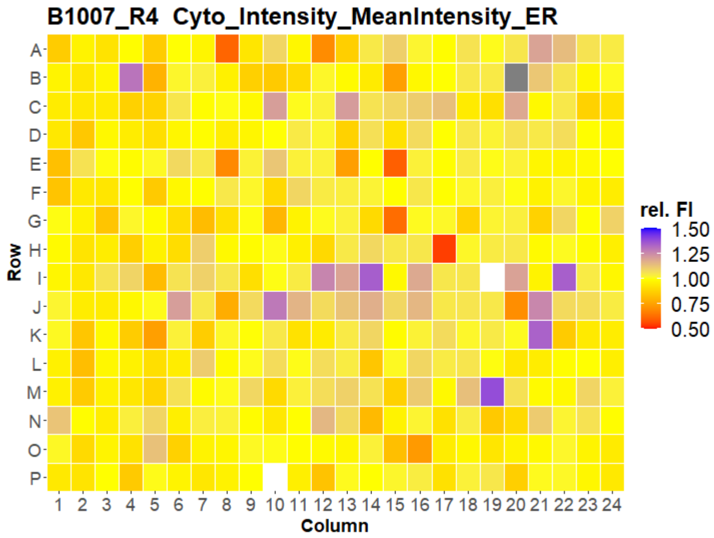
B1007_R4 Cyto_Intensity_MeanIntensity_ER
Row
A B C D E F G H I J K L M N O P
1 2 3 4 5 6 7 8 9 10 11 12 13 14 15 16 17 18 19 20 21 22 23 24
Column
rel. FI
1.50
1.25
1.00
0.75
0.50

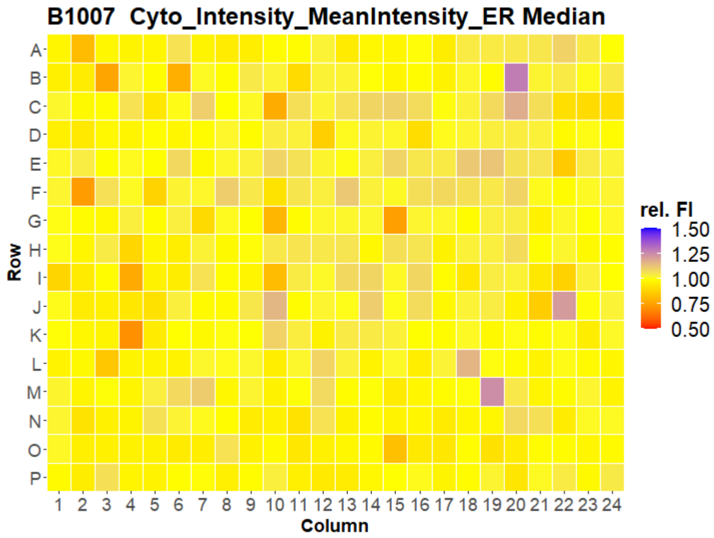
B1007 Cyto_Intensity_MeanIntensity_ER Median
Row
A B C D E F G H I J K L M N O P
1 2 3 4 5 6 7 8 9 10 11 12 13 14 15 16 17 18 19 20 21 22 23 24
Column
rel. FI
1.50
1.25
1.00
0.75
0.50

### HepG2 IMTM



B1001_R1 Cyto_Intensity_MeanIntensity_Mito

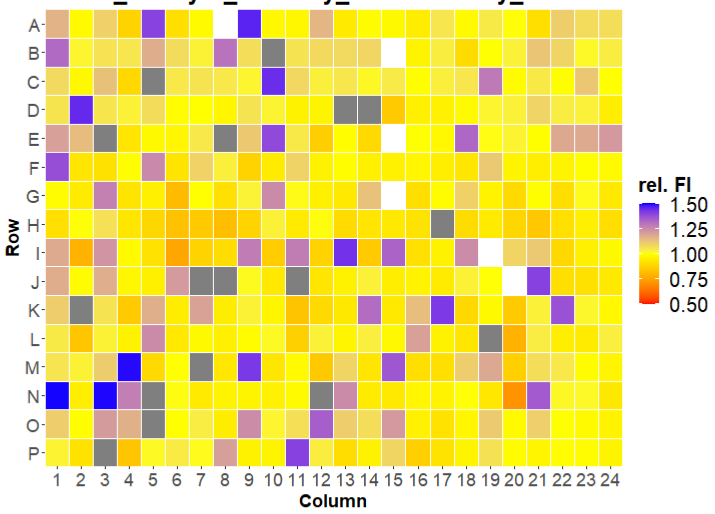


B1001_R2 Cyto_Intensity_MeanIntensity_Mito

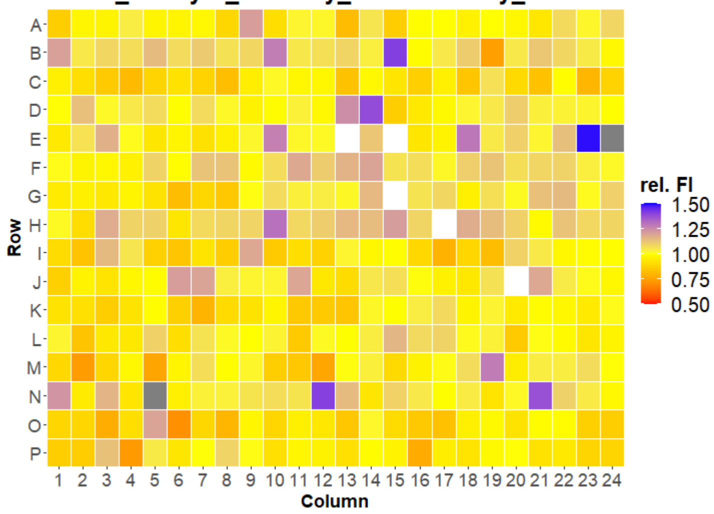


B1001_R3 Cyto_Intensity_MeanIntensity_Mito

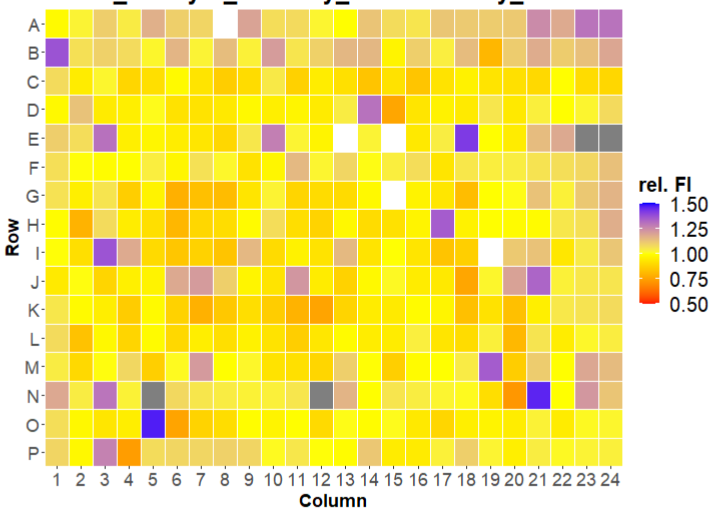


B1001_R4 Cyto_Intensity_MeanIntensity_Mito

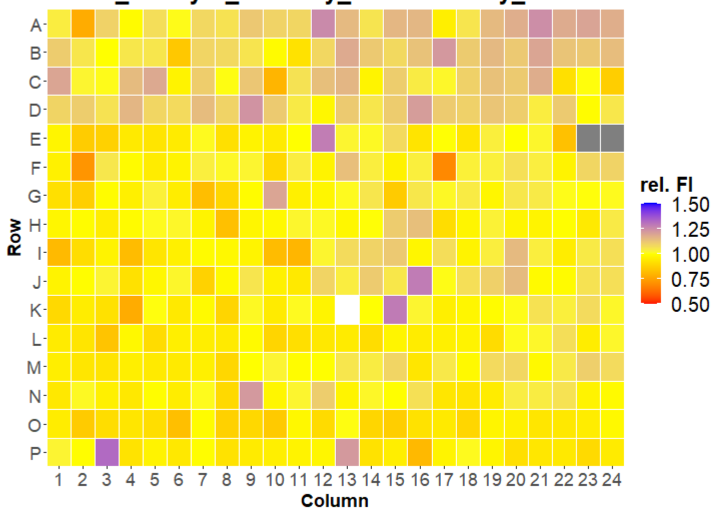


B1001 Cyto_Intensity_MeanIntensity_Mito Median

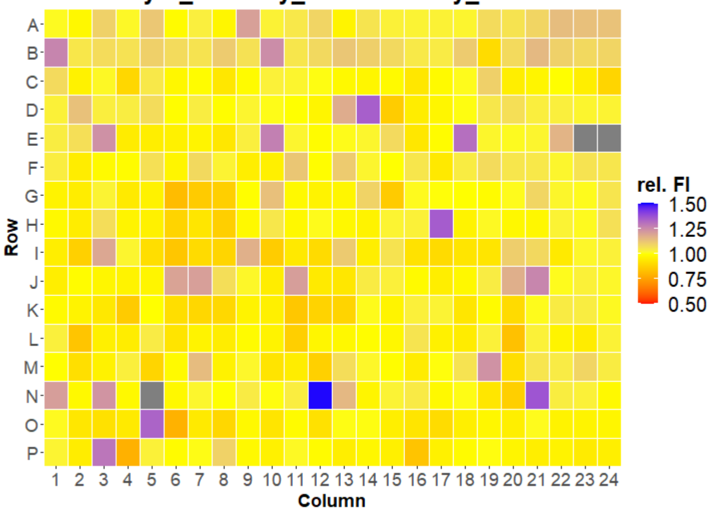

### HepG2 IMTM

16

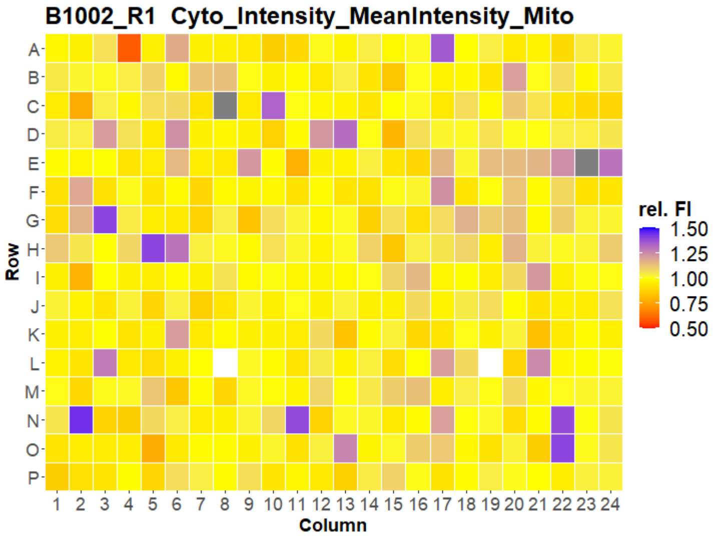


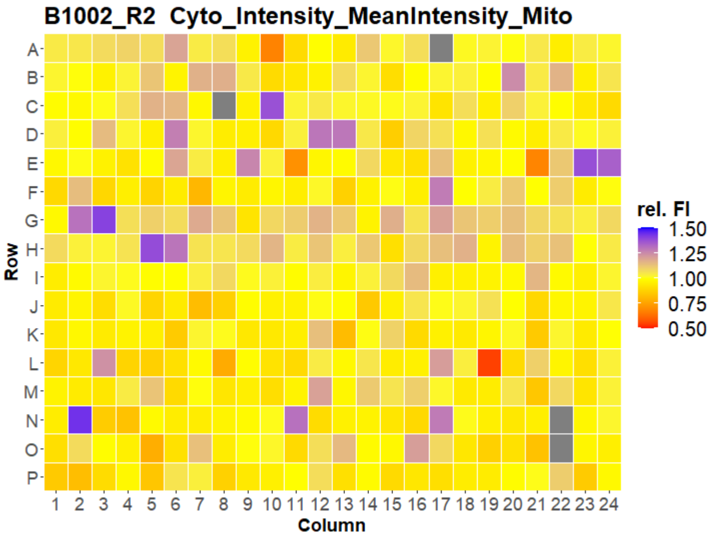


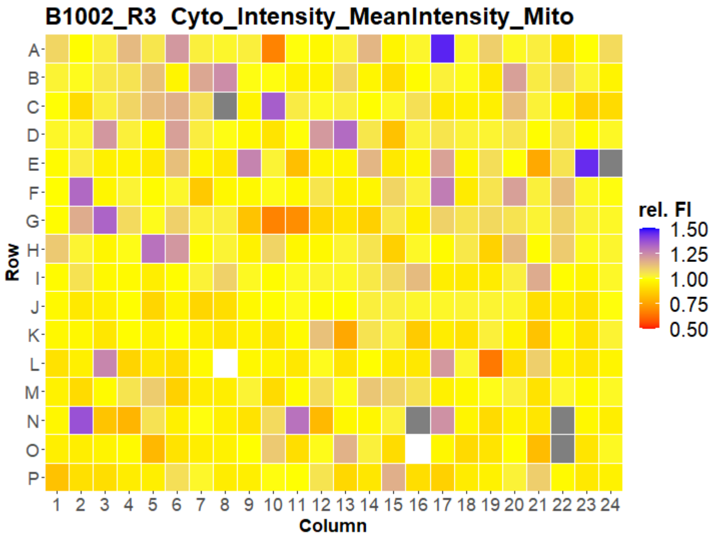


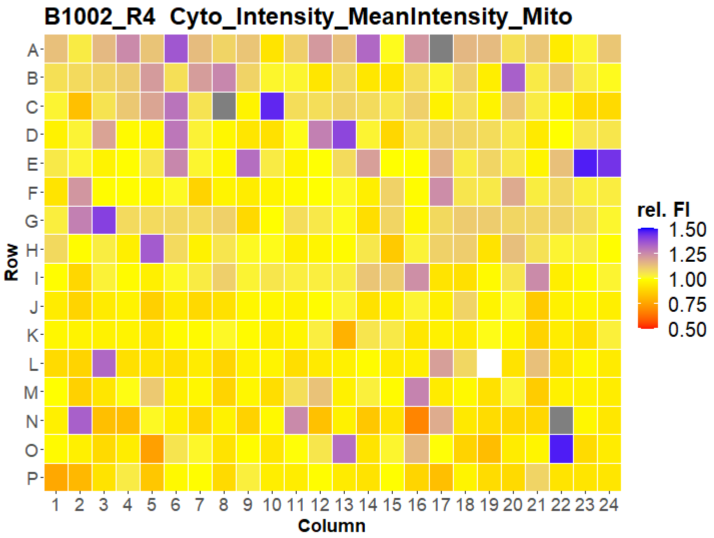


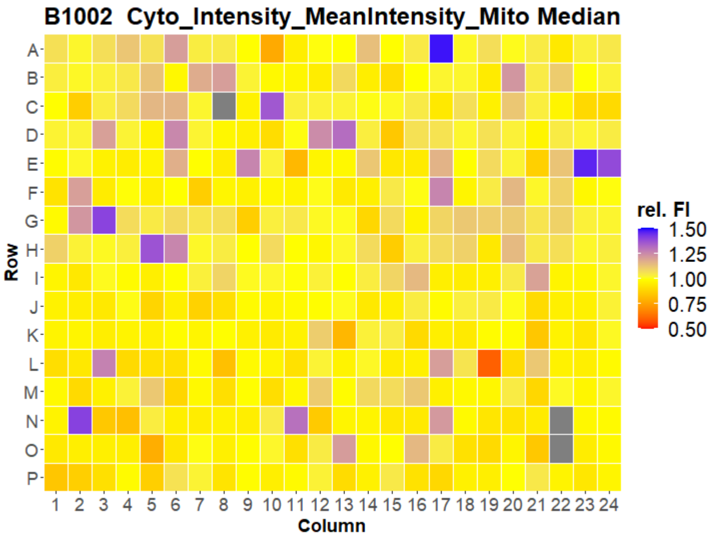

### HepG2 IMTM

17

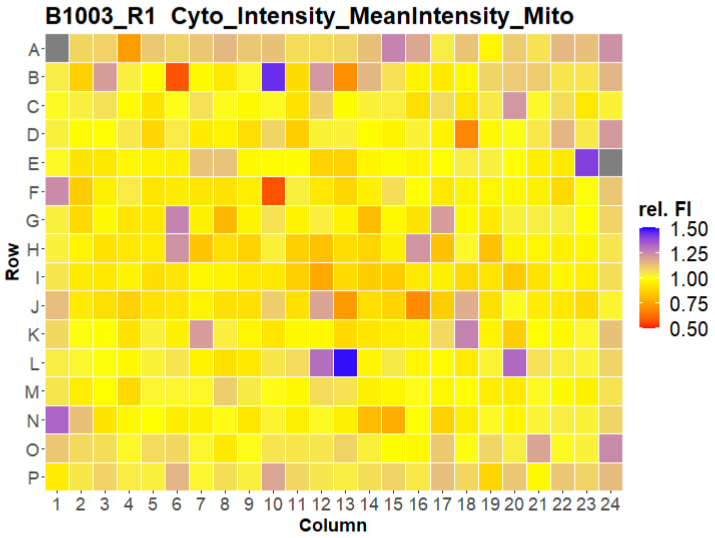


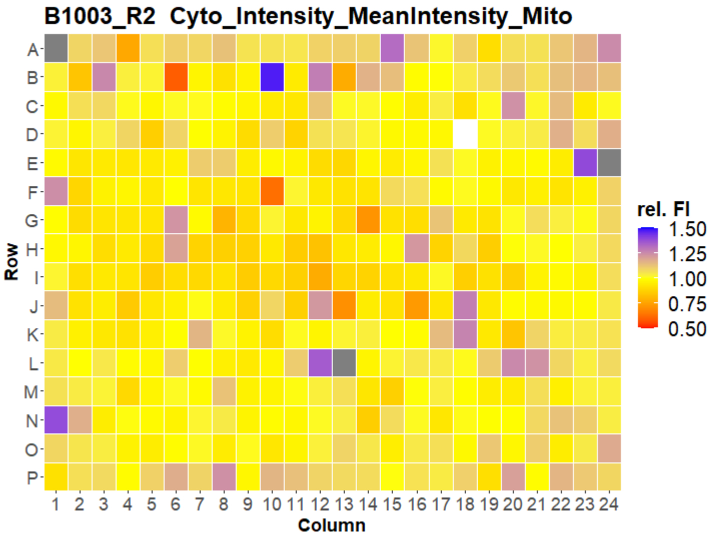


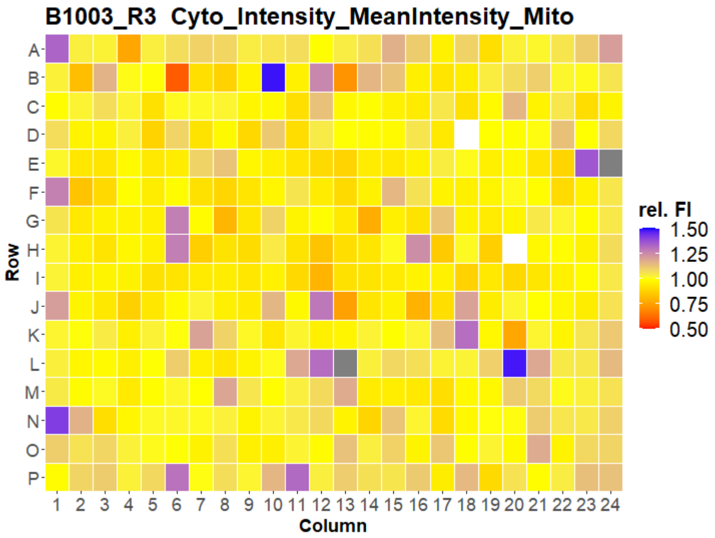


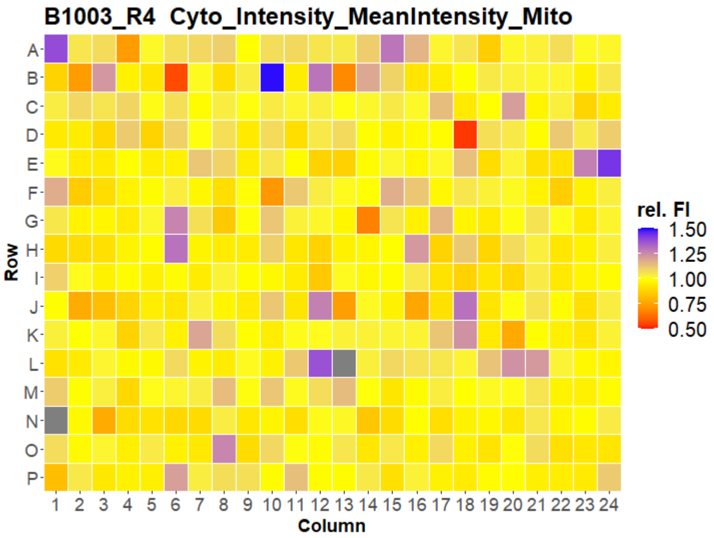


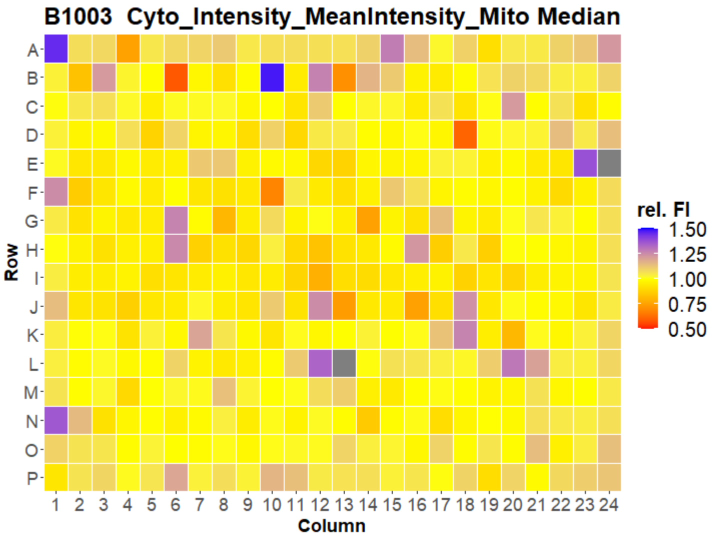

### HepG2 IMTM

18

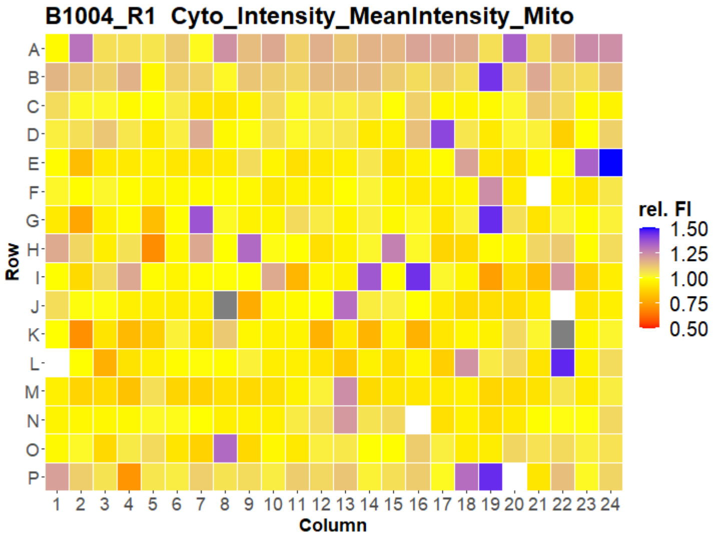


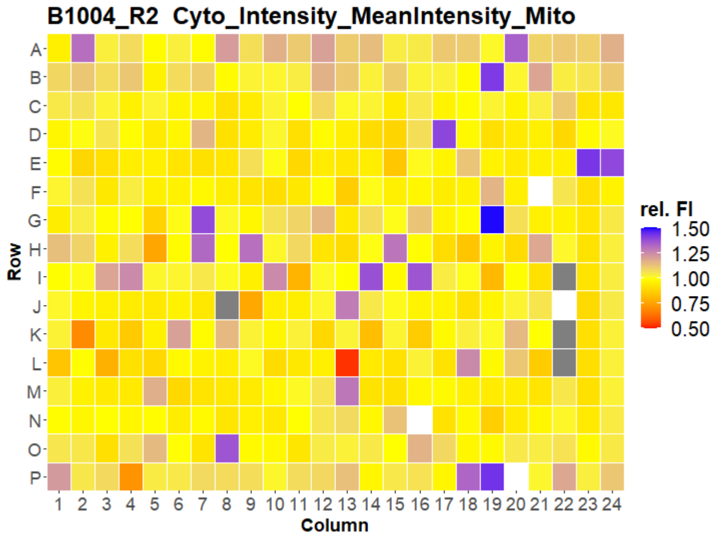


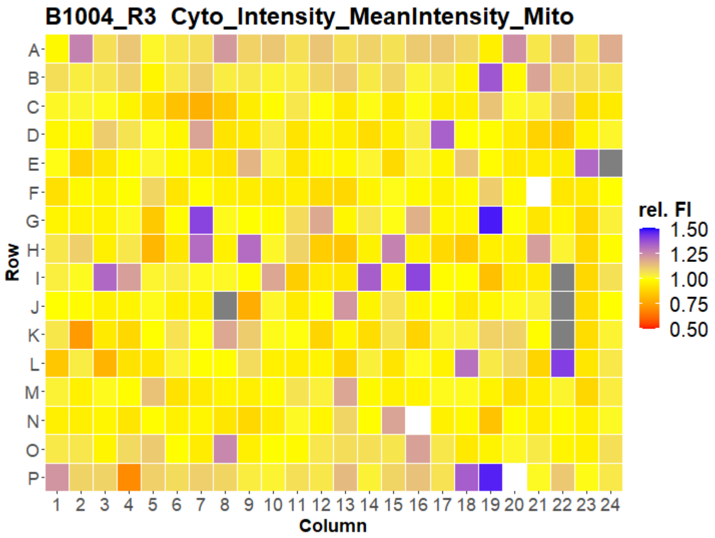


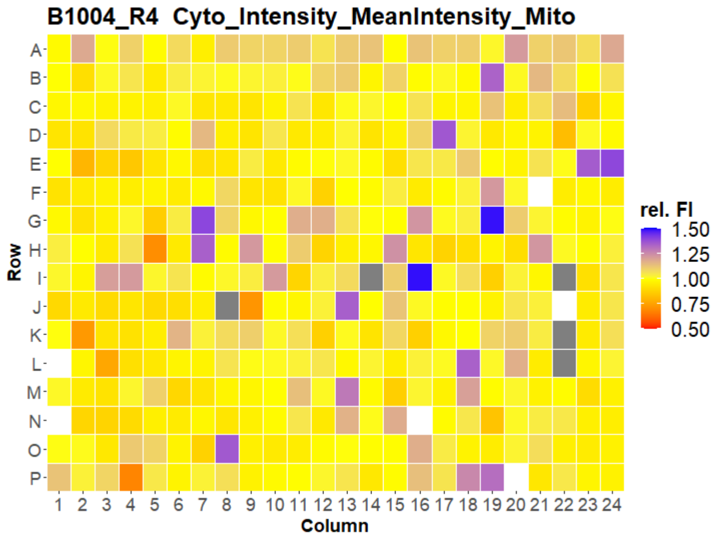


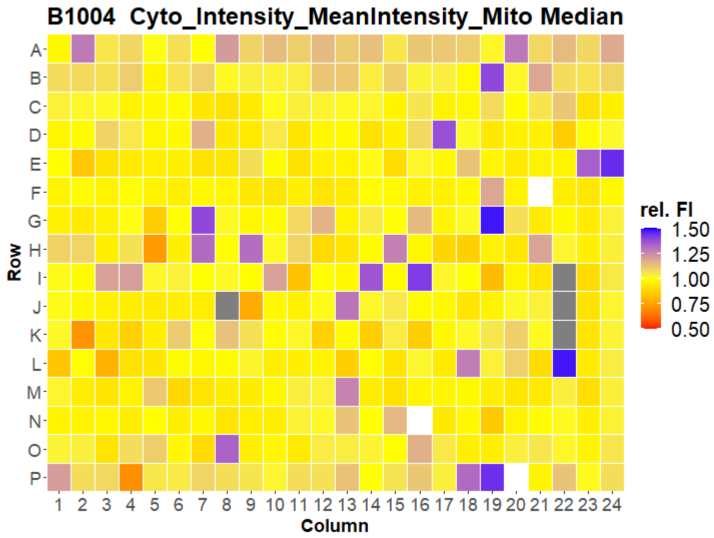


Created with KNIME Analytics Platform www.knime.com
Open for Innovation KNIME

### HepG2 IMTM

19

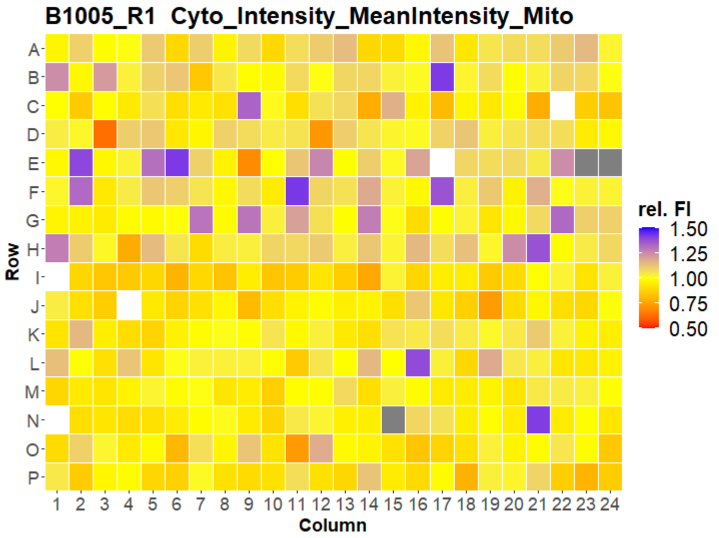


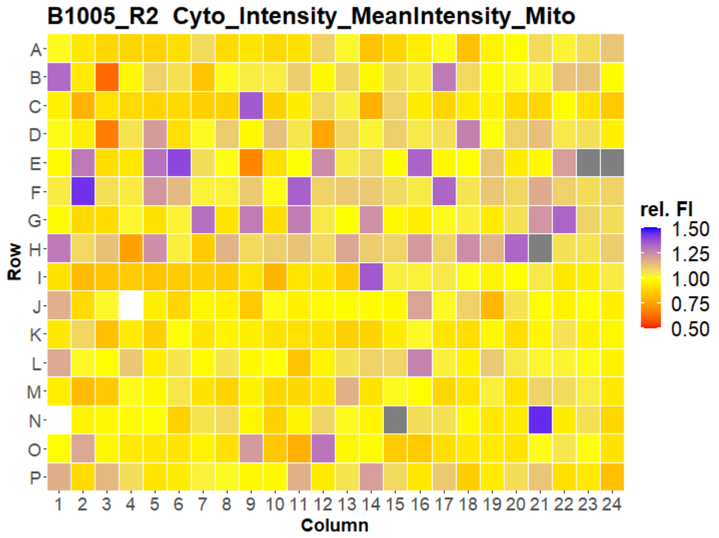


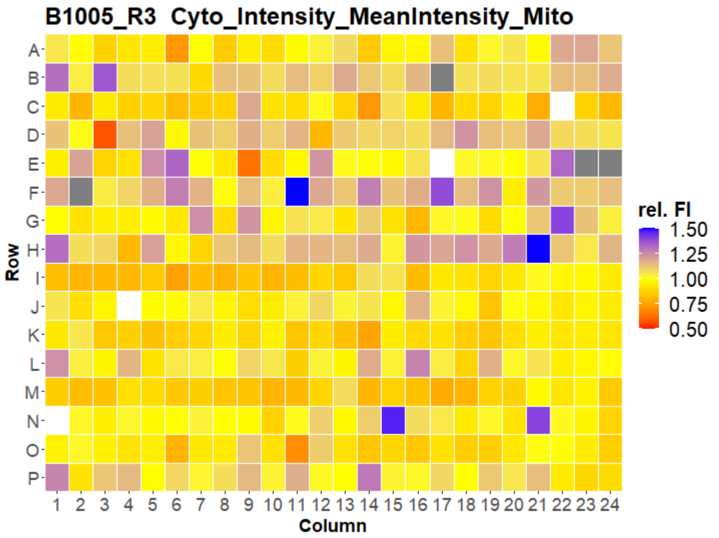


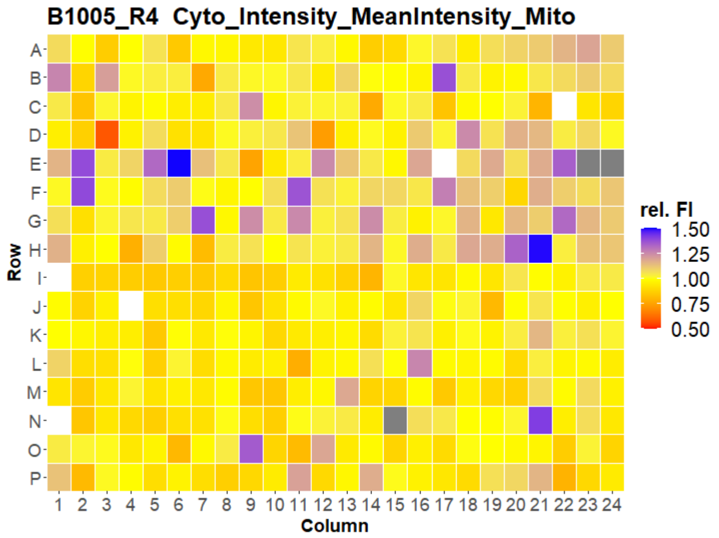


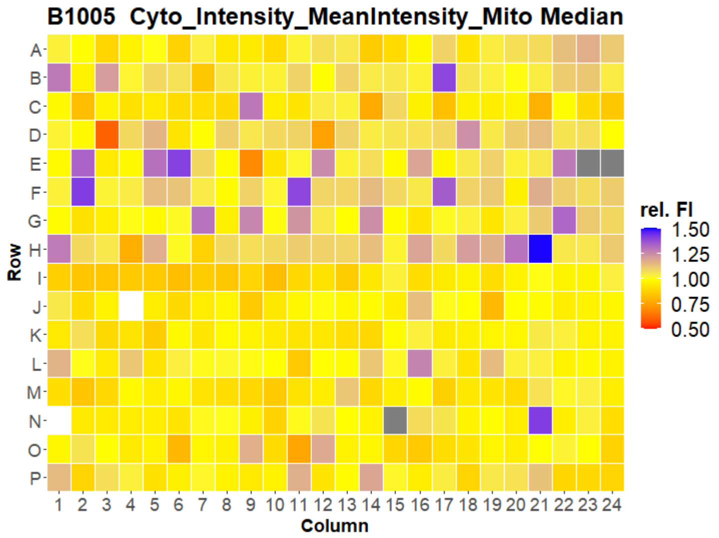

### HepG2 IMTM



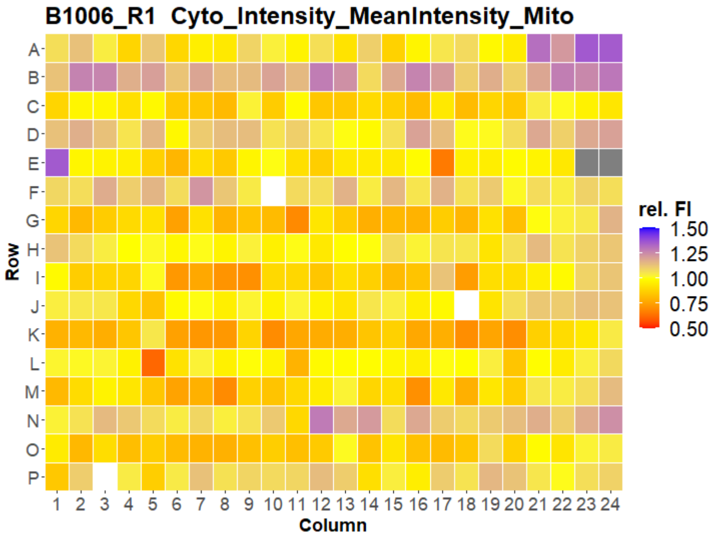
B1006_R1 Cyto_Intensity_MeanIntensity_Mito
Row
Column
rel. FI
1.50
1.25
1.00
0.75
0.50

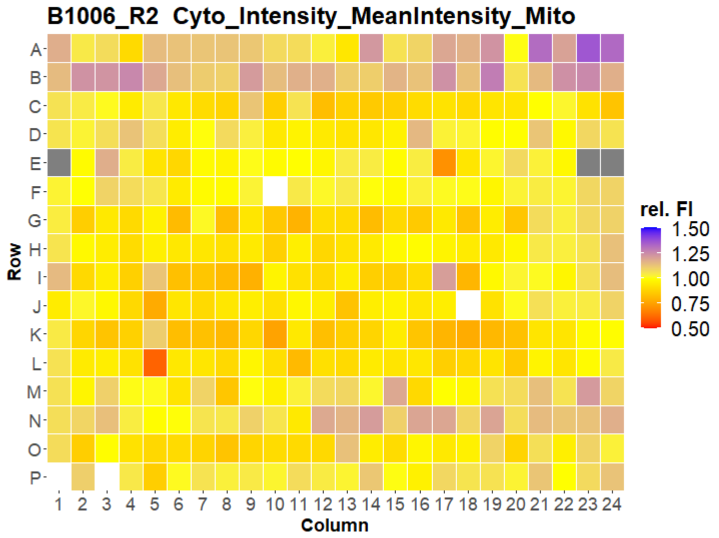
B1006_R2 Cyto_Intensity_MeanIntensity_Mito
Row
Column
rel. FI
1.50
1.25
1.00
0.75
0.50

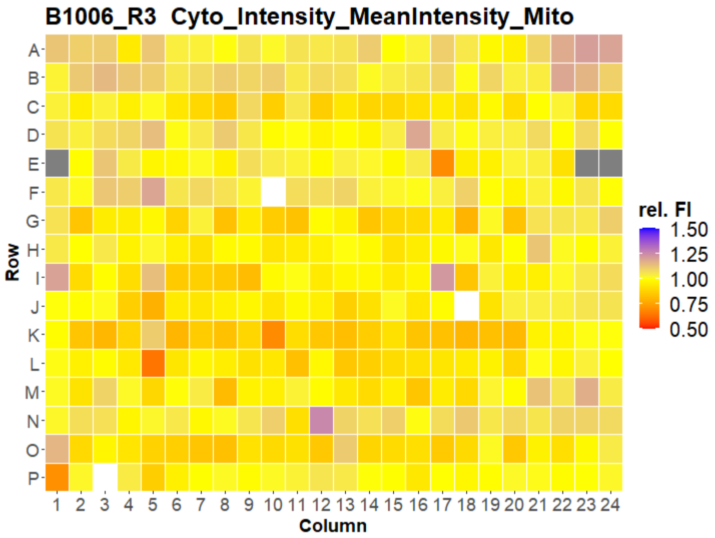
B1006_R3 Cyto_Intensity_MeanIntensity_Mito
Row
Column
rel. FI
1.50
1.25
1.00
0.75
0.50

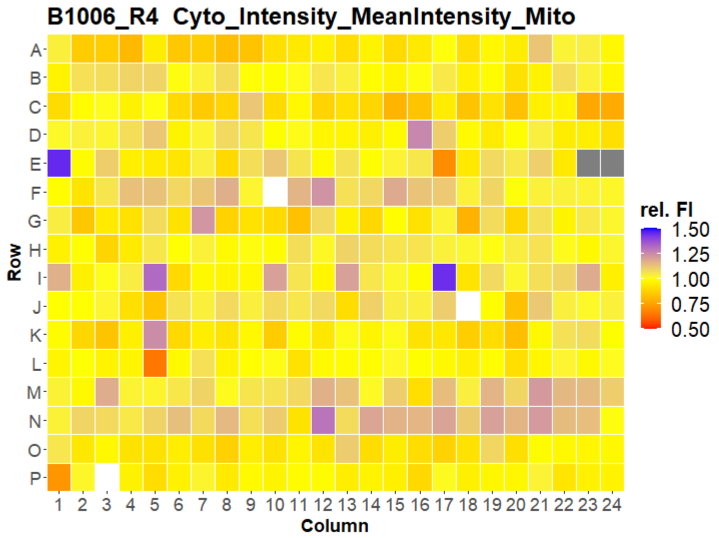
B1006_R4 Cyto_Intensity_MeanIntensity_Mito
Row
Column
rel. FI
1.50
1.25
1.00
0.75
0.50

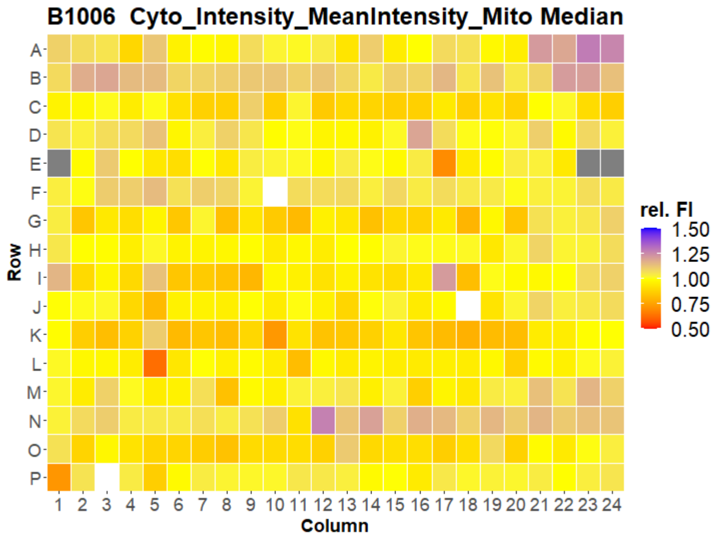
B1006 Cyto_Intensity_MeanIntensity_Mito Median
Row
Column
rel. FI
1.50
1.25
1.00
0.75
0.50

### HepG2 IMTM



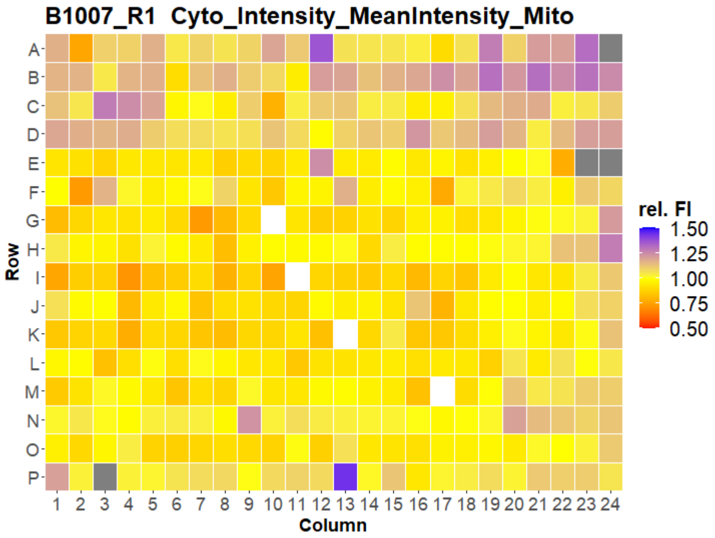
B1007_R1 Cyto_Intensity_MeanIntensity_Mito
Row
Column
rel. FI
1.50
1.25
1.00
0.75
0.50

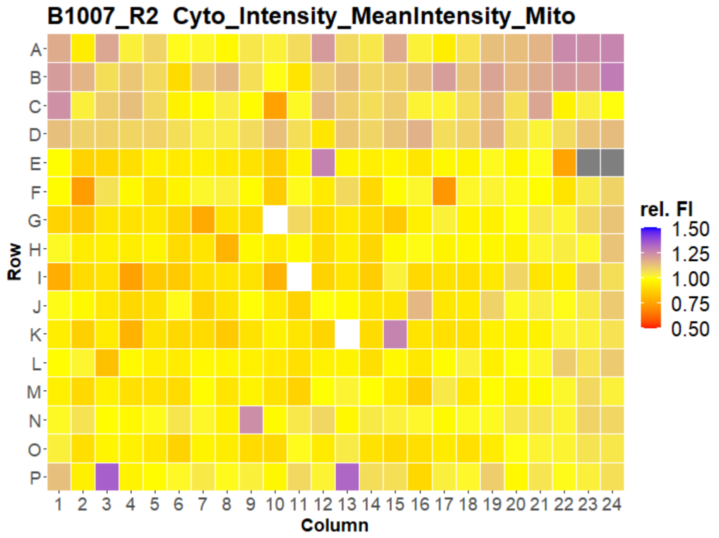
B1007_R2 Cyto_Intensity_MeanIntensity_Mito
Row
Column
rel. FI
1.50
1.25
1.00
0.75
0.50

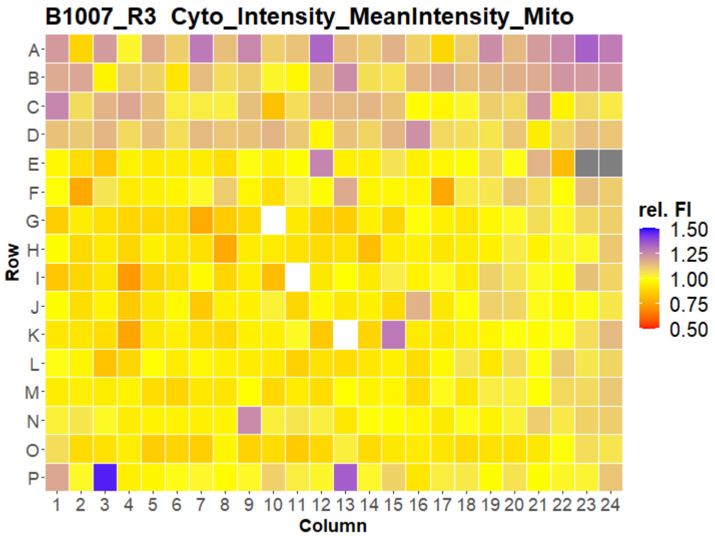
B1007_R3 Cyto_Intensity_MeanIntensity_Mito
Row
Column
rel. FI
1.50
1.25
1.00
0.75
0.50

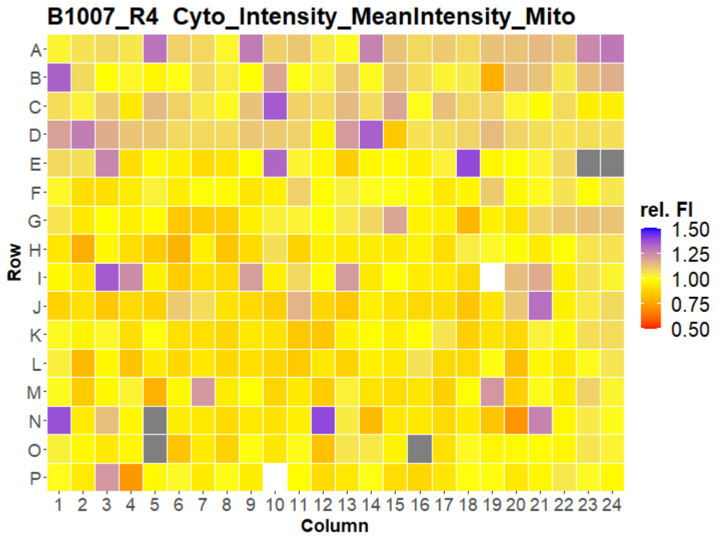
B1007_R4 Cyto_Intensity_MeanIntensity_Mito
Row
Column
rel. FI
1.50
1.25
1.00
0.75
0.50

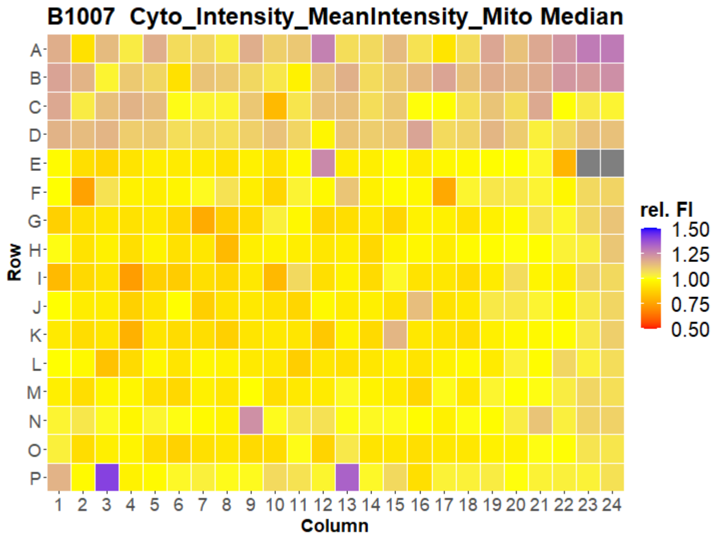
B1007 Cyto_Intensity_MeanIntensity_Mito Median
Row
Column
rel. FI
1.50
1.25
1.00
0.75
0.50

### HepG2 IMTM



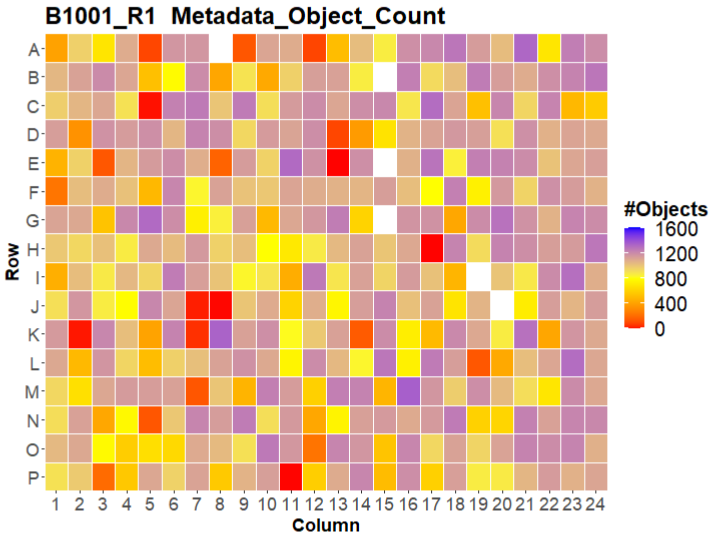
B1001_R1 Metadata_Object_Count
#Objects
1600
1200
800
400
0
Row
Column

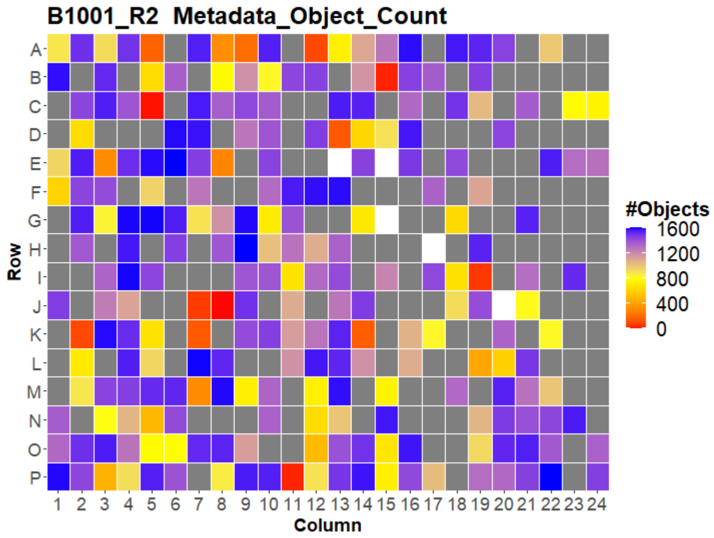
B1001_R2 Metadata_Object_Count
#Objects
1600
1200
800
400
0
Row
Column

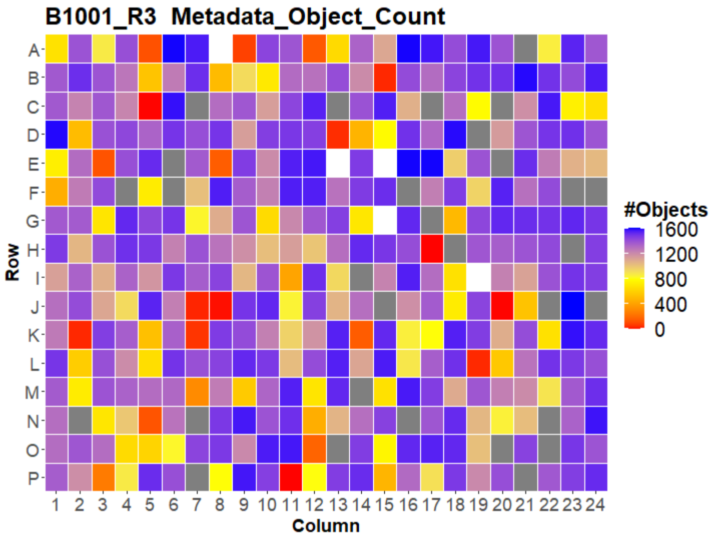
B1001_R3 Metadata_Object_Count
#Objects
1600
1200
800
400
0
Row
Column

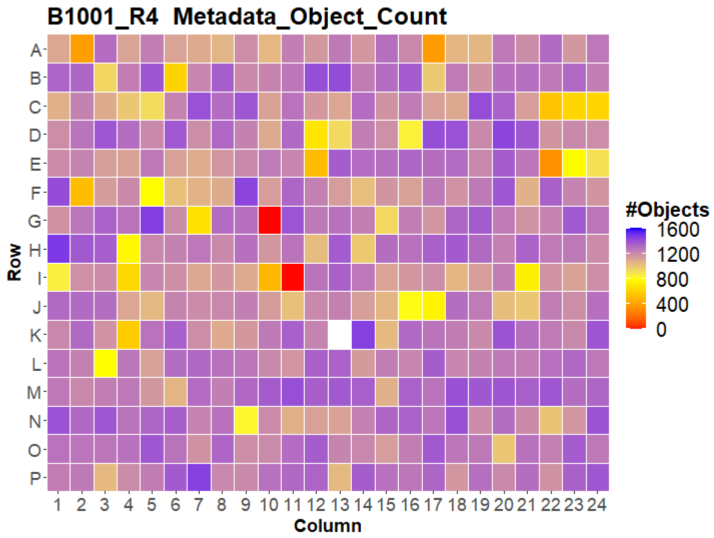
B1001_R4 Metadata_Object_Count
#Objects
1600
1200
800
400
0
Row
Column

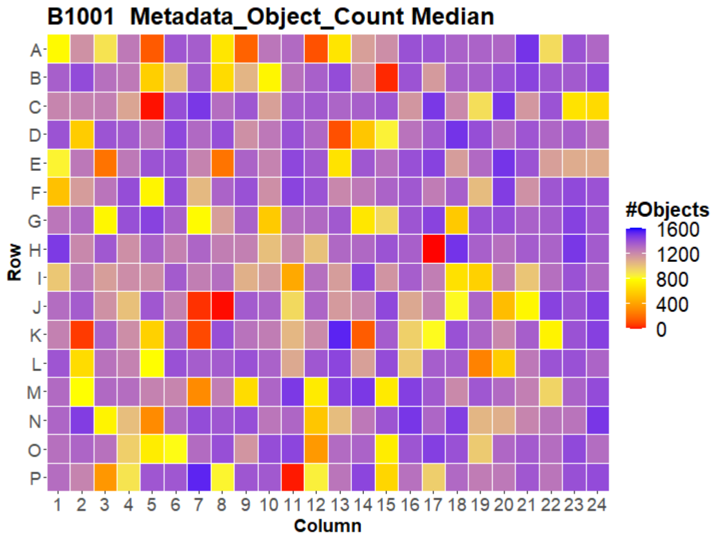
B1001 Metadata_Object_Count Median
#Objects
1600
1200
800
400
0
Row
Column

### HepG2 IMTM

23

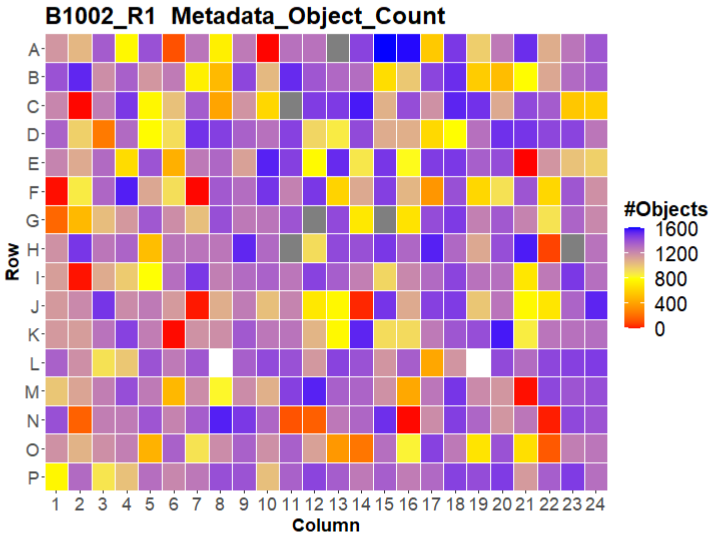


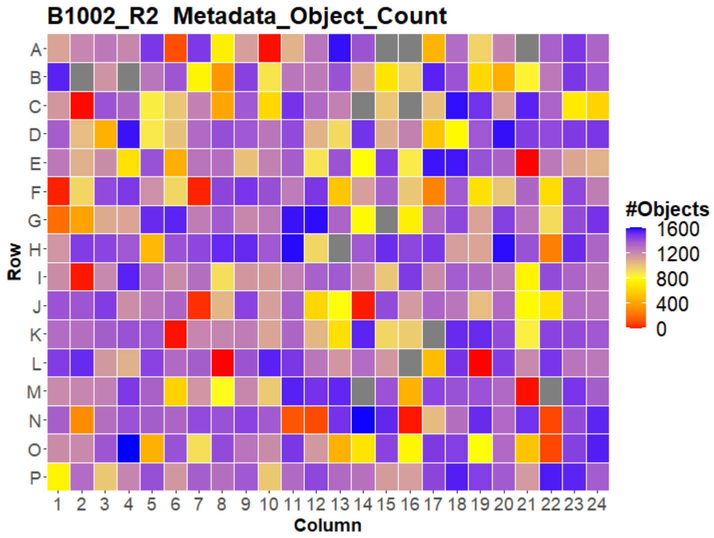


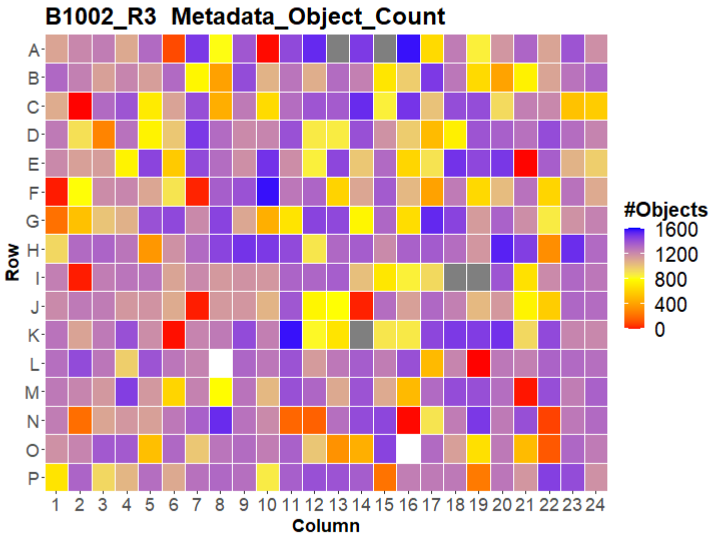


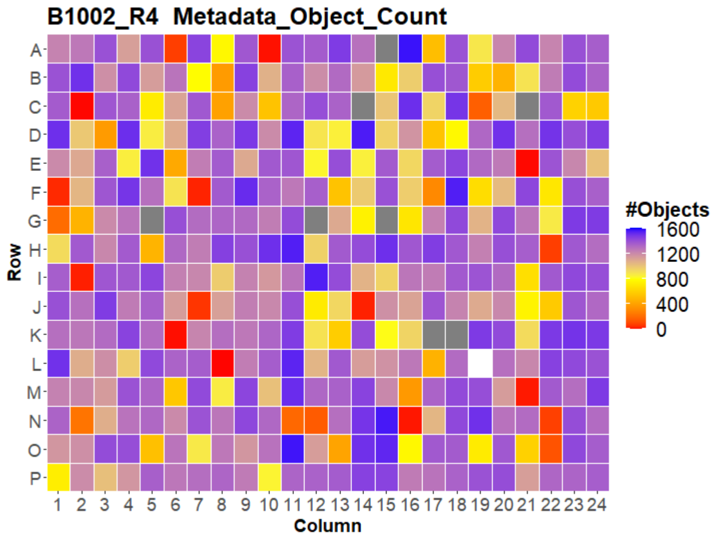


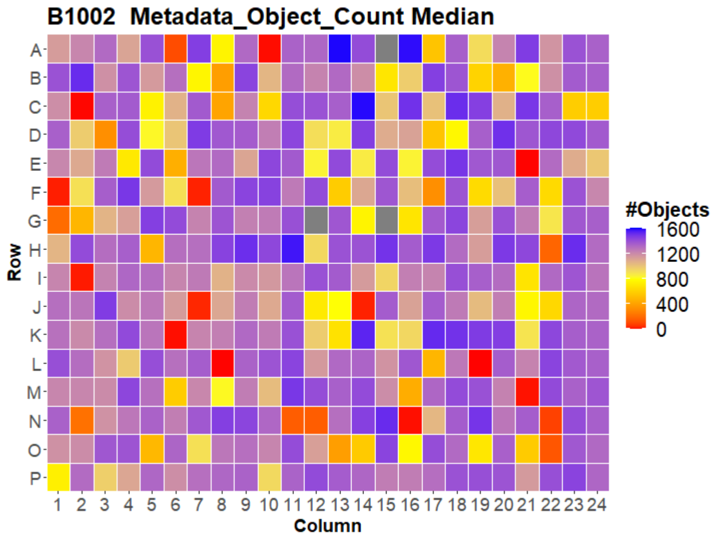

### HepG2 IMTM



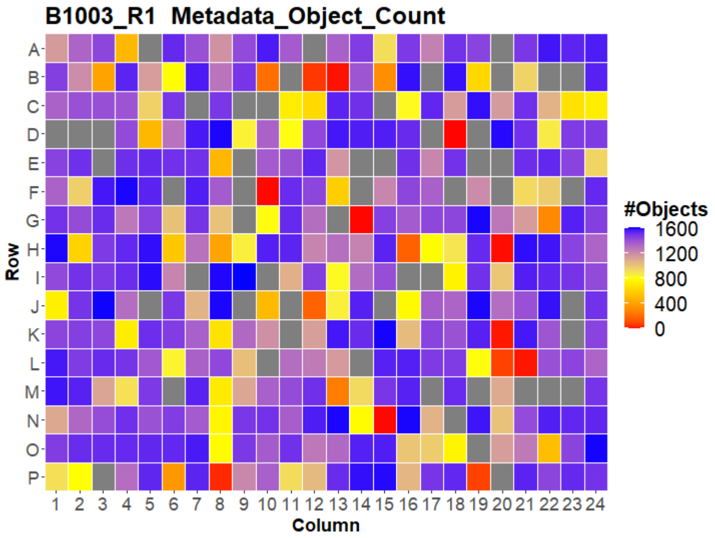
B1003_R1 Metadata_Object_Count
Row
A B C D E F G H I J K L M N O P
Column
1 2 3 4 5 6 7 8 9 10 11 12 13 14 15 16 17 18 19 20 21 22 23 24
#Objects
1600
1200
800
400
0

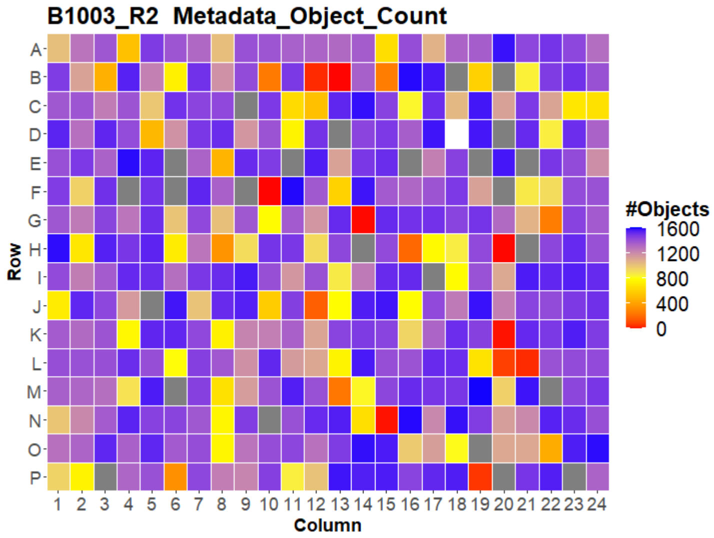
B1003_R2 Metadata_Object_Count
Row
A B C D E F G H I J K L M N O P
Column
1 2 3 4 5 6 7 8 9 10 11 12 13 14 15 16 17 18 19 20 21 22 23 24
#Objects
1600
1200
800
400
0

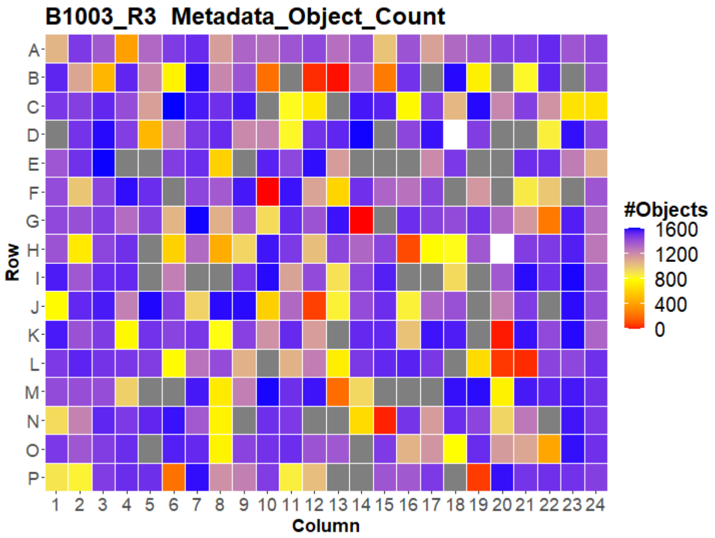
B1003_R3 Metadata_Object_Count
Row
A B C D E F G H I J K L M N O P
Column
1 2 3 4 5 6 7 8 9 10 11 12 13 14 15 16 17 18 19 20 21 22 23 24
#Objects
1600
1200
800
400
0

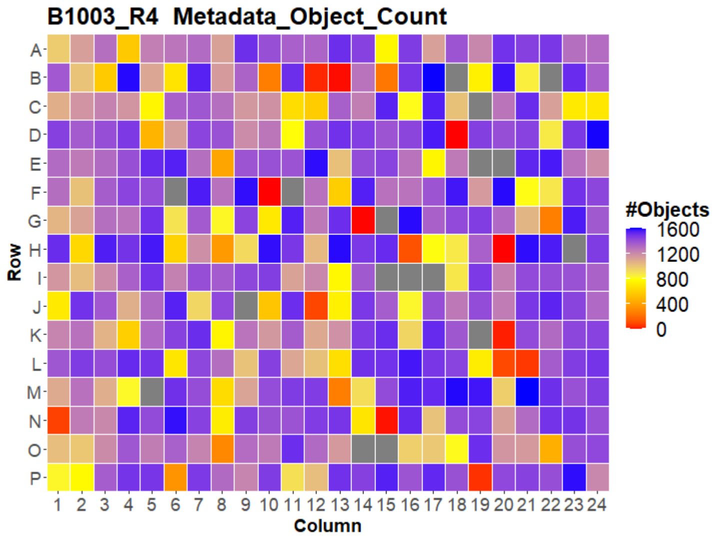
B1003_R4 Metadata_Object_Count
Row
A B C D E F G H I J K L M N O P
Column
1 2 3 4 5 6 7 8 9 10 11 12 13 14 15 16 17 18 19 20 21 22 23 24
#Objects
1600
1200
800
400
0

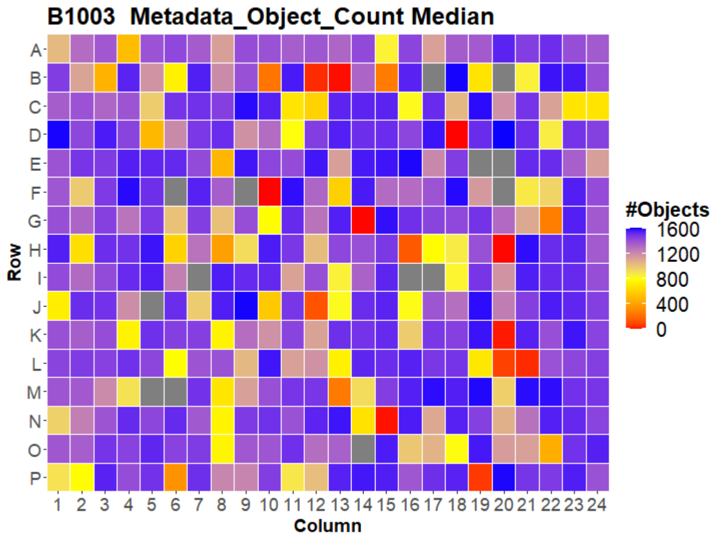
B1003 Metadata_Object_Count Median
Row
A B C D E F G H I J K L M N O P
Column
1 2 3 4 5 6 7 8 9 10 11 12 13 14 15 16 17 18 19 20 21 22 23 24
#Objects
1600
1200
800
400
0

### HepG2 IMTM



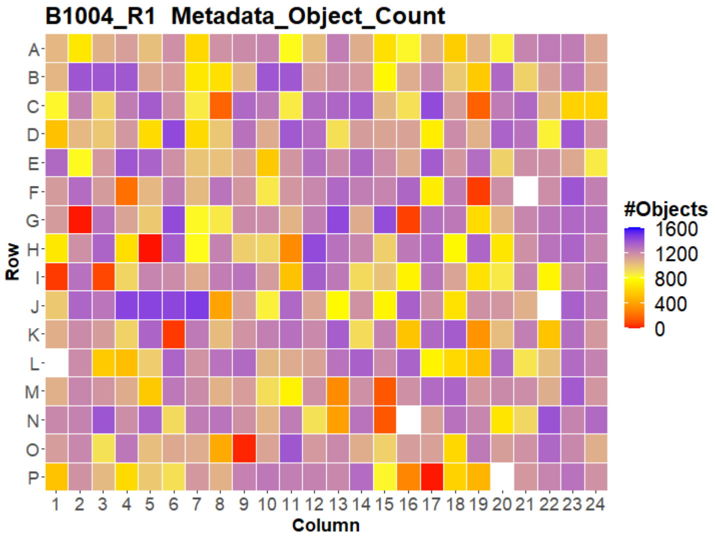
B1004_R1 Metadata_Object_Count
Row
Column
#Objects
1600
1200
800
400
0

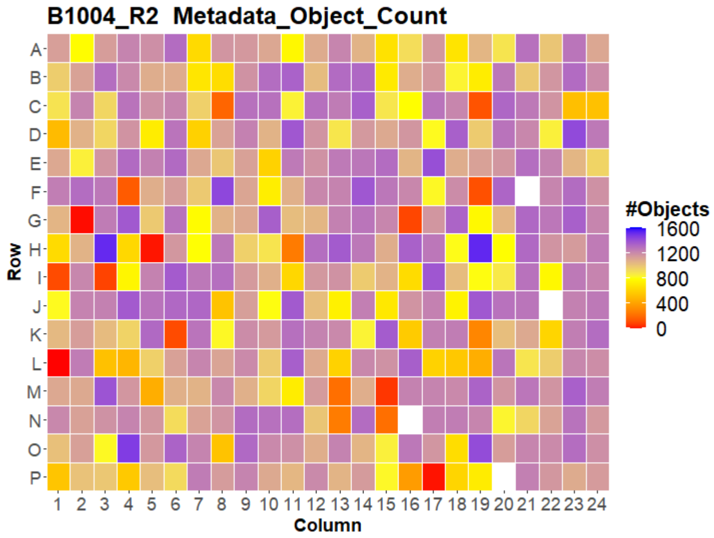
B1004_R2 Metadata_Object_Count
Row
Column
#Objects
1600
1200
800
400
0

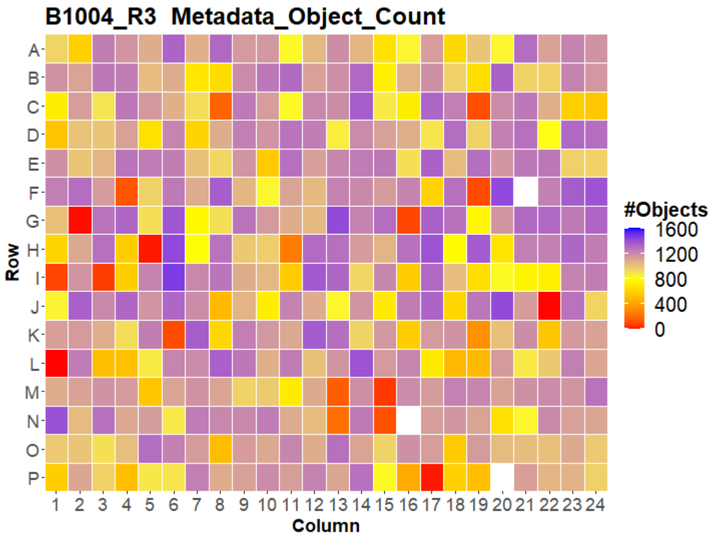
B1004_R3 Metadata_Object_Count
Row
Column
#Objects
1600
1200
800
400
0

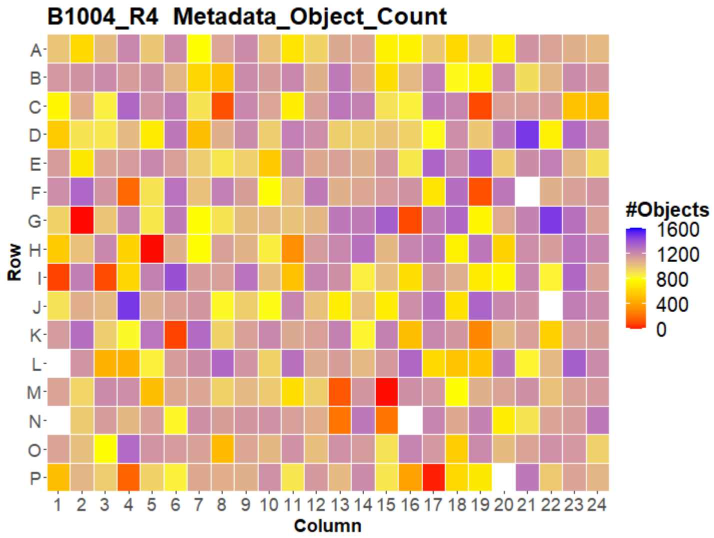
B1004_R4 Metadata_Object_Count
Row
Column
#Objects
1600
1200
800
400
0

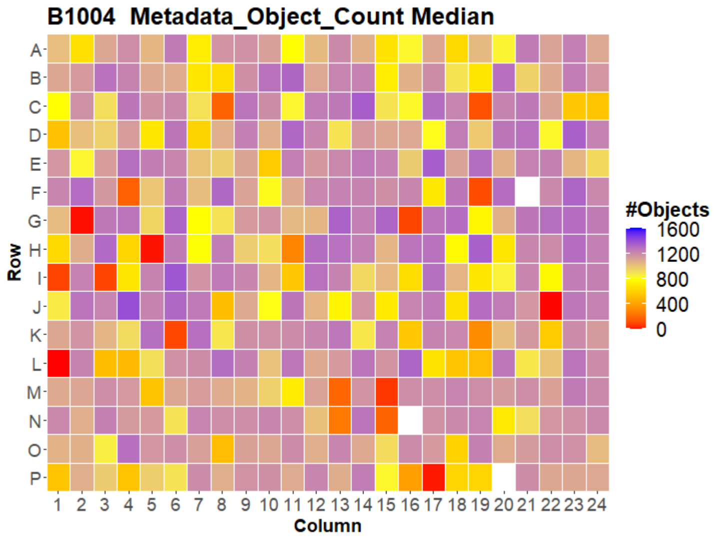
B1004 Metadata_Object_Count Median
Row
Column
#Objects
1600
1200
800
400
0

### HepG2 IMTM



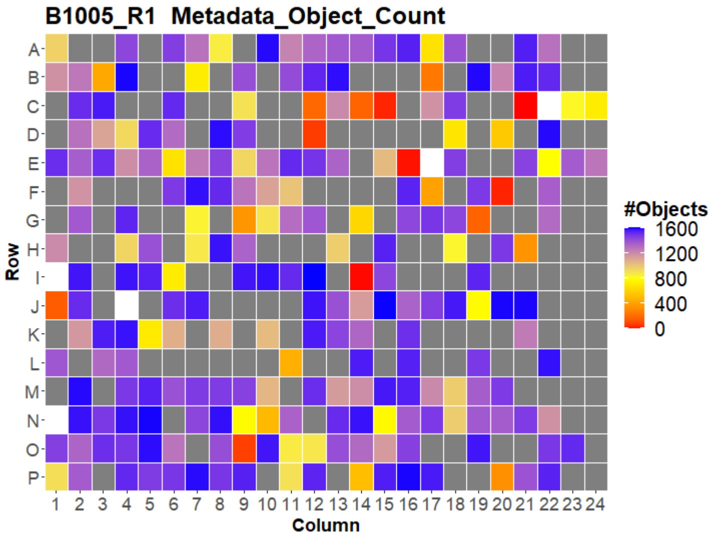
B1005_R1 Metadata_Object_Count
Row
Column
#Objects
1600
1200
800
400
0

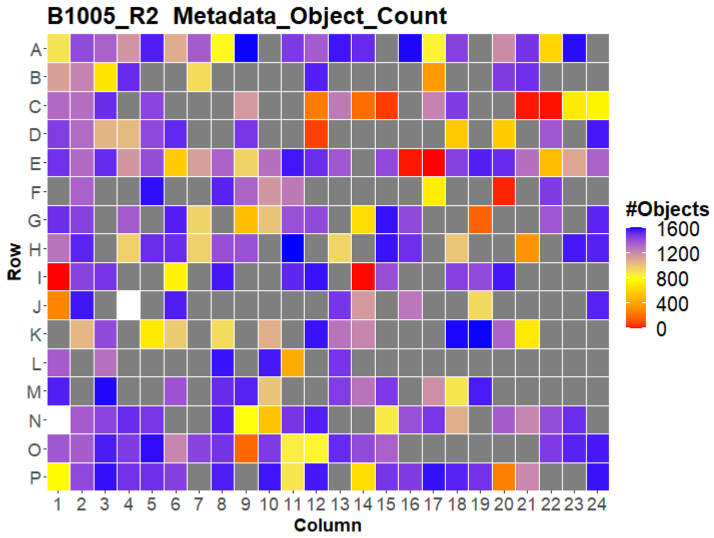
B1005_R2 Metadata_Object_Count
Row
Column
#Objects
1600
1200
800
400
0

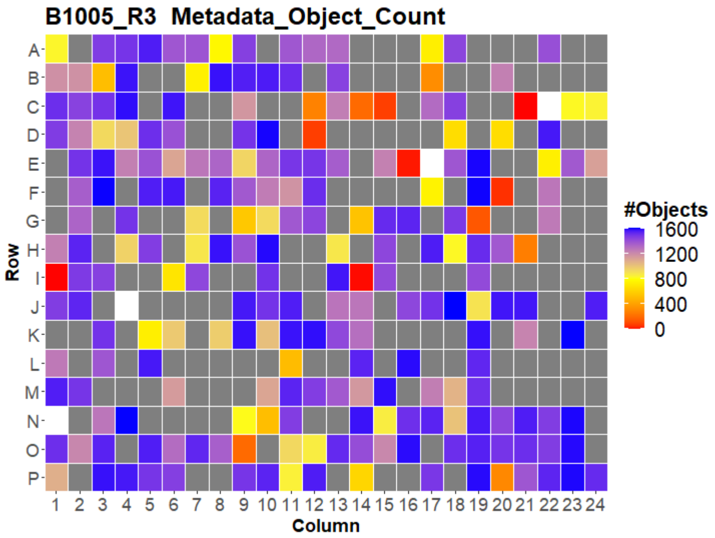
B1005_R3 Metadata_Object_Count
Row
Column
#Objects
1600
1200
800
400
0

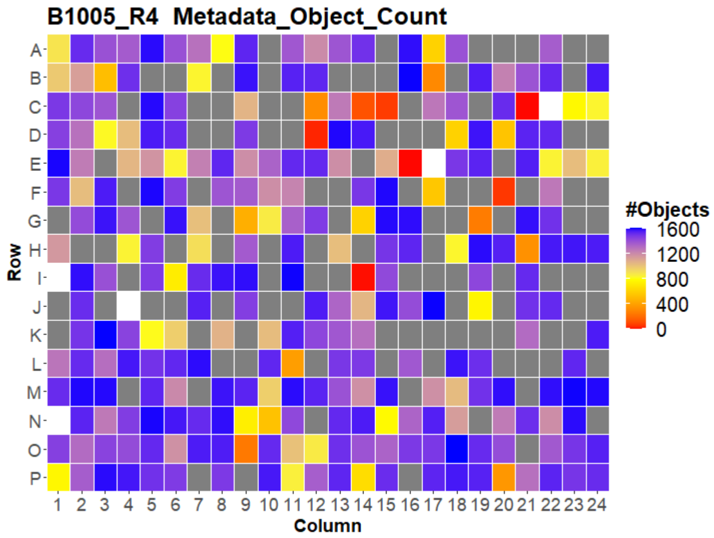
B1005_R4 Metadata_Object_Count
Row
Column
#Objects
1600
1200
800
400
0

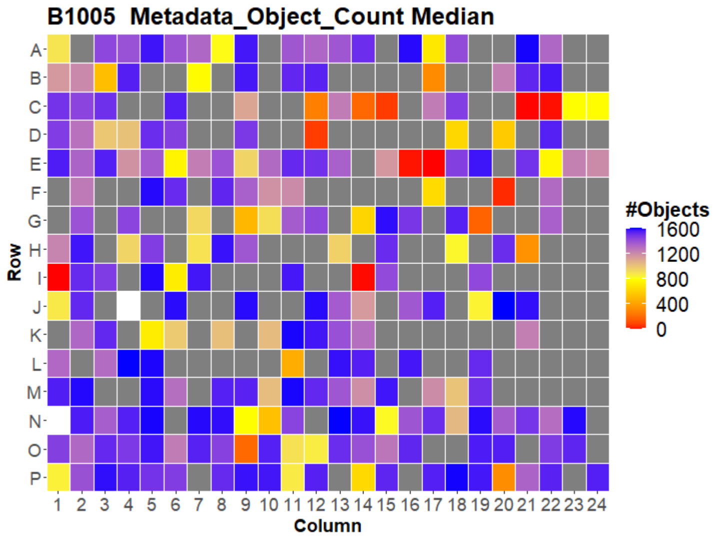
B1005 Metadata_Object_Count Median
Row
Column
#Objects
1600
1200
800
400
0

### HepG2 IMTM



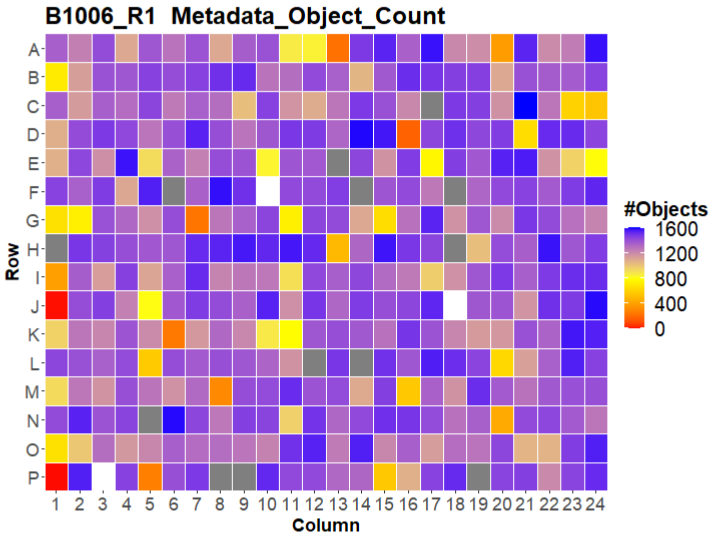


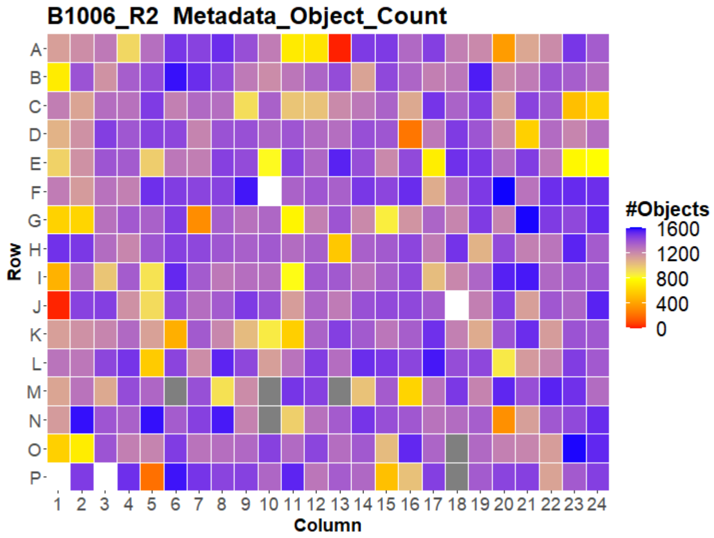


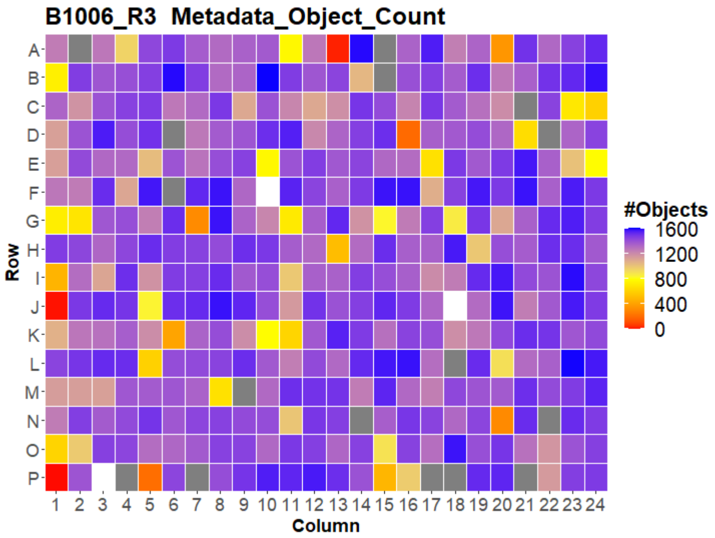


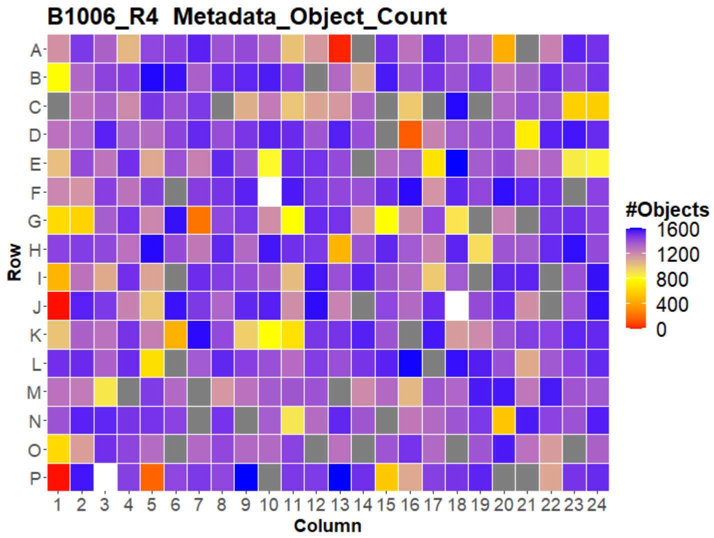


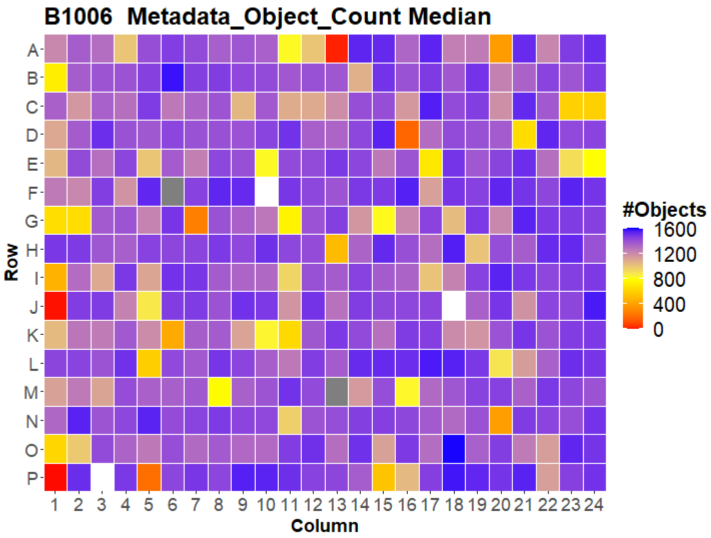



Open for Innovation KNIME

### HepG2 IMTM



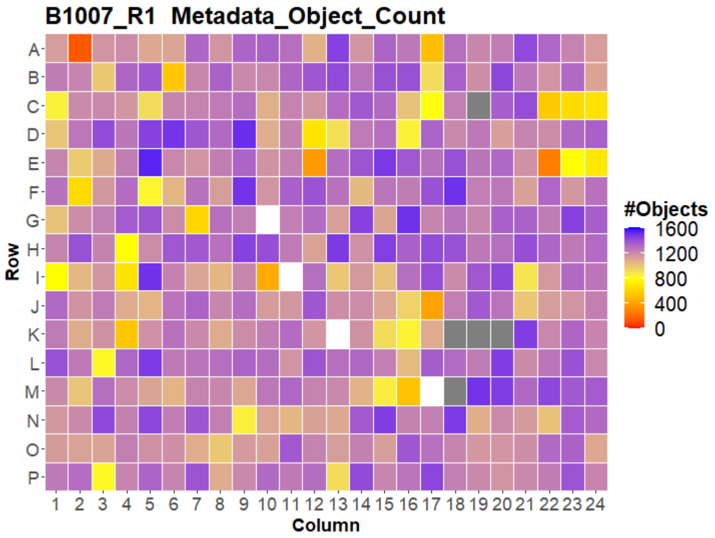


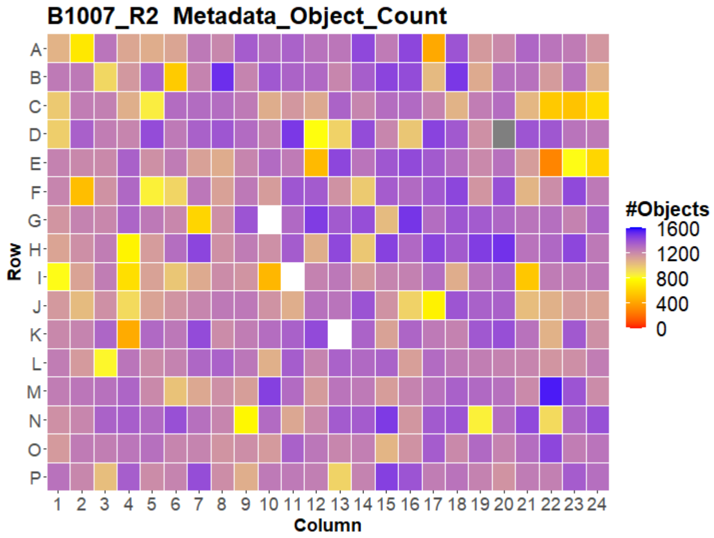


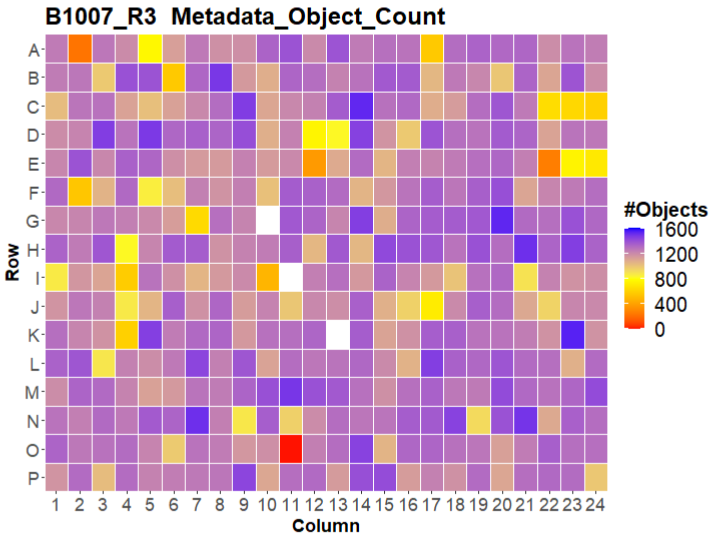


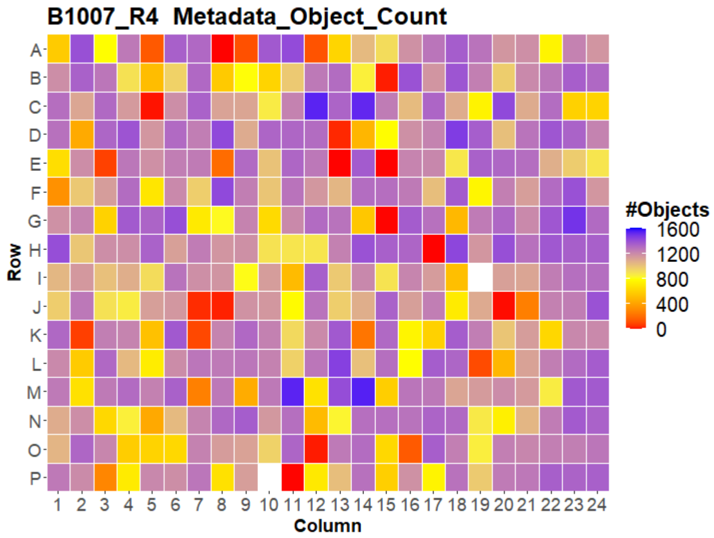


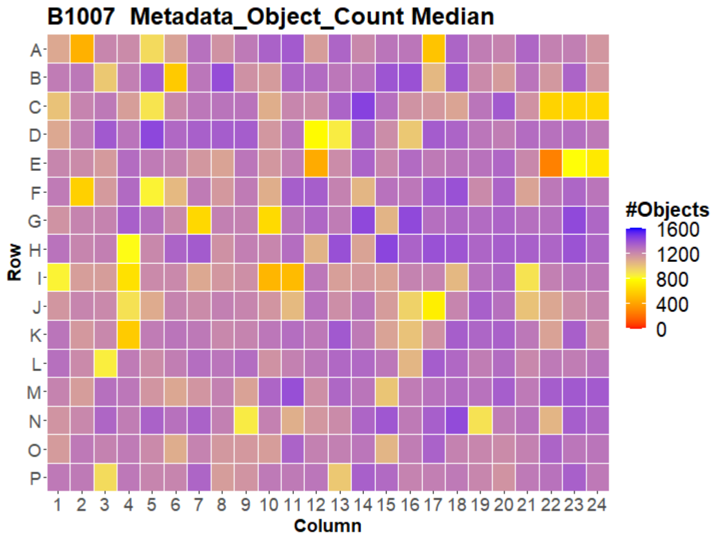




Open for Innovation KNIME

### HepG2 IMTM

29

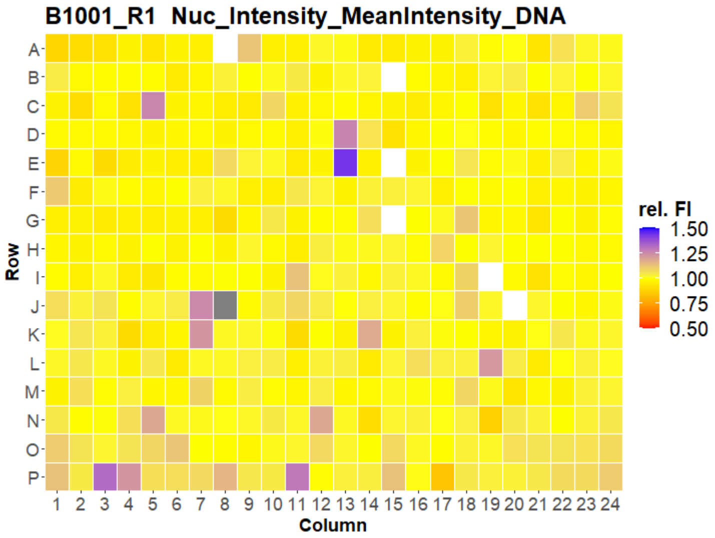


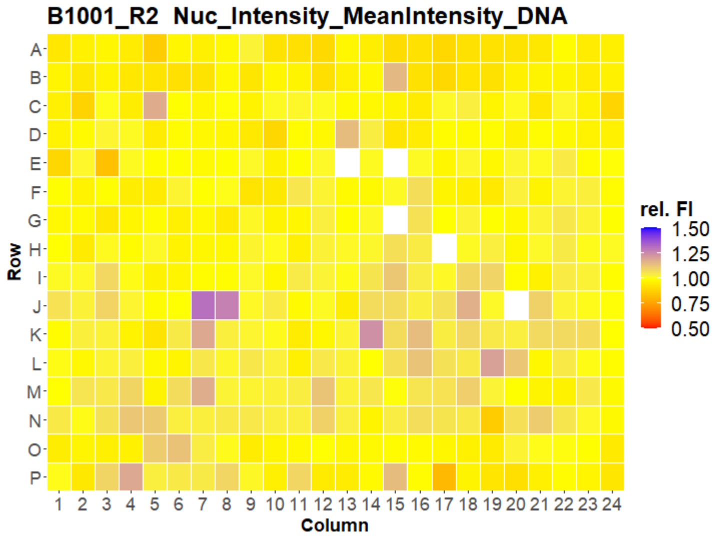


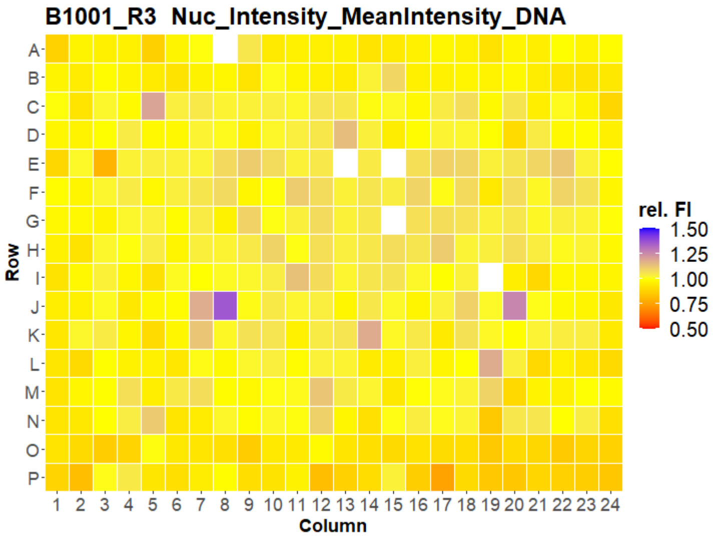


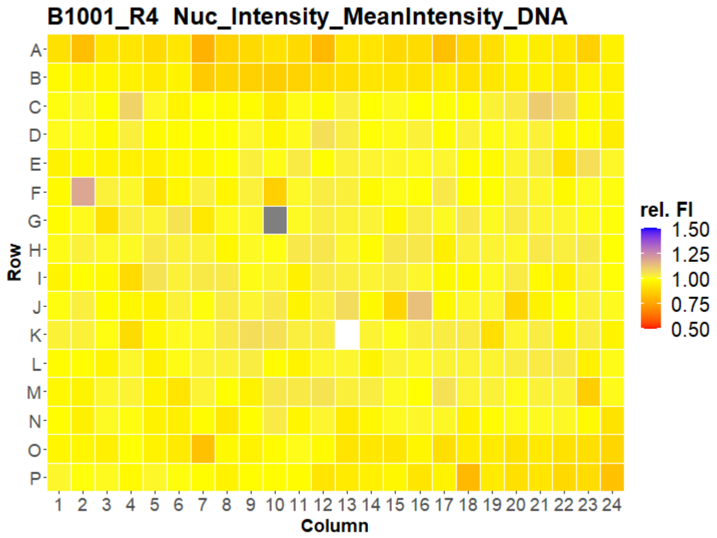


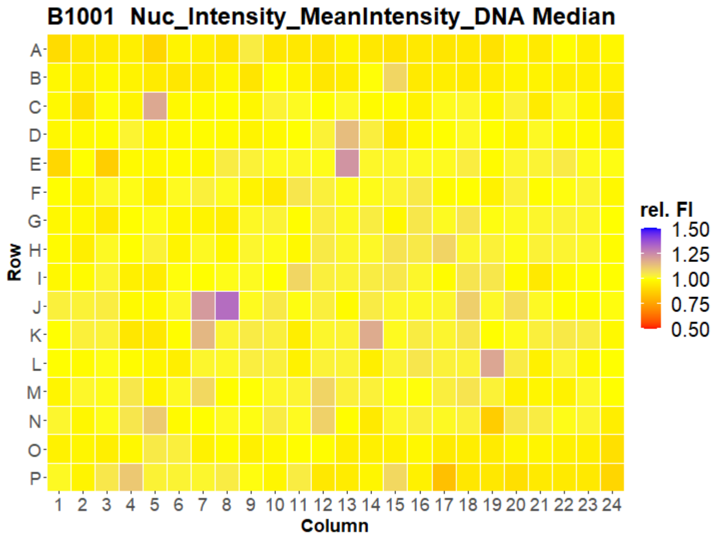

### HepG2 IMTM

30

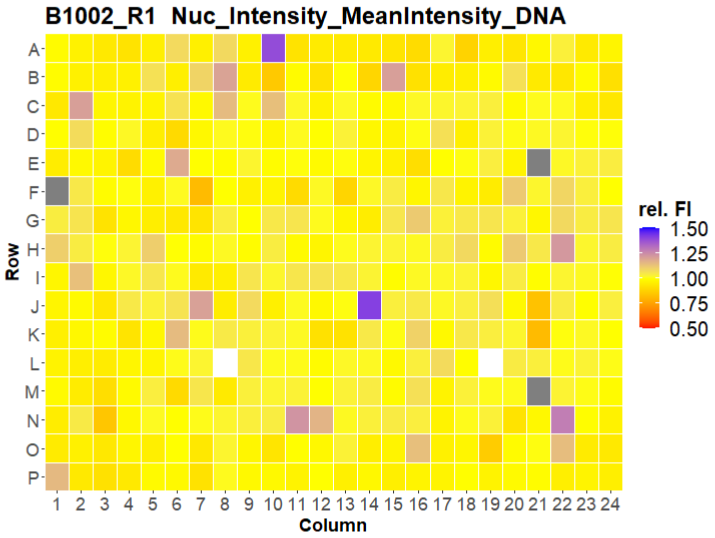


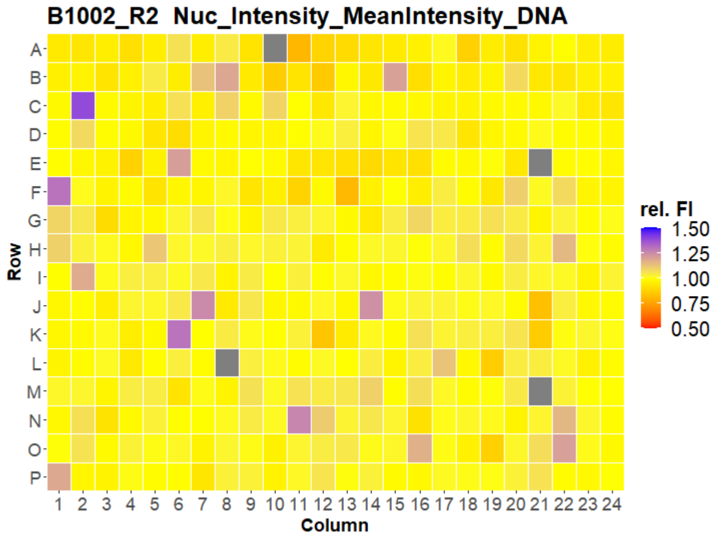


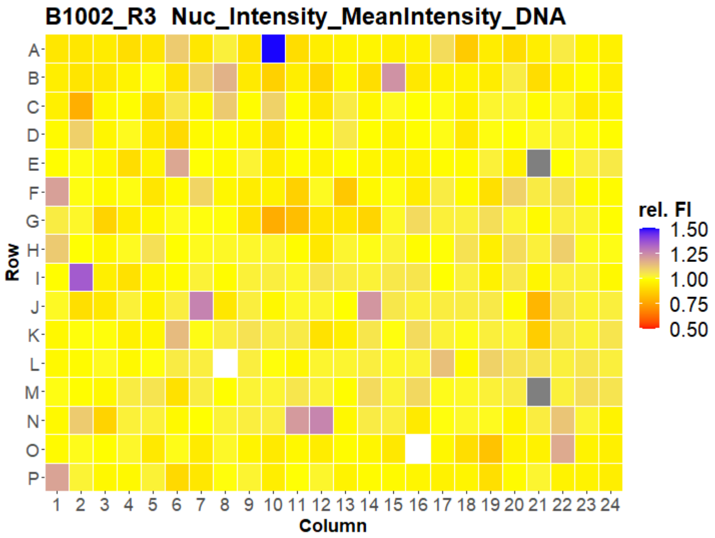


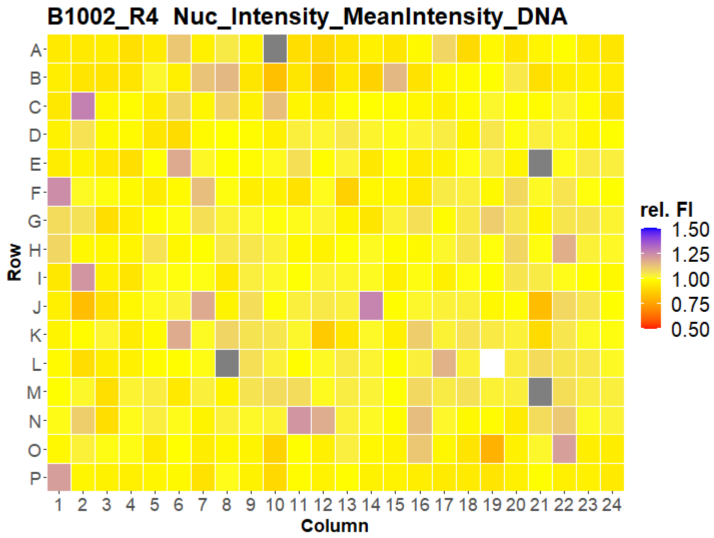


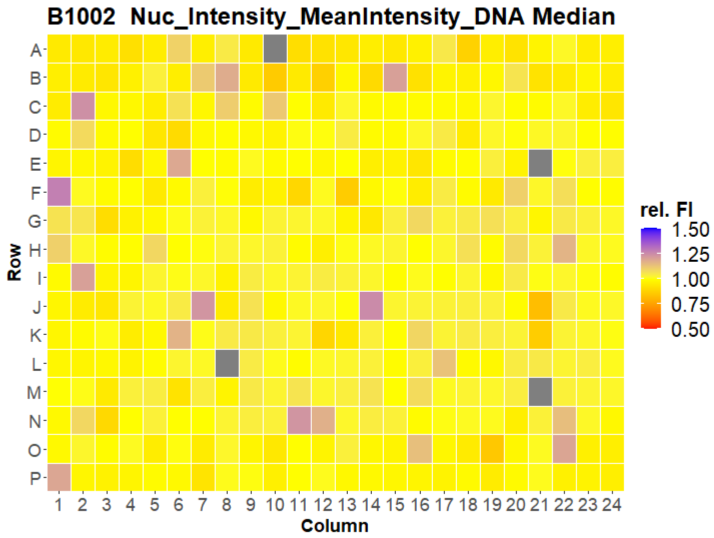

### HepG2 IMTM

31

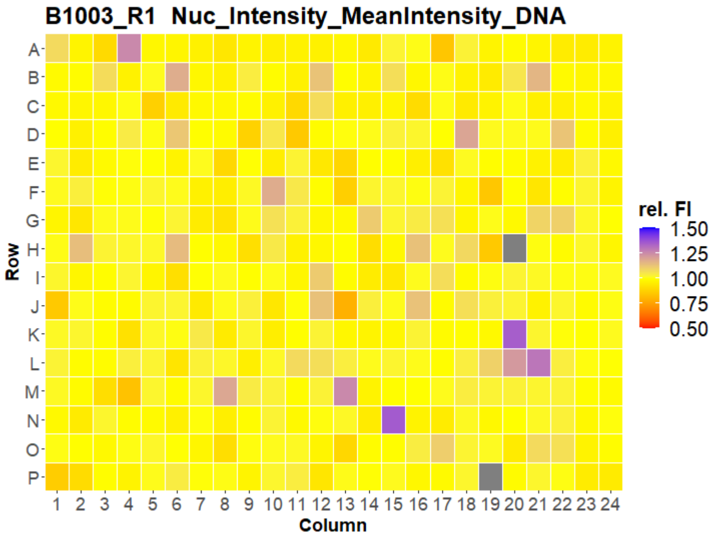


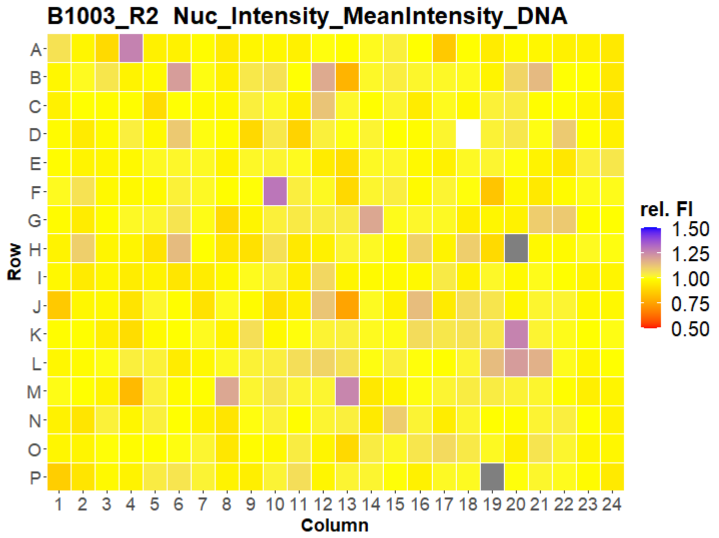


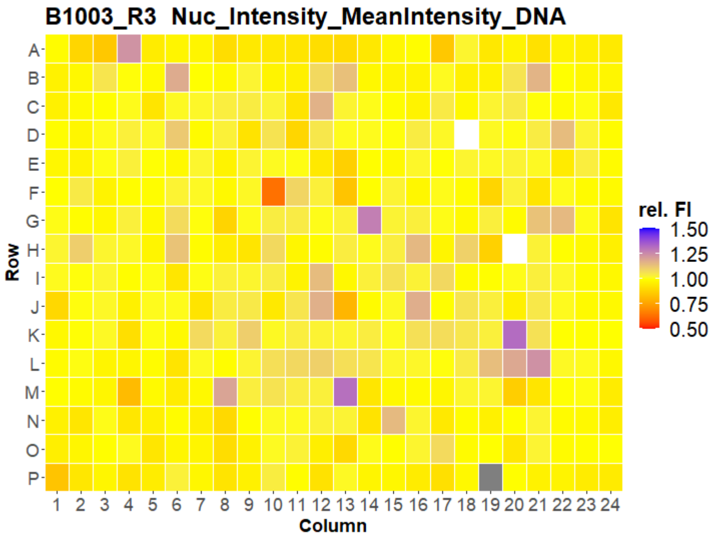


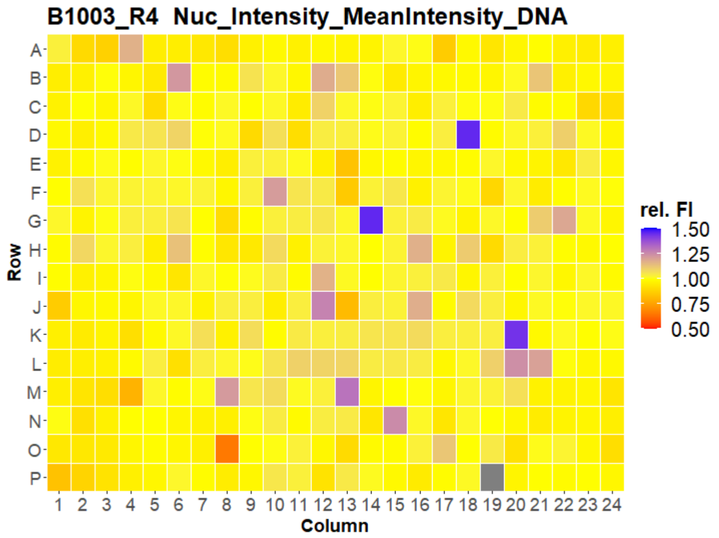


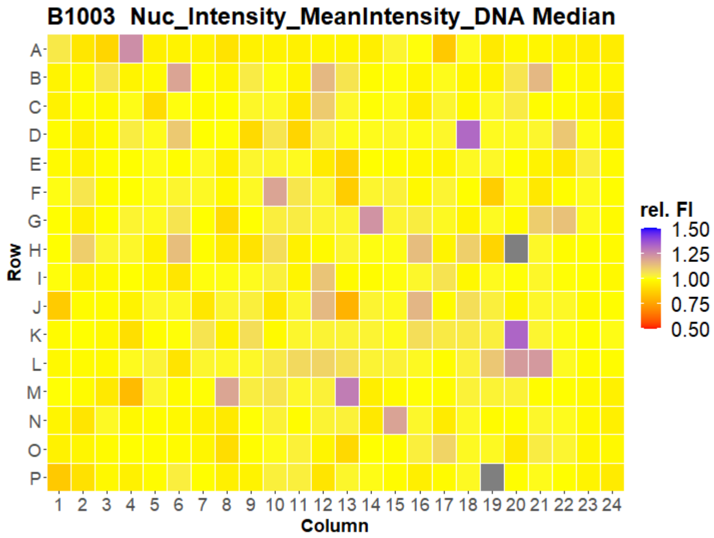

### HepG2 IMTM

32

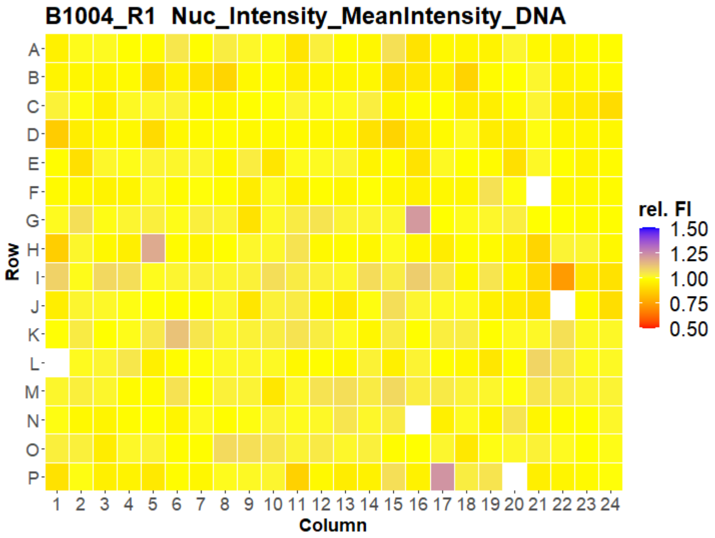


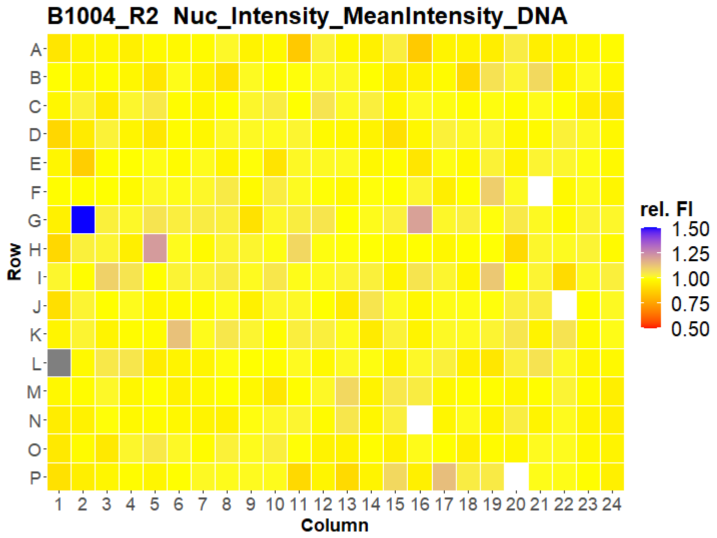


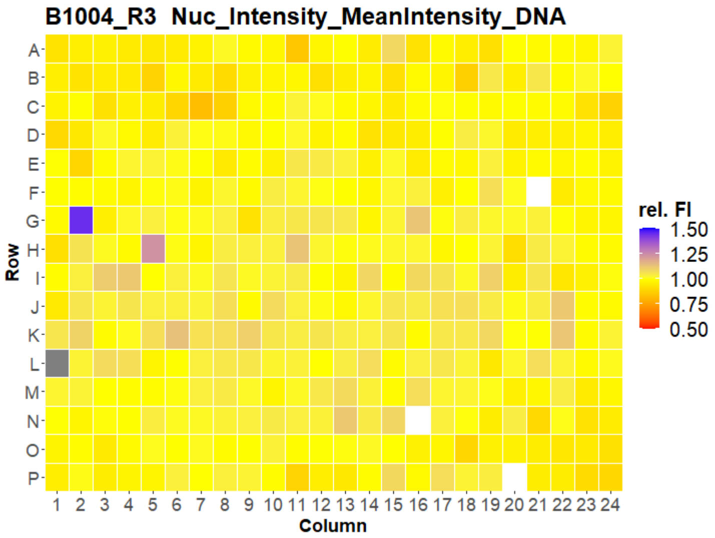


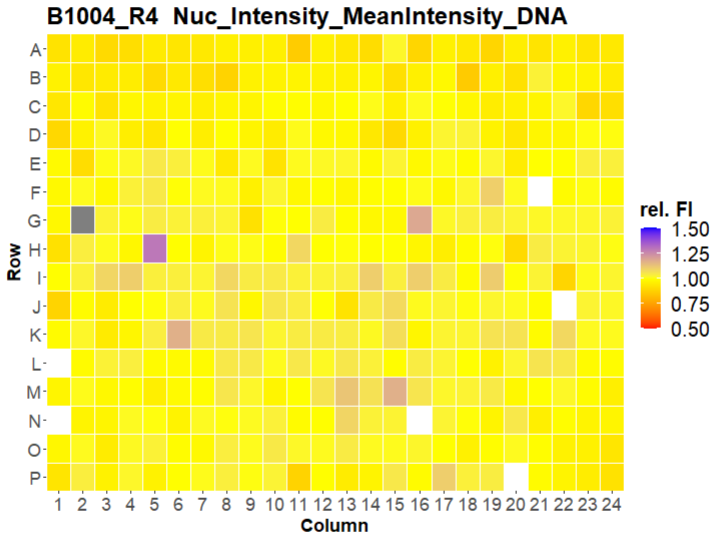


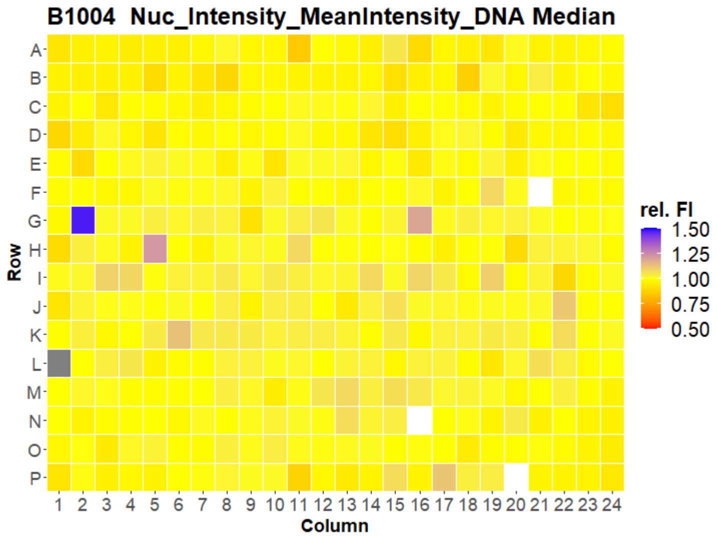

### HepG2 IMTM



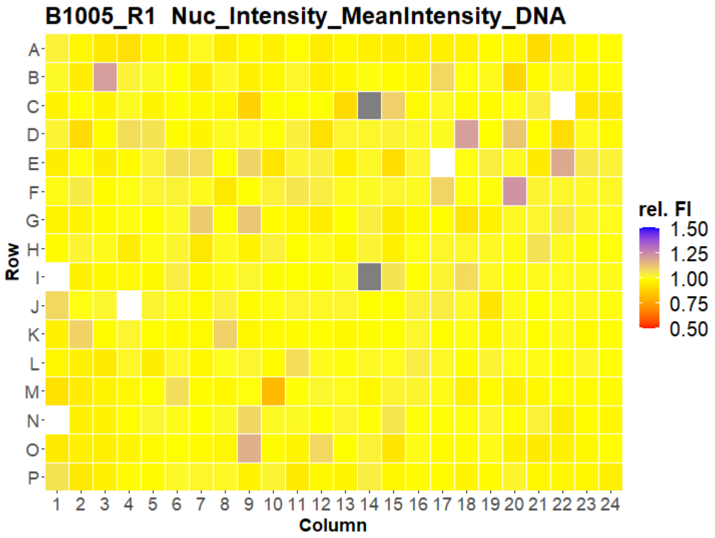
B1005_R1 Nuc_Intensity_MeanIntensity_DNA
Row
Column
rel. FI
1.50
1.25
1.00
0.75
0.50

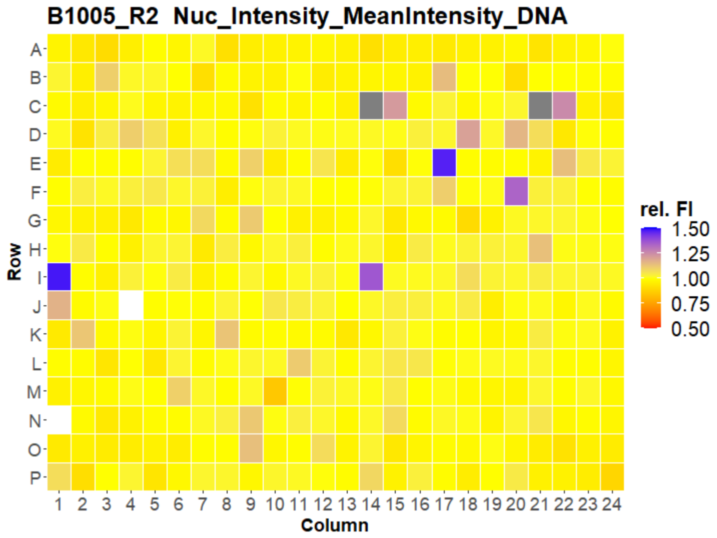
B1005_R2 Nuc_Intensity_MeanIntensity_DNA
Row
Column
rel. FI
1.50
1.25
1.00
0.75
0.50

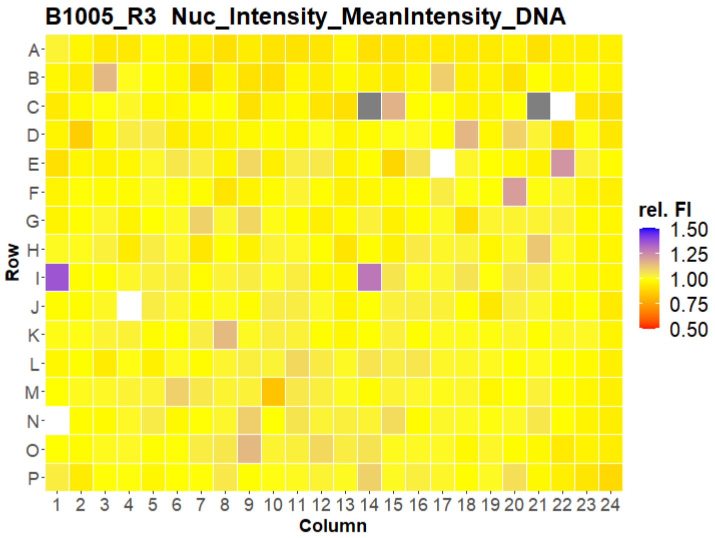
B1005_R3 Nuc_Intensity_MeanIntensity_DNA
Row
Column
rel. FI
1.50
1.25
1.00
0.75
0.50

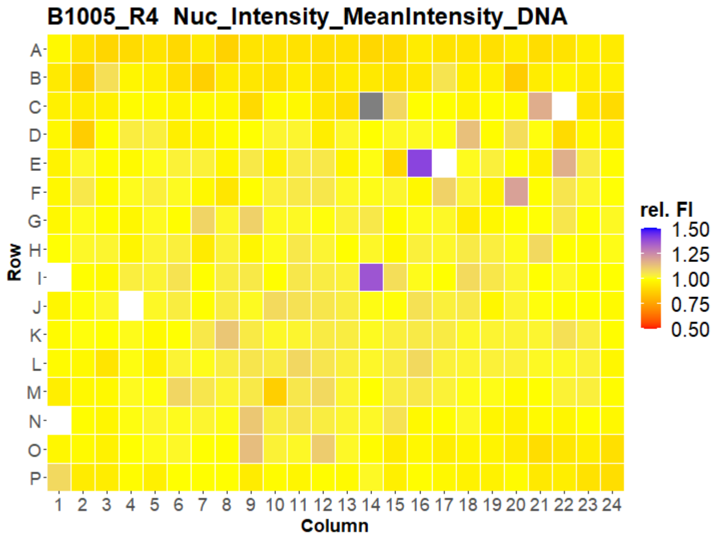
B1005_R4 Nuc_Intensity_MeanIntensity_DNA
Row
Column
rel. FI
1.50
1.25
1.00
0.75
0.50

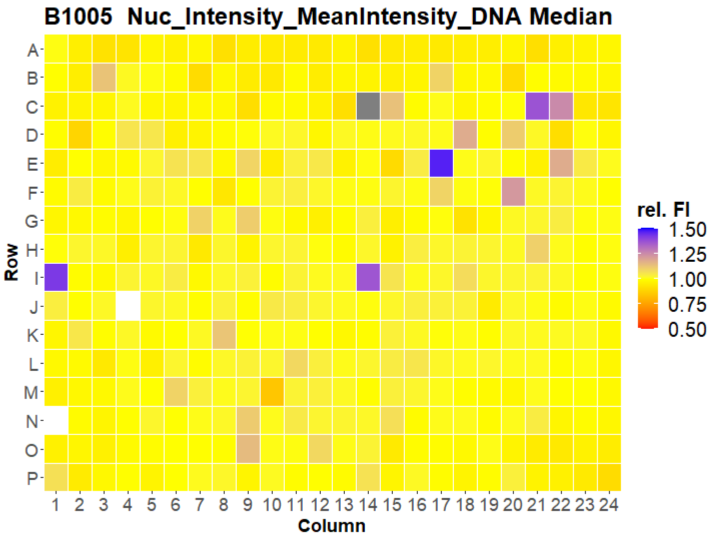
B1005 Nuc_Intensity_MeanIntensity_DNA Median
Row
Column
rel. FI
1.50
1.25
1.00
0.75
0.50

### HepG2 IMTM



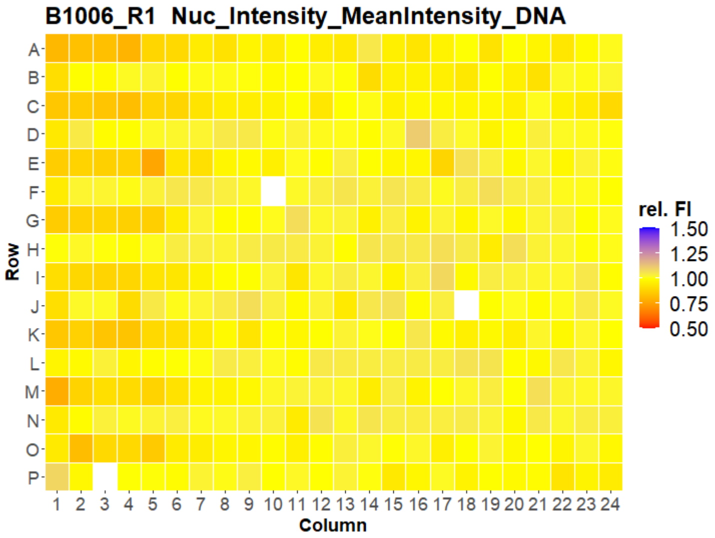


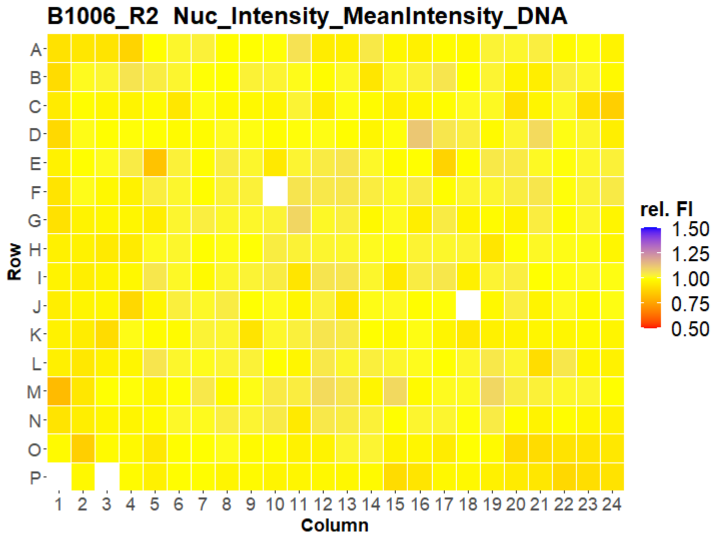


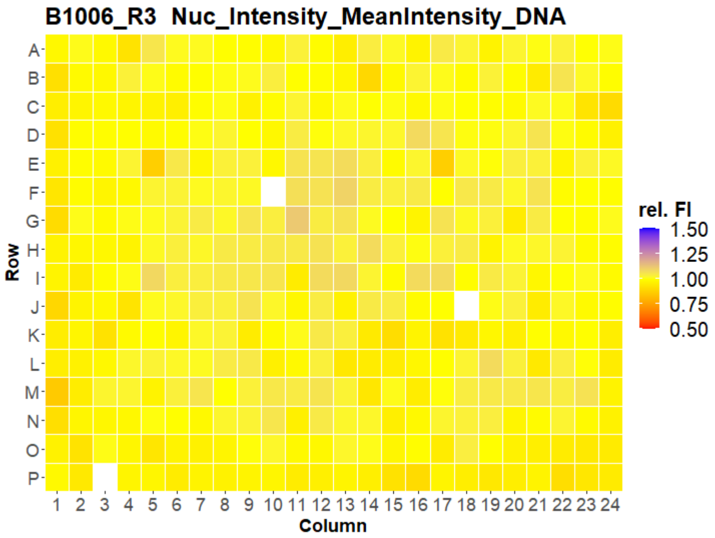


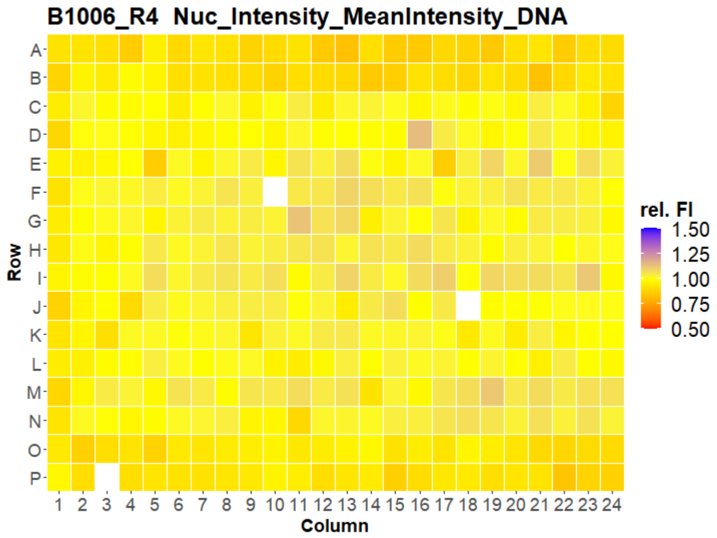


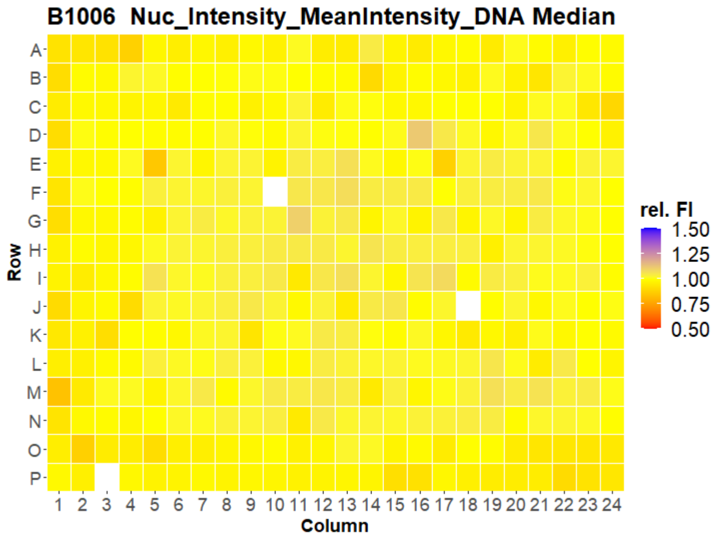



Open for Innovation KNIME

### HepG2 IMTM

35

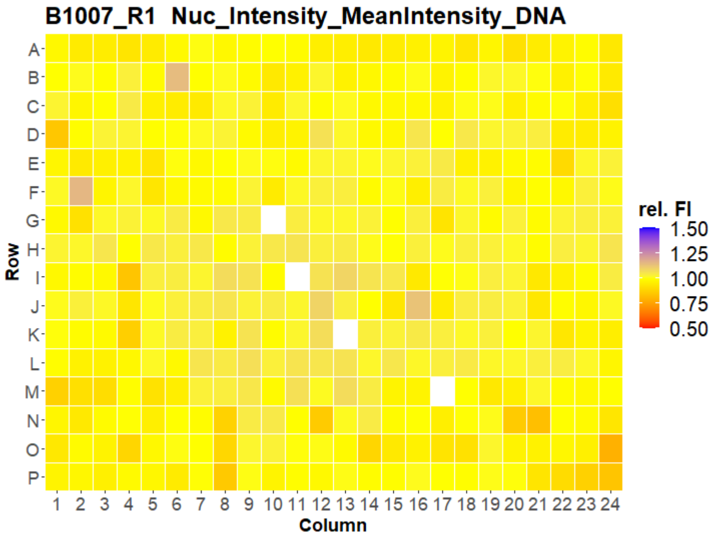


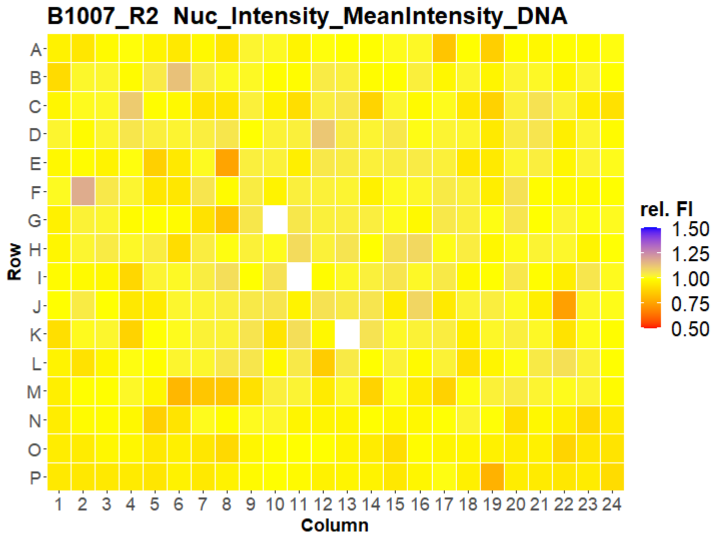


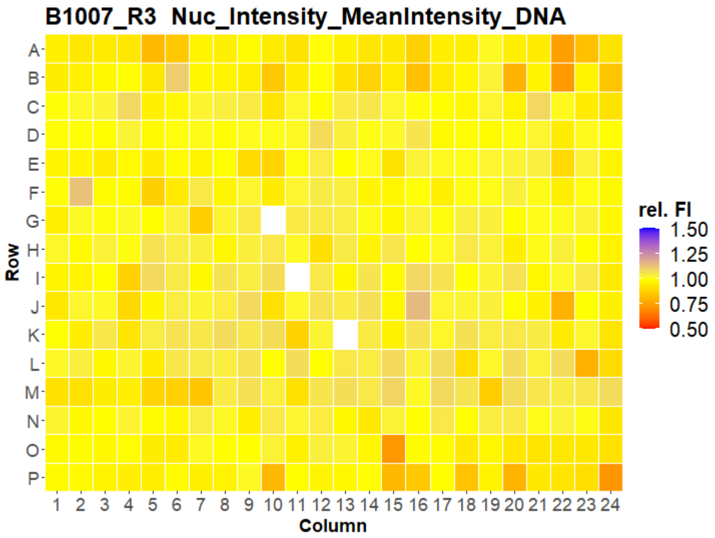


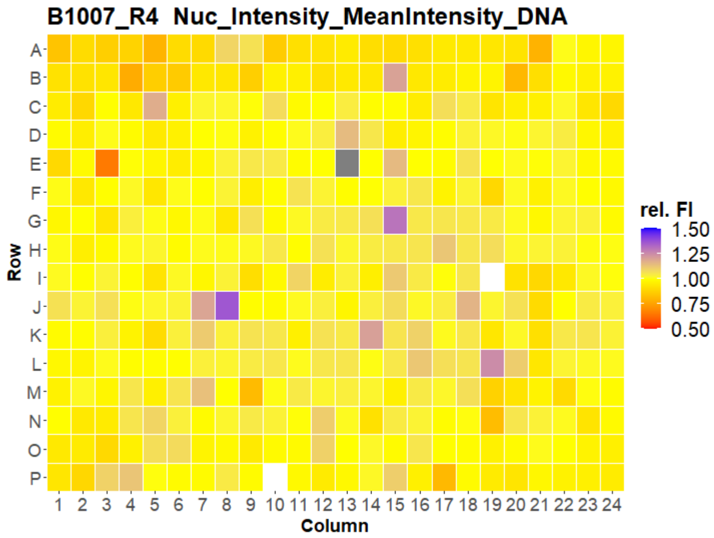


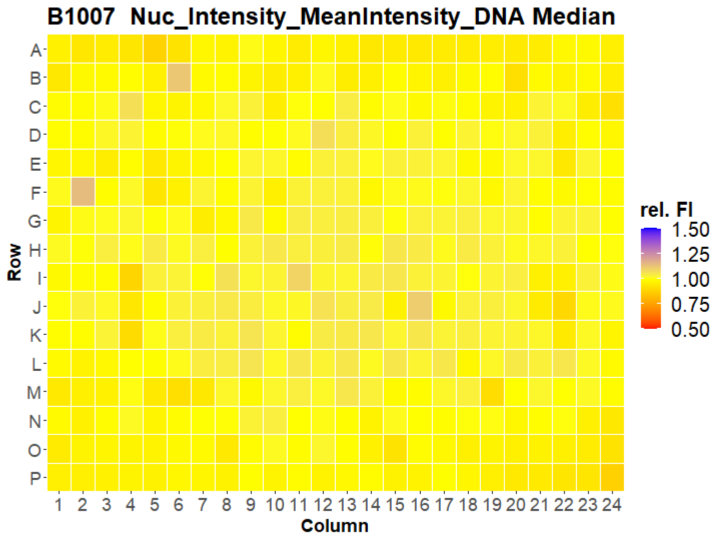

### HepG2 IMTM



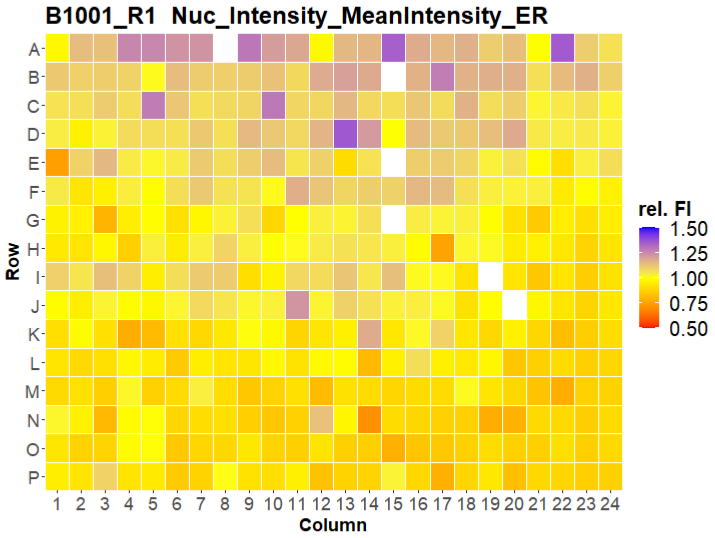
B1001_R1 Nuc_Intensity_MeanIntensity_ER
Row
Column
rel. FI
1.50
1.25
1.00
0.75
0.50

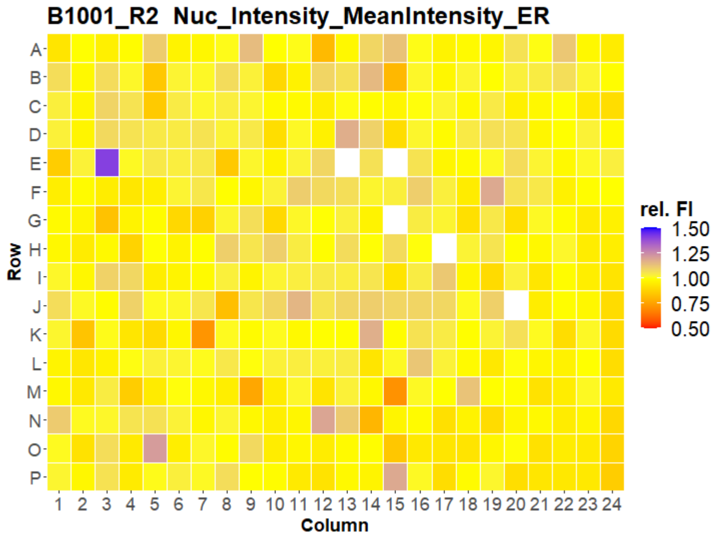
B1001_R2 Nuc_Intensity_MeanIntensity_ER
Row
Column
rel. FI
1.50
1.25
1.00
0.75
0.50

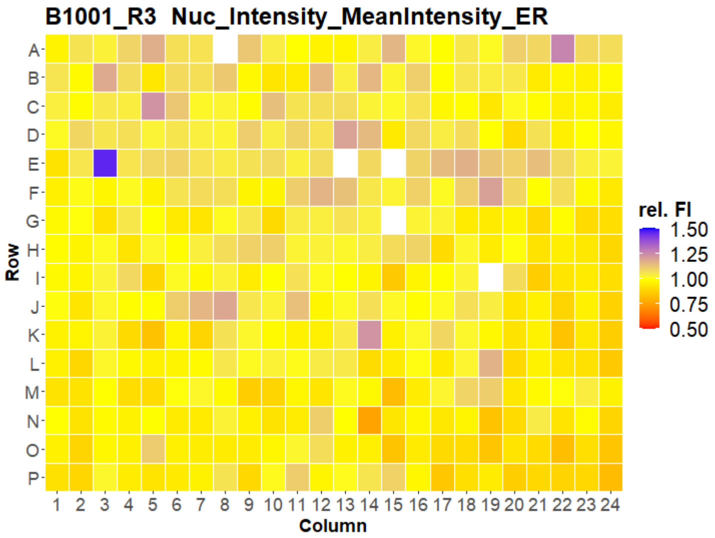
B1001_R3 Nuc_Intensity_MeanIntensity_ER
Row
Column
rel. FI
1.50
1.25
1.00
0.75
0.50

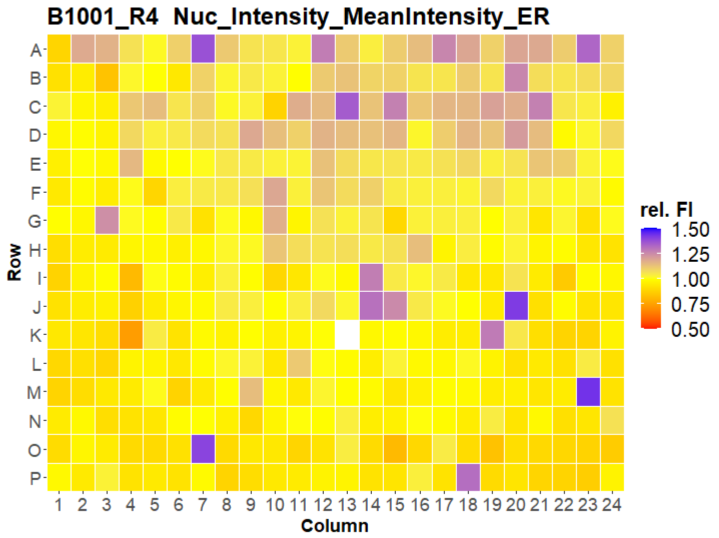
B1001_R4 Nuc_Intensity_MeanIntensity_ER
Row
Column
rel. FI
1.50
1.25
1.00
0.75
0.50

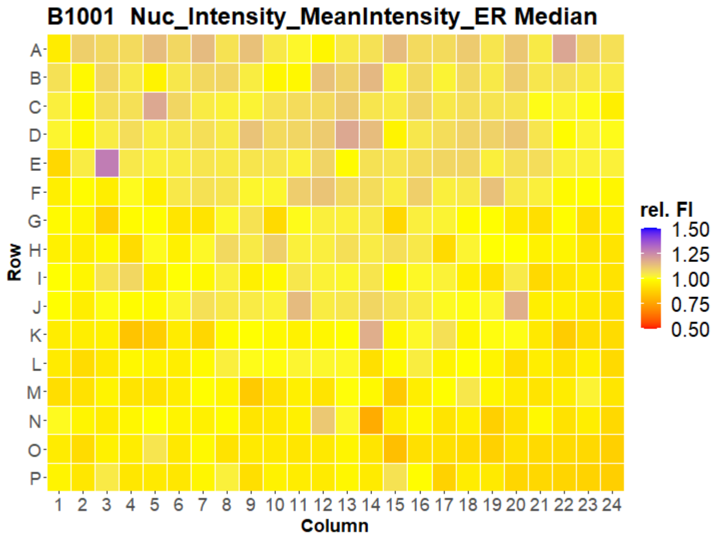
B1001 Nuc_Intensity_MeanIntensity_ER Median
Row
Column
rel. FI
1.50
1.25
1.00
0.75
0.50

### HepG2 IMTM

37

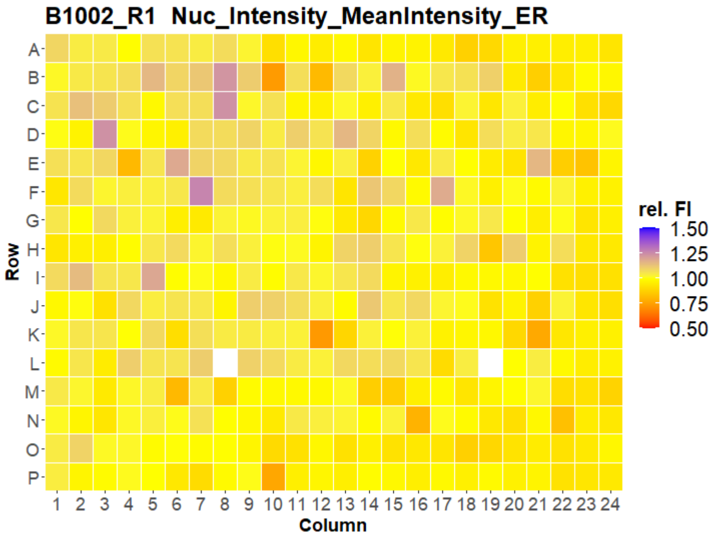


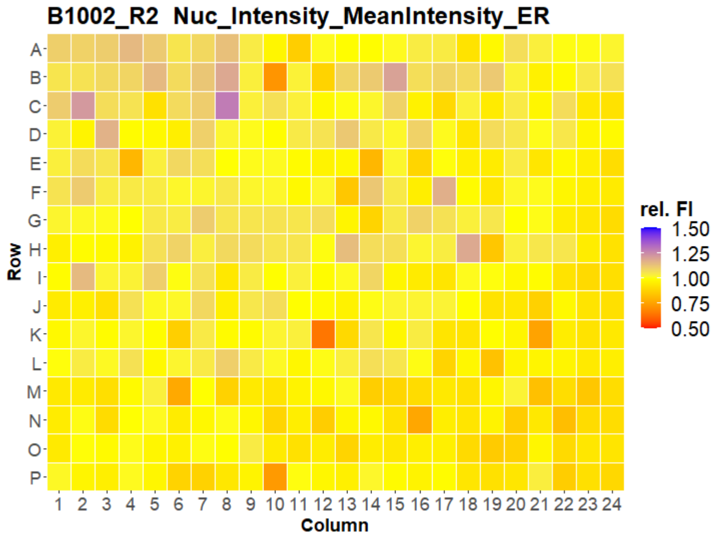


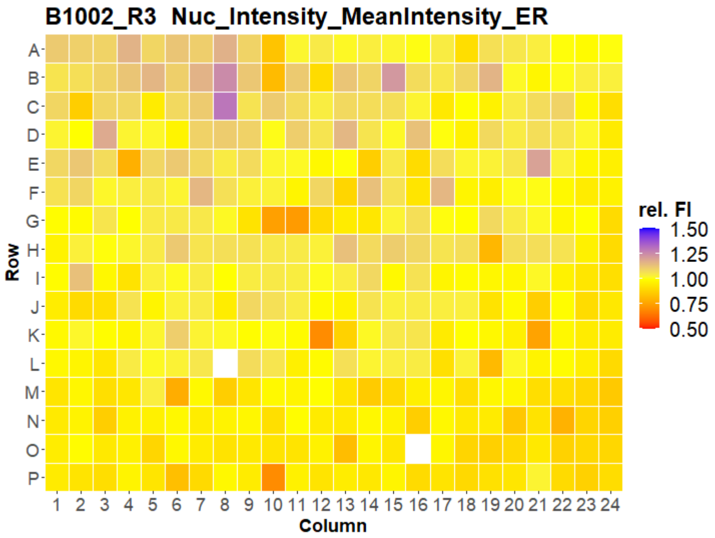


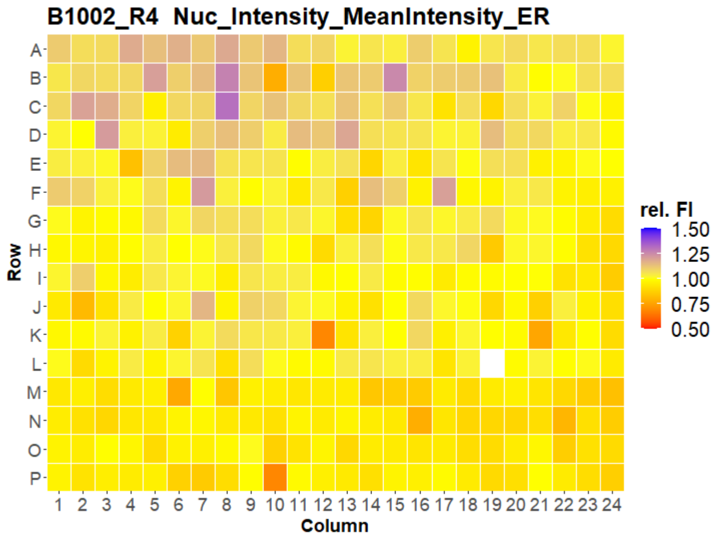


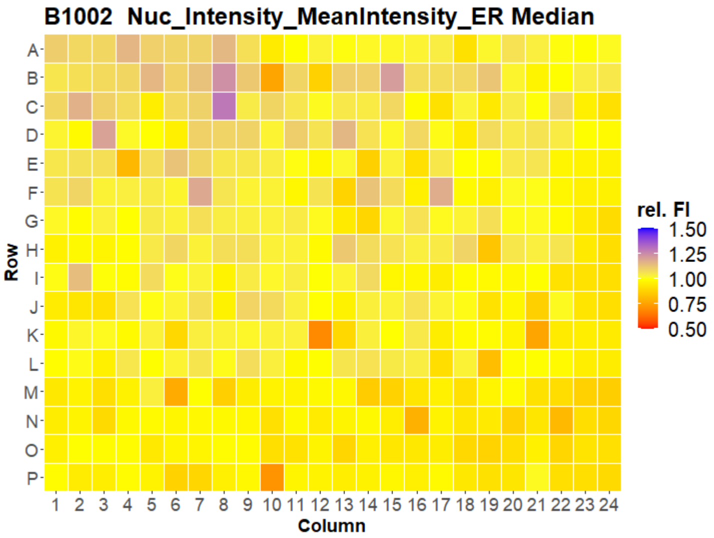

### HepG2 IMTM

38

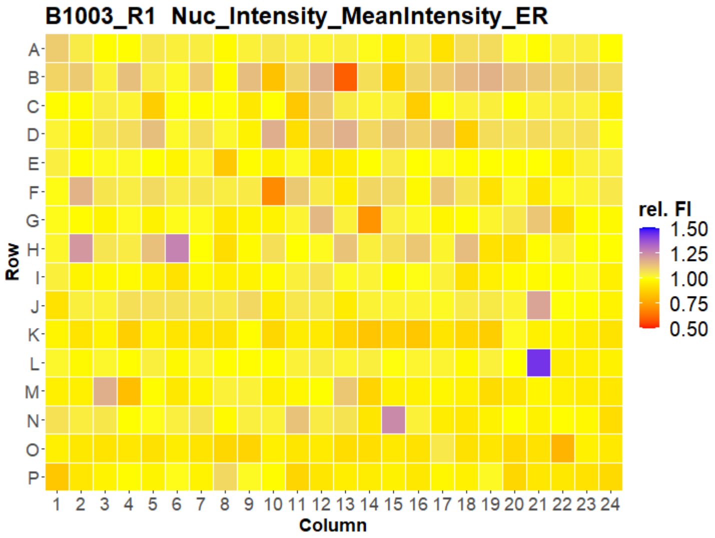


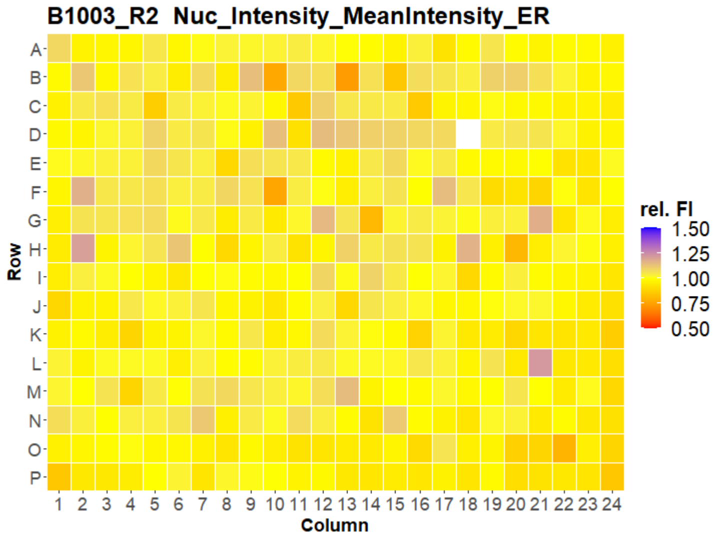


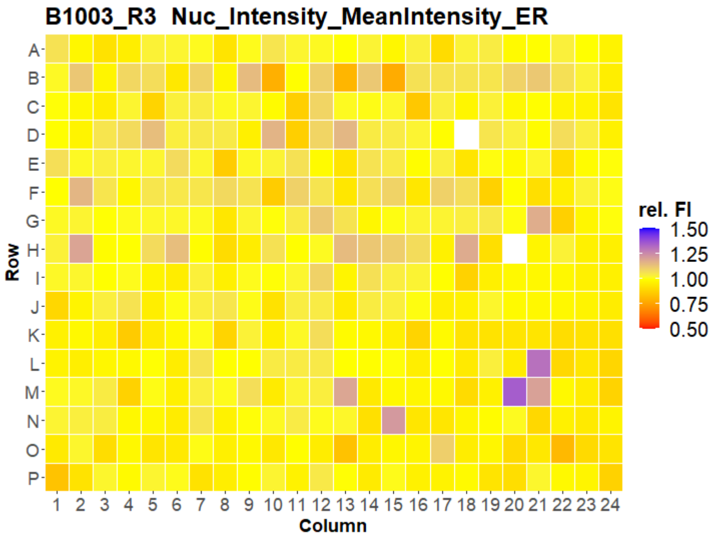


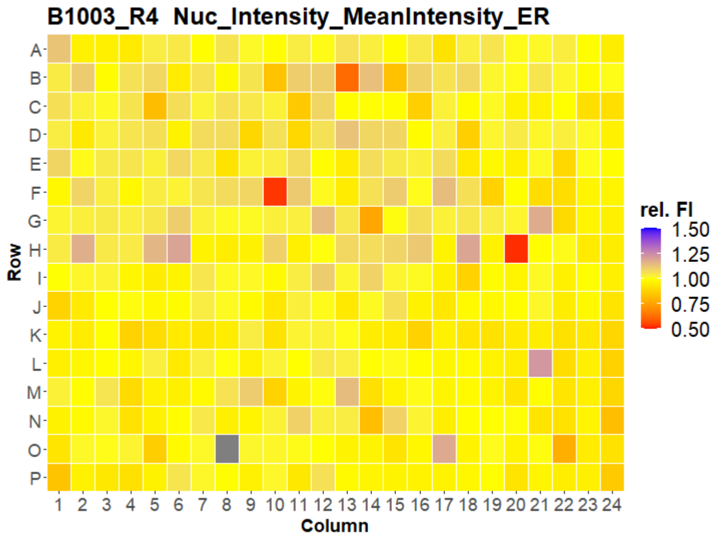


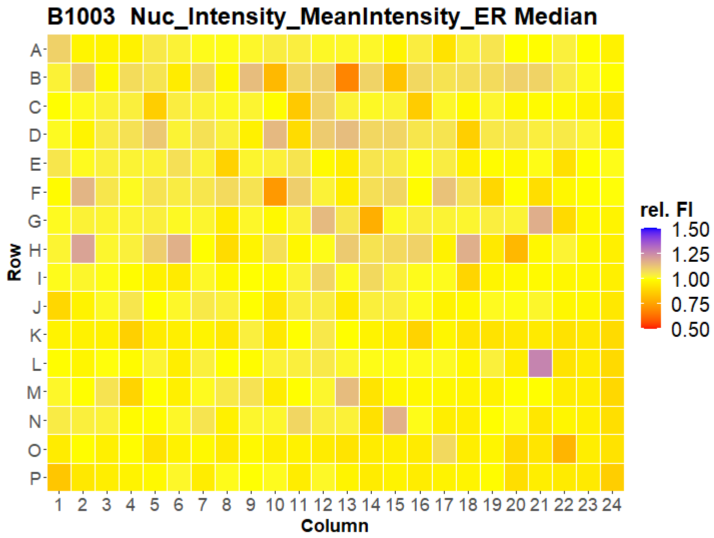



Open for Innovation KNIME

### HepG2 IMTM

39

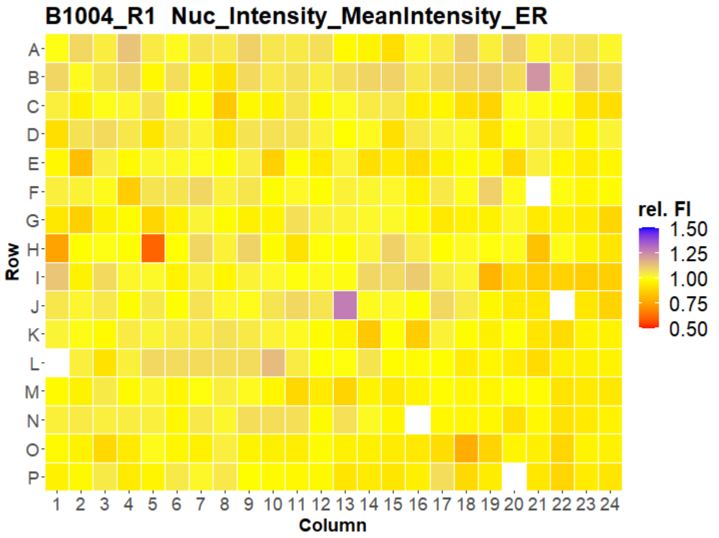


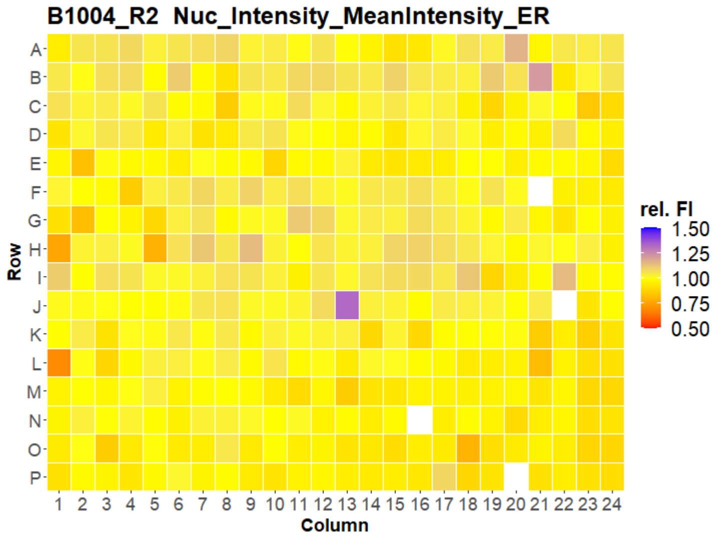


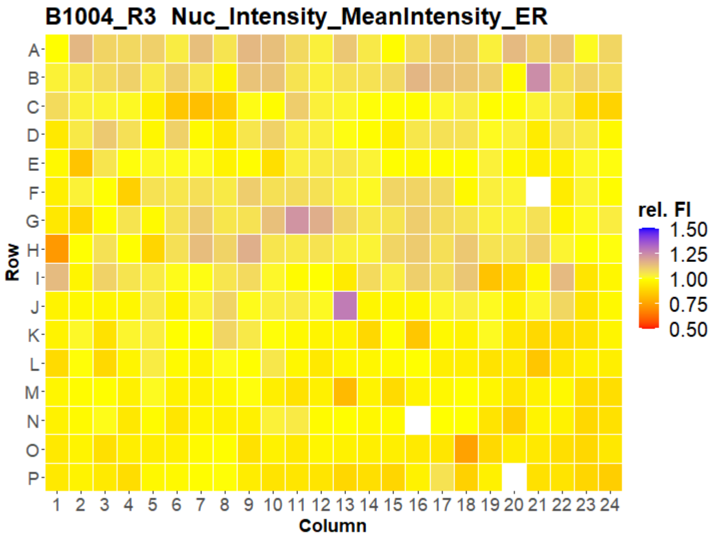


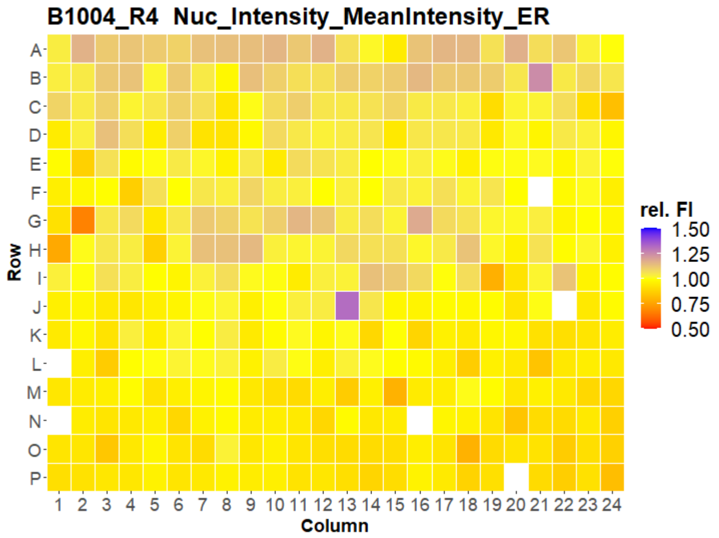


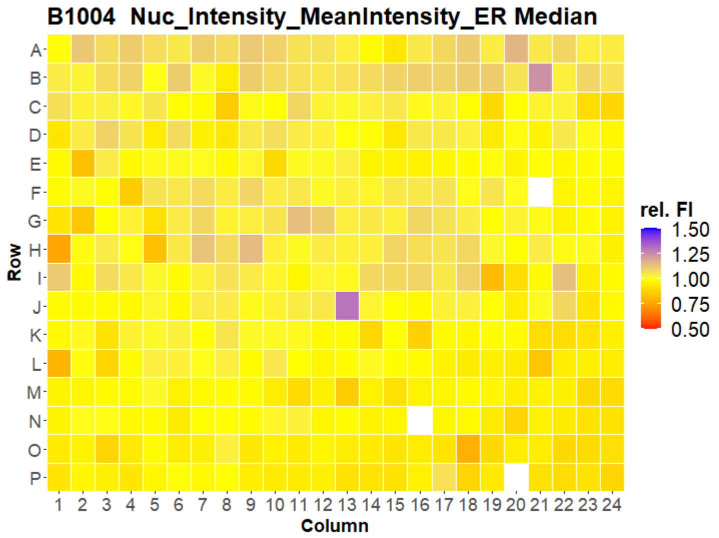

### HepG2 IMTM



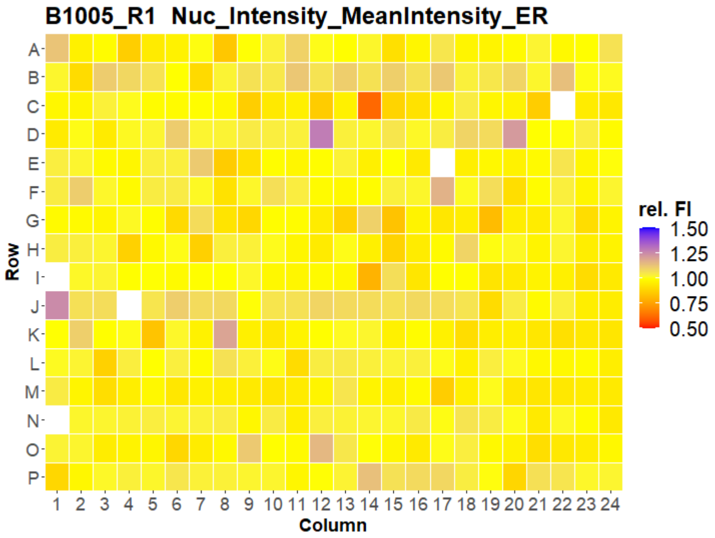


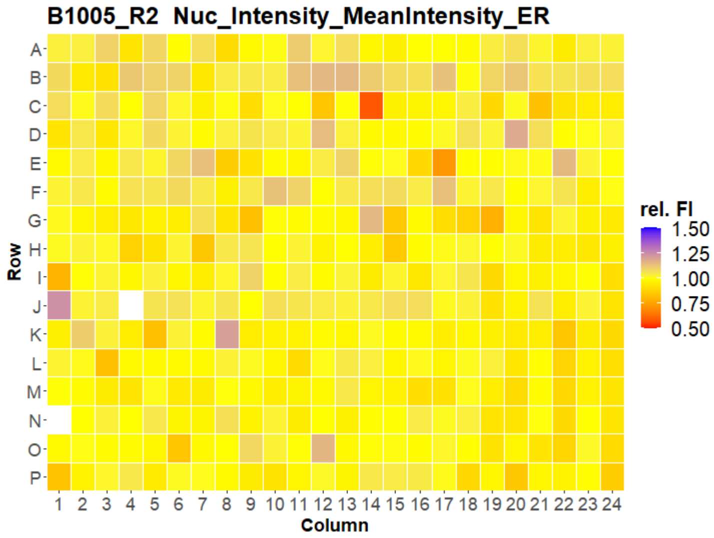


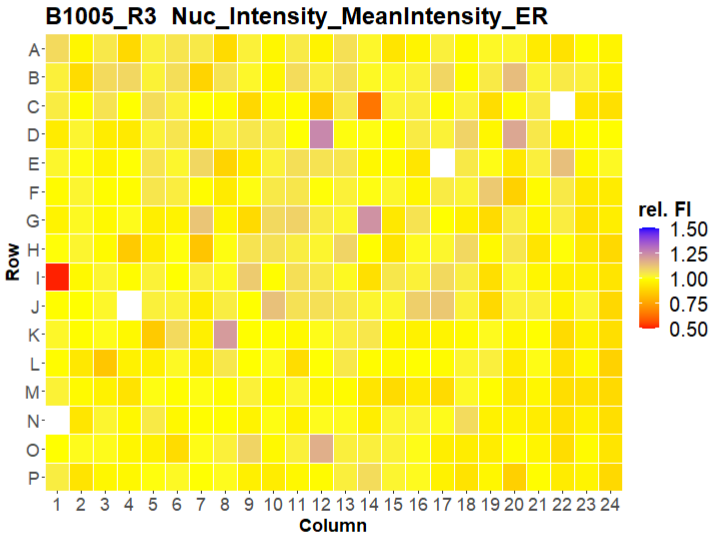


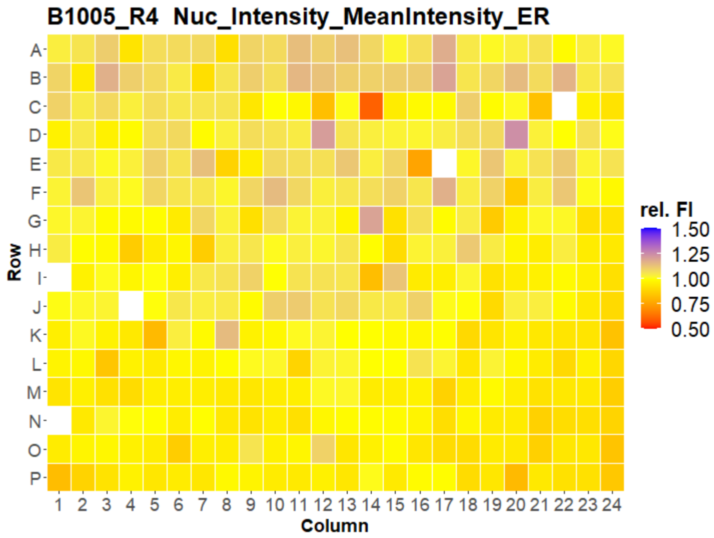


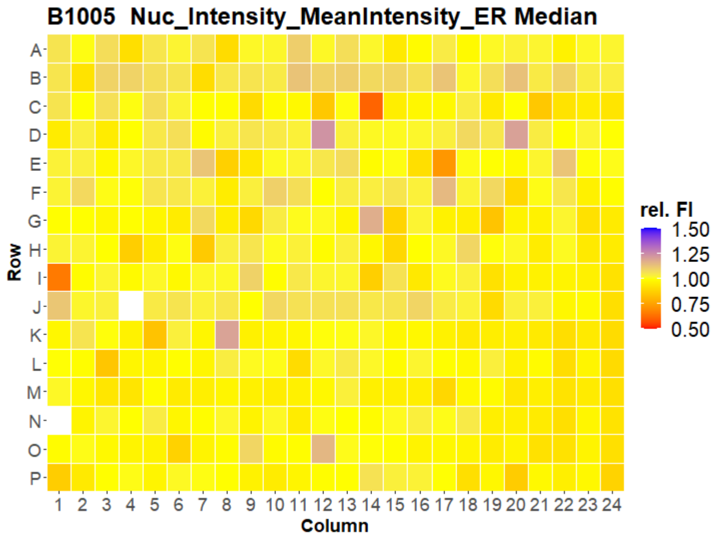



Open for Innovation KNIME

### HepG2 IMTM

41

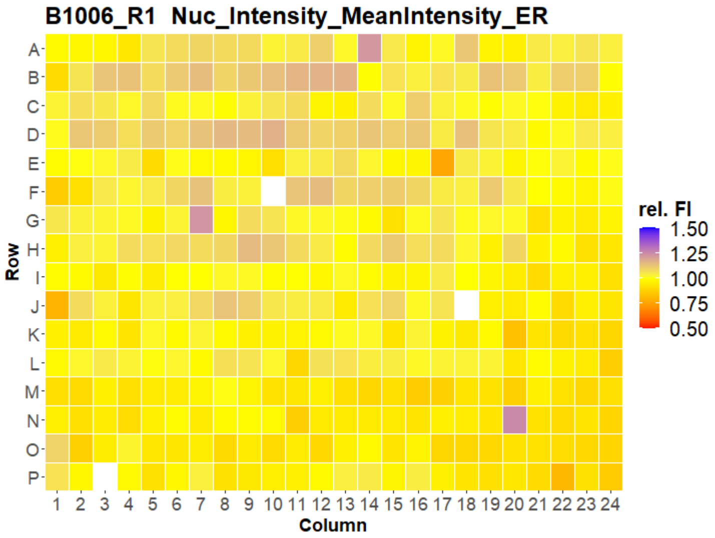


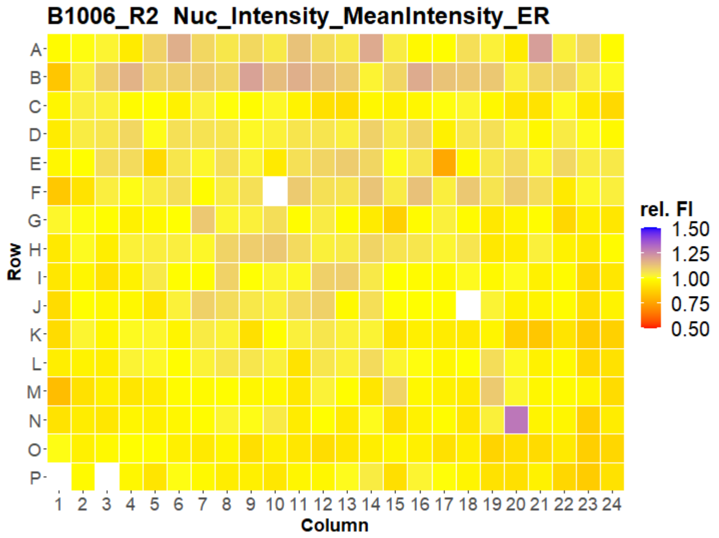


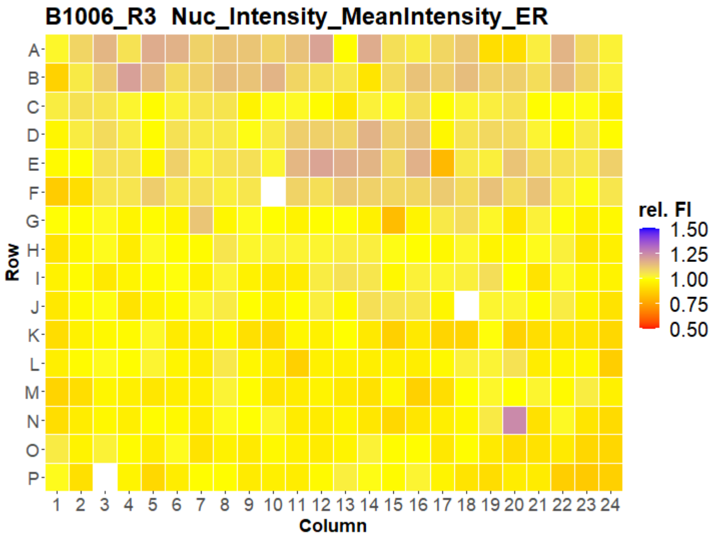


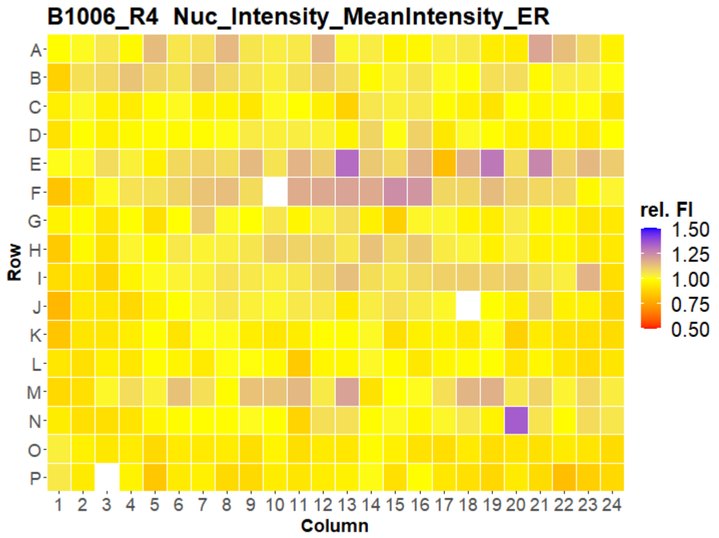


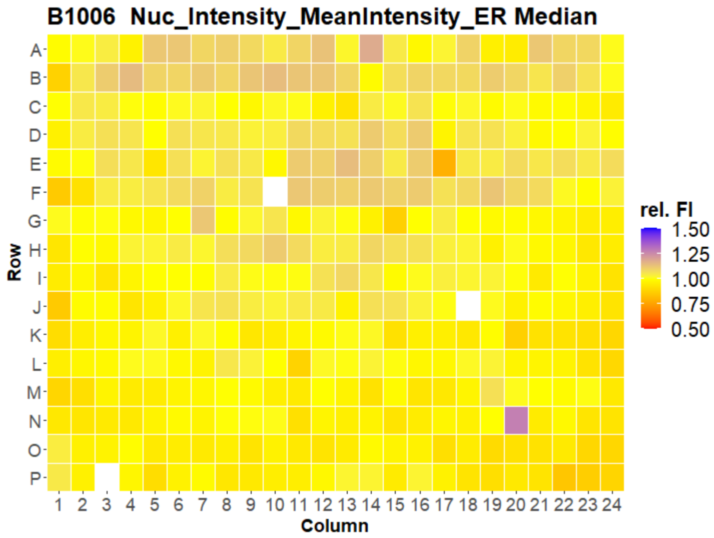

### HepG2 IMTM



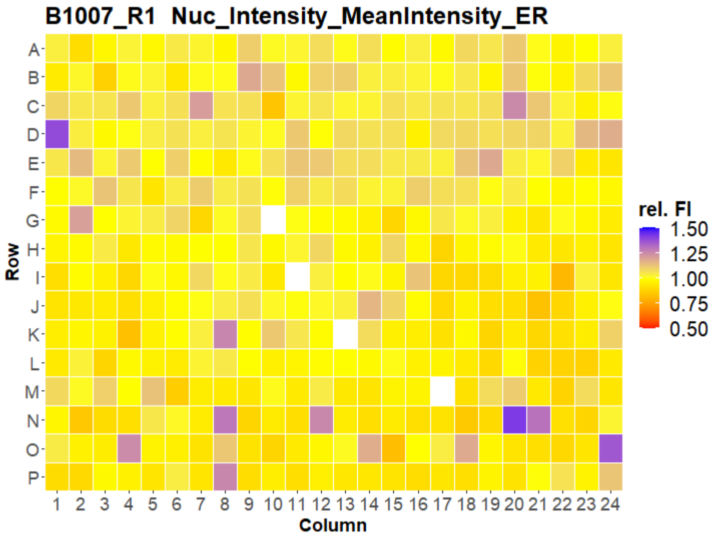
B1007_R1 Nuc_Intensity_MeanIntensity_ER
rel. FI
1.50
1.25
1.00
0.75
0.50
Row
A B C D E F G H I J K L M N O P
1 2 3 4 5 6 7 8 9 10 11 12 13 14 15 16 17 18 19 20 21 22 23 24
Column

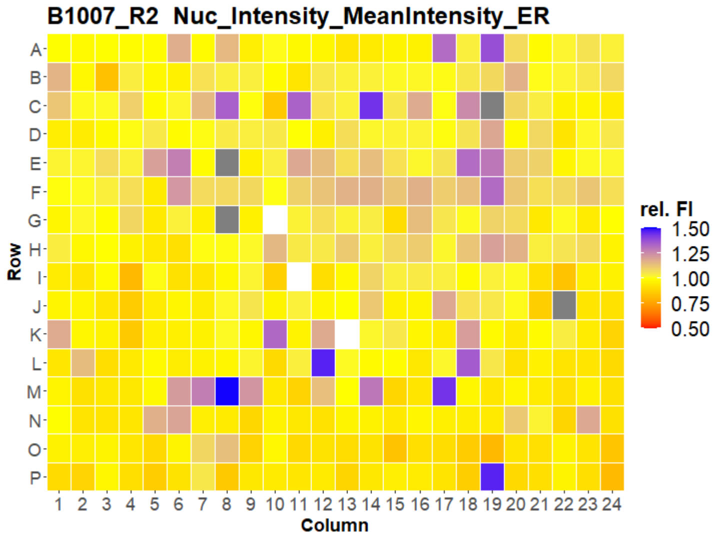
B1007_R2 Nuc_Intensity_MeanIntensity_ER
rel. FI
1.50
1.25
1.00
0.75
0.50
Row
A B C D E F G H I J K L M N O P
1 2 3 4 5 6 7 8 9 10 11 12 13 14 15 16 17 18 19 20 21 22 23 24
Column

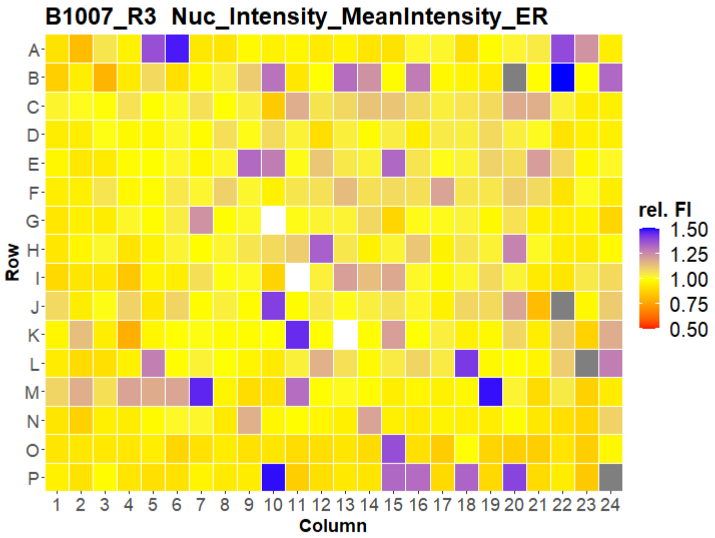
B1007_R3 Nuc_Intensity_MeanIntensity_ER
rel. FI
1.50
1.25
1.00
0.75
0.50
Row
A B C D E F G H I J K L M N O P
1 2 3 4 5 6 7 8 9 10 11 12 13 14 15 16 17 18 19 20 21 22 23 24
Column

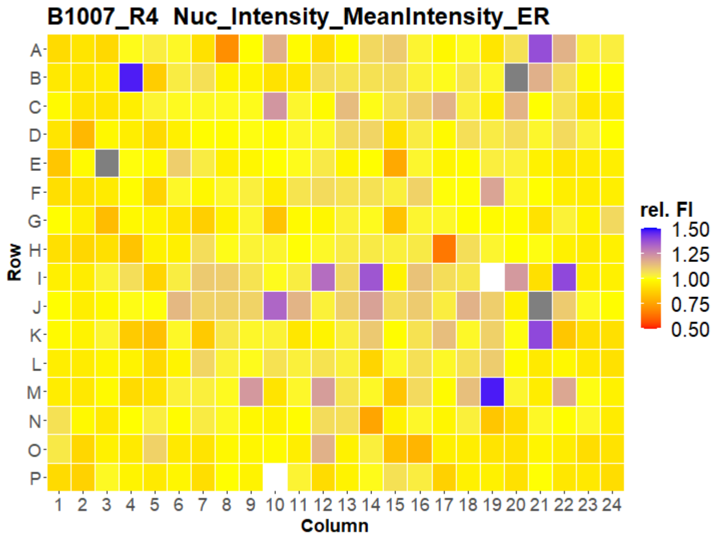
B1007_R4 Nuc_Intensity_MeanIntensity_ER
rel. FI
1.50
1.25
1.00
0.75
0.50
Row
A B C D E F G H I J K L M N O P
1 2 3 4 5 6 7 8 9 10 11 12 13 14 15 16 17 18 19 20 21 22 23 24
Column

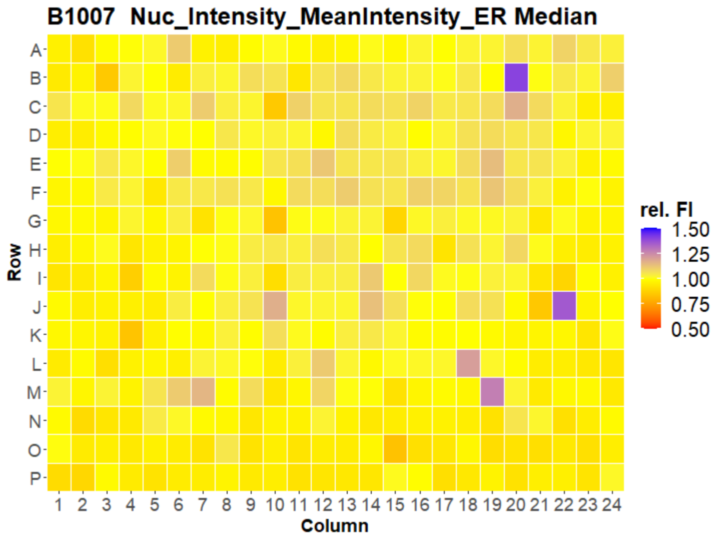
B1007 Nuc_Intensity_MeanIntensity_ER Median
rel. FI
1.50
1.25
1.00
0.75
0.50
Row
A B C D E F G H I J K L M N O P
1 2 3 4 5 6 7 8 9 10 11 12 13 14 15 16 17 18 19 20 21 22 23 24
Column

### HepG2 IMTM
