## Supplementary material for "Morphological Profiling Dataset of EU-OPENSCREEN Bioactive Compounds Over Multiple Imaging Sites and Cell Lines": Suppl_QC_MEDI_HepG2.pdf

### HepG2 Medina

#### B1001 Metadata_Object_Count

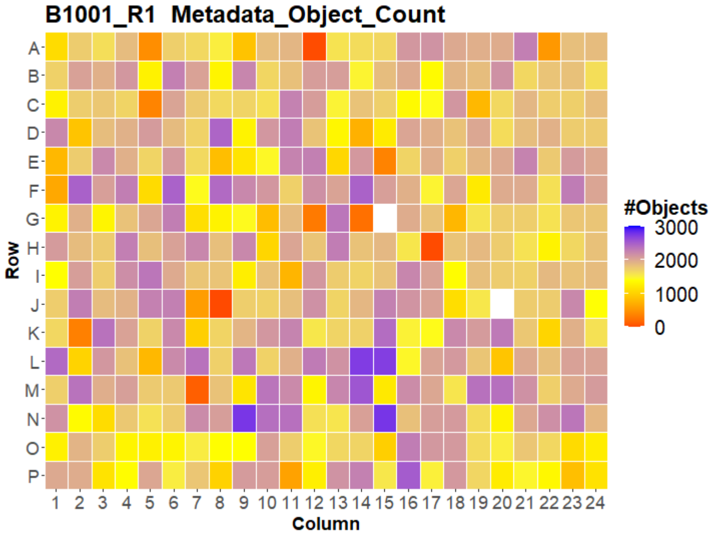


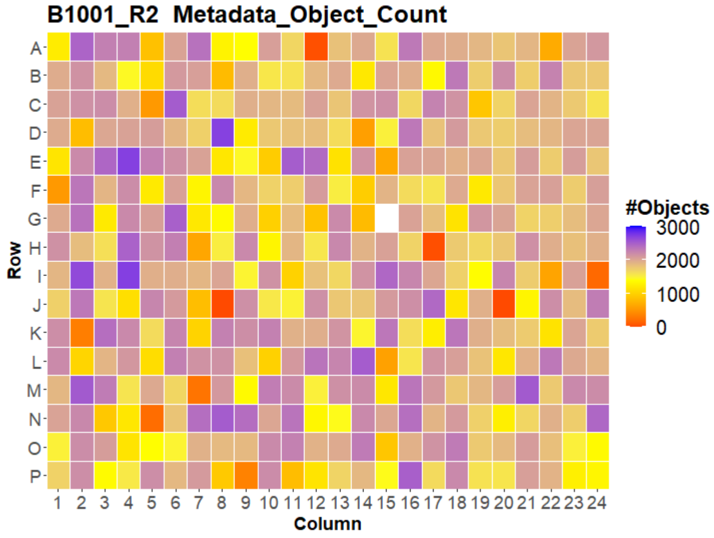


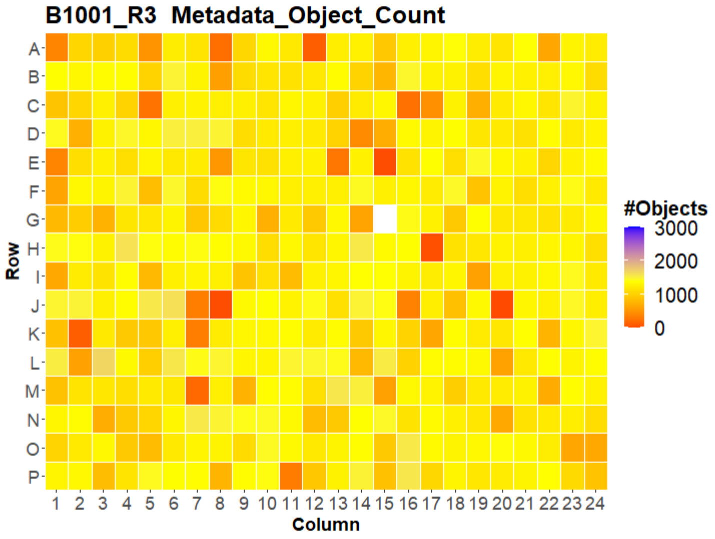


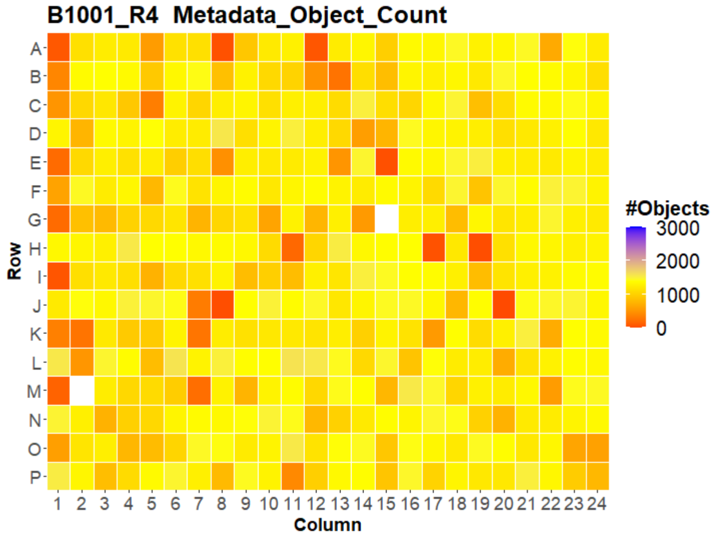


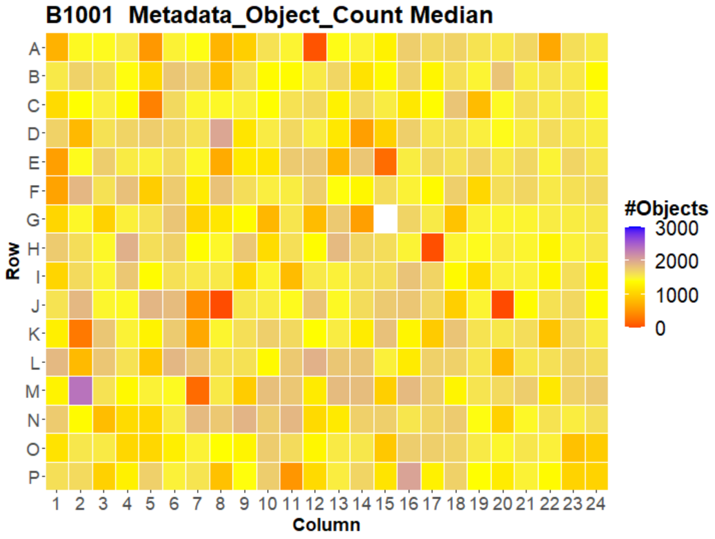

### HepG2 Medina

#### B1002 Metadata_Object_Count

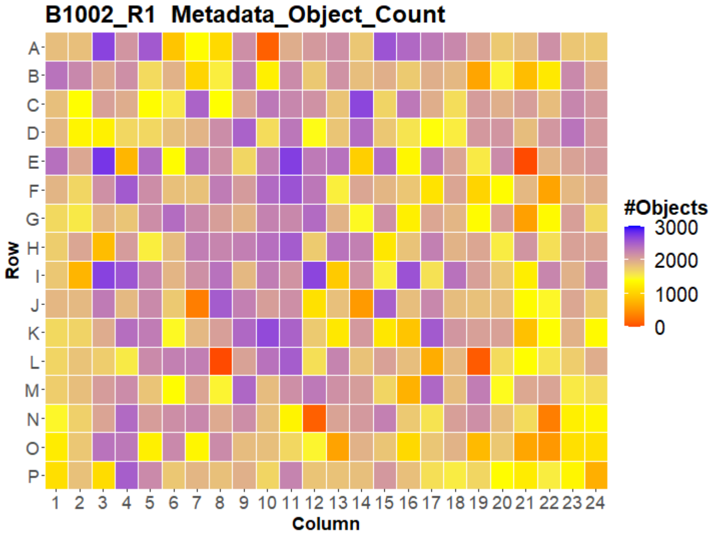


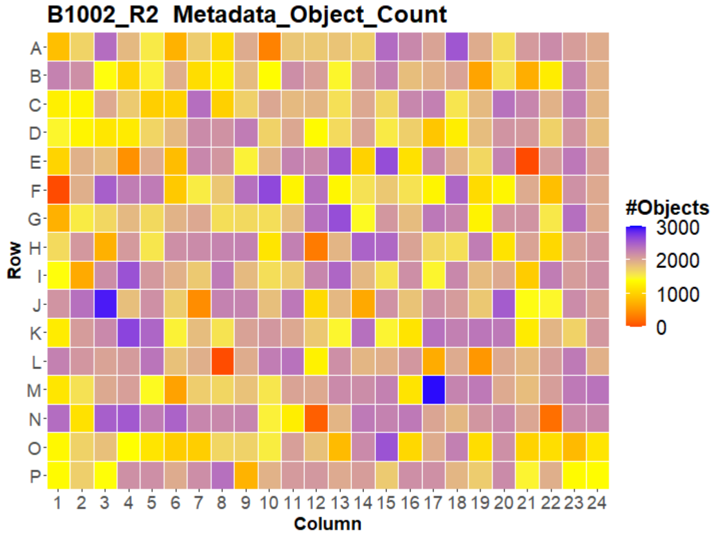


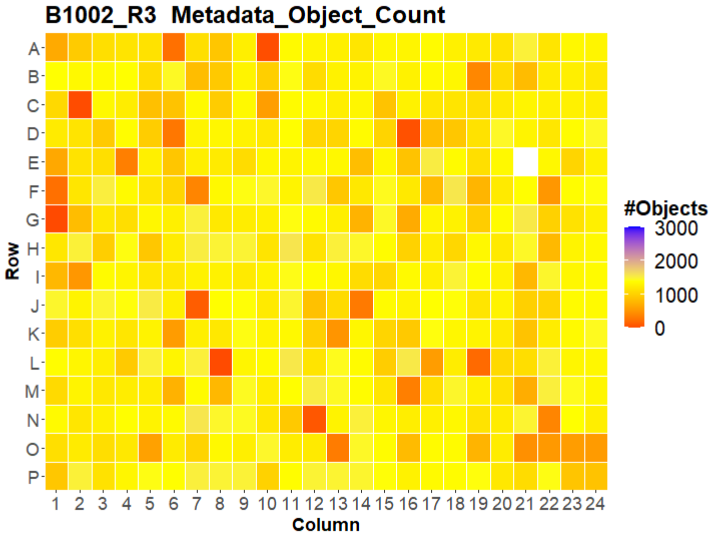


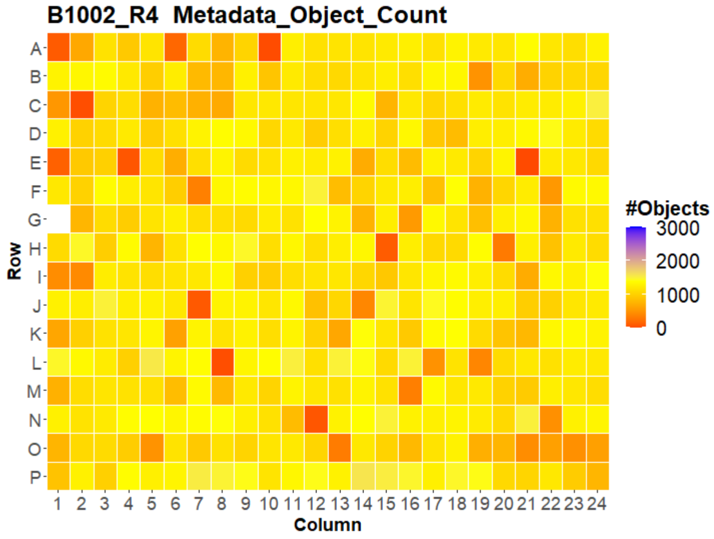


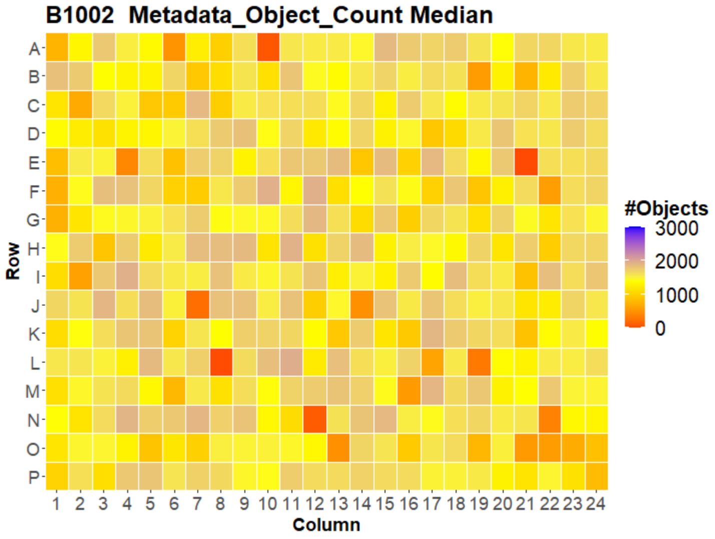

### HepG2 Medina

#### B1003 Metadata_Object_Count

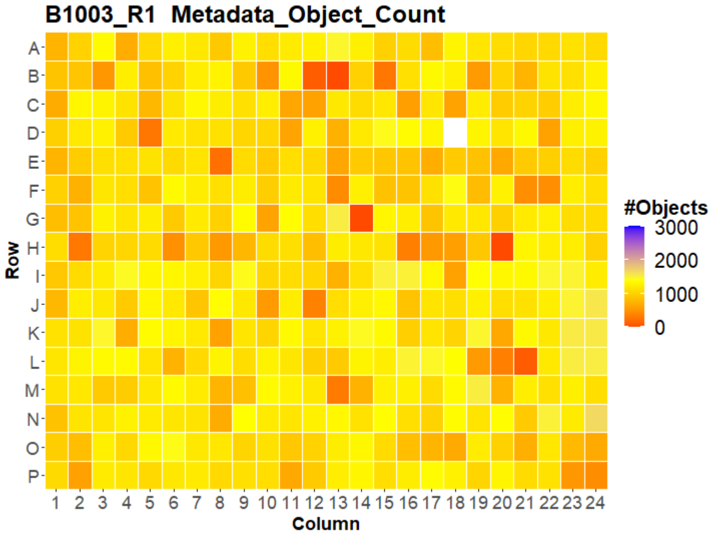


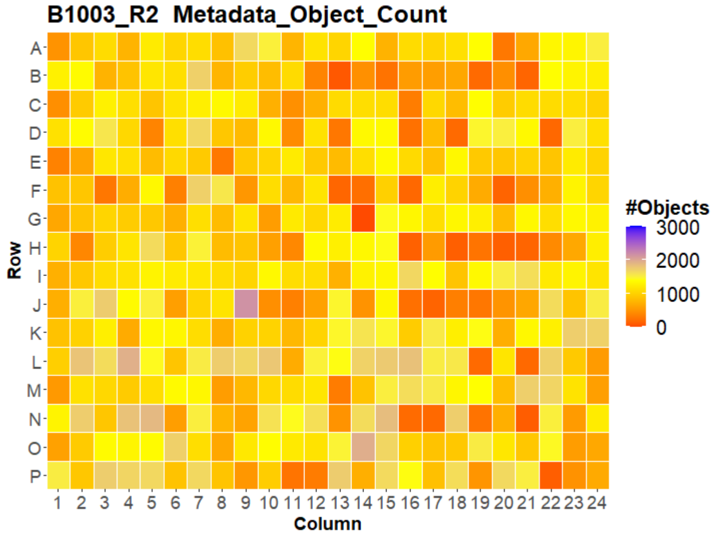


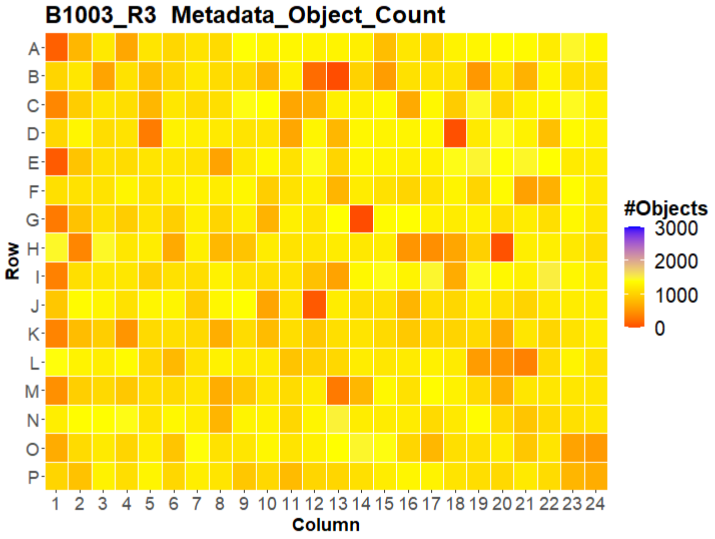


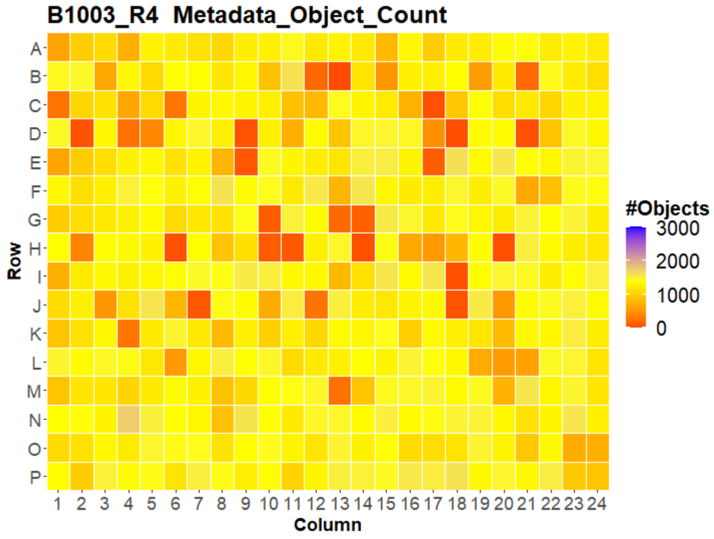


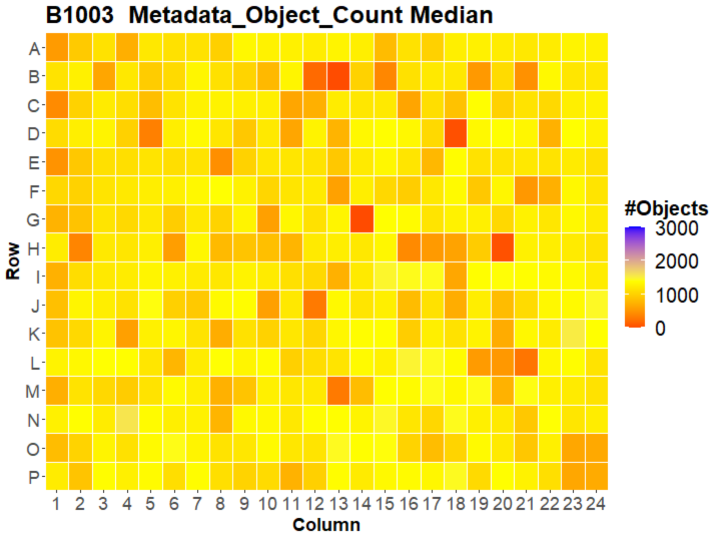

### HepG2 Medina

#### B1004 Metadata_Object_Count

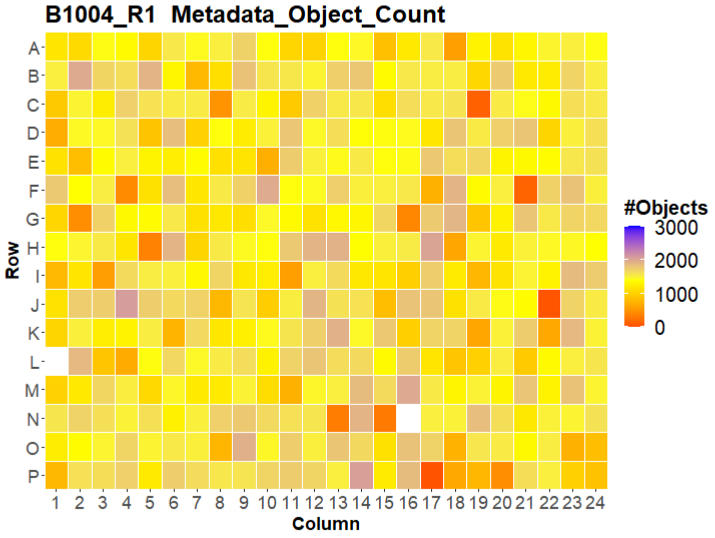


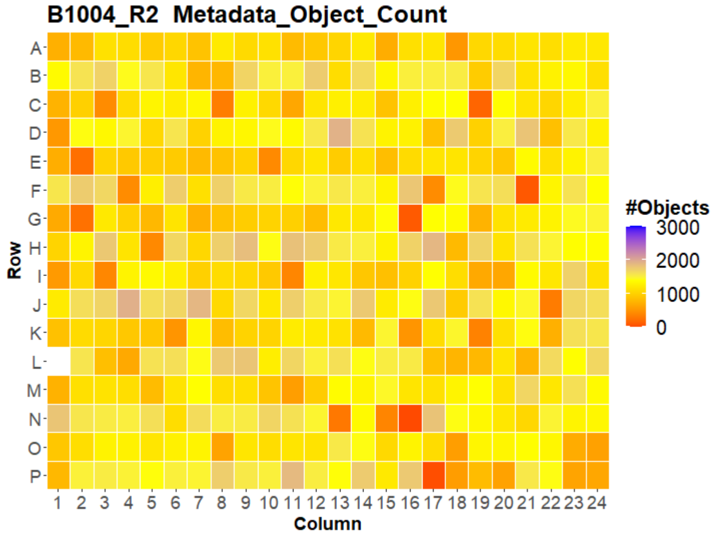


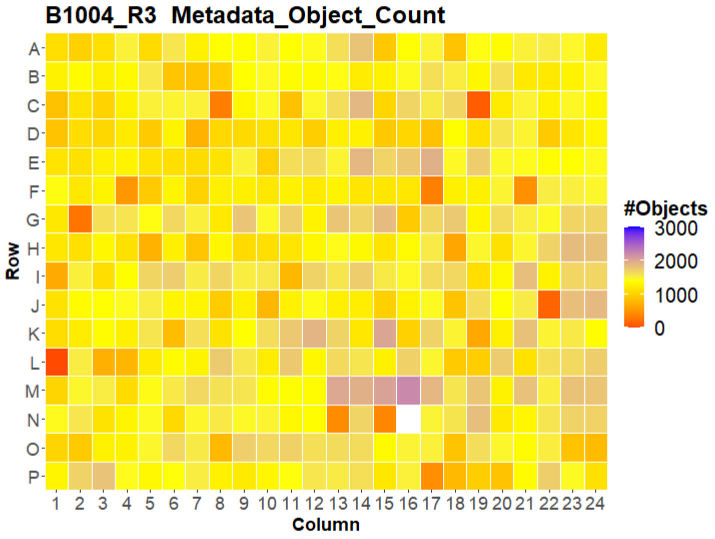


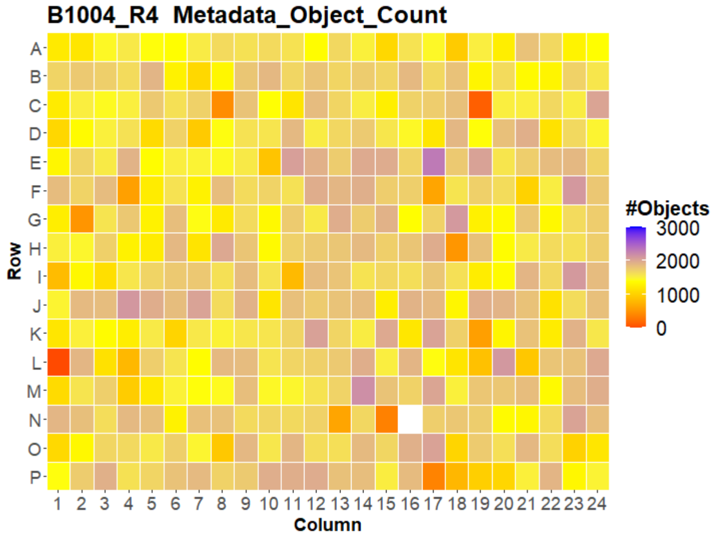


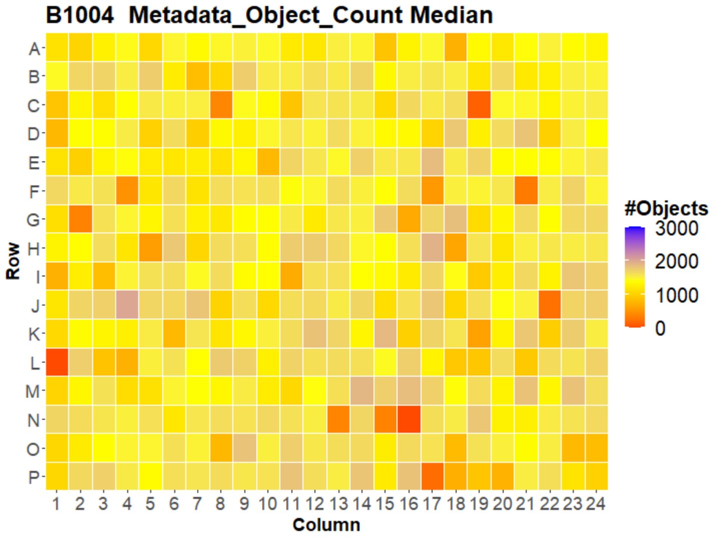

### HepG2 Medina

#### B1005 Metadata_Object_Count

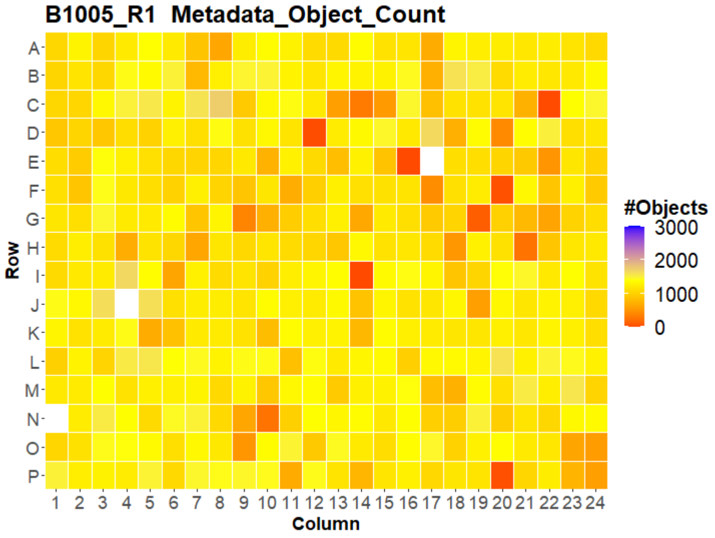


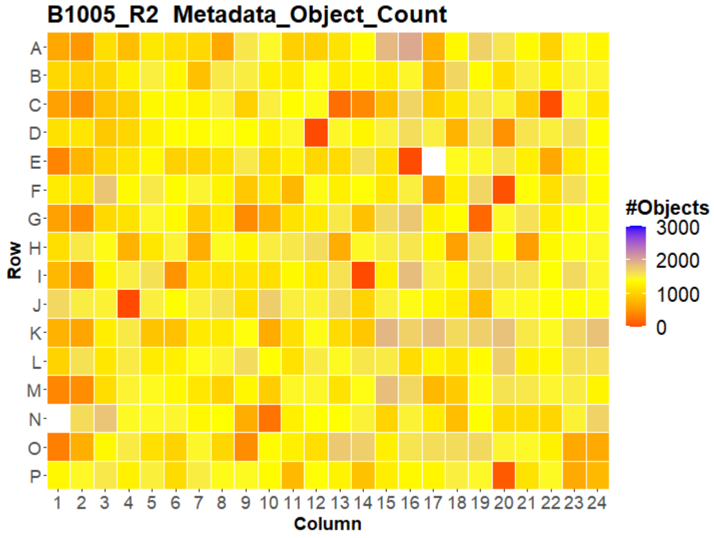


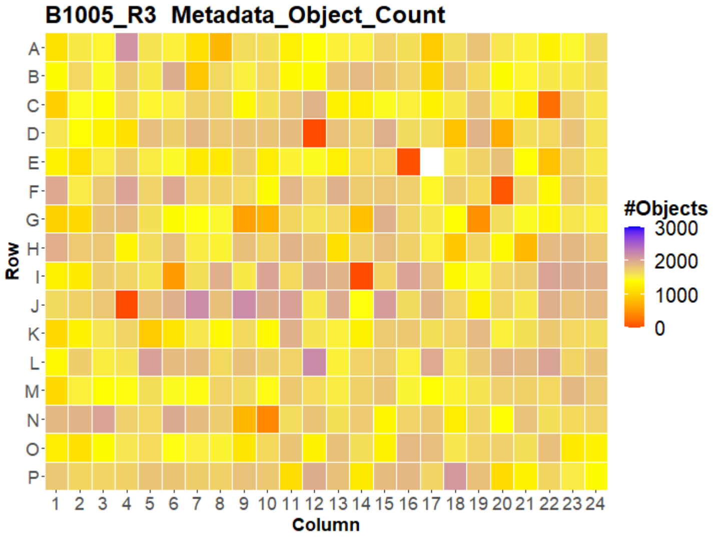


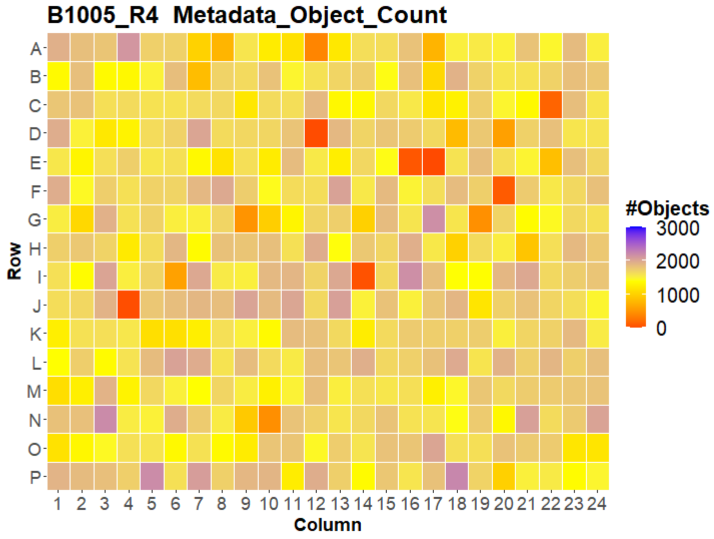


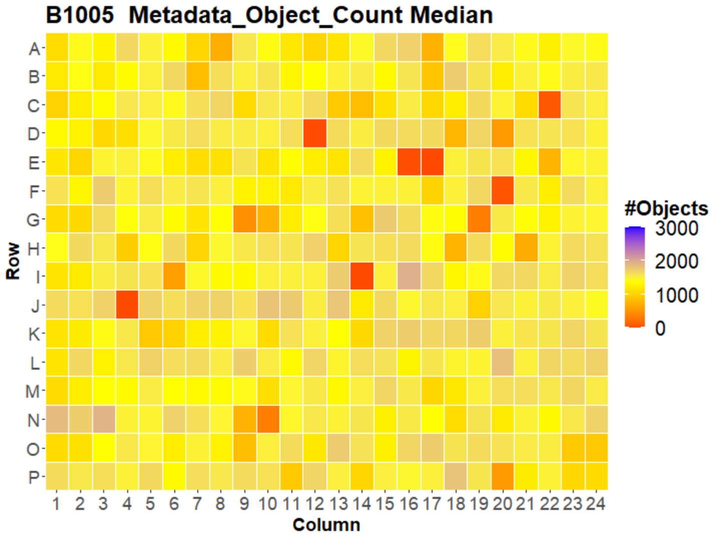


LEIBNIZ FORSCHUNGSINSTITUT FÜR MOLEKULARE PHARMAKOLOGIE

Open for Innovation KNIME

### HepG2 Medina

#### B1006 Metadata_Object_Count

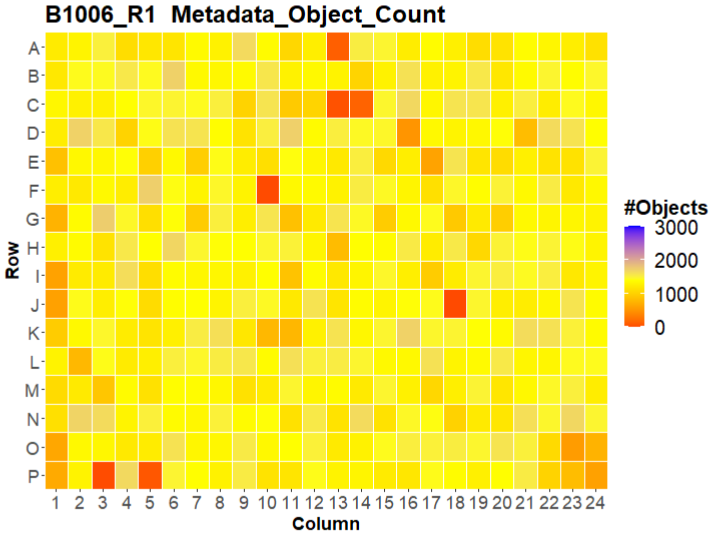


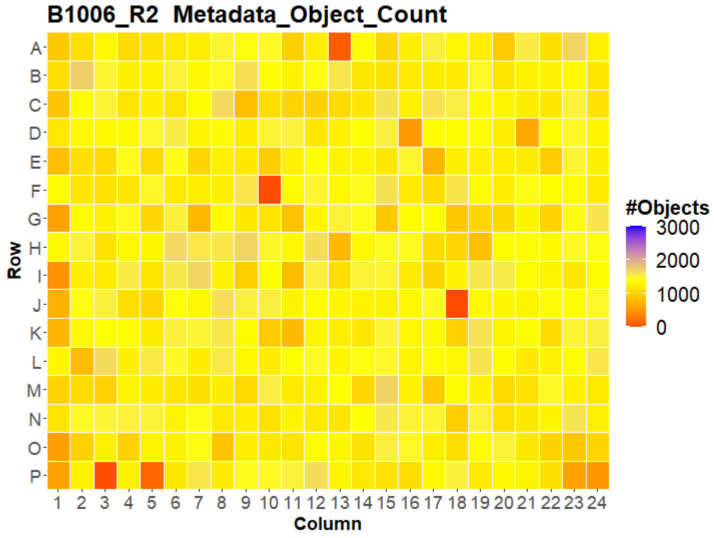


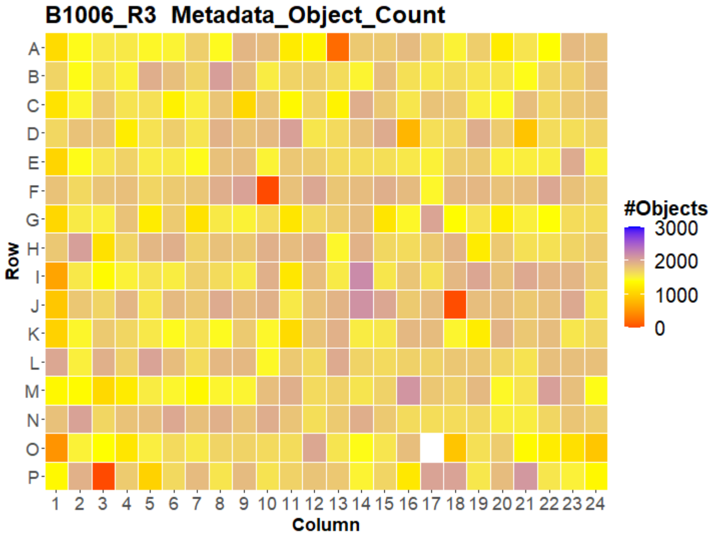


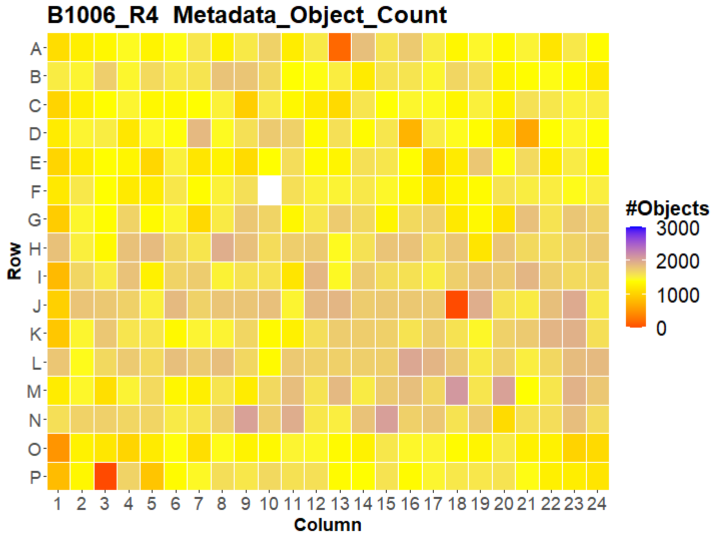


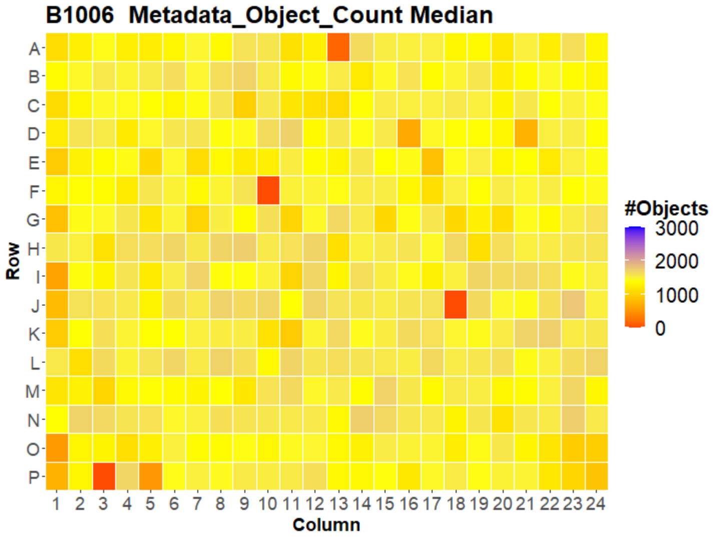

### HepG2 Medina

#### B1007 Metadata_Object_Count

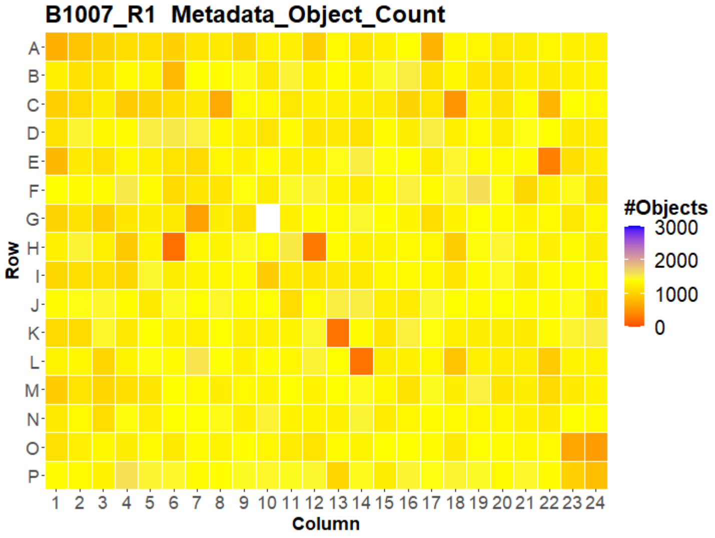


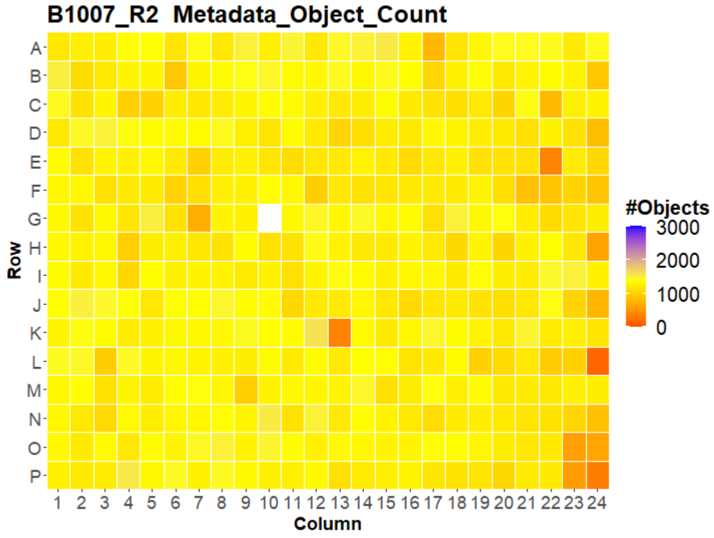


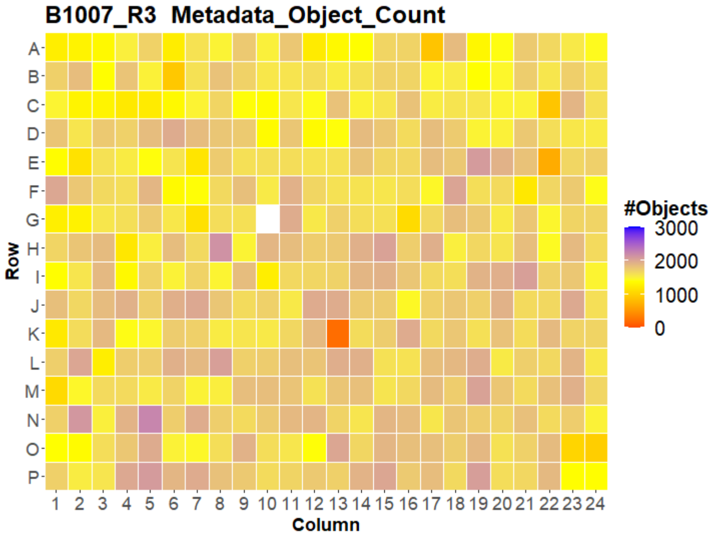


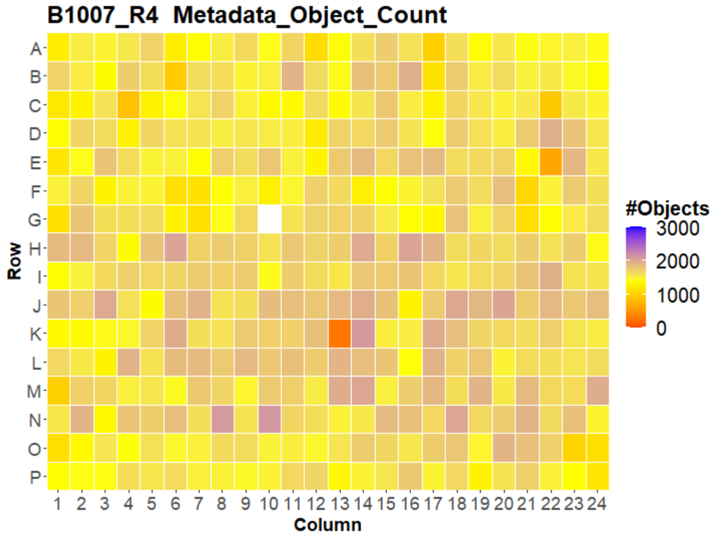


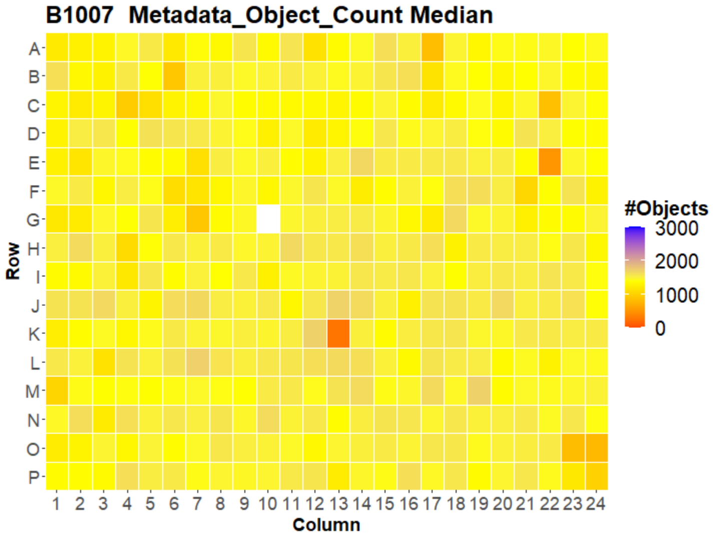

### HepG2 Medina

#### B1001 Cyto_Intensity_MeanIntensity_AGP

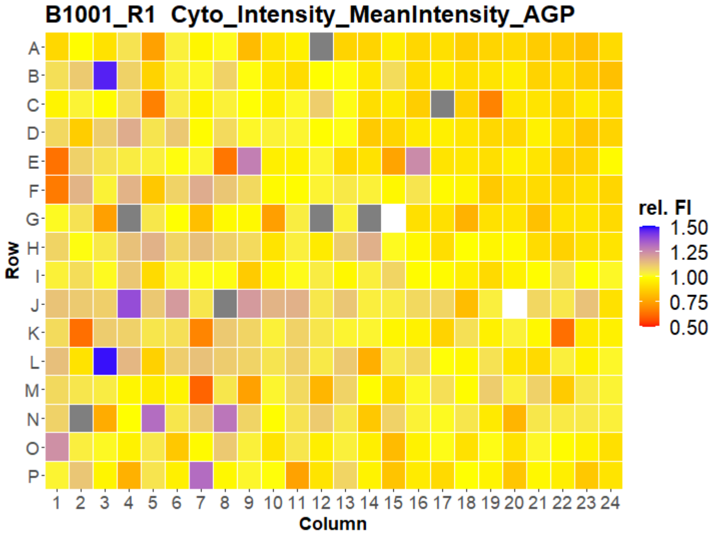


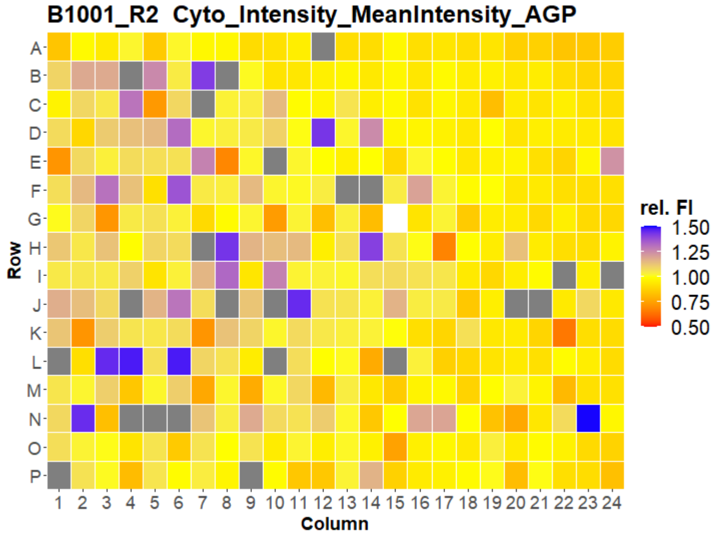


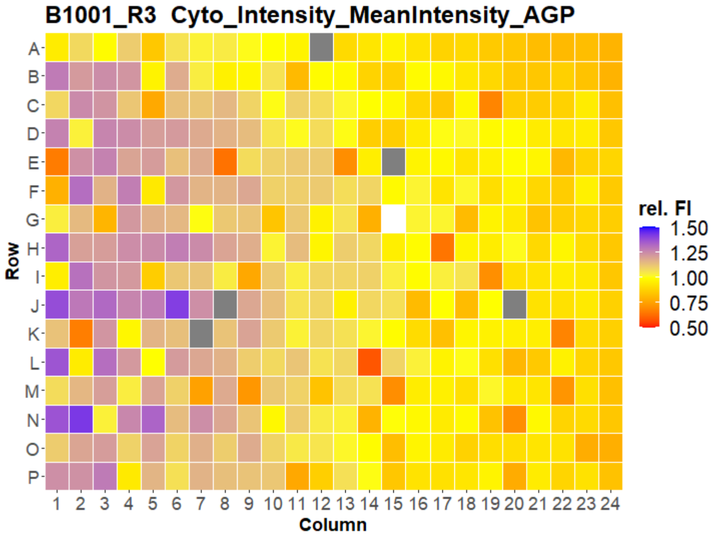


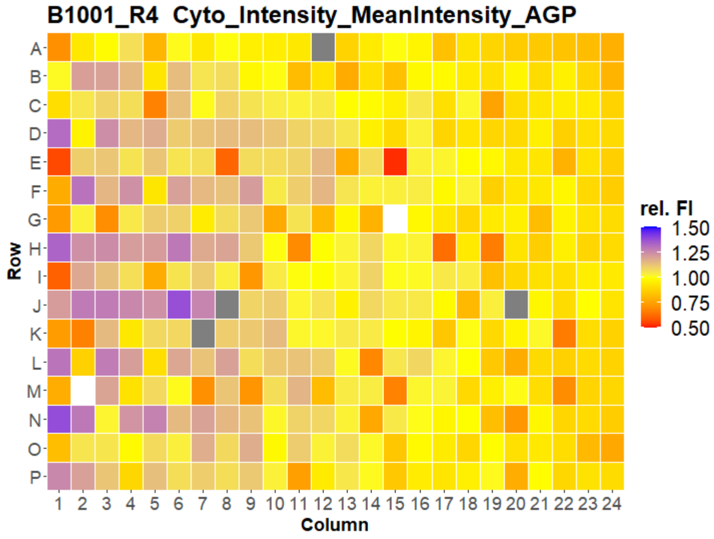


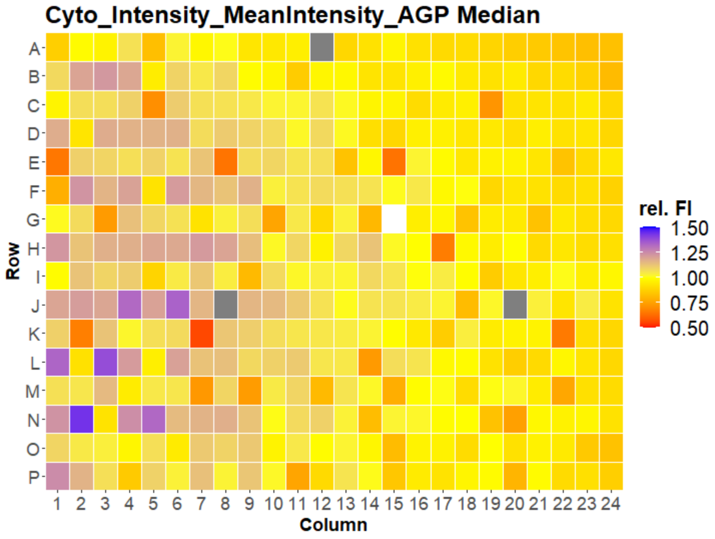

### HepG2 Medina

#### B1002 Cyto_Intensity_MeanIntensity_AGP

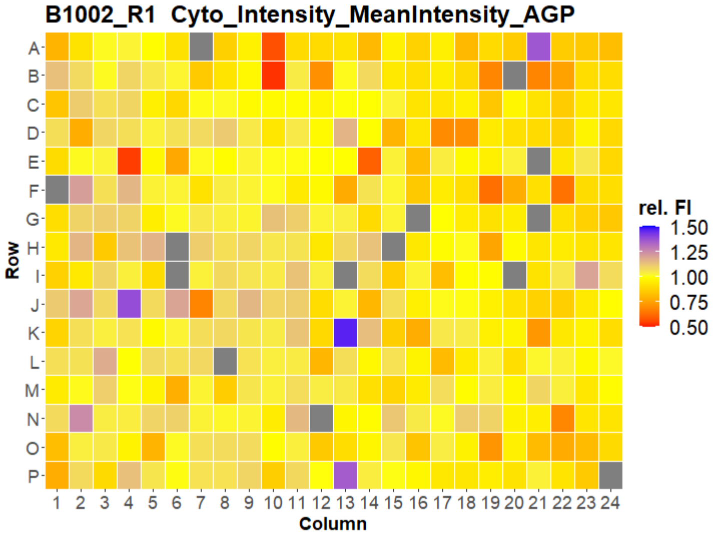


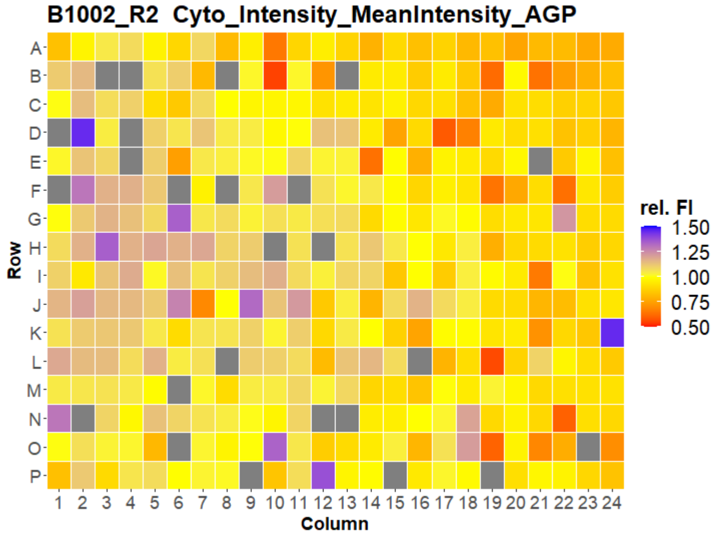


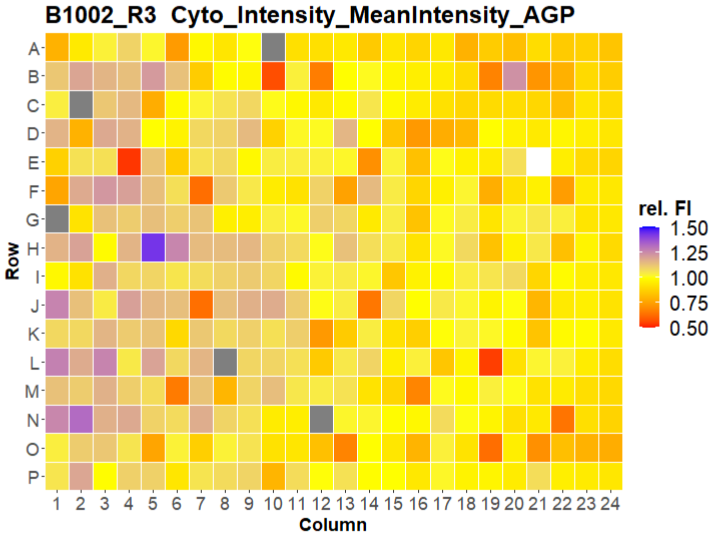


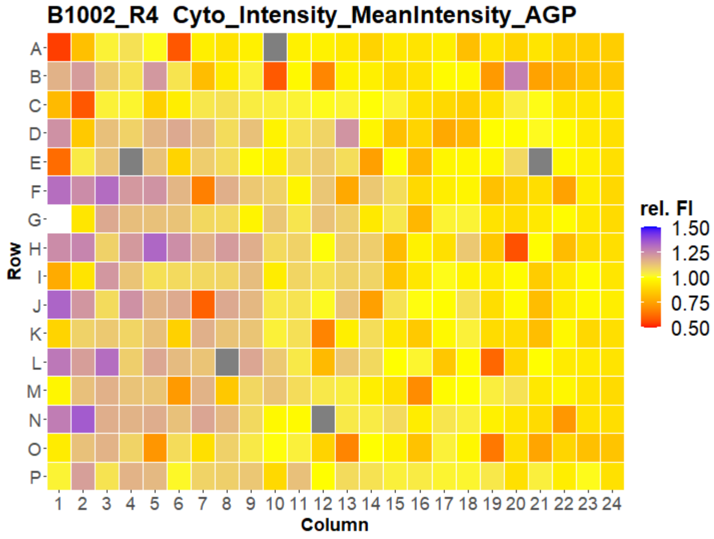


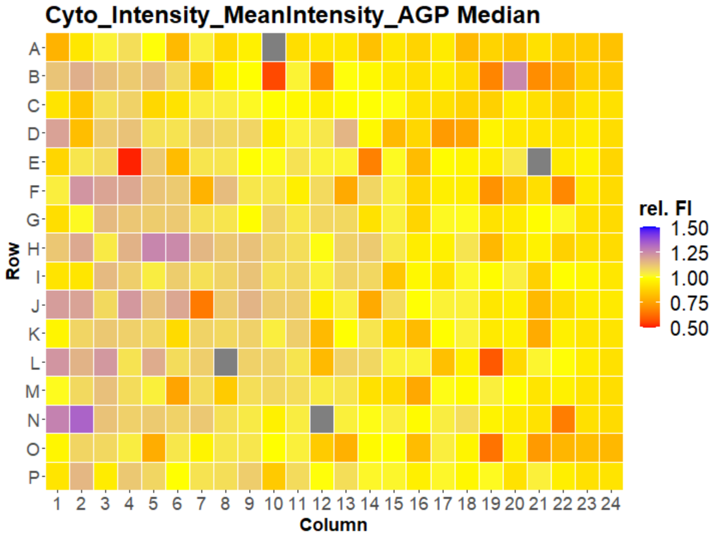

### HepG2 Medina

#### B1003 Cyto_Intensity_MeanIntensity_AGP

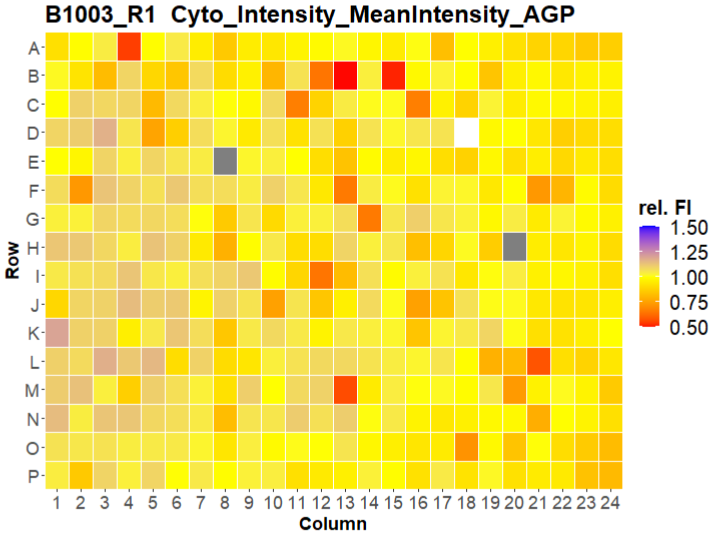


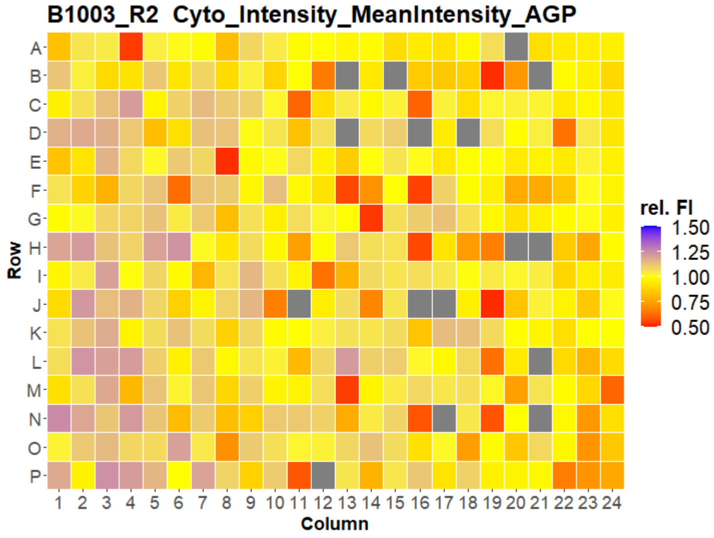


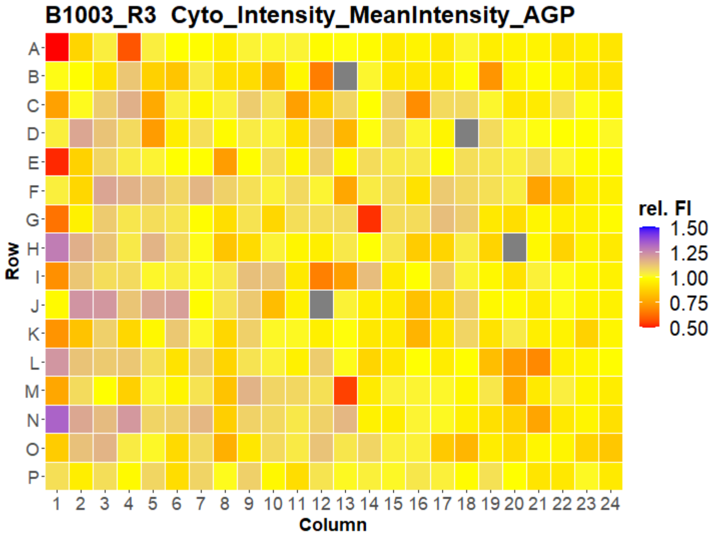


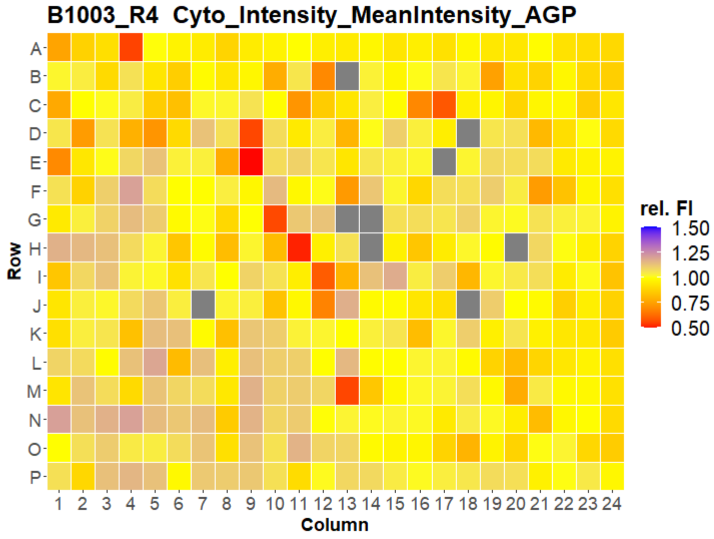


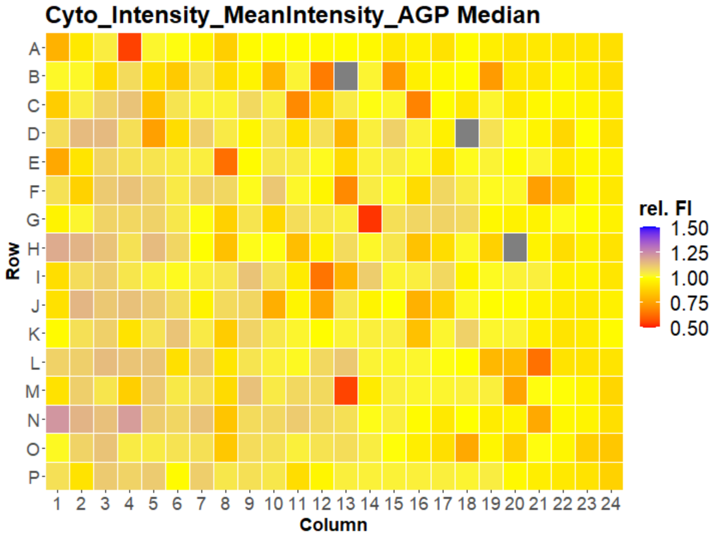

### HepG2 Medina

#### B1004 Cyto_Intensity_MeanIntensity_AGP

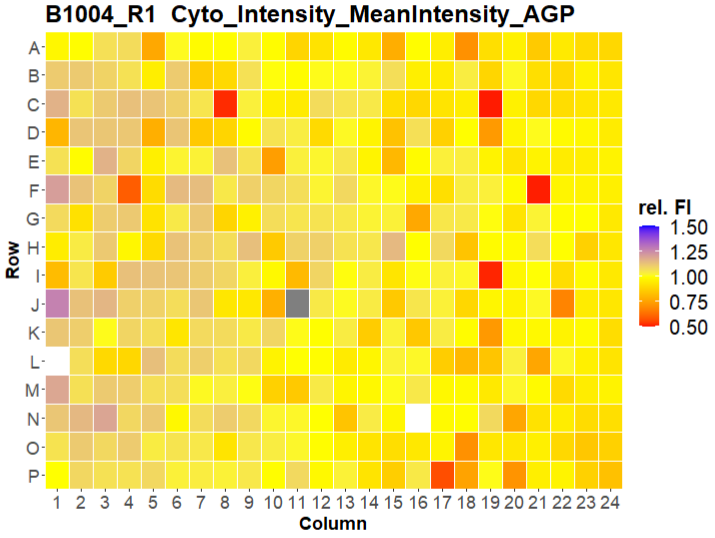


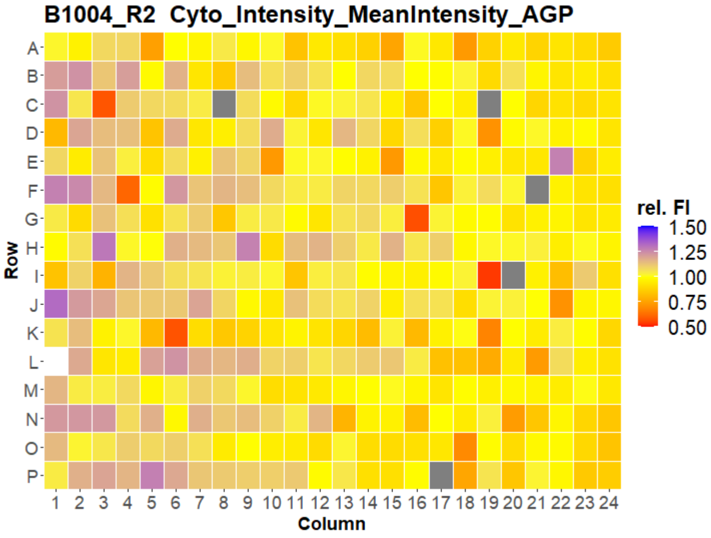


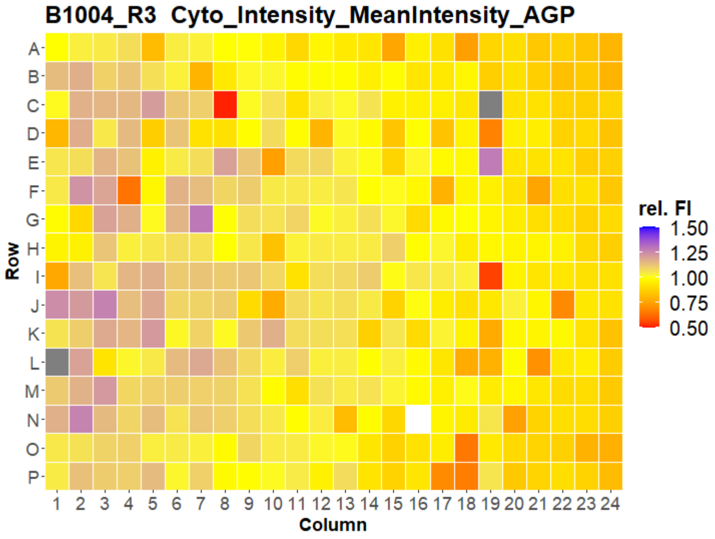


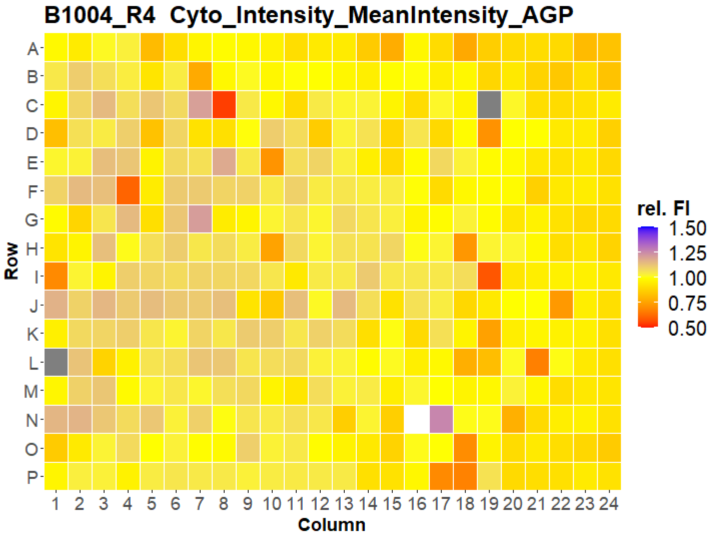


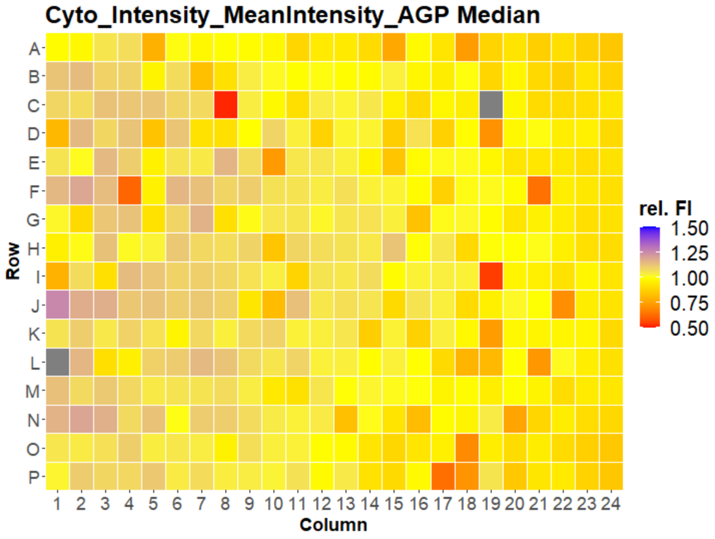

### HepG2 Medina

#### B1005 Cyto_Intensity_MeanIntensity_AGP

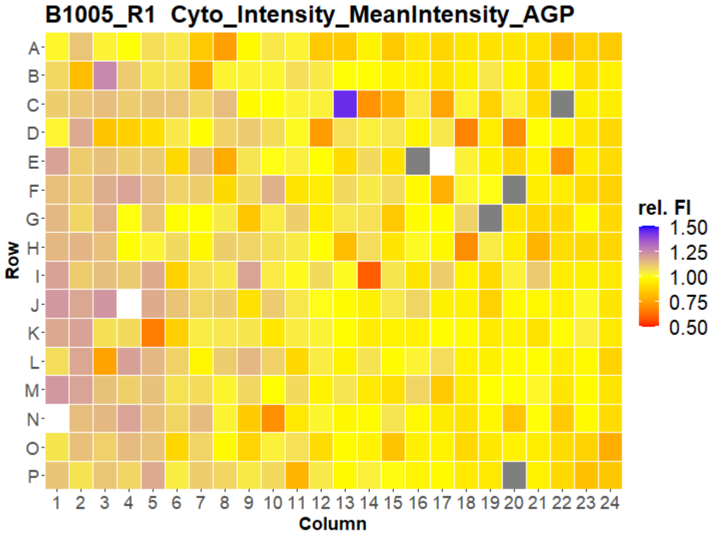


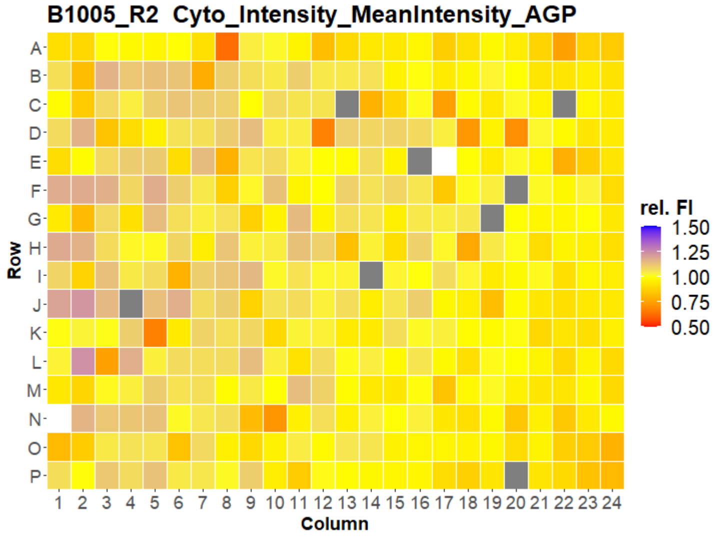


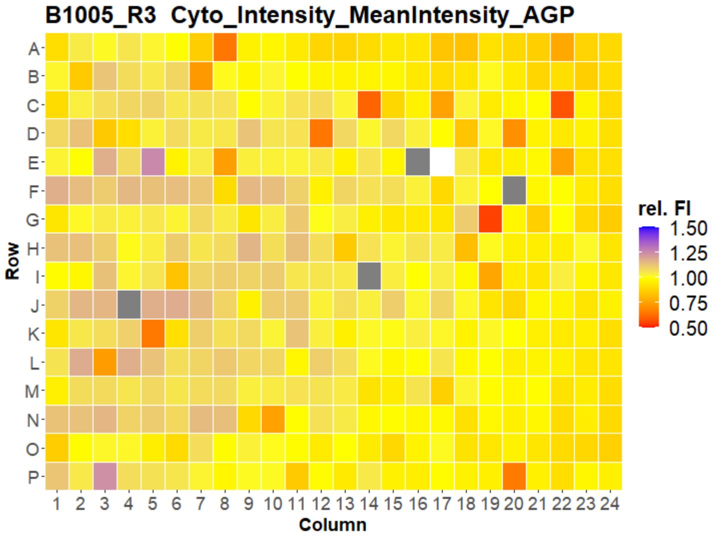


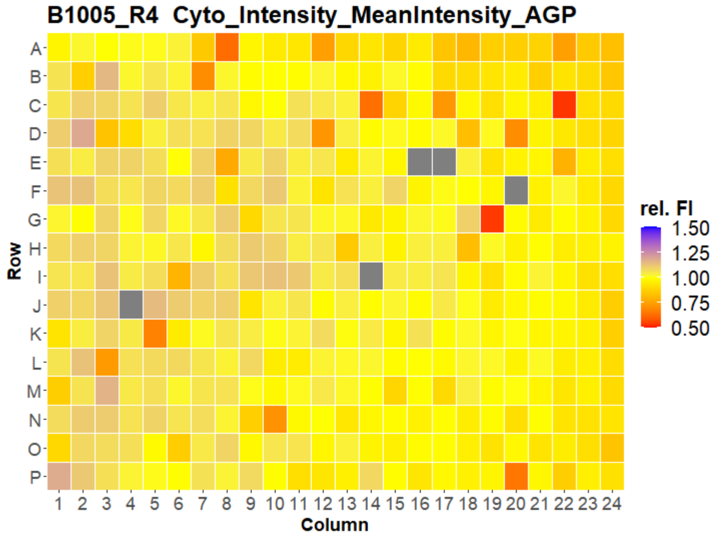


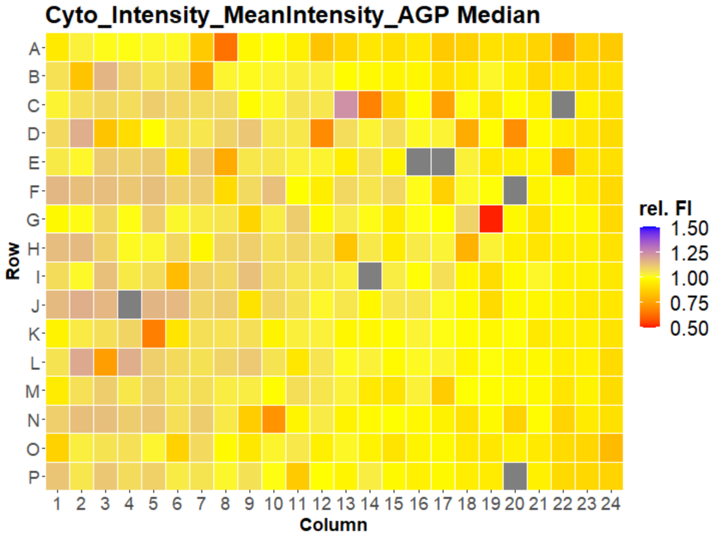

### HepG2 Medina

#### B1006 Cyto_Intensity_MeanIntensity_AGP

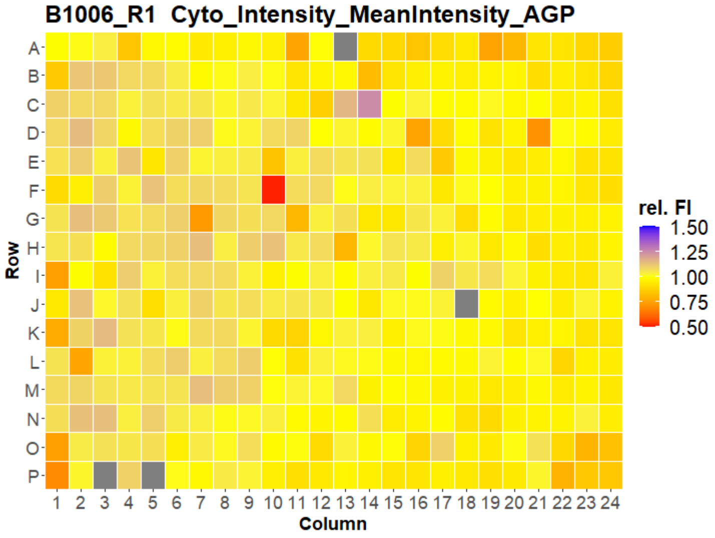


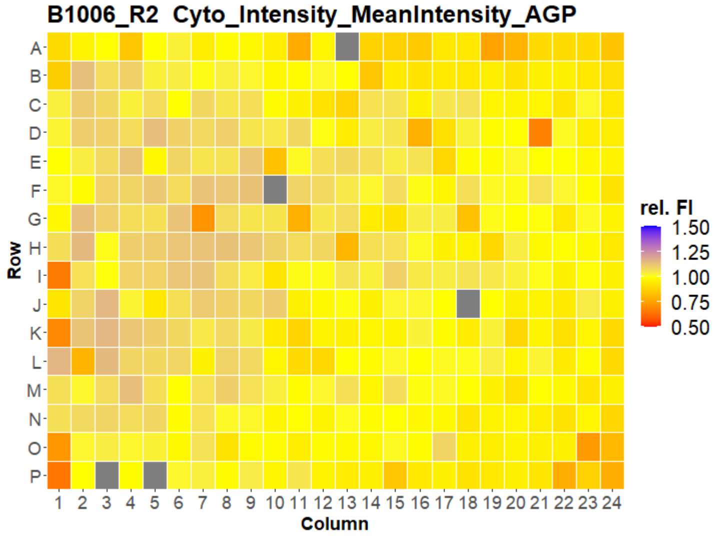


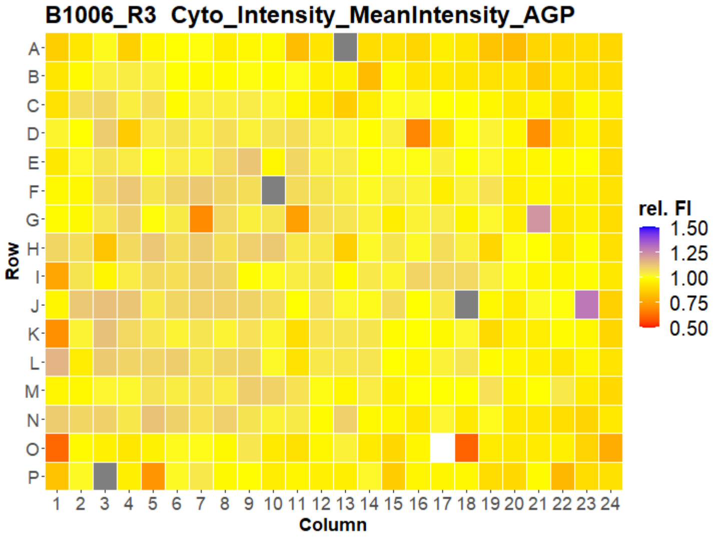


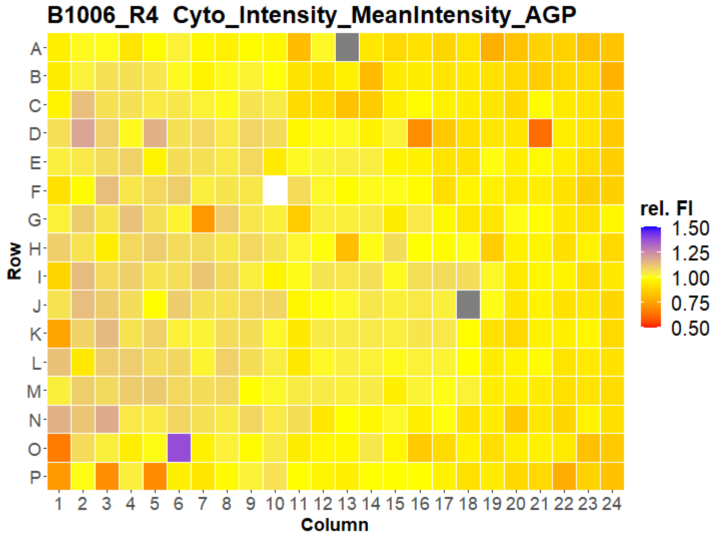


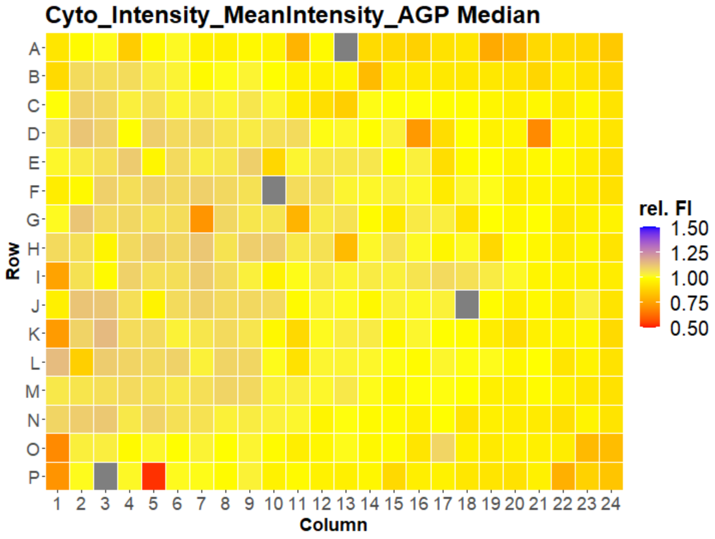

### HepG2 Medina

#### B1007 Cyto_Intensity_MeanIntensity_AGP

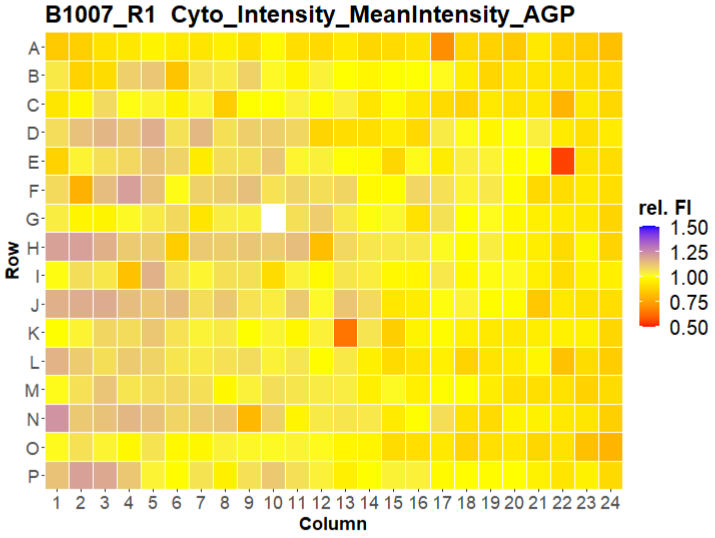


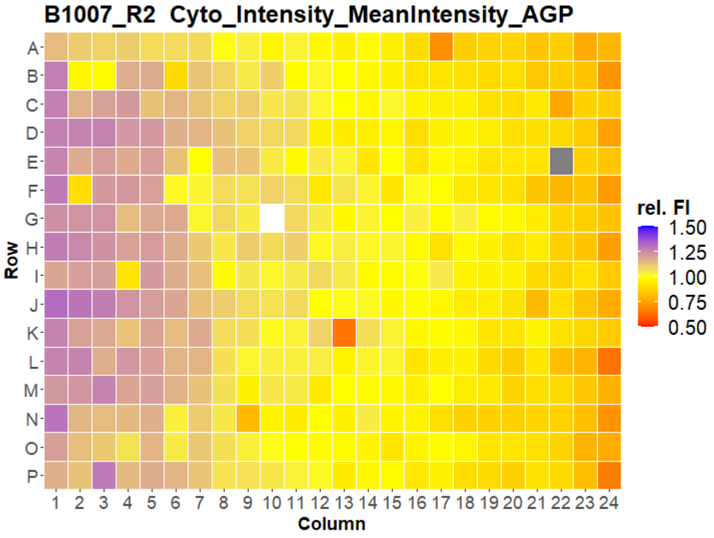


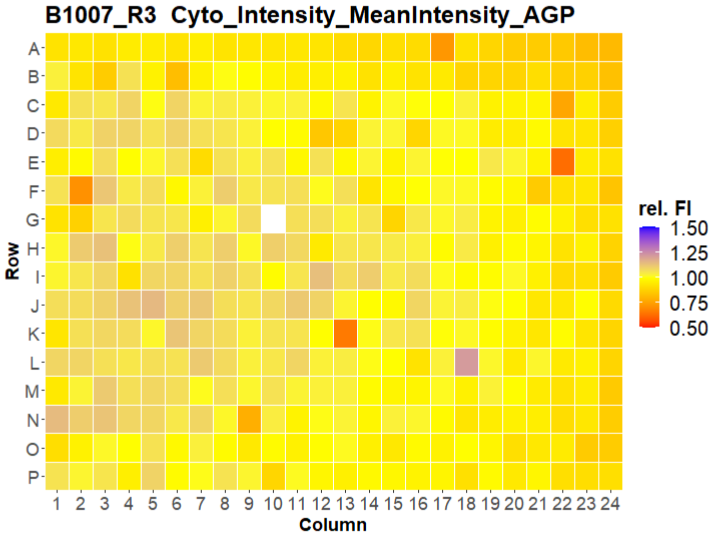


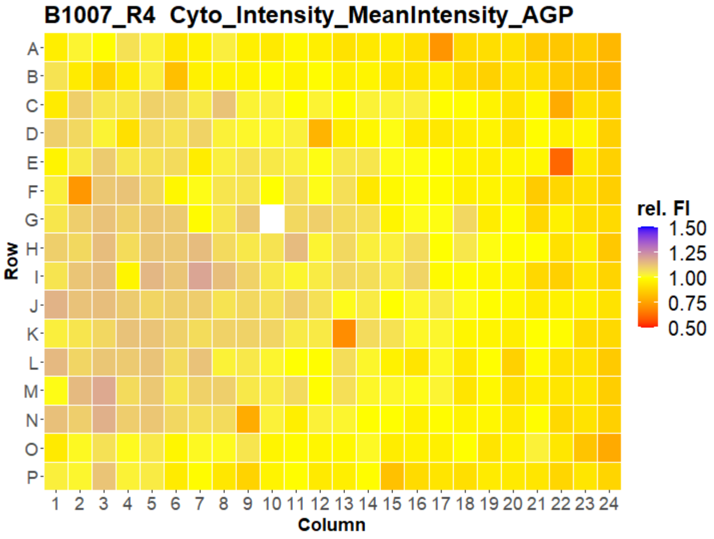


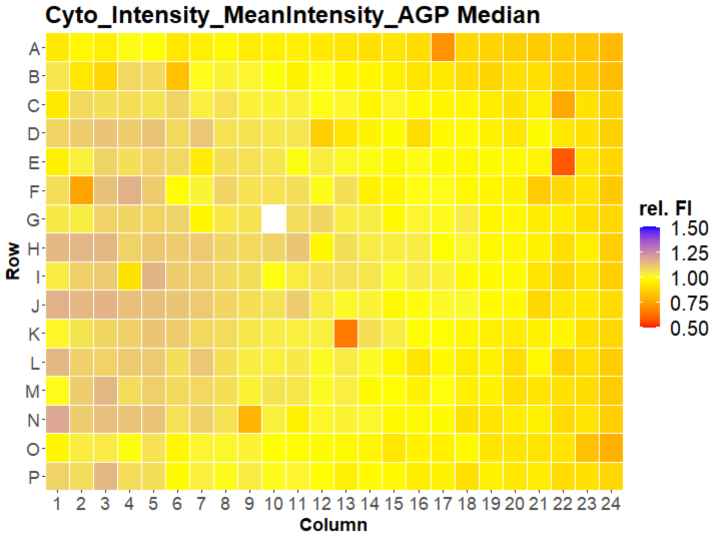

### HepG2 Medina

#### B1001 Cyto_Intensity_MeanIntensity_ER

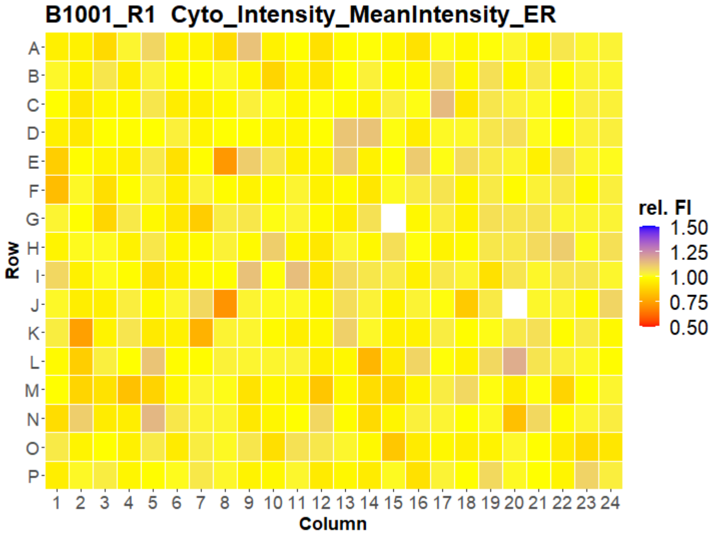


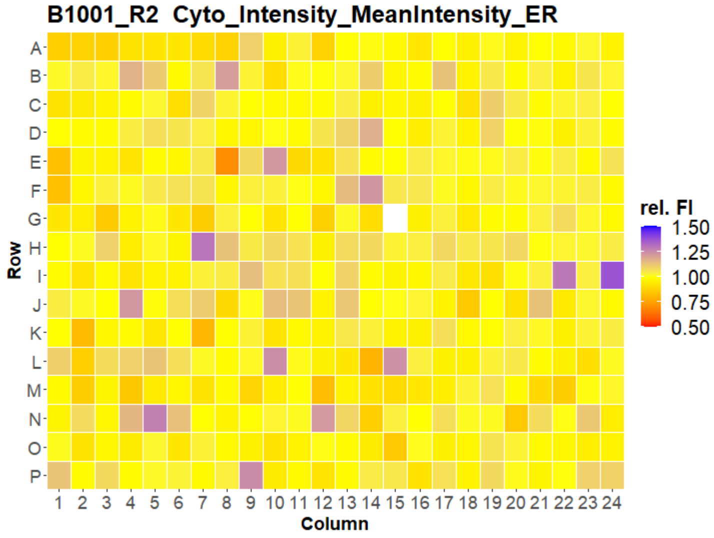


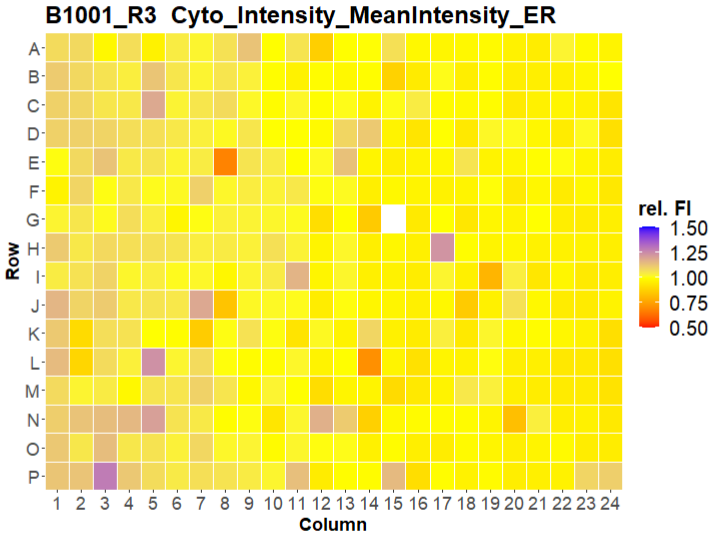


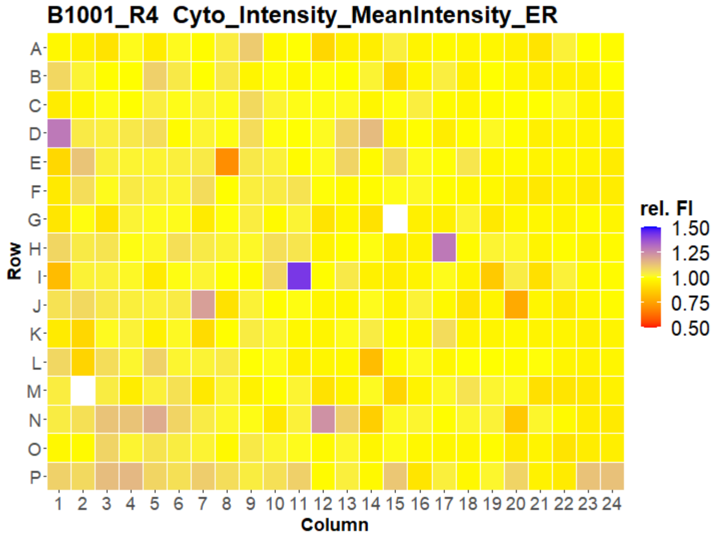


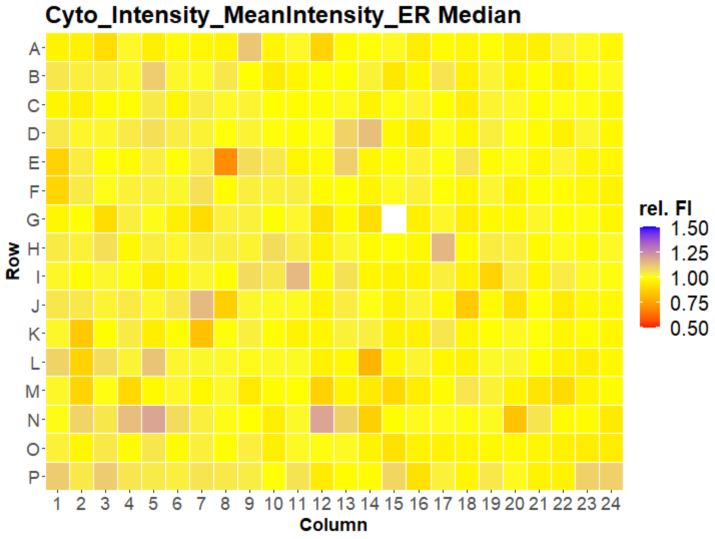

### HepG2 Medina

#### B1002 Cyto_Intensity_MeanIntensity_ER

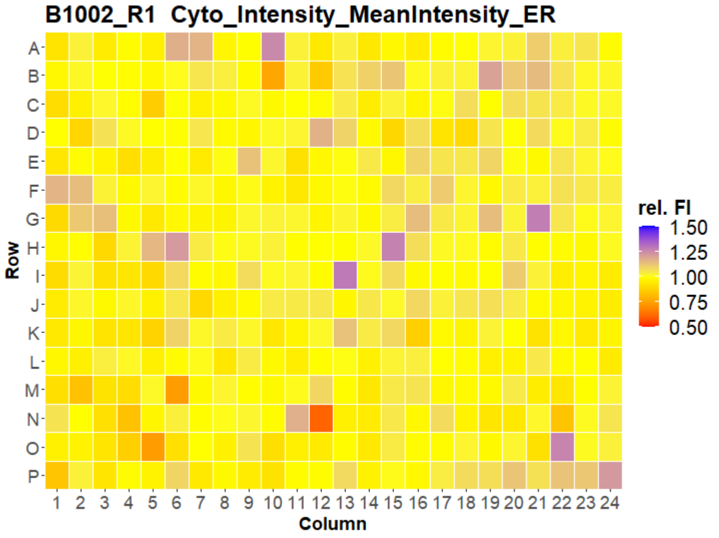


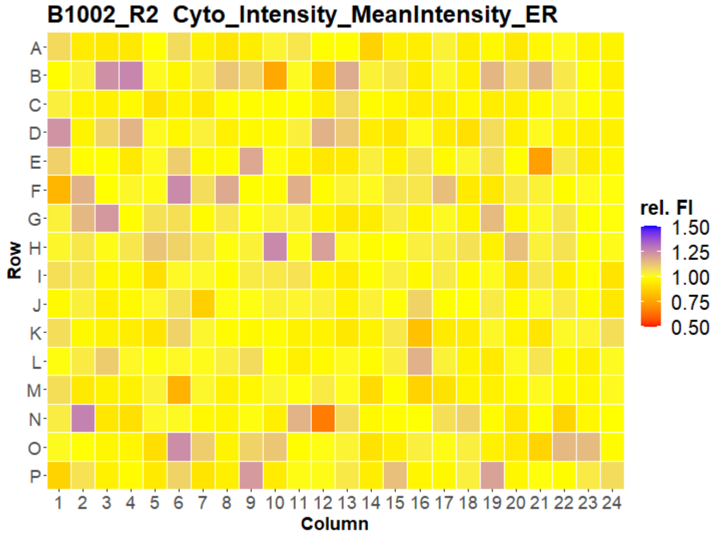


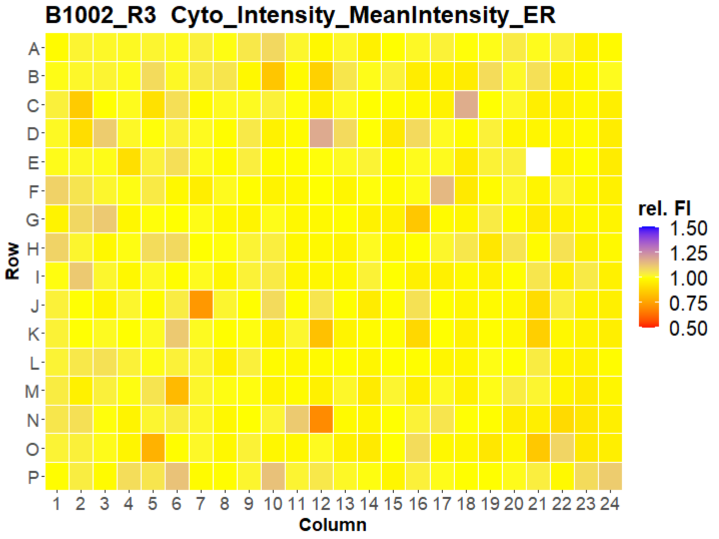


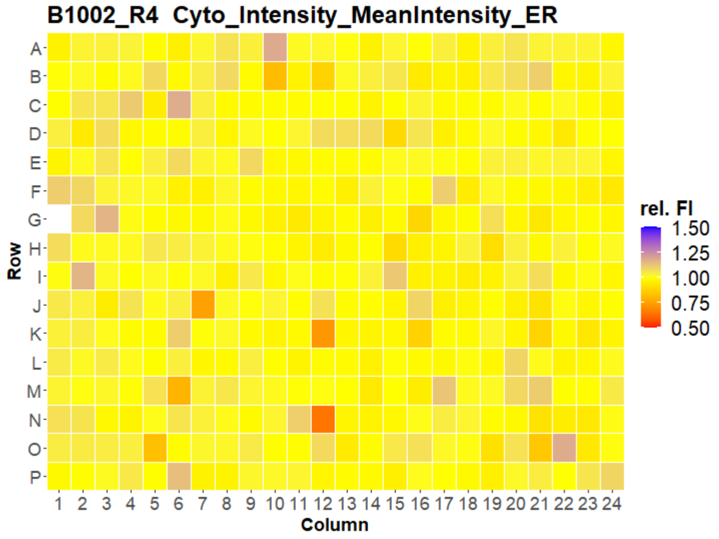


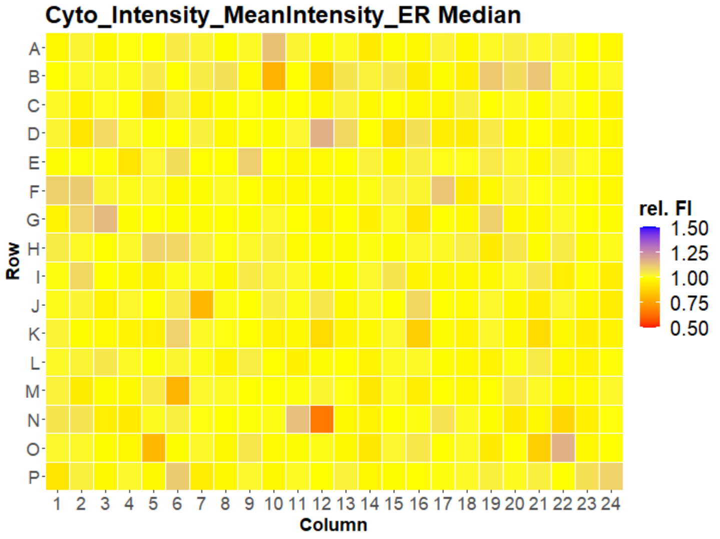

### HepG2 Medina

#### B1003 Cyto_Intensity_MeanIntensity_ER

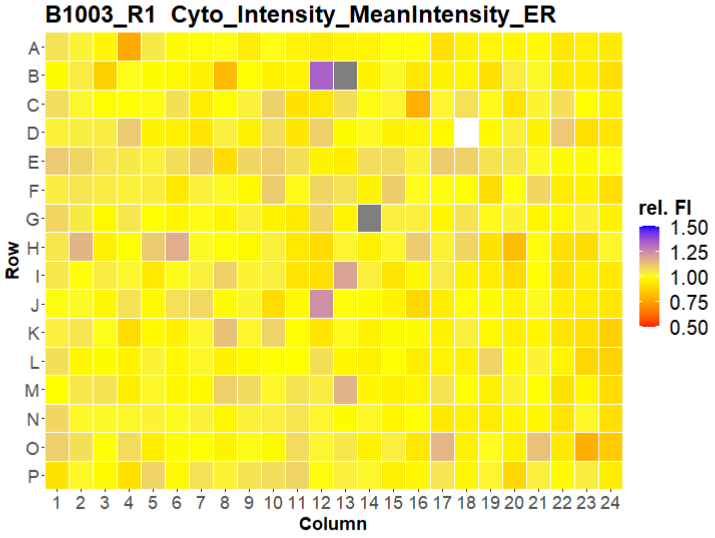


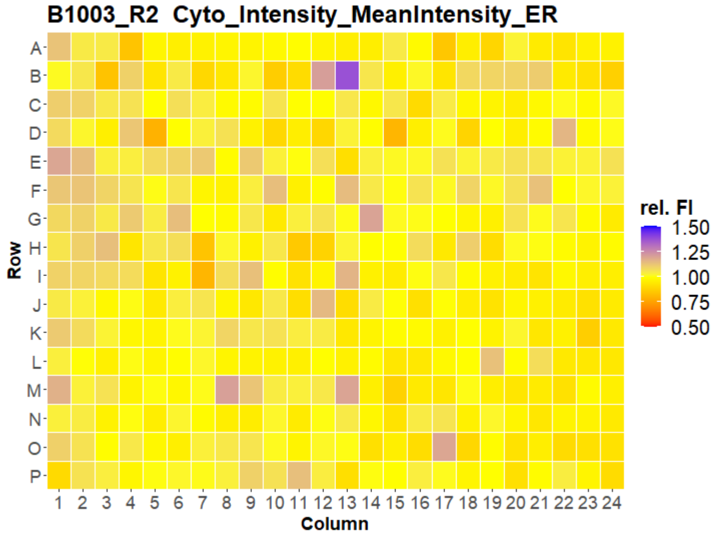


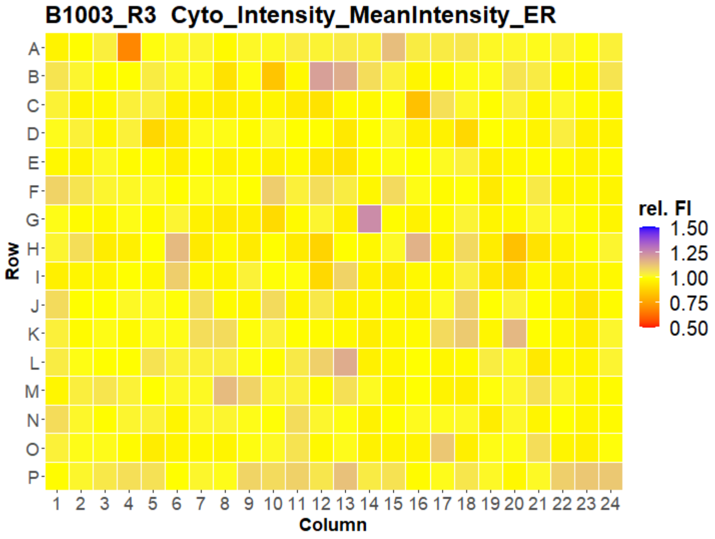


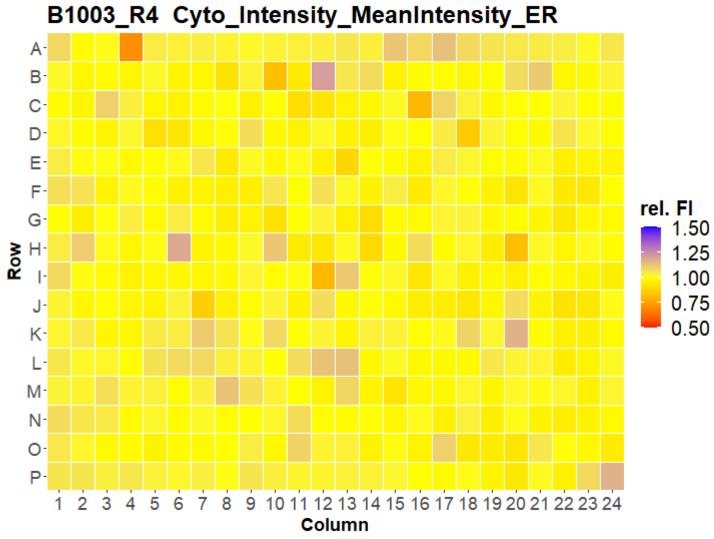


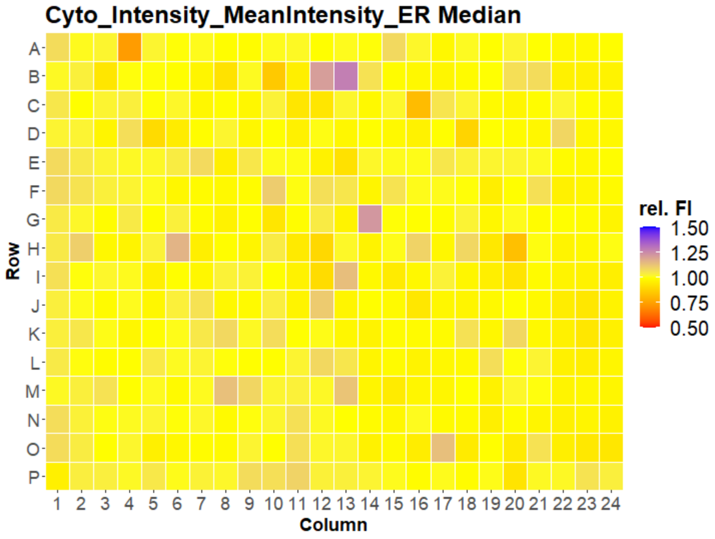

### HepG2 Medina

#### B1004 Cyto_Intensity_MeanIntensity_ER

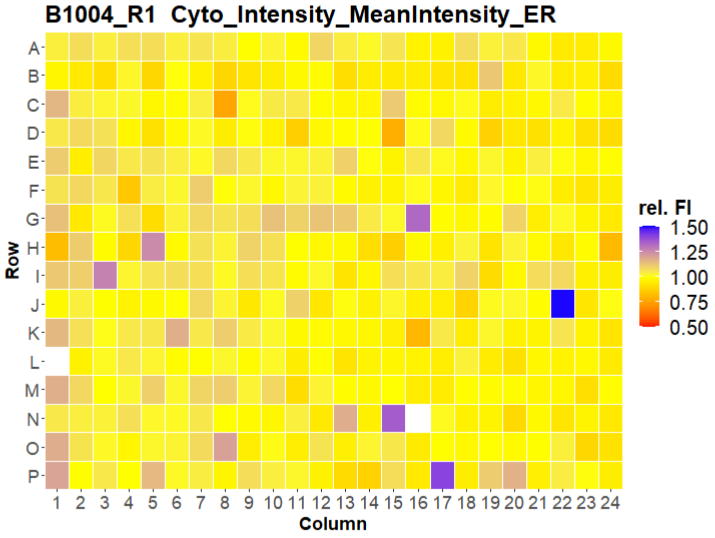


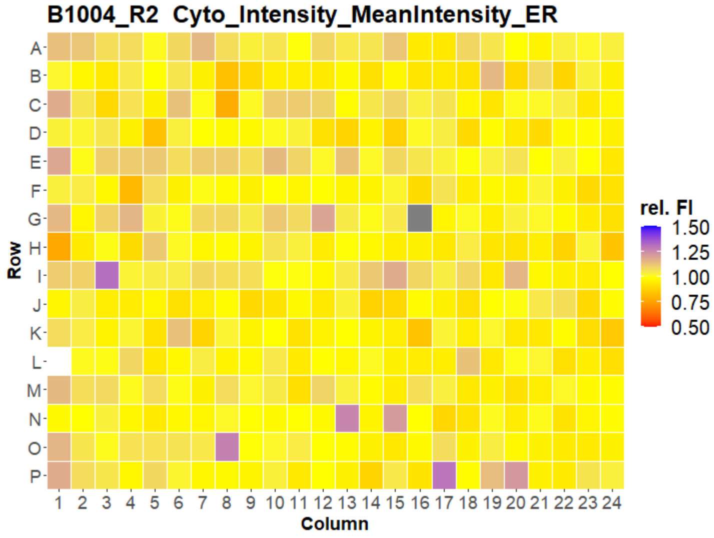


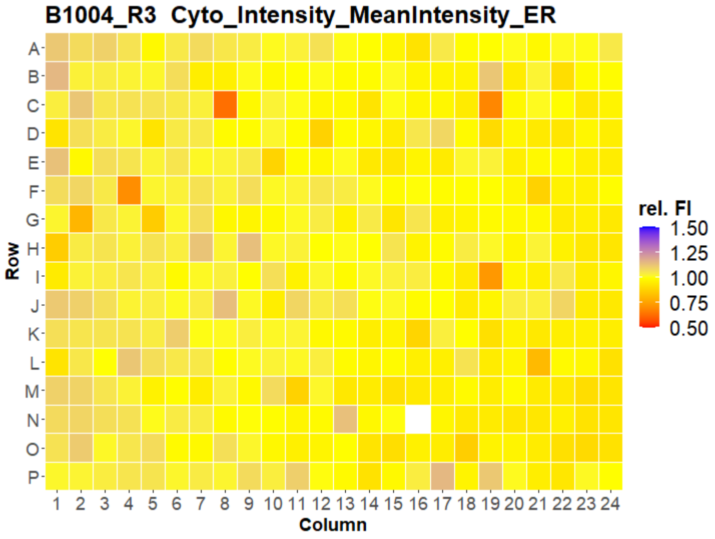


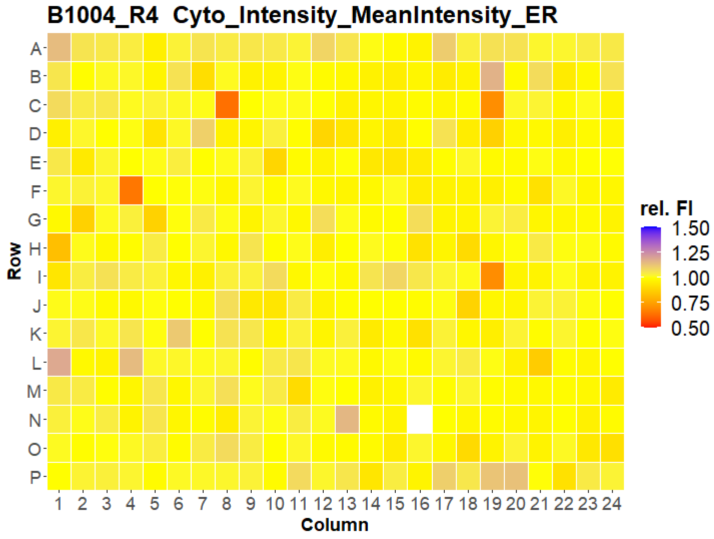


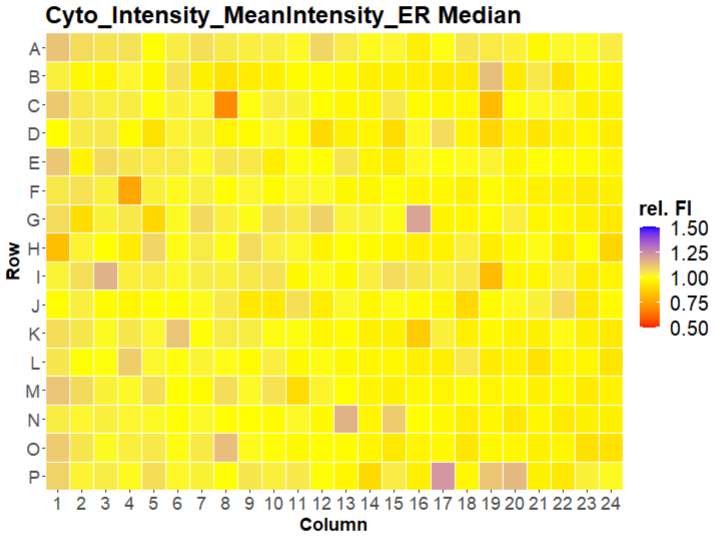

### HepG2 Medina

#### B1005 Cyto_Intensity_MeanIntensity_ER

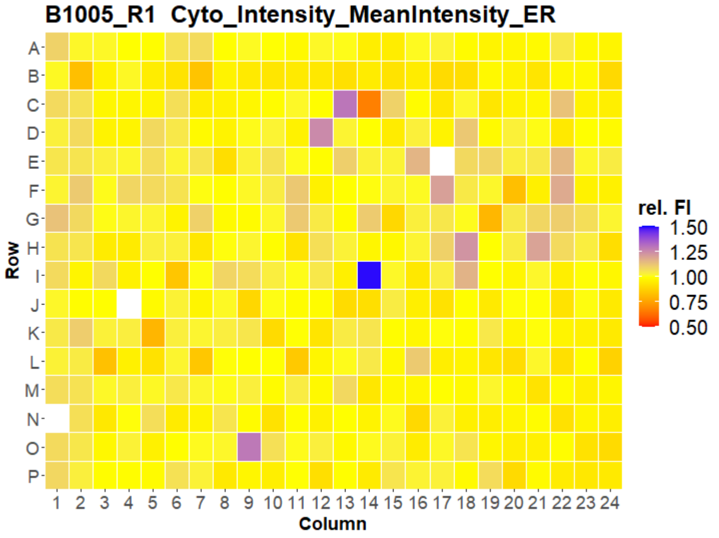


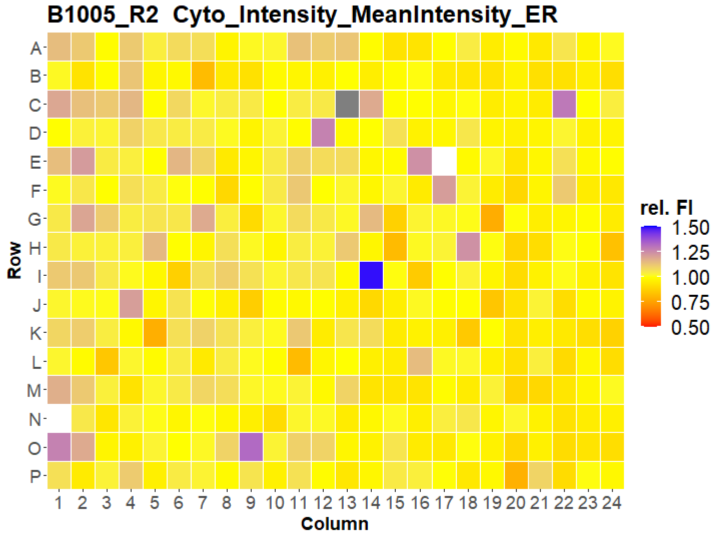


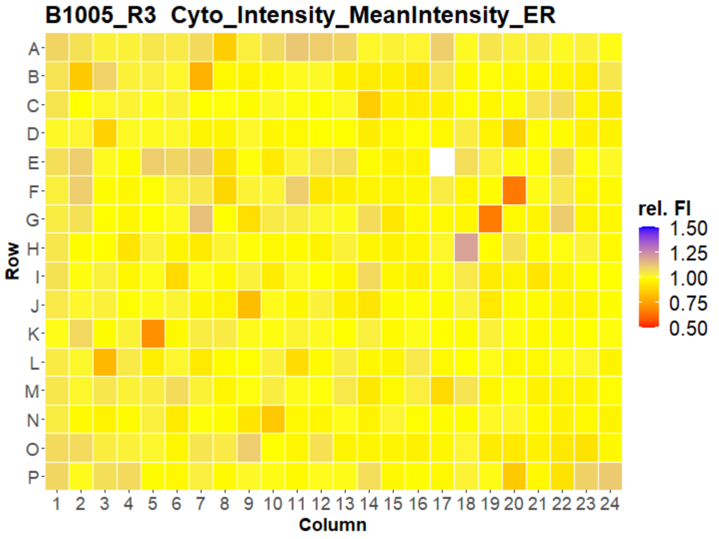


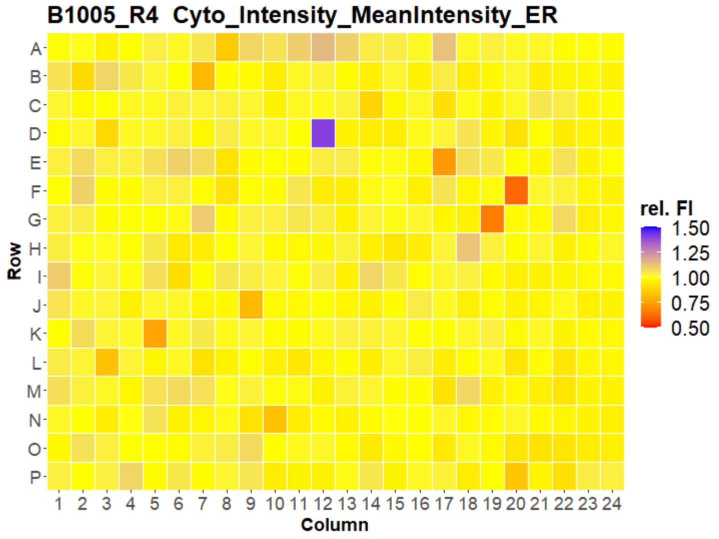


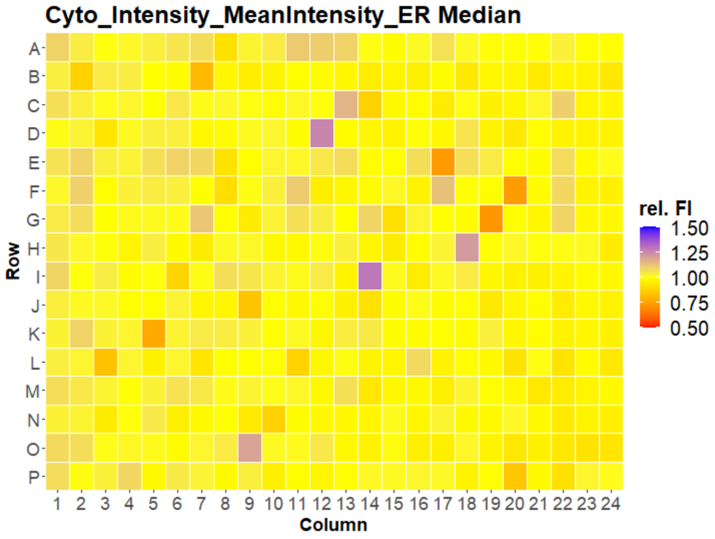


LEIBNIZ FORSCHUNGSINSTITUT FÜR MOLEKULARE PHARMAKOLOGIE
Open for Innovation KNIME

### HepG2 Medina

#### B1006 Cyto_Intensity_MeanIntensity_ER

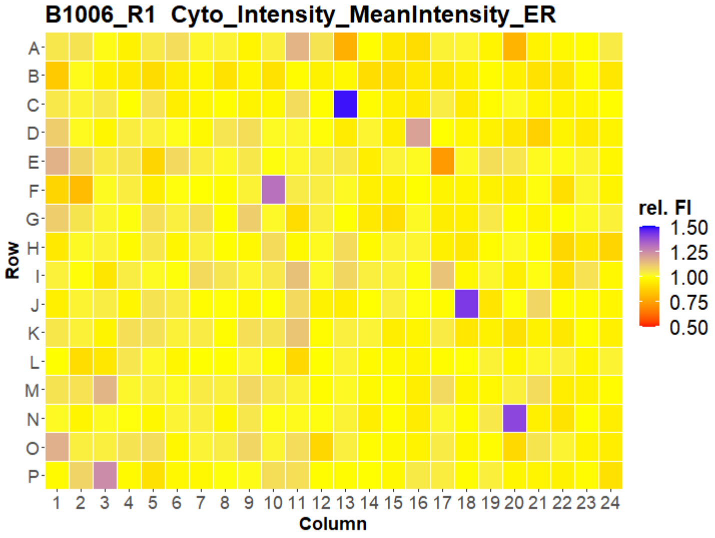


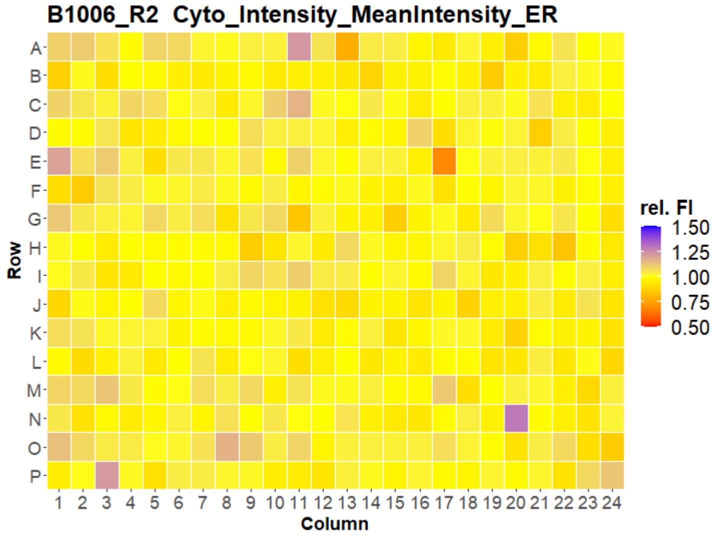


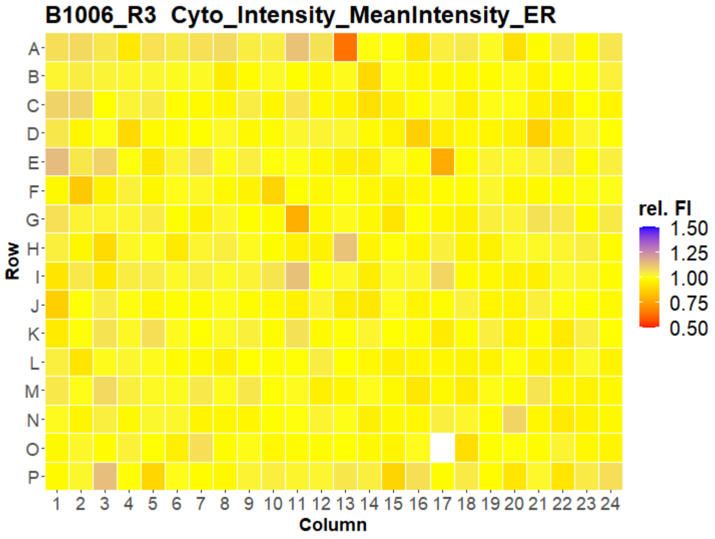


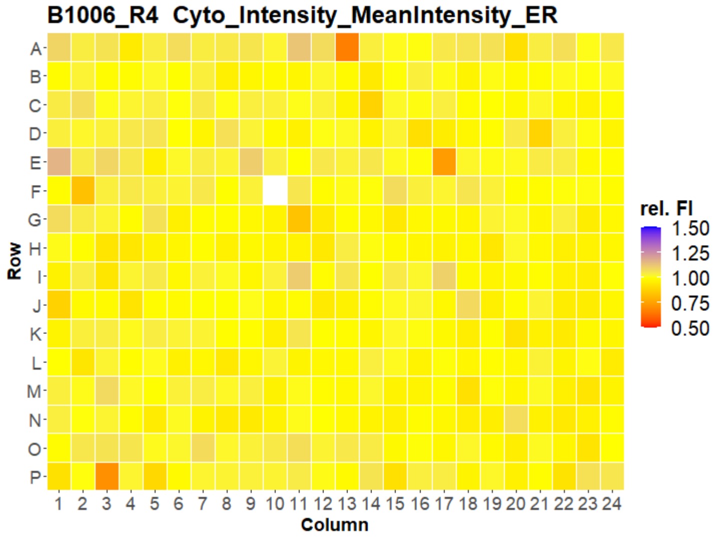


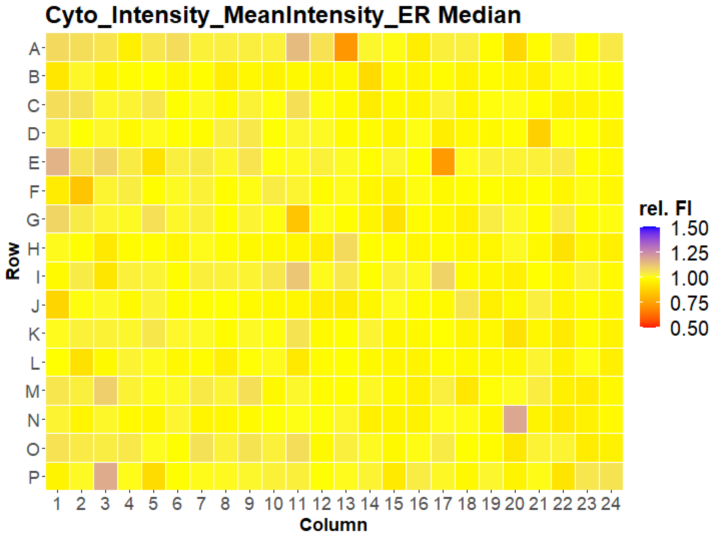

### HepG2 Medina

#### B1007 Cyto_Intensity_MeanIntensity_ER

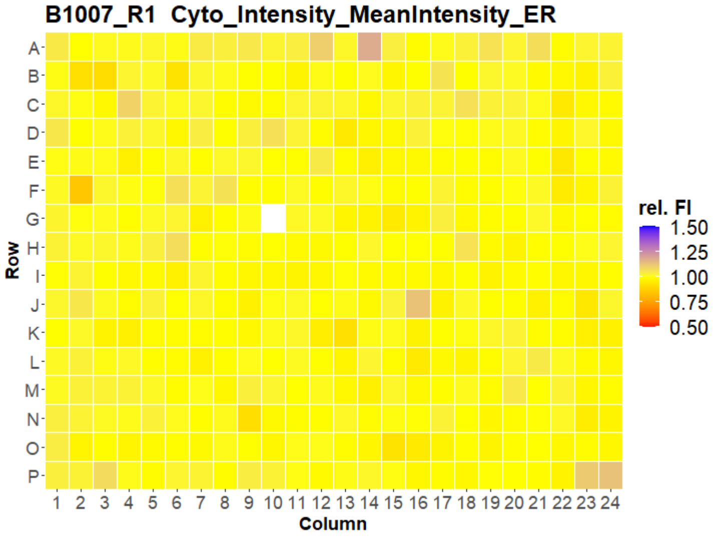


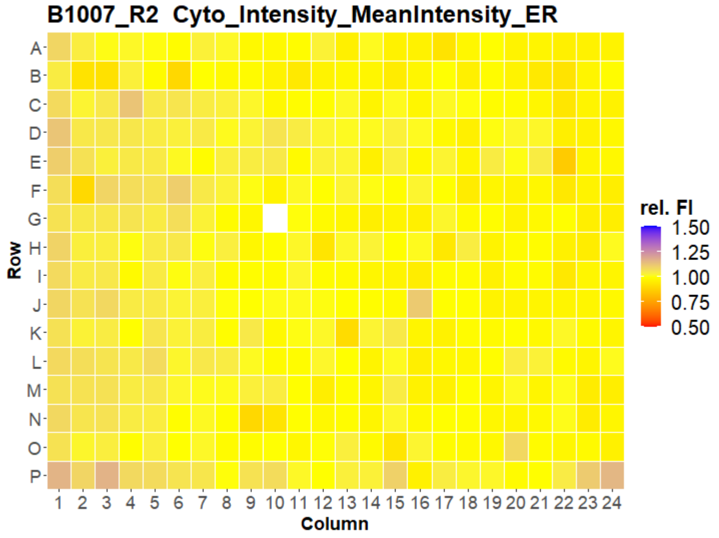


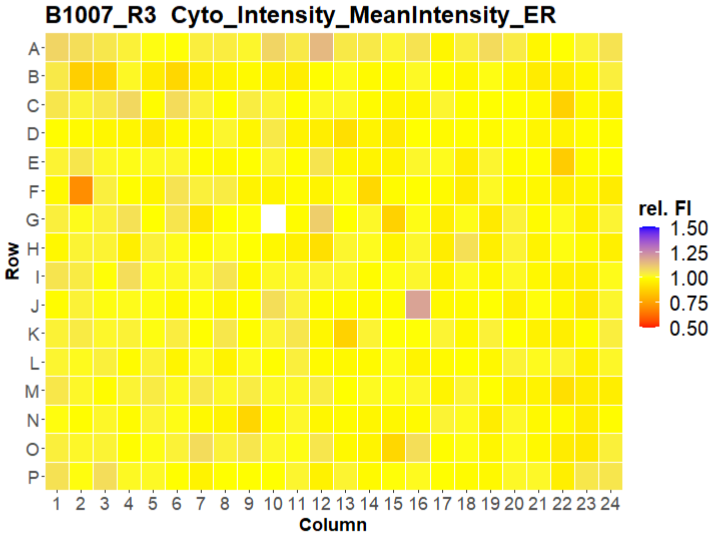


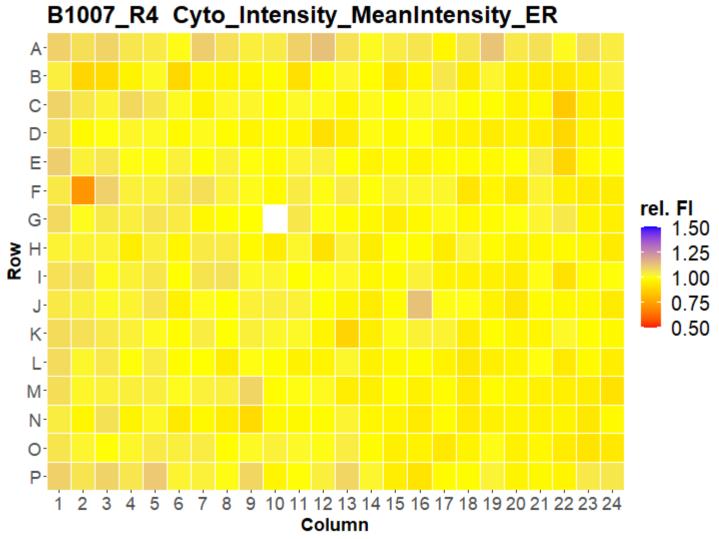


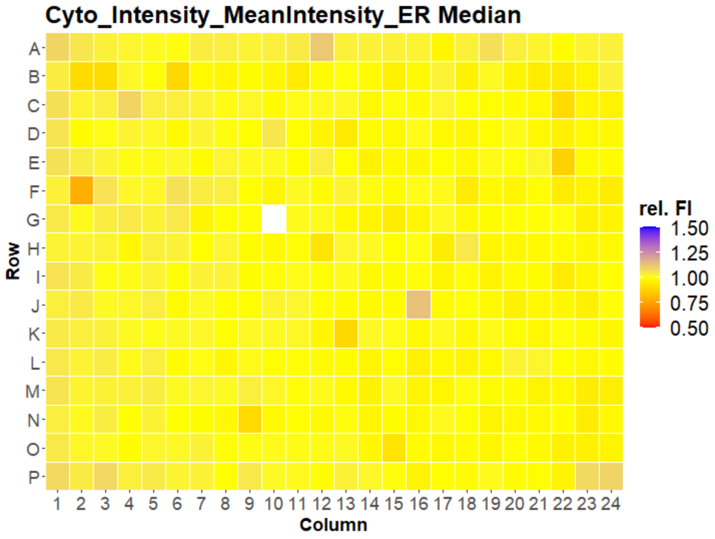

### HepG2 Medina

#### B1001 Cyto_Intensity_MeanIntensity_Mito

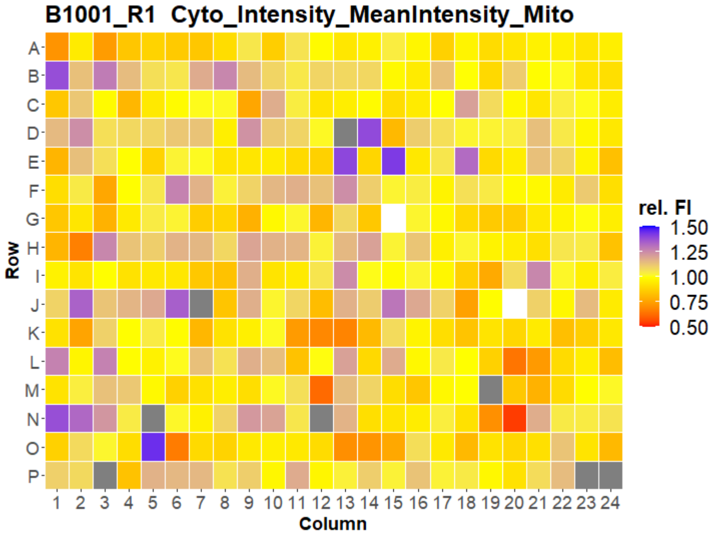


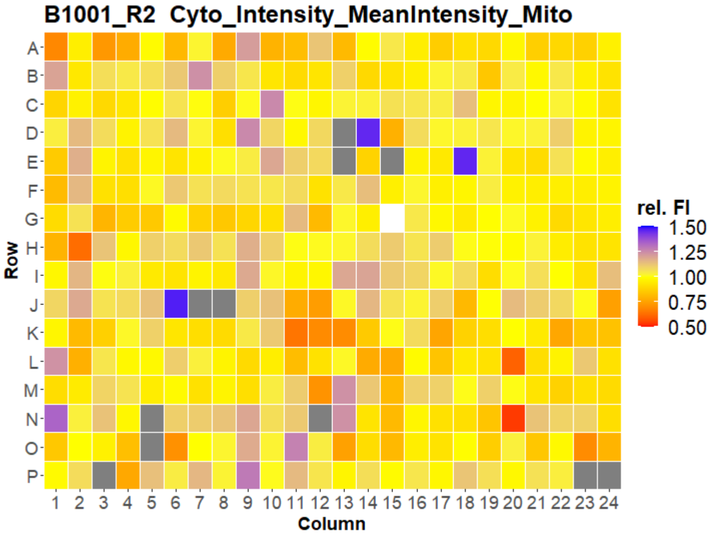


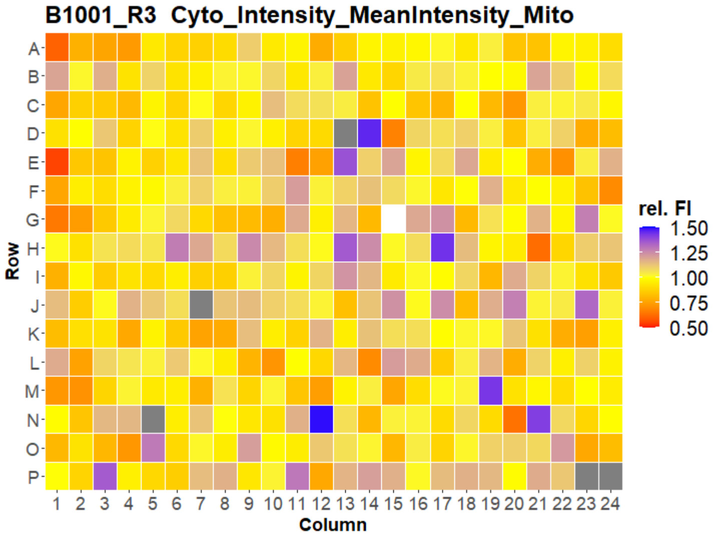


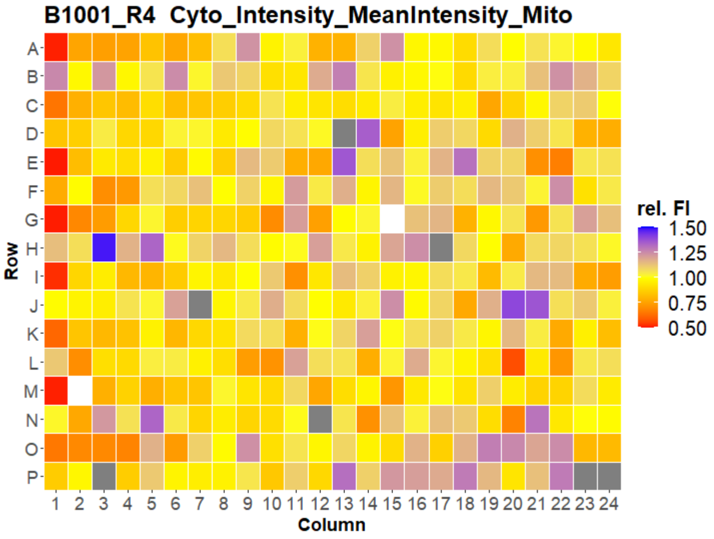


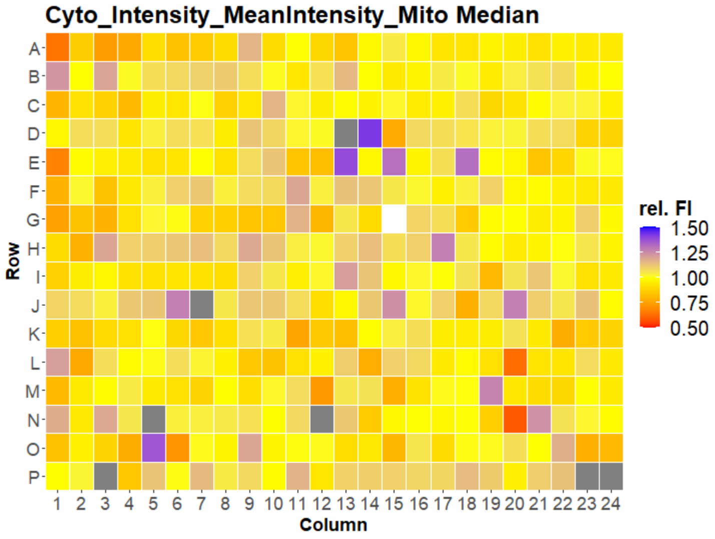

### HepG2 Medina

#### B1002 Cyto_Intensity_MeanIntensity_Mito

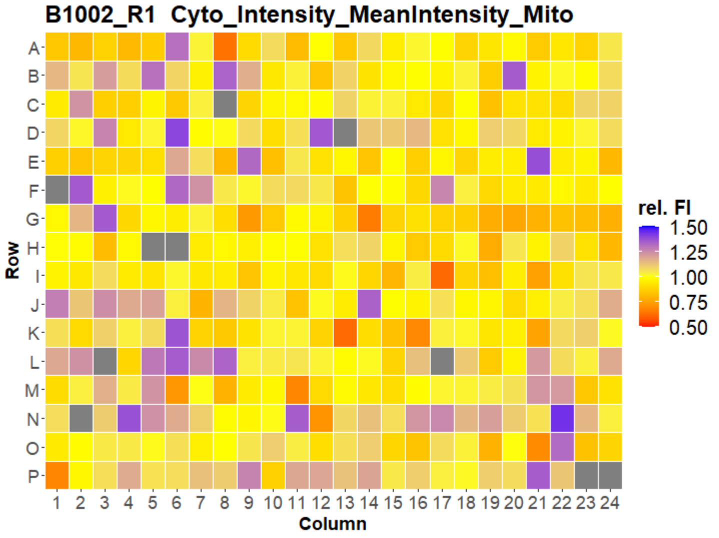


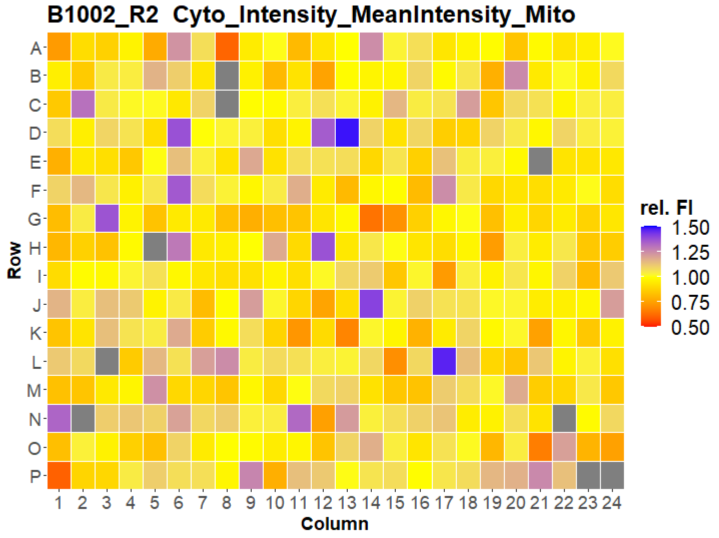


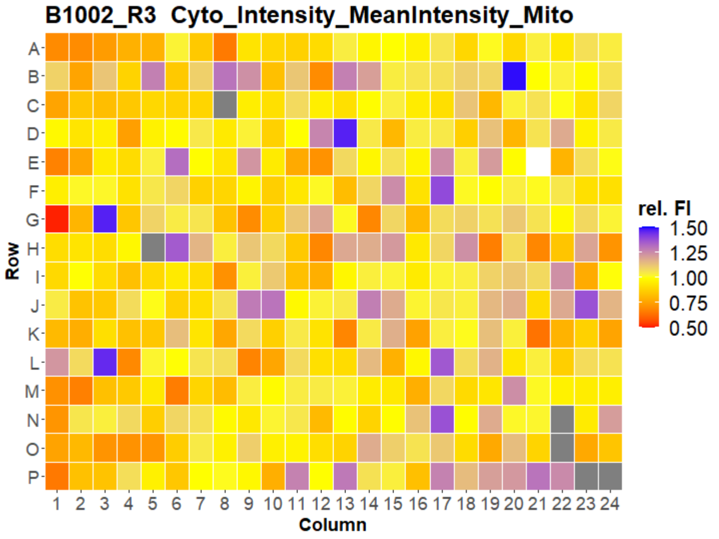


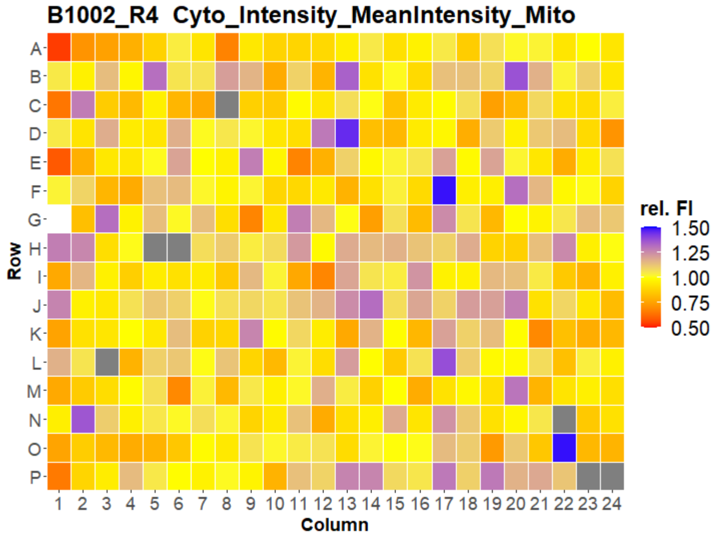


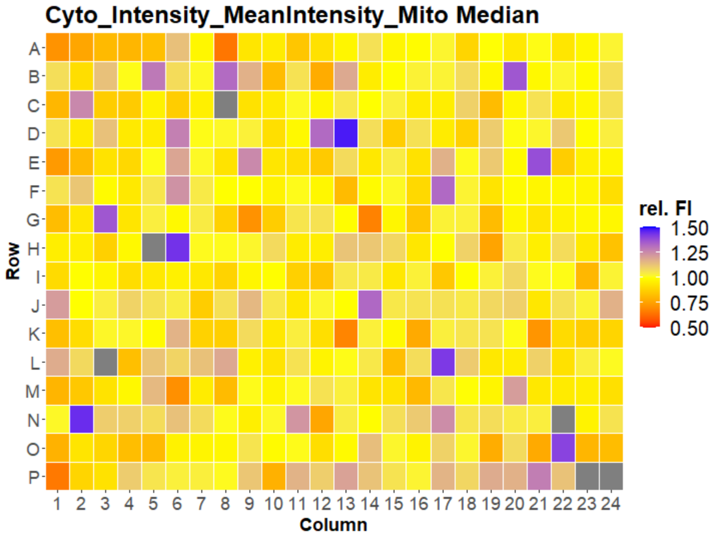

### HepG2 Medina

#### B1003 Cyto_Intensity_MeanIntensity_Mito

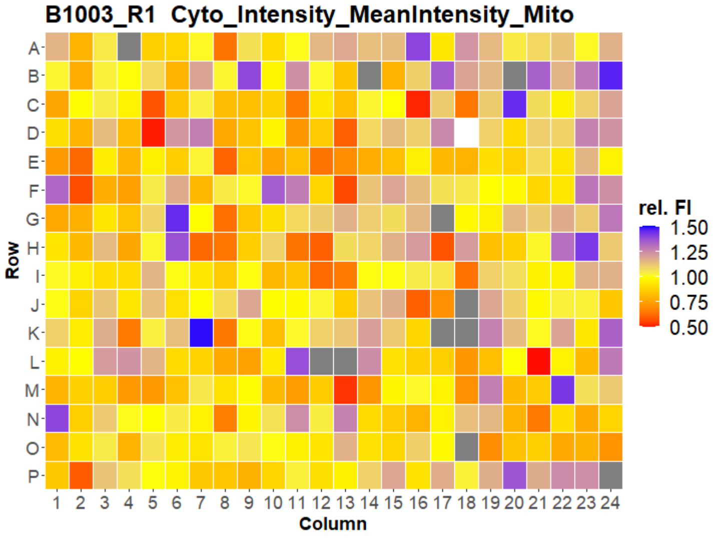


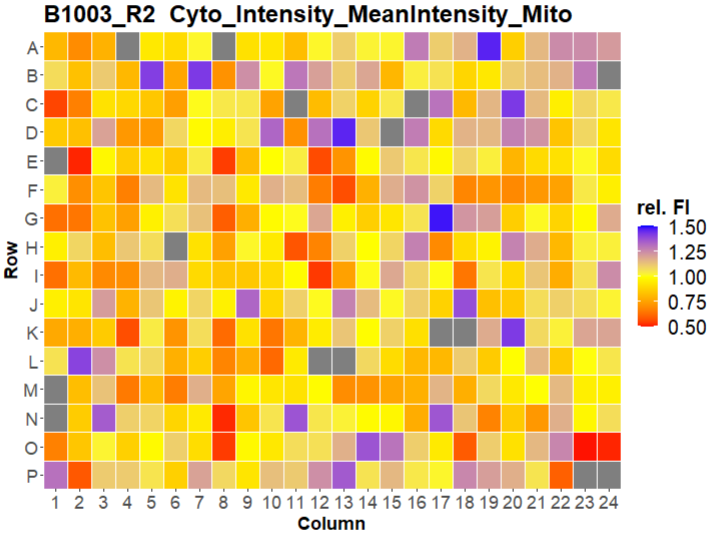


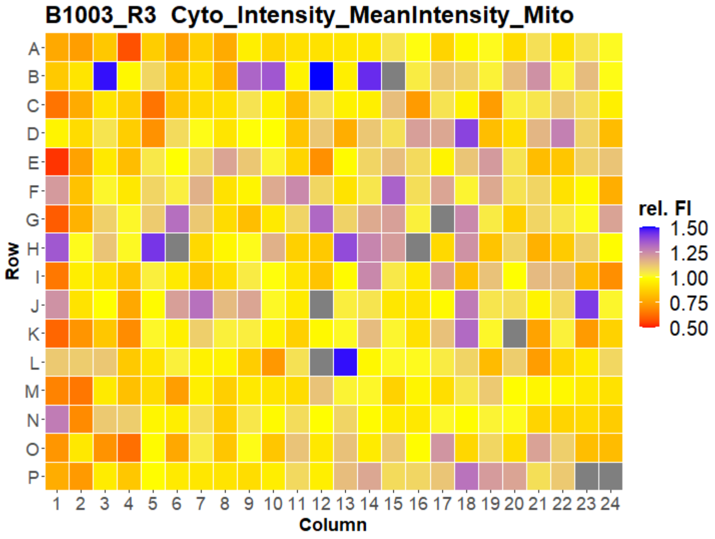


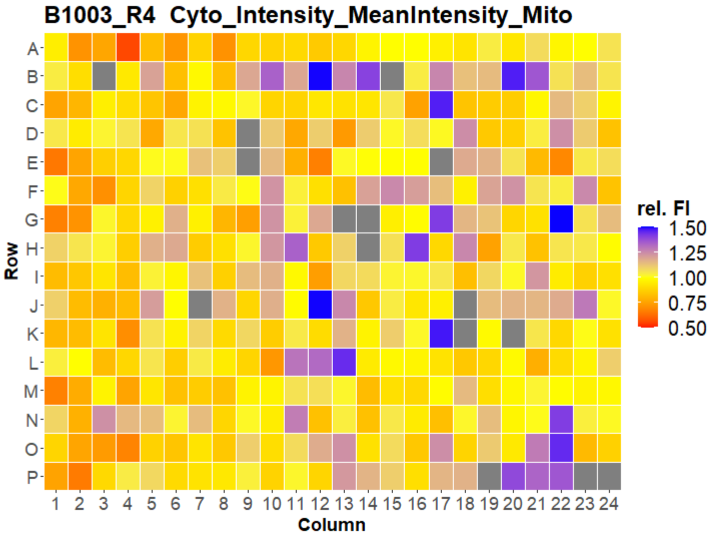


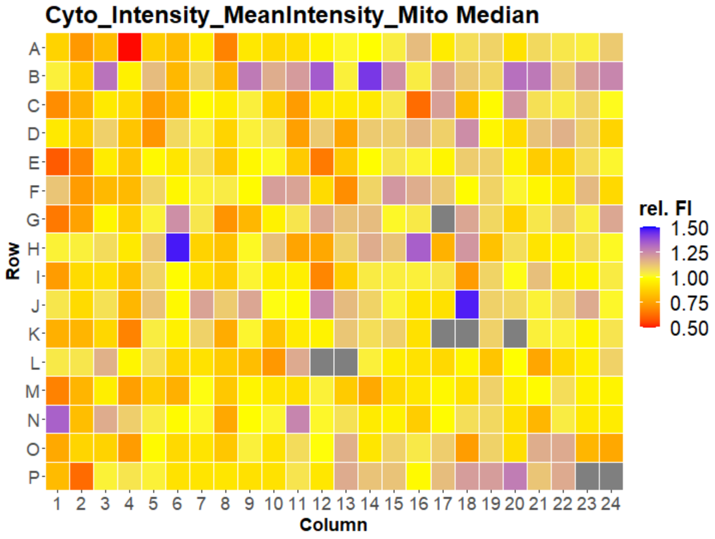

### HepG2 Medina

#### B1004 Cyto_Intensity_MeanIntensity_Mito

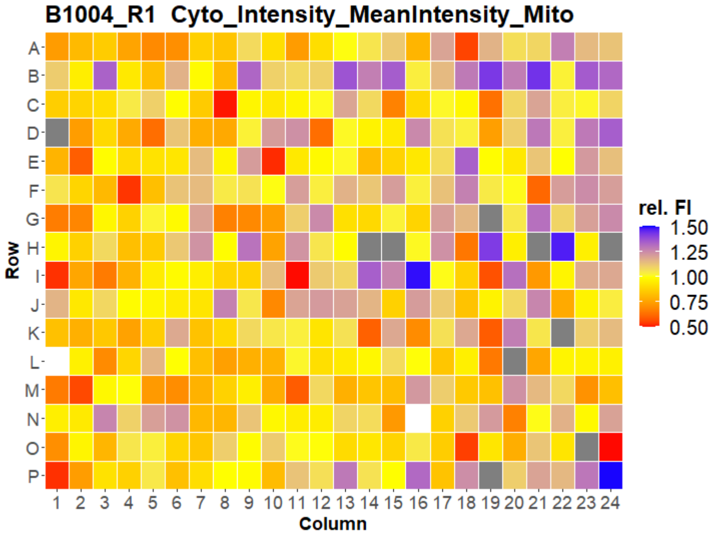


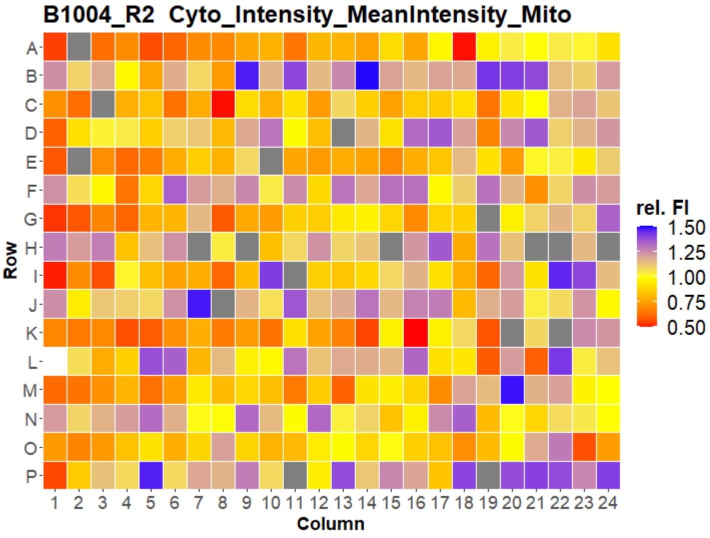


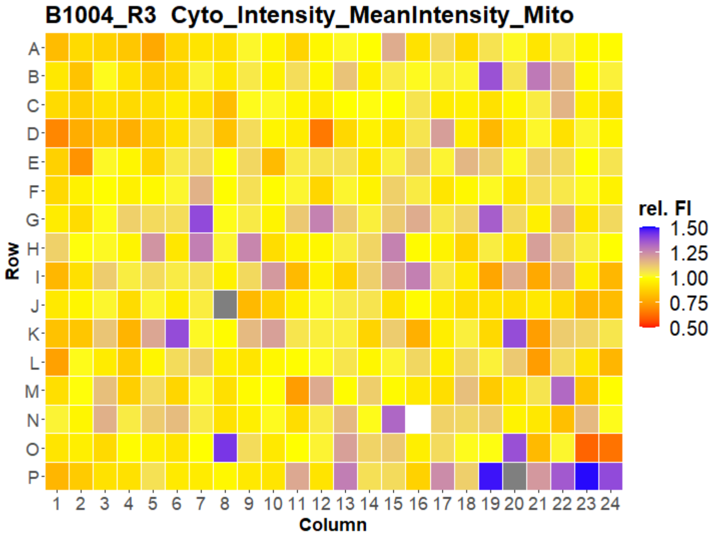


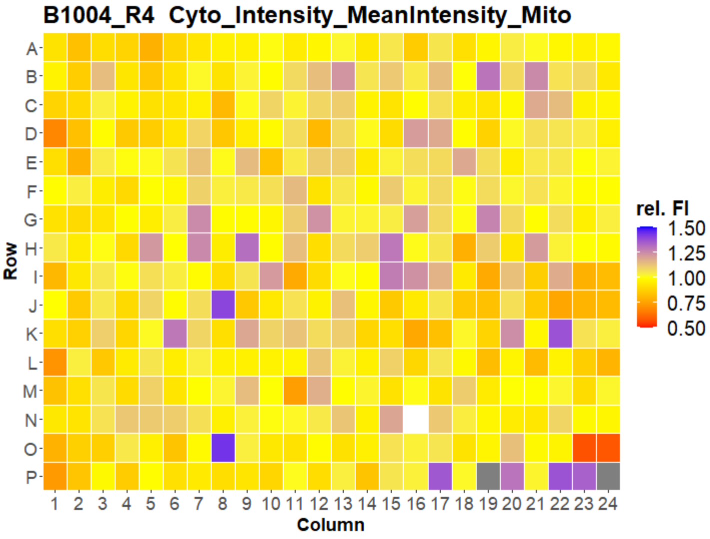


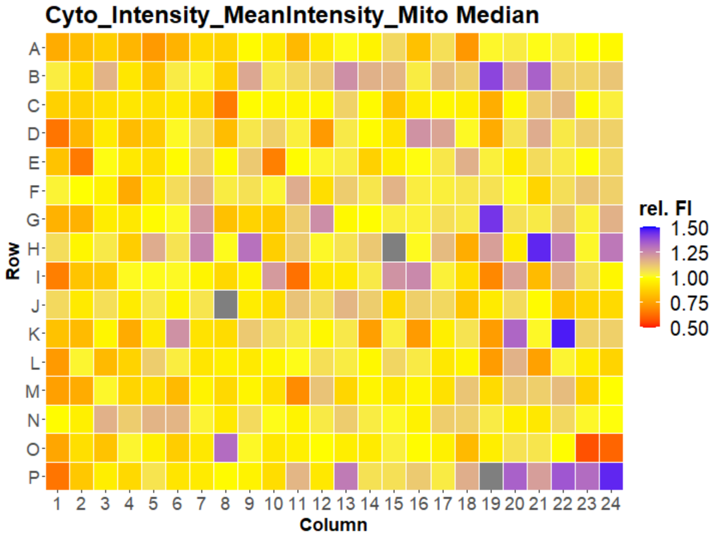

### HepG2 Medina

#### B1005 Cyto_Intensity_MeanIntensity_Mito

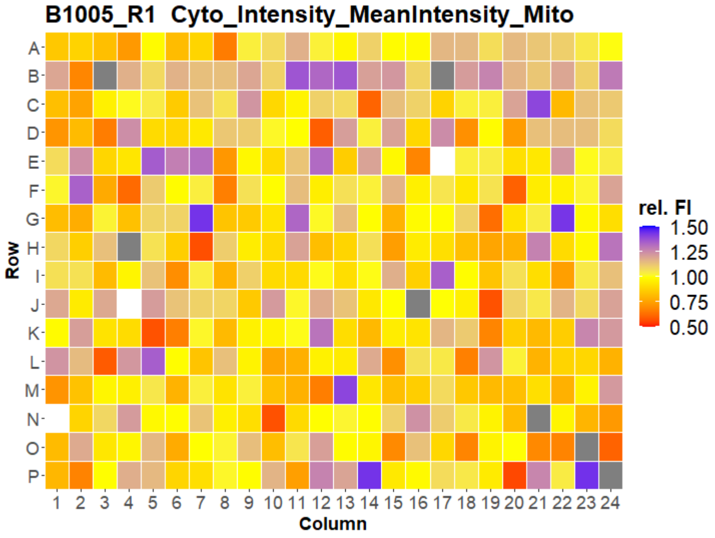


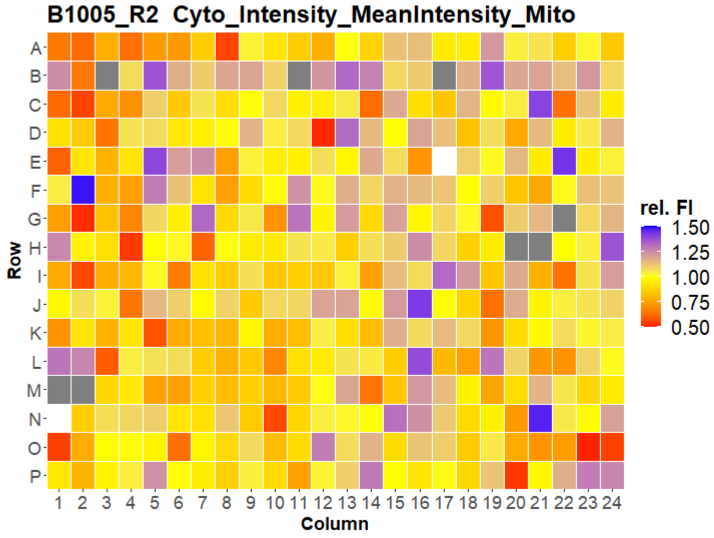


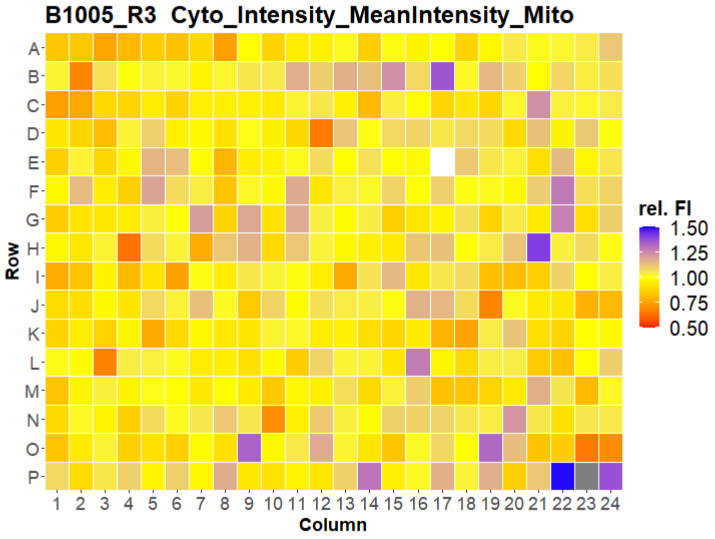


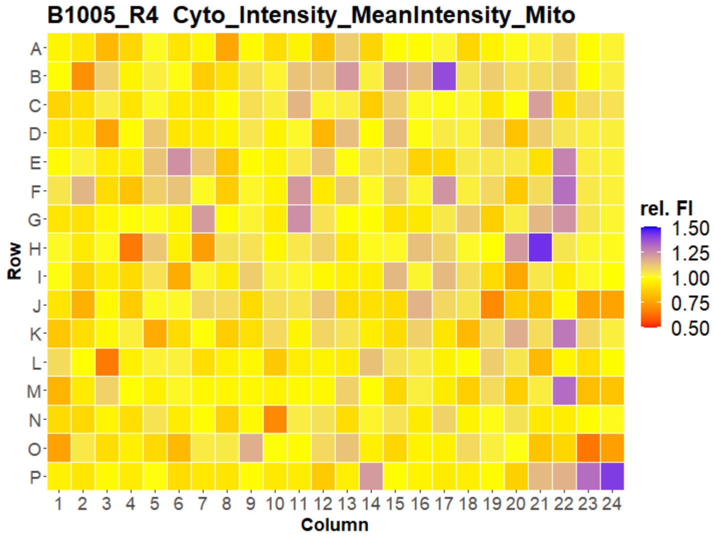


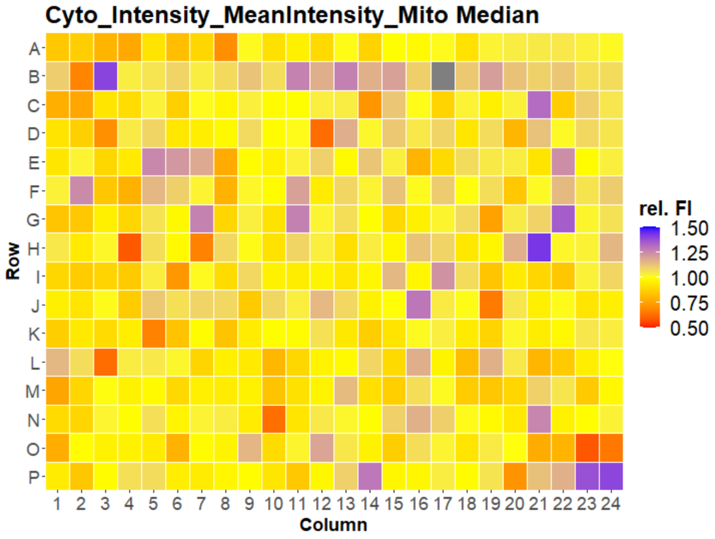

### HepG2 Medina

#### B1006 Cyto_Intensity_MeanIntensity_Mito

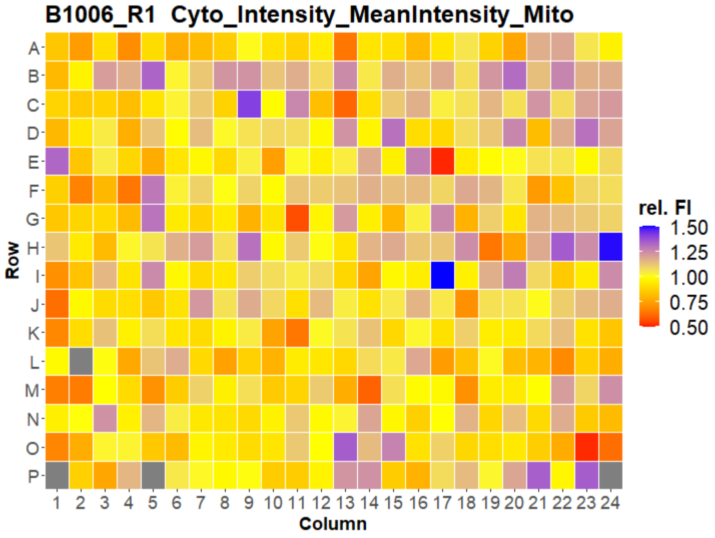


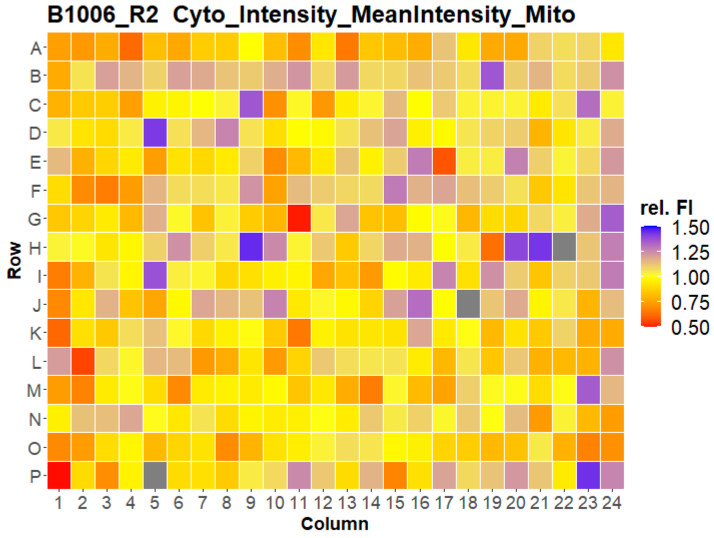


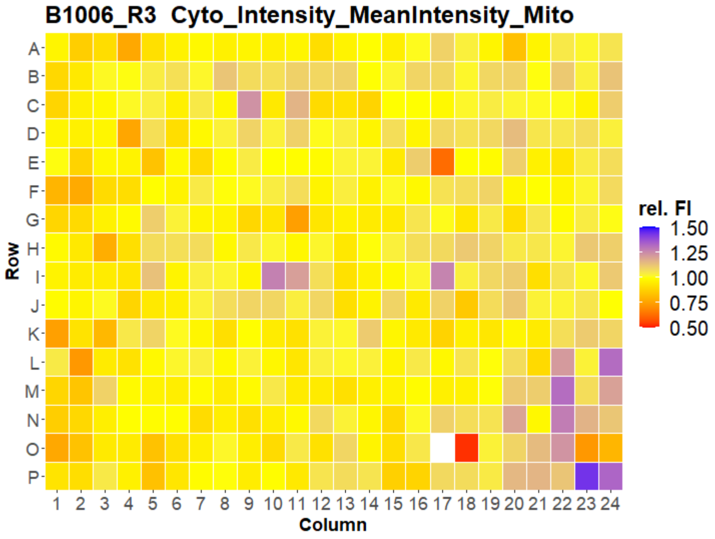


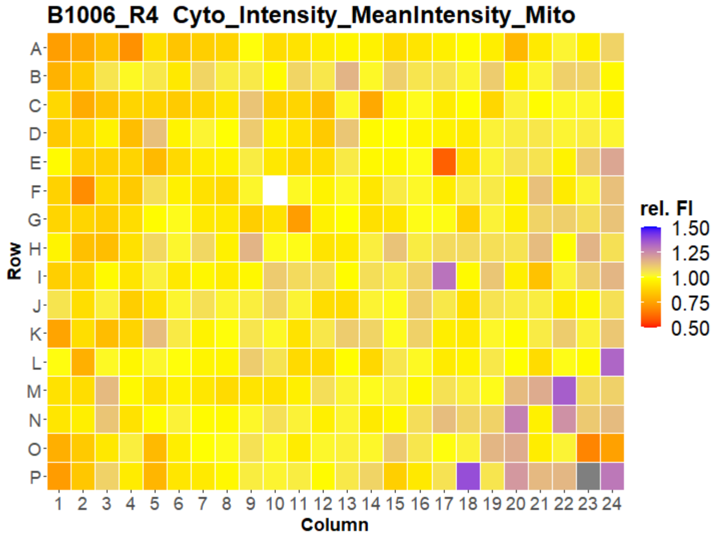


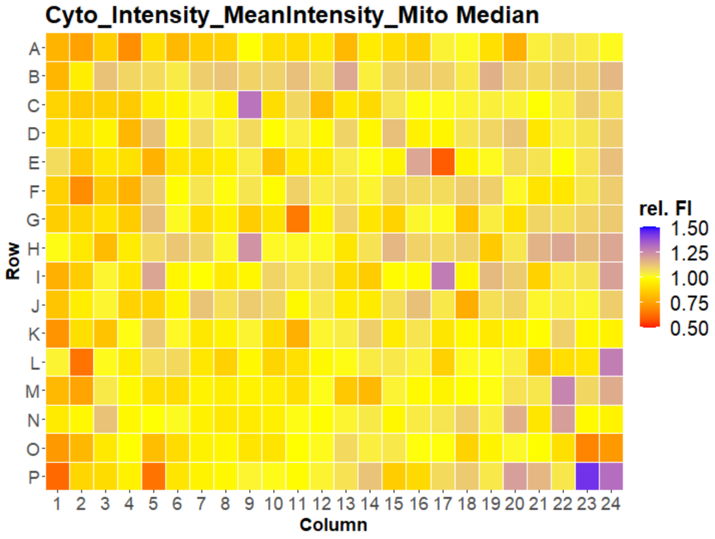

### HepG2 Medina

#### B1007 Cyto_Intensity_MeanIntensity_Mito

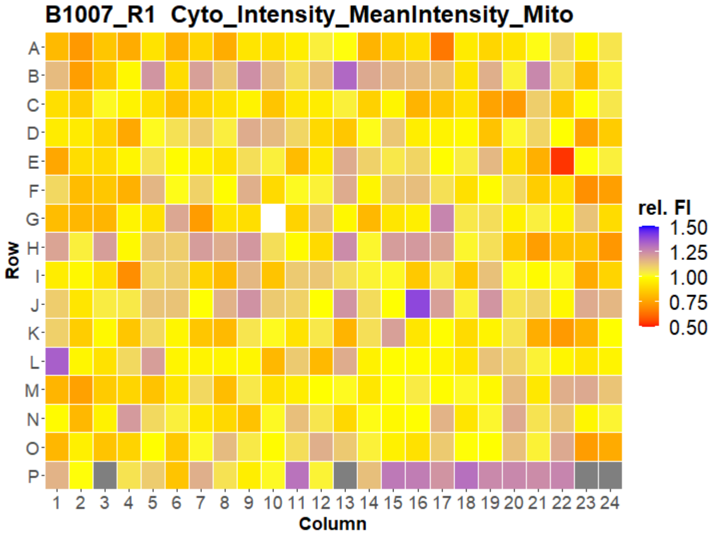


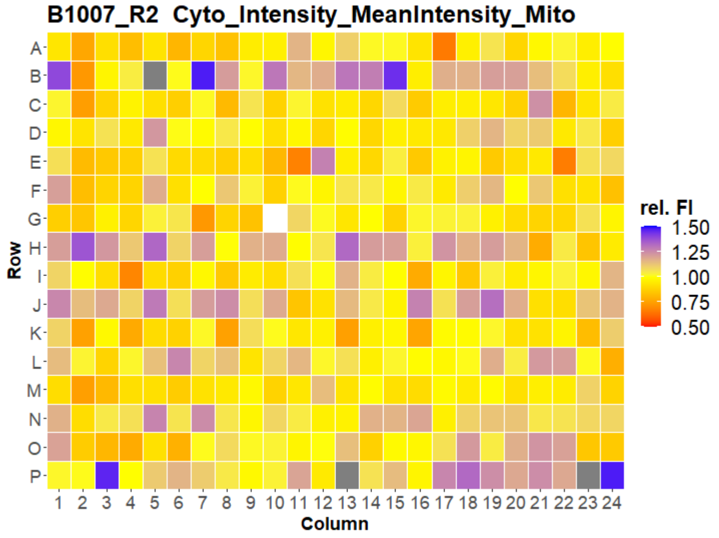


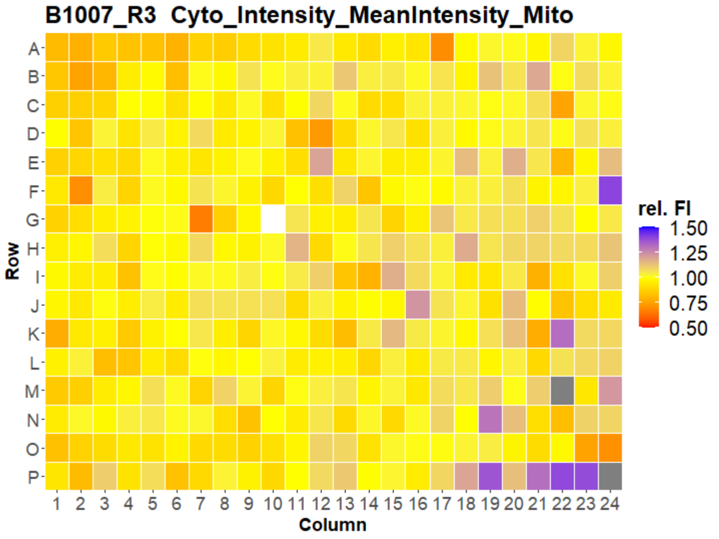


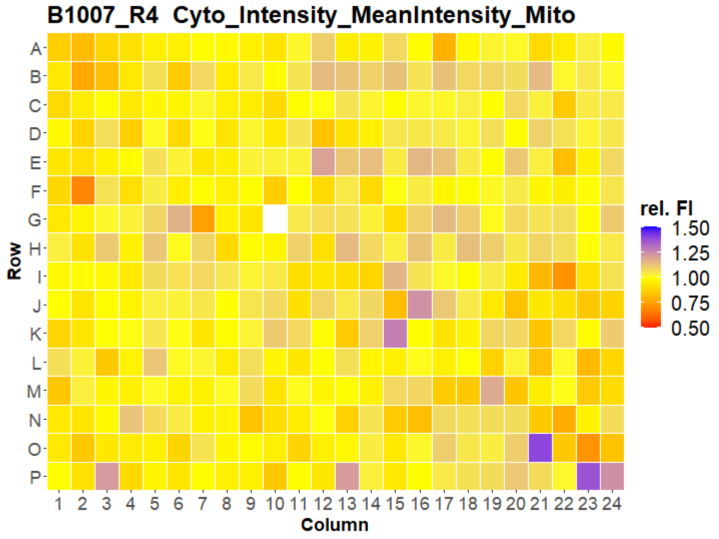


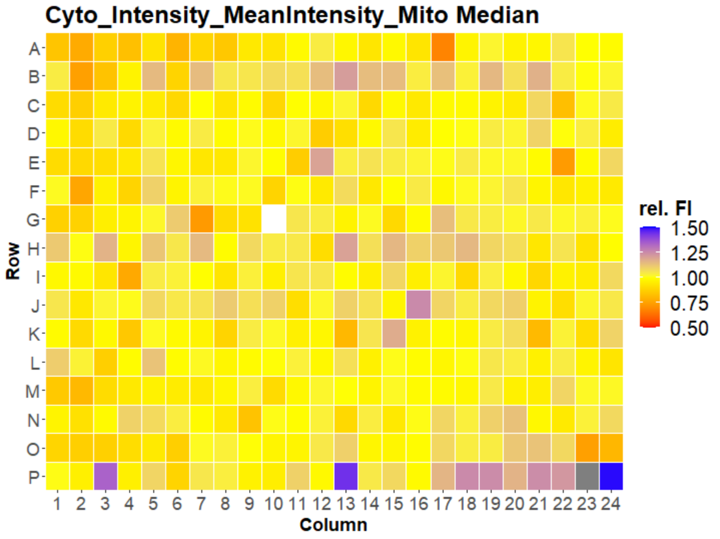


LEIBNIZ FORSCHUNGSINSTITUT FÜR MOLEKULARE PHARMAKOLOGIE
Open for Innovation KNIME

### HepG2 Medina

#### B1001 Nuc_Intensity_MeanIntensity_DNA

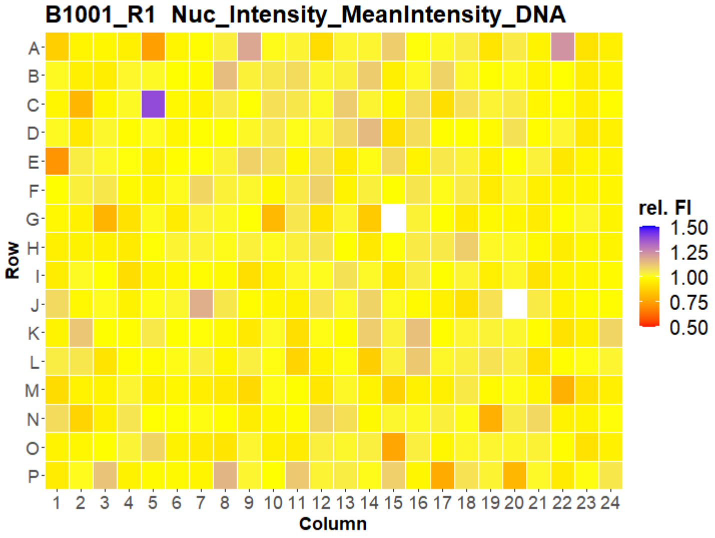


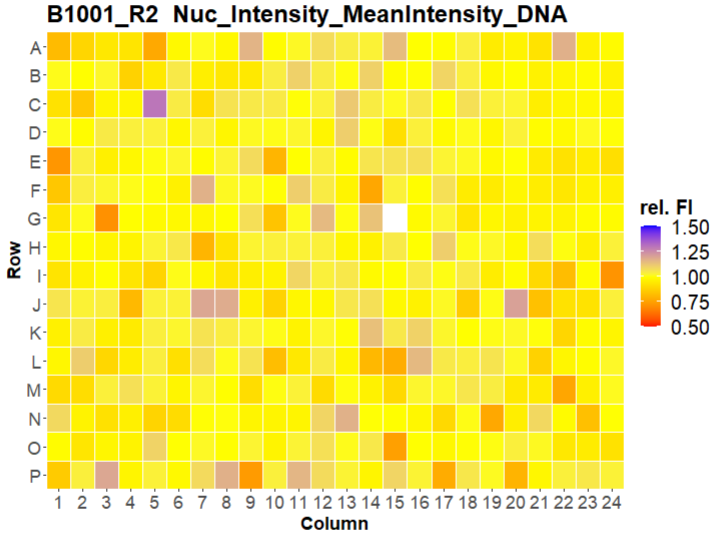


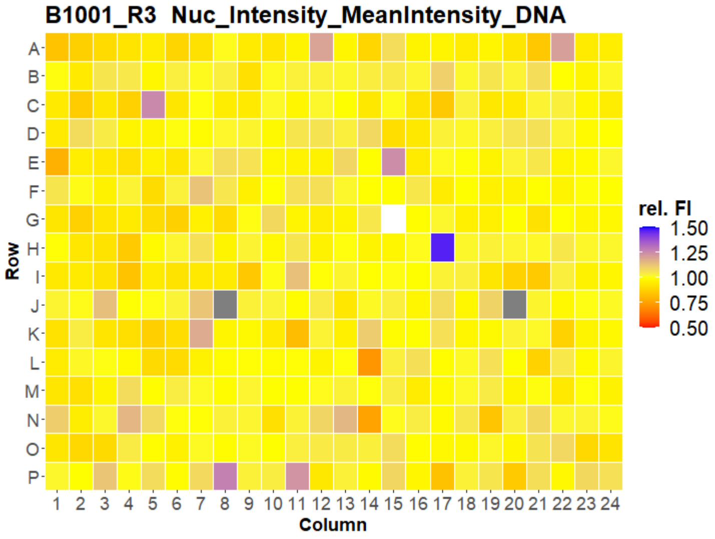


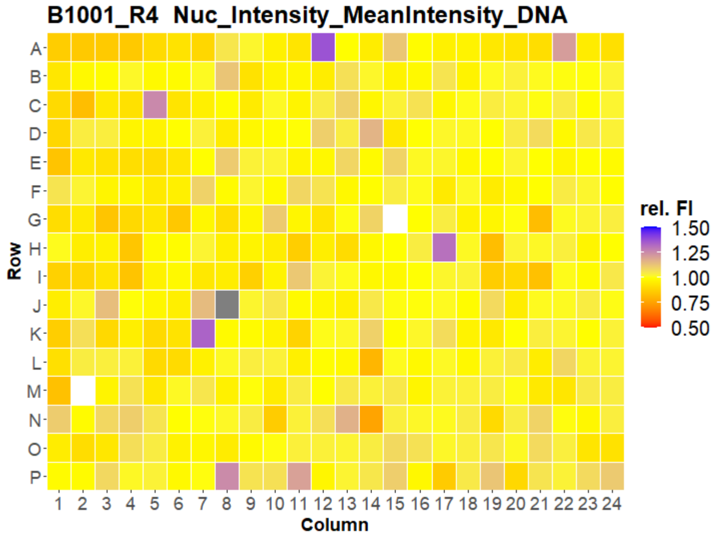


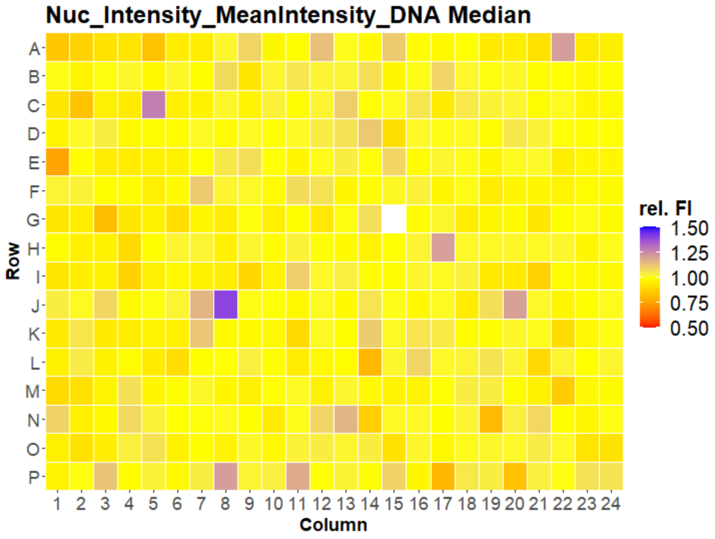

### HepG2 Medina

#### B1002 Nuc_Intensity_MeanIntensity_DNA

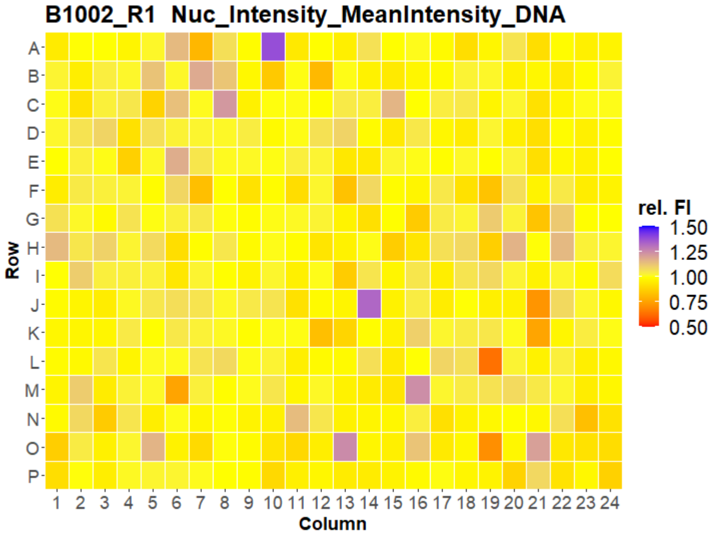


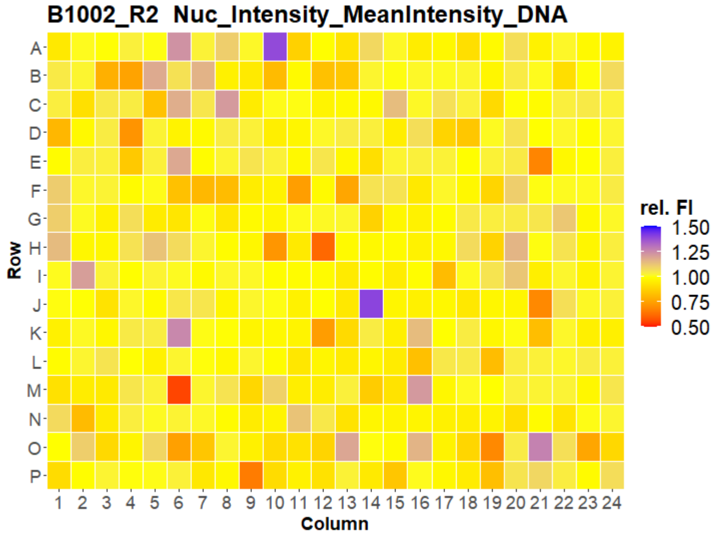


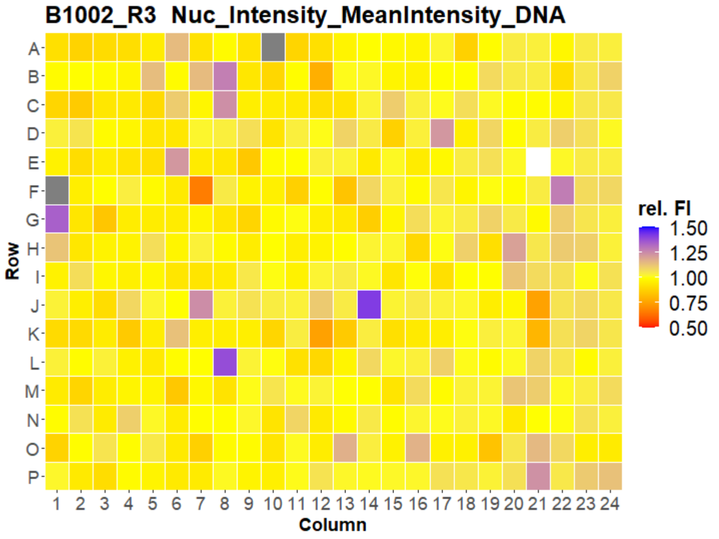


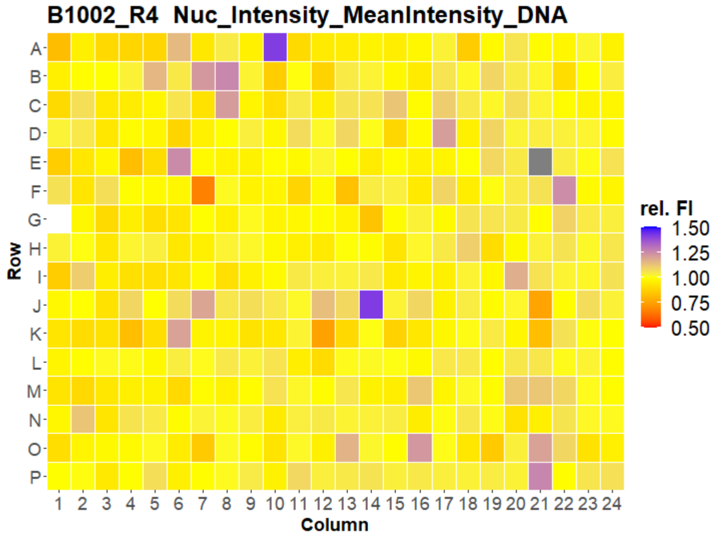


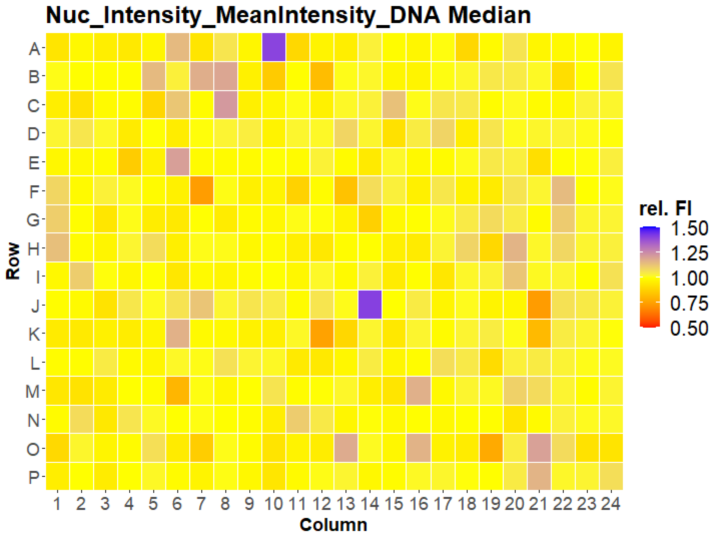

### HepG2 Medina

#### B1003 Nuc_Intensity_MeanIntensity_DNA

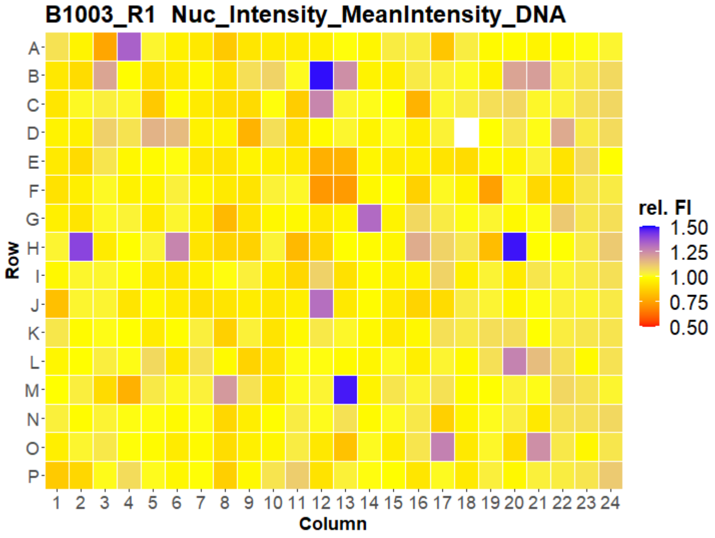


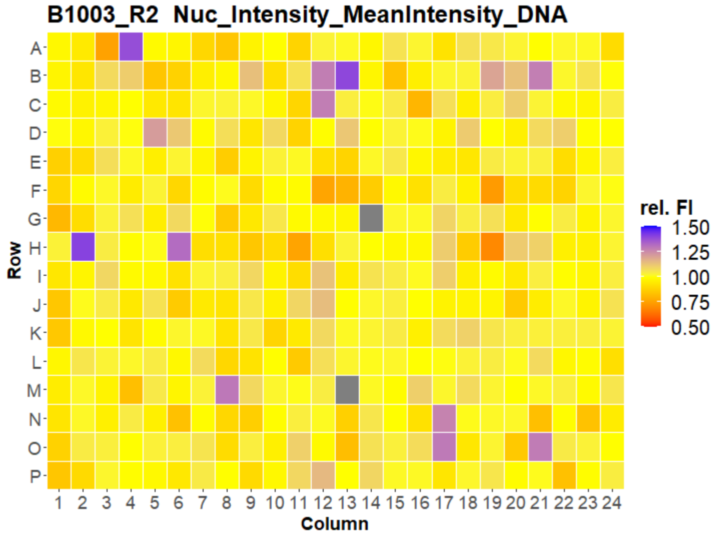


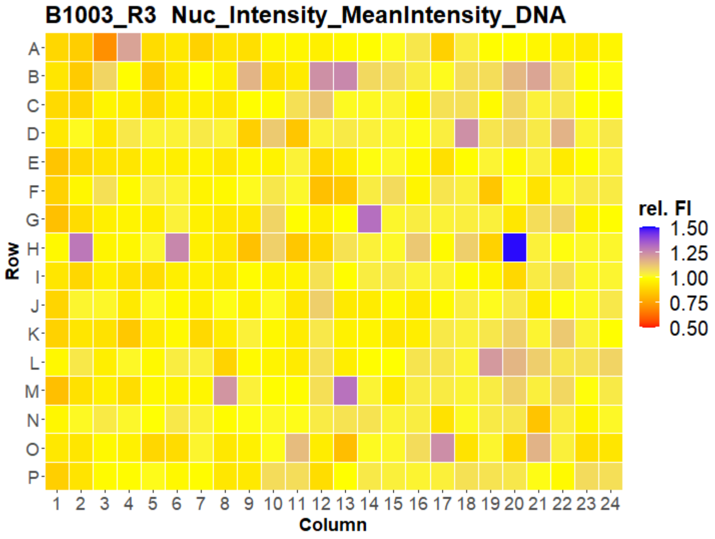


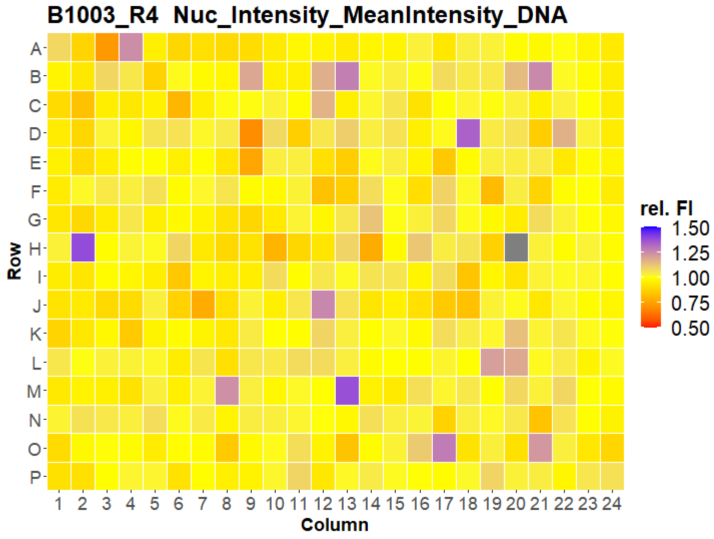


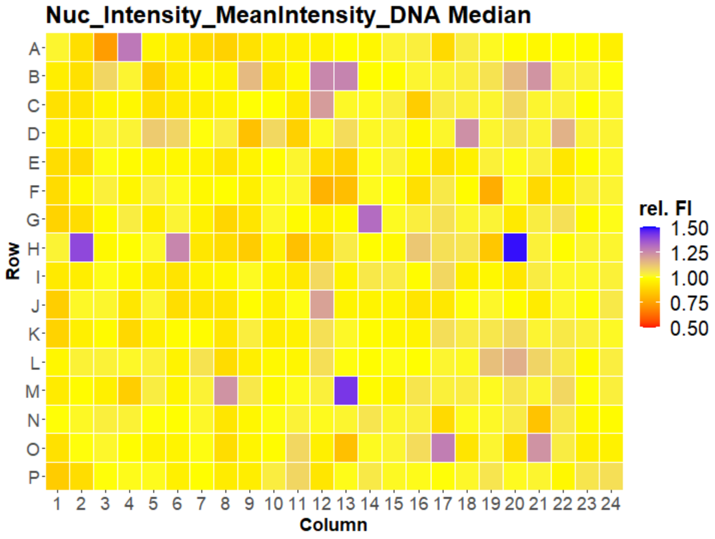

### HepG2 Medina

#### B1004 Nuc_Intensity_MeanIntensity_DNA

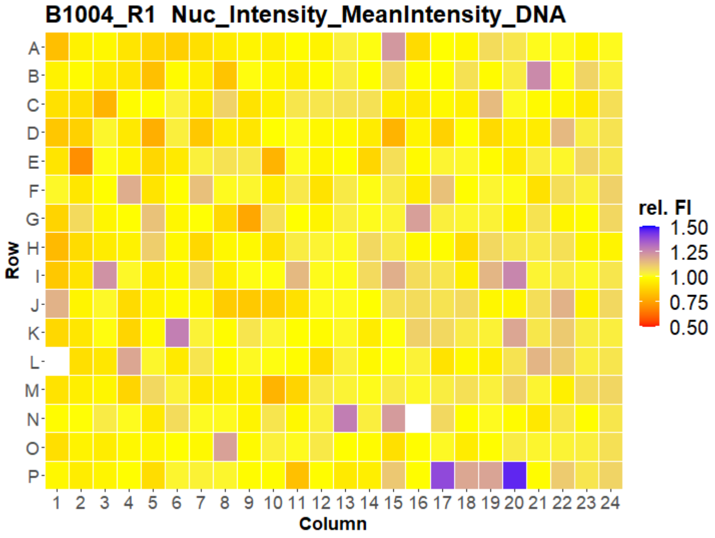


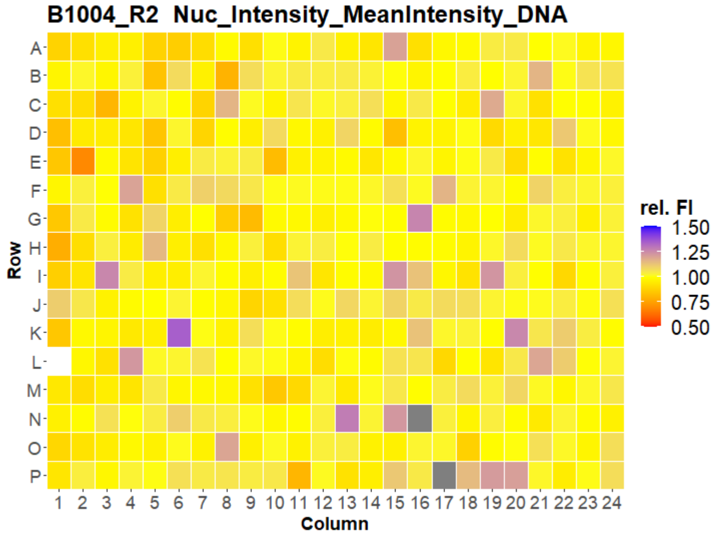


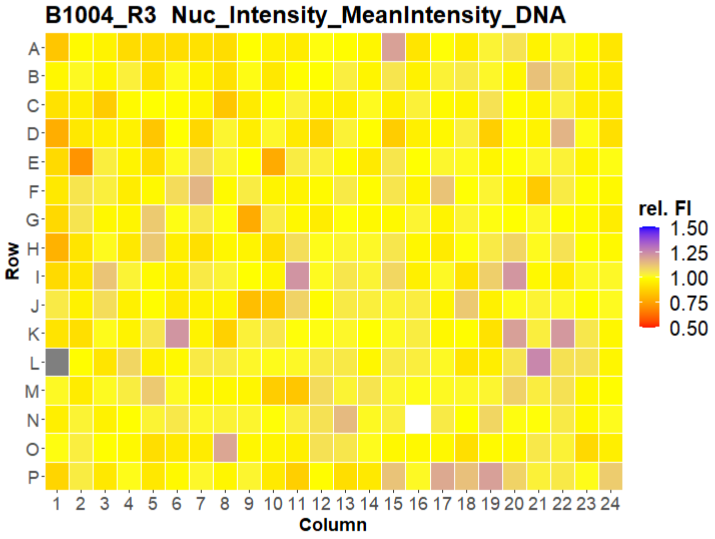


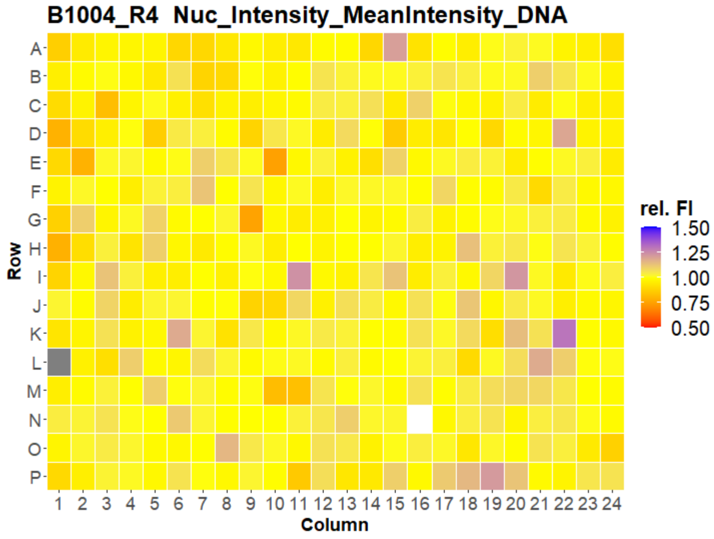


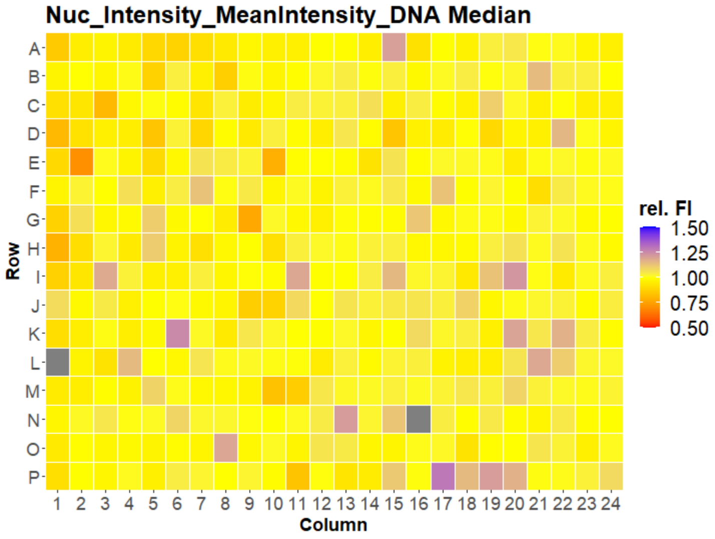

### HepG2 Medina

#### B1005 Nuc_Intensity_MeanIntensity_DNA

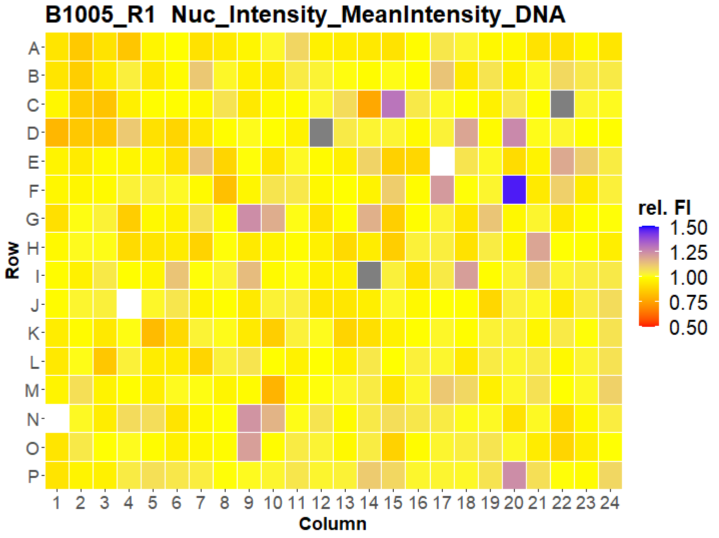


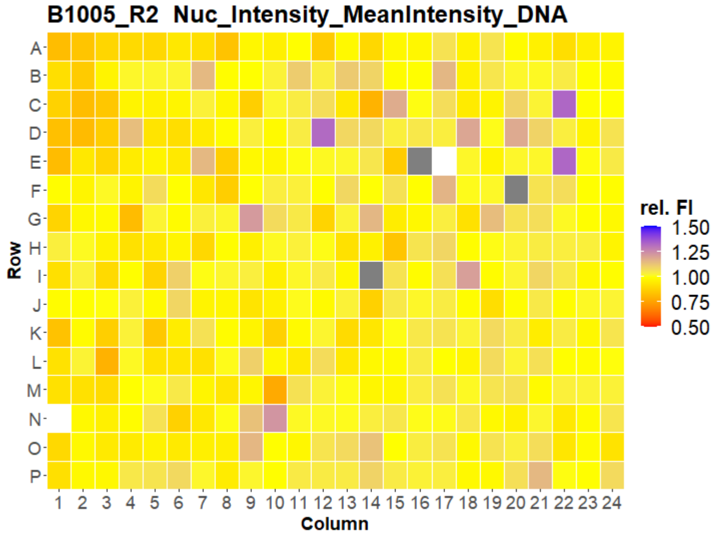


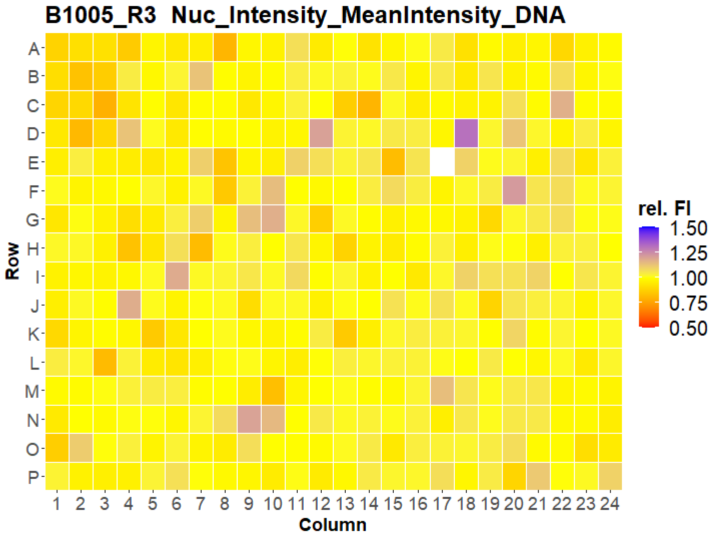


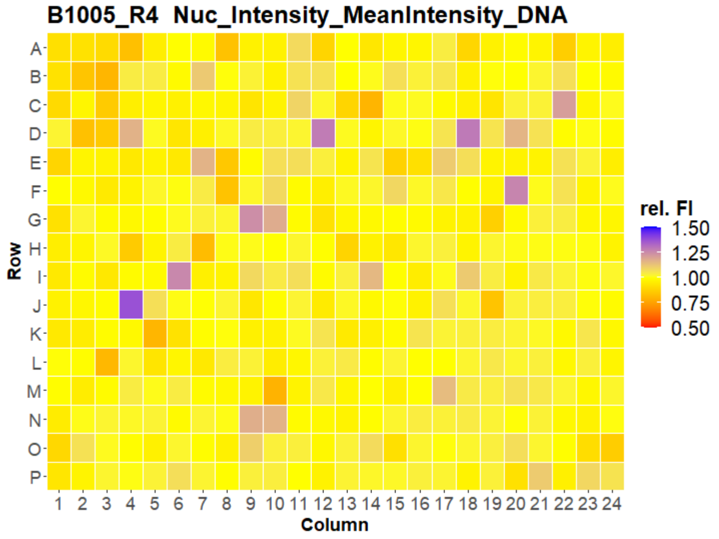


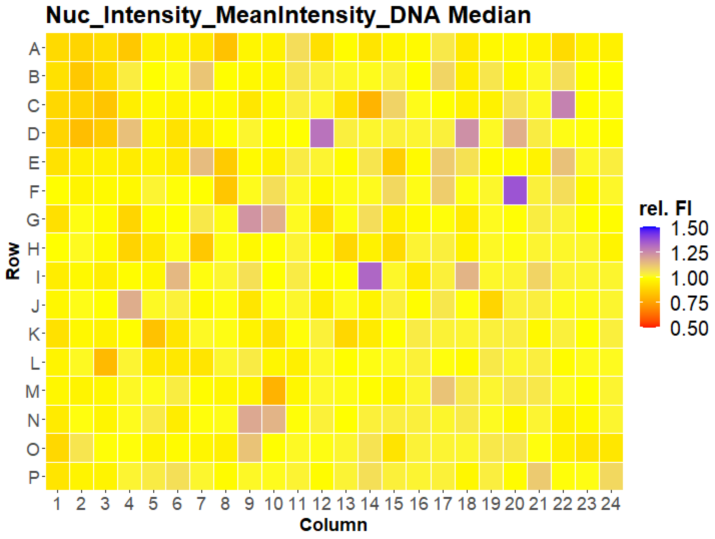

### HepG2 Medina

#### B1006 Nuc_Intensity_MeanIntensity_DNA

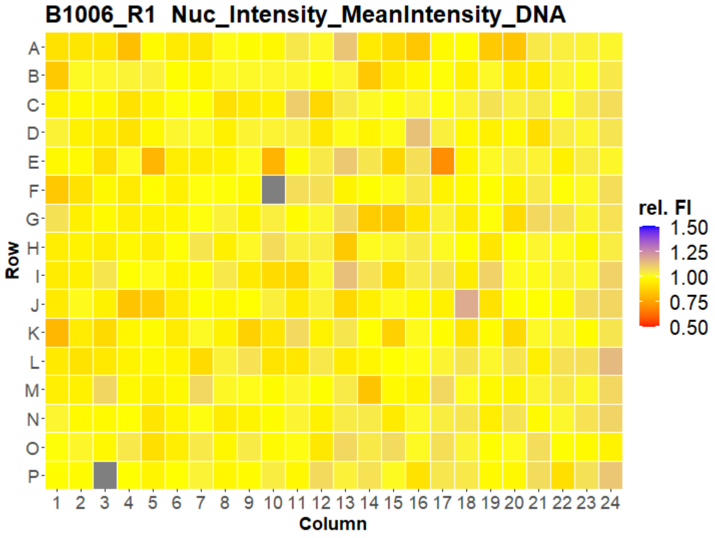


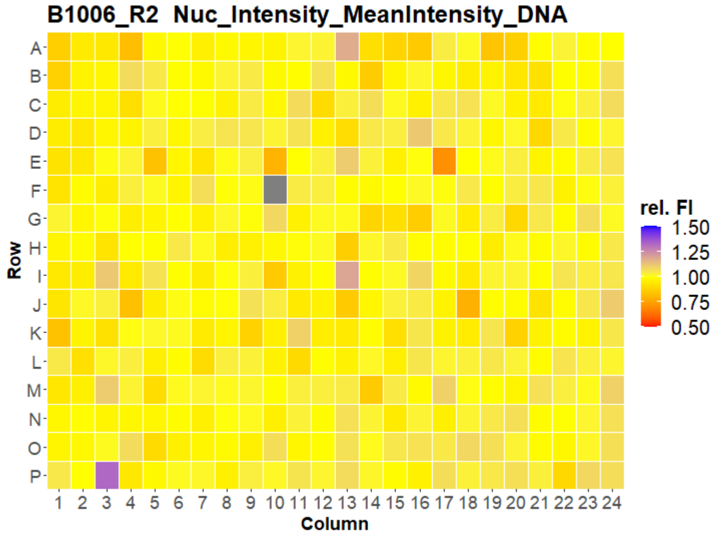


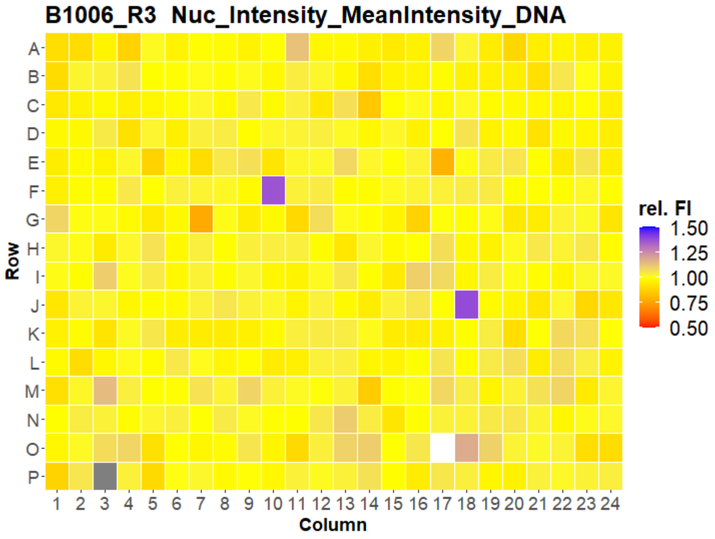


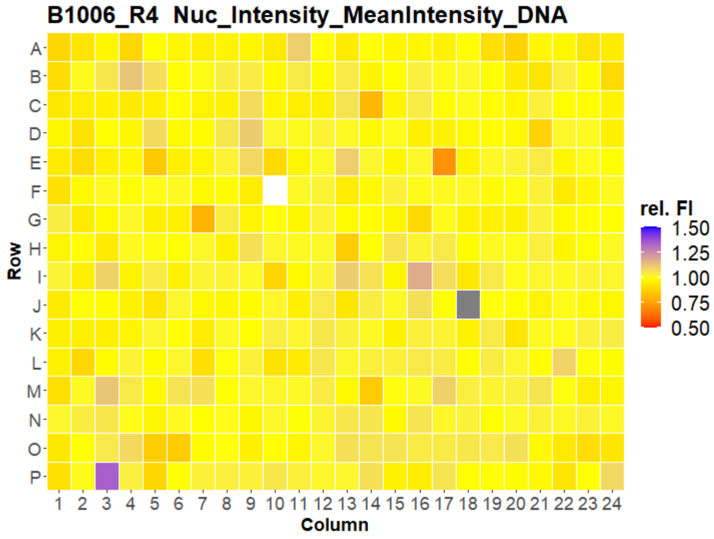


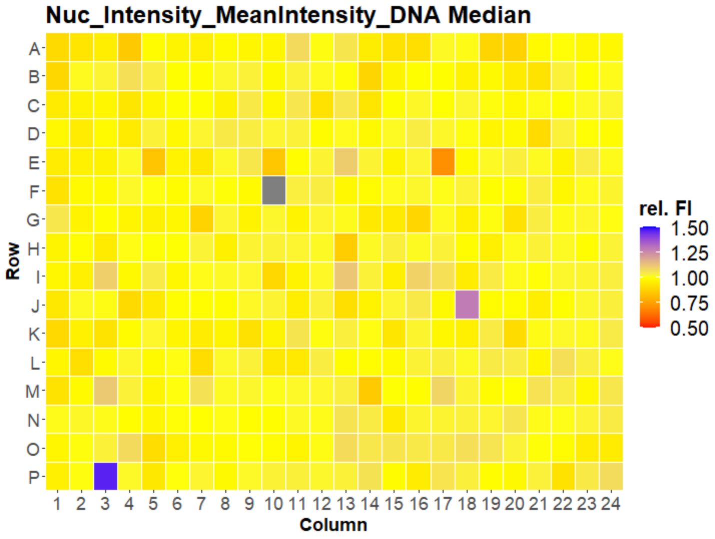

### HepG2 Medina

#### B1007 Nuc_Intensity_MeanIntensity_DNA

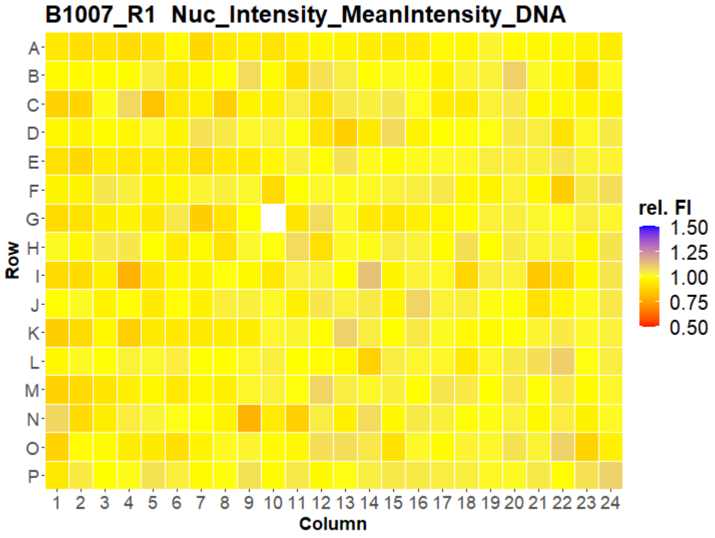


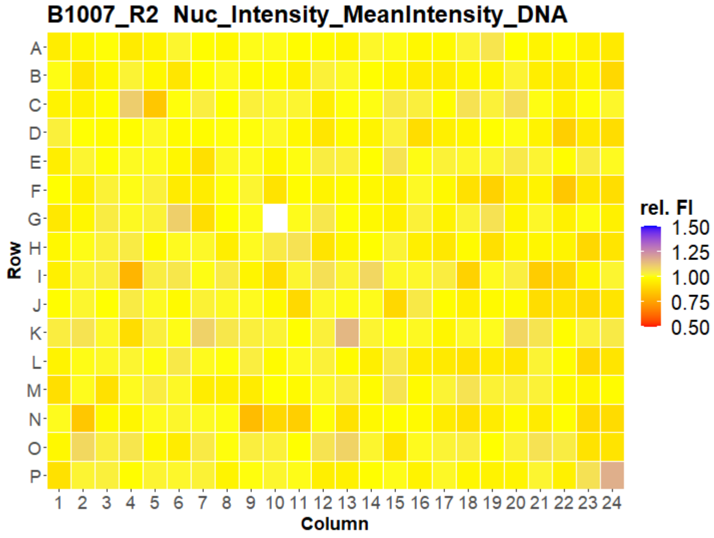


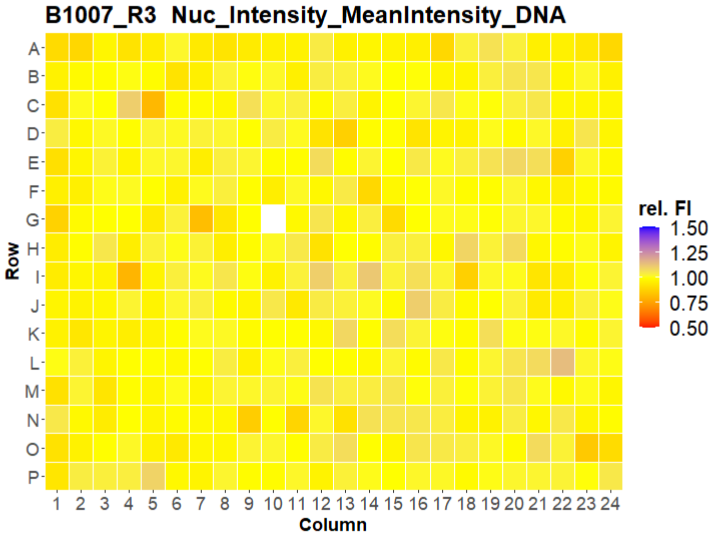


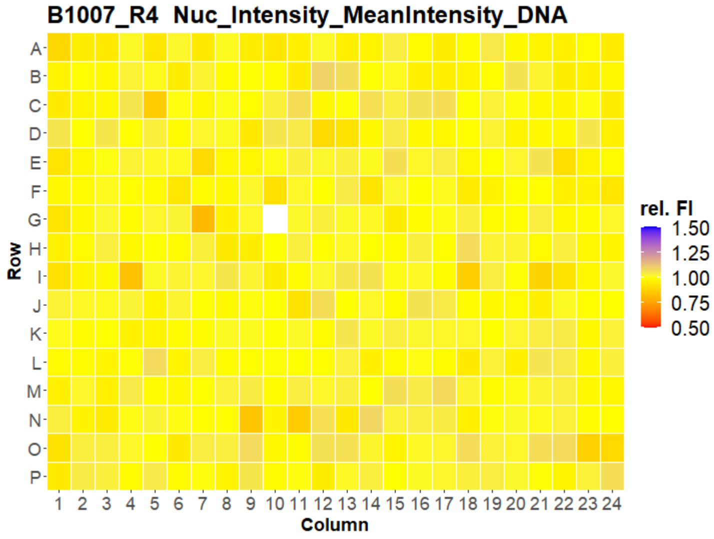


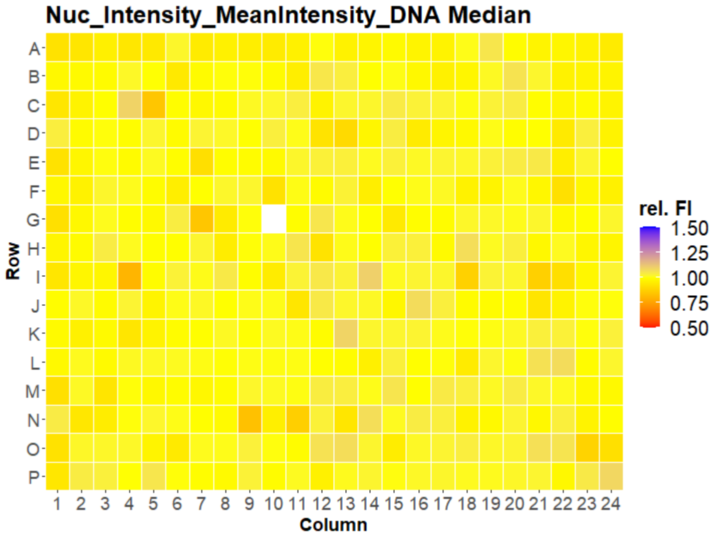

### HepG2 Medina

#### B1001 Nuc_Intensity_MeanIntensity_ER

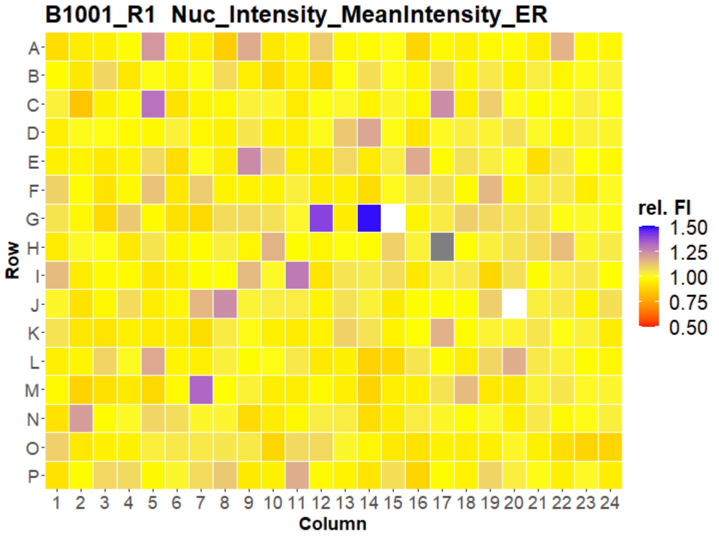


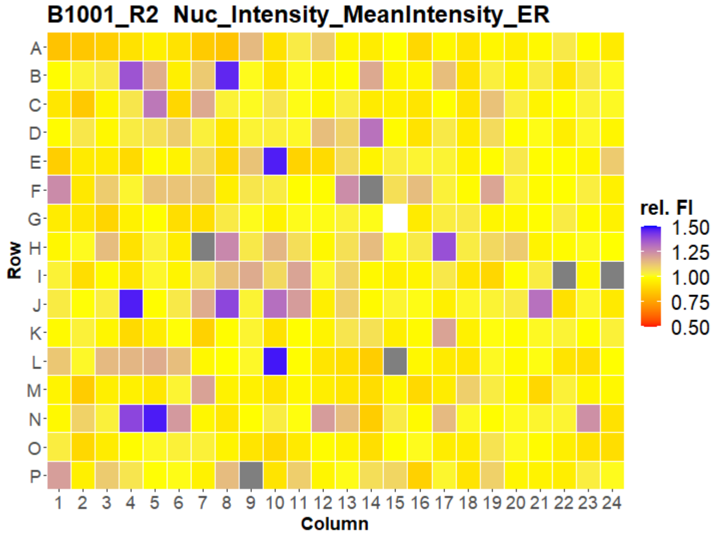


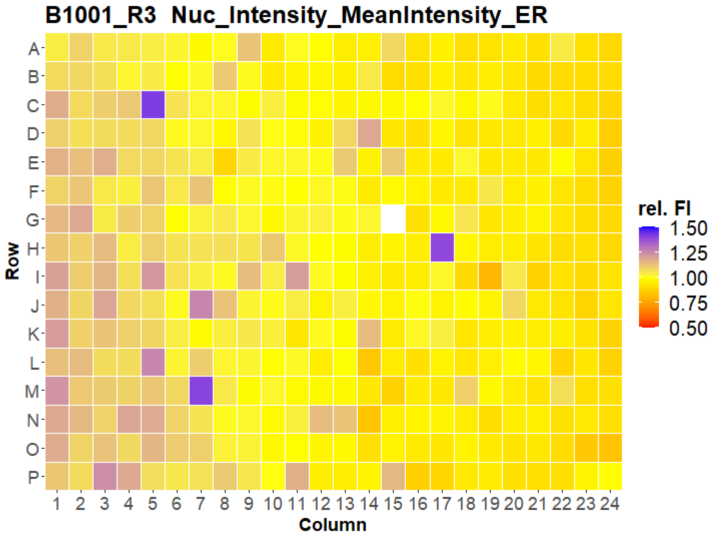


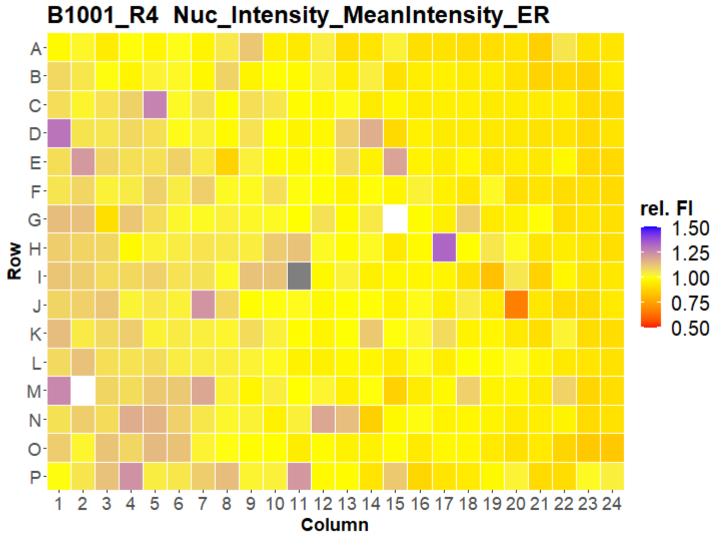


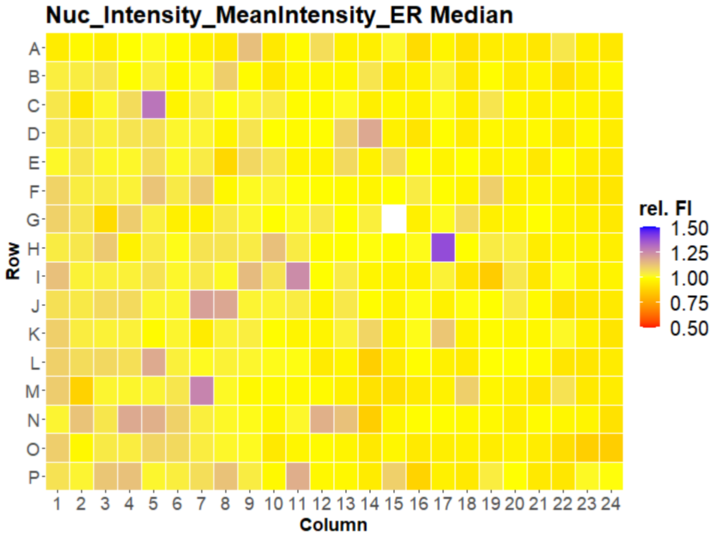

### HepG2 Medina

#### B1002 Nuc_Intensity_MeanIntensity_ER

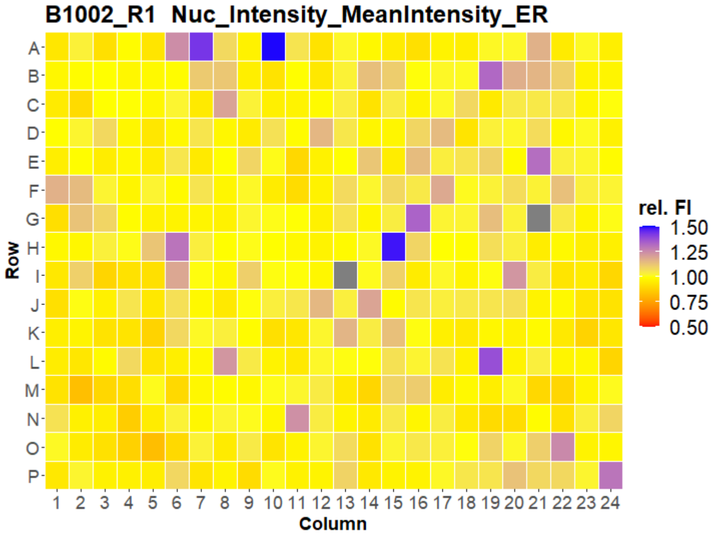


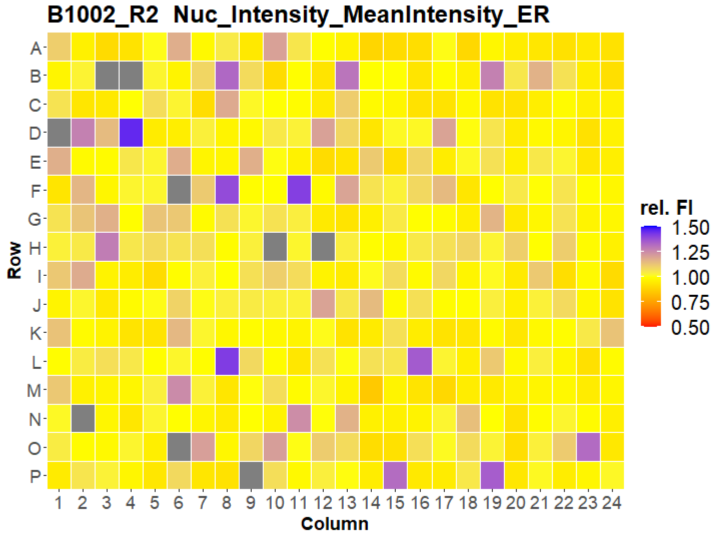


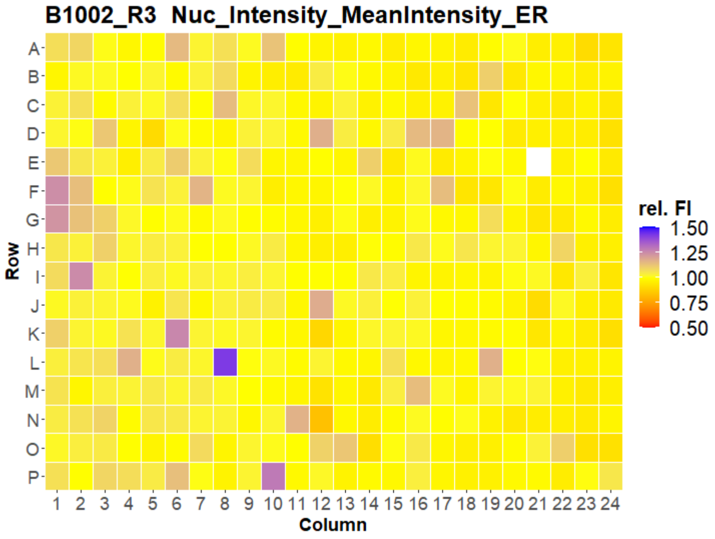


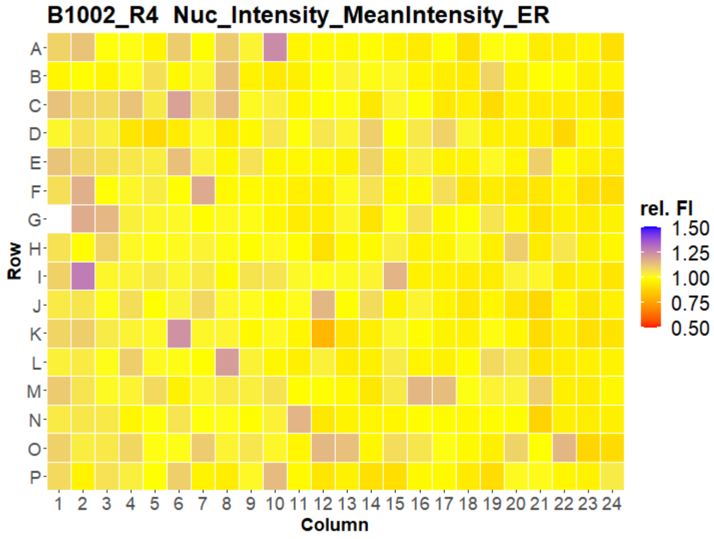


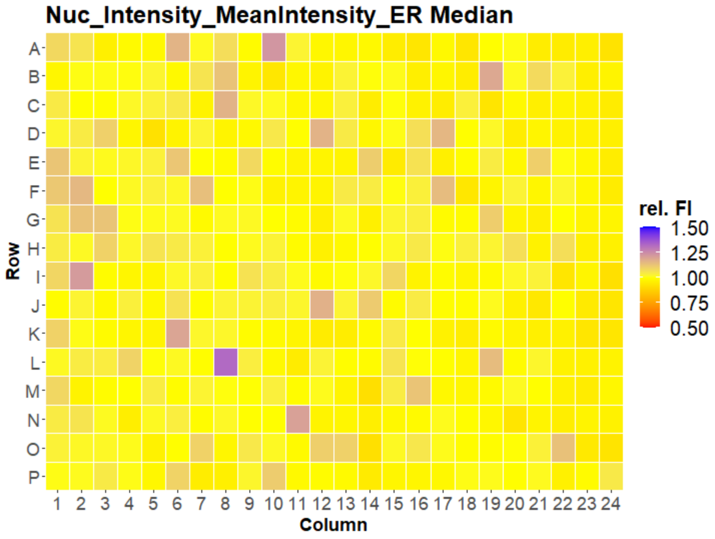

### HepG2 Medina

#### B1003 Nuc_Intensity_MeanIntensity_ER

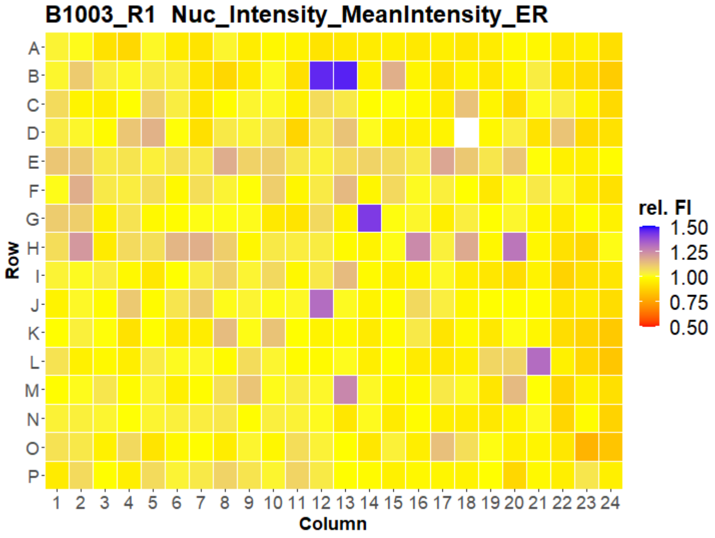


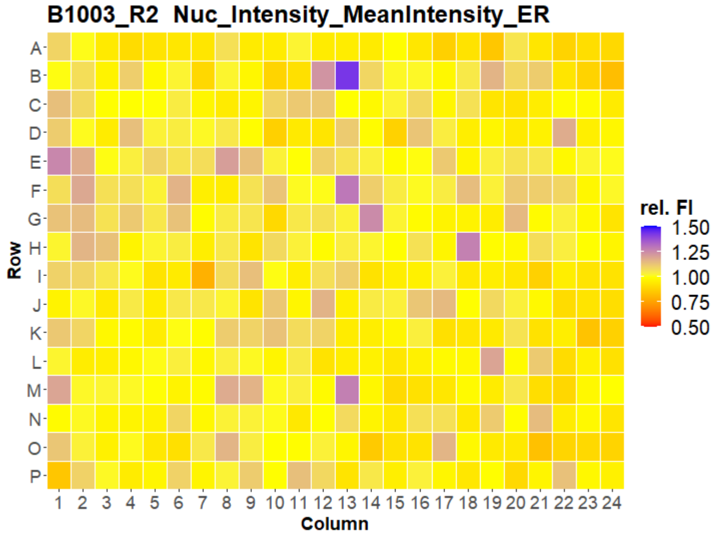


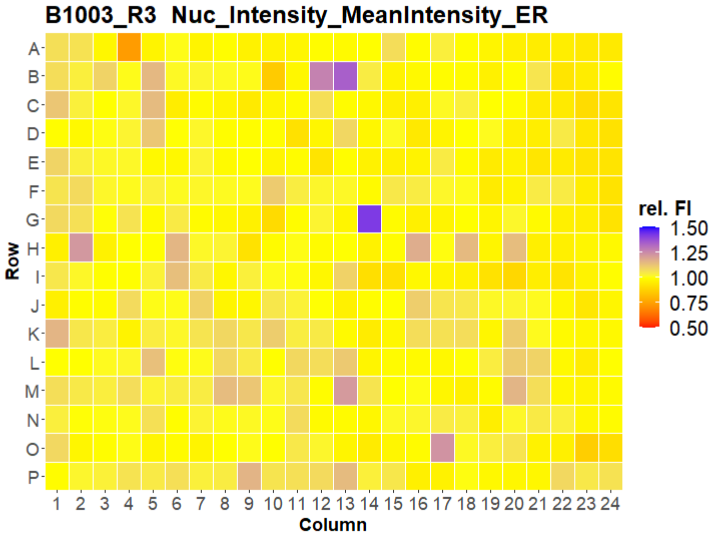


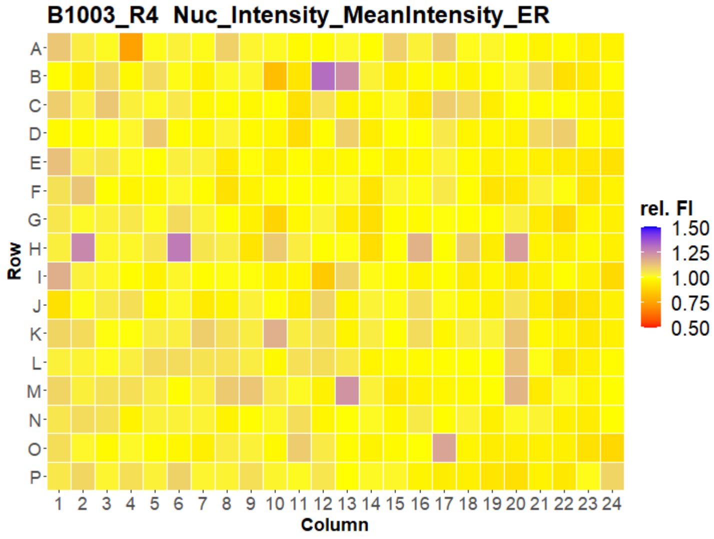


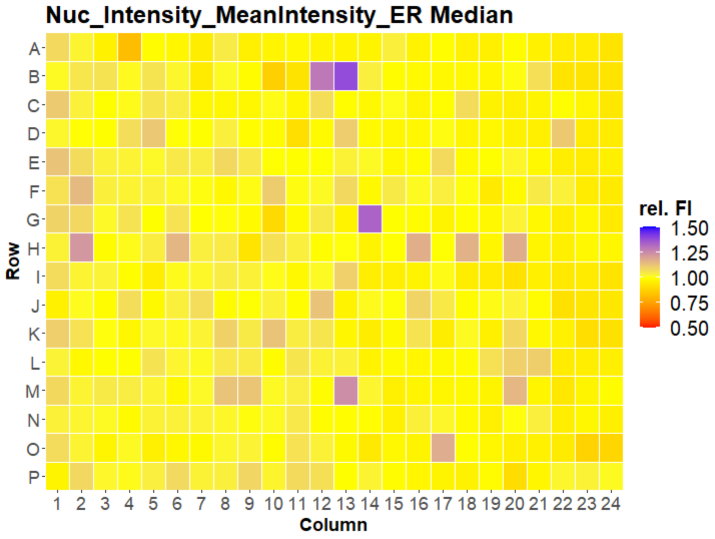

### HepG2 Medina

#### B1004 Nuc_Intensity_MeanIntensity_ER

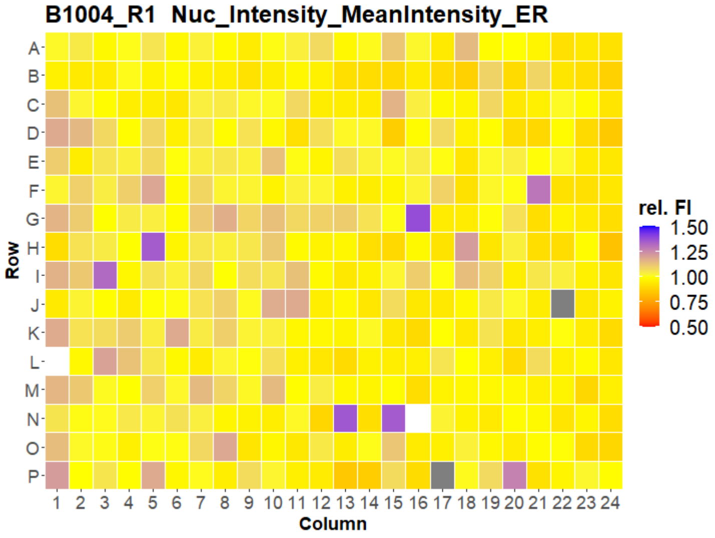


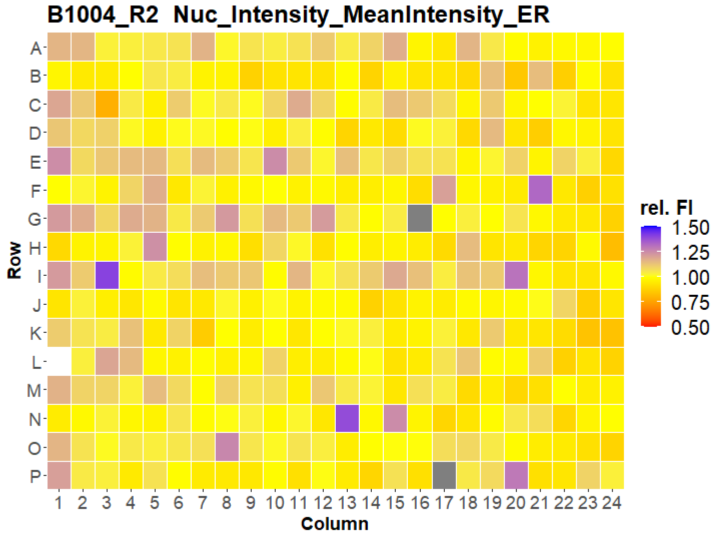


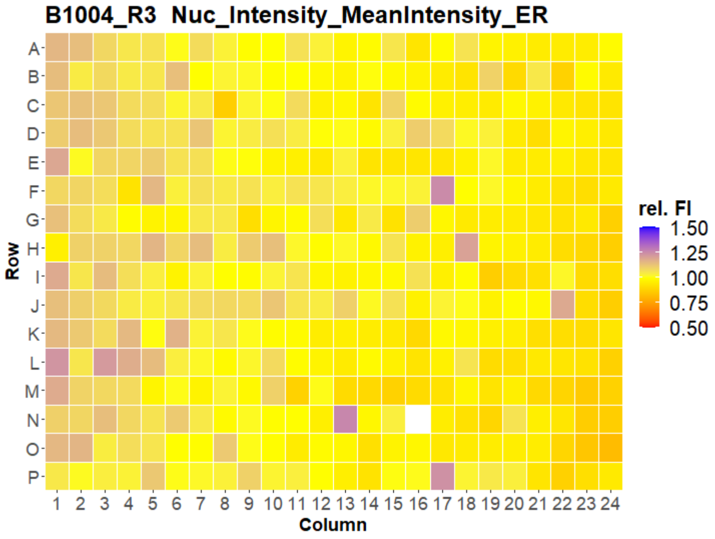


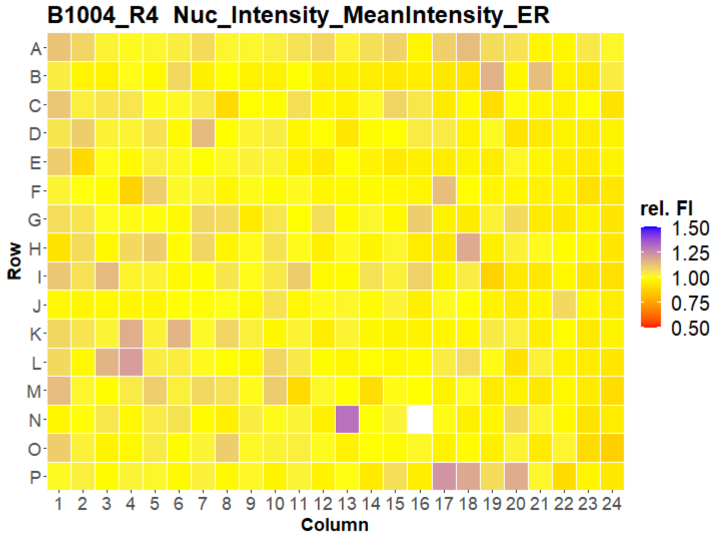


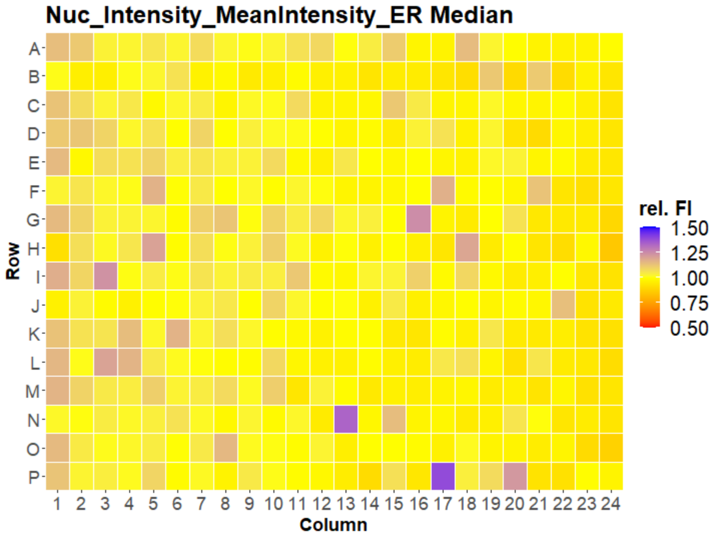

### HepG2 Medina

#### B1005 Nuc_Intensity_MeanIntensity_ER

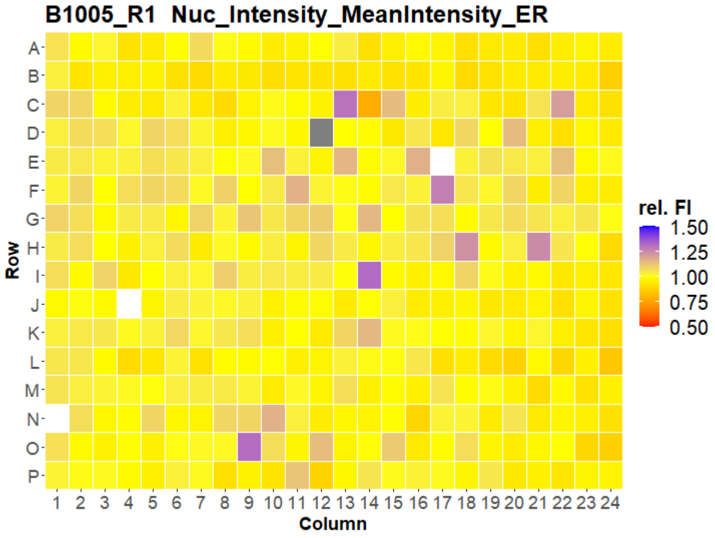


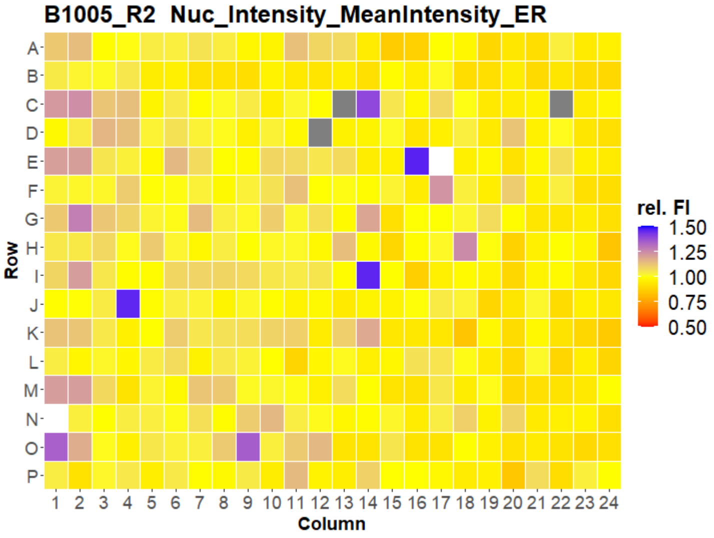


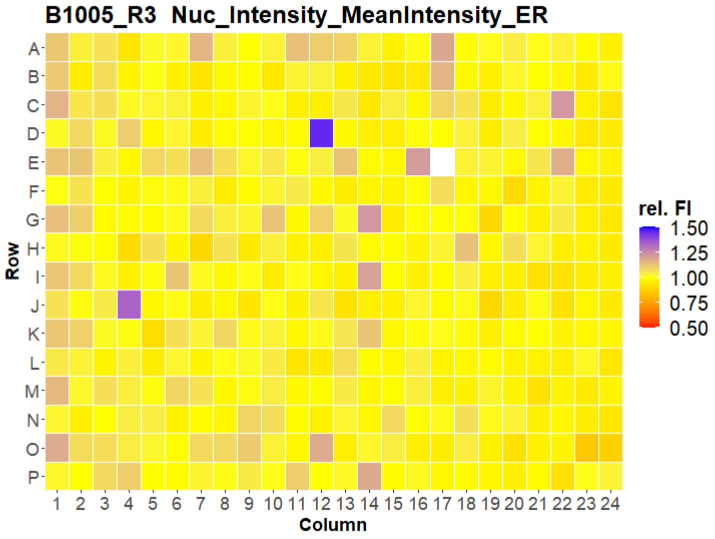


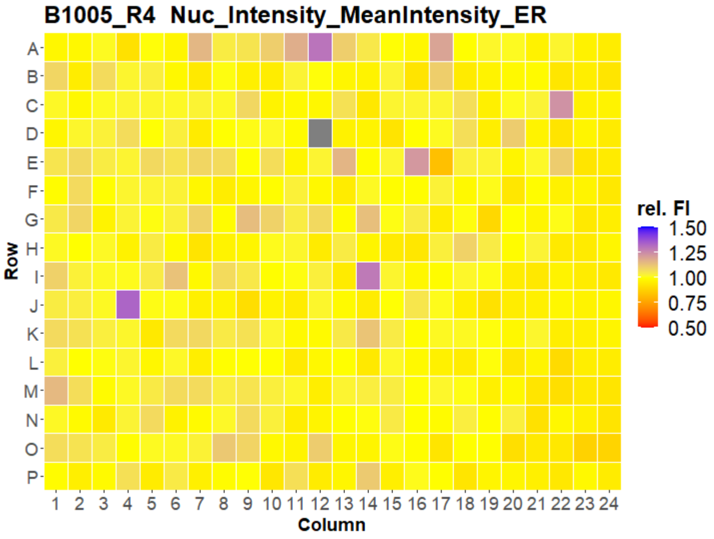


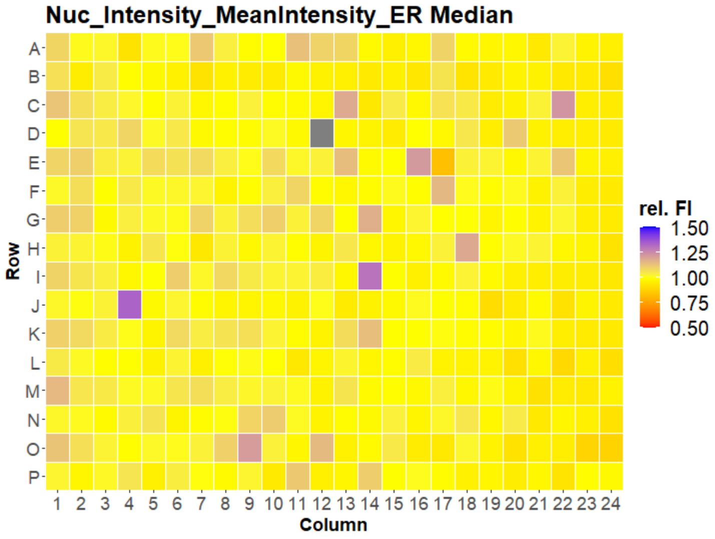

### HepG2 Medina

#### B1006 Nuc_Intensity_MeanIntensity_ER

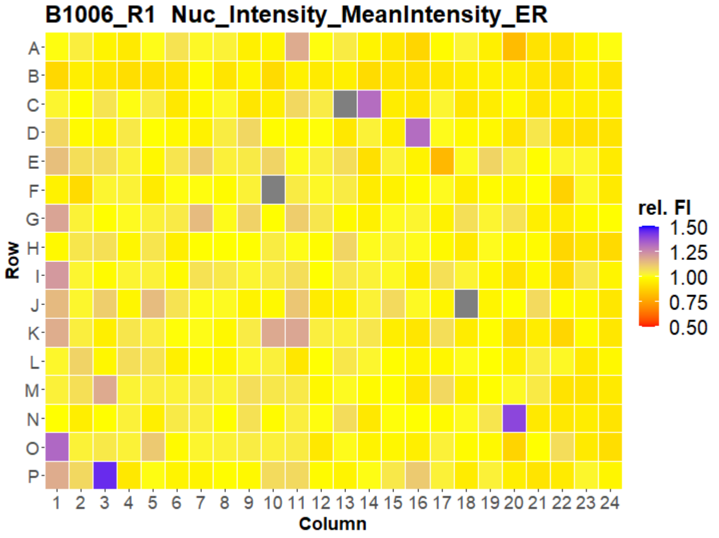


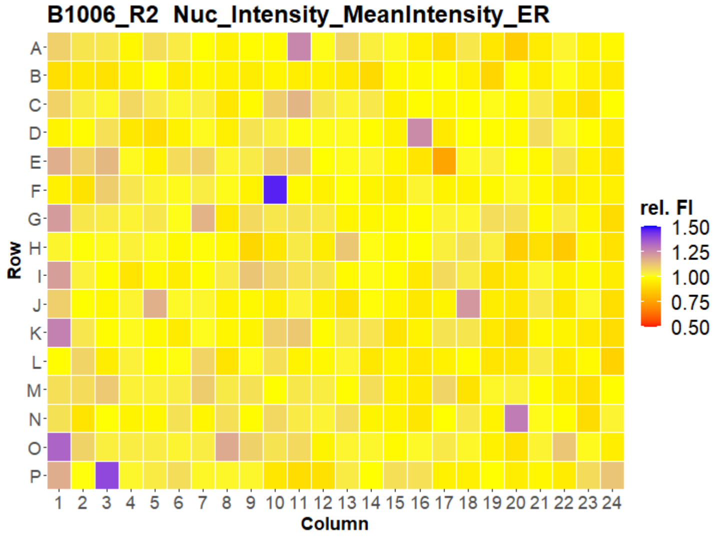


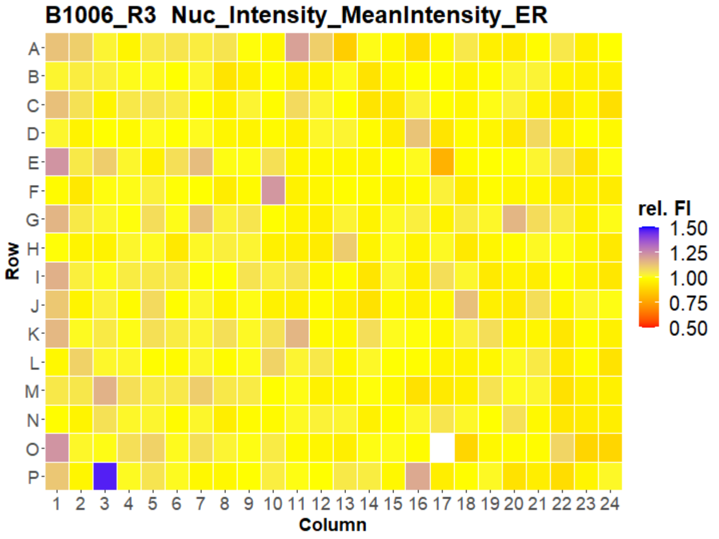


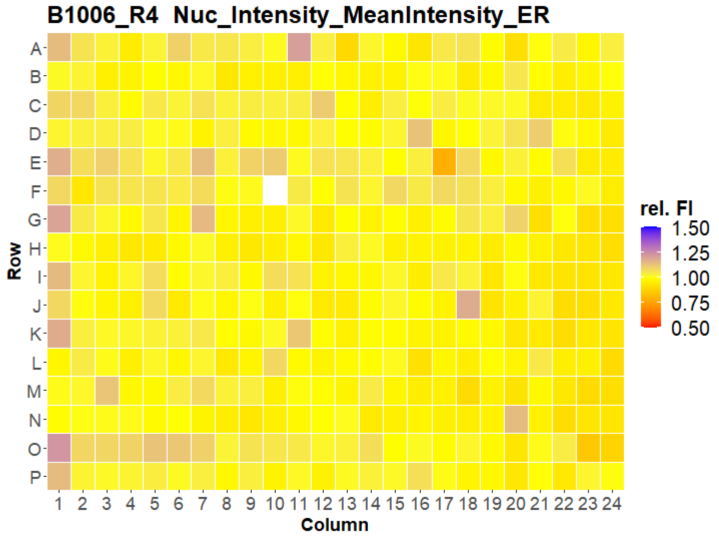


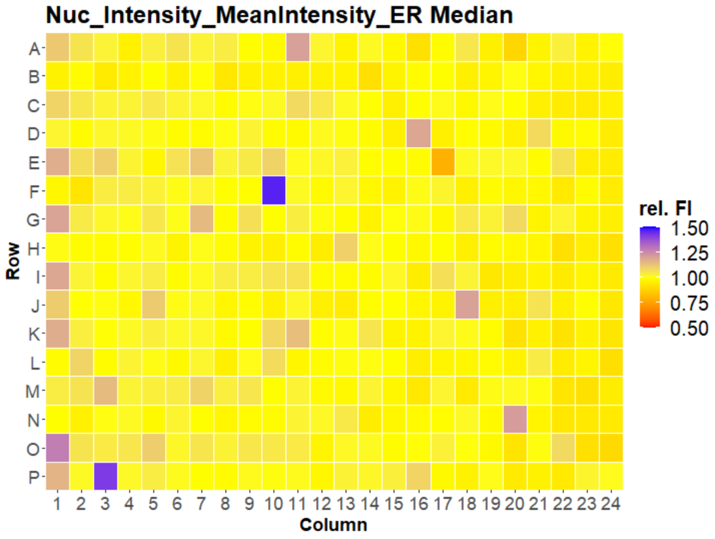

### HepG2 Medina

#### B1007 Nuc_Intensity_MeanIntensity_ER

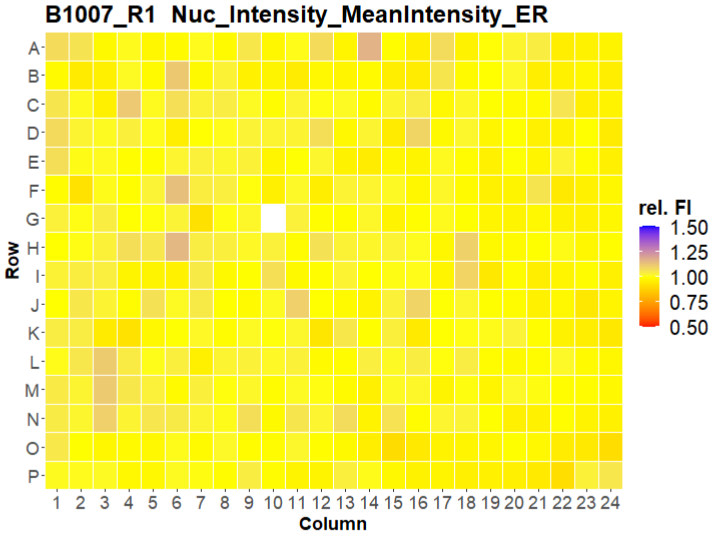


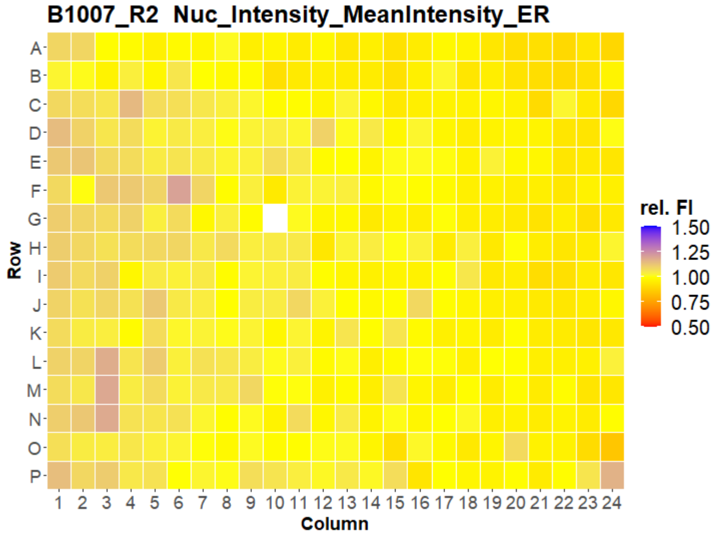


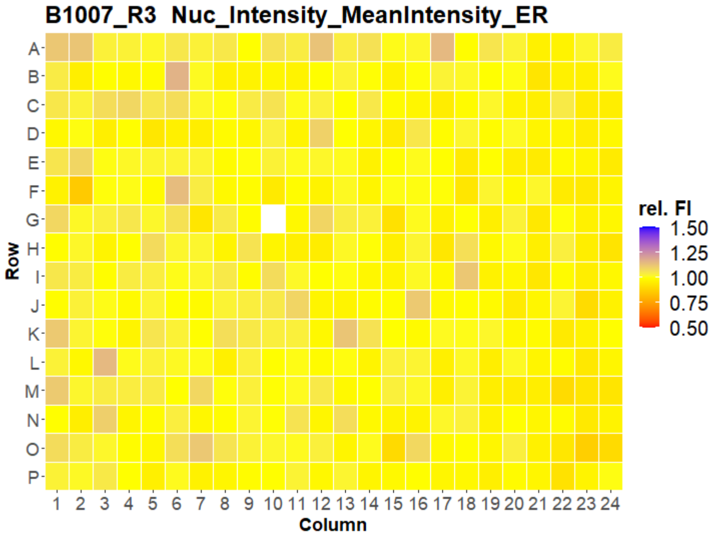


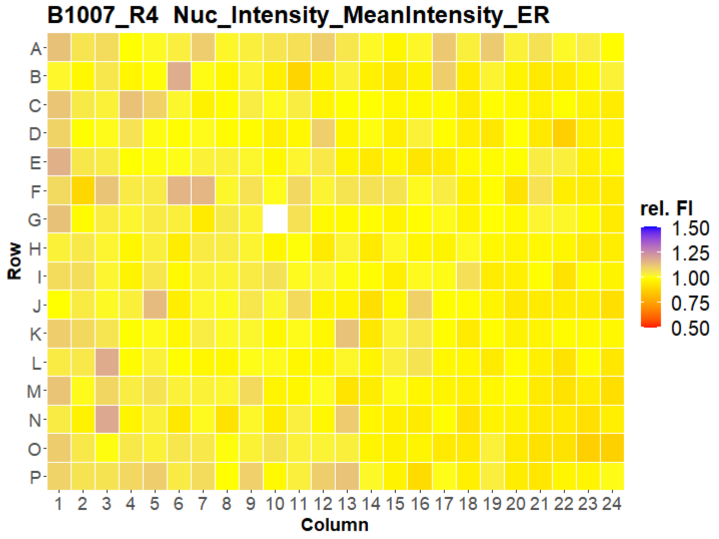


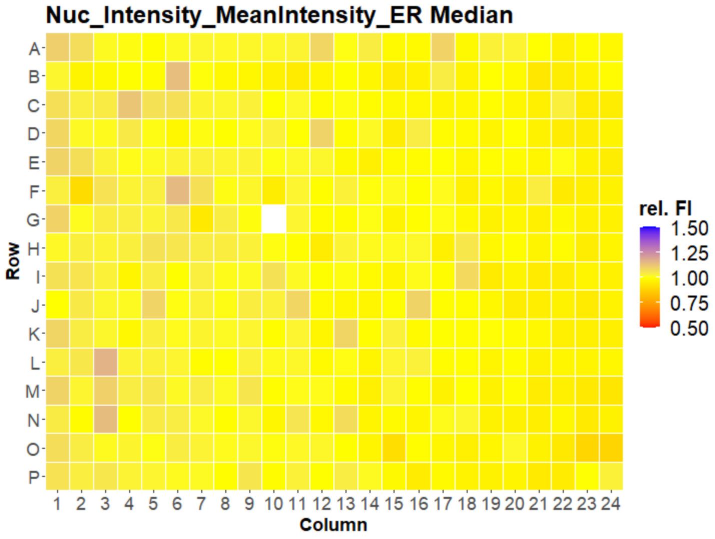

Scales:
for #Objects, the maximum object number of all plates was assigned to blue, the median object number of all plates was assigned to yellow, and zero objects were assigned to red.

for relative fluorescence intensity (rel. FI), the median fluorescence intensity of all wells of a single plate was assigned to yellow, 1.5-fold higher intensity was assigned to blue, 0.5-fold lower intensity was assigned to red.

Values that are outside of the scale values are painted in grey.
